# LIBR Methamphetamine and Opioid Cue Database (LIBR MOCD): Development and Validation

**DOI:** 10.1101/731331

**Authors:** Hamed Ekhtiari, Rayus Kuplicki, Asheema Pruthi, Martin Paulus

**Affiliations:** Laureate Institute for Brain Research (LIBR), Tulsa, OK, US; School of Community Medicine, University of Oklahoma, Tulsa, OK, US

**Keywords:** Drug Cue, Craving, Validation, Image, Database, Methamphetamine, Opioid, Heroin

## Abstract

**Introduction:** Drug cue reactivity (DCR) is widely used in experimental settings for both assessment and intervention. There is no validated database of pictorial cues available for methamphetamine and opioids.

**Methods:** 360 images in three-groups (methamphetamine, opioid and neutral (control)) matched for their content (objects, hands, faces and actions) were selected in an initial development phase. 28 participants with a history of both methamphetamine and opioid use (37.1 ± 8.11 years old, 12 female) with over six months of abstinence were asked to rate images for craving, valence, arousal, typicality and relatedness.

**Results:** All drug images were differentiated from neutral images. Drug related images received higher arousal and lower valence ratings compared to neutral images (craving (0-100) for neutral (11.5±21.9), opioid (87.7±18.5), and methamphetamine (88±18), arousal (1-9) for neutral (2.4±1.9), opioid (4.6±2.7), and methamphetamine (4.6±2.6), and valence (1-9) for neutral (4.8±1.3), opioid (4.4±1.9), and methamphetamine (4.4±1.8)). There is no difference between methamphetamine and opioid images in craving, arousal and valence. There is a significant positive relationship between the amount of time that participants spent on drug-related images and the craving they reported for the image. Every 10 points of craving were associated with an increased response time of 383millisecond. Three image sets were automatically selected for equivalent fMRI tasks (methamphetamine and opioids) from the database (tasks are available at github).

**Conclusion:** LIBR MOCD provides a resource of validated images/tasks for future DCR studies. Additionally, researchers can select several sets of unique but equivalent images based-on their psychological/physical characteristics for multiple assessments/interventions.

## 1. Introduction

Emotional and motivational response (reactivity) to conditioned stimuli (cues) is a transdiagnostic phenomenon in substance use and behavioral addictive disorders (Ekhtiari, Nasseri, Yavari, Mokri, & Monterosso, 2016). There are many published studies that have used drug related images, videos, imagery scripts, words, paraphernalia, etc., within drug cue reactivity (DCR) paradigms as an assessment or intervention. DCR can effectively induce subjective craving response (Grodin, Courtney, & Ray, 2019) and also neural activations in many brain areas associated with reward processing and decision making (Ekhtiari, Faghiri, Oghabian, & Paulus, 2016; Zilverstand, Huang, Alia-Klein, & Goldstein, 2018). DCR during abstinence has shown the potential to predict relapse (Allenby et al., 2019; Guillem & Ahmed, 2018; Li et al., 2015; Zakiniaeiz, Scheinost, Seo, Sinha, & Constable, 2017). DCR is also being explored as an intervention within exposure therapy (extinction) and memory reconsolidation paradigms (Konova & Goldstein, 2019; Liu, Tian, & Li, 2018; Torregrossa & Taylor, 2016).

Many DCR studies have used drug related images that were not properly validated before (Billieux et al., 2011). There are also many studies using image databases that are not available for use by other research groups. Meanwhile, designing cue exposure tasks with blocks of drug cues without significant differences in their psychological or physical characteristics requires normative values for dimensions like content, craving, valence, arousal, and image physical characteristics (value, hue and saturation).

International affective image system (IAPS) (Lang, Bradley, & Cuthbert, 1997) provides very few drug related images (image numbers for cigarette: 2715 and 2749, alcohol: 2600 and 2749, cocaine: 9101 and heroin: 9120) with normative values on valence, arousal and dominance. The first published database specified for appetitive images, the Normative Appetitive Picture System (NAPS), includes 18 alcohol, 6 cigarette, 12 food and 12 non-alcoholic beverage related images with the same normative values as IAPS (Stritzke, Breiner, Curtin, & Lang, 2004). In two separate studies, Khazaal, et al. asked participants to rate 60 smoking-related images (Khazaal, Zullino, & Billieux, 2012) and 60 alcohol-related images (Billieux et al., 2011) for valence, arousal and dominance. In two other studies, Ekhtiari et al, asked participants to report induced-craving for 60 methamphetamine (Ekhtiari, Alam-Mehrjerdi, Nouri, George, & Mokri, 2010) and 50 opioid related images (Ekhtiari et al., 2008).

However, there is still no published study/database (June 2019) containing drug-related and control (neutral) images available for the research community with values for craving, valence, arousal, and typicality. In this validation study, we have developed a database with 120 methamphetamine and 120 opioid related images to be rated by people with history of both methamphetamine and opioid use. Adding 120 neutral images matched for their content (objects, hands, faces and actions) with drug related images increases the potential for this database to be used on experimental DCR tasks. Our experience in designing three equivalent block-design fMRI tasks from this database for both opioids and methamphetamine is reported as well. We hope this database (LIBR MOCD) and its fMRI tasks will open doors for future developments, harmonization and collaborations with DCR paradigms using shared protocols and tasks.

## 2. Methods

### a. Initial image database development

Two hundred images representing each group (opioid, methamphetamine, and neutral) were initially collected from publicly available resources (without copy right), previous published studies (Ekhtiari et al., 2010) and the Shutterstock website (purchased with standard image license). After focused group discussions with peer-group counsellors with previous methamphetamine and opioid use, the list was reduced to 120 images (Figure S1) for each group (opioid, methamphetamine, and neutral). Each group is further divided in to 6 categories based on the image content: (i) object/drug (20 images), (ii) object/drug with hand (12 images), (iii) tools/instruments (36 images) (iv) tools/instruments with hands (20 images), (v) tools/instruments with hands in action (12 images), (vi) neutral/drug-related activities including faces (20 images).

### b. Participants

Flyers were distributed in the addiction recovery community centers in Tulsa, Oklahoma and twenty-eight participants enrolled in the study. Subjects were phone screened before being invited for further screening and consenting at the study site (LIBR). All participants signed the informed consent approved by Western IRB (protocol number: 20171742) and were appropriately compensated for their time. The Inclusion criteria consisted of having a history of both methamphetamine and opioid use and being abstinent after receiving treatment for substance use disorder for more than 6 months. Details of the demographic and clinical characteristics of the participants are summarized in table 1.

**Table 1.**
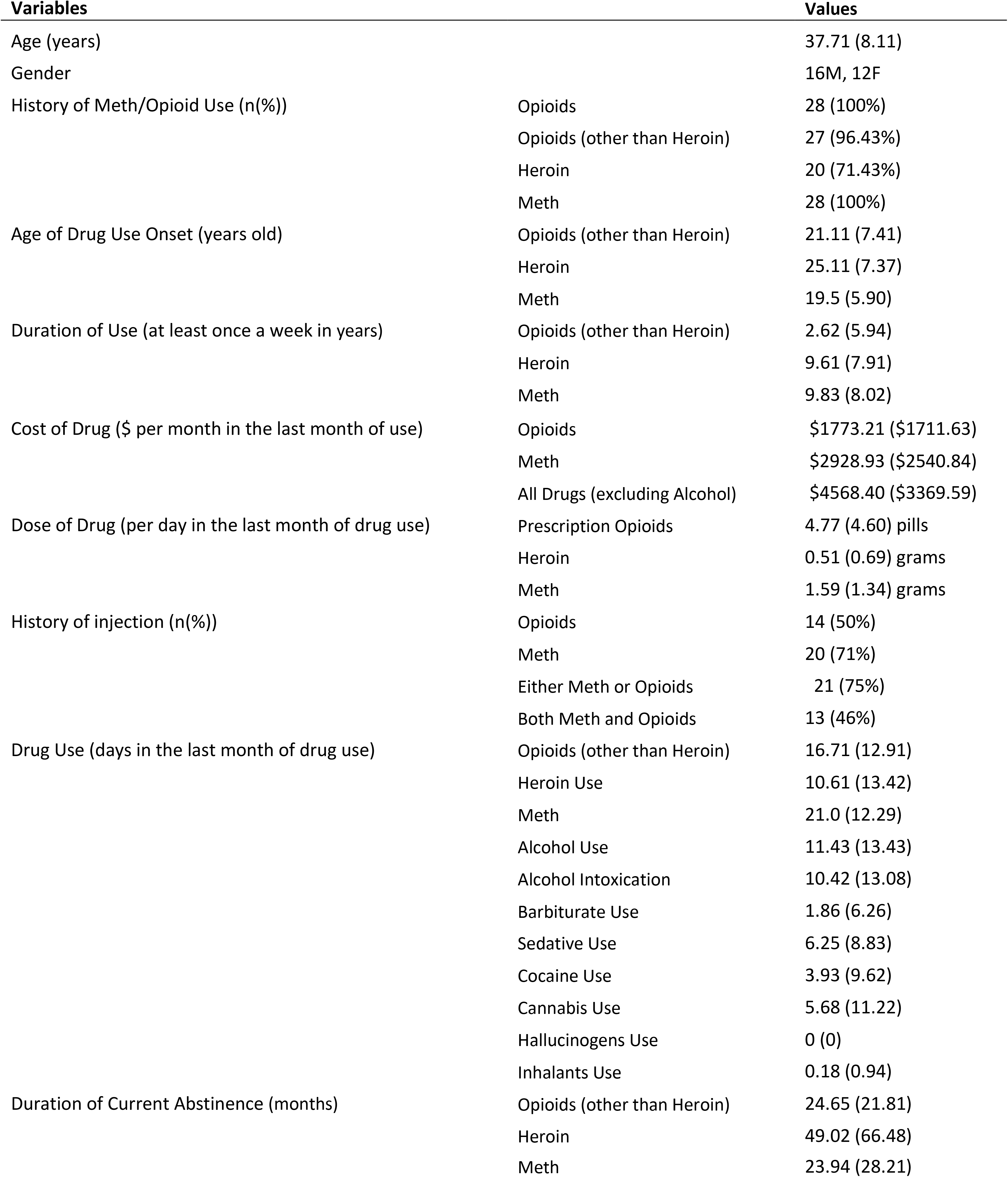
Demographics and substance use profile of participants (n=28). Values are reported as mean (SD)/frequency (Percent%)

### c. Procedure

Participants were instructed to look at different images displayed on the screen in front of them. They were asked to rate each image in terms of feeling and craving associated with each image. The participants were then asked the following questions presented consecutively under each image in 5 domains:

#### 1. Relatedness

“Is this picture related to meth, opioids, both, or none of them?” Options were meth, opioid, both, or none.

#### 2. Valence (pleasantness)

“After seeing this picture, please describe your mood, from negative to positive.” This included the Self Assessment Manikin (SAM) for valence (Bradley & Lang, 1994) with text anchors for Negative, Neutral, and Positive and numeric anchors from 1 to 9.

#### 3. Arousal (excitement)

“After seeing this picture, please describe level of arousal, from calm to excited.” This included the SAM for arousal (Bradley & Lang, 1994) with text anchors for Calm, Middle, and Excited and numeric anchors from 1 to 9.

#### 4. Craving

“How much can this picture induce drug craving in an active drug user?” This was rated using a visual analogue scale ranging from 0 (NOT AT ALL) to 100 (EXTREMELY).

#### 5. Typicality

“How frequently does an active opioid or methamphetamine user see scenes like this image during his/her drug use?” This was rated using a visual analogue scale ranging from 0 (NOT AT ALL) to 100 (EXTREMELY FREQUENT).

Each participant rated 180 images that had been randomly selected from the 360-image database and presented on the screen in each session. There was no limitation for the time subjects were allowed to spend on each question. The two cue rating sessions were separated by at least one day to reduce fatigue. A sample set of responses from one of the study participants is presented in figure 1. Participants rated their momentary drug craving before and after each sessions on the 13 item Desire for Drug Questionnaire (DDQ) (Franken, Hendriksa, & van den Brink, 2002). The DDQ includes three sub scores, (1) desire and intention with questions such as “I would consider using drugs now”, (2) negative reinforcement with questions like “Even major problems in my life would not bother me if I used drugs now” and (3) control with questions like “I could easily limit how much substance I would use if I used now”. Participants rated their response to each item in a Likert scale from 1 (strongly disagree) to 7 (strongly agree).

**Figure 1.**
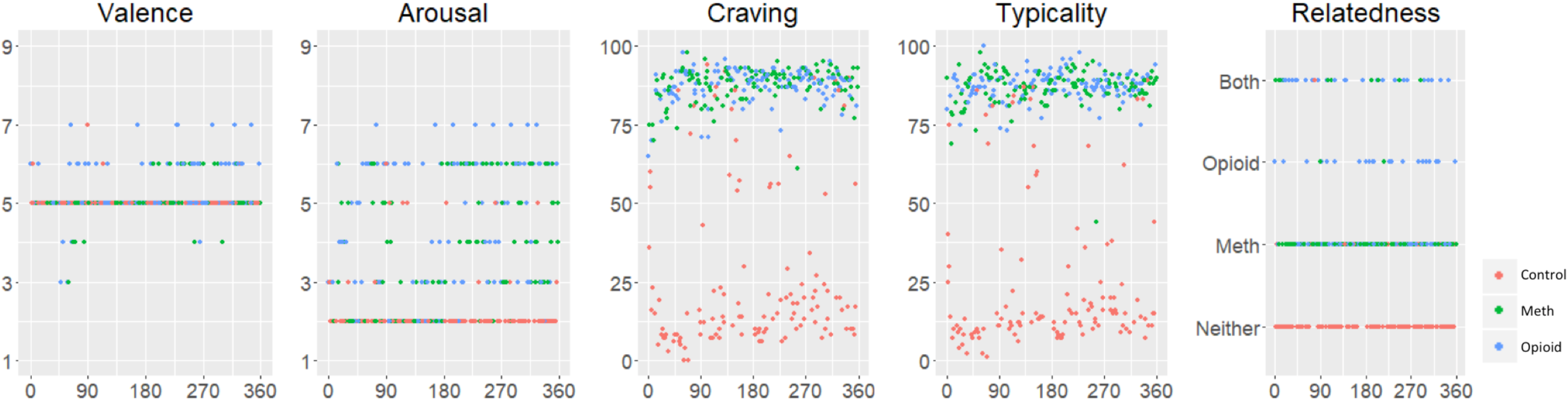
Records from a sample participant. Five questions were asked for each of the 360 pictures (120 control, 120 meth and 120 opioid, randomly presented) on valence (self assessment manikin (SAM) with 9: extremely positive, 5: neutral and 1: extremely negative), arousal (SAM with 1: calm and 9: extremely aroused), craving (visual analogues scale (VAS) 0 to 100), typicality (VAS 0-100), and relatedness.

### d. Data analysis

We have defined a derived measure, “MethToOpioid”, between 1 and −1 with considering reports for opioid as 1, methamphetamine as −1, none and both as 0 in the relatedness question. We also measured “absolute valence” as the distance between valence and 5 (neutral) (distance to mean).

A total of six linear mixed effects models were fit to investigate the effects of demographics, practice/fatigue, and clinical history on ratings and reaction times. Specifically, outcome measures were either craving rating, reaction time to the craving question, or total time to complete all five questions for the image, separately for meth and opioid images. Independent variables included age, sex, time (coded as within-session image number), visit (session 1 and 2), the time by visit interaction (Time*Visit), age of drug use onset, number of months of drug use, number of days of drug use in the last month before abstinence, amount of money spent on drugs in the month before abstinence, and days since last use, where these values were specific to the drug category. All linear mixed effect (LME) models were fit in LME4 using R and took the general form “Y ~ Age*Sex + Time*Visit + DrugOnsetAge + DrugMonthsUsed + DrugUseDaysLastMonth + MonthlyDrugCost + DrugDaysAbstinant + (1|id)”, replacing Y with the rating or reaction time and Drug with Meth or Opioid.

Effects on craving were investigated for meth and opioid groups separately using two linear mixed effects models. The outcome was craving with fixed effects for age, sex, the age by sex interaction, injection use, category (six possible values), the injection use by category interaction, as well as time, visit, and the time by visit interaction as before. The primary effects of interest were category and the injection use by category interaction, with the idea that injection users might give different ratings for injection related images than non-injection users. Explicitly, the LME4 formula was “Y ~ Time * Visit + Age * Sex + InjectionUser * Category + (1|id)” (Figure S2). In order to test for an overall task effect on the participants’ current craving state, we fit linear mixed effects models for each of the three DDQ subscales. These were collected before and after each image rating session, and the only fixed effect in the model was Time, so that a main effect of Time would indicate a change in DDQ subscale from baseline.

To design 3 equivalent fMRI tasks, 3 sets of images were generated with 24 neutral, 24 methamphetamine, and 24 opioid related images. Each image set had 4 images from each of the six categories, for a total of 24 images per set. This was accomplished in a two-step process by first selecting 72 images from each category that were matched on hue/saturation/value using an empirically derived cost function and then randomly diving them into three sets and testing for group differences until there were none.

## 3. Results

### a. Clinical Features

Demographic and clinical features of participants are summarized in table 1. There is no relationship between demographic (age and sex) and clinical features (age of methamphetamine or opioid use onset, duration of use at least once a week, days of drug use during last month before starting current recovery, monthly drug cost and duration of abstinence) of participants with the craving ratings, time spent for craving rating and total time spent for ratings with LME models explained in the data analysis section (2.d) following the correction for multiple comparison (Figure S2). There was also no significant change in craving self-reports in DDQ sub scores from pre to post rating sessions (Time F-values all < 1.9 and p-values all > 0.13).

### b. Picture Ratings

Mean and standard deviation of the responses to each image by all participants and participants with or without drug injection history along with mean and standard deviation of contrast, hue and saturation values of pixels within the image is reported in a database (supplementary material 1). Final product of all images including their validation data is provided in supplementary materials 2 to 4 for neutral (control), methamphetamine and opioid images consecutively.

Mean typicality for each image was highly correlated with mean craving in drug related images (Pearson correlation coefficient (R)=0.89) (Figure 2). Distribution of responses to craving, valence and arousal are represented in figure 3. There is a two-peak distribution for arousal, one around 1 (“Calm”) and the other around 5 (“Middle”). Valence has a single peak around 5 (“Neutral”). Both methamphetamine and opioids have significantly higher craving, lower valence and higher arousal compared to neutral images (Table S1). There is no difference between methamphetamine and opioid images in craving, arousal and valence (Table S1). Reported values for images in each category are presented in table 2. Image category had main effects on craving ratings to opioid (F=6.4, p-value<0.001) and meth (F=3.5, p-value=0.004), but there was no category by injection use interaction for either image set (F= 0.004, p-value=0.94 and F=0.035, p-value=0.85).

**Table 2.** Craving, arousal, valence, typicality and relatedness responses in six categories of the pictures. Values are reported as mean (standard deviation). Relatedness question has four options whether the picture relates to meth, opioid, both or neither. We have reported average frequency of response for these four options in Meth, Opioid, Both and Neither columns (The values add up to 1 in each row). MethToOpioid derived measure for drug pictures will represent 1 if all subjects relate a picture to opioids and −1 if all subjects relate a picture to meth.

| Categories (number) |  | Craving<br>(0-100) | Valence<br>(1-9) | Arousal<br>(1-9) | Typicality<br>(0-100) | Relatedness |  |  |  |  |
| --- | --- | --- | --- | --- | --- | --- | --- | --- | --- | --- |
|  |  |  |  |  |  | Meth | Opioid | Both | Neither | MethToOpioid |
|  | **Neutral Pictures (120)** | **11.59 (21.93)** | **4.87 (1.37)** | **2.49 (1.95)** | **12.29 (21.11)** | **0.013** | **0.006** | **0.006** | **0.973** |  |
| 1 | Neutral Objects (20) | 12.23 (22.97) | 4.91 (1.33) | 2.5 (1.97) | 11.68 (21.26) | 0.002 | 0.022 | 0.013 | 0.964 |  |
| 2 | Neutral Objects with Hands (12) | 13.38 (25.21) | 4.95 (1.39) | 2.56 (2.07) | 12.19 (21.79) | 0.009 | 0.015 | 0.006 | 0.970 |  |
| 3 | Neutral Tools (36) | 10.8 (20.43) | 4.86 (1.38) | 2.49 (1.91) | 12.39 (20.76) | 0.020 | 0.000 | 0.005 | 0.975 |  |
| 4 | Neutral Tools with Hands (20) | 10.98 (21.02) | 4.86 (1.38) | 2.51 (1.97) | 12.51 (21.06) | 0.016 | 0.002 | 0.004 | 0.978 |  |
| 5 | Neutral Tools, Hands and Actions (12) | 10.16 (20.52) | 4.85 (1.4) | 2.4 (1.9) | 10.25 (18.11) | 0.003 | 0.000 | 0.003 | 0.994 |  |
| 6 | Faces and Neutral Activities (20) | 12.8 (22.96) | 4.84 (1.31) | 2.48 (1.95) | 13.78 (22.79) | 0.020 | 0.004 | 0.007 | 0.969 |  |
|  | **Meth Pictures (120)** | **87.74 (18.56)** | **4.43 (1.918)** | **4.66 (2.72)** | **89.27 (17.81)** | **0.68** | **0.035** | **0.269** | **0.014** | **-0.644** |
| 1 | Meth Drug (20) | 89.83 (16.5) | 4.46 (2.02) | 4.73 (2.79) | 90.63 (16.57) | 0.955 | 0.015 | 0.029 | 0.002 | -0.940 |
| 2 | Meth with Hands (12) | 88.32 (16.74) | 4.47 (1.9) | 4.68 (2.74) | 90.4(15.3) | 0.671 | 0.035 | 0.285 | 0.009 | -0.636 |
| 3 | Meth Use Instruments (36) | 87.8 (17.57) | 4.54 (1.87) | 4.66 (2.72) | 89.51 (16.86) | 0.663 | 0.082 | 0.255 | 0.000 | -0.581 |
| 4 | Meth Use Instruments with Hands (20) | 88.86 (17.17) | 4.16 (1.98) | 4.81 (2.72) | 89.85 (17.23) | 0.122 | 0.046 | 0.824 | 0.009 | -0.076 |
| 5 | Meth related Instruments, Hands and Actions (12) | 86.54 (20.33) | 4.44 (1.87) | 4.59 (2.69) | 88.24 (19.49) | 0.658 | 0.034 | 0.276 | 0.031 | -0.624 |
| 6 | Face and Meth Related Activities (20) | 86.49 (20.05) | 4.43 (1.87) | 4.63 (2.68) | 88.17 (18.93) | 0.800 | 0.027 | 0.160 | 0.013 | -0.773 |
|  | **Opioid Pictures (120)** | **88.08 (18.09)** | **4.43 (1.872)** | **4.66 (2.69)** | **89.52 (17.09)** | **0.113** | **0.443** | **0.43** | **0.013** | **0.33** |
| 1 | Opioid Drug (20) | 86.45 (18.32) | 4.65 (1.78) | 4.49 (2.67) | 88.45 (17.38) | 0.082 | 0.725 | 0.184 | 0.009 | 0.644 |
| 2 | Opioid with Hands (12) | 89.42 (16.83) | 4.31 (1.93) | 4.72 (2.69) | 90.53 (15.9) | 0.105 | 0.307 | 0.580 | 0.007 | 0.202 |
| 3 | Opioid Use Instruments (36) | 85.41 (19.09) | 4.7 (1.69) | 4.48 (2.62) | 87 (19) | 0.200 | 0.606 | 0.164 | 0.030 | 0.406 |
| 4 | Opioid Use Instruments with Hands (20) | 90.54 (15.98) | 4.16 (2.05) | 4.89 (2.75) | 91.23 (15.48) | 0.085 | 0.255 | 0.655 | 0.006 | 0.170 |
| 5 | Opioid related Instruments, Hands and Actions (12) | 87.46 (19.69) | 4.4 (1.87) | 4.67 (2.7) | 89.35 (17.89) | 0.112 | 0.402 | 0.468 | 0.018 | 0.290 |
| 6 | Faces and Opioid Related Activities (20) | 89.6 (16.2) | 4.37 (1.84) | 4.7 (2.7) | 90.36 (15.99) | 0.118 | 0.389 | 0.485 | 0.007 | 0.271 |

**Figure 2.**
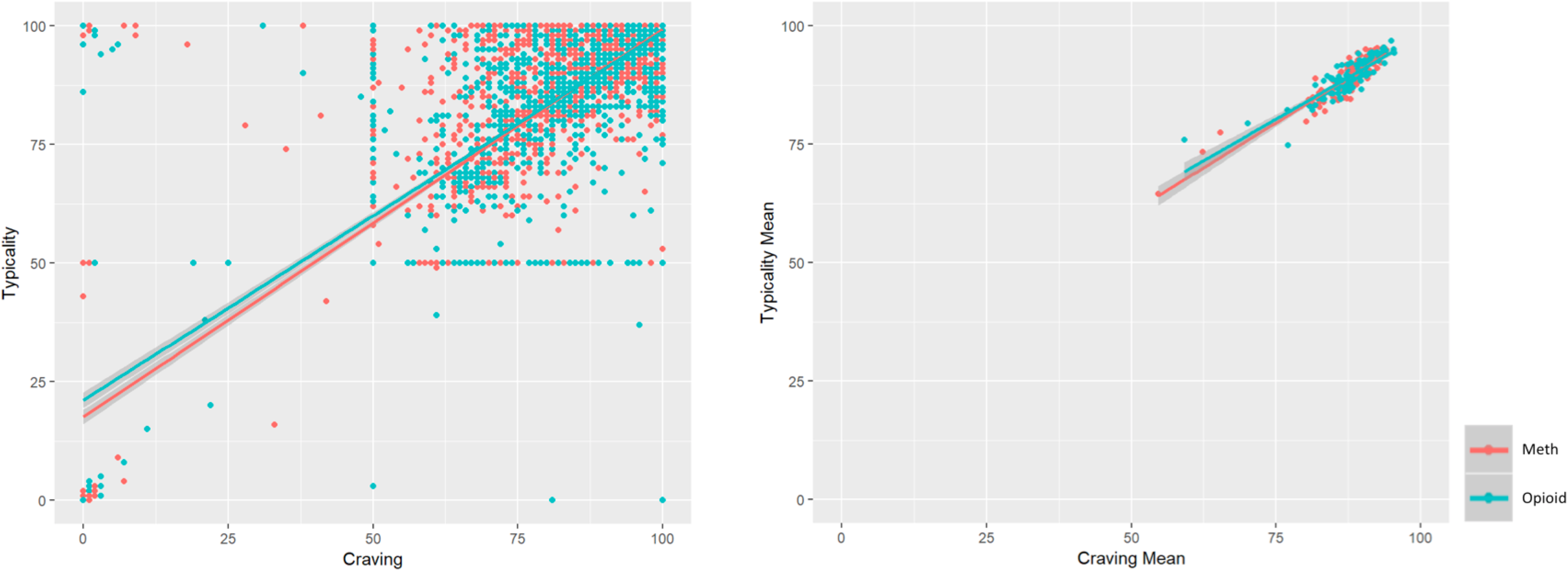
Distribution of craving and typicality responses in meth and opioid pictures. The left scatter plot represents all individual responses and the right scatter plot represents average in each individual picture (Pearson correlation R= 0.89).

**Figure 3.**
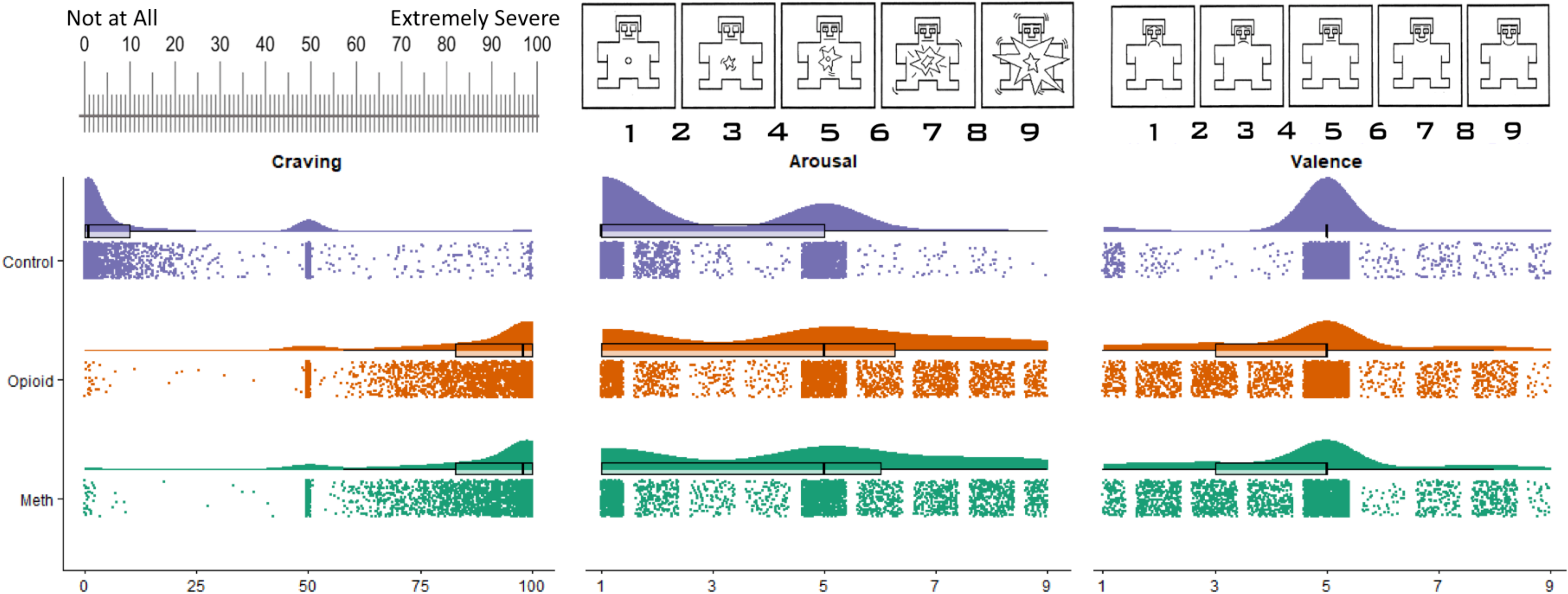
Distribution of responses to pictures. Self assessment manikin (SAM) and visual analogue scale (VAS) used for the ratings are also provided. Boxpolts represent quartiles. Values jittered for better visualization.

Neutral images are all differentiated from the drug related images with over 92 percent of reports on “none” in the relatedness question (neither related to methamphetamine nor opioid) except for 5 images (39, 62, 85, 89, 89 percent, the highest report for “neither” in drug related images is 28 percent). Two neutral images with highest percent of reporting as drug related images contain light bulbs (potentially due to the use of heat resistant light bulb glass to create pipe for drug use) (Figure 4, first two neutral images with highest craving report). Sample drug related images with highest reported craving are presented in figure 4.

**Figure 4.**
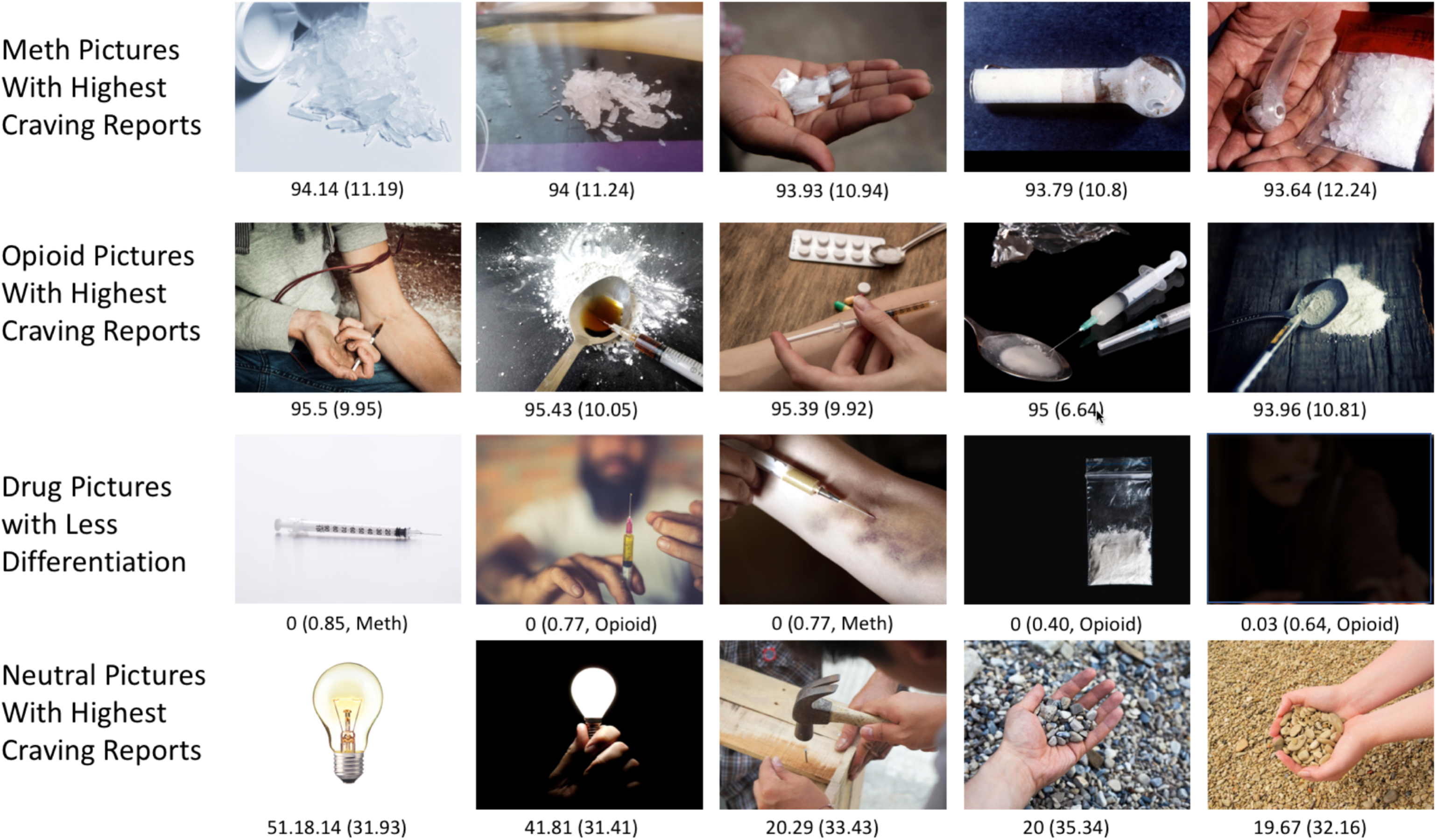
Sample pictures from the database. Five meth pictures and 5 opioid pictures with highest reported craving (mean (standard deviation)), 5 meth or opioid pictures that are less reported to be related to just opioid or meth categories (relatedness score, −1: meth, 1: opioid, in parenthesis average frequency of receiving “Both Meth and Opioid” response in relatedness question and the original category reported as well) and 5 neutral pictures with highest reported craving. We removed few elements in this figure in this version of the manuscript to respect biorxiv’s policy “to avoid the inclusion of photographs and any other identifying information of people, whether it be patients, participants, test volunteers or experimental stimuli, because verification of their consent is incompatible with the rapid and automated nature of preprint posting”.

For some of the image categories, there is a high prevalence of relating images to both methamphetamine and opioid (“both” response in the relatedness question) (82% in the “instruments and hands” category in the methamphetamine images and 58% for “drugs and hands” category in the opioid images). The methamphetamine (drug alone) image category received −.94 on average in MethToOpioid (defined in the data analysis section (2.d) and the opioid category received 0.64. Sample images with the least differentiation between methamphetamine and opioids (MethToOpioid close to 0) are presented in figure 4.

As depicted in figure 5, craving and arousal have a highly positively correlation in both methamphetamine (Pearson correlation coefficient (R)=0.74) and opioid (R=0.66) images without significant difference between methamphetamine and opioid images. Craving and valence are negatively correlated in opioid (R=−0.43) images (for methamphetamine R=−0.16, p-value=0.08) and opioid images have significantly higher negative correlation between craving and valence compared to methamphetamine images (difference in Z scores between methamphetamine and opioid=2.27) after FDR correction (Figure 5).

**Figure 5.**
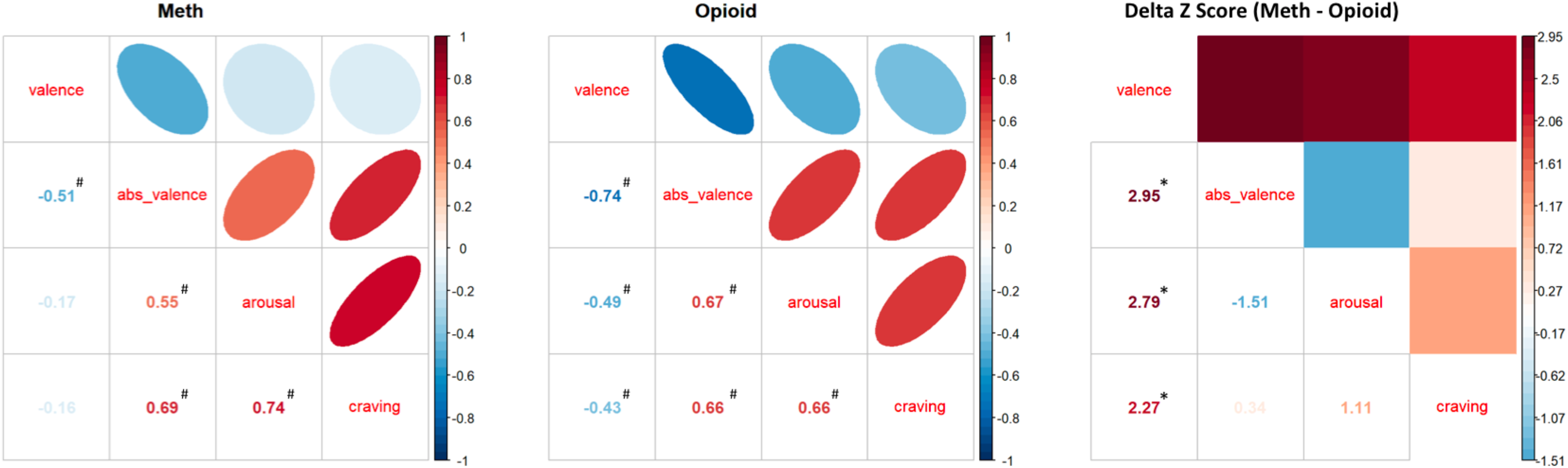
Correlation matrix between craving, valence, absolute valence and arousal. Right matrix represents difference in Z values between meth and opioid images. Values with * and ^#^ passed 0.05 and 0.0001 FDR corrected p value threshold respectively.

There is a 53% and 25% shared variance between absolute valence (as defined in the “data analysis” (2.d) section and valence in opioid and methamphetamine images (opioid: R=−0.73; meth: R=−0.504; control: R=−0.0003, p-value=0.99) (Figure S3). Absolute valence is positively correlated with craving in both methamphetamine (R=0.69) and opioid (R=0.66) images without any significant difference between methamphetamine and opioid images.

### c. Response Durations

Participants spent 79.8 (27.5) and 59.7 (18) minutes in session one and two, 26.62 (9.18) and 19.90 (6) seconds per image. After removing datapoints with over 100 seconds spent for image completion (total rating time) (1% of datapoints) or 20 seconds for craving rating time (0.6% of datapoints), participants rated meth and opioid images 253 and 376 milliseconds slower than neutral images (p<0.001 and p<0.001), which is an increase of about 10 percent. They also rated opioid images significantly slower than methamphetamine images (123 millisecond, p-value=0.0096). Interestingly, there is a significant positive correlation between craving rating and both the time spent for the craving rating questions (craving rating time) and the time spent to answer all questions related to each image (rating completion time) (Figure 6). After controlling for drug type, time, visit, and the time by visit interaction, every 10 points of craving were associated with an increased response time of 383 millisecond (p-value<0.001).

**Figure 6.**
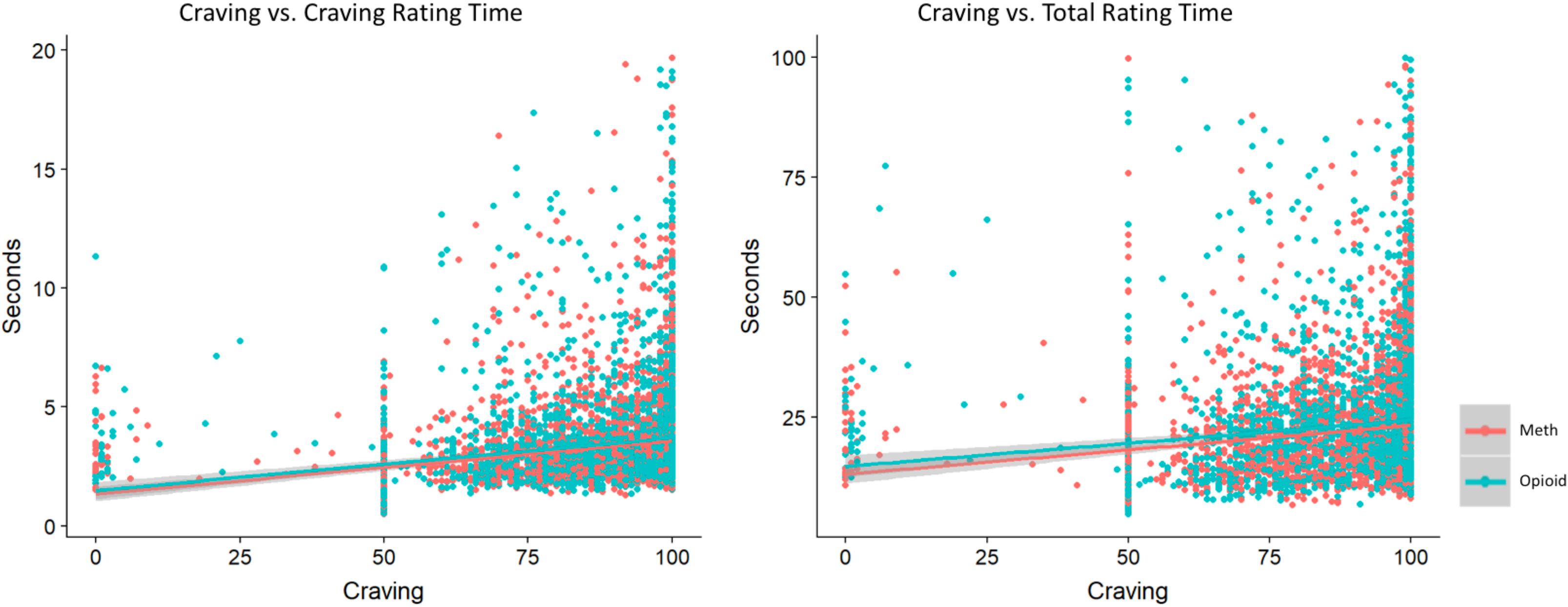
Correlation between craving rating and time spent for rating. Left scatterplot represents time spent for the craving rating questions (craving rating time) and the right scatterplot represents time spent to answer all questions related to each image (rating completion time) after removing datapoints with over 100 seconds spent for completion (1% of datapoints) or 20 seconds for craving rating (0.06% of datapoints). This association will remain significant if we add time, session, sex and age of the participants in the model (further details in figure S2).

There were significant practice or fatigue effects for both craving ratings and response times. For craving ratings, there were negative effects for time among both meth (beta=−0.02, p-value=0.0026) and opioid (beta=−0.38, p-value<0.001) images, meaning people rated images as less craving-inducing over time. For opioid craving ratings, there was a main effect of visit (beta=−3.3, p-value<0.001), indicating overall lower ratings in the second session, while for meth images there was a negative visit by time interaction (beta=−0.0453, p-value<0.001), meaning the decrease in rating over time was greater in the second session. Total image completion times and craving rating reaction times were negatively correlated with both visit and time for both meth and opioid images, indicating that within each session subjects responded more quickly as time went on, and that they responded overall faster in the second session (all p-value<0.001 for all tests).

### d. Equivalent fMRI Tasks

As we described in the methods, data analysis section, we extracted 3 equivalent image sets, 72 images each (3 subsets with 24 methamphetamine, 24 opioid and 24 control (neutral) images) from the LIBR MOCD. Each set is selected for two fMRI tasks, one for methamphetamine and one for opioids, with 4 drug related blocks (6 images for each block, one image from each category) and 4 neutral (control) blocks. There is no significant difference between blocks within each sub set in terms of craving, valence, arousal and physical features (value, hue and contrast) (Tables S2, S3 and S4). Among the 3 equivalent image sets, there is no significant difference between methamphetamine image sub-sets for craving, valence, arousal and physical features (value, hue and contrast) (Table S5). This is true between opioid image subsets and also between neutral (control) image sets as well (Table S5). Furthermore, there is no significant difference in physical features (value, hue and contrast) between drug and neutral subsets within each set (Table S6). Images in each set are provided in Figures 7, S4 and S5). The list of selected images for each sets, subsets and blocks are provided in supplementary material 5. The Psychopy codes for these 6 fMRI tasks are also provided in this link (https://github.com/rkuplicki/LIBR_MOCD).

**Figure 7.**
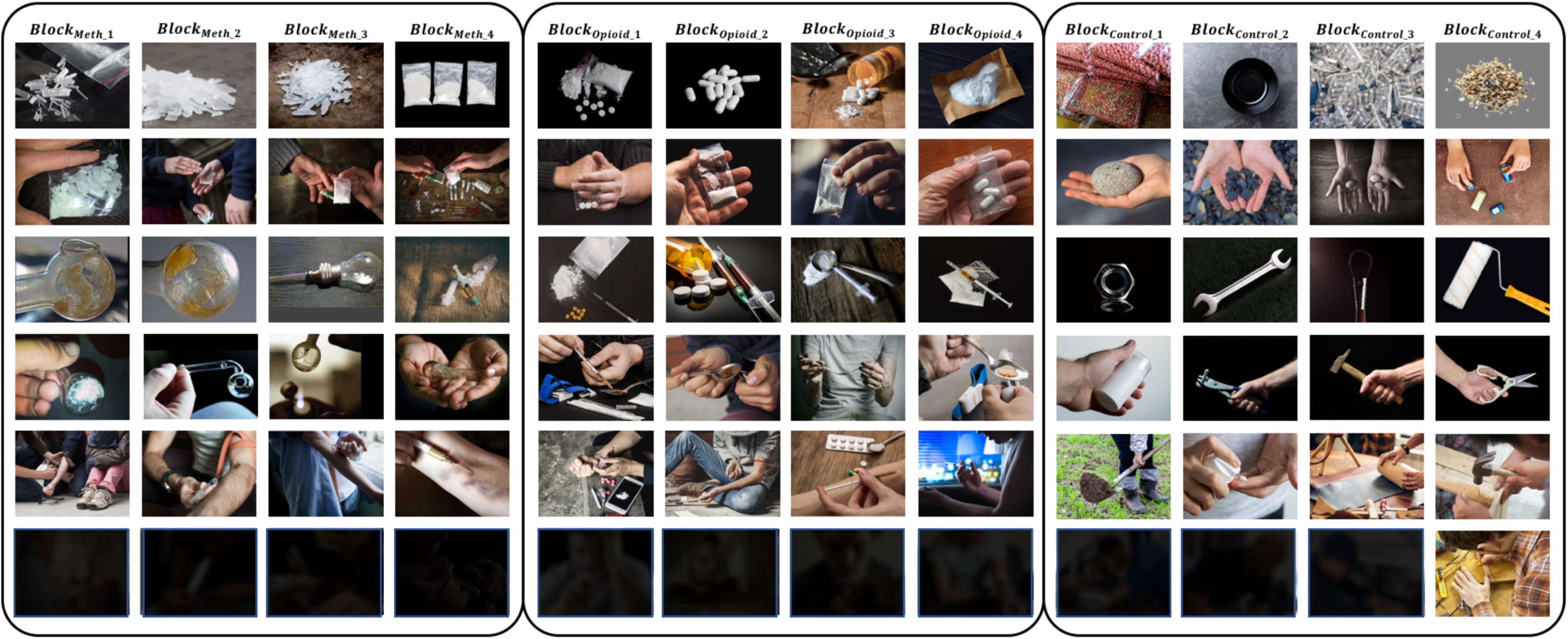
The first image set (methamphetamine, opioid and control (neutral) subsets) from 3 equivalent image sets extracted from LIBR MOCD. There is no significant difference between blocks within each sub set in terms of craving, valence, arousal and physical features (value, hue and contrast) (Tables S2, S3 and S4). Between the 3 equivalent image sets, there is no significant difference between methamphetamine image sub-sets for craving, valence, arousal and physical features (value, hue and contrast). This is true between opioid image subsets and also between neutral (control) image sets as well (Table S5). Furthermore, there is no significant difference in physical features (value, hue and contrast) between drug and neutral subsets within each set (Table S6). We removed few elements in this figure in this version of the manuscript to respect biorxiv’s policy “to avoid the inclusion of photographs and any other identifying information of people, whether it be patients, participants, test volunteers or experimental stimuli, because verification of their consent is incompatible with the rapid and automated nature of preprint posting”.

## 4. Discussion

Drug cue reactivity (DCR) paradigm provides unique opportunities to examine response to conditioned cues in a controlled experimental environment. Ecological validity and feasibility of the pictorial DCR even in extreme experimental settings, like inside an fMRI scanner, makes pictorial DCR a common practice in addiction neuroscience research in human subjects (Ekhtiari, Nasseri, et al., 2016). But there is a significant variability in pictorial cues that researchers have recruited thus far (Ekhtiari, Faghiri, et al., 2016). The LIBR Methamphetamine and Opioid Cue Database (LIBR MOCD) can help to reduce this variability and increase harmony and replicability in future studies. As presented in this article, LIBR MOC database can be used as a source to select equivalent sets of drug cues and their neutral controls based on both psychological and physical characteristics for multiple assessments/interventions. With images that relate to both methamphetamine and opioid use (MethToOpioid score close to 0), researchers may consider designing a task that can be used in methamphetamine and/or opioid users.

There was no significant association between clinical features and craving reports, time spent for craving and time spent for all ratings for images. In this study, the participants were asked to rate how much the images can induce drug craving in an “active methamphetamine or opioid user”. This might explain the lack of correlation between “clinical features of participants” and their rating on how much they expect the picture can induce craving in an “active methamphetamine or opioid user”. There is also no significant change in craving reported after seeing pictorial cues (induced craving) among our participants. Our participants were successfully abstinent from methamphetamine and opioids for more than 2 years on average and the lack of craving self-report seems reasonable.

In this study, there are small but interesting differences between methamphetamine and opioid images in terms of the relationship between craving, valence and arousal that needs further exploration. Opioid drugs with narcotic effects and significant withdrawal syndrome compared to methamphetamine with stimulant effects without a serious withdrawal syndrome can provide a background for this difference. Variation in both subjective and objective (including both neural and peripheral) conditioned response to methamphetamine and opioid drugs may predict variations in response to different interventions.

There is an interesting relationship, within the drug related images, between the craving report and the time participants spent on each image. After controlling for drug type, time, visit, and the time by visit interaction, every 10 points of craving were associated with an increased response time of 383milliseconds. Spending more time and utilizing more zoom-in features for salient images (food, drug, sex and etc.) has been well explained previously in the approach-avoidance paradigm describing cognitive bias (Gladwin, Wiers, & Wiers, 2016). Our results suggest that the amount of time spent with images might be a more sensitive measure for craving than subjective self-report after the cue exposure.

This study has several limitations. (1) We recruited people with history of methamphetamine and opioid use (75% injection history of opioids or methamphetamine) successfully abstinent for more than 6 months (2 years in average). Participants were asked to rate how much craving the images can induce in an active methamphetamine or opioid user. Exploring potential differences in response among people with shorter duration of abstinence and even while they are still active users or during different treatment modalities like methadone maintenance will be interesting. We have not found significantly different craving report for images between people with or without injection history (supplementary material 1). However, further exploration on the variation within methamphetamine and opioid users (injection or not, pill versus heroin and etc.) can be interesting. (2) There is a significant effect of time, session number, and their interaction on reported craving ratings as participants reported lower craving over time. It is hard to differentiate the effect of habituation from simple fatigue due to a long rating task. There are negative results with initial attempts for desensitization interventions with cue exposure in drug addiction (Marissen, Franken, Blanken, van den Brink, & Hendriks, 2007), but a large pictorial database like LIBR MOCD can provide a great resource for future cue exposure interventions with multiple sessions including more precise measures of habituation/desensitization. Recent successful results with memory reconsolidation interventions (Germeroth et al., 2017; Xue et al., 2017) can provide experimental insights for future cue exposure studies in combination with pharmacological/neuromodulation/cognitive interventions. (3) Figures 109 and 118 in the opioid set are identical by error. So, we ended up having 119 unique opioid images instead of 120 (Figure S1). Furthermore, we have 6 images with written text among the opioid images (bottles of pills and boxes of pills) and 4 among neutral images. We avoided using them in the image sets for the fMRI tasks to control for brain activations related to language/reading, and (4) We haven’t done any test retest study in this database. Furthermore, validation with other subjective or objective measures and recruitment of healthy controls will increase our understanding about the psychometric characteristics of the LIBR MOCD.

LIBR MOCD is one of the first attempts to provide a large database of validated pictorial cues for methamphetamine, opioids and neutral images. We hope this study and its associated database, codes and resources will help addiction scientists to design future assessments and interventions for DCR. Making a consensus for optimum fMRI tasks using this database will help us to share our data within larger databases to be used for development of biomarkers from DCR. Harmonized DCR fMRI tasks will be an important step towards bringing functional neuroimaging to the daily clinical practice in addiction medicine.

## Acknowledgements

Authors would like to thank Kyle Rogers for designing the computerized web-based version of the rating task. Authors also highly appreciated scientific advices from Rajita Sinha and Dongju Seo in designing the study and feedbacks from Peyman Ghobadi Azbari in the first draft of the manuscript.

---

**Supplementary Table 1.**
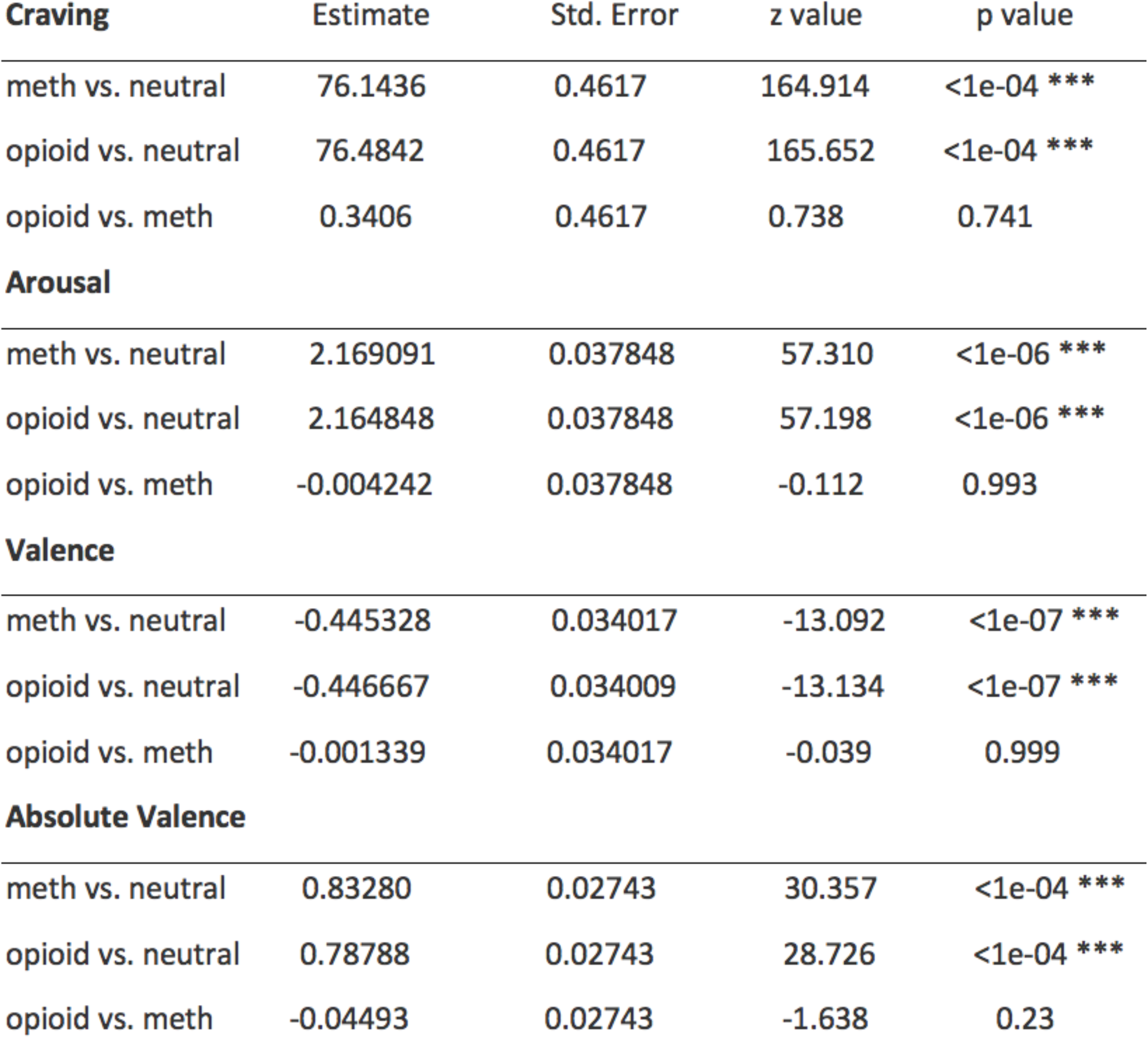
Post hoc Tukey test for linear mixed effect models including ratings (craving, arousal, valence, and absolute valence) as dependent variables and age, sex and drug types as dependent variables.

**Supplementary Table 2.** The first image set (methamphetamine, opioid and control (neutral) subsets) from 3 equivalent image sets extracted from LIBR MOCD. No significant difference in arousal, craving, typicality, valence, hue, saturation, and value between blocks within the image set 0 (reported p values are uncorrected).

| Variable | Between Sets | Block_1<br>(N=6) | Block_2<br>(N=6) | Block_3<br>(N=6) | Block_4<br>(N=6) | F | P-value |
| --- | --- | --- | --- | --- | --- | --- | --- |
|  |  | Mean±SD | Mean±SD | Mean±SD | Mean±SD |  |  |
| **Arousal** | *Methamphetamine Task_0* | 4.69±0.27 | 4.66±0.18 | 4.71±0.29 | 4.78±0.15 | 0.33 | 0.80 |
|  | *Opioid Task_0* | 4.77±0.26 | 4.66±0.17 | 4.77±0.28 | 4.56±0.20 | 1.10 | 0.37 |
|  | *Control Task_0* | 2.65±0.28 | 2.48±0.27 | 2.58±0.13 | 2.29±0.22 | 2.67 | 0.08 |
| **Craving** | *Methamphetamine Task_0* | 87.71±2.53 | 87.45±2.58 | 87.21±2.81 | 87.04±3.54 | 0.06 | 0.98 |
|  | *Opioid Task_0* | 87.96±3.15 | 89.67±1.17 | 90.44±3.73 | 87.36±4.15 | 1.17 | 0.35 |
|  | *Control Task_0* | 11.61±4.86 | 11.41±1.62 | 11.11±2.38 | 10.54±5.09 | 0.09 | 0.96 |
| **Typicality** | *Methamphetamine Task_0* | 88.98±2.92 | 88.00±2.75 | 89.01±2.55 | 87.37±3.74 | 0.42 | 0.74 |
|  | *Opioid Task_0* | 90.45±3.23 | 90.79±1.42 | 91.04±3.95 | 88.02±2.48 | 1.35 | 0.29 |
|  | *Control Task_0* | 12.54±5.67 | 13.32±1.74 | 12.51±2.60 | 9.87±1.43 | 1.24 | 0.32 |
| **Valence** | *Methamphetamine Task_0* | 4.45±0.25 | 4.33±0.22 | 4.48±0.20 | 4.40±0.34 | 0.42 | 0.74 |
|  | *Opioid Task_0* | 4.39±0.26 | 4.35±0.29 | 4.46±0.22 | 4.42±0.19 | 0.21 | 0.89 |
|  | *Control Task_0* | 4.98±0.22 | 4.90±0.10 | 4.85±0.10 | 4.77±0.13 | 2.39 | 0.10 |
| **Hue** | *Methamphetamine Task_0* | 0.30±0.10 | 0.29±0.12 | 0.23±0.14 | 0.16±0.07 | 2.27 | 0.11 |
|  | *Opioid Task_0* | 0.20±0.06 | 0.16±0.12 | 0.22±0.16 | 0.32±0.21 | 1.30 | 0.30 |
|  | *Control Task_0* | 0.31±0.11 | 0.35±0.19 | 0.28±0.24 | 0.17±0.23 | 0.86 | 0.48 |
| **Saturation** | *Methamphetamine Task_0* | 0.23±0.08 | 0.26±0.12 | 0.33±0.14 | 0.34±0.17 | 1.03 | 0.40 |
|  | *Opioid Task_0* | 0.17±0.08 | 0.22±0.11 | 0.32±0.11 | 0.30±0.14 | 2.34 | 0.10 |
|  | *Control Task_0* | 0.32±0.12 | 0.22±0.08 | 0.22±0.10 | 0.28±0.19 | 0.83 | 0.49 |
| **Value** | *Methamphetamine Task_0* | 0.42±0.06 | 0.37±0.15 | 0.35±0.08 | 0.30±0.11 | 1.20 | 0.33 |
|  | *Opioid Task_0* | 0.34±0.12 | 0.35±0.13 | 0.43±0.11 | 0.42±0.16 | 0.67 | 0.58 |
|  | *Control Task_0* | 0.47±0.22 | 0.36±0.20 | 0.38±0.24 | 0.47±0.20 | 0.42 | 0.74 |

**Supplementary Table 3.** The second image set (methamphetamine, opioid and control (neutral) subsets) from 3 equivalent image sets extracted from LIBR MOCD. No significant difference in arousal, craving, typicality, valence, hue, saturation, and value between blocks within the image set 1 (reported p values are uncorrected).

| Variable | Between Sets | Block_1<br>(N=6) | Block_2<br>(N=6) | Block_3<br>(N=6) | Block_4<br>(N=6) | F | P-value |
| --- | --- | --- | --- | --- | --- | --- | --- |
|  |  | Mean±SD | Mean±SD | Mean±SD | Mean±SD |  |  |
| **Arousal** | *Methamphetamine Task_1* | 4.77±0.16 | 4.80±0.18 | 4.76±0.21 | 4.64±0.09 | 1.08 | 0.38 |
|  | *Opioid Task_1* | 4.68±0.28 | 4.61±0.17 | 4.66±0.36 | 4.64±0.22 | 0.09 | 0.97 |
|  | *Control Task_1* | 2.73±0.38 | 2.42±0.23 | 2.51±0.26 | 2.56±0.23 | 1.32 | 0.29 |
| **Craving** | *Methamphetamine Task_1* | 90.32±2.42 | 90.21±3.94 | 88.23±3.72 | 88.26±4.41 | 0.60 | 0.62 |
|  | *Opioid Task_1* | 90.13±3.79 | 90.54±2.17 | 86.71±6.56 | 87.50±2.19 | 1.29 | 0.30 |
|  | *Control Task_1* | 12.23±3.57 | 10.50±3.90 | 10.50±2.93 | 13.24±4.45 | 0.78 | 0.52 |
| **Typicality** | *Methamphetamine Task_1* | 90.01±2.43 | 91.53±3.08 | 89.38±4.21 | 90.27±3.86 | 0.40 | 0.75 |
|  | *Opioid Task_1* | 91.97±3.84 | 90.84±2.13 | 87.15±7.12 | 87.33±1.56 | 1.99 | 0.15 |
|  | *Control Task_1* | 13.46±4.26 | 11.20±2.80 | 10.66±2.66 | 11.45±1.55 | 1.01 | 0.41 |
| **Valence** | *Methamphetamine Task_1* | 4.33±0.09 | 4.49±0.22 | 4.43±0.10 | 4.36±0.14 | 1.55 | 0.23 |
|  | *Opioid Task_1* | 4.47±0.33 | 4.44±0.16 | 4.47±0.30 | 4.55±0.31 | 0.18 | 0.91 |
|  | *Control Task_1* | 4.88±0.09 | 4.84±0.21 | 4.88±0.22 | 5.02±0.14 | 1.35 | 0.29 |
| **Hue** | *Methamphetamine Task_1* | 0.23±0.10 | 0.39±0.22 | 0.32±0.19 | 0.27±0.21 | 0.82 | 0.49 |
|  | *Opioid Task_1* | 0.24±0.08 | 0.21±0.06 | 0.19±0.09 | 0.32±0.14 | 1.98 | 0.15 |
|  | *Control Task_1* | 0.27±0.17 | 0.26±0.17 | 0.15±0.17 | 0.20±0.23 | 0.55 | 0.65 |
| **Saturation** | *Methamphetamine Task_1* | 0.27±0.12 | 0.24±0.10 | 0.31±0.16 | 0.37±0.13 | 1.30 | 0.30 |
|  | *Opioid Task_1* | 0.26±0.08 | 0.24±0.16 | 0.33±0.13 | 0.27±0.14 | 0.55 | 0.66 |
|  | *Control Task_1* | 0.34±0.17 | 0.25±0.21 | 0.22±0.15 | 0.21±0.16 | 0.68 | 0.58 |
| **Value** | *Methamphetamine Task_1* | 0.47±0.12 | 0.39±0.15 | 0.55±0.17 | 0.32±0.16 | 2.66 | 0.07 |
|  | *Opioid Task_1* | 0.35±0.07 | 0.36±0.12 | 0.44±0.19 | 0.55±0.22 | 1.88 | 0.16 |
|  | *Control Task_1* | 0.28±0.14 | 0.45±0.10 | 0.34±0.26 | 0.50±0.24 | 1.52 | 0.24 |

**Supplementary Table 4.** The third image set (methamphetamine, opioid and control (neutral) subsets) from 3 equivalent image sets extracted from LIBR MOCD. No significant difference in arousal, craving, typicality, valence, hue, saturation, and value between blocks within the image set 2 (reported p values are uncorrected).

| Variable | Between Sets | Block_1<br>(N=6) | Block_2<br>(N=6) | Block_3<br>(N=6) | Block_4<br>(N=6) | F | P-value |
| --- | --- | --- | --- | --- | --- | --- | --- |
|  |  | Mean±SD | Mean±SD | Mean±SD | Mean±SD |  |  |
| **Arousal** | *Methamphetamine Task_2* | 4.71±0.10 | 4.77±0.19 | 4.60±0.21 | 4.72±0.13 | 1.27 | 0.31 |
|  | *Opioid Task_2* | 4.38±0.20 | 4.73±0.26 | 4.70±0.28 | 4.67±0.21 | 2.68 | 0.07 |
|  | *Control Task_2* | 2.58±0.23 | 2.60±0.21 | 2.41±0.11 | 2.58±0.29 | 1.05 | 0.39 |
| **Craving** | *Methamphetamine Task_2* | 88.35±3.14 | 89.12±2.41 | 88.09±2.98 | 90.43±2.37 | 0.88 | 0.47 |
|  | *Opioid Task_2* | 85.91±4.44 | 87.68±2.50 | 89.53±3.82 | 87.29±3.11 | 1.06 | 0.39 |
|  | *Control Task_2* | 10.92±2.56 | 10.47±2.32 | 13.58±3.49 | 16.39±12.99 | 0.93 | 0.44 |
| **Typicality** | *Methamphetamine Task_2* | 89.61±2.67 | 91.28±2.92 | 90.29±3.49 | 90.65±3.15 | 0.31 | 0.82 |
|  | *Opioid Task_2* | 87.43±4.28 | 89.65±1.96 | 91.01±3.69 | 88.84±2.23 | 1.33 | 0.29 |
|  | *Control Task_2* | 11.40±2.17 | 11.40±2.80 | 12.13±1.72 | 16.75±12.55 | 0.92 | 0.45 |
| **Valence** | *Methamphetamine Task_2* | 4.39±0.14 | 4.52±0.11 | 4.47±0.14 | 4.37±0.26 | 0.94 | 0.44 |
|  | *Opioid Task_2* | 4.50±0.24 | 4.45±0.25 | 4.35±0.27 | 4.46±0.34 | 0.33 | 0.80 |
|  | *Control Task_2* | 4.91±0.14 | 4.82±0.17 | 4.93±0.02 | 4.94±0.12 | 1.09 | 0.38 |
| **Hue** | *Methamphetamine Task_2* | 0.27±0.05 | 0.27±0.17 | 0.28±0.16 | 0.24±0.11 | 0.09 | 0.96 |
|  | *Opioid Task_2* | 0.22±0.11 | 0.24±0.06 | 0.24±0.18 | 0.28±0.14 | 0.22 | 0.88 |
|  | *Control Task_2* | 0.30±0.13 | 0.21±0.16 | 0.20±0.15 | 0.16±0.13 | 1.08 | 0.38 |
| **Saturation** | *Methamphetamine Task_2* | 0.28±0.11 | 0.28±0.13 | 0.32±0.15 | 0.40±0.05 | 1.50 | 0.24 |
|  | *Opioid Task_2* | 0.27±0.09 | 0.37±0.09 | 0.23±0.17 | 0.32±0.19 | 1.13 | 0.36 |
|  | *Control Task_2* | 0.31±0.11 | 0.31±0.08 | 0.22±0.07 | 0.22±0.15 | 1.24 | 0.32 |
| **Value** | *Methamphetamine Task_2* | 0.48±0.12 | 0.38±0.09 | 0.33±0.18 | 0.31±0.18 | 1.51 | 0.24 |
|  | *Opioid Task_2* | 0.42±0.07 | 0.33±0.15 | 0.37±0.20 | 0.50±0.18 | 1.28 | 0.31 |
|  | *Control Task_2* | 0.38±0.16 | 0.47±0.17 | 0.47±0.22 | 0.50±0.25 | 0.43 | 0.73 |

**Supplementary Table 5.**
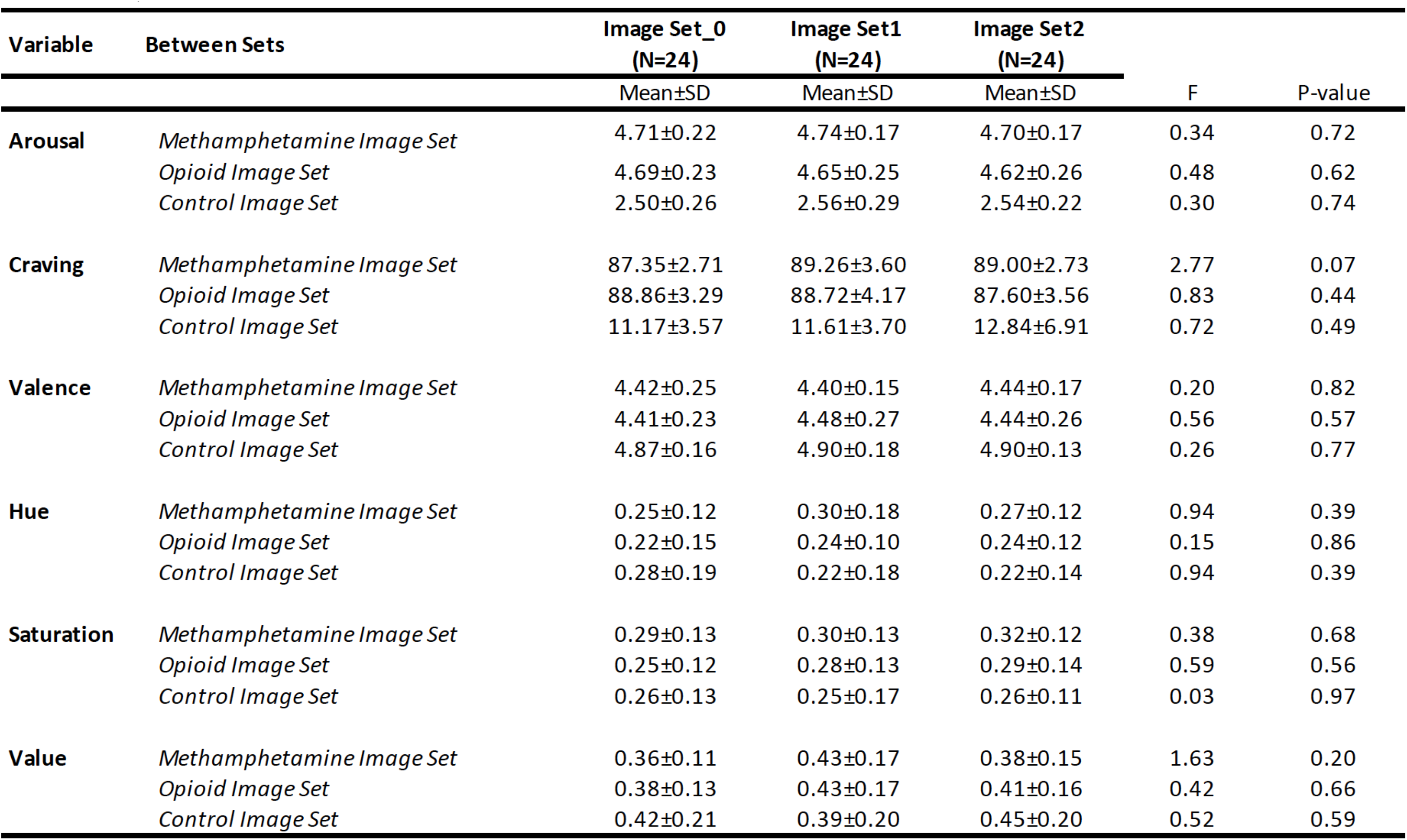
Testing equivalency between sets. No significant difference in arousal, craving, typicality, valence, hue, saturation, and value between methamphetamine, opioid, and control image subsets between three equivalent image sets (reported p values are uncorrected).

| Variable | Between Sets | Image Set_0<br>(N=24) | Image Set1<br>(N=24) | Image Set2<br>(N=24) | F | P-value |
| --- | --- | --- | --- | --- | --- | --- |
|  |  | Mean±SD | Mean±SD | Mean±SD |  |  |
| Arousal | *Methamphetamine Image Set* | 4.71±0.22 | 4.74±0.17 | 4.70±0.17 | 0.34 | 0.72 |
|  | *Opioid Image Set* | 4.69±0.23 | 4.65±0.25 | 4.62±0.26 | 0.48 | 0.62 |
|  | *Control Image Set* | 2.50±0.26 | 2.56±0.29 | 2.54±0.22 | 0.30 | 0.74 |
| Craving | *Methamphetamine Image Set* | 87.35±2.71 | 89.26±3.60 | 89.00±2.73 | 2.77 | 0.07 |
|  | *Opioid Image Set* | 88.86±3.29 | 88.72±4.17 | 87.60±3.56 | 0.83 | 0.44 |
|  | *Control Image Set* | 11.17±3.57 | 11.61±3.70 | 12.84±6.91 | 0.72 | 0.49 |
| Valence | *Methamphetamine Image Set* | 4.42±0.25 | 4.40±0.15 | 4.44±0.17 | 0.20 | 0.82 |
|  | *Opioid Image Set* | 4.41±0.23 | 4.48±0.27 | 4.44±0.26 | 0.56 | 0.57 |
|  | *Control Image Set* | 4.87±0.16 | 4.90±0.18 | 4.90±0.13 | 0.26 | 0.77 |
| Hue | *Methamphetamine Image Set* | 0.25±0.12 | 0.30±0.18 | 0.27±0.12 | 0.94 | 0.39 |
|  | *Opioid Image Set* | 0.22±0.15 | 0.24±0.10 | 0.24±0.12 | 0.15 | 0.86 |
|  | *Control Image Set* | 0.28±0.19 | 0.22±0.18 | 0.22±0.14 | 0.94 | 0.39 |
| Saturation | *Methamphetamine Image Set* | 0.29±0.13 | 0.30±0.13 | 0.32±0.12 | 0.38 | 0.68 |
|  | *Opioid Image Set* | 0.25±0.12 | 0.28±0.13 | 0.29±0.14 | 0.59 | 0.56 |
|  | *Control Image Set* | 0.26±0.13 | 0.25±0.17 | 0.26±0.11 | 0.03 | 0.97 |
| Value | *Methamphetamine Image Set* | 0.36±0.11 | 0.43±0.17 | 0.38±0.15 | 1.63 | 0.20 |
|  | *Opioid Image Set* | 0.38±0.13 | 0.43±0.17 | 0.41±0.16 | 0.42 | 0.66 |
|  | *Control Image Set* | 0.42±0.21 | 0.39±0.20 | 0.45±0.20 | 0.52 | 0.59 |

**Supplementary Table 6.** Testing difference in psychophysics characteristics between image subsets in each set. No significant difference in valence, hue, saturation, and value between methamphetamine, opioid, and control subsets within 3 image sets (reported p values are uncorrected).

| Variable | Image Set | Control<br>(N=24) | Meth<br>(N=24) | Opioid<br>(N=24) | F | P-value |
| --- | --- | --- | --- | --- | --- | --- |
|  |  | Mean±SD | Mean±SD | Mean±SD |  |  |
| Hue | *Image Set_0* | 0.28±0.19 | 0.25±0.12 | 0.22±0.15 | 0.67 | 0.51 |
|  | *Image Set_1* | 0.22±0.18 | 0.30±0.18 | 0.24±0.10 | 1.72 | 0.19 |
|  | *Image Set_2* | 0.22±0.14 | 0.27±0.12 | 0.24±0.12 | 0.92 | 0.40 |
| Saturation | *Image Set_0* | 0.26±0.13 | 0.29±0.13 | 0.25±0.12 | 0.51 | 0.60 |
|  | *Image Set_1* | 0.25±0.17 | 0.30±0.13 | 0.28±0.13 | 0.57 | 0.57 |
|  | *Image Set_2* | 0.26±0.11 | 0.32±0.12 | 0.29±0.14 | 1.24 | 0.30 |
| Value | *Image Set_0* | 0.42±0.21 | 0.36±0.11 | 0.38±0.13 | 0.81 | 0.45 |
|  | *Image Set_1* | 0.39±0.20 | 0.43±0.17 | 0.43±0.17 | 0.34 | 0.71 |
|  | *Image Set_2* | 0.45±0.20 | 0.38±0.15 | 0.41±0.16 | 1.12 | 0.33 |

**Supplementary Figure 1.**
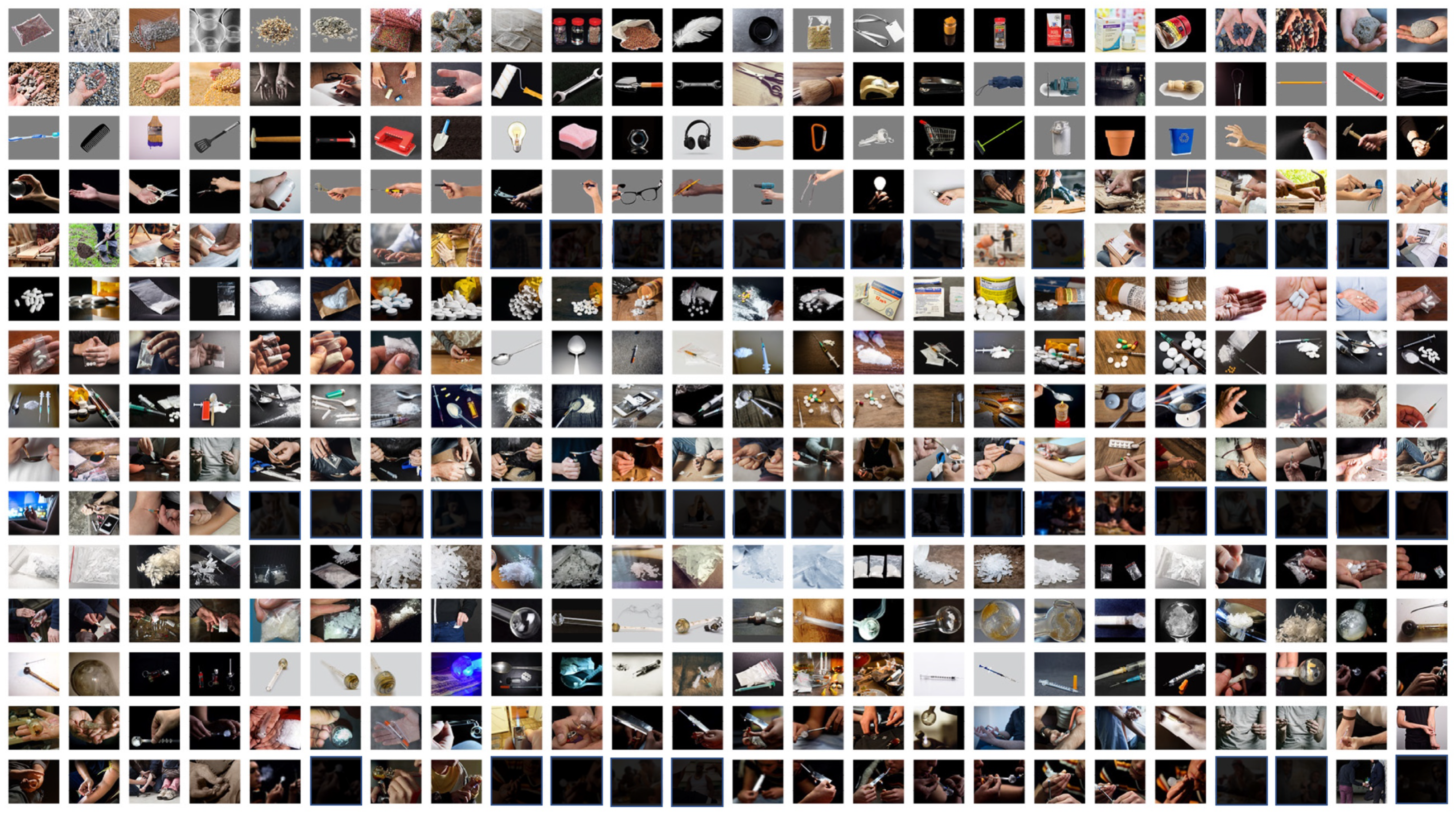
Three hundred and sixty images in the database. Figures 109 and 118 in the opioid set are identical by error. So, we ended up having 119 unique opioid images instead of 120. We removed few elements in this figure in this version of the manuscript to respect biorxiv’s policy “to avoid the inclusion of photographs and any other identifying information of people, whether it be patients, participants, test volunteers or experimental stimuli, because verification of their consent is incompatible with the rapid and automated nature of preprint posting”.

**Supplementary Figure 2.**
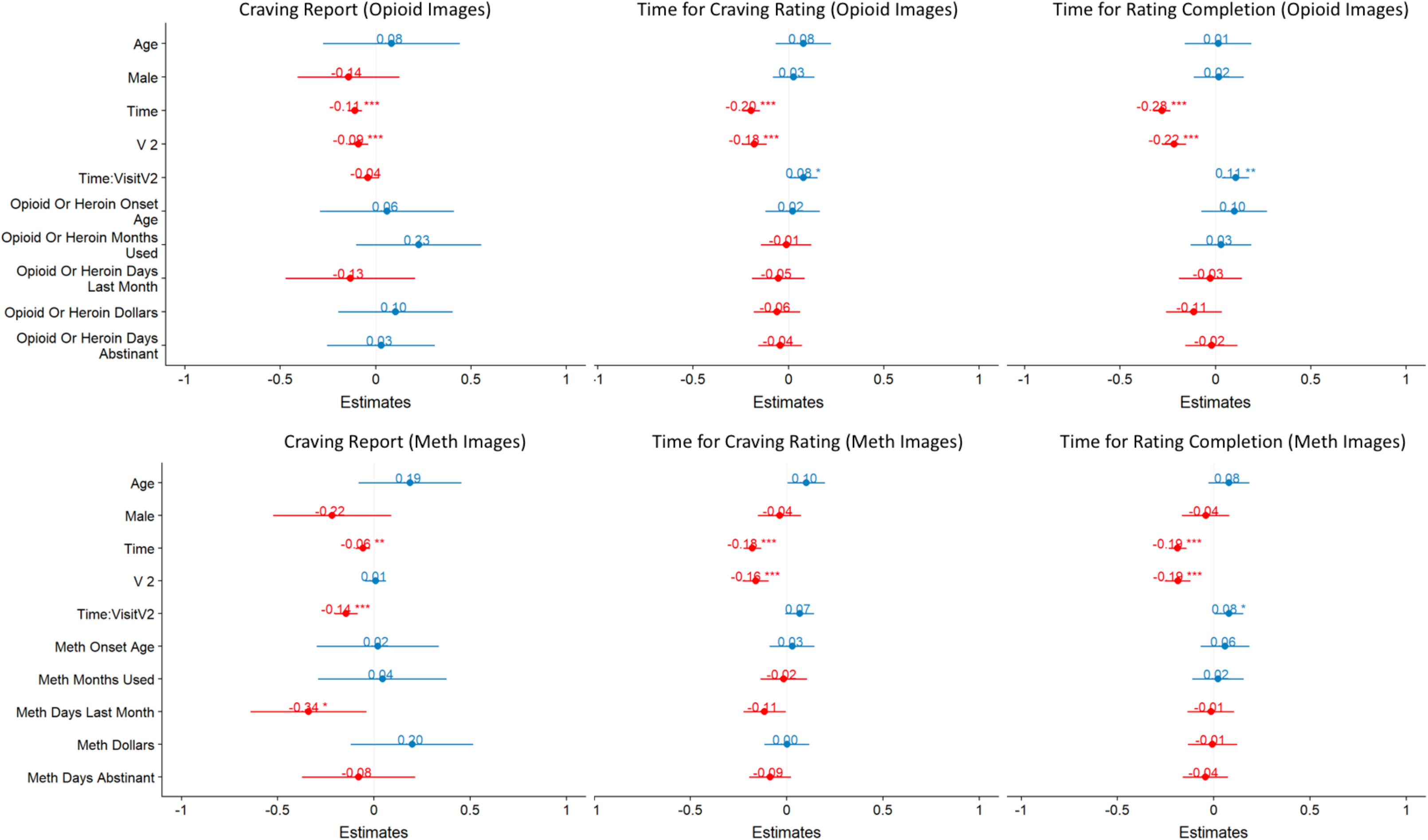
Relationship between demographic and clinical variables and craving self-report, time spent for craving rating and time spent for all ratings (completion time) controlled by age, sex, presentation time and session (visit: V1 and V2). Linear mixed effect model with mean craving reports and time variables as the dependent variable. Standardized beta coefficients are reported in Y axis. None of the demographic and clinical variables will pass the multiple comparison corrected p value threshold of 0.05. There is a significant reduction in craving report over time (standardized estimate in the LME model = − 0.11, p value<0.001 for opioids and −0.06 p value<0.01 for methamphetamine). The slope of this negative effect is more negative in the second session in meth images (time: visitV2 interaction).

**Supplementary Figure 3.**
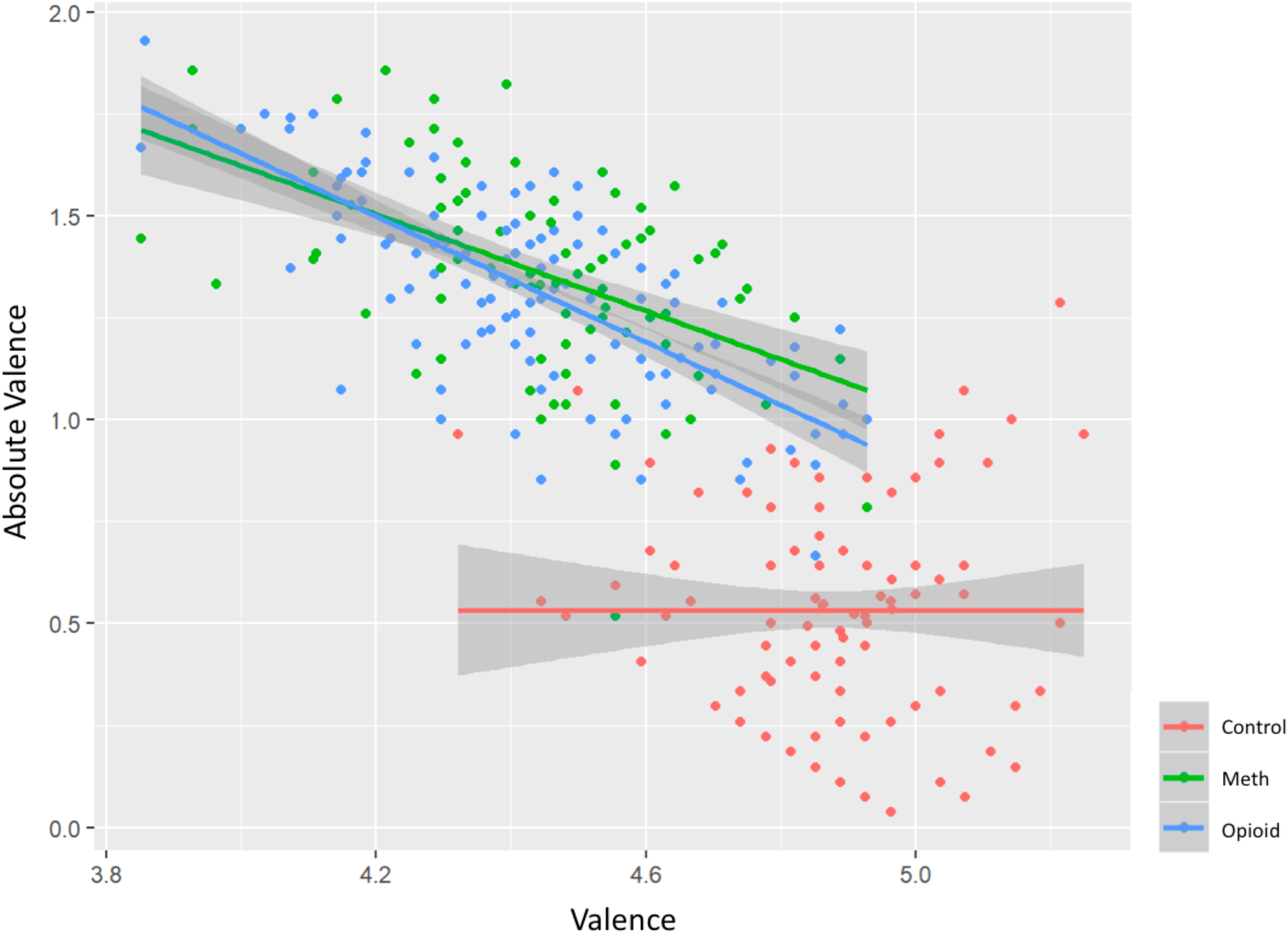
Scatterplot between valence and absolute valence (distance to mean, 5: neutral valence) for each picture.

**Supplementary Figure 4.**
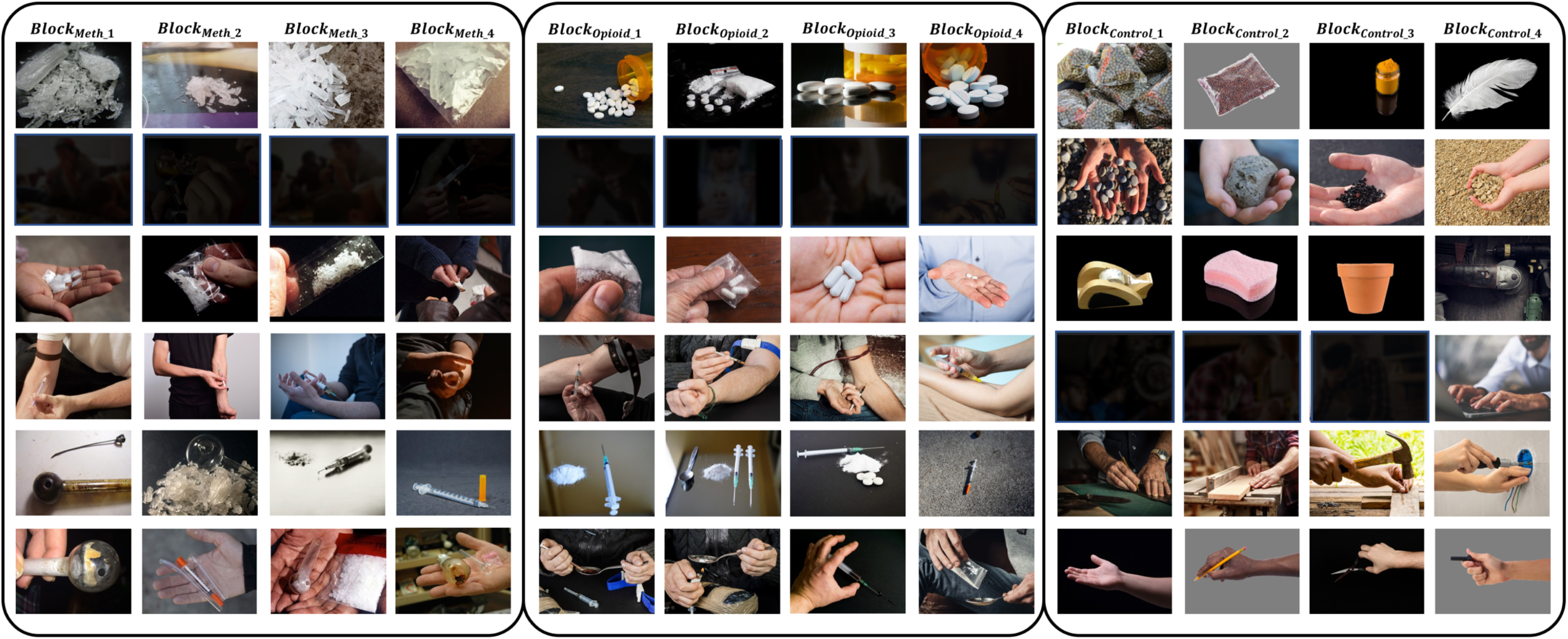
The second image set (methamphetamine, opioid and control (neutral) subsets) from 3 equivalent image sets extracted from LIBR MOCD. We removed few elements in this figure in this version of the manuscript to respect biorxiv’s policy “to avoid the inclusion of photographs and any other identifying information of people, whether it be patients, participants, test volunteers or experimental stimuli, because verification of their consent is incompatible with the rapid and automated nature of preprint posting”.

**Supplementary Figure 5.**
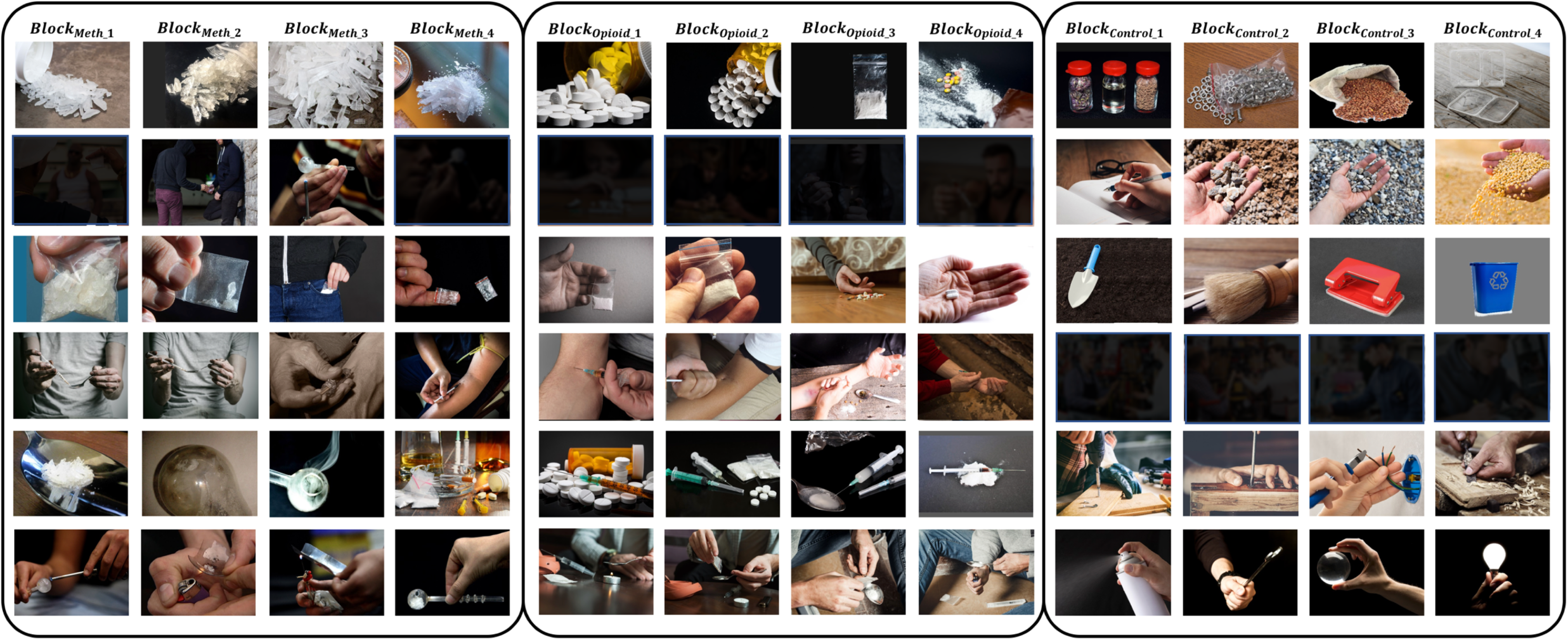
The third image set (methamphetamine, opioid and control (neutral) subsets) from 3 equivalent image sets extracted from LIBR MOCD. We removed few elements in this figure in this version of the manuscript to respect biorxiv’s policy “to avoid the inclusion of photographs and any other identifying information of people, whether it be patients, participants, test volunteers or experimental stimuli, because verification of their consent is incompatible with the rapid and automated nature of preprint posting”.

