## Supplementary Material 2 for "LIBR Methamphetamine and Opioid Cue Database (LIBR MOCD): Development and Validation"

### Meth and Opioid Cue Database

#### Neutral Image

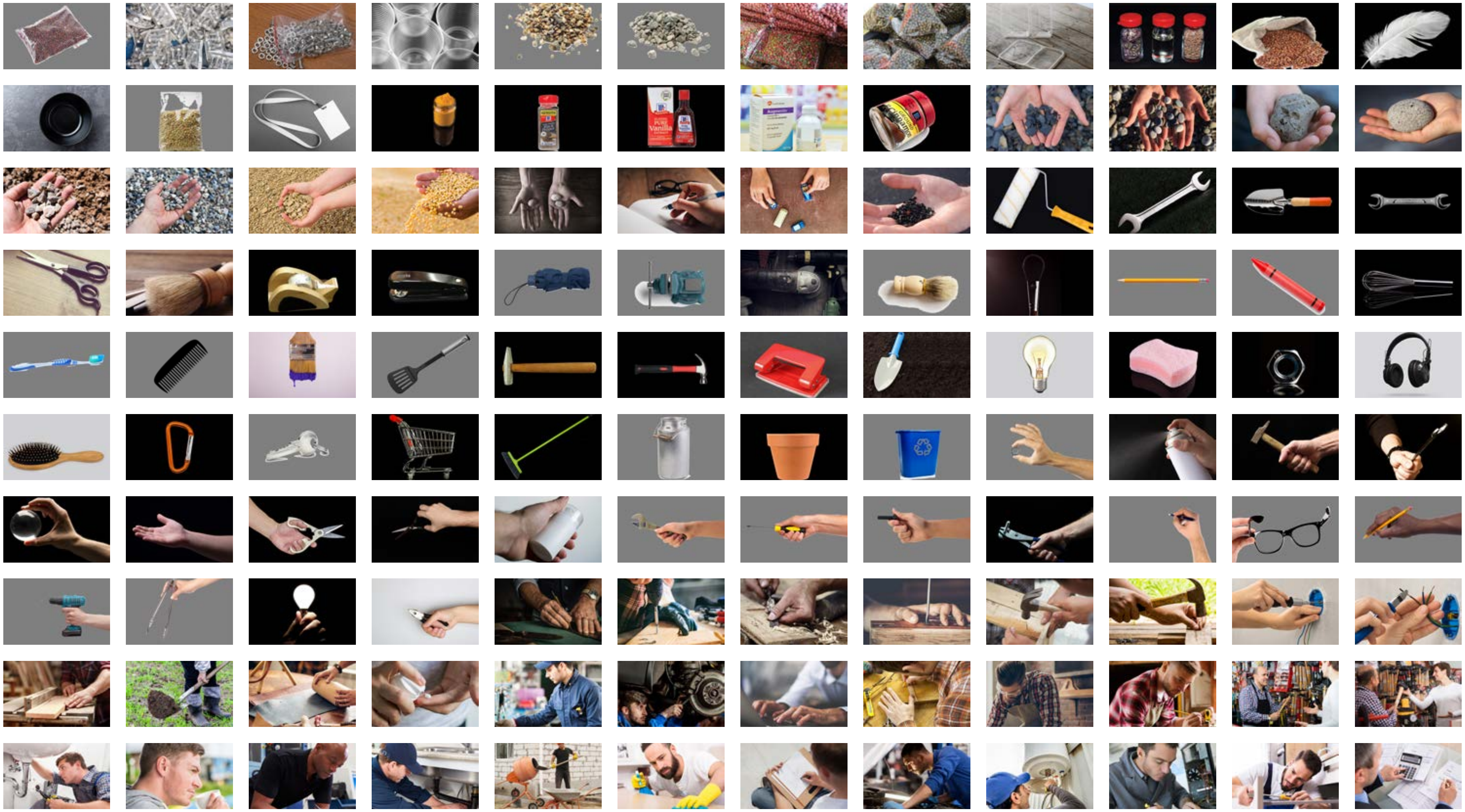

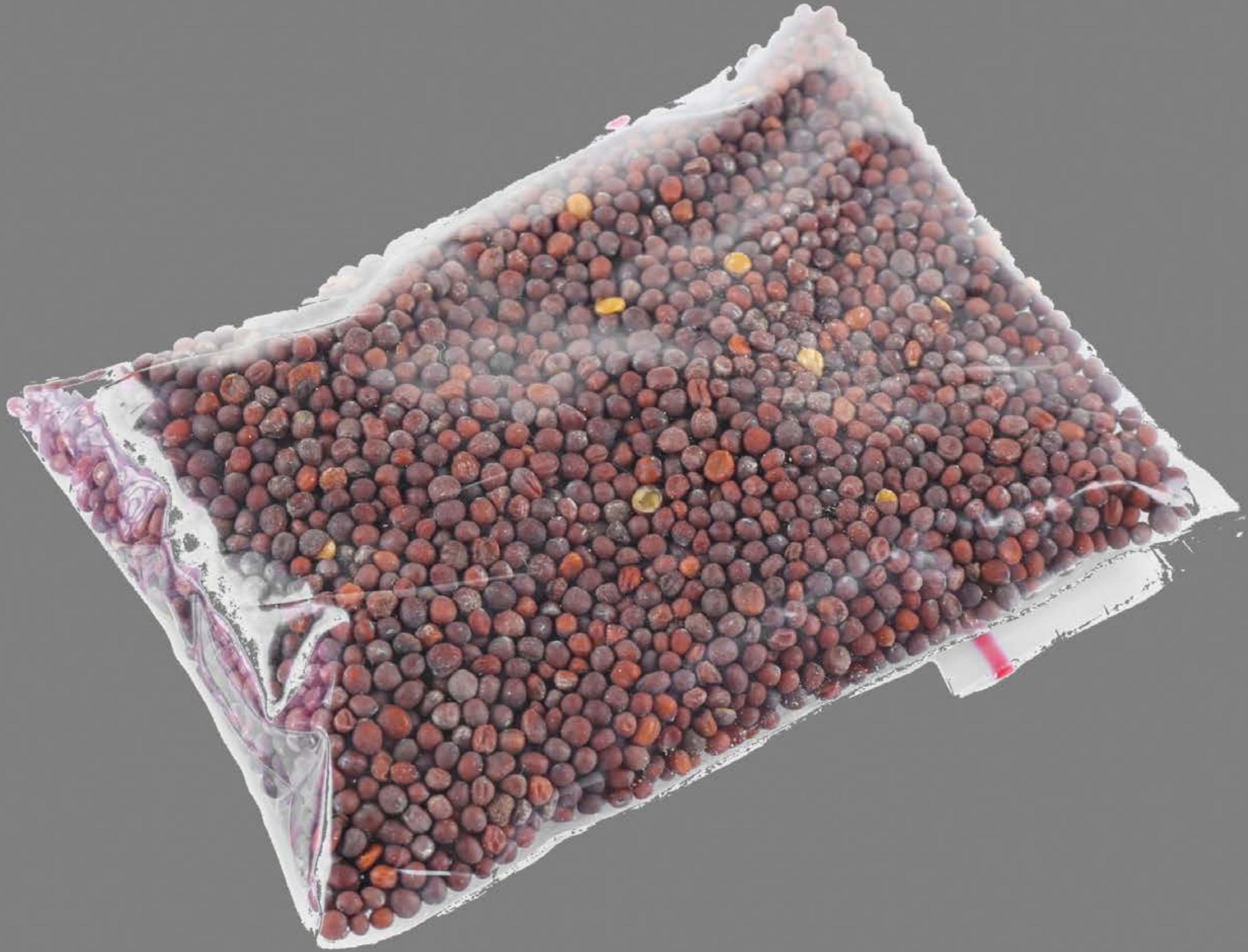

| Category | Craving (0-100) | Valence (1-9) | Arousal (1-9) | Typicality (0-100) | Relatedness | | | | | HSV | | |
|---|---|---|---|---|---|---|---|---|---|---|---|---|
| | | | | | Meth | Opioid | Both | Neither | MethToOpioid | Hue | Saturation | Value |
| Neutral objects | 14.25 (22) | 5.07 (1.412) | 2.36 (2) | 11.14 (20.88) | 0 | 0.07143 | 0.03571 | 0.89286 | 0.07143 | 0.176 (0.364) | 0.053 (0.11) | 0.516 (0.1) |

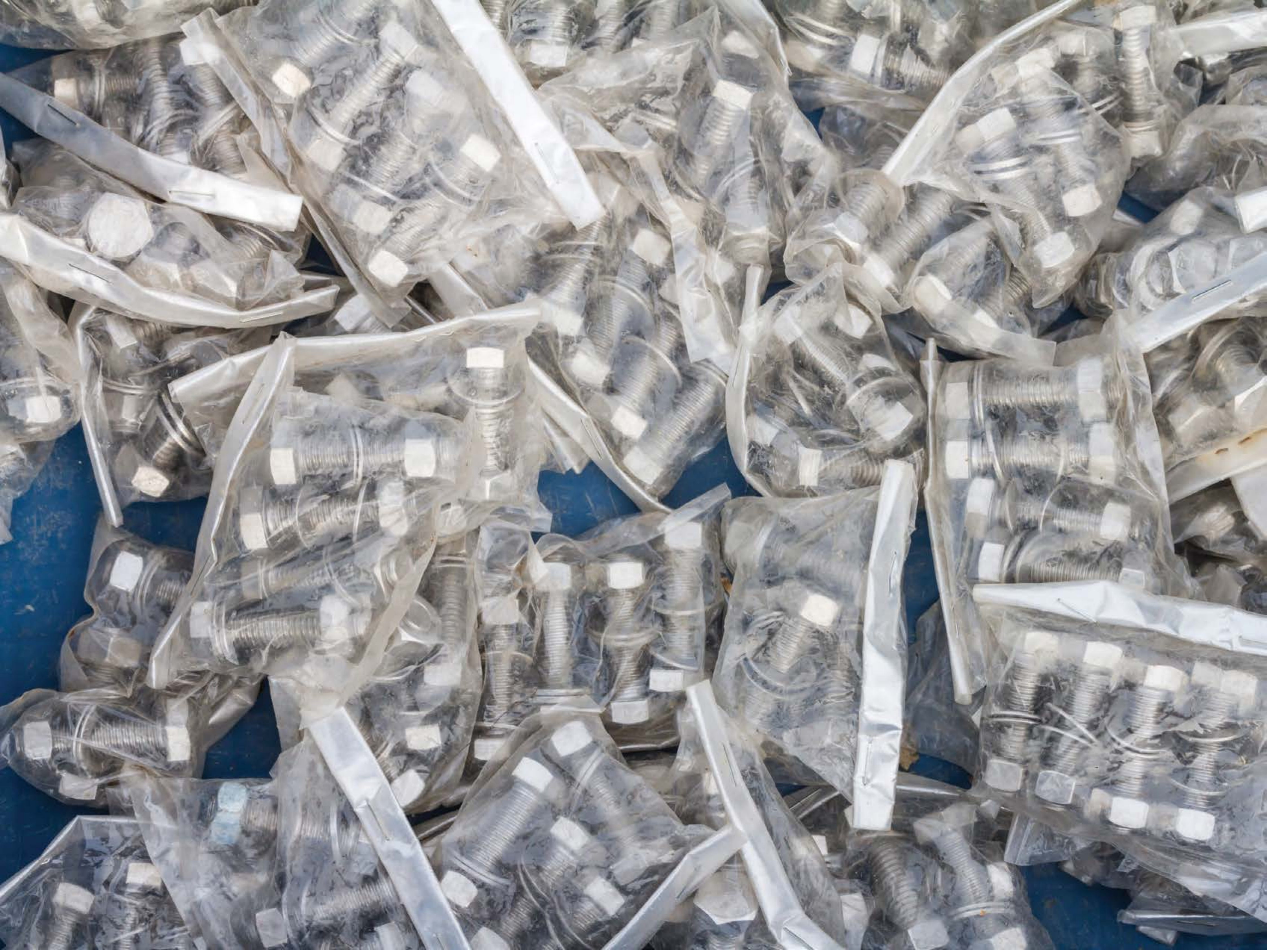

| Category | Craving (0-100) | Valence (1-9) | Arousal (1-9) | Typicality (0-100) | Relatedness | | | | | HSV | | |
|---|---|---|---|---|---|---|---|---|---|---|---|---|
| | | | | | Meth | Opioid | Both | Neither | MethToOpioid | Hue | Saturation | Value |
| Neutral objects | 13.93 (23.27) | 4.96 (0.898) | 2.67 (2.13) | 16.11 (25.08) | 0 | 0.03704 | 0 | 0.96296 | 0.03704 | 0.393(0.298) | 0.09(0.15) | 0.611(0.175) |

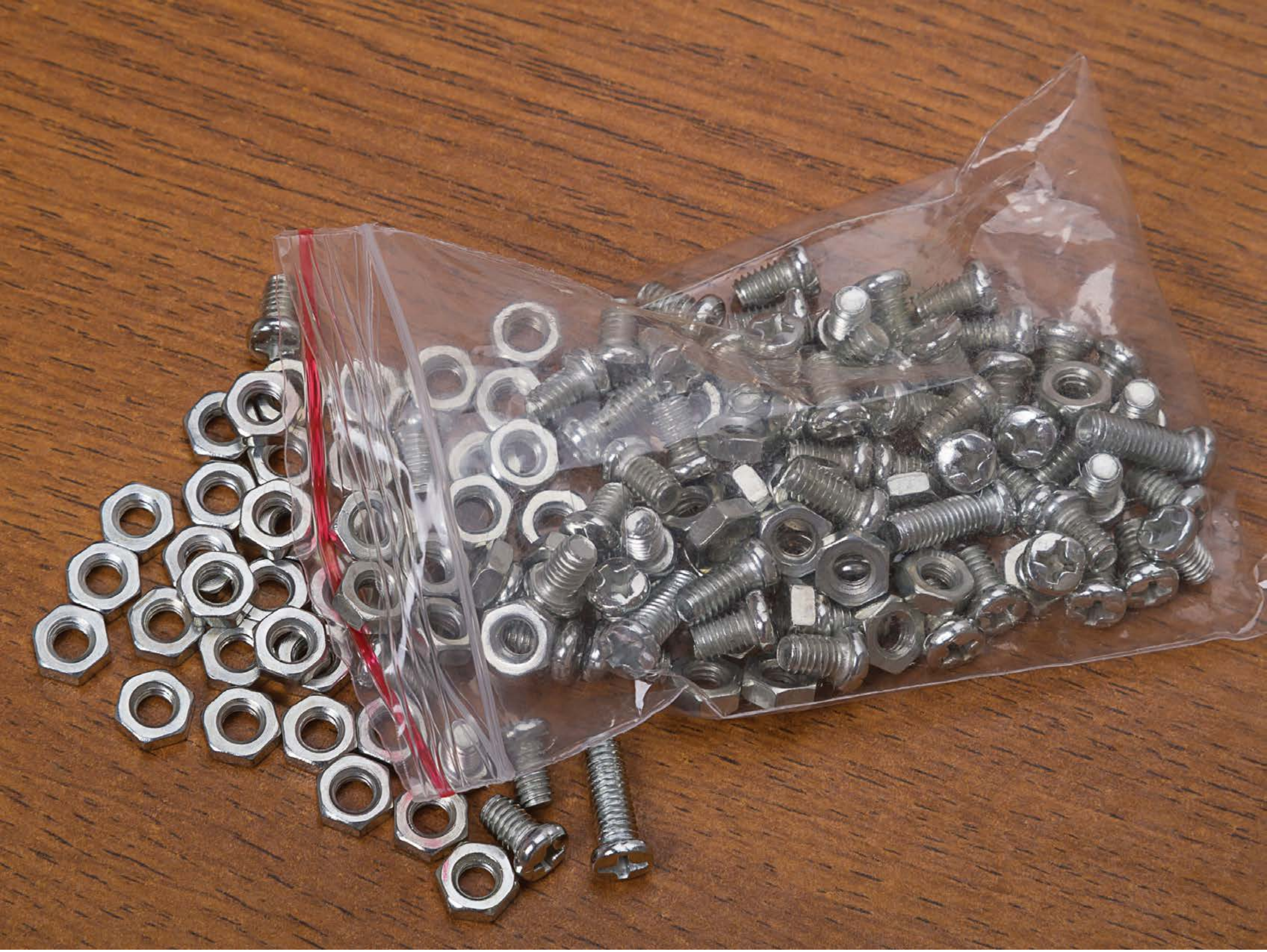

| Category | Craving (0-100) | Valence (1-9) | Arousal (1-9) | Typicality (0-100) | Relatedness | | | | | HSV | | |
|---|---|---|---|---|---|---|---|---|---|---|---|---|
| | | | | | Meth | Opioid | Both | Neither | MethToOpioid | Hue | Saturation | Value |
| Neutral objects | 13.5 (26.65) | 4.96 (1.478) | 2.68 (2.11) | 14.57 (26.98) | 0 | 0.03571 | 0.03571 | 0.92857 | 0.03571 | 0.182(0.275) | 0.394(0.247) | 0.529(0.135) |

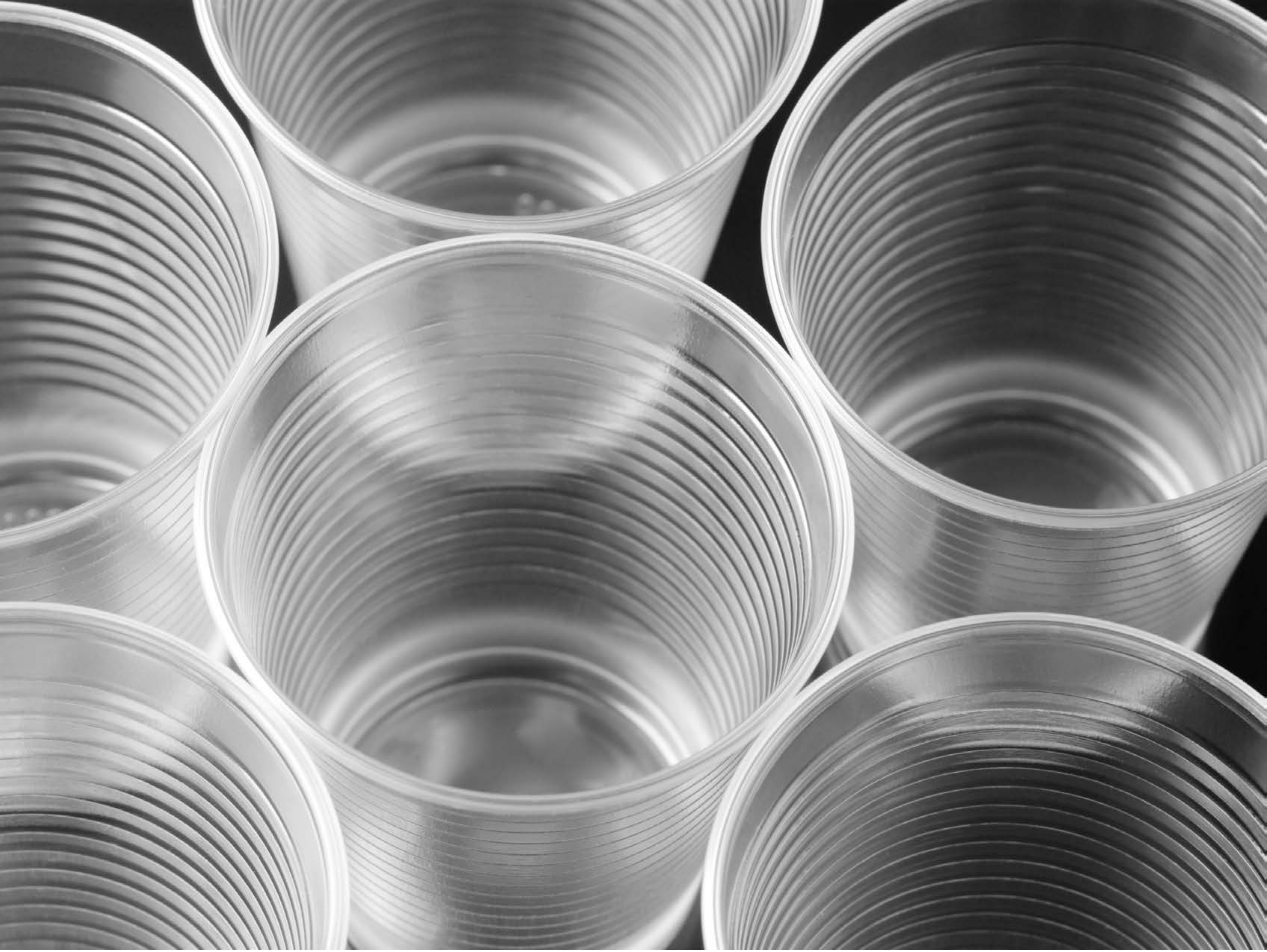

| Category | Craving (0-100) | Valence (1-9) | Arousal (1-9) | Typicality (0-100) | Relatedness | | | | | HSV | | |
|---|---|---|---|---|---|---|---|---|---|---|---|---|
| | | | | | Meth | Opioid | Both | Neither | MethToOpioid | Hue | Saturation | Value |
| Neutral objects | 7.78 (15.8) | 4.59 (1.185) | 2.37 (1.98) | 8.48 (16.11) | 0 | 0 | 0 | 1 | 0 | 0(0.002) | 0(0) | 0.544(0.219) |

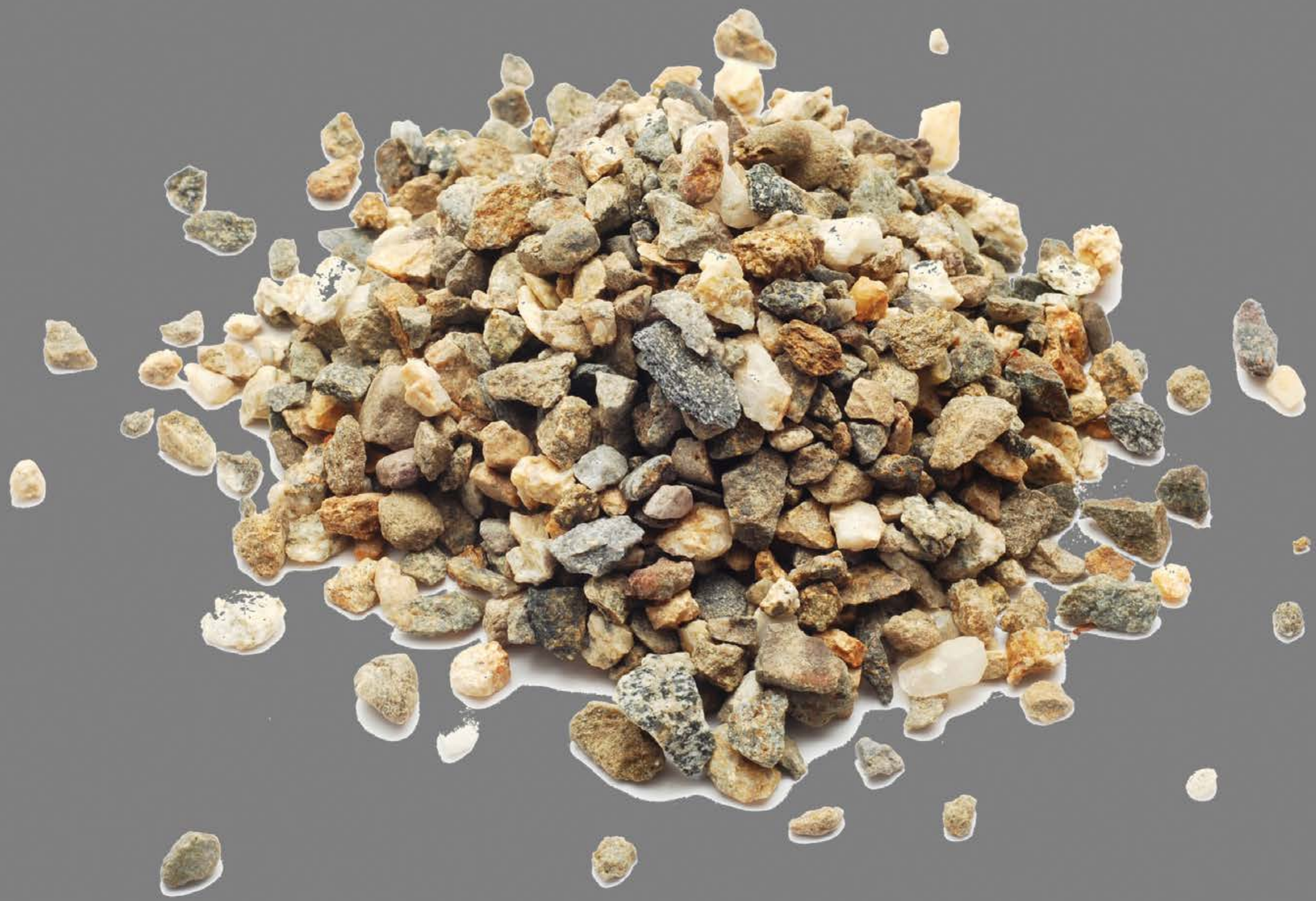

| Category | Craving (0-100) | Valence (1-9) | Arousal (1-9) | Typicality (0-100) | Relatedness | | | | | HSV | | |
|---|---|---|---|---|---|---|---|---|---|---|---|---|
| | | | | | Meth | Opioid | Both | Neither | MethToOpioid | Hue | Saturation | Value |
| Neutral objects | 6.71 (15.51) | 4.61 (1.595) | 1.93 (1.54) | 7.54 (15.66) | 0 | 0 | 0 | 1 | 0 | 0.036(0.097) | 0.083(0.161) | 0.539(0.142) |

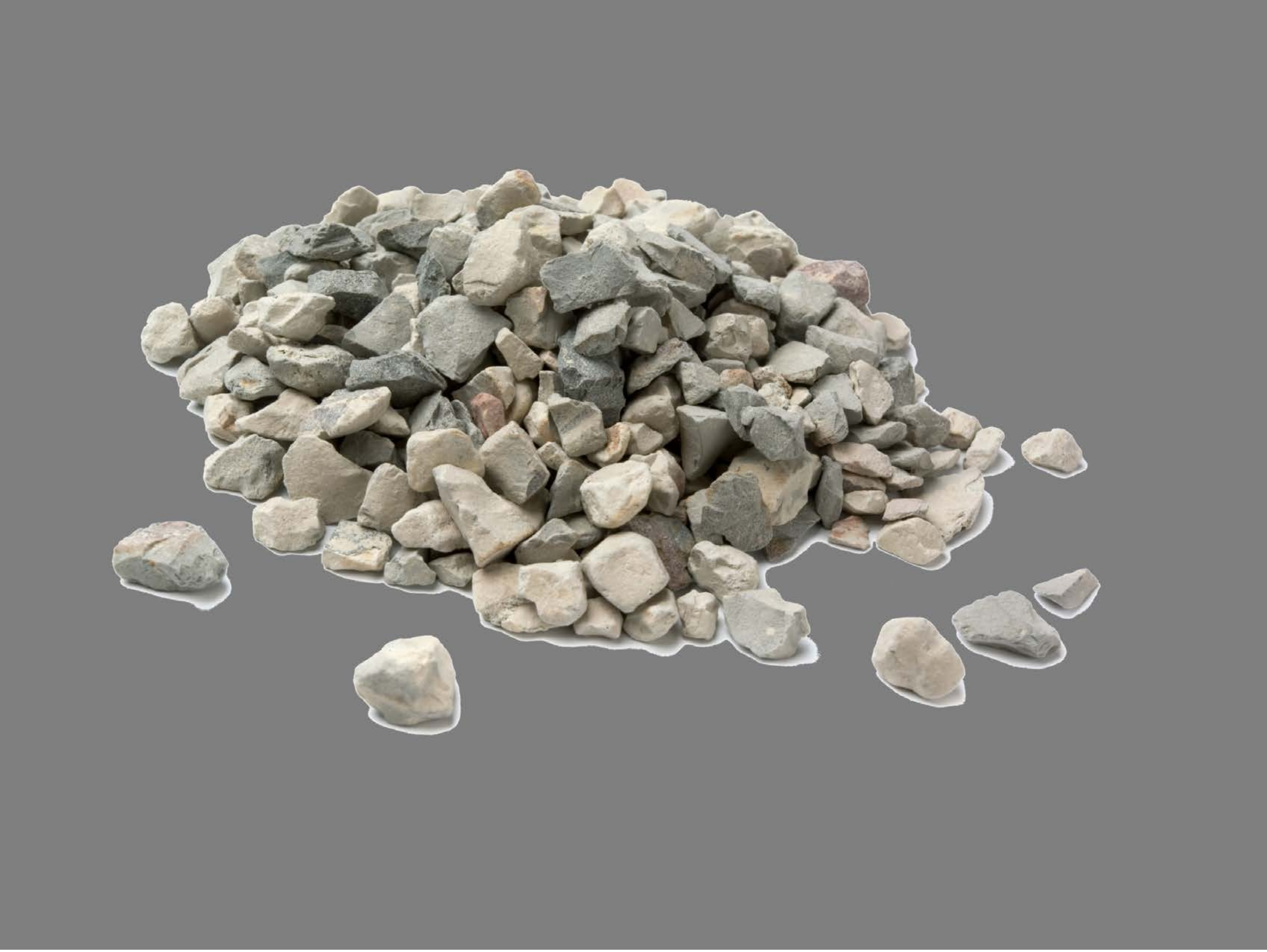

| Category | Craving (0-100) | Valence (1-9) | Arousal (1-9) | Typicality (0-100) | Relatedness | | | | | HSV | | |
|---|---|---|---|---|---|---|---|---|---|---|---|---|
| | | | | | Meth | Opioid | Both | Neither | MethToOpioid | Hue | Saturation | Value |
| Neutral objects | 12.96 (27.4) | 4.96 (1.753) | 2.11 (1.91) | 12.11 (22.68) | 0 | 0 | 0.03571 | 0.96429 | 0 | 0.038(0.078) | 0.041(0.094) | 0.525(0.116) |

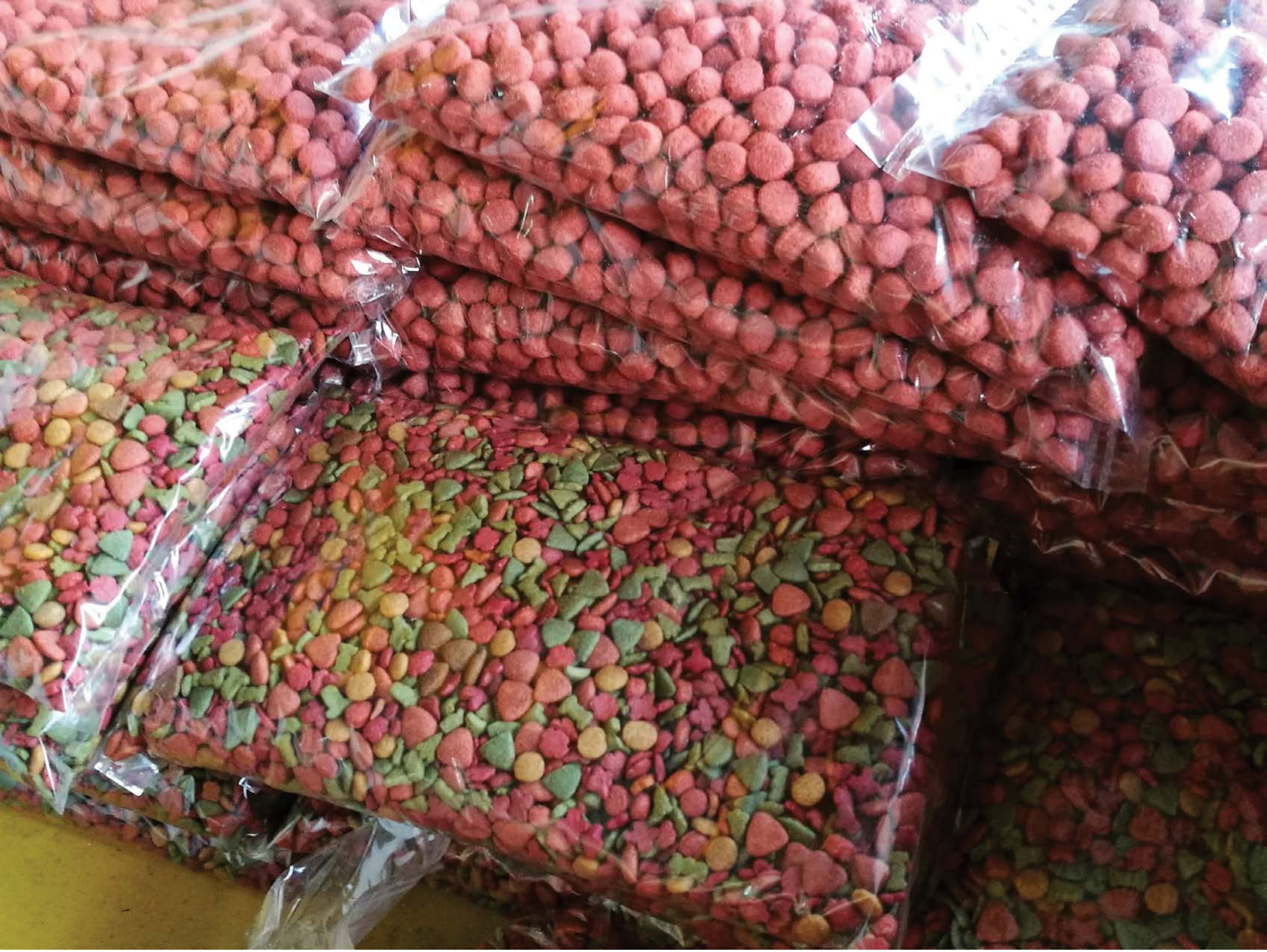

| Category | Craving (0-100) | Valence (1-9) | Arousal (1-9) | Typicality (0-100) | Relatedness | | | | | HSV | | |
|---|---|---|---|---|---|---|---|---|---|---|---|---|
| | | | | | Meth | Opioid | Both | Neither | MethToOpioid | Hue | Saturation | Value |
| Neutral objects | 13.89 (27.42) | 5.11 (0.577) | 2.89 (2.12) | 15.67 (28.19) | 0 | 0.03704 | 0 | 0.96296 | 0.03704 | 0.3 (0.399) | 0.513 (0.227) | 0.517 (0.26) |

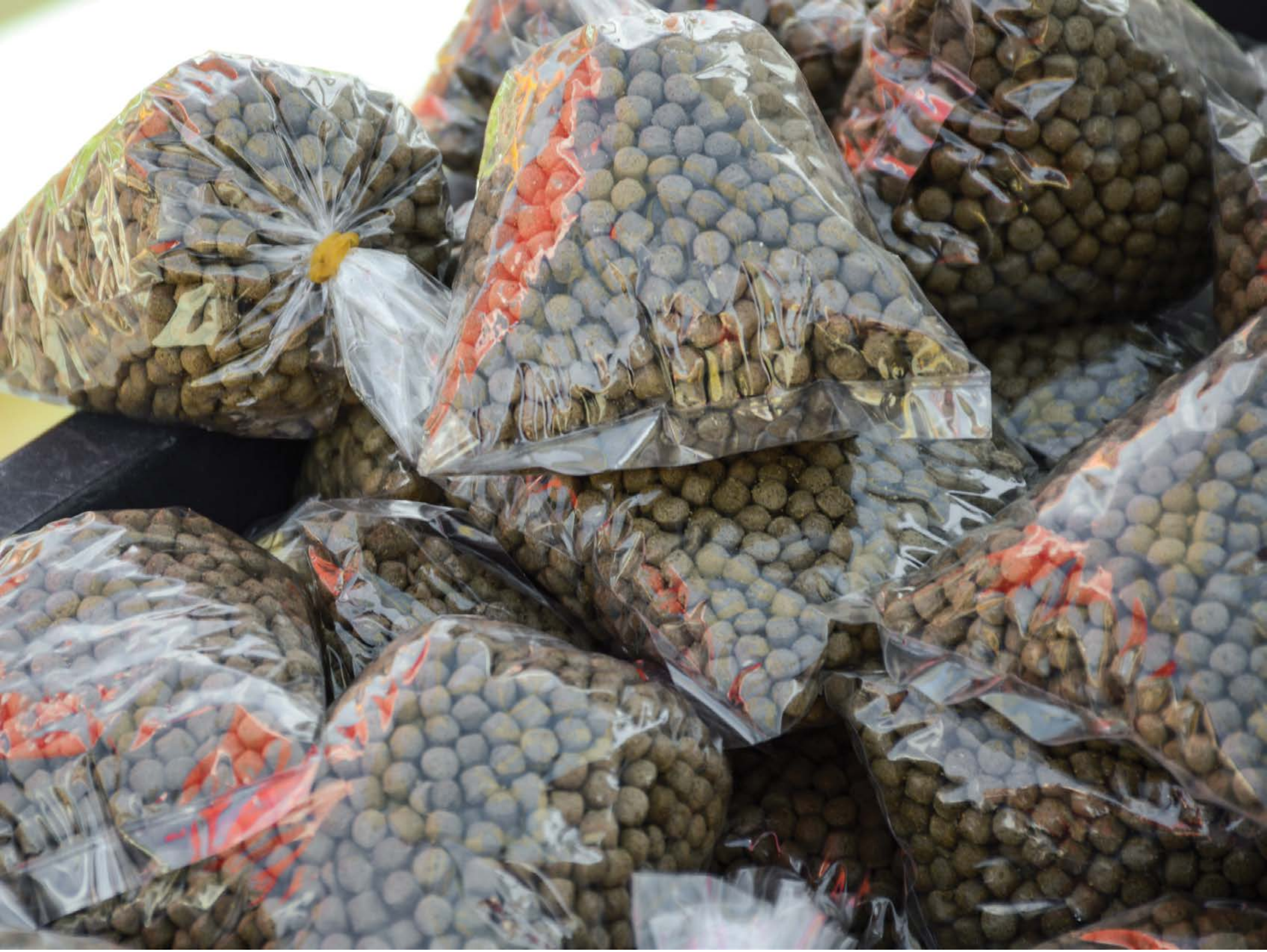

| Category | Craving (0-100) | Valence (1-9) | Arousal (1-9) | Typicality (0-100) | Relatedness | | | | | HSV | | |
|---|---|---|---|---|---|---|---|---|---|---|---|---|
| | | | | | Meth | Opioid | Both | Neither | MethToOpioid | Hue | Saturation | Value |
| Neutral objects | 12.74 (22.41) | 4.81 (1.241) | 3.37 (2.32) | 14.15 (23.59) | 0 | 0.03704 | 0 | 0.96296 | 0.03704 | 0.289 (0.261) | 0.192 (0.158) | 0.491(0.236) |

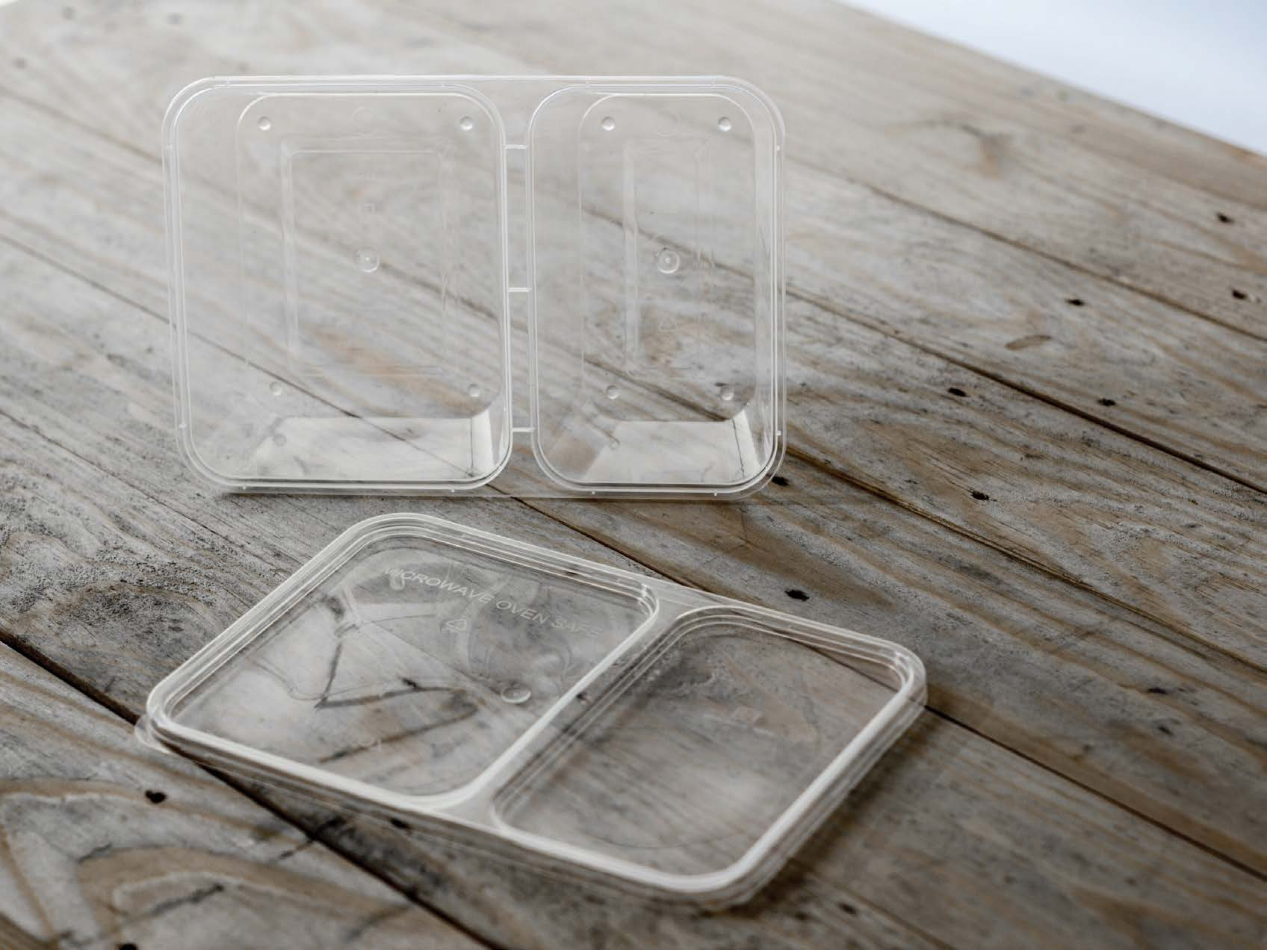


| Category | Craving (0-100) | Valence (1-9) | Arousal (1-9) | Typicality (0-100) | Relatedness | | | | | HSV | | |
|---|---|---|---|---|---|---|---|---|---|---|---|---|
| | | | | | Meth | Opioid | Both | Neither | MethToOpioid | Hue | Saturation | Value |
| Neutral objects | 16.63 (26.4) | 4.74 (0.944) | 2.67 (1.98) | 11.19(17.79) | 0 | 0 | 0.03704 | 0.96296 | 0 | 0.131 (0.149) | 0.092 (0.123) | 0.591 (0.168) |

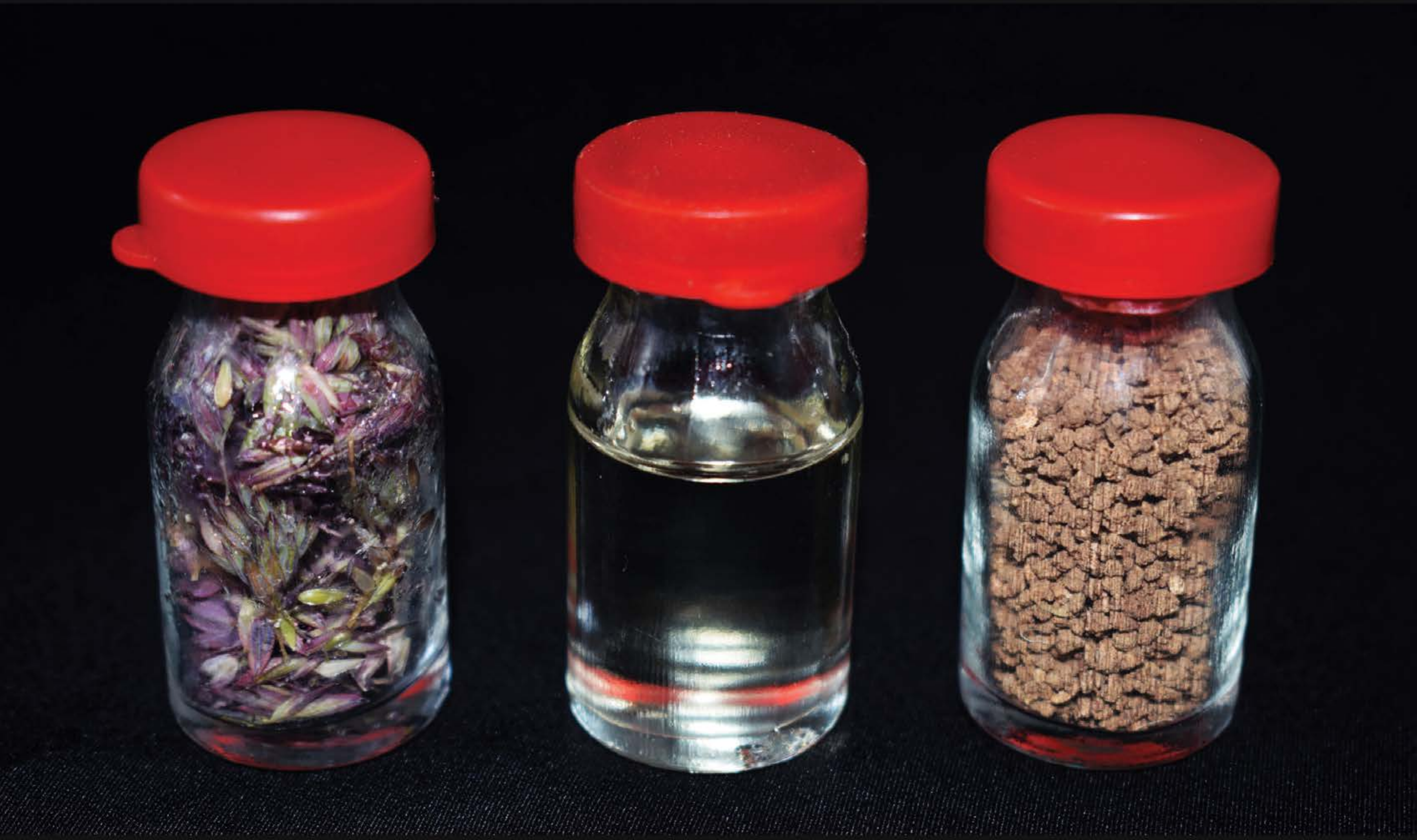

| Category | Craving (0-100) | Valence (1-9) | Arousal (1-9) | Typicality (0-100) | Relatedness | | | | | HSV | | |
|---|---|---|---|---|---|---|---|---|---|---|---|---|
| | | | | | Meth | Opioid | Both | Neither | MethToOpioid | Hue | Saturation | Value |
| Neutral objects | 13 (19.87) | 4.96 (1.372) | 2.63 (1.96) | 12 (19.4) | 0 | 0 | 0.03704 | 0.96296 | 0 | 0.414 (0.338) | 0.447 (0.342) | 0.212(0.271) |

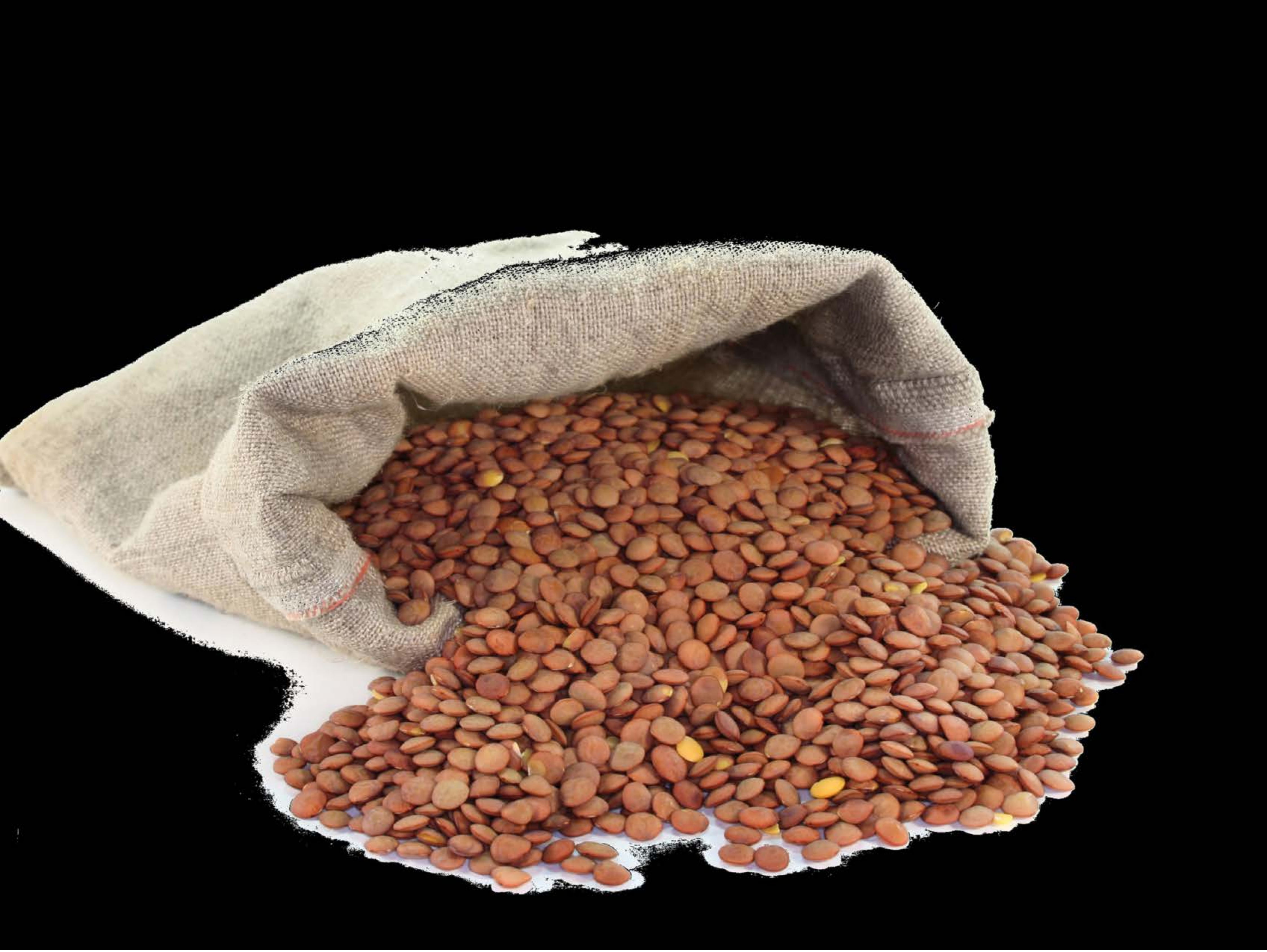

| Category | Craving (0-100) | Valence (1-9) | Arousal (1-9) | Typicality (0-100) | Relatedness | | | | | HSV | | |
|---|---|---|---|---|---|---|---|---|---|---|---|---|
| | | | | | Meth | Opioid | Both | Neither | MethToOpioid | Hue | Saturation | Value |
| Neutral objects | 11.29 (22.26) | 4.93 (1.654) | 2.46 (1.95) | 12.43 (22.28) | 0 | 0.03571 | 0 | 0.96429 | 0.03571 | 0.043 (0.107) | 0.176 (0.251) | 0.279 (0.343) |

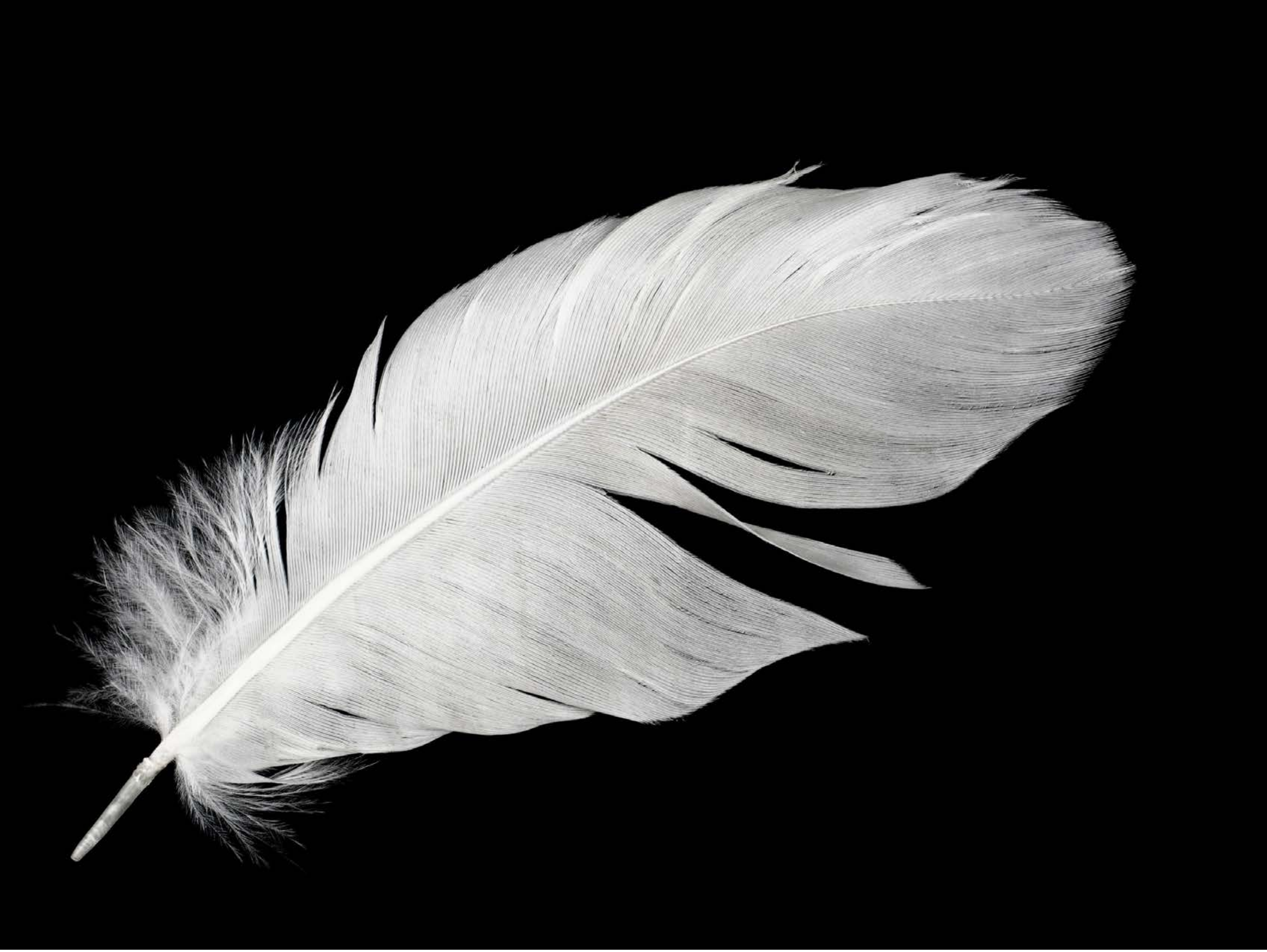

| Category | Craving (0-100) | Valence (1-9) | Arousal (1-9) | Typicality (0-100) | Relatedness | | | | | HSV | | |
|---|---|---|---|---|---|---|---|---|---|---|---|---|
| | | | | | Meth | Opioid | Both | Neither | MethToOpioid | Hue | Saturation | Value |
| Neutral objects | 15.46 (29.28) | 5.25 (1.818) | 2.39 (1.77) | 9.43 (17.38) | 0 | 0 | 0 | 1 | 0 | 0.059 (0.16) | 0.008 (0.061) | 0.234 (0.358) |

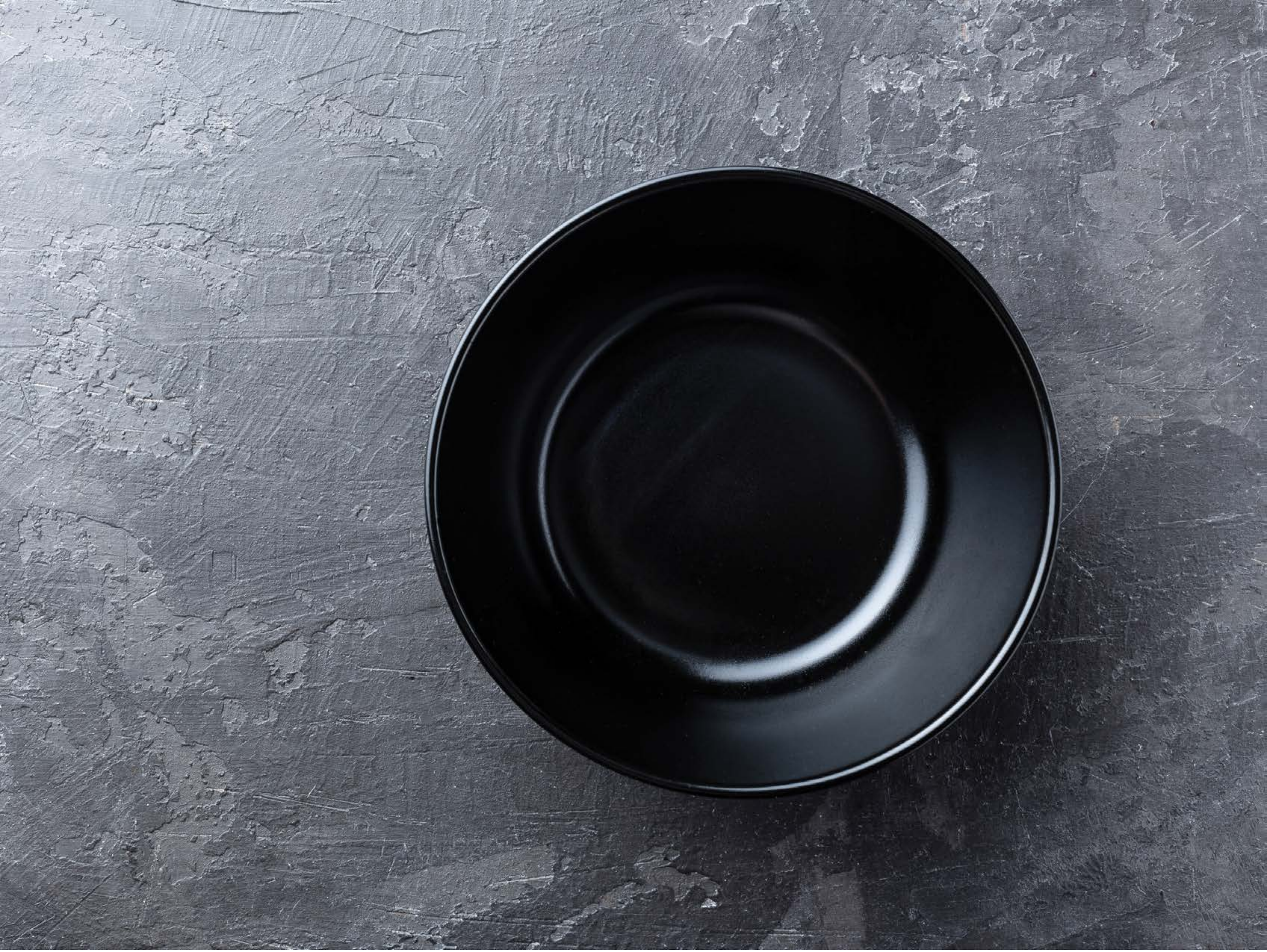

| Category | Craving (0-100) | Valence (1-9) | Arousal (1-9) | Typicality (0-100) | Relatedness | | | | | HSV | | |
|---|---|---|---|---|---|---|---|---|---|---|---|---|
| | | | | | Meth | Opioid | Both | Neither | MethToOpioid | Hue | Saturation | Value |
| Neutral objects | 10.25 (22.02) | 4.79 (1.548) | 2.29 (1.98) | 14 (25.01) | 0 | 0.07143 | 0 | 0.92857 | 0.07143 | 0.6 (0.134) | 0.154 (0.117) | 0.402 (0.256) |

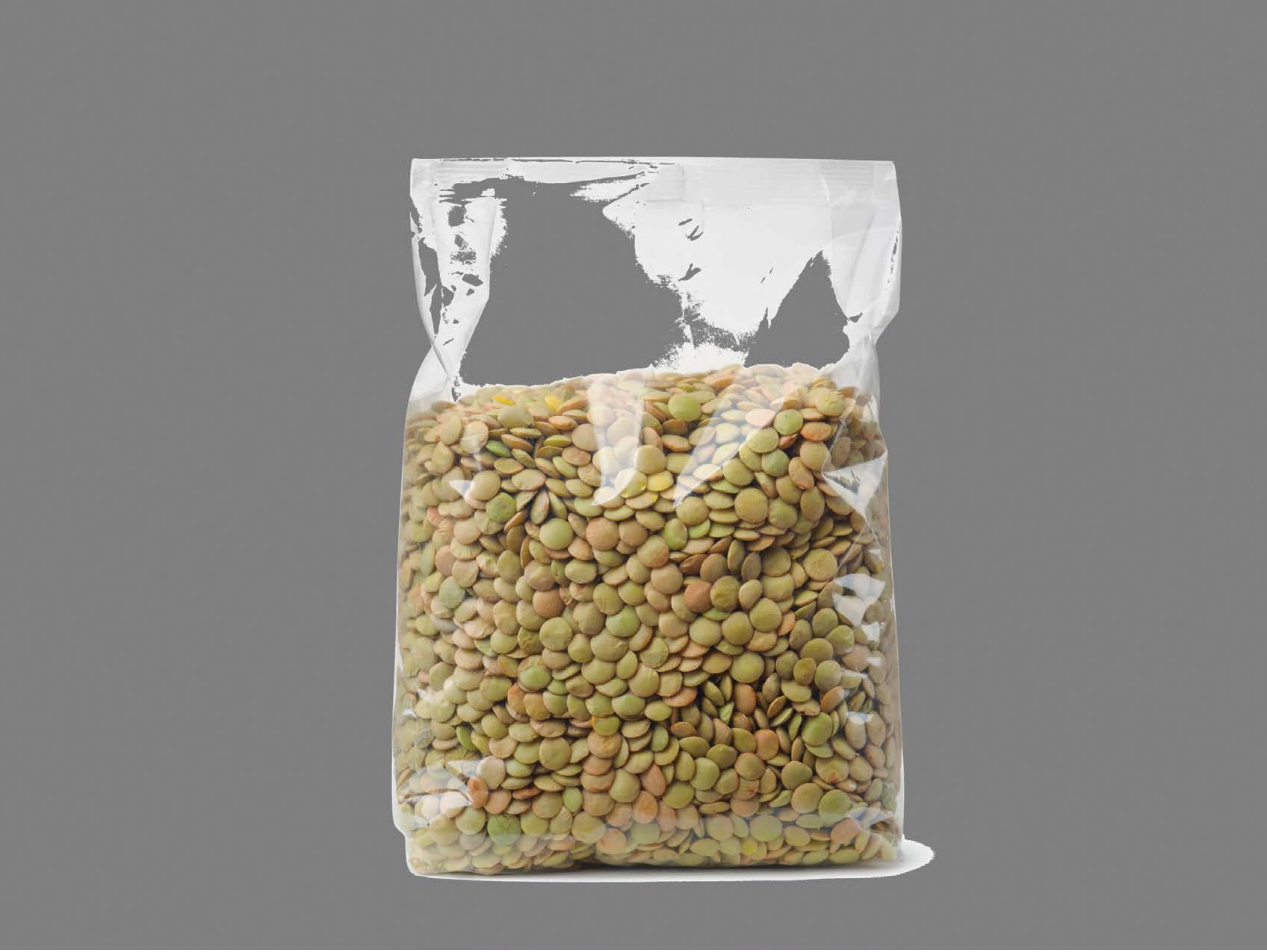

| Category | Craving (0-100) | Valence (1-9) | Arousal (1-9) | Typicality (0-100) | Relatedness | | | | | HSV | | |
|---|---|---|---|---|---|---|---|---|---|---|---|---|
| | | | | | Meth | Opioid | Both | Neither | MethToOpioid | Hue | Saturation | Value |
| Neutral objects | 13.86 (27.67) | 5.04 (1.503) | 2.11 (1.97) | 10.54 (21.89) | 0 | 0.03571 | 0 | 0.96429 | 0.03571 | 0.028 (0.071) | 0.08 (0.175) | 0.543(0.121) |

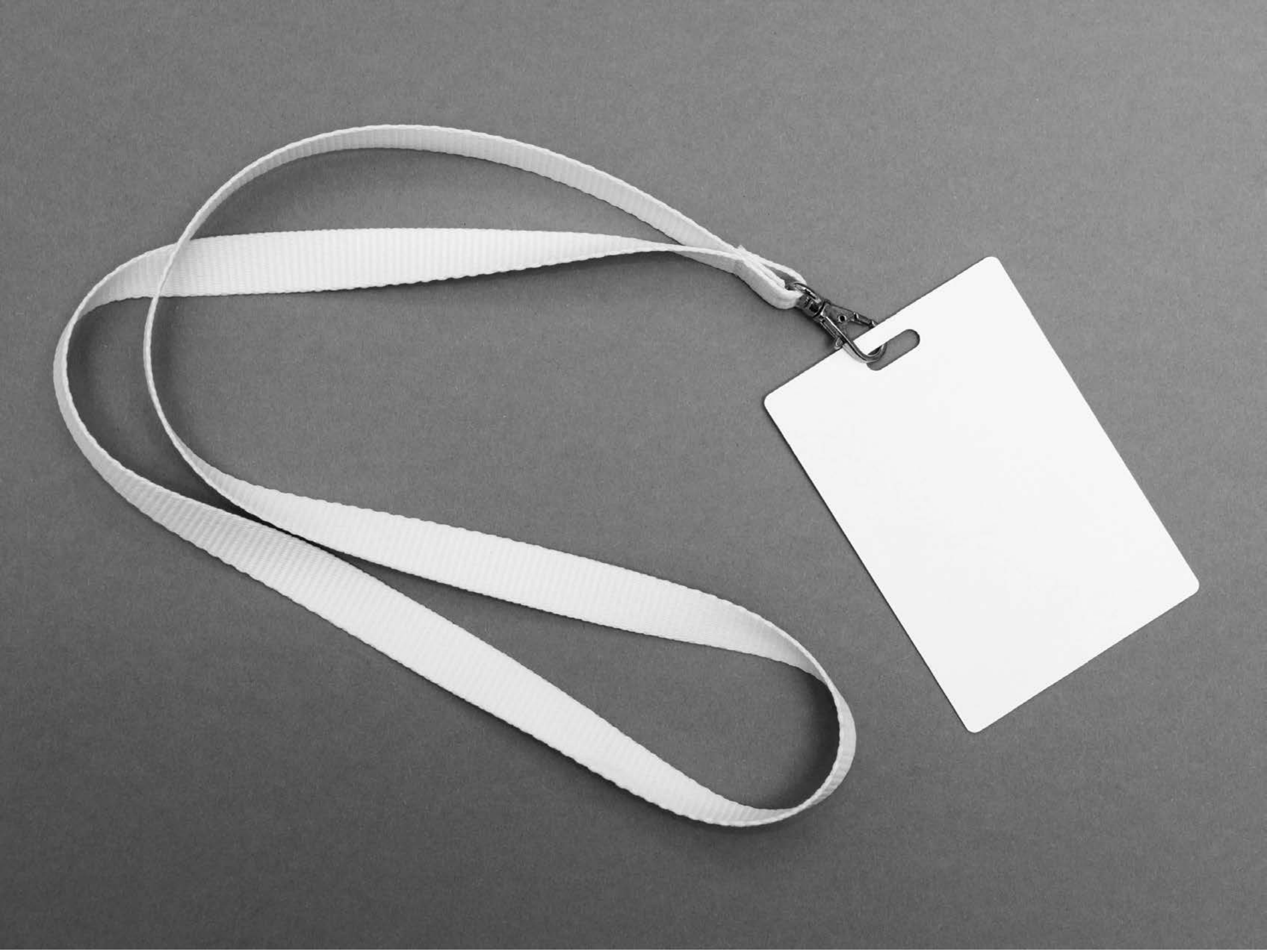

| Category | Craving (0-100) | Valence (1-9) | Arousal (1-9) | Typicality (0-100) | Relatedness | | | | | HSV | | |
|---|---|---|---|---|---|---|---|---|---|---|---|---|
| | | | | | Meth | Opioid | Both | Neither | MethToOpioid | Hue | Saturation | Value |
| Neutral objects | 12.78 (24.83) | 4.74 (1.095) | 2.48 (1.87) | 9.67 (17.58) | 0 | 0 | 0 | 1 | 0 | 0 (0) | 0 (0) | 0.527 (0.197) |

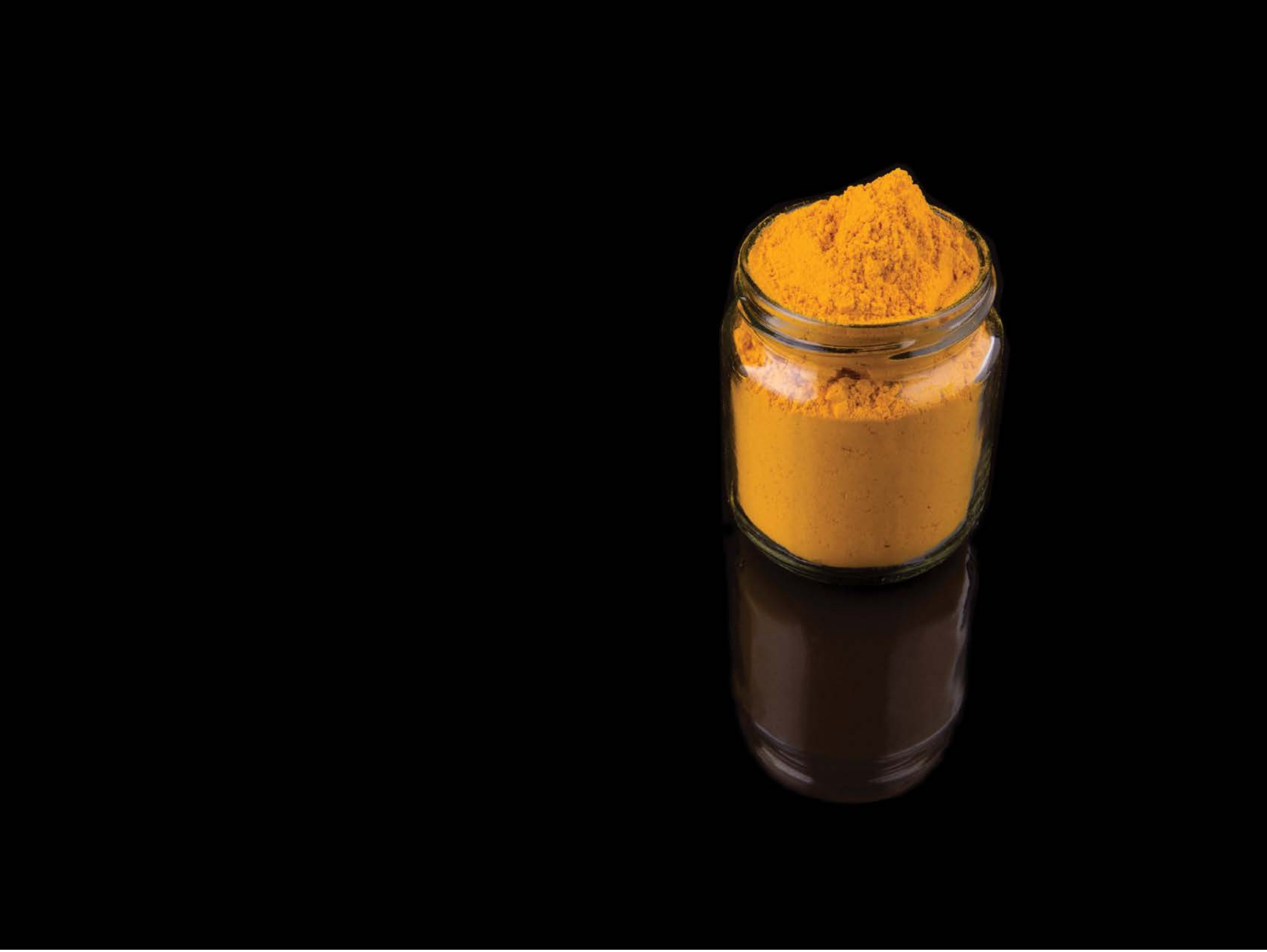

| Category | Craving (0-100) | Valence (1-9) | Arousal (1-9) | Typicality (0-100) | Relatedness | | | | | HSV | | |
|---|---|---|---|---|---|---|---|---|---|---|---|---|
| | | | | | Meth | Opioid | Both | Neither | MethToOpioid | Hue | Saturation | Value |
| Neutral objects | 12.37 (21.58) | 5.15 (0.949) | 2.93 (2.02) | 10.04 (19.8) | 0.03704 | 0 | 0 | 0.96296 | -0.03704 | 0.014 (0.067) | 0.099 (0.274) | 0.062 (0.201) |

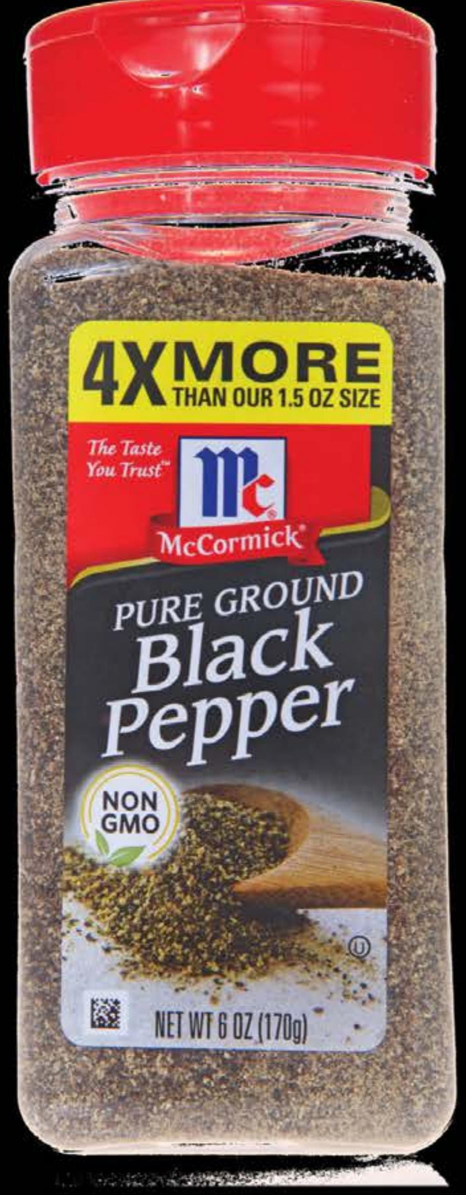


| Category | Craving (0-100) | Valence (1-9) | Arousal (1-9) | Typicality (0-100) | Relatedness | | | | | HSV | | |
|---|---|---|---|---|---|---|---|---|---|---|---|---|
| | | | | | Meth | Opioid | Both | Neither | MethToOpioid | Hue | Saturation | Value |
| Neutral objects | 11.19 (23.45) | 4.85 (1.064) | 2.48 (1.87) | 12.15 (23.84) | 0 | 0 | 0 | 1 | 0 | 0.062 (0.213) | 0.054 (0.182) | 0.084 (0.235) |

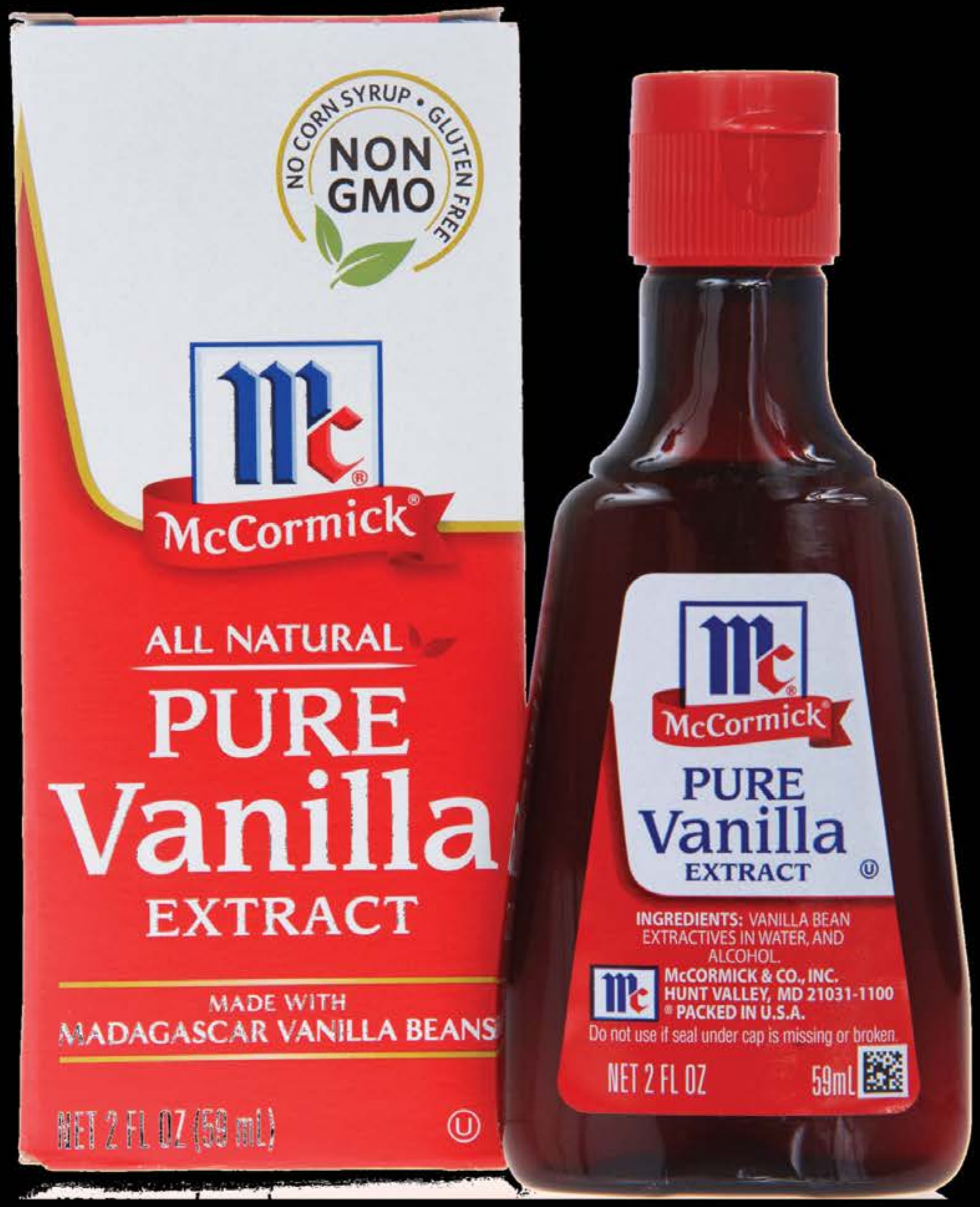


| | | | | | Relatedness | | | | | HSV | | |
|---|---|---|---|---|---|---|---|---|---|---|---|---|
| **Category** | **Craving (0-100)** | **Valence (1-9)** | **Arousal (1-9)** | **Typicality (0-100)** | Meth | Opioid | Both | Neither | MethToOpioid | Hue | Saturation | Value |
| Neutral objects | 10.71 (19.55) | 5 (1.785) | 2.36 (2.11) | 10.54 (18.99) | 0 | 0 | 0 | 1 | 0 | 0.181 (0.35) | 0.148 (0.295) | 0.219 (0.37) |

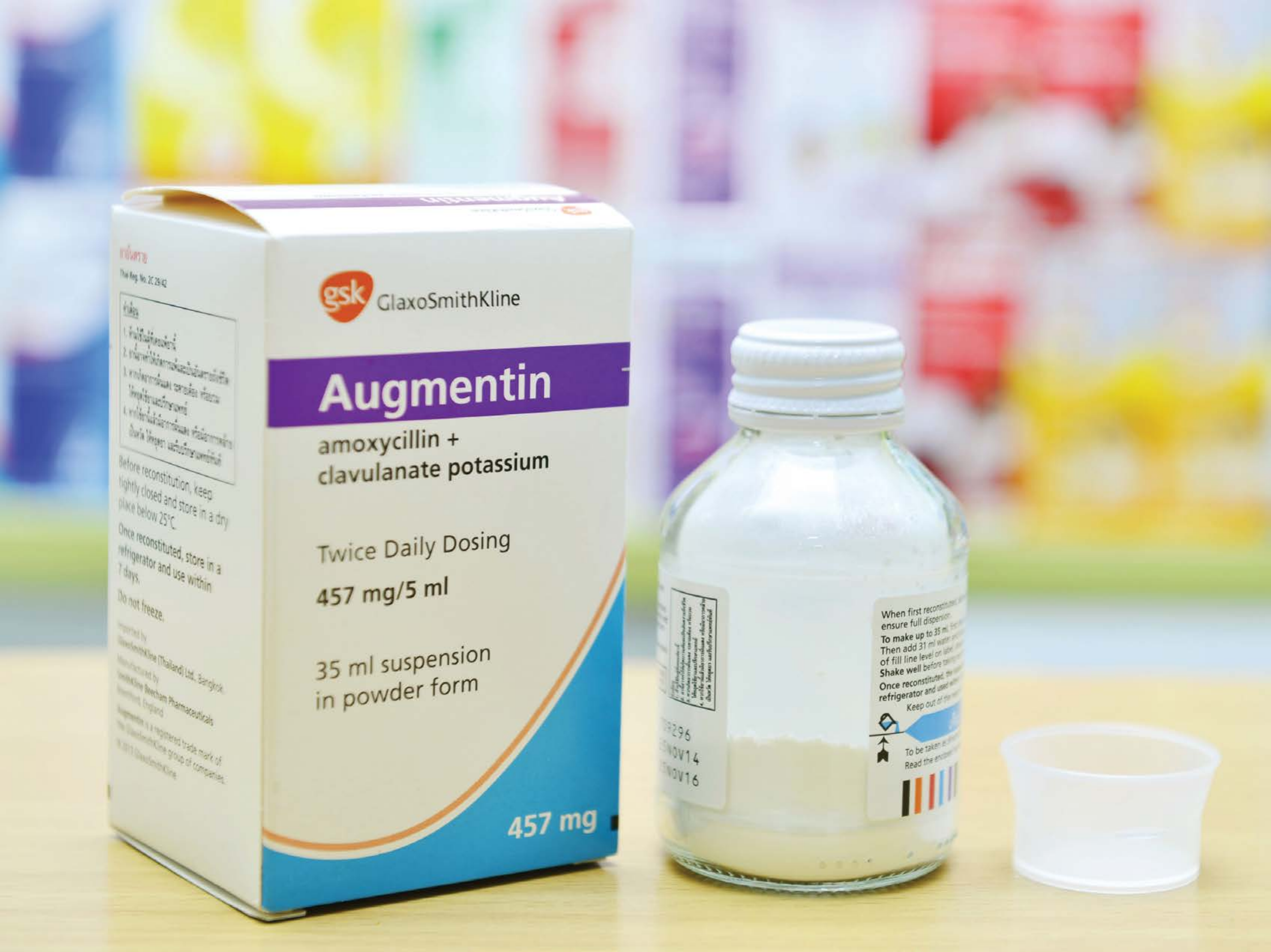


| Category | Craving (0-100) | Valence (1-9) | Arousal (1-9) | Typicality (0-100) | Relatedness | | | | | HSV | | |
|---|---|---|---|---|---|---|---|---|---|---|---|---|
| | | | | | Meth | Opioid | Both | Neither | MethToOpioid | Hue | Saturation | Value |
| Neutral objects | 13.04 (24.56) | 4.79 (1.101) | 2.46 (2.05) | 13.79 (24.52) | 0 | 0.07143 | 0.07143 | 0.85714 | 0.07143 | 0.368 (0.262) | 0.209 (0.207) | 0.858 (0.096) |

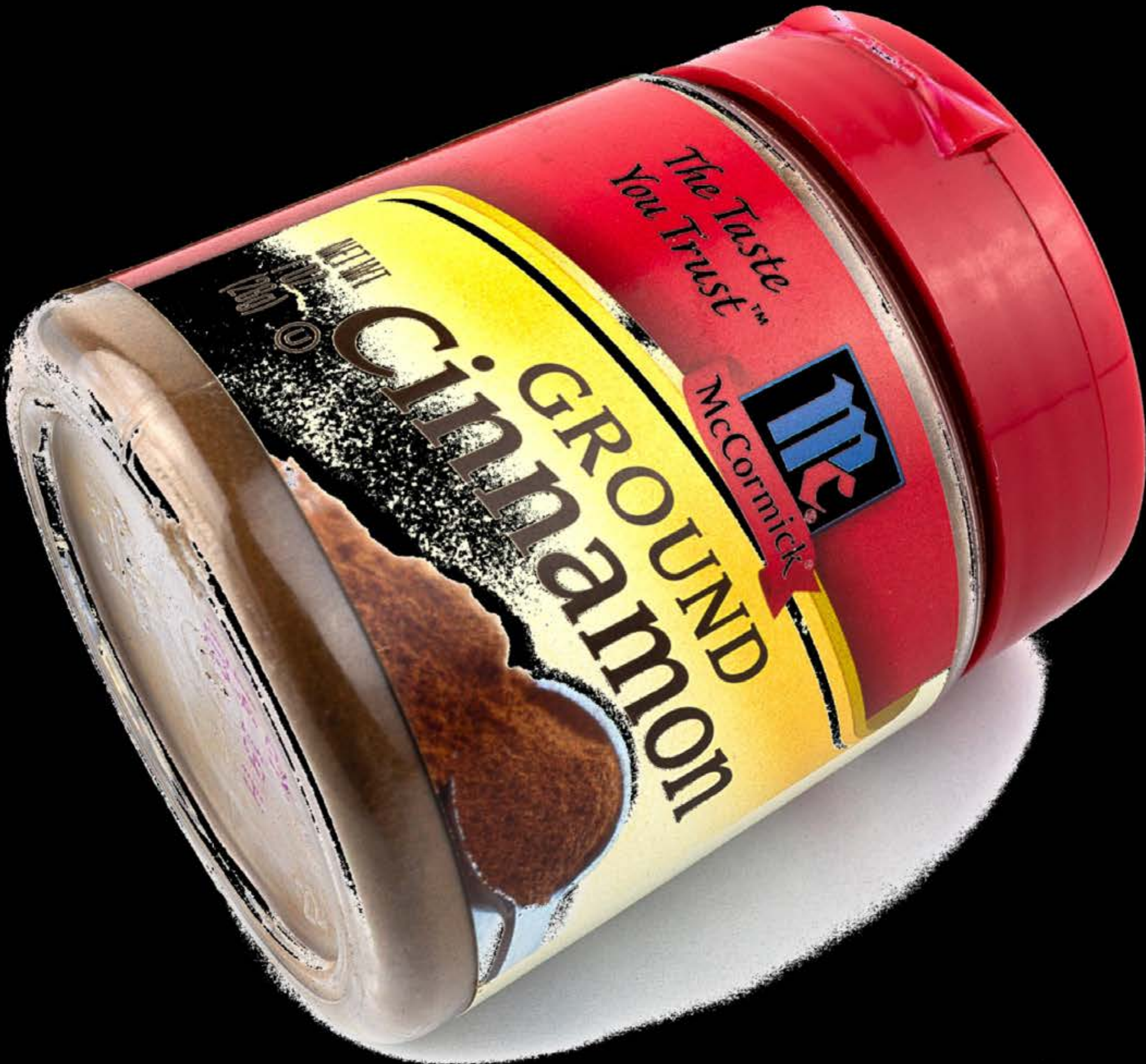


| Category | Craving (0-100) | Valence (1-9) | Arousal (1-9) | Typicality (0-100) | Relatedness | | | | | HSV | | |
|---|---|---|---|---|---|---|---|---|---|---|---|---|
| | | | | | Meth | Opioid | Both | Neither | MethToOpioid | Hue | Saturation | Value |
| Neutral objects | 8.26 (15.78) | 4.85 (1.134) | 2.48 (1.85) | 8.11 (15.6) | 0 | 0 | 0 | 1 | 0 | 0.132 (0.302) | 0.147 (0.276) | 0.219 (0.352) |

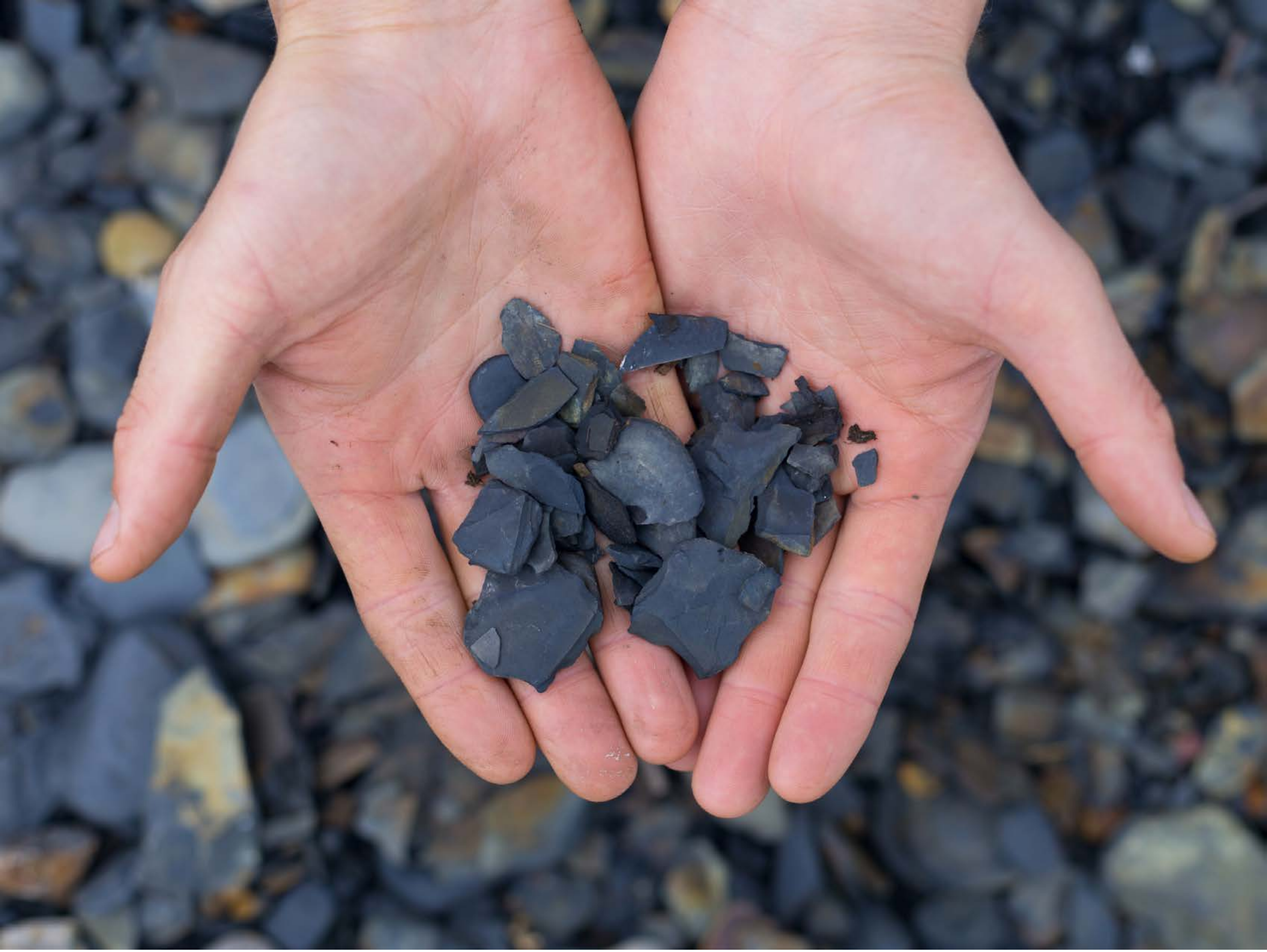

| Category | Craving (0-100) | Valence (1-9) | Arousal (1-9) | Typicality (0-100) | Relatedness | | | | | HSV | | |
|---|---|---|---|---|---|---|---|---|---|---|---|---|
| | | | | | Meth | Opioid | Both | Neither | MethToOpioid | Hue | Saturation | Value |
| Neutral objects hand | 11.43 (21.41) | 4.96 (1.427) | 2.39 (1.81) | 12.18 (21.84) | 0.03571 | 0 | 0 | 0.96429 | -0.03571 | 0.428 (0.329) | 0.273 (0.12) | 0.522 (0.254) |

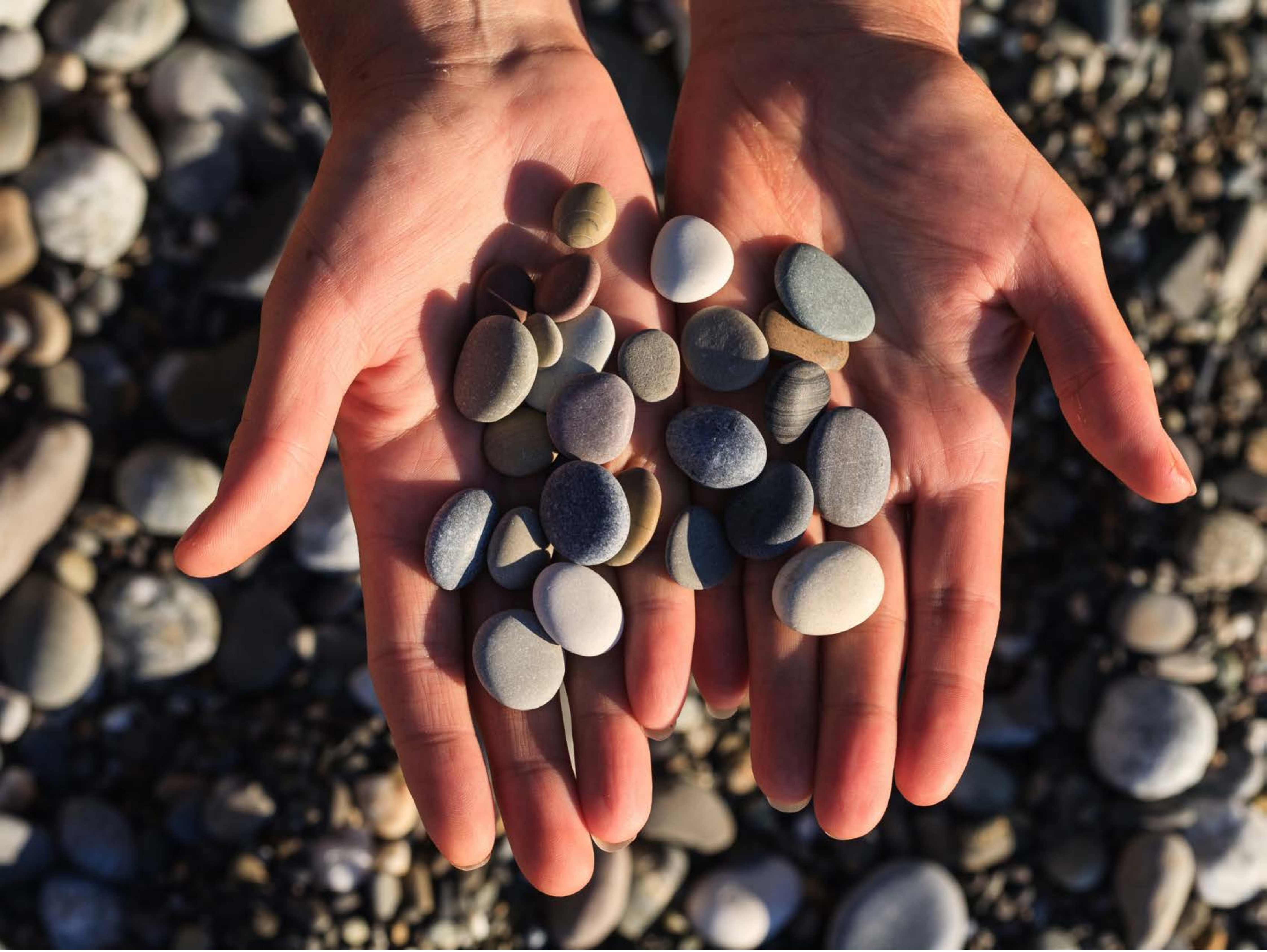

| Category | Craving (0-100) | Valence (1-9) | Arousal (1-9) | Typicality (0-100) | Relatedness | | | | | HSV | | |
|---|---|---|---|---|---|---|---|---|---|---|---|---|
| | | | | | Meth | Opioid | Both | Neither | MethToOpioid | Hue | Saturation | Value |
| Neutral objects hand | 18.19 (28.57) | 5.04 (1.055) | 2.7 (2.15) | 15 (24.05) | | 0.03704 | | 0.96296 | 0.03704 | 0.302 (0.352) | 0.316 (0.174) | 0.416 (0.3) |

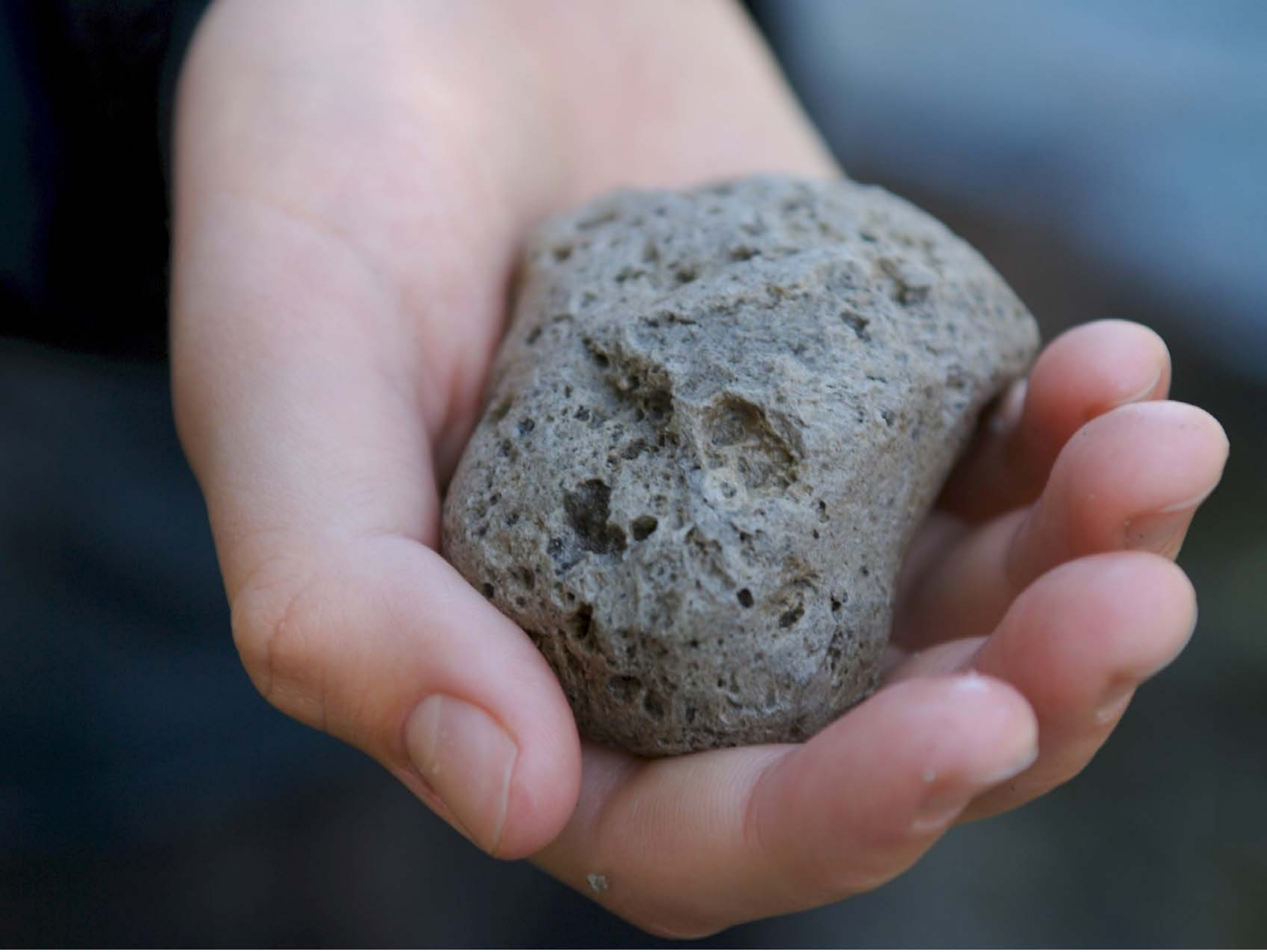

| Category | Craving (0-100) | Valence (1-9) | Arousal (1-9) | Typicality (0-100) | Relatedness | | | | | HSV | | |
|---|---|---|---|---|---|---|---|---|---|---|---|---|
| | | | | | Meth | Opioid | Both | Neither | MethToOpioid | Hue | Saturation | Value |
| Neutral objects hand | 9.18 (20.78) | 5.04 (1.774) | 2.54 (2.12) | 8.79 (19.86) | 0.03571 | 0 | 0 | 0.96429 | -0.03571 | 0.533 (0.316) | 0.257 (0.205) | 0.453 (0.247) |

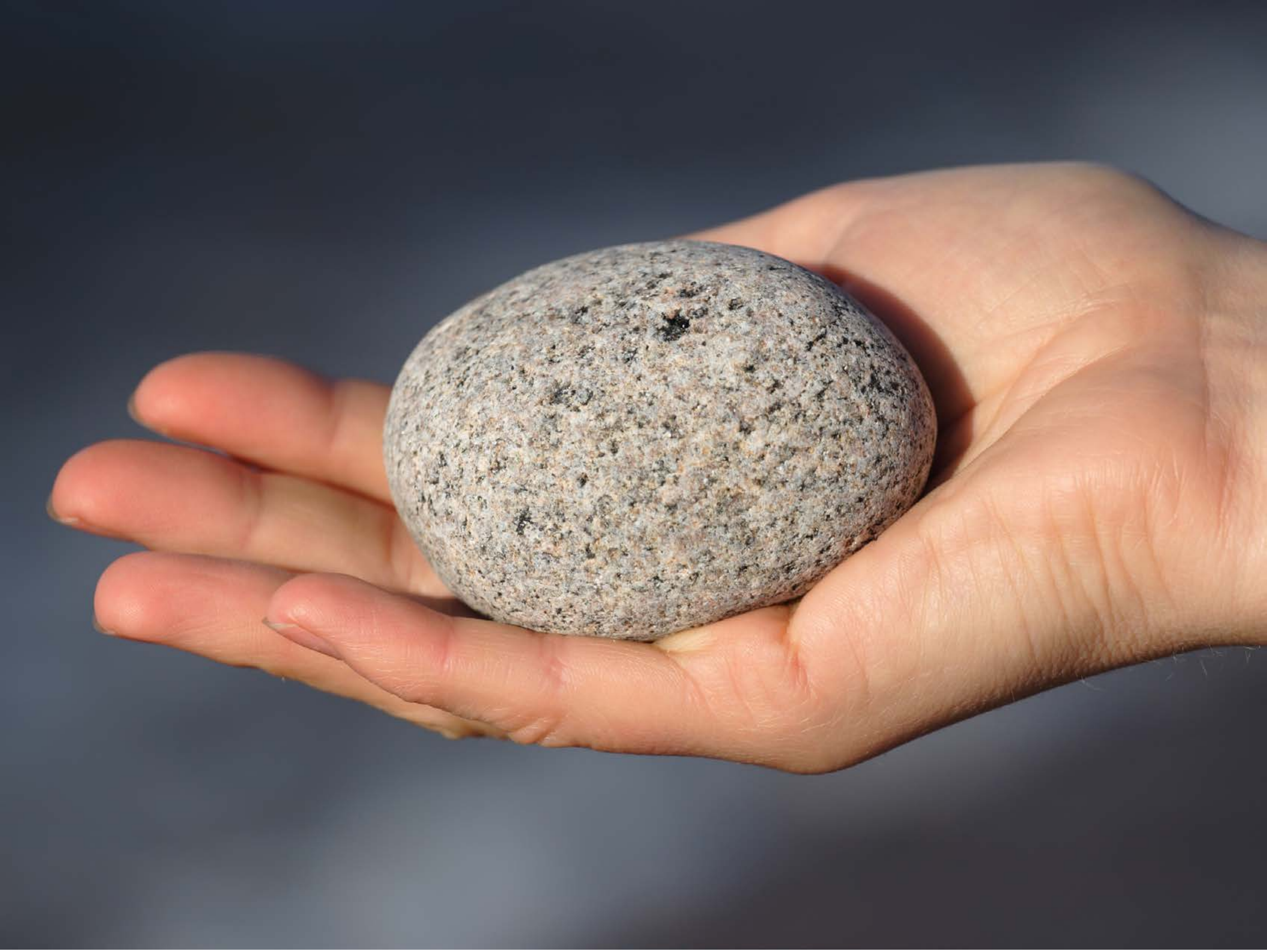

| Category | Craving (0-100) | Valence (1-9) | Arousal (1-9) | Typicality (0-100) | Relatedness | | | | | HSV | | |
|---|---|---|---|---|---|---|---|---|---|---|---|---|
| | | | | | Meth | Opioid | Both | Neither | MethToOpioid | Hue | Saturation | Value |
| Neutral objects hand | 7.37 (15.71) | 5.19 (1.039) | 2.7 (2.25) | 8.33 (15.73) | 0 | 0 | 0 | 1 | 0 | 0.387 (0.285) | 0.261 (0.133) | 0.504 (0.258) |

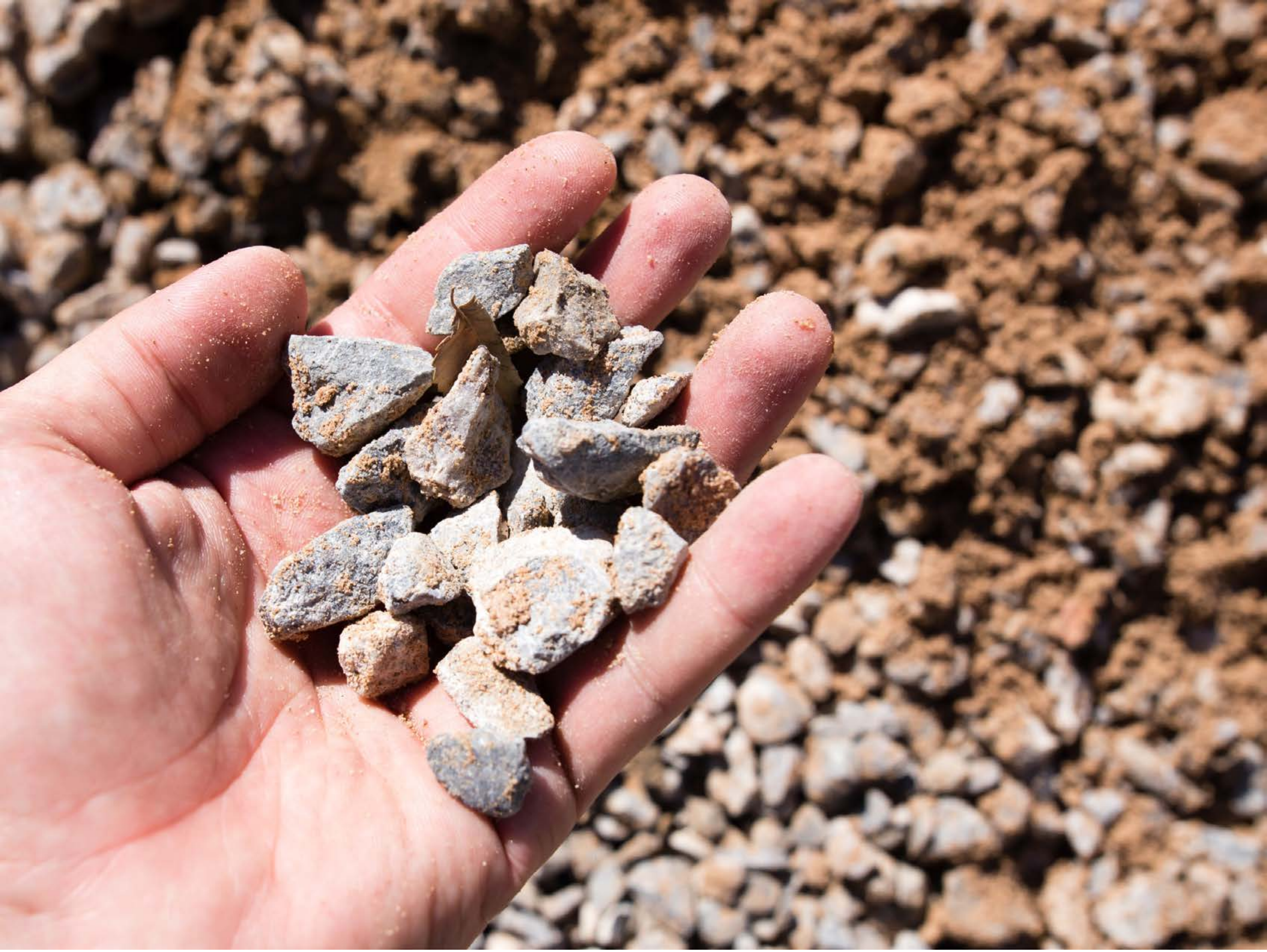

| Category | Craving (0-100) | Valence (1-9) | Arousal (1-9) | Typicality (0-100) | Relatedness: Meth | Opioid | Both | Neither | MethToOpioid | HSV: Hue | Saturation | Value |
|---|---|---|---|---|---|---|---|---|---|---|---|---|
| Neutral objects hand | 10.78 (22.01) | 4.89 (1.155) | 2.89 (2.21) | 10.93 (22.29) | 0 | 0.03704 | 0 | 0.96296 | 0.03704 | 0.104 (0.215) | 0.354 (0.178) | 0.63 (0.287) |

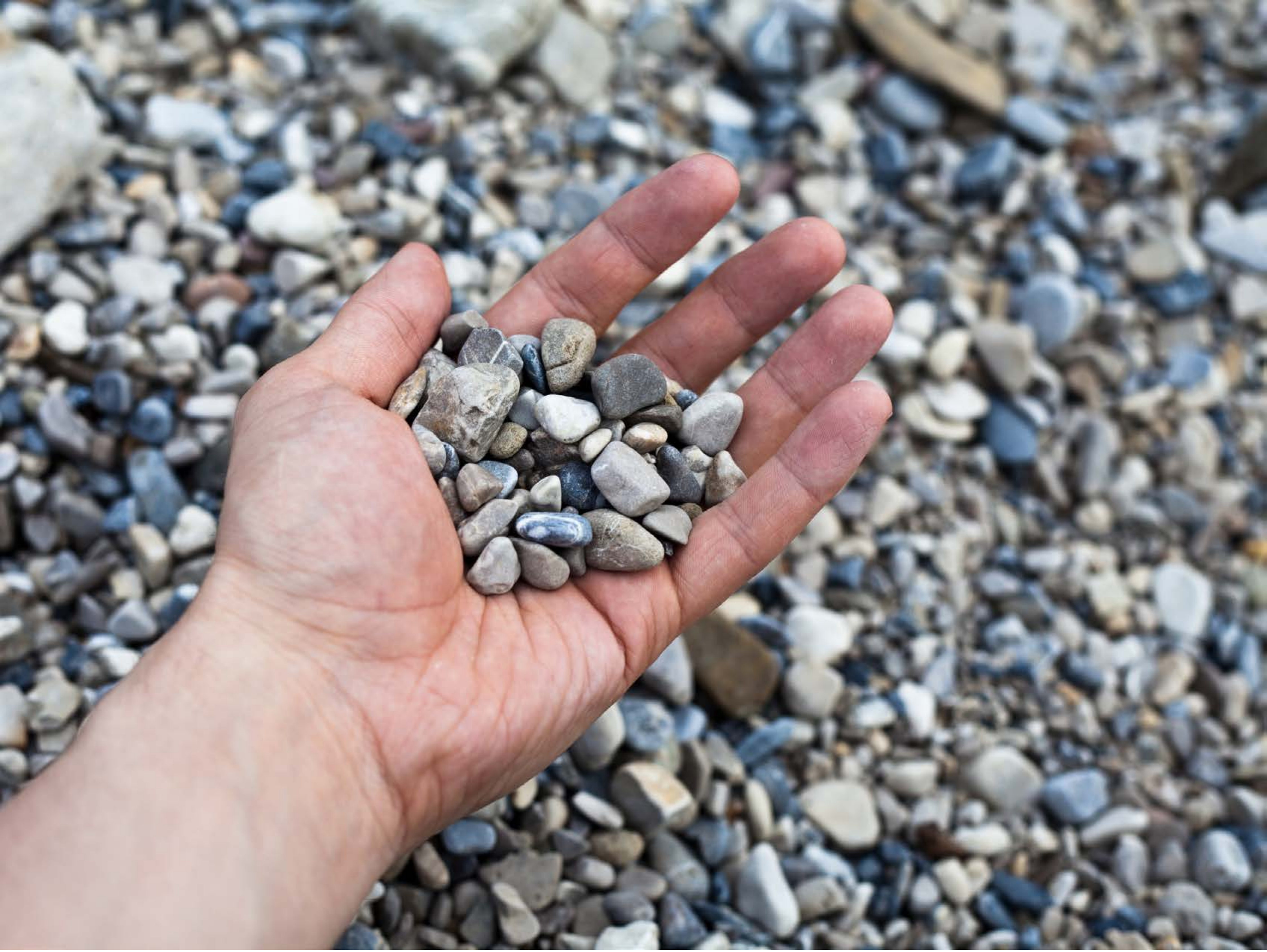

| Category | Craving (0-100) | Valence (1-9) | Arousal (1-9) | Typicality (0-100) | Relatedness | | | | | HSV | | |
|---|---|---|---|---|---|---|---|---|---|---|---|---|
| | | | | | Meth | Opioid | Both | Neither | MethToOpioid | Hue | Saturation | Value |
| Neutral objects hand | 20 (35.34) | 4.93 (1.386) | 2.36 (1.93) | 11.71 (21.53) | 0.03571 | 0 | 0 | 0.96429 | -0.03571 | 0.358 (0.291) | 0.151 (0.123) | 0.588 (0.226) |

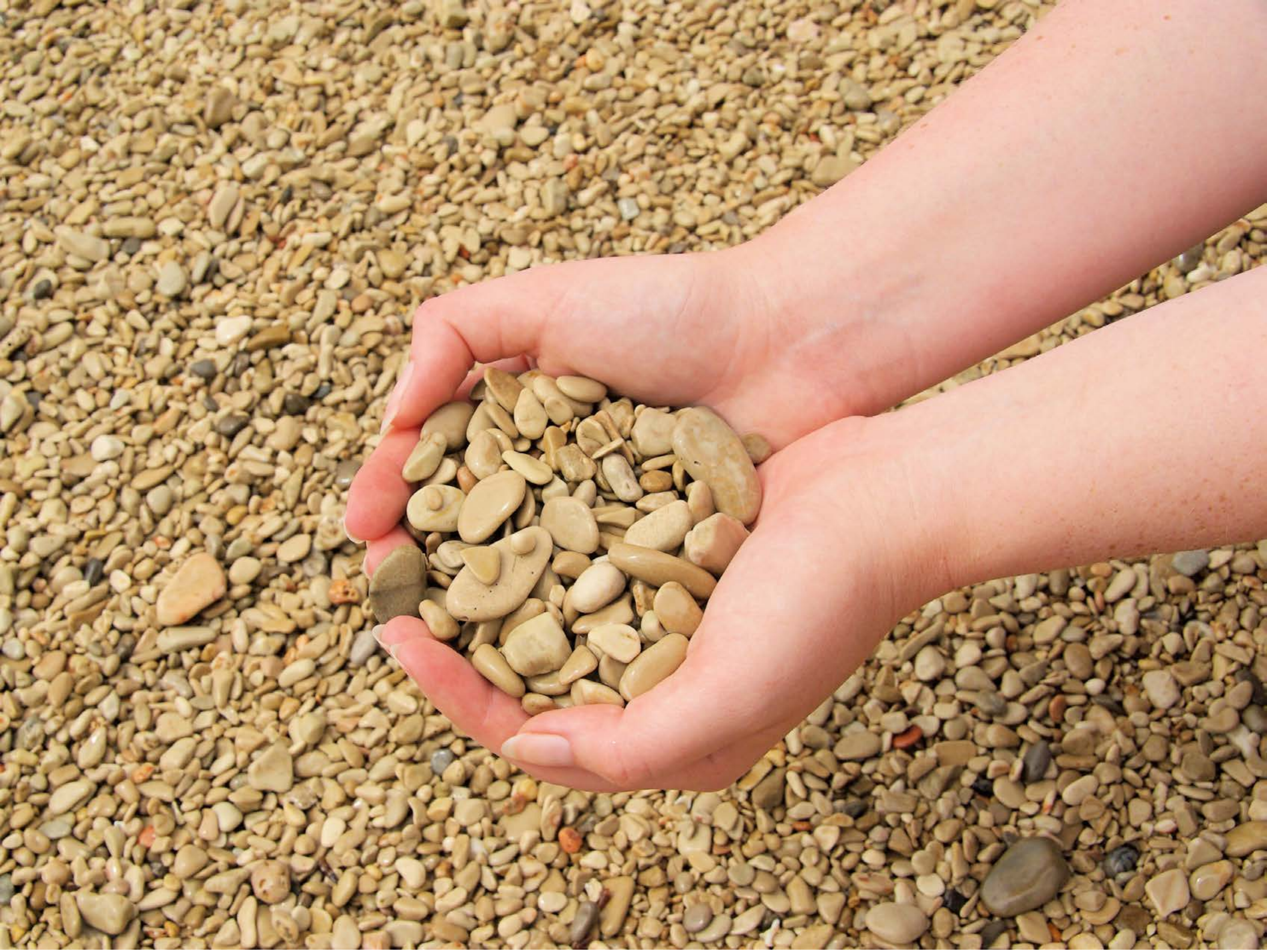

| Category | Craving (0-100) | Valence (1-9) | Arousal (1-9) | Typicality (0-100) | Relatedness | | | | | HSV | | |
|---|---|---|---|---|---|---|---|---|---|---|---|---|
| | | | | | Meth | Opioid | Both | Neither | MethToOpioid | Hue | Saturation | Value |
| Neutral objects hand | 19.67 (32.16) | 5.04 (0.437) | 2.89 (2.17) | 12.63 (22.41) | 0 | 0.03704 | 0 | 0.96296 | 0.03704 | 0.082 (0.041) | 0.429 (0.133) | 0.743 (0.193) |

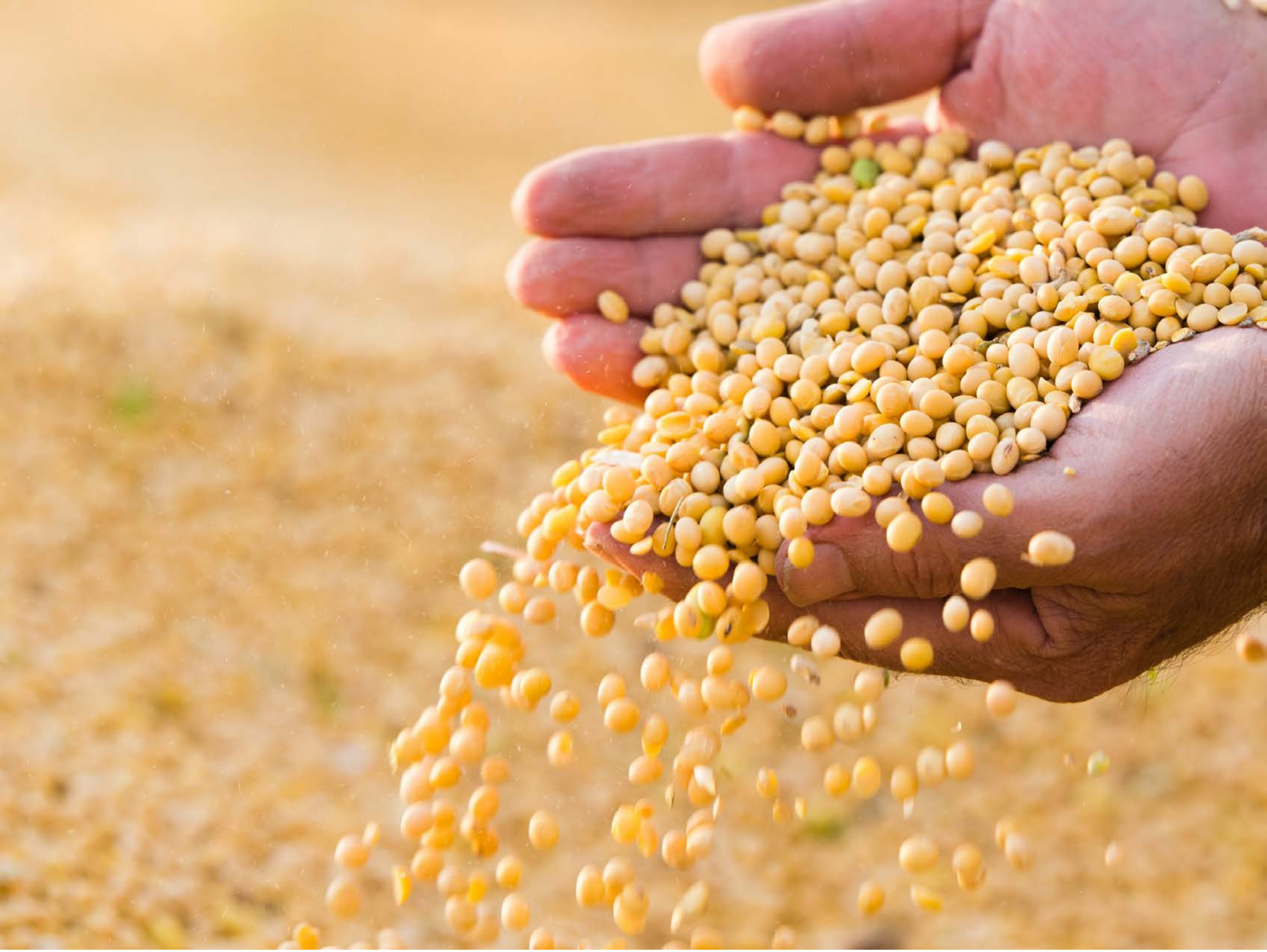

| Category | Craving (0-100) | Valence (1-9) | Arousal (1-9) | Typicality (0-100) | Relatedness | | | | | HSV | | |
|---|---|---|---|---|---|---|---|---|---|---|---|---|
| | | | | | Meth | Opioid | Both | Neither | MethToOpioid | Hue | Saturation | Value |
| Neutral objects hand | 13.15 (24) | 4.96 (0.98) | 2.67 (2.15) | 14.07 (23.86) | 0 | 0.03704 | 0 | 0.96296 | 0.03704 | 0.136 (0.206) | 0.481 (0.173) | 0.849 (0.139) |

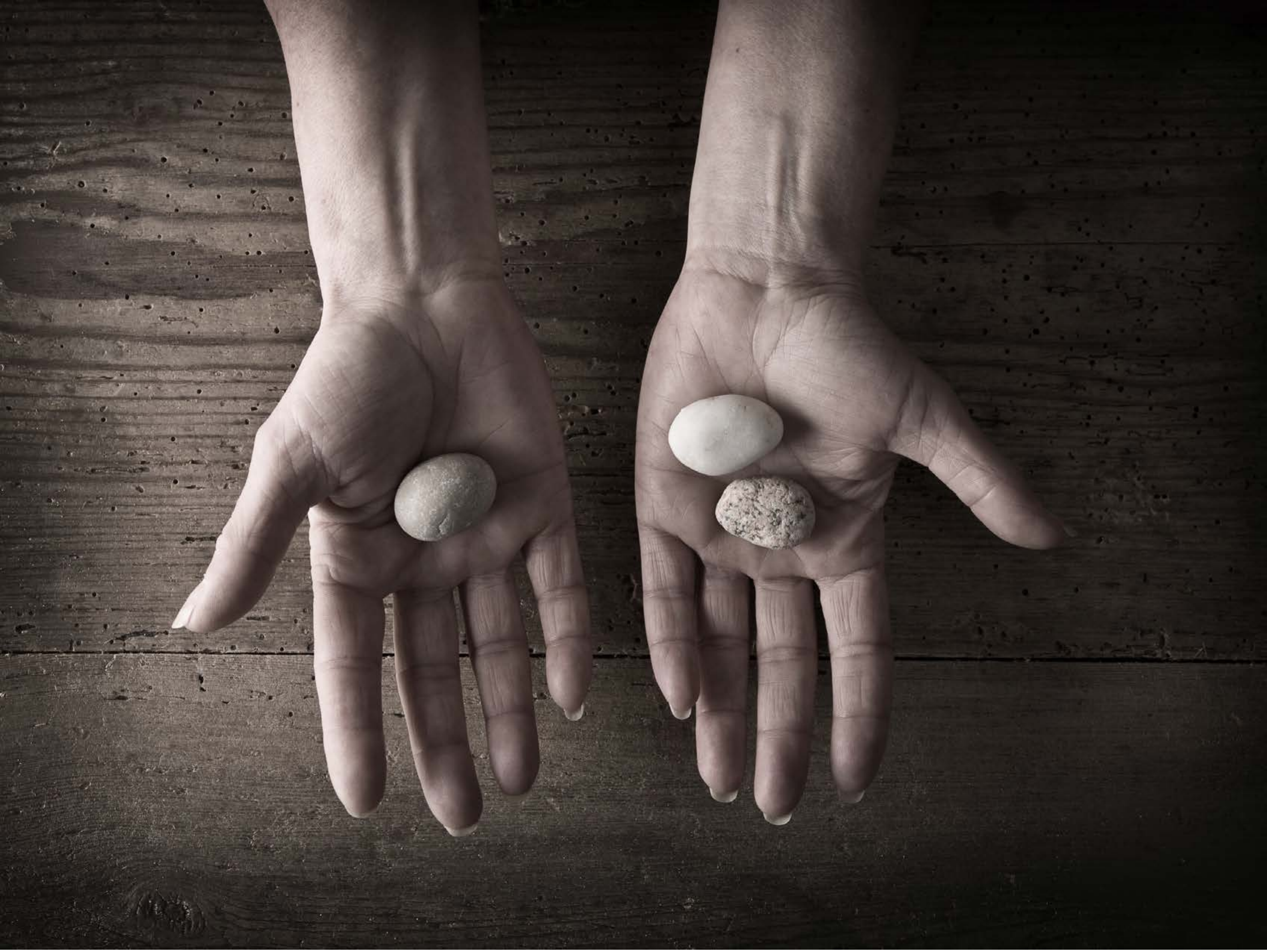

| Category | Craving (0-100) | Valence (1-9) | Arousal (1-9) | Typicality (0-100) | Relatedness | | | | | HSV | | |
|---|---|---|---|---|---|---|---|---|---|---|---|---|
| | | | | | Meth | Opioid | Both | Neither | MethToOpioid | Hue | Saturation | Value |
| Neutral objects hand | 13.43 (28.15) | 4.86 (1.627) | 2.36 (2.06) | 15.18 (28) | 0 | 0 | 0.03571 | 0.96429 | 0 | 0.064 (0.053) | 0.219 (0.081) | 0.274 (0.206) |

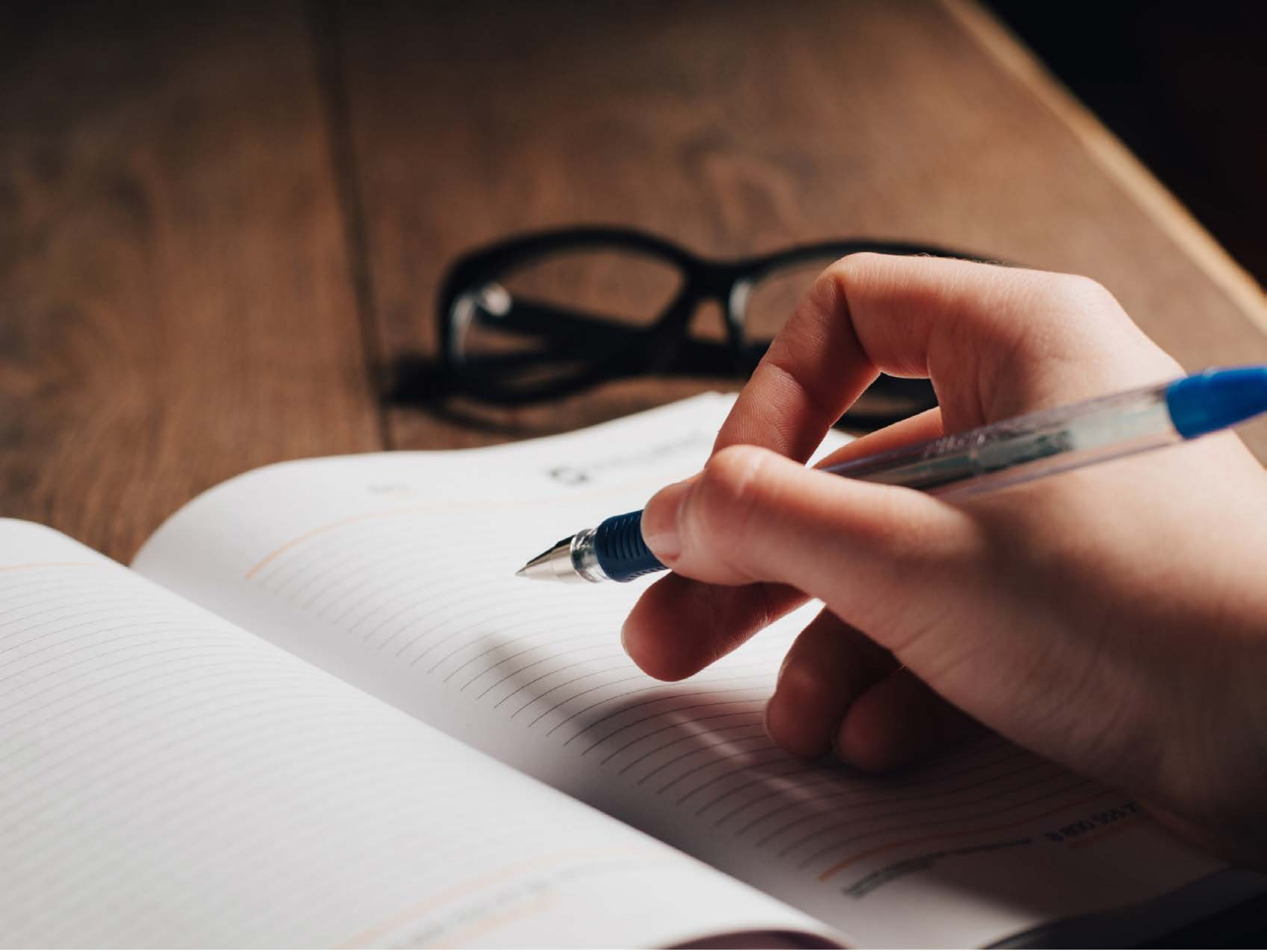

| Category | Craving (0-100) | Valence (1-9) | Arousal (1-9) | Typicality (0-100) | Relatedness | | | | | HSV | | |
|---|---|---|---|---|---|---|---|---|---|---|---|---|
| | | | | | Meth | Opioid | Both | Neither | MethToOpioid | Hue | Saturation | Value |
| Neutral objects hand | 10.64 (23.27) | 5.04 (1.815) | 2.43 (2.18) | 13.54 (22.93) | 0 | 0 | 0.03571 | 0.96429 | 0 | 0.233 (0.354) | 0.307 (0.215) | 0.497 (0.279) |

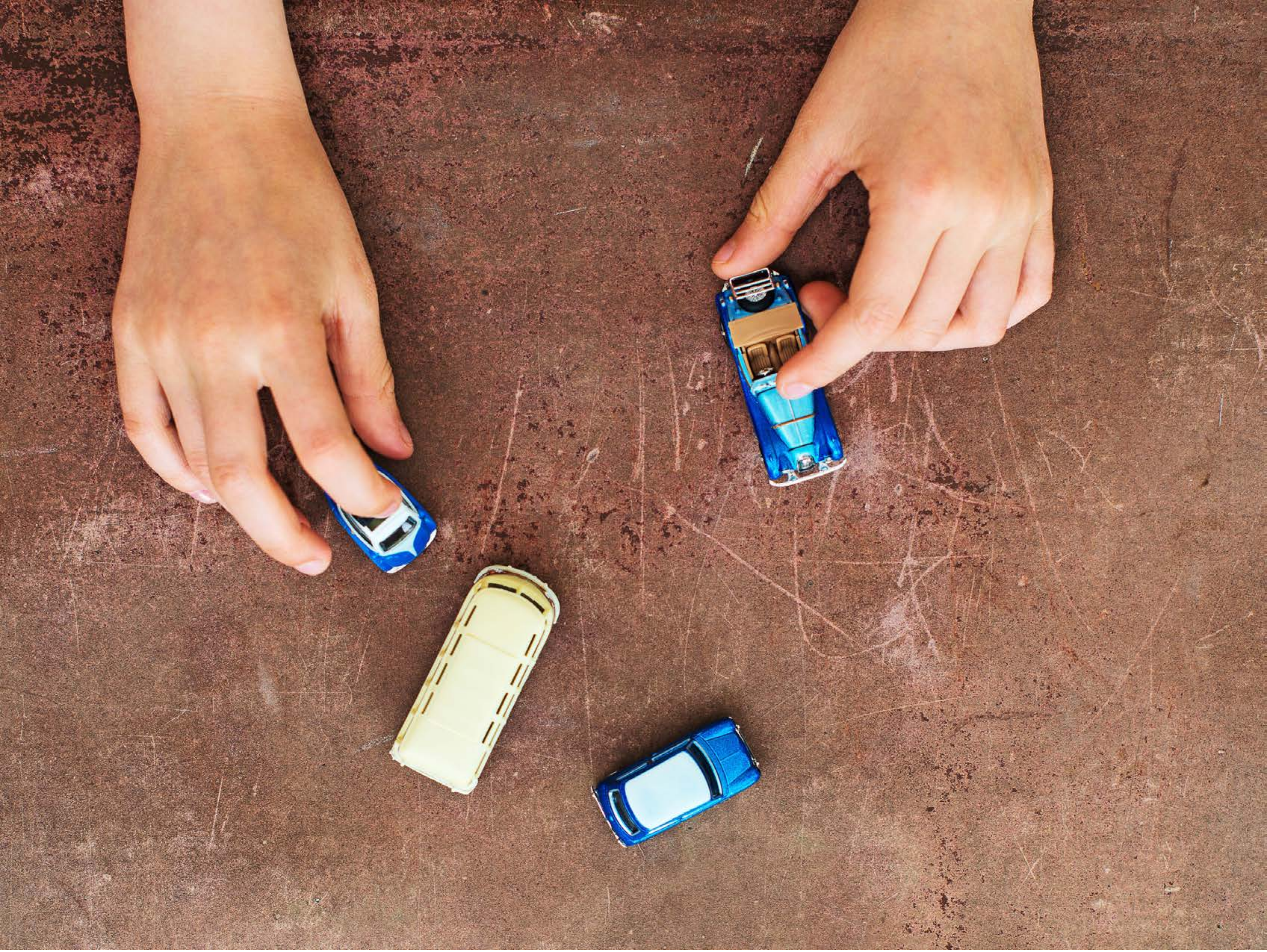

| Category | Craving (0-100) | Valence (1-9) | Arousal (1-9) | Typicality (0-100) | Relatedness | | | | | HSV | | |
|---|---|---|---|---|---|---|---|---|---|---|---|---|
| | | | | | Meth | Opioid | Both | Neither | MethToOpioid | Hue | Saturation | Value |
| Neutral objects hand | 11.71 (23.19) | 4.82 (2.056) | 2.25 (1.96) | 9.07 (14.86) | 0 | 0 | 0 | 1 | 0 | 0.068 (0.106) | 0.437 (0.124) | 0.623 (0.186) |

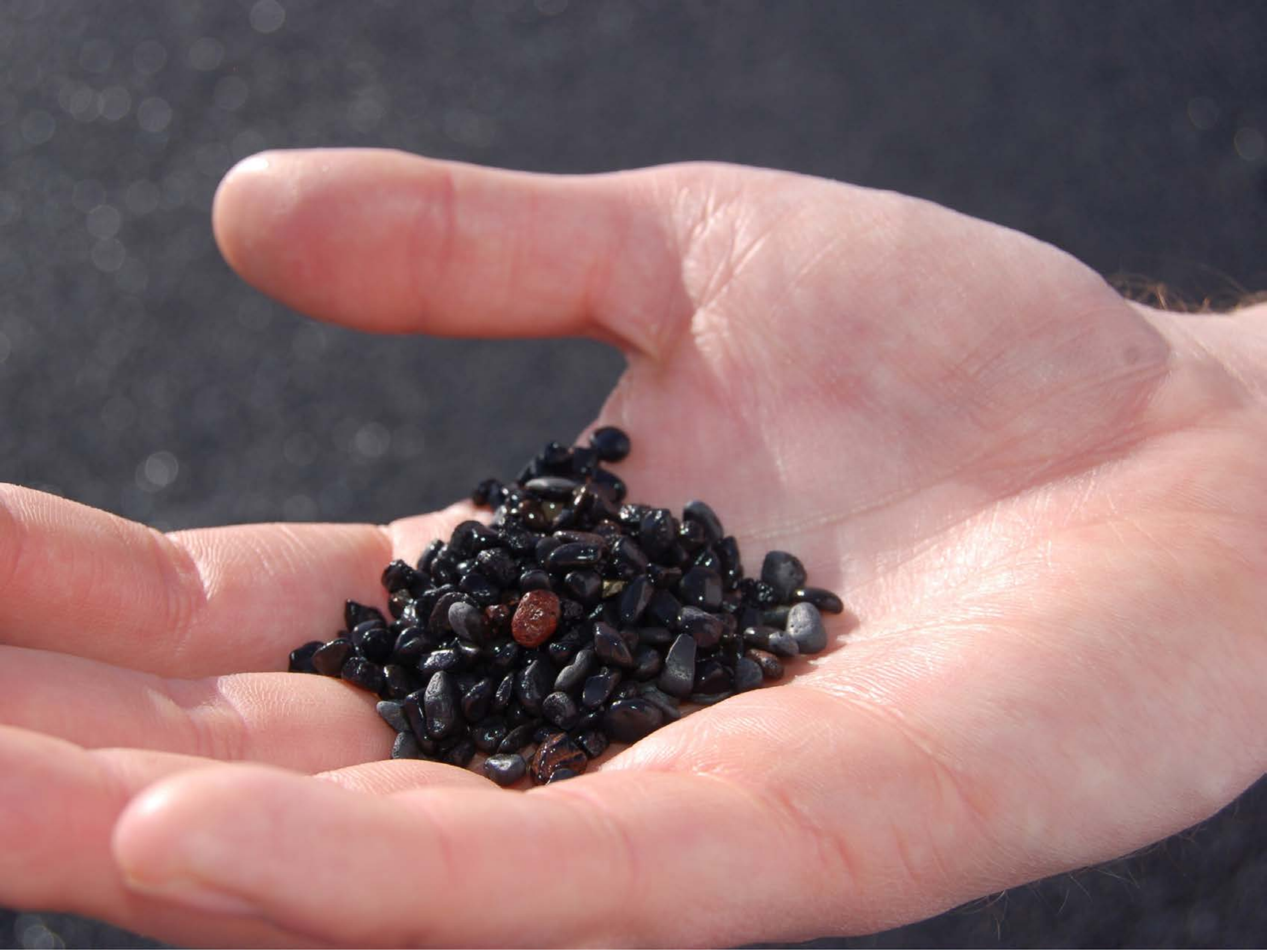

| Category | Craving (0-100) | Valence (1-9) | Arousal (1-9) | Typicality (0-100) | Relatedness | | | | | HSV | | |
|---|---|---|---|---|---|---|---|---|---|---|---|---|
| | | | | | Meth | Opioid | Both | Neither | MethToOpioid | Hue | Saturation | Value |
| Neutral objects hand | 15.15 (22.94) | 4.63 (1.363) | 2.63 (2.08) | 15 (23.82) | 0 | 0.03704 | 0 | 0.96296 | 0.03704 | 0.374 (0.371) | 0.233 (0.113) | 0.562 (0.272) |

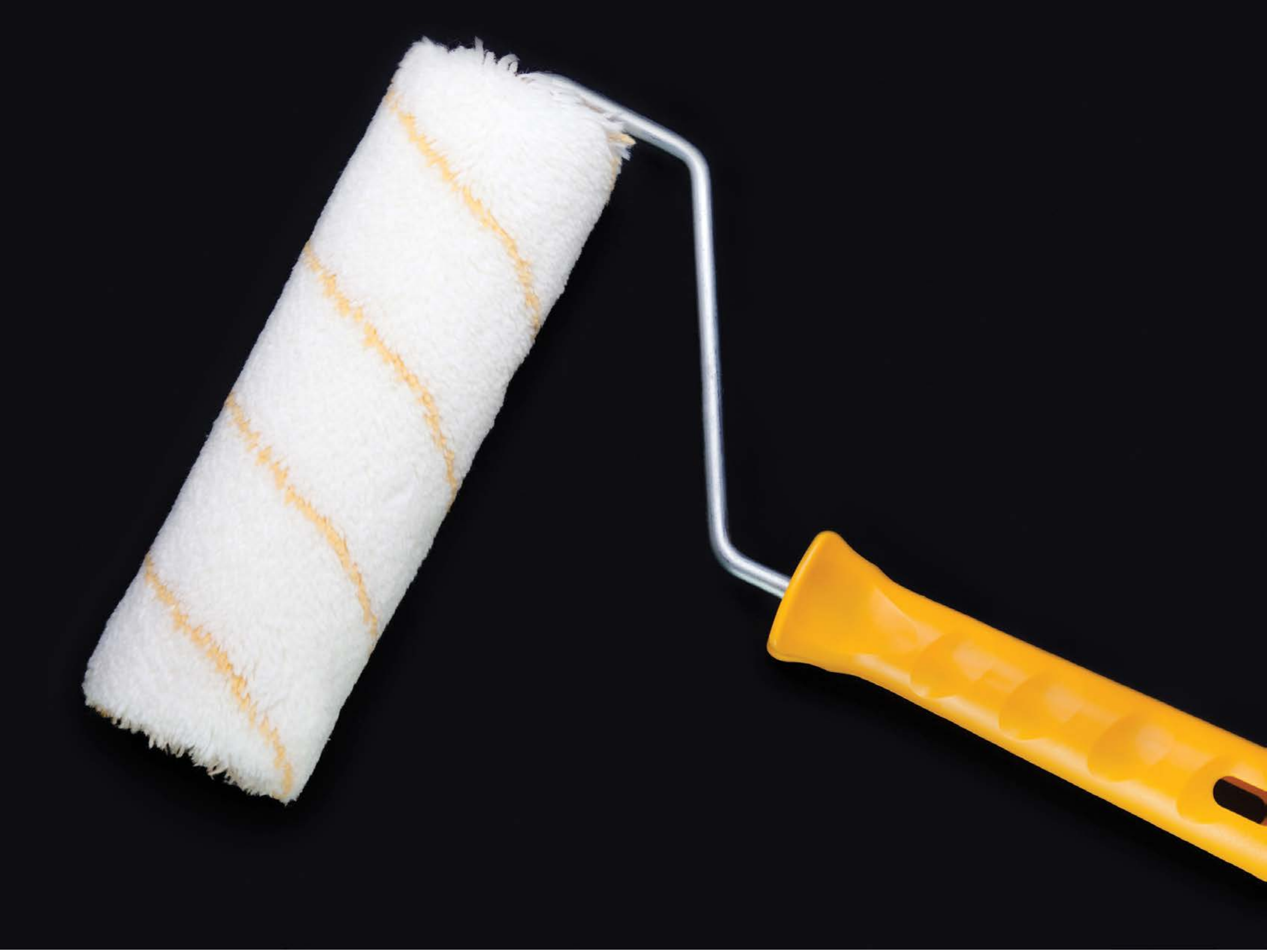

| Category | Craving (0-100) | Valence (1-9) | Arousal (1-9) | Typicality (0-100) | Relatedness | | | | | HSV | | |
|---|---|---|---|---|---|---|---|---|---|---|---|---|
| | | | | | Meth | Opioid | Both | Neither | MethToOpioid | Hue | Saturation | Value |
| Neutral tool | 7.78 (15.89) | 4.78 (1.155) | 2.52 (1.95) | 10.81 (17.63) | 0 | 0 | 0 | 1 | 0 | 0.617 (0.204) | 0.317 (0.184) | 0.241 (0.348) |

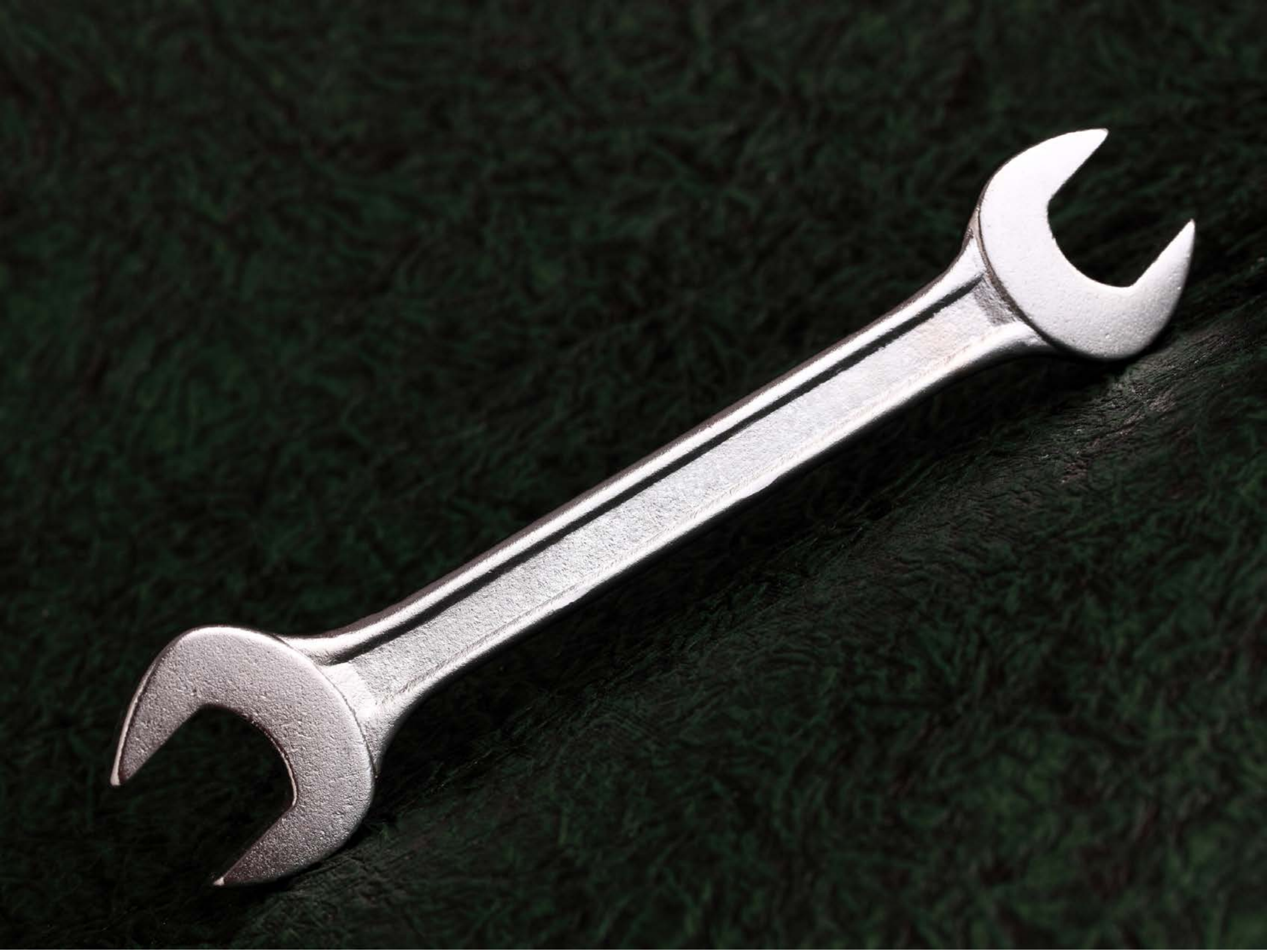

| Category | Craving (0-100) | Valence (1-9) | Arousal (1-9) | Typicality (0-100) | Relatedness | | | | | HSV | | |
|---|---|---|---|---|---|---|---|---|---|---|---|---|
| | | | | | Meth | Opioid | Both | Neither | MethToOpioid | Hue | Saturation | Value |
| Neutral tool | 13.25 (22.71) | 5 (1.515) | 2.93 (2.14) | 12.79 (21.4) | 0.03571 | 0 | 0 | 0.96429 | -0.03571 | 0.292 (0.208) | 0.227 (0.11) | 0.163 (0.22) |

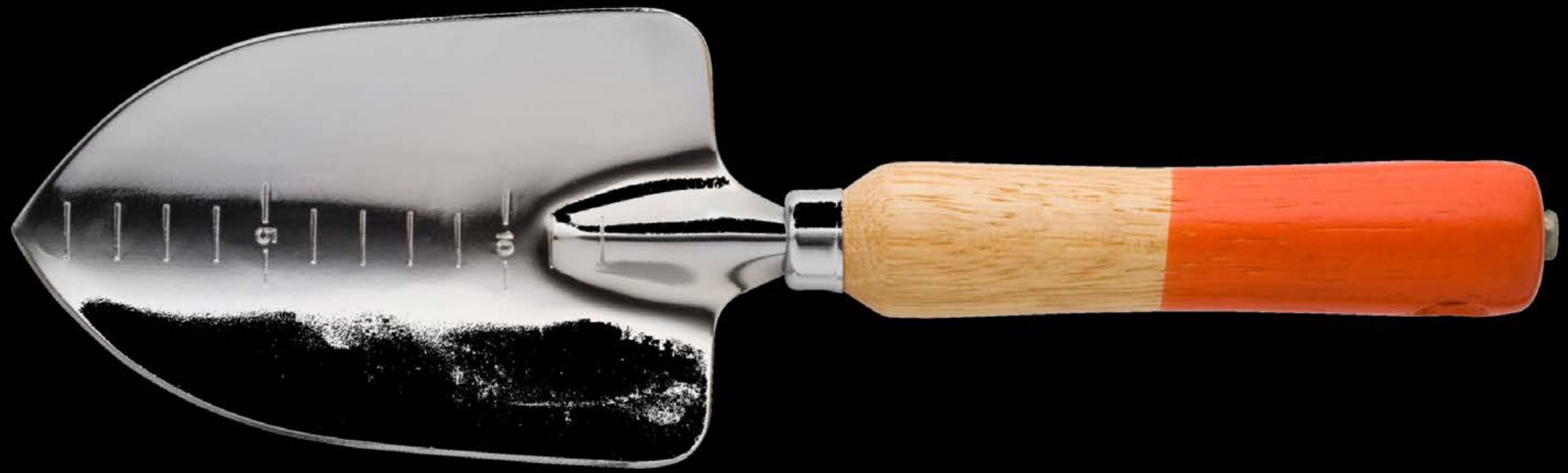


| Category | Craving (0-100) | Valence (1-9) | Arousal (1-9) | Typicality (0-100) | Relatedness | | | | | HSV | | |
|---|---|---|---|---|---|---|---|---|---|---|---|---|
| | | | | | Meth | Opioid | Both | Neither | MethToOpioid | Hue | Saturation | Value |
| Neutral tool | 5.32 (13) | 4.68 (1.744) | 2.11 (1.81) | 8.04 (15.65) | 0 | 0 | 0 | 1 | 0 | 0.025 (0.127) | 0.032 (0.147) | 0.085 (0.241) |

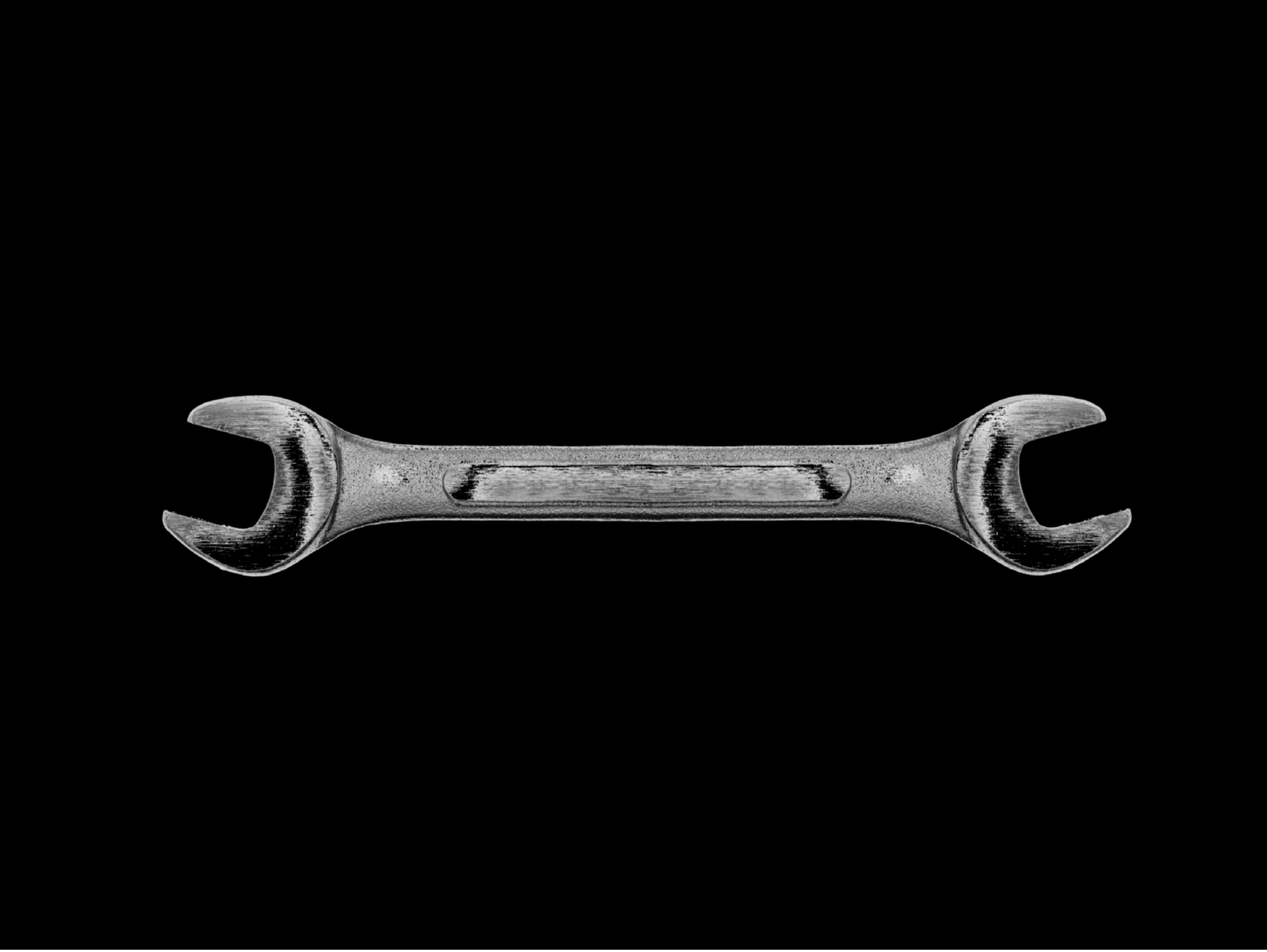

| Category | Craving (0-100) | Valence (1-9) | Arousal (1-9) | Typicality (0-100) | Relatedness | | | | | HSV | | |
|---|---|---|---|---|---|---|---|---|---|---|---|---|
| | | | | | Meth | Opioid | Both | Neither | MethToOpioid | Hue | Saturation | Value |
| Neutral tool | 15.78 (25.74) | 5 (1.109) | 2.85 (2.14) | 15.89 (25.45) | 0.03704 | 0 | 0.96296 | 0 | -0.03704 | 0 (0) | 0 (0) | 0.033 (0.135) |

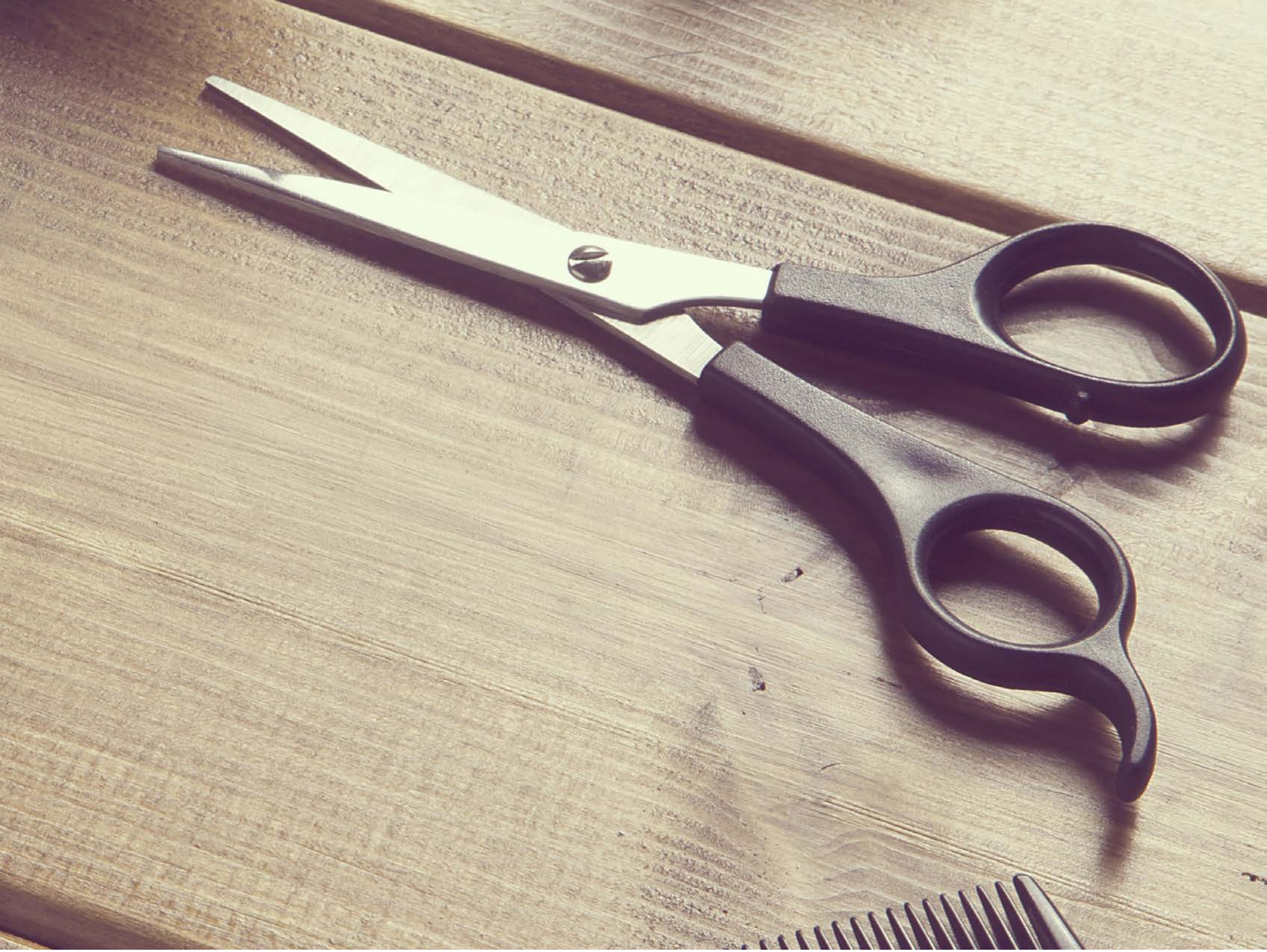

| Category | Craving (0-100) | Valence (1-9) | Arousal (1-9) | Typicality (0-100) | Relatedness | | | | | HSV | | |
|---|---|---|---|---|---|---|---|---|---|---|---|---|
| | | | | | Meth | Opioid | Both | Neither | MethToOpioid | Hue | Saturation | Value |
| Neutral tool | 6.46 (13.57) | 4.75 (1.691) | 2.14 (1.72) | 7.18 (11.09) | 0 | 0 | 0 | 1 | 0 | 0.247 (0.298) | 0.204 (0.083) | 0.719 (0.19) |

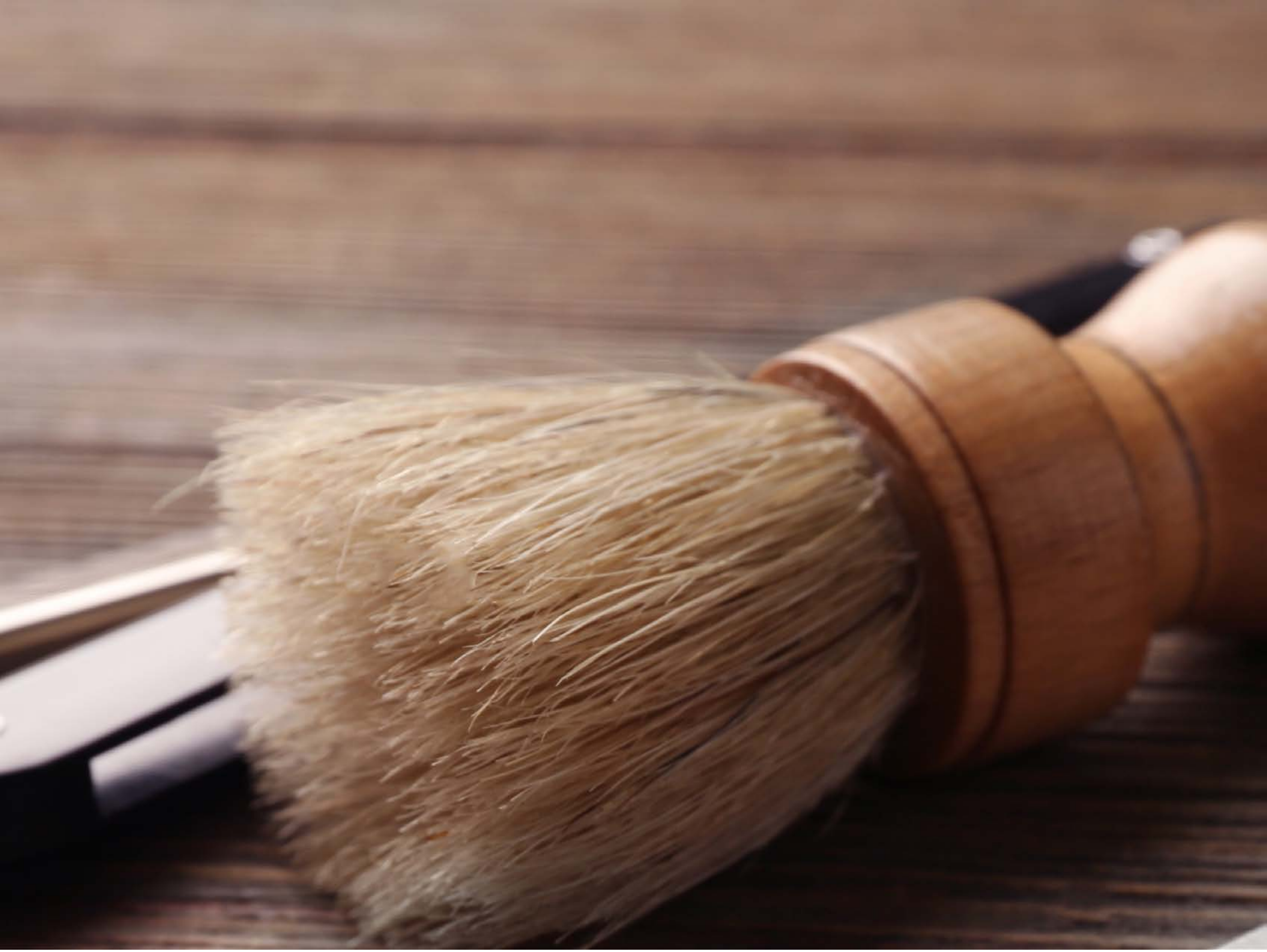

| Category | Craving (0-100) | Valence (1-9) | Arousal (1-9) | Typicality (0-100) | Relatedness | | | | | HSV | | |
|---|---|---|---|---|---|---|---|---|---|---|---|---|
| | | | | | Meth | Opioid | Both | Neither | MethToOpioid | Hue | Saturation | Value |
| Neutral tool | 9.11 (17.39) | 4.86 (1.557) | 2.36 (1.87) | 9.11 (17.28) | 0 | 0 | 0 | 1 | 0 | 0.101 (0.197) | 0.366 (0.161) | 0.508 (0.233) |

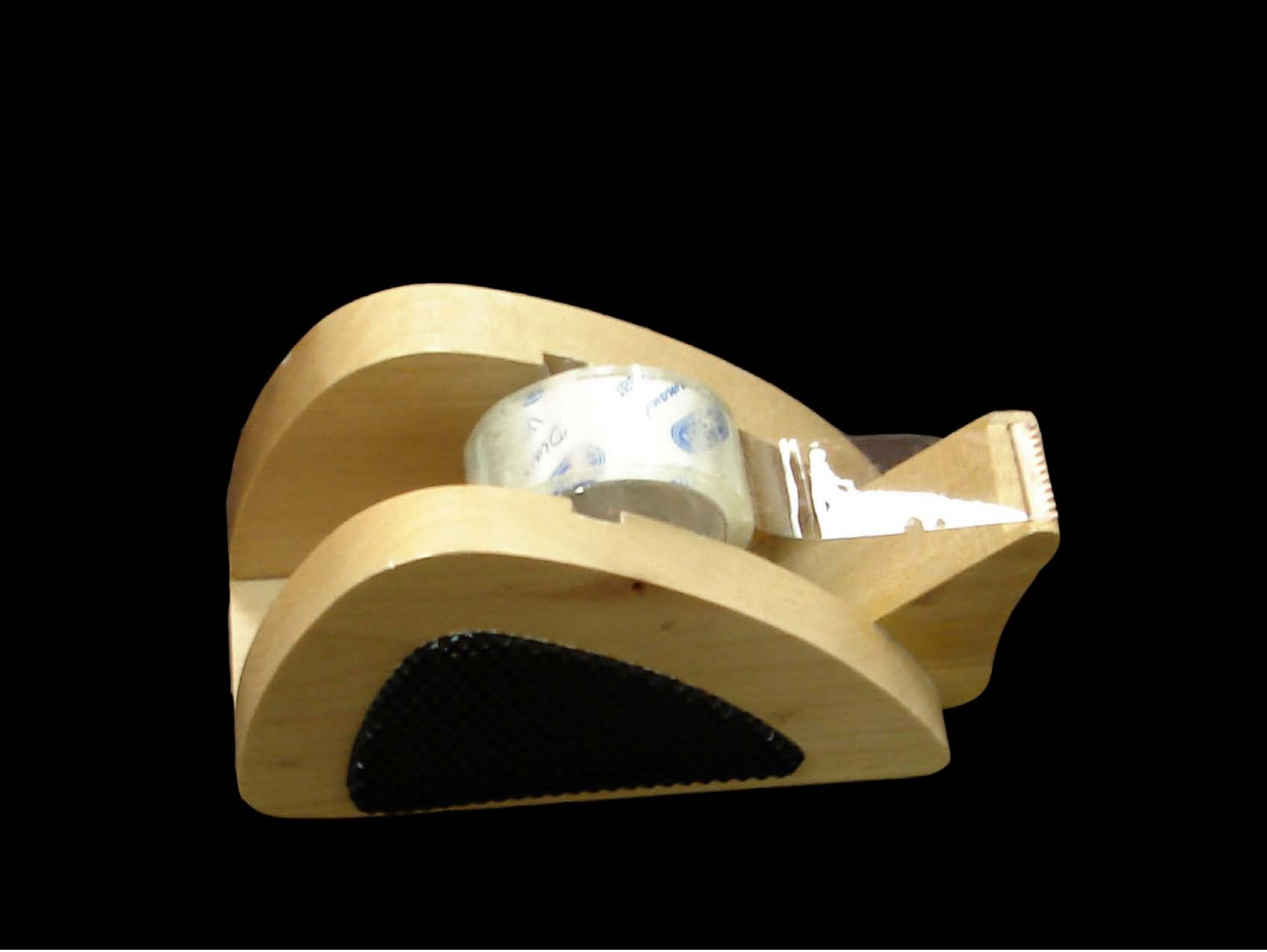

| Category | Craving (0-100) | Valence (1-9) | Arousal (1-9) | Typicality (0-100) | Relatedness | | | | | HSV | | |
|---|---|---|---|---|---|---|---|---|---|---|---|---|
| | | | | | Meth | Opioid | Both | Neither | MethToOpioid | Hue | Saturation | Value |
| Neutral tool | 9.63 (17.93) | 4.85 (1.35) | 2.37 (1.84) | 10.04 (17.91) | 0 | 0 | 0 | 1 | 0 | 0.038 (0.085) | 0.127 (0.238) | 0.166 (0.309) |

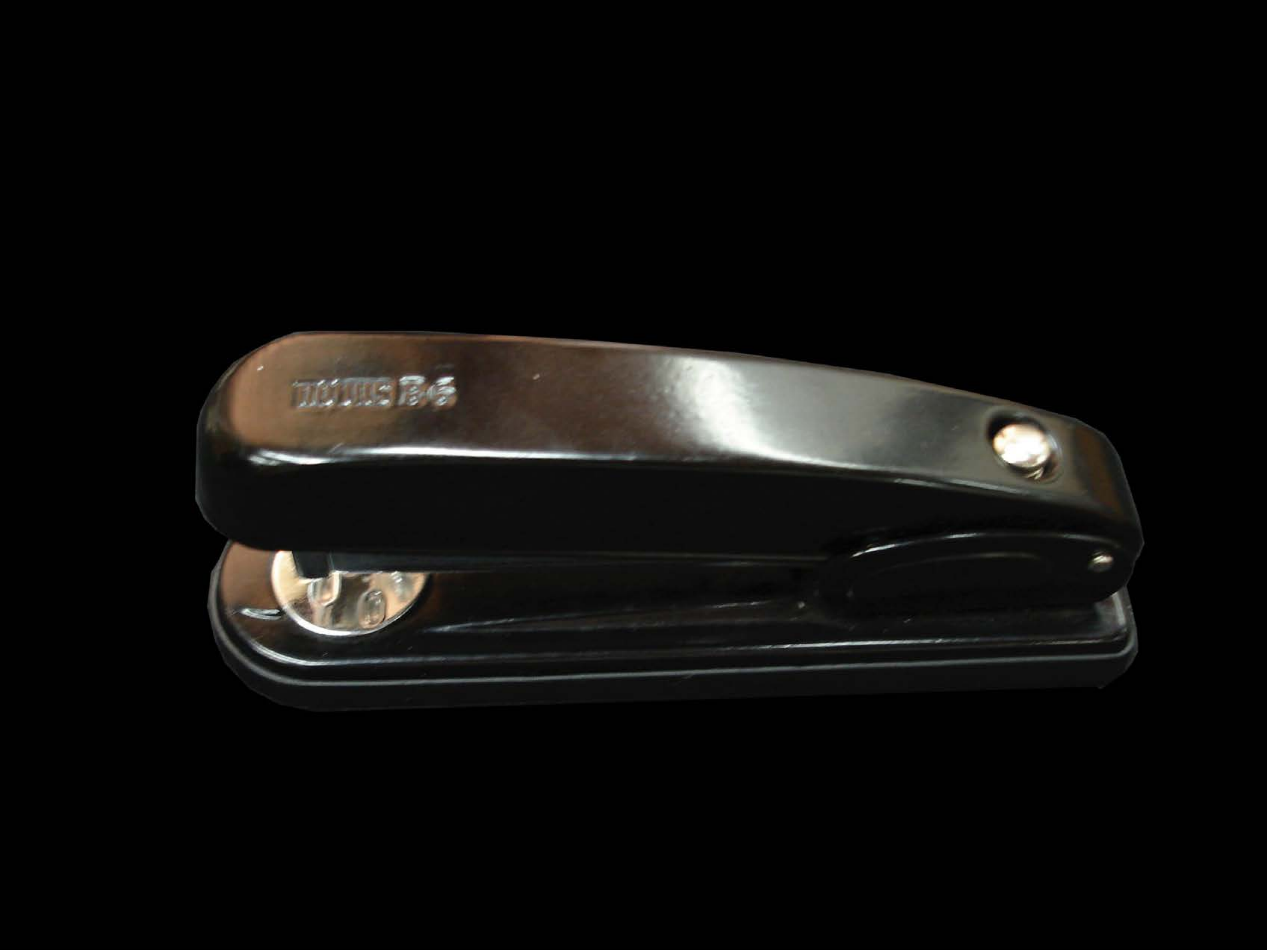

| | | | | | Relatedness | | | | | HSV | | |
|---|---|---|---|---|---|---|---|---|---|---|---|---|
| **Category** | **Craving (0-100)** | **Valence (1-9)** | **Arousal (1-9)** | **Typicality (0-100)** | Meth | Opioid | Both | Neither | MethToOpioid | Hue | Saturation | Value |
| Neutral tool | 12.59 (24.85) | 4.78 (0.801) | 2.59 (1.91) | 14.93 (25.51) | 0 | 0 | 0 | 1 | 0 | 0.045 (0.13) | 0.037 (0.101) | 0.056 (0.149) |

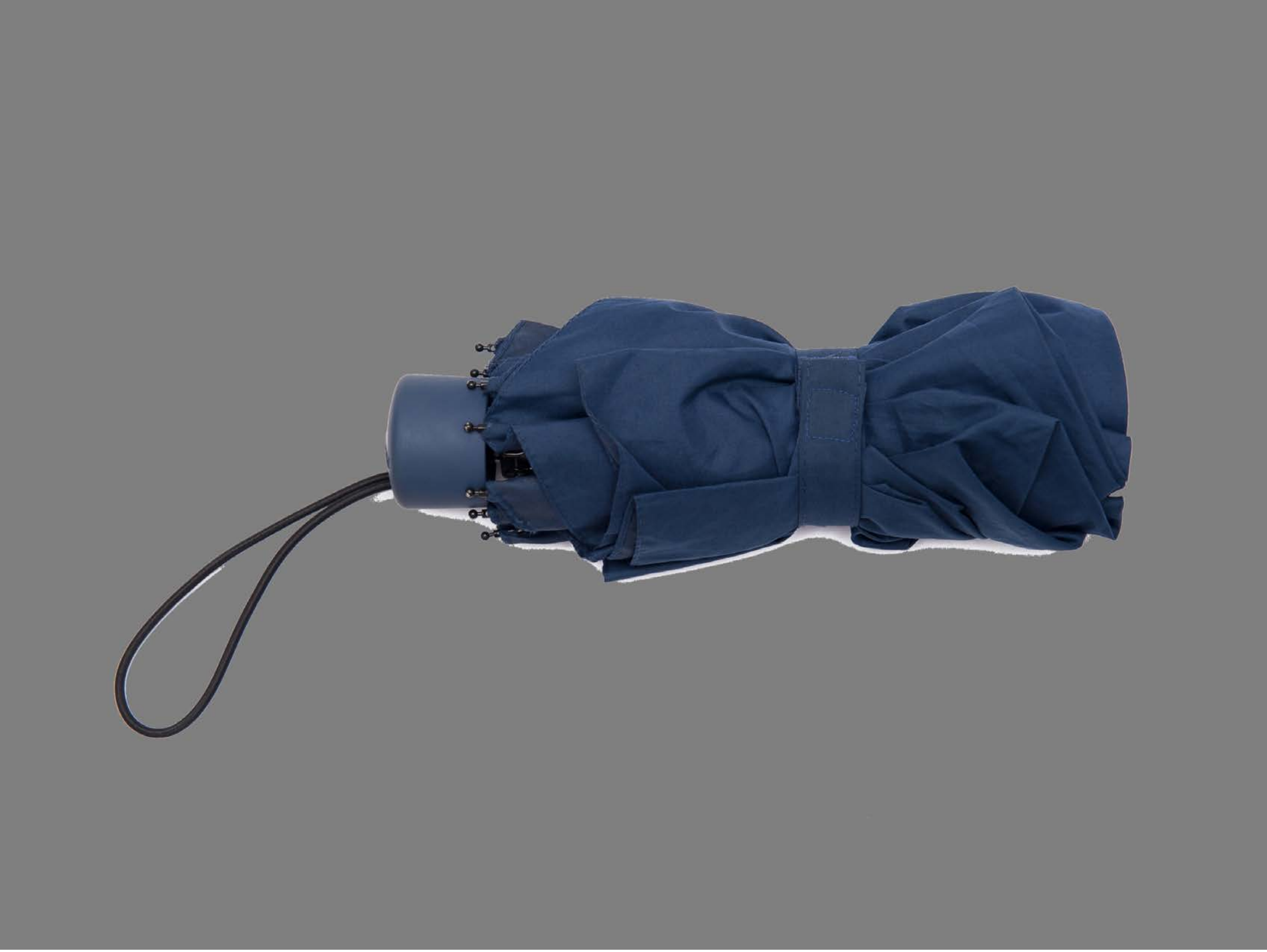

| Category | Craving (0-100) | Valence (1-9) | Arousal (1-9) | Typicality (0-100) | Relatedness | | | | | HSV | | |
|---|---|---|---|---|---|---|---|---|---|---|---|---|
| | | | | | Meth | Opioid | Both | Neither | MethToOpioid | Hue | Saturation | Value |
| Neutral tool | 11.33 (19.24) | 4.7 (1.068) | 2.44 (1.89) | 13.81 (20.26) | 0 | 0 | 0 | 1 | 0 | 0.092 (0.221) | 0.075 (0.192) | 0.474 (0.074) |

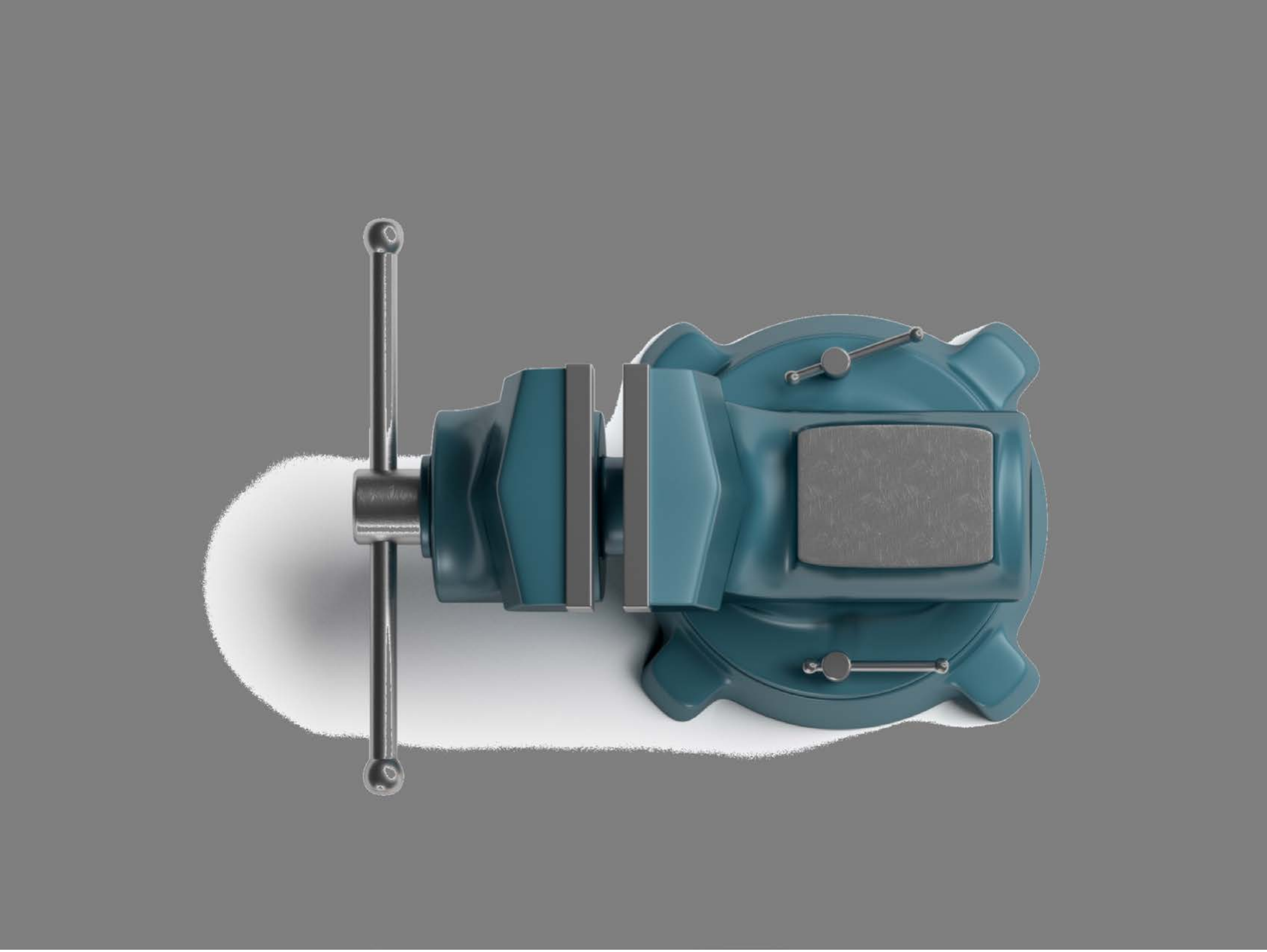

| Category | Craving (0-100) | Valence (1-9) | Arousal (1-9) | Typicality (0-100) | Relatedness | | | | | HSV | | |
|---|---|---|---|---|---|---|---|---|---|---|---|---|
| | | | | | Meth | Opioid | Both | Neither | MethToOpioid | Hue | Saturation | Value |
| Neutral tool | 14.19 (21.69) | 4.96 (0.192) | 2.7 (1.96) | 12.56 (20.52) | 0 | 0 | 0 | 1 | 0 | 0.099 (0.214) | 0.058 (0.156) | 0.504 (0.102) |

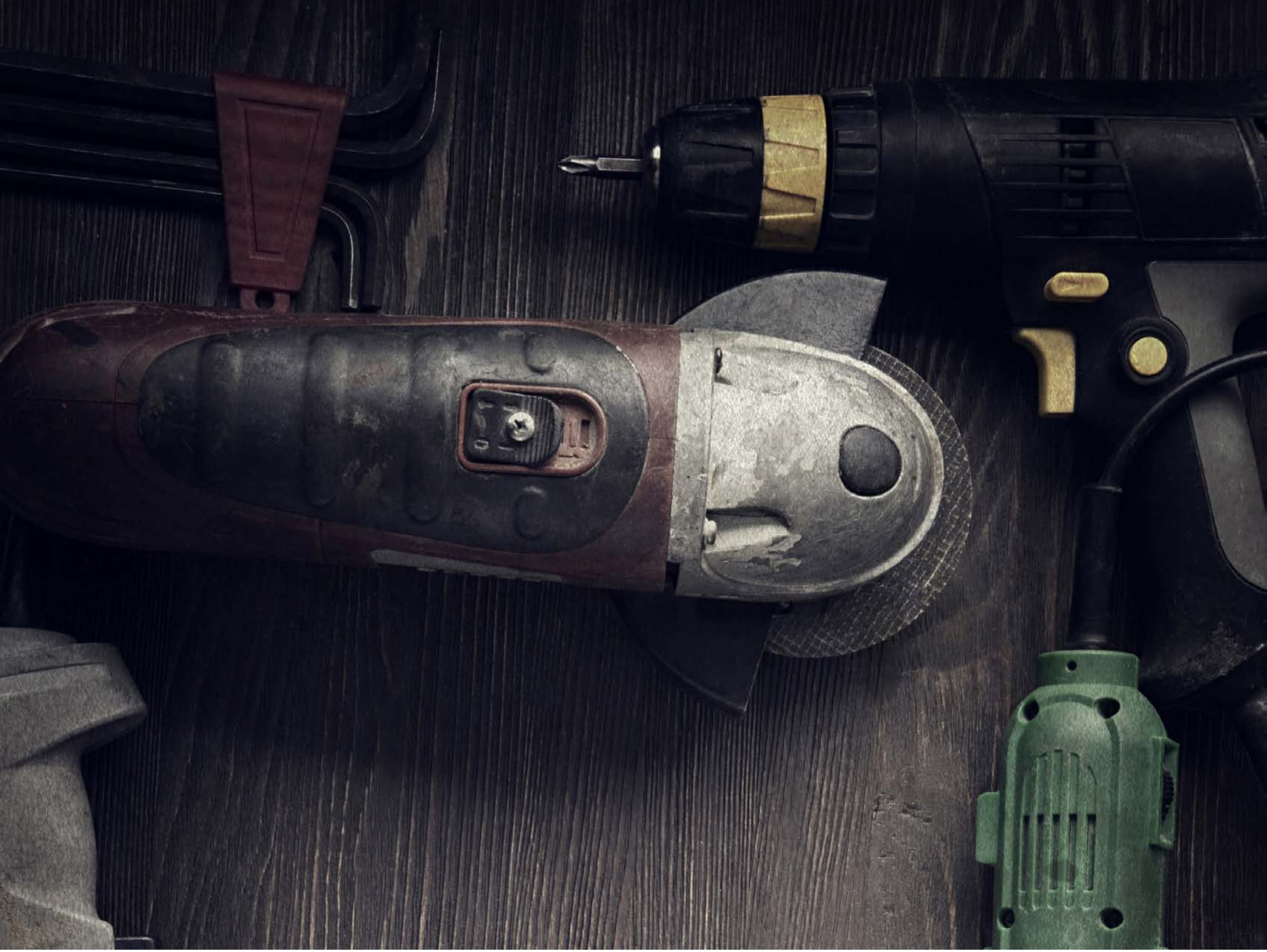

| Category | Craving (0-100) | Valence (1-9) | Arousal (1-9) | Typicality (0-100) | Relatedness | | | | | HSV | | |
|---|---|---|---|---|---|---|---|---|---|---|---|---|
| | | | | | Meth | Opioid | Both | Neither | MethToOpioid | Hue | Saturation | Value |
| Neutral tool | 8.64 (17.34) | 4.89 (1.343) | 2.39 (1.87) | 10.21 (17.32) | 0 | 0 | 0 | 1 | 0 | 0.613 (0.226) | 0.336 (0.245) | 0.201 (0.155) |

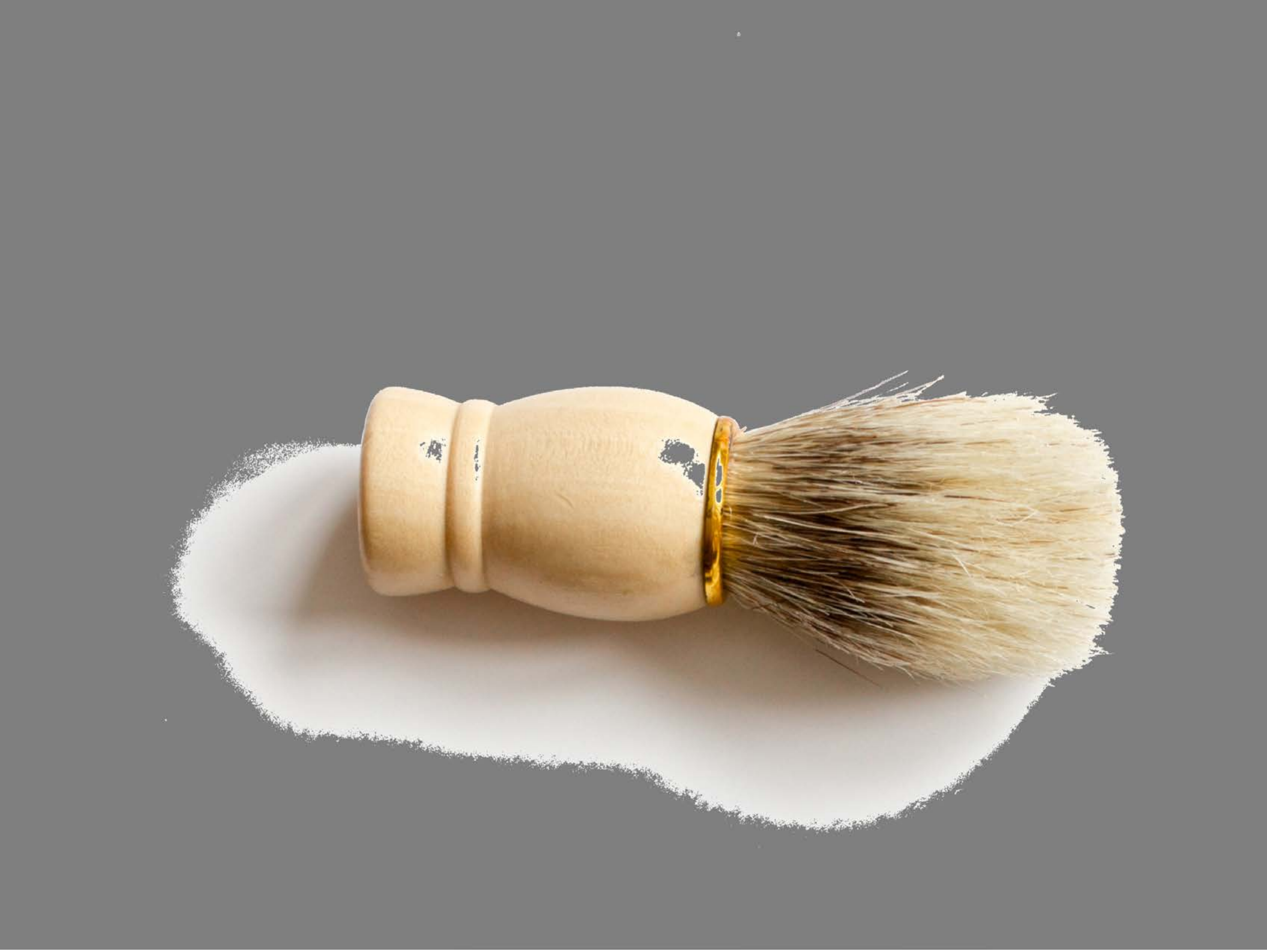

| Category | Craving (0-100) | Valence (1-9) | Arousal (1-9) | Typicality (0-100) | Relatedness | | | | | HSV | | |
|---|---|---|---|---|---|---|---|---|---|---|---|---|
| | | | | | Meth | Opioid | Both | Neither | MethToOpioid | Hue | Saturation | Value |
| Neutral tool | 11.96 (19.52) | 5.15 (0.77) | 2.52 (1.95) | 12.19 (19.5) | 0 | 0 | 0 | 1 | 0 | 0.027 (0.062) | 0.07 (0.162) | 0.566 (0.146) |

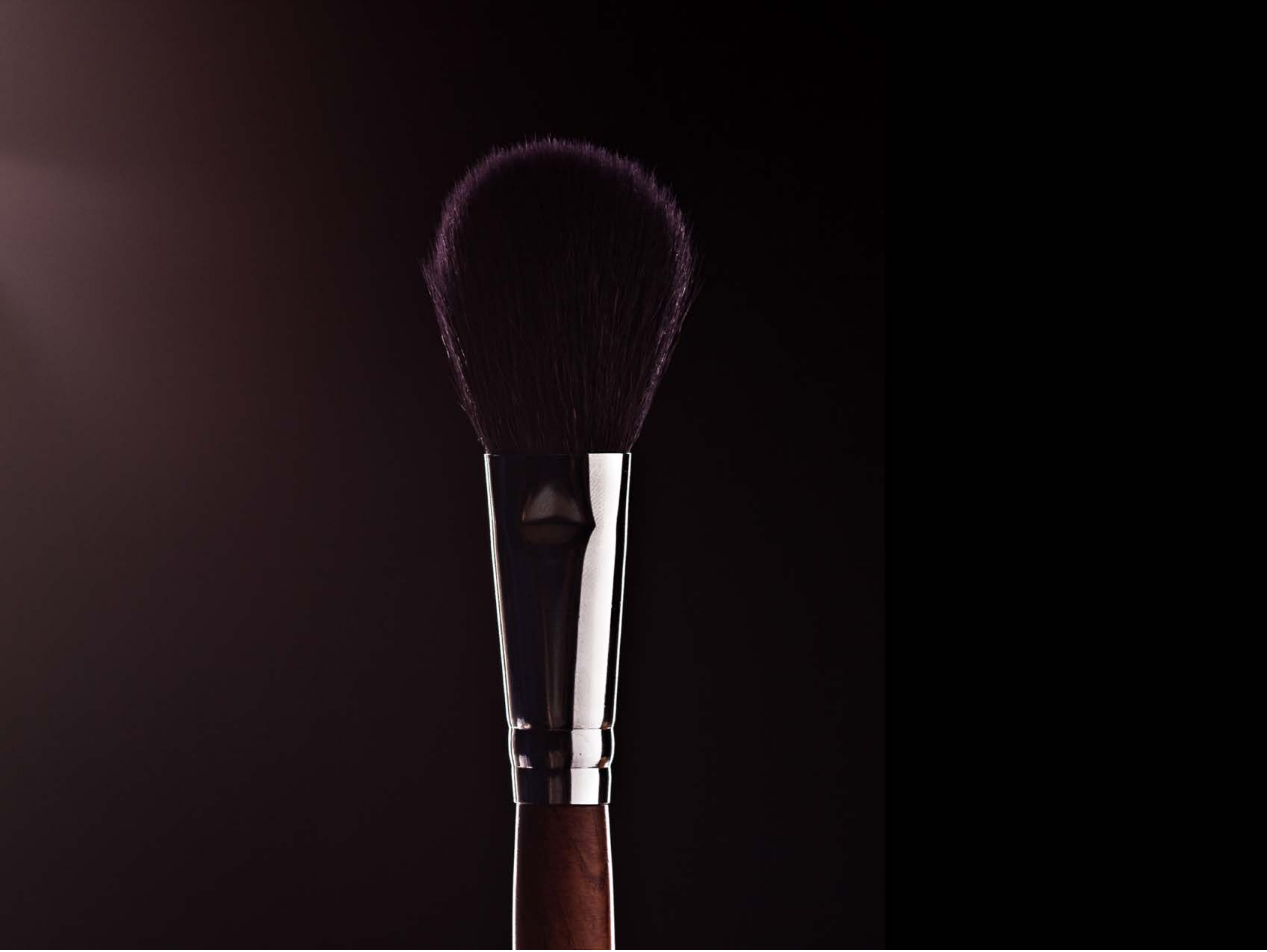

| Category | Craving (0-100) | Valence (1-9) | Arousal (1-9) | Typicality (0-100) | Relatedness | | | | | HSV | | |
|---|---|---|---|---|---|---|---|---|---|---|---|---|
| | | | | | Meth | Opioid | Both | Neither | MethToOpioid | Hue | Saturation | Value |
| Neutral tool | 11.59 (19.79) | 4.93 (0.385) | 2.67 (1.98) | 12.11 (19.49) | 0.03704 | 0 | 0.03704 | 0.92593 | -0.03704 | 0.647 (0.441) | 0.303 (0.231) | 0.086 (0.119) |

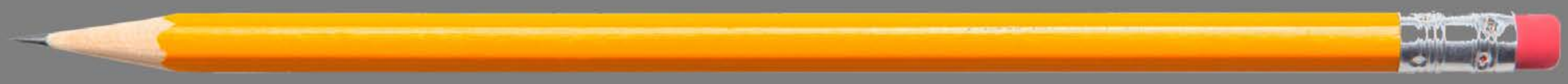

| Category | Craving (0-100) | Valence (1-9) | Arousal (1-9) | Typicality (0-100) | Relatedness | | | | | HSV | | |
|---|---|---|---|---|---|---|---|---|---|---|---|---|
| | | | | | Meth | Opioid | Both | Neither | MethToOpioid | Hue | Saturation | Value |
| Neutral tool | 7.5 (20.58) | 4.5 (1.934) | 2.04 (1.69) | 6.96 (13.02) | 0 | 0 | 0 | 1 | 0 | 0.007 (0.057) | 0.03 (0.164) | 0.513 (0.082) |

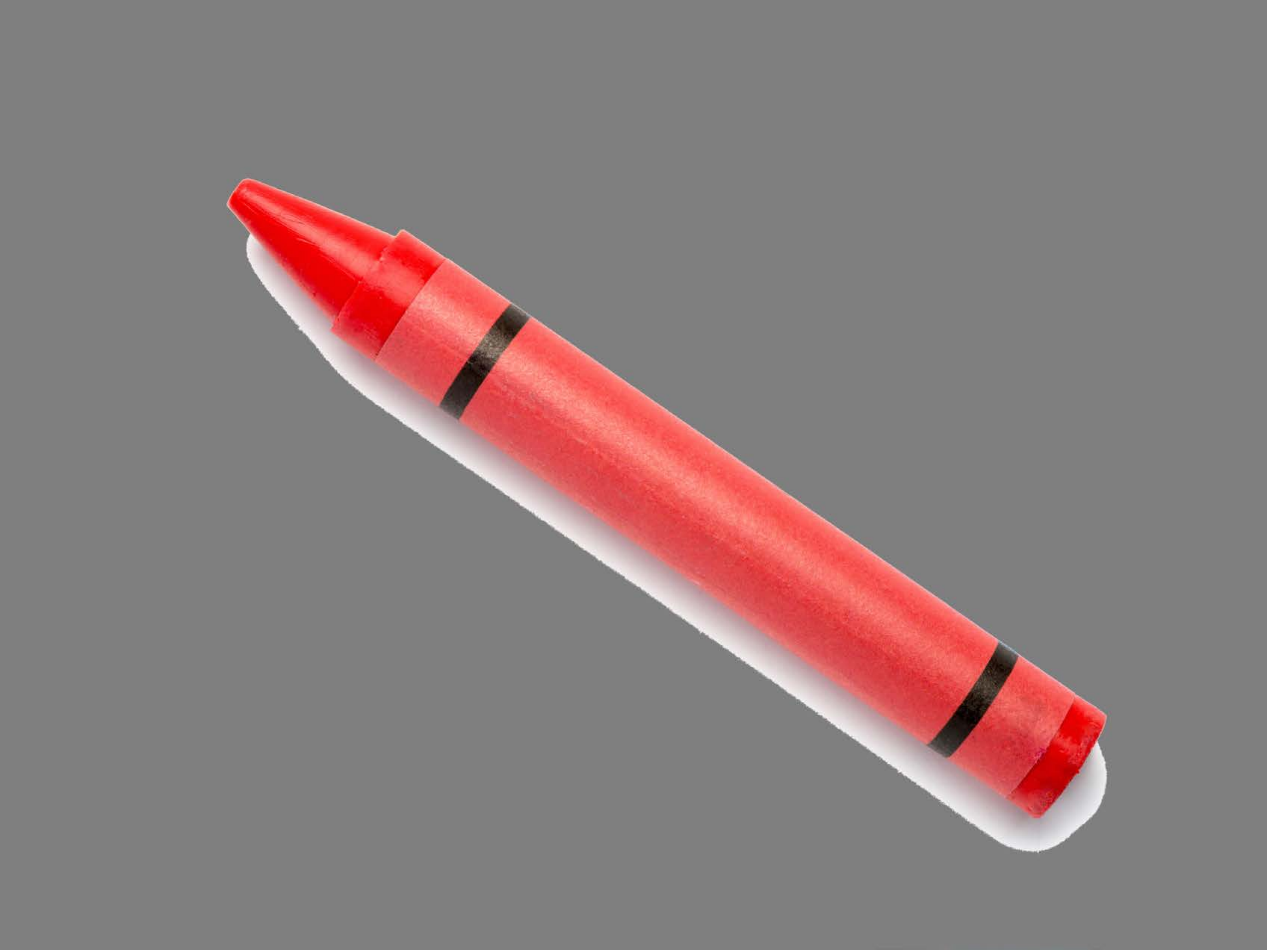

| Category | Craving (0-100) | Valence (1-9) | Arousal (1-9) | Typicality (0-100) | Relatedness | | | | | HSV | | |
|---|---|---|---|---|---|---|---|---|---|---|---|---|
| | | | | | Meth | Opioid | Both | Neither | MethToOpioid | Hue | Saturation | Value |
| Neutral tool | 10.07 (21.18) | 4.86 (1.779) | 2.46 (1.99) | 12.54 (21.92) | 0 | 0 | 0 | 1 | 0 | 0.069 (0.249) | 0.068 (0.201) | 0.548 (0.139) |

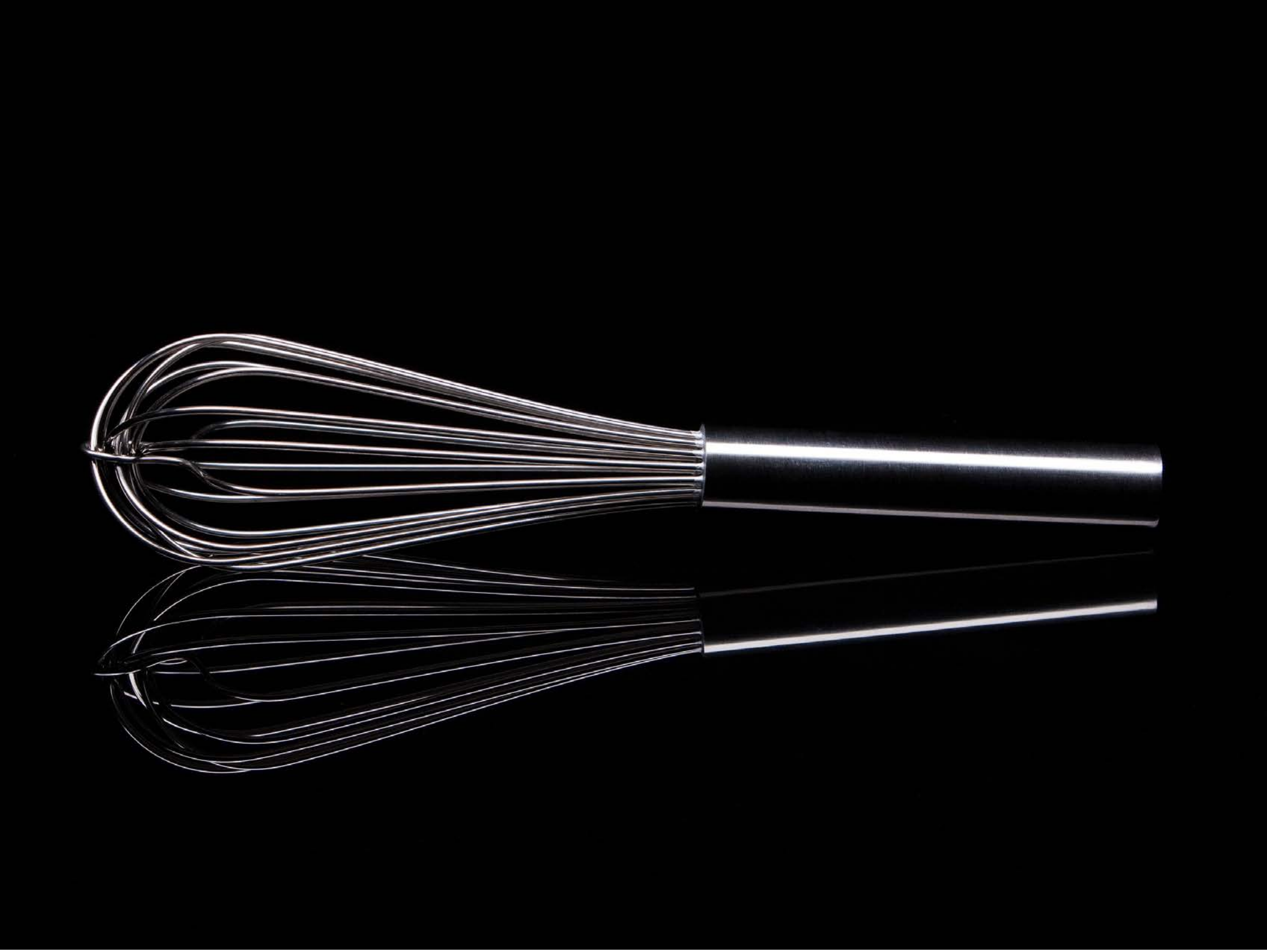

| Category | Craving (0-100) | Valence (1-9) | Arousal (1-9) | Typicality (0-100) | Relatedness | | | | | HSV | | |
|---|---|---|---|---|---|---|---|---|---|---|---|---|
| | | | | | Meth | Opioid | Both | Neither | MethToOpioid | Hue | Saturation | Value |
| Neutral tool | 7.57 (15.84) | 5 (1.466) | 2.21 (1.75) | 8.04 (15.49) | 0 | 0 | 0 | 1 | 0 | 0.099 (0.26) | 0.064 (0.198) | 0.04 (0.151) |

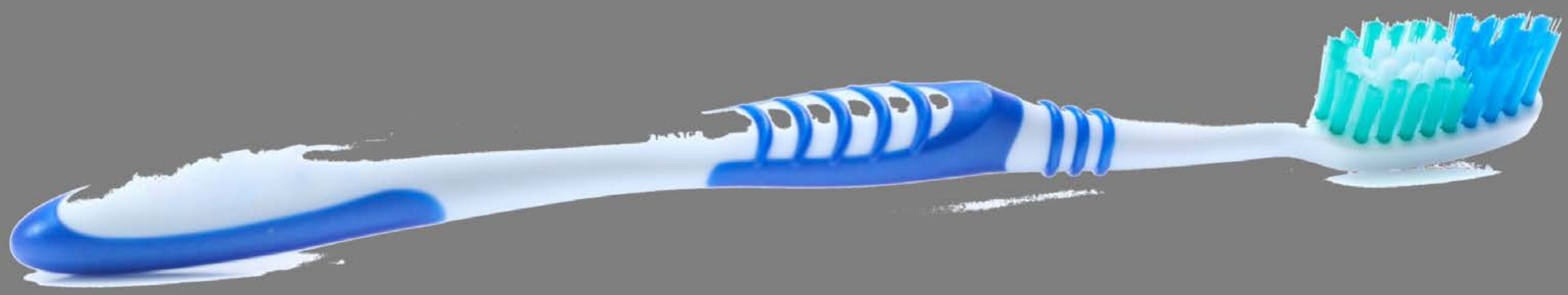

| Category | Craving (0-100) | Valence (1-9) | Arousal (1-9) | Typicality (0-100) | Relatedness | | | | | HSV | | |
|---|---|---|---|---|---|---|---|---|---|---|---|---|
| | | | | | Meth | Opioid | Both | Neither | MethToOpioid | Hue | Saturation | Value |
| Neutral tool | 13.18 (24.22) | 5.21 (2.217) | 2.43 (1.91) | 10.61 (17.52) | 0 | 0 | 0.03571 | 0.96429 | 0 | 0.038 (0.145) | 0.026 (0.132) | 0.522 (0.097) |

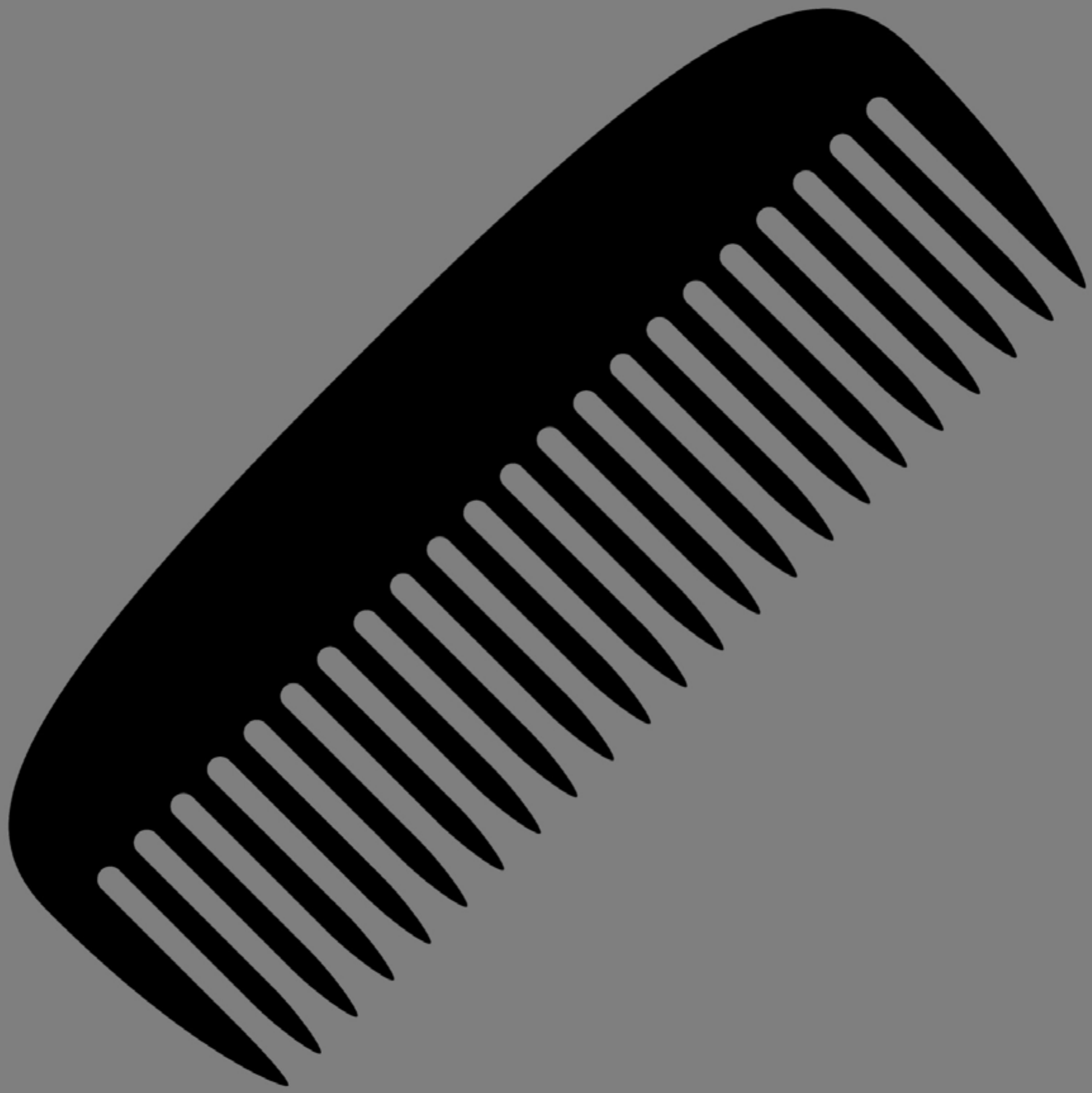

| Category | Craving (0-100) | Valence (1-9) | Arousal (1-9) | Typicality (0-100) | Relatedness | | | | | HSV | | |
|---|---|---|---|---|---|---|---|---|---|---|---|---|
| | | | | | Meth | Opioid | Both | Neither | MethToOpioid | Hue | Saturation | Value |
| Neutral tool | 7.33 (15.77) | 4.89 (0.847) | 2.37 (1.9) | 12.19 (18.84) | 0 | 0 | 0 | 1 | 0 | 0 (0) | 0 (0) | 0.446 (0.151) |

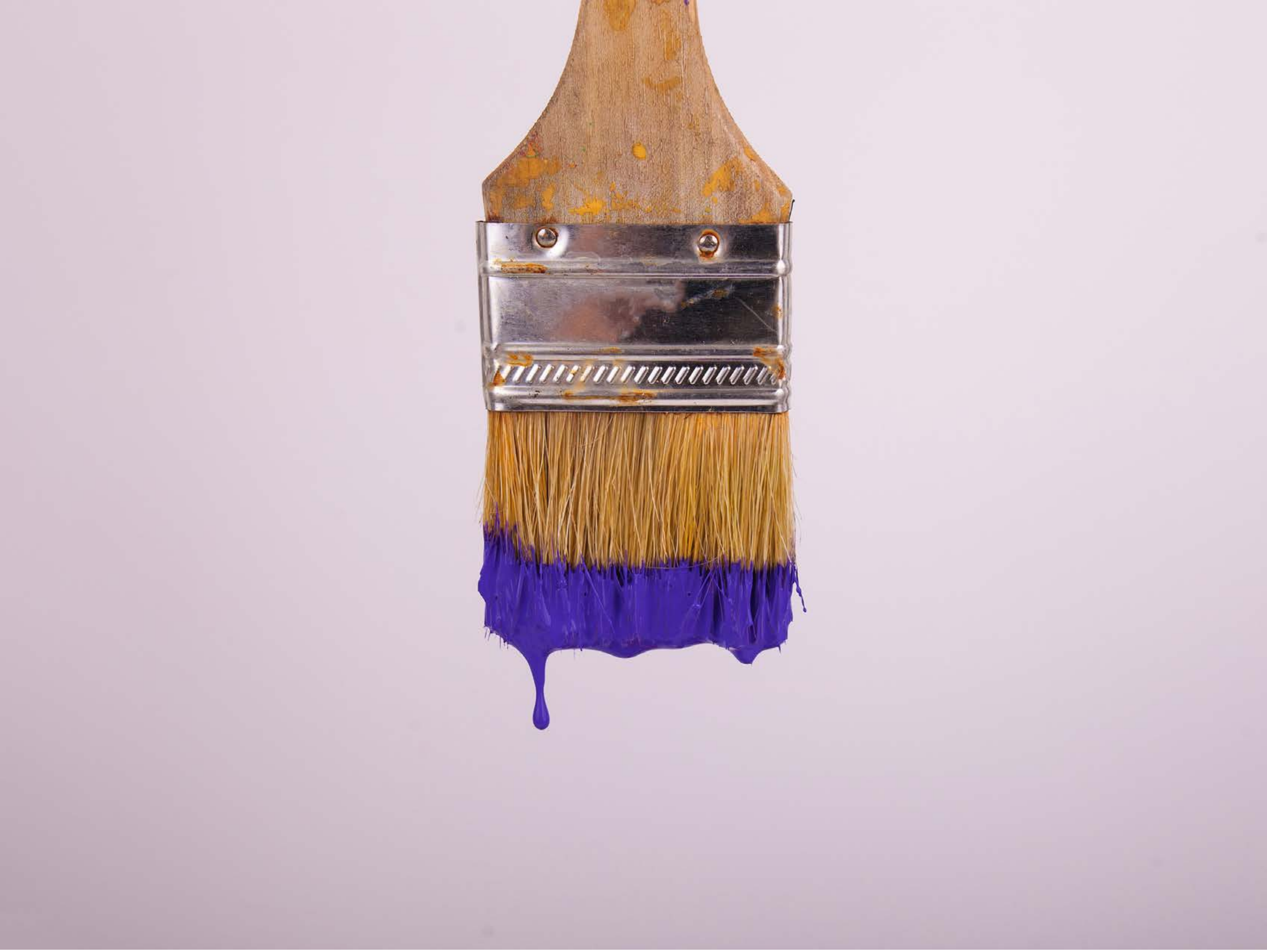

| Category | Craving (0-100) | Valence (1-9) | Arousal (1-9) | Typicality (0-100) | Relatedness | | | | | HSV | | |
|---|---|---|---|---|---|---|---|---|---|---|---|---|
| | | | | | Meth | Opioid | Both | Neither | MethToOpioid | Hue | Saturation | Value |
| Neutral tool | 5.89 (13.07) | 5.07 (1.562) | 2.39 (1.81) | 7.32 (15.45) | 0 | 0 | 0 | 1 | 0 | 0.789 (0.241) | 0.124 (0.174) | 0.821 (0.118) |

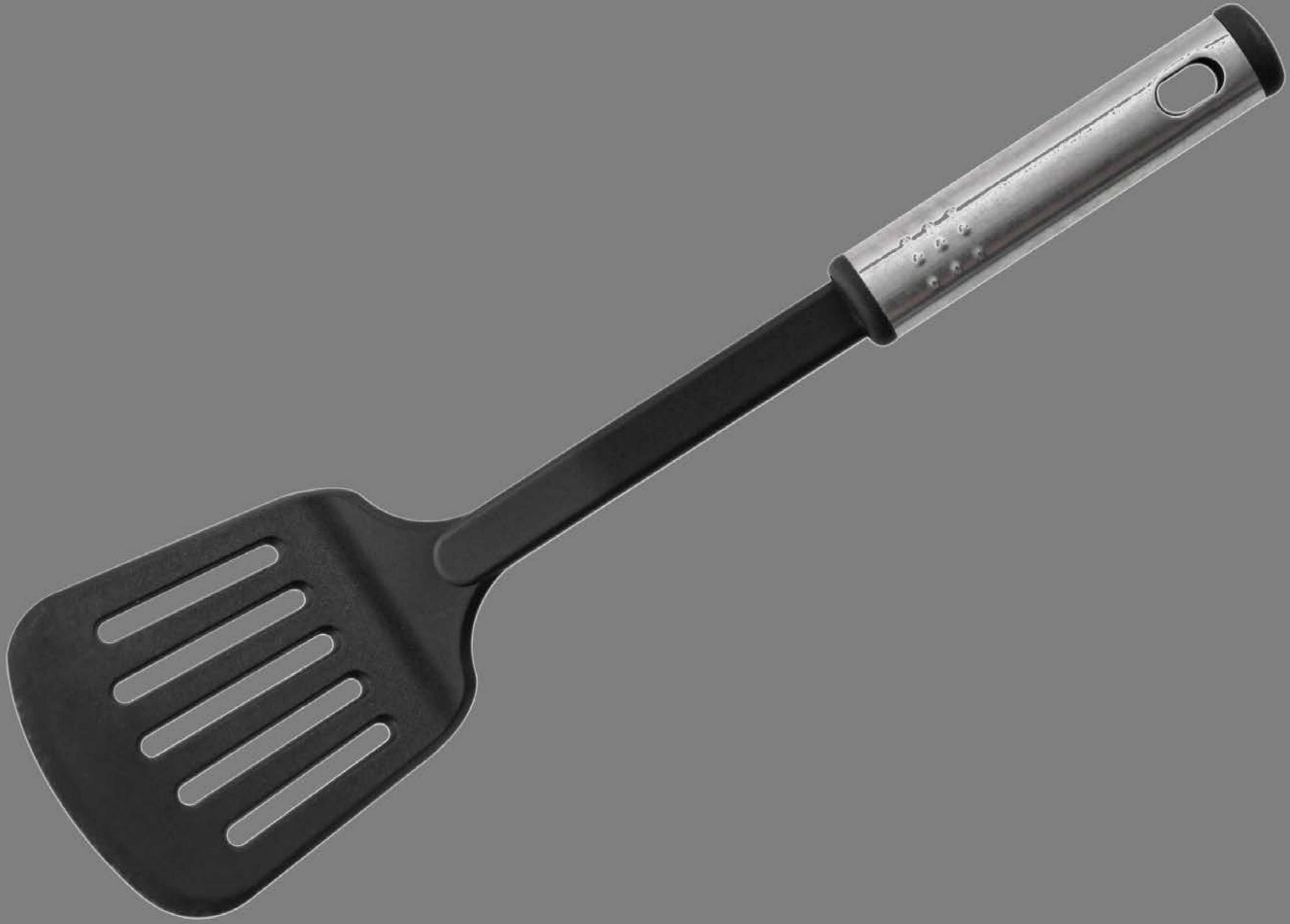

| Category | Craving (0-100) | Valence (1-9) | Arousal (1-9) | Typicality (0-100) | Relatedness | | | | | HSV | | |
|---|---|---|---|---|---|---|---|---|---|---|---|---|
| | | | | | Meth | Opioid | Both | Neither | MethToOpioid | Hue | Saturation | Value |
| Neutral tool | 8.85 (17.75) | 4.89 (1.311) | 2.63 (2.17) | 10.67 (17.47) | 0 | 0 | 0 | 1 | 0 | 0.008 (0.078) | 0 (0.005) | 0.478 (0.086) |

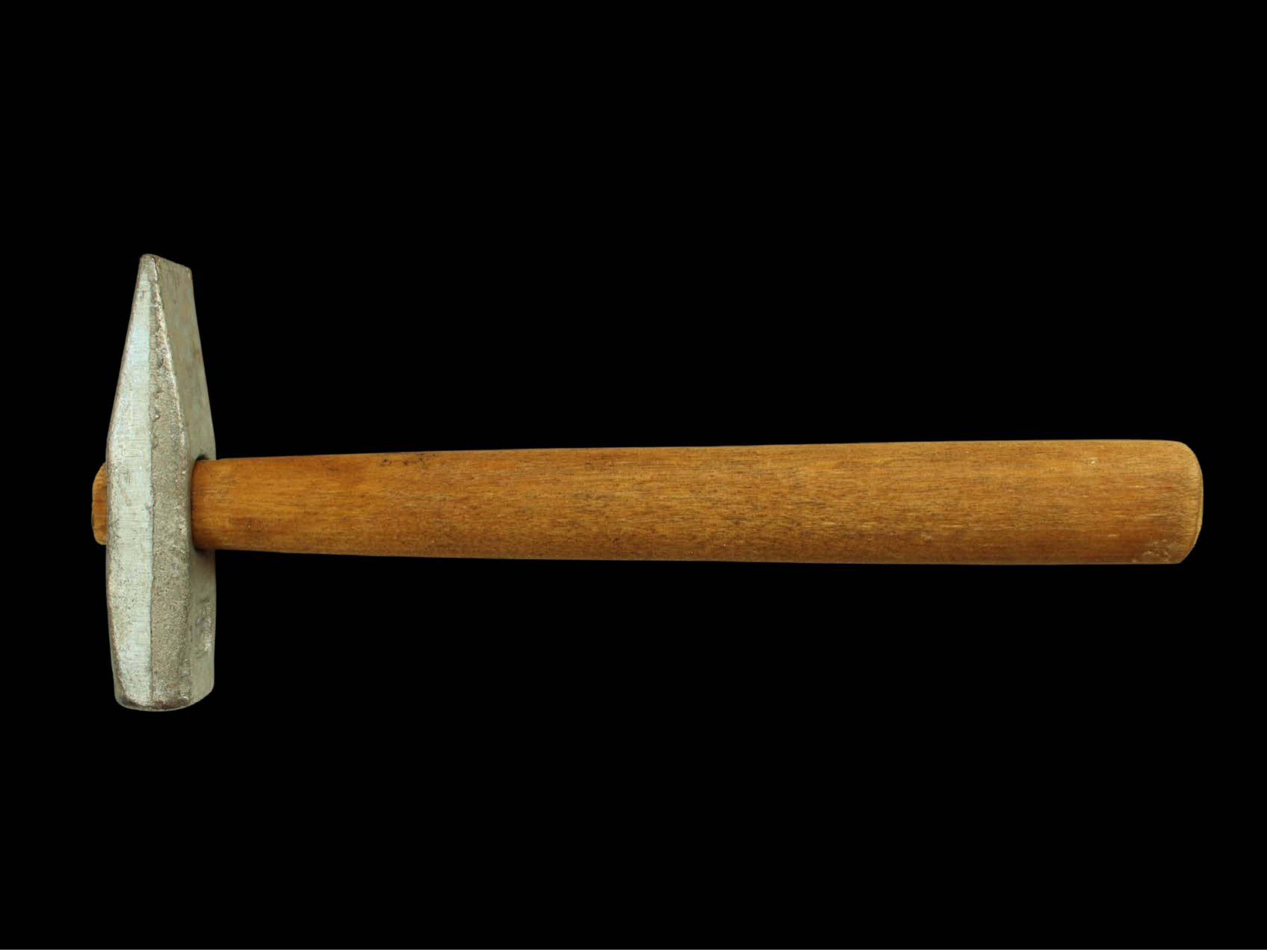

| Category | Craving (0-100) | Valence (1-9) | Arousal (1-9) | Typicality (0-100) | Relatedness | | | | | HSV | | |
|---|---|---|---|---|---|---|---|---|---|---|---|---|
| | | | | | Meth | Opioid | Both | Neither | MethToOpioid | Hue | Saturation | Value |
| Neutral tool | 12 (24.17) | 4.79 (1.893) | 2.29 (1.86) | 12.79 (24.31) | 0 | 0 | 0 | 1 | 0 | 0.017 (0.053) | 0.086 (0.238) | 0.079 (0.209) |

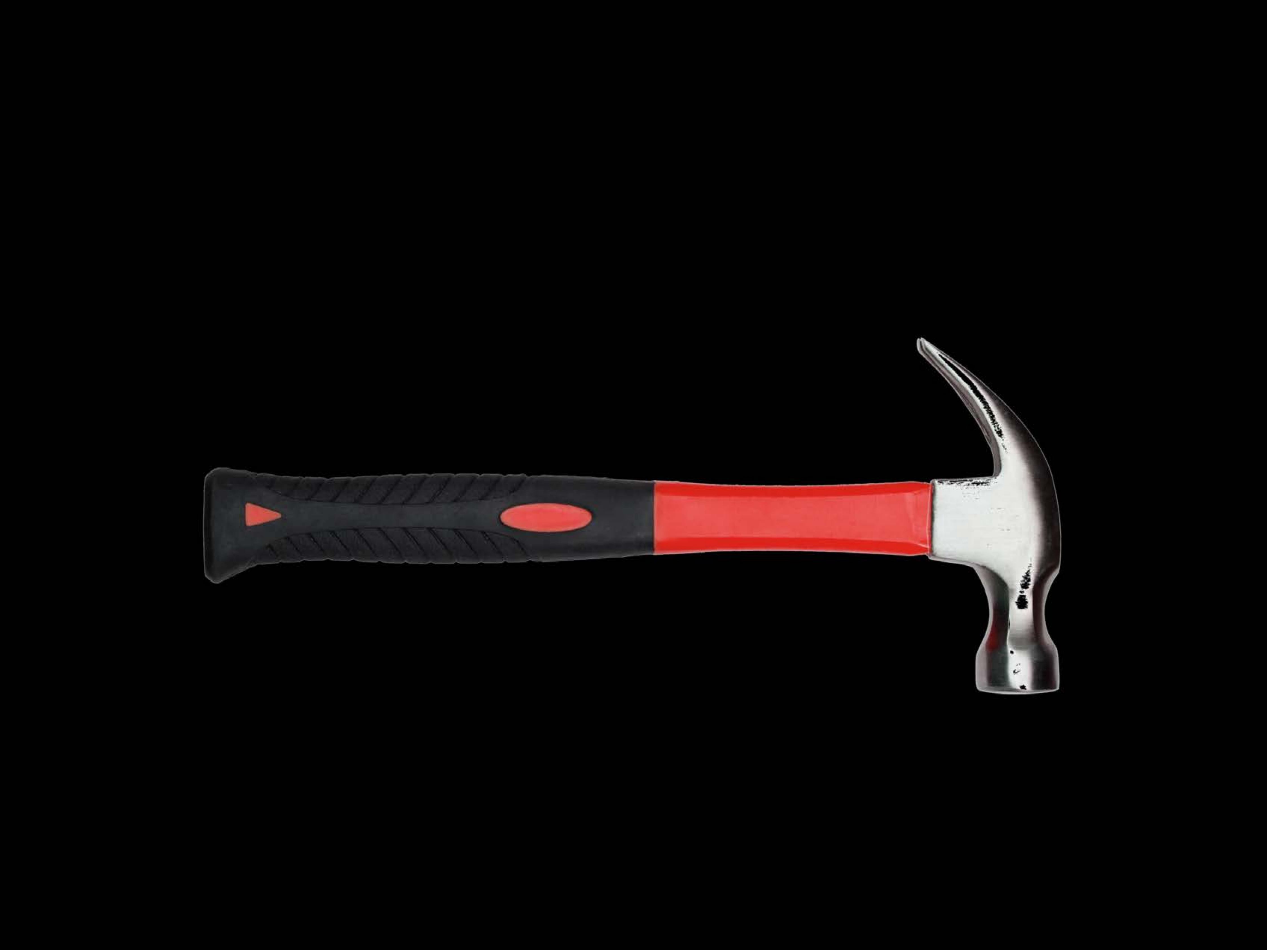

| Category | Craving (0-100) | Valence (1-9) | Arousal (1-9) | Typicality (0-100) | Relatedness | | | | | HSV | | |
|---|---|---|---|---|---|---|---|---|---|---|---|---|
| | | | | | Meth | Opioid | Both | Neither | MethToOpioid | Hue | Saturation | Value |
| Neutral tool | 9.71 (19.17) | 4.32 (1.786) | 2.54 (1.91) | 9.25 (17.23) | 0.03571 | 0 | 0 | 0.96429 | -0.03571 | 0.028 (0.134) | 0.018 (0.105) | 0.033 (0.151) |

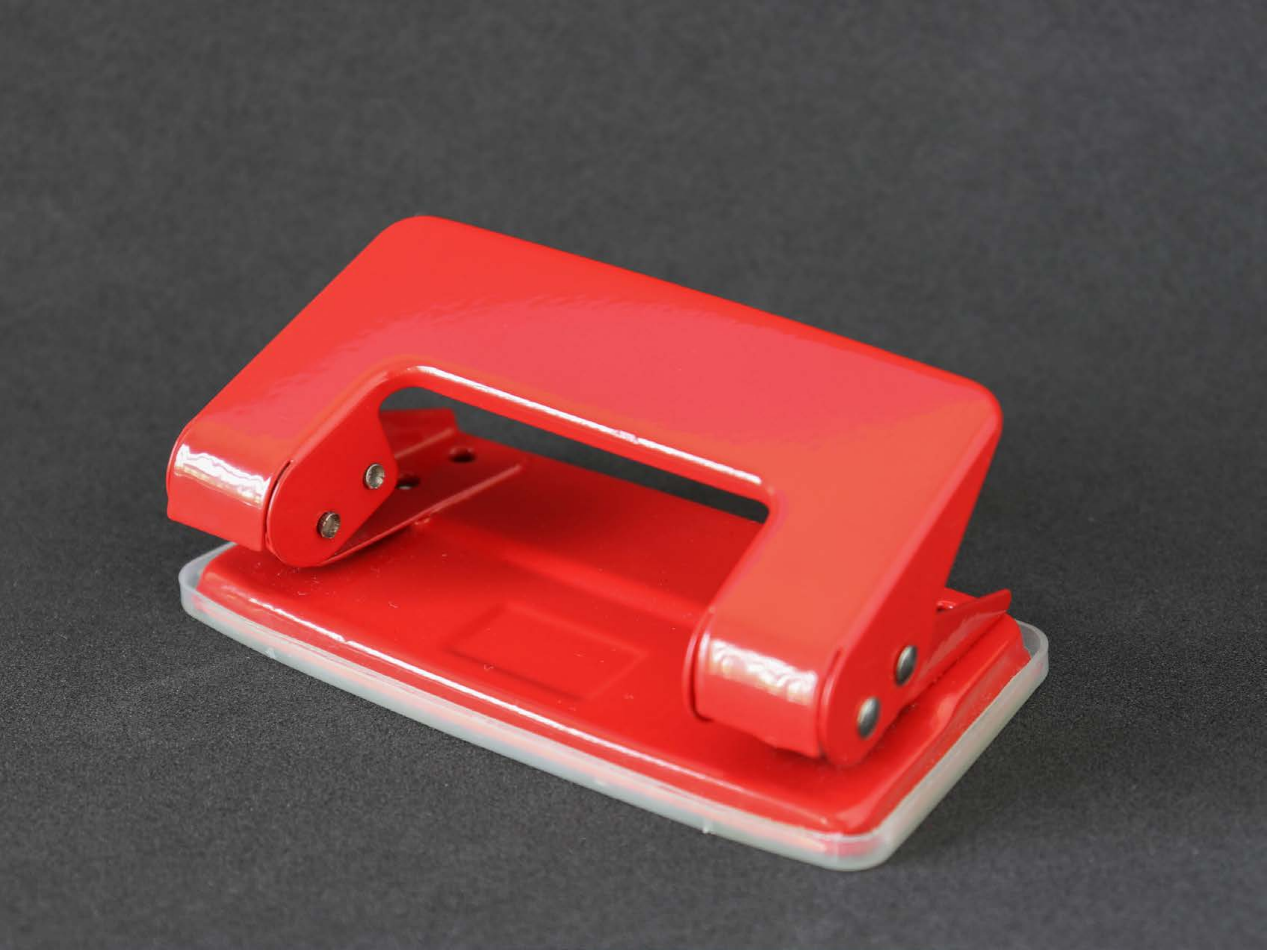

| Category | Craving (0-100) | Valence (1-9) | Arousal (1-9) | Typicality (0-100) | Relatedness | | | | | HSV | | |
|---|---|---|---|---|---|---|---|---|---|---|---|---|
| | | | | | Meth | Opioid | Both | Neither | MethToOpioid | Hue | Saturation | Value |
| Neutral tool | 10 (18) | 4.93 (1.412) | 2.26 (1.83) | 9.59 (17.77) | 0 | 0 | 0 | 1 | 0 | 0.23 (0.332) | 0.225 (0.338) | 0.463 (0.278) |

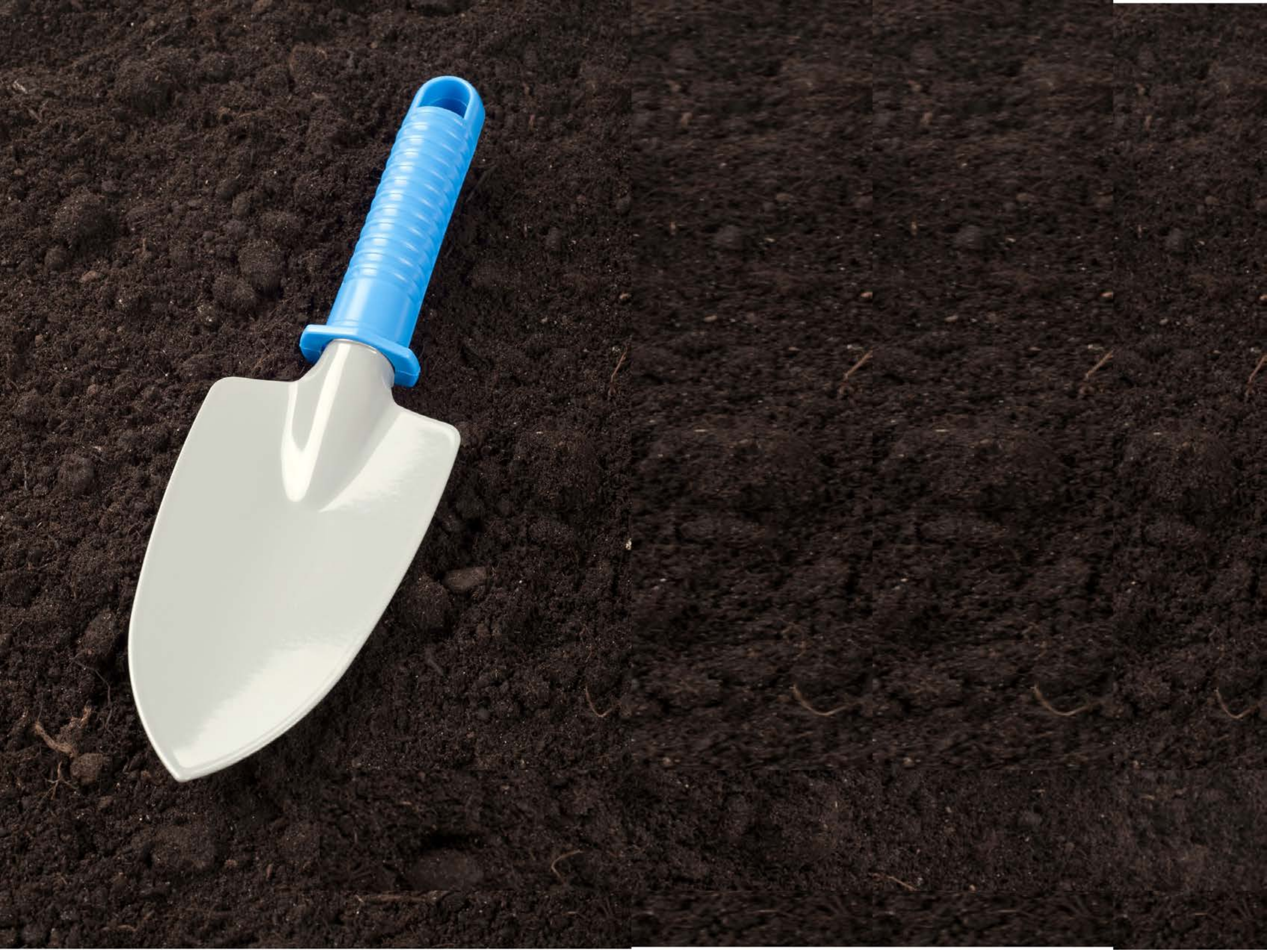

| Category | Craving (0-100) | Valence (1-9) | Arousal (1-9) | Typicality (0-100) | Relatedness | | | | | HSV | | |
|---|---|---|---|---|---|---|---|---|---|---|---|---|
| | | | | | Meth | Opioid | Both | Neither | MethToOpioid | Hue | Saturation | Value |
| Neutral tool | 6.33 (16.03) | 4.85 (1.35) | 2.26 (1.83) | 7.48 (15.7) | 0 | 0 | 0 | 1 | 0 | 0.087 (0.162) | 0.331 (0.152) | 0.222 (0.223) |

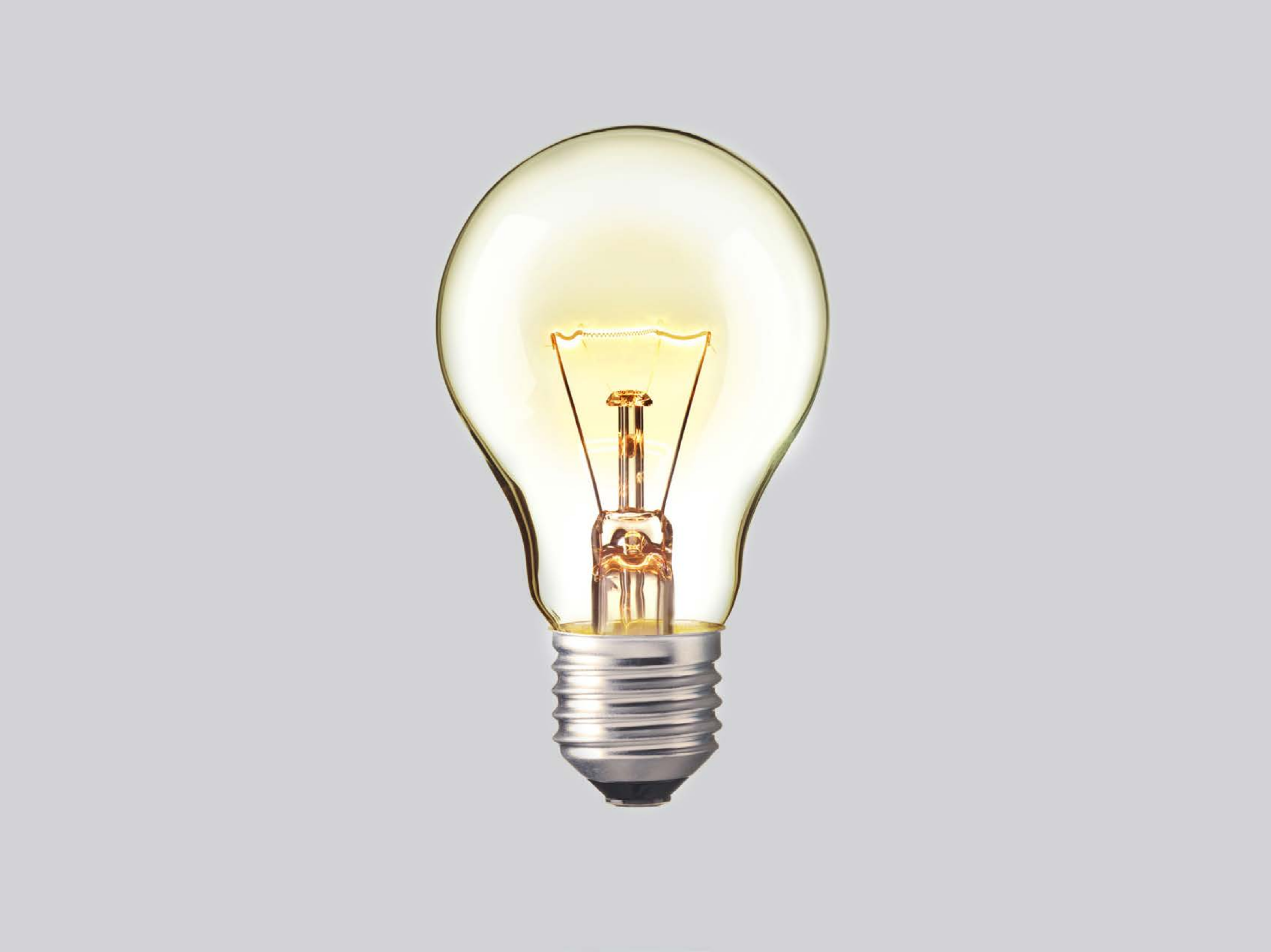

| Category | Craving (0-100) | Valence (1-9) | Arousal (1-9) | Typicality (0-100) | Relatedness | | | | | HSV | | |
|---|---|---|---|---|---|---|---|---|---|---|---|---|
| | | | | | Meth | Opioid | Both | Neither | MethToOpioid | Hue | Saturation | Value |
| Neutral tool | 51.18 (31.93) | 5.07 (1.184) | 3.75 (2.43) | 55.43 (30.92) | 0.53571 | 0 | 0.07143 | 0.39286 | -0.53571 | 0.028 (0.098) | 0.02 (0.074) | 0.984 (0.078) |

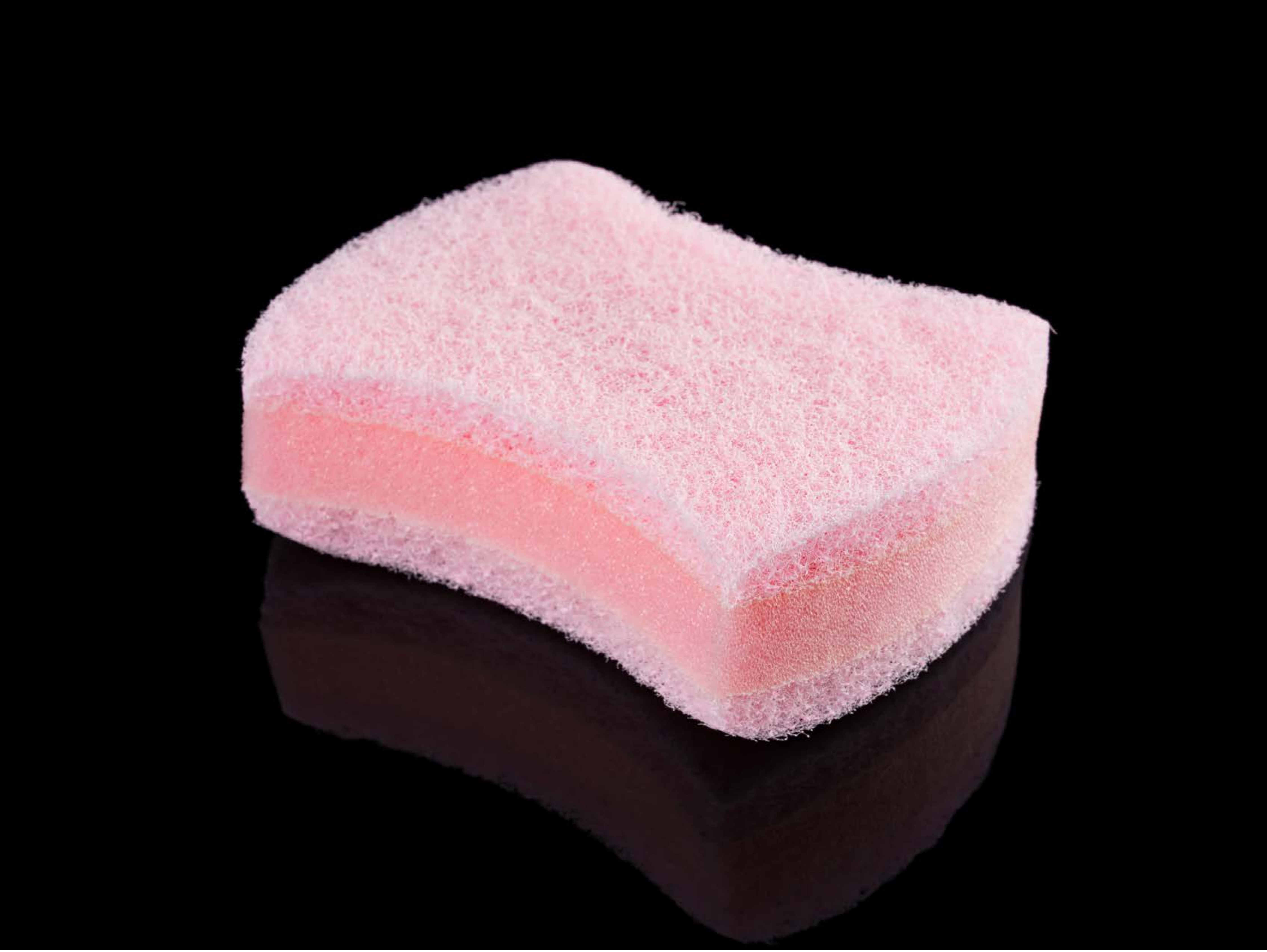

| Category | Craving (0-100) | Valence (1-9) | Arousal (1-9) | Typicality (0-100) | Relatedness | | | | | HSV | | |
|---|---|---|---|---|---|---|---|---|---|---|---|---|
| | | | | | Meth | Opioid | Both | Neither | MethToOpioid | Hue | Saturation | Value |
| Neutral tool | 7.39 (15.92) | 4.86 (1.649) | 2.21 (1.89) | 14.18 (21.12) | 0 | 0 | 0 | 1 | 0 | 0.363 (0.463) | 0.108 (0.163) | 0.254 (0.393) |

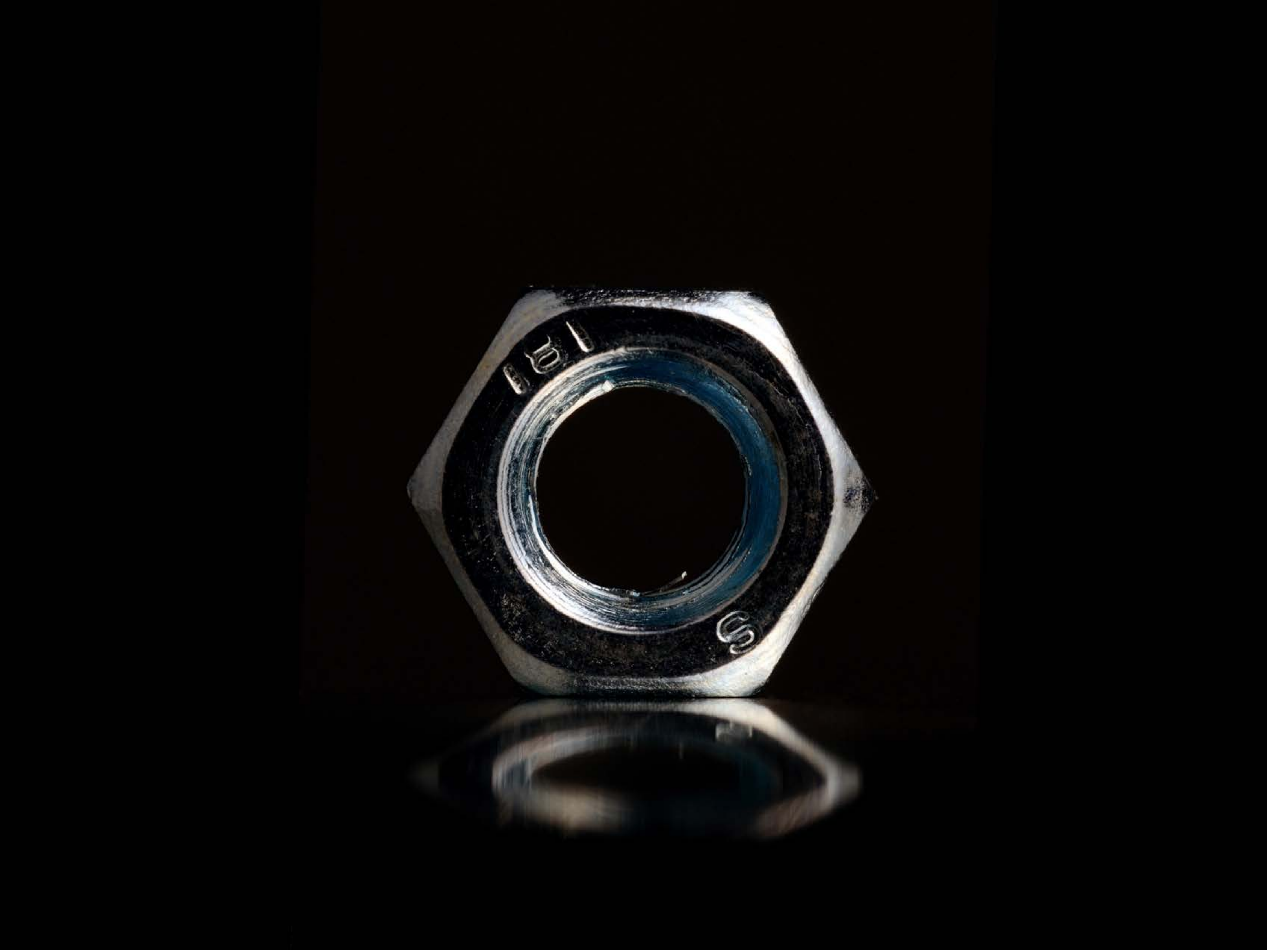

| Category | Craving (0-100) | Valence (1-9) | Arousal (1-9) | Typicality (0-100) | Relatedness | | | | | HSV | | |
|---|---|---|---|---|---|---|---|---|---|---|---|---|
| | | | | | Meth | Opioid | Both | Neither | MethToOpioid | Hue | Saturation | Value |
| Neutral tool | 9.37 (17.64) | 4.7 (1.068) | 2.41 (1.89) | 7.81 (15.72) | 0 | 0 | 0 | 1 | 0 | 0.096 (0.223) | 0.261 (0.351) | 0.04 (0.125) |

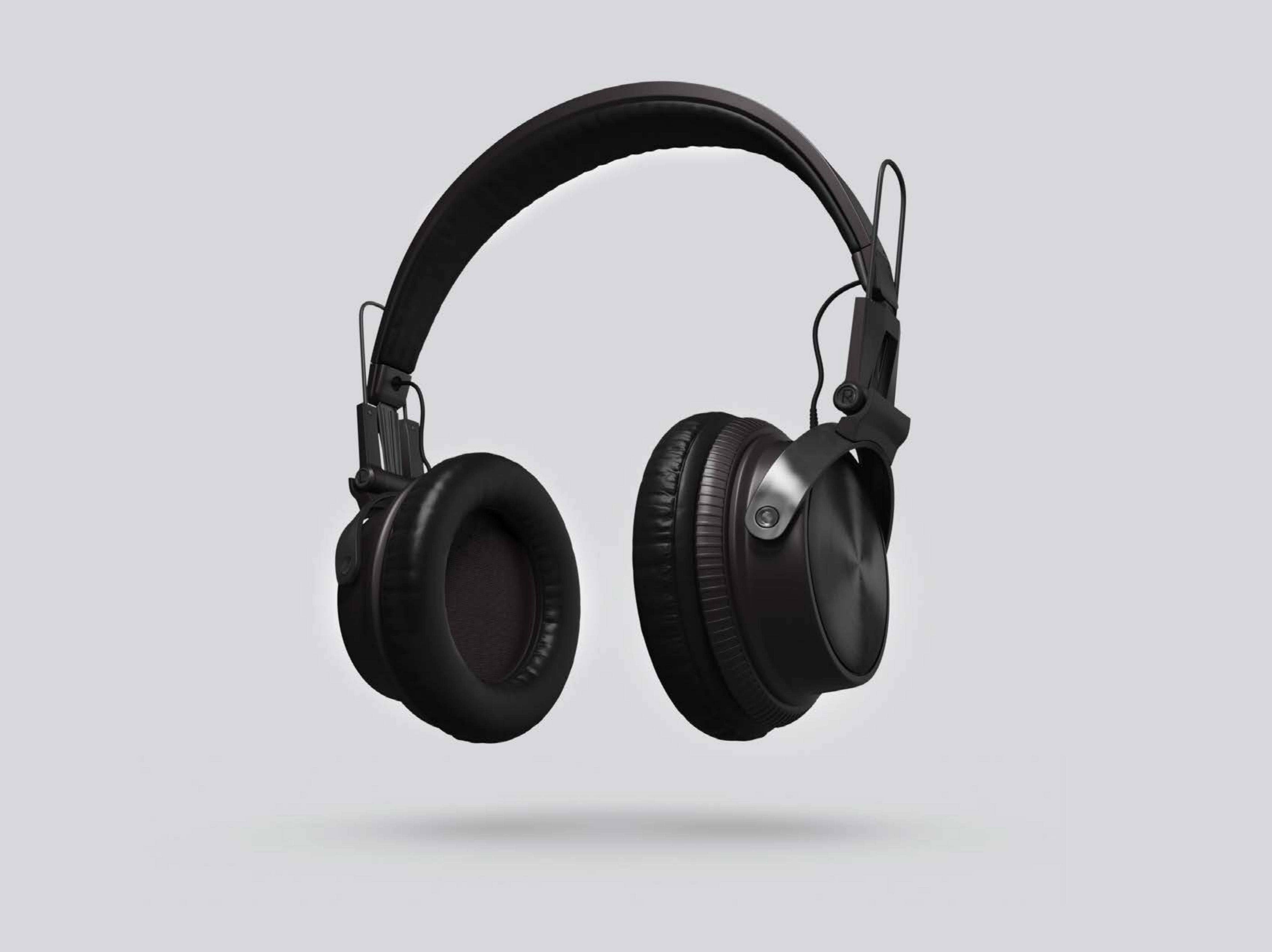

| Category | Craving (0-100) | Valence (1-9) | Arousal (1-9) | Typicality (0-100) | Relatedness | | | | | HSV | | |
|---|---|---|---|---|---|---|---|---|---|---|---|---|
| | | | | | Meth | Opioid | Both | Neither | MethToOpioid | Hue | Saturation | Value |
| Neutral tool | 7.14 (14.85) | 5.07 (1.864) | 2.29 (1.7) | 13.36 (18.86) | 0 | 0 | 0.03571 | 0.96429 | 0 | 0.039 (0.186) | 0.003 (0.017) | 0.871 (0.303) |

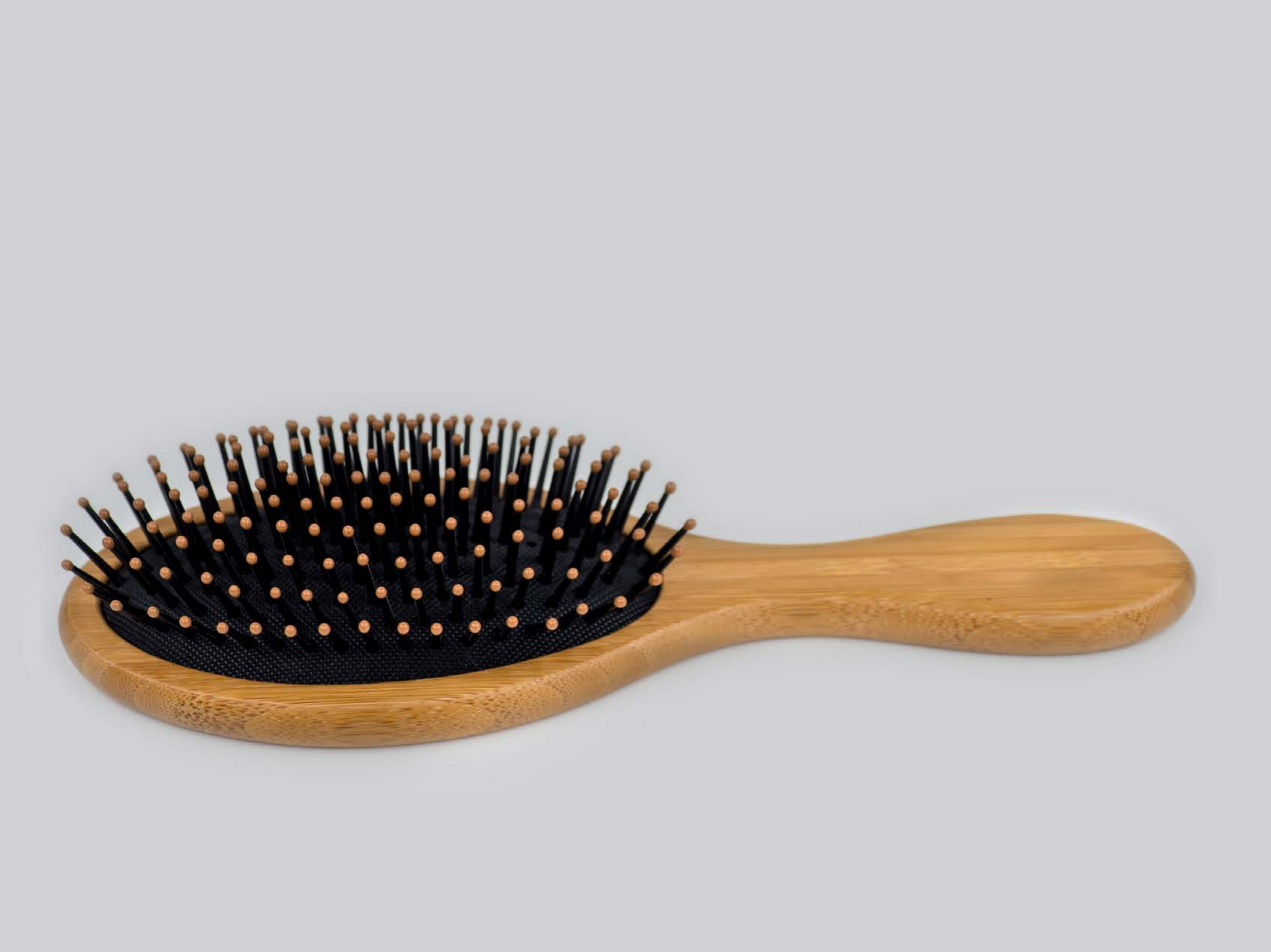

| Category | Craving (0-100) | Valence (1-9) | Arousal (1-9) | Typicality (0-100) | Relatedness | | | | | HSV | | |
|---|---|---|---|---|---|---|---|---|---|---|---|---|
| | | | | | Meth | Opioid | Both | Neither | MethToOpioid | Hue | Saturation | Value |
| Neutral tool | 10.81 (19.32) | 4.89 (1.311) | 2.96 (2.01) | 16.22 (25.19) | 0 | 0 | 0 | 1 | 0 | 0.065 (0.184) | 0.087 (0.212) | 0.898 (0.257) |

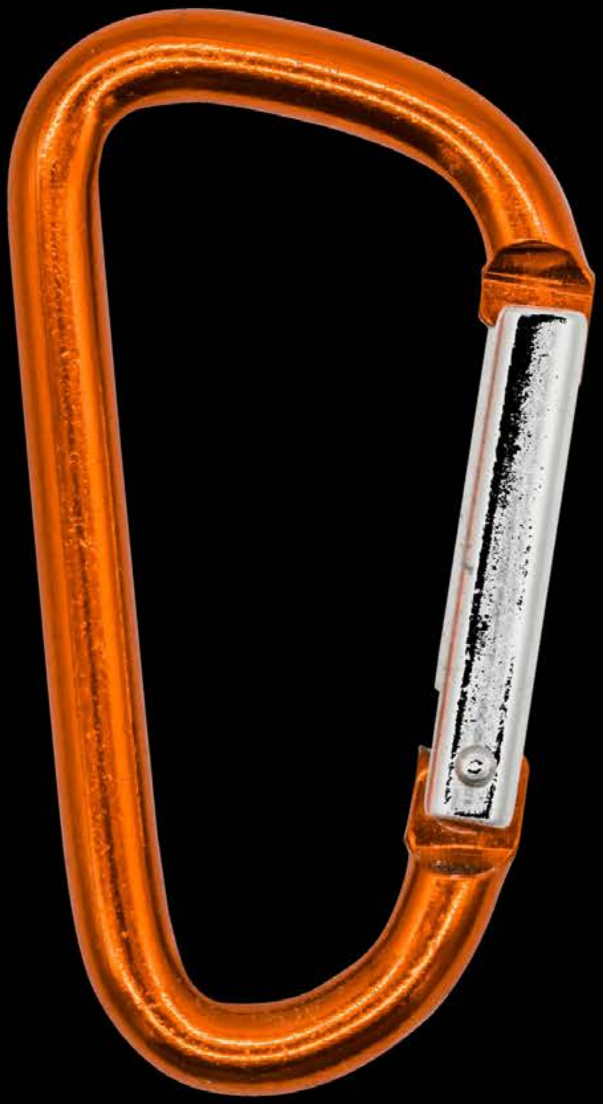

| Category | Craving (0-100) | Valence (1-9) | Arousal (1-9) | Typicality (0-100) | Relatedness | | | | | HSV | | |
|---|---|---|---|---|---|---|---|---|---|---|---|---|
| | | | | | Meth | Opioid | Both | Neither | MethToOpioid | Hue | Saturation | Value |
| Neutral tool | 10.18 (19.02) | 5.04 (1.453) | 2.64 (1.95) | 11.43 (19.05) | 0 | 0 | 0 | 1 | 0 | 0.007 (0.049) | 0.053 (0.215) | 0.046 (0.186) |

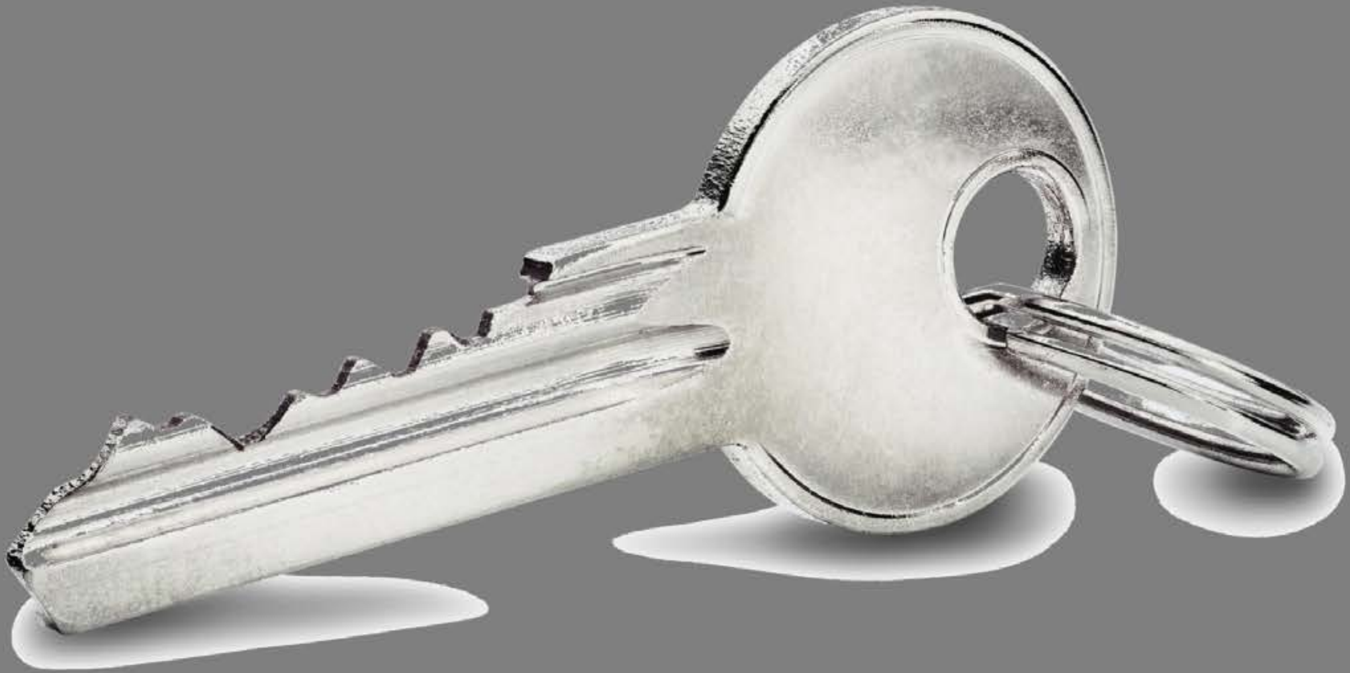

| Category | Craving (0-100) | Valence (1-9) | Arousal (1-9) | Typicality (0-100) | Relatedness | | | | | HSV | | |
|---|---|---|---|---|---|---|---|---|---|---|---|---|
| | | | | | Meth | Opioid | Both | Neither | MethToOpioid | Hue | Saturation | Value |
| Neutral tool | 11.33 (19.52) | 4.81 (0.786) | 2.7 (1.96) | 18.67 (23.01) | 0 | 0 | 0 | 1 | 0 | 0.014 (0.069) | 0.003 (0.012) | 0.525 (0.097) |

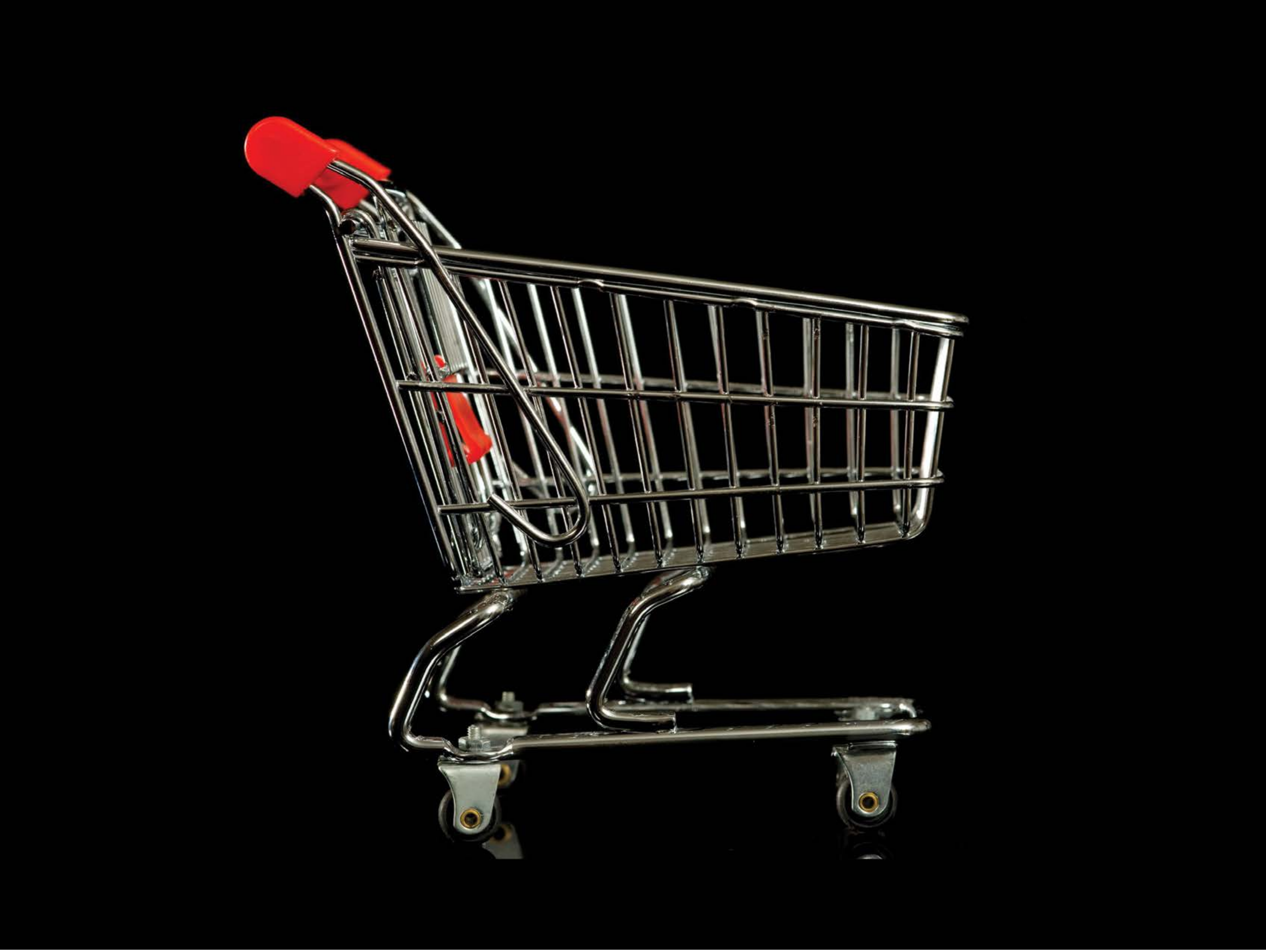

| Category | Craving (0-100) | Valence (1-9) | Arousal (1-9) | Typicality (0-100) | Relatedness | | | | | HSV | | |
|---|---|---|---|---|---|---|---|---|---|---|---|---|
| | | | | | Meth | Opioid | Both | Neither | MethToOpioid | Hue | Saturation | Value |
| Neutral tool | 13.48 (24.77) | 4.7 (1.068) | 2.7 (1.96) | 14.26 (24.65) | 0 | 0 | 0 | 1 | 0 | 0.036 (0.113) | 0.072 (0.216) | 0.062 (0.182) |

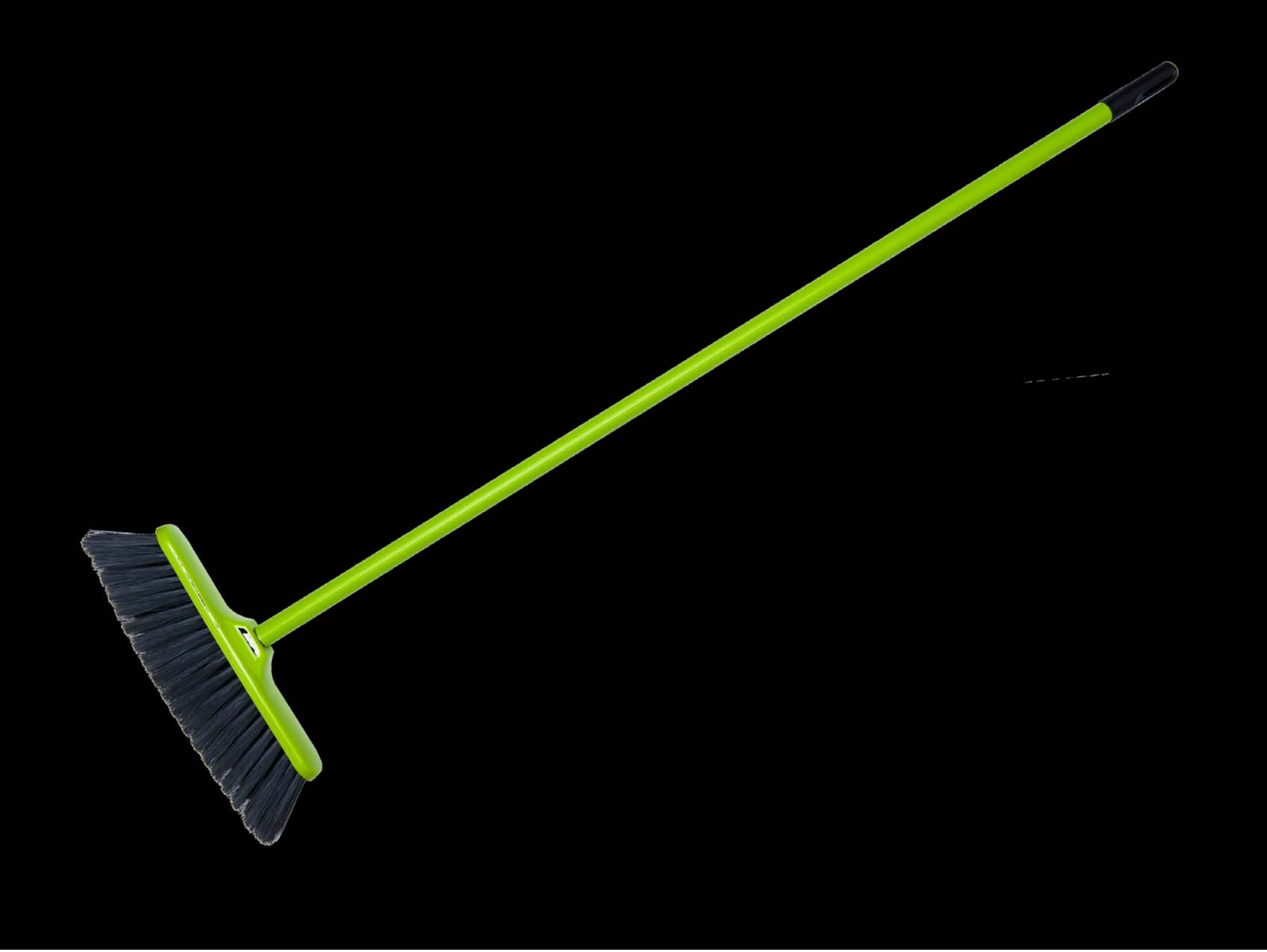

| Category | Craving (0-100) | Valence (1-9) | Arousal (1-9) | Typicality (0-100) | Relatedness | | | | | HSV | | |
|---|---|---|---|---|---|---|---|---|---|---|---|---|
| | | | | | Meth | Opioid | Both | Neither | MethToOpioid | Hue | Saturation | Value |
| Neutral tool | 7.26 (15.64) | 4.74 (1.095) | 2.26 (1.83) | 10.3 (17.62) | 0 | 0 | 0 | 1 | 0 | 0.022 (0.102) | 0.034 (0.163) | 0.025 (0.126) |

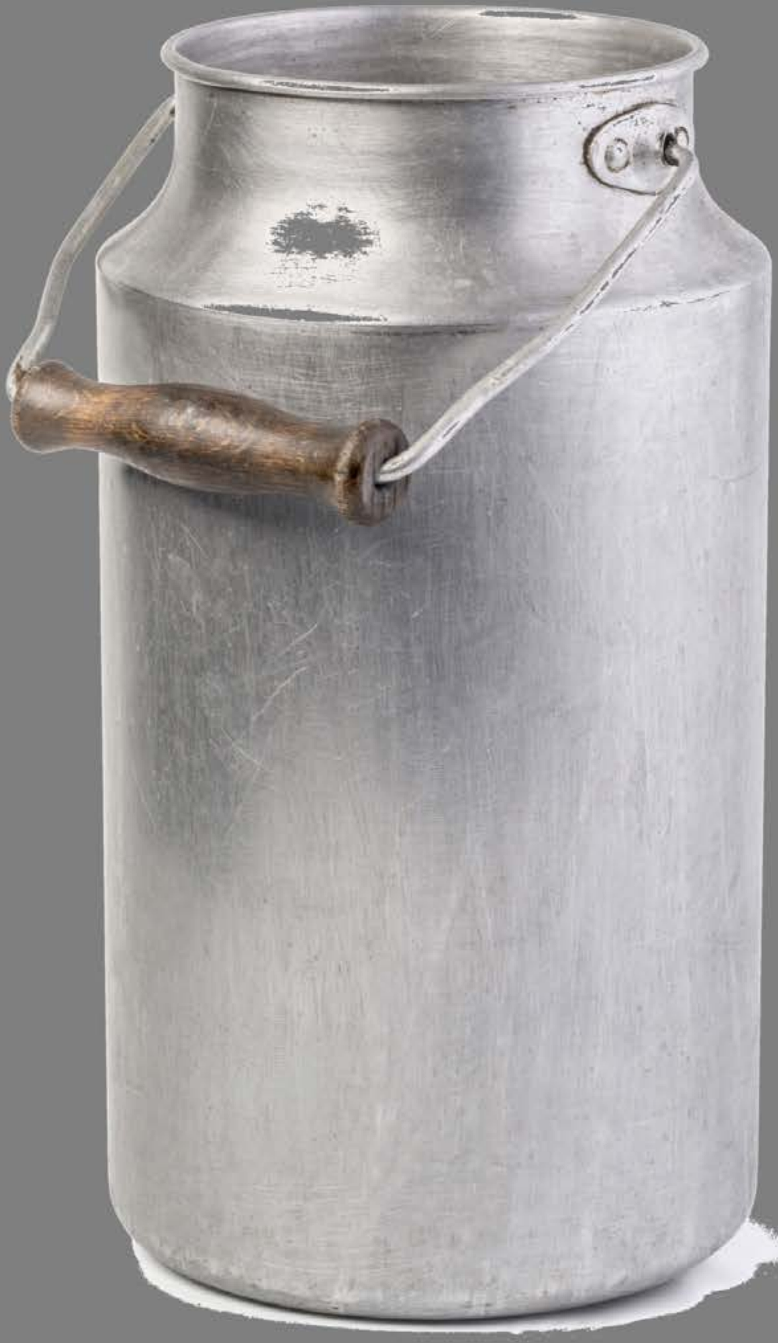

| Category | Craving (0-100) | Valence (1-9) | Arousal (1-9) | Typicality (0-100) | Relatedness | | | | | HSV | | |
|---|---|---|---|---|---|---|---|---|---|---|---|---|
| | | | | | Meth | Opioid | Both | Neither | MethToOpioid | Hue | Saturation | Value |
| Neutral tool | 5.57 (13.07) | 4.79 (1.572) | 2.14 (1.72) | 6.46 (13.74) | 0 | 0 | 0 | 1 | 0 | 0.1 (0.247) | 0.009 (0.029) | 0.542 (0.103) |

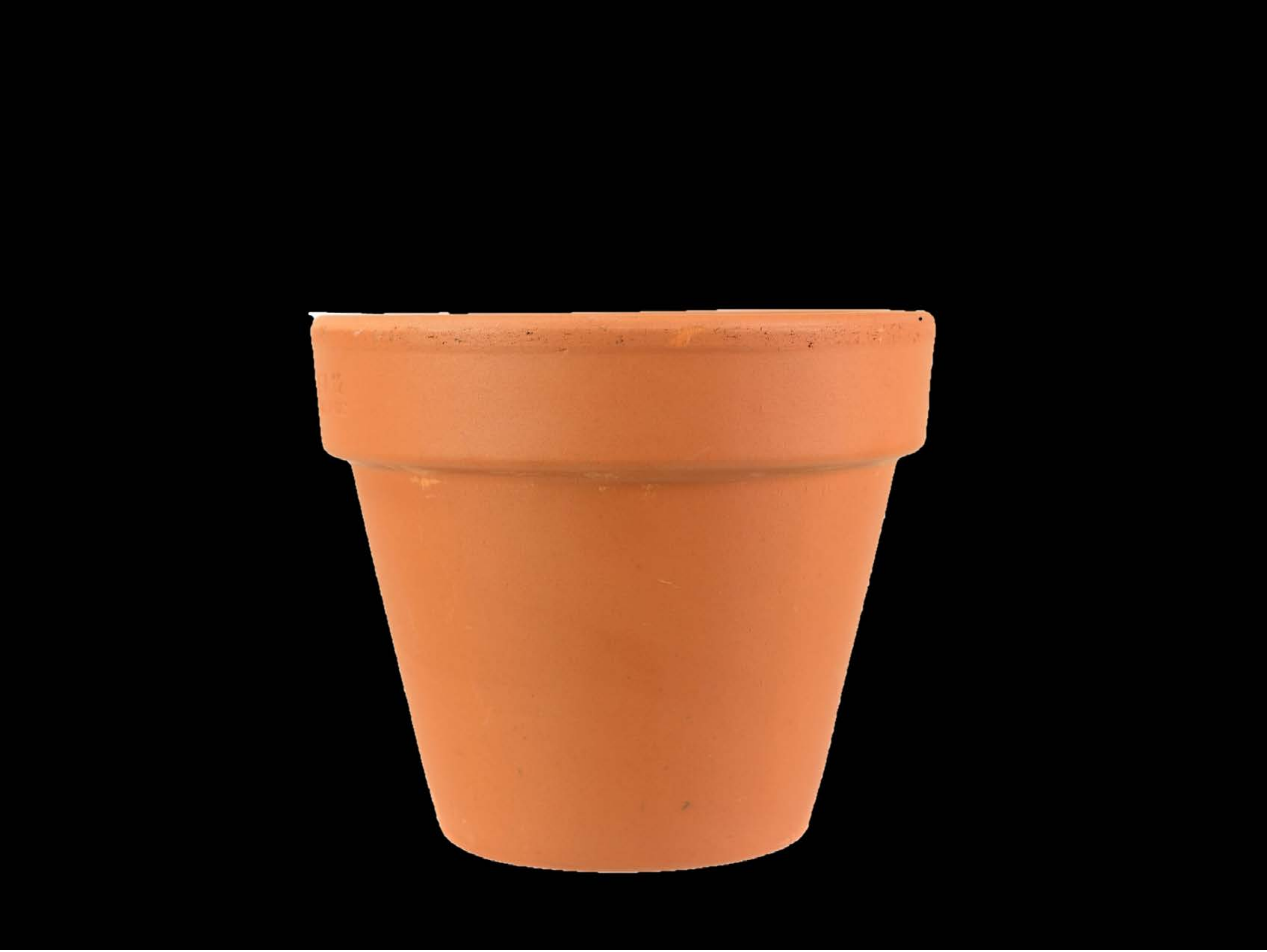

| Category | Craving (0-100) | Valence (1-9) | Arousal (1-9) | Typicality (0-100) | Relatedness | | | | | HSV | | |
|---|---|---|---|---|---|---|---|---|---|---|---|---|
| | | | | | Meth | Opioid | Both | Neither | MethToOpioid | Hue | Saturation | Value |
| Neutral tool | 6.96 (15.73) | 4.7 (1.068) | 2.52 (1.95) | 7.15 (15.77) | 0 | 0 | 0 | 1 | 0 | 0.015 (0.039) | 0.153 (0.281) | 0.192 (0.355 |

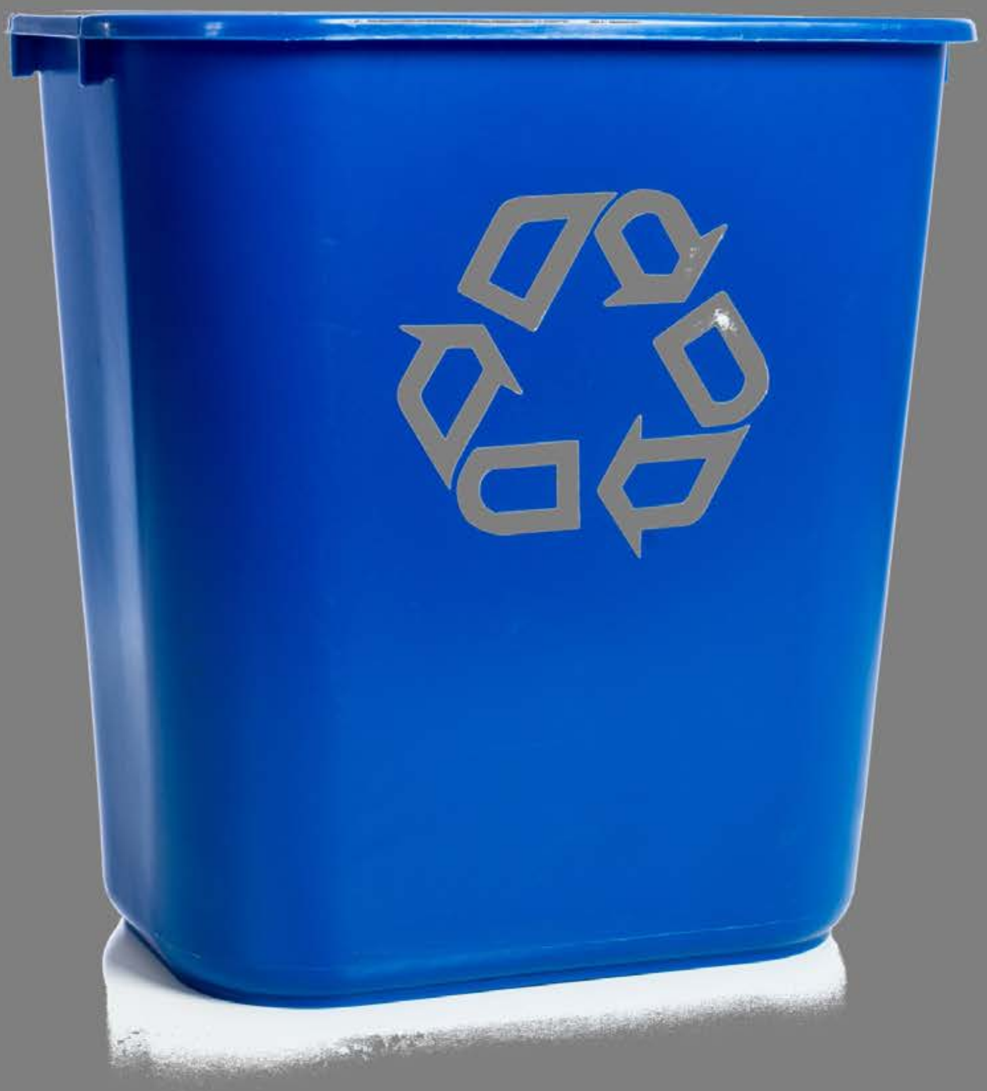

| Category | Craving (0-100) | Valence (1-9) | Arousal (1-9) | Typicality (0-100) | Relatedness | | | | | HSV | | |
|---|---|---|---|---|---|---|---|---|---|---|---|---|
| | | | | | Meth | Opioid | Both | Neither | MethToOpioid | Hue | Saturation | Value |
| Neutral tool | 11.68( 20.86) | 5 (1.515) | 2.79 (1.97) | 13.32 (19.68) | 0.03571 | 0 | 0 | 0.96429 | -0.03571 | 0.133 (0.249) | 0.19 (0.384) | 0.54 (0.097) |

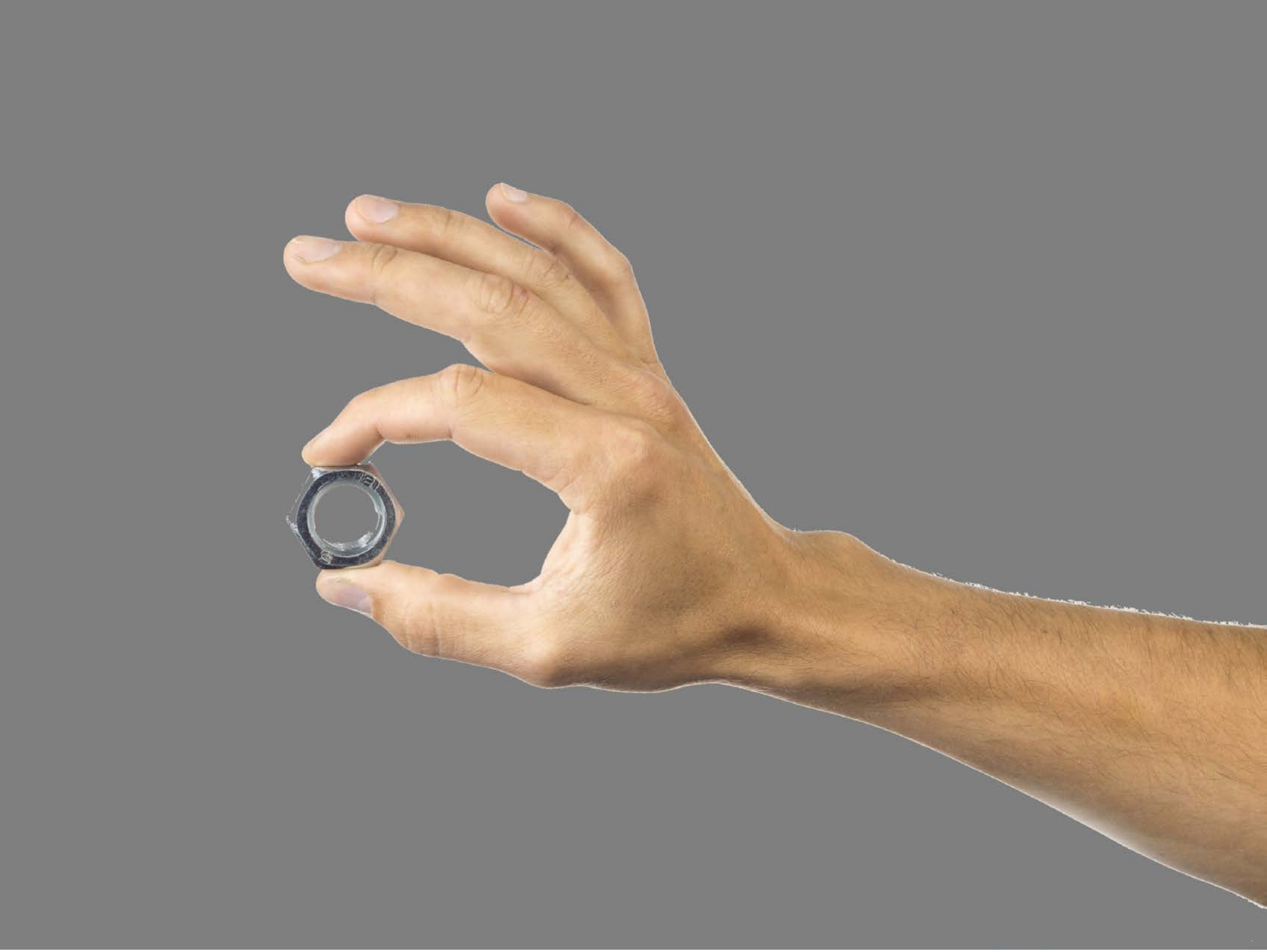

| Category | Craving (0-100) | Valence (1-9) | Arousal (1-9) | Typicality (0-100) | Relatedness | | | | | HSV | | |
|---|---|---|---|---|---|---|---|---|---|---|---|---|
| | | | | | Meth | Opioid | Both | Neither | MethToOpioid | Hue | Saturation | Value |
| Neutral tool hand simple | 13.86 (27.64) | 4.86 (1.627) | 2.25 (1.97) | 10.93 (22) | 0 | 0 | 0 | 1 | 0 | 0.022 (0.067) | 0.082 (0.166) | 0.553 (0.121) |

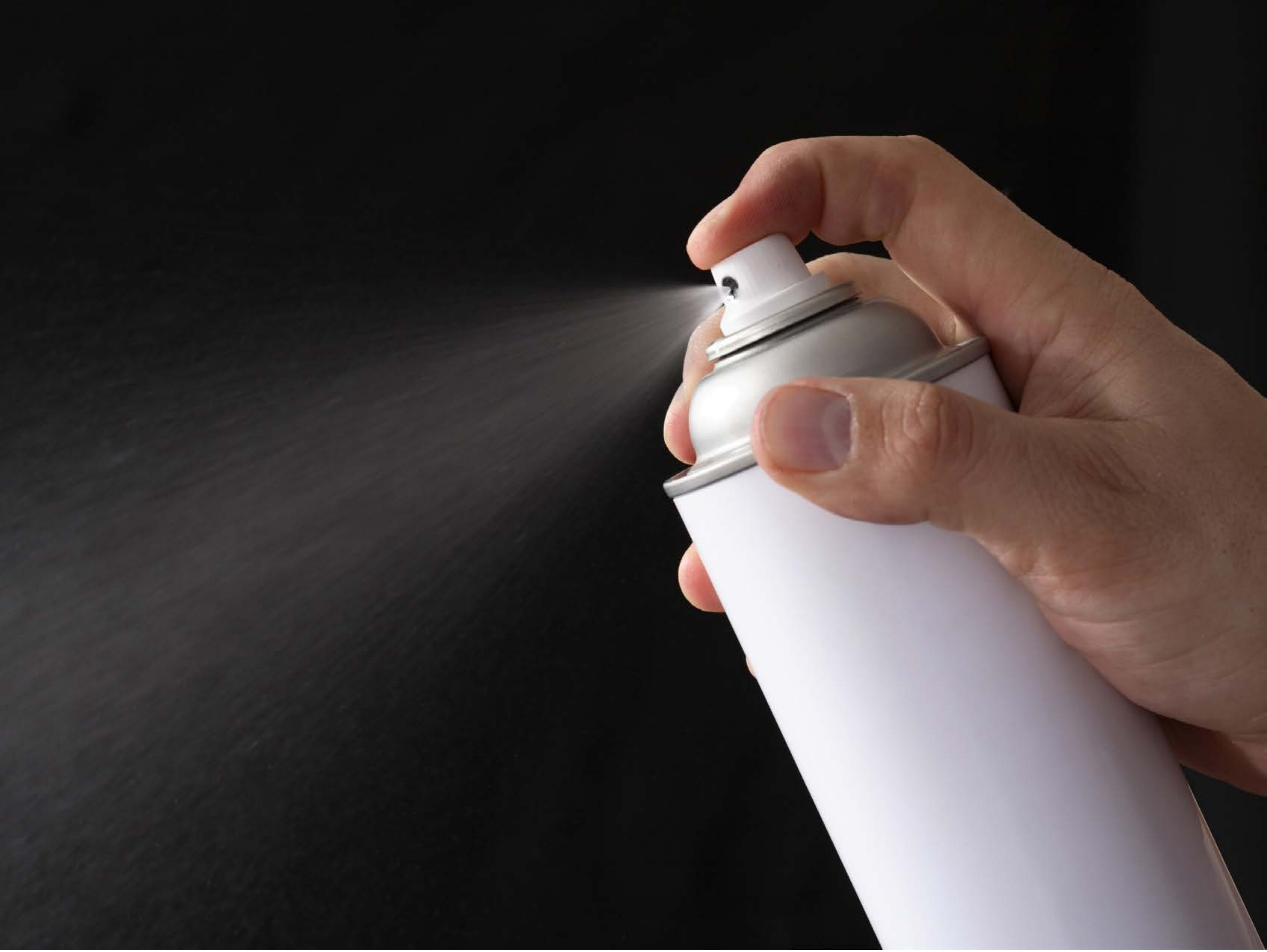

| Category | Craving (0-100) | Valence (1-9) | Arousal (1-9) | Typicality (0-100) | Relatedness | | | | | HSV | | |
|---|---|---|---|---|---|---|---|---|---|---|---|---|
| | | | | | Meth | Opioid | Both | Neither | MethToOpioid | Hue | Saturation | Value |
| Neutral tool hand simple | 12.93 (23.2) | 4.67 (1.441) | 2.89 (2.22) | 11.52 (21.77) | 0 | 0 | 0 | 1 | 0 | 0.379 (0.411) | 0.123 (0.153) | 0.258 (0.264) |

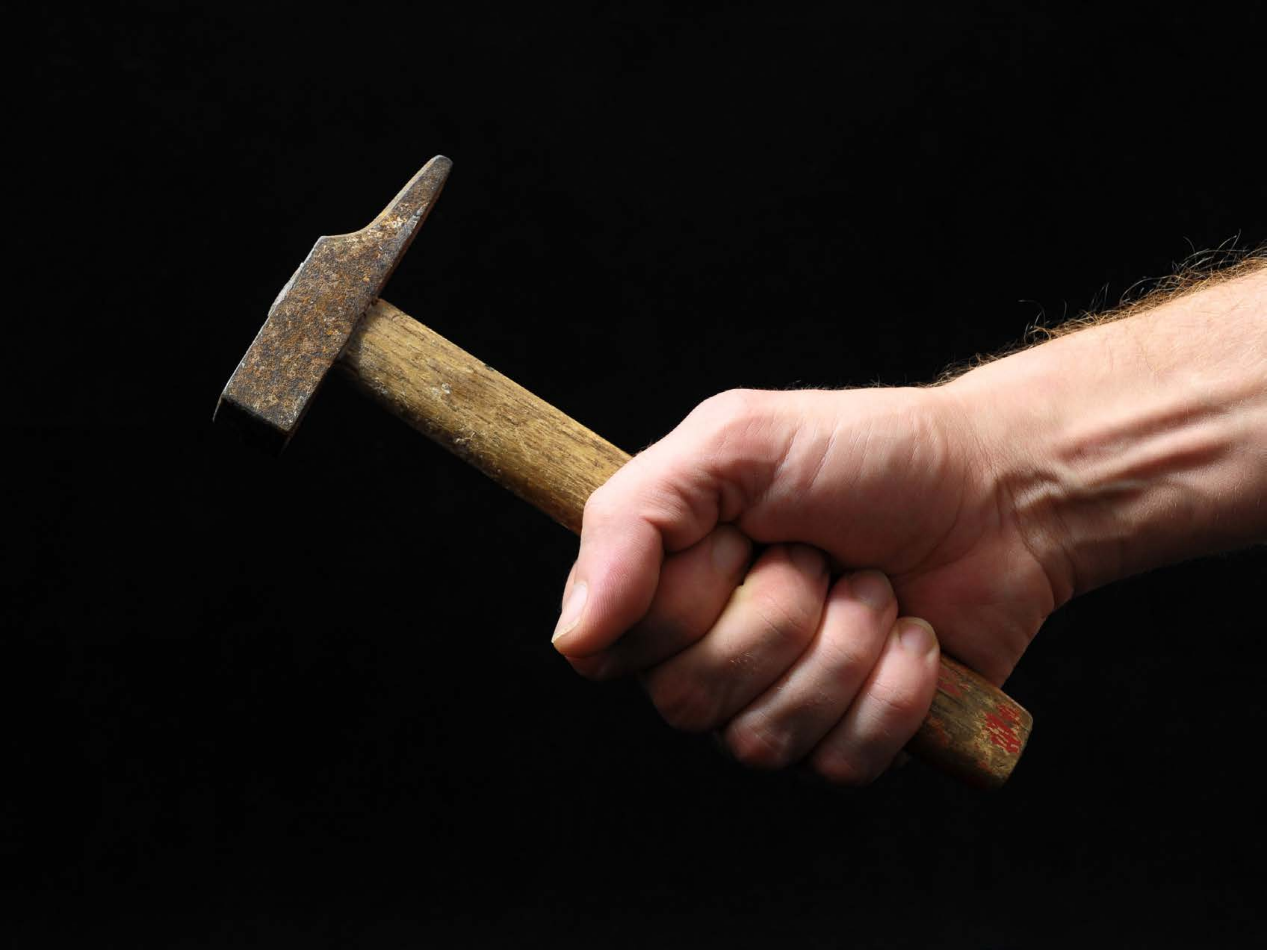

| Category | Craving (0-100) | Valence (1-9) | Arousal (1-9) | Typicality (0-100) | Relatedness | | | | | HSV | | |
|---|---|---|---|---|---|---|---|---|---|---|---|---|
| | | | | | Meth | Opioid | Both | Neither | MethToOpioid | Hue | Saturation | Value |
| Neutral tool hand simple | 10.26 (19.39) | 4.85 (0.77) | 2.56 (1.93) | 11.11 (19.26) | 0 | 0 | 0 | 1 | 0 | 0.016 (0.064) | 0.105 (0.208) | 0.138 (0.26) |

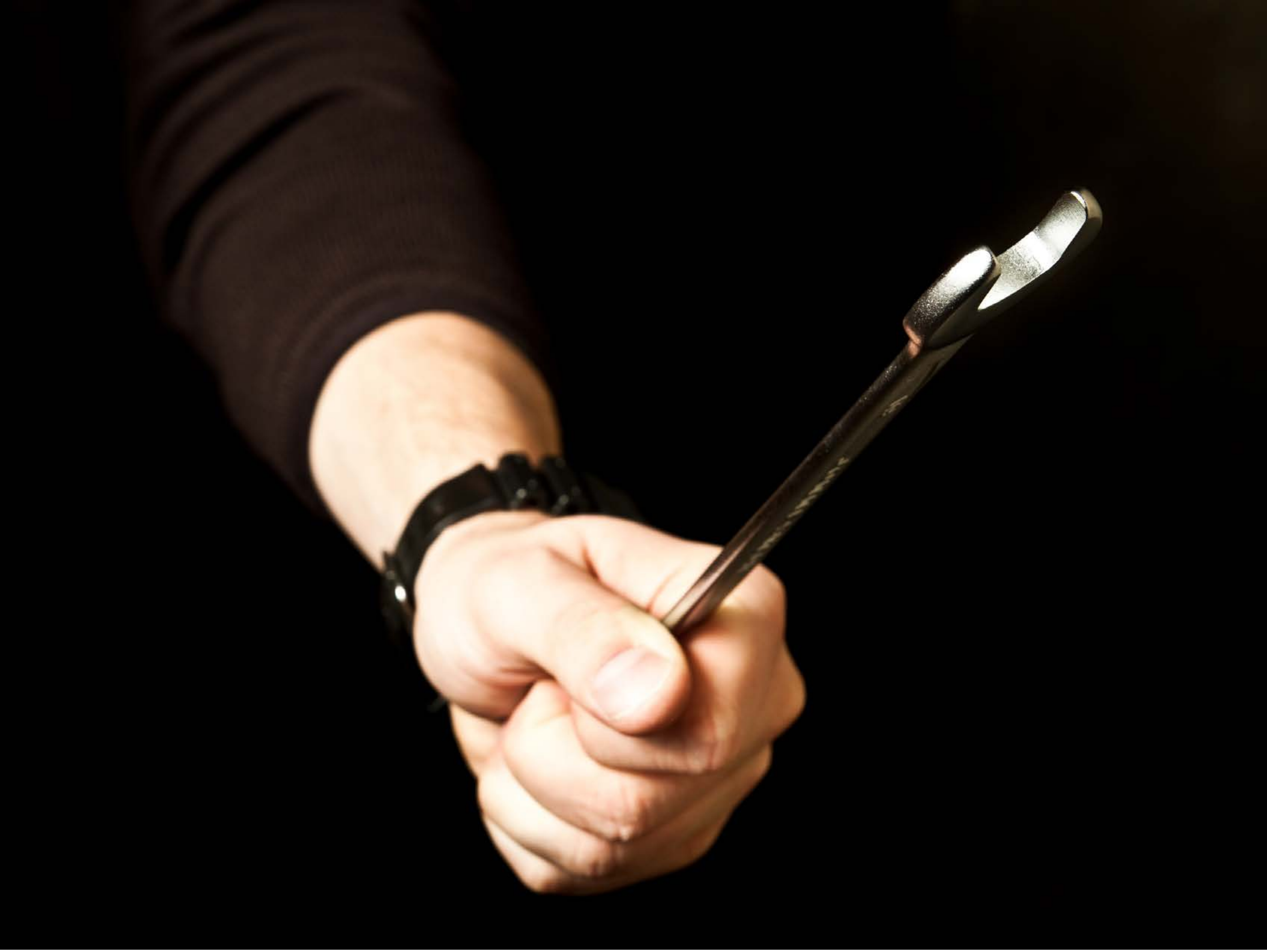

| Category | Craving (0-100) | Valence (1-9) | Arousal (1-9) | Typicality (0-100) | Relatedness | | | | | HSV | | |
|---|---|---|---|---|---|---|---|---|---|---|---|---|
| | | | | | Meth | Opioid | Both | Neither | MethToOpioid | Hue | Saturation | Value |
| Neutral tool hand simple | 12.78 (23.46) | 4.93 (1.269) | 2.7 (2.13) | 15.19 (22.85) | 0.03704 | 0 | 0 | 0.96296 | -0.03704 | 0.038 (0.112) | 0.217 (0.334) | 0.133 (0.292) |

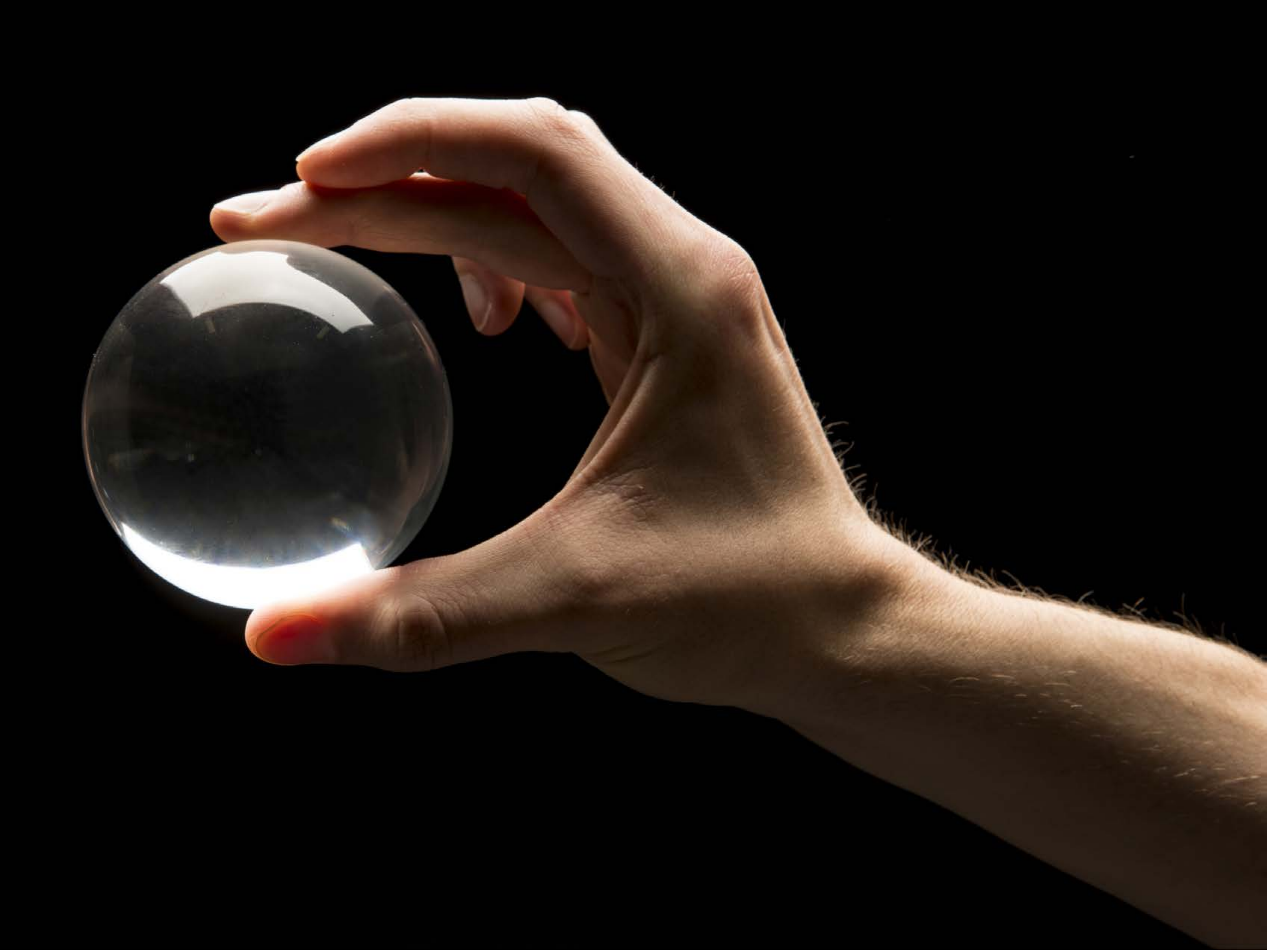

| Category | Craving (0-100) | Valence (1-9) | Arousal (1-9) | Typicality (0-100) | Relatedness | | | | | HSV | | |
|---|---|---|---|---|---|---|---|---|---|---|---|---|
| | | | | | Meth | Opioid | Both | Neither | MethToOpioid | Hue | Saturation | Value |
| Neutral tool hand simple | 12.68 (19.37) | 4.89 (1.343) | 2.32 (1.81) | 11.57 (19.54) | 0.03571 | 0 | 0 | 0.96429 | -0.03571 | 0.037 (0.094) | 0.16 (0.258) | 0.134 (0.244) |

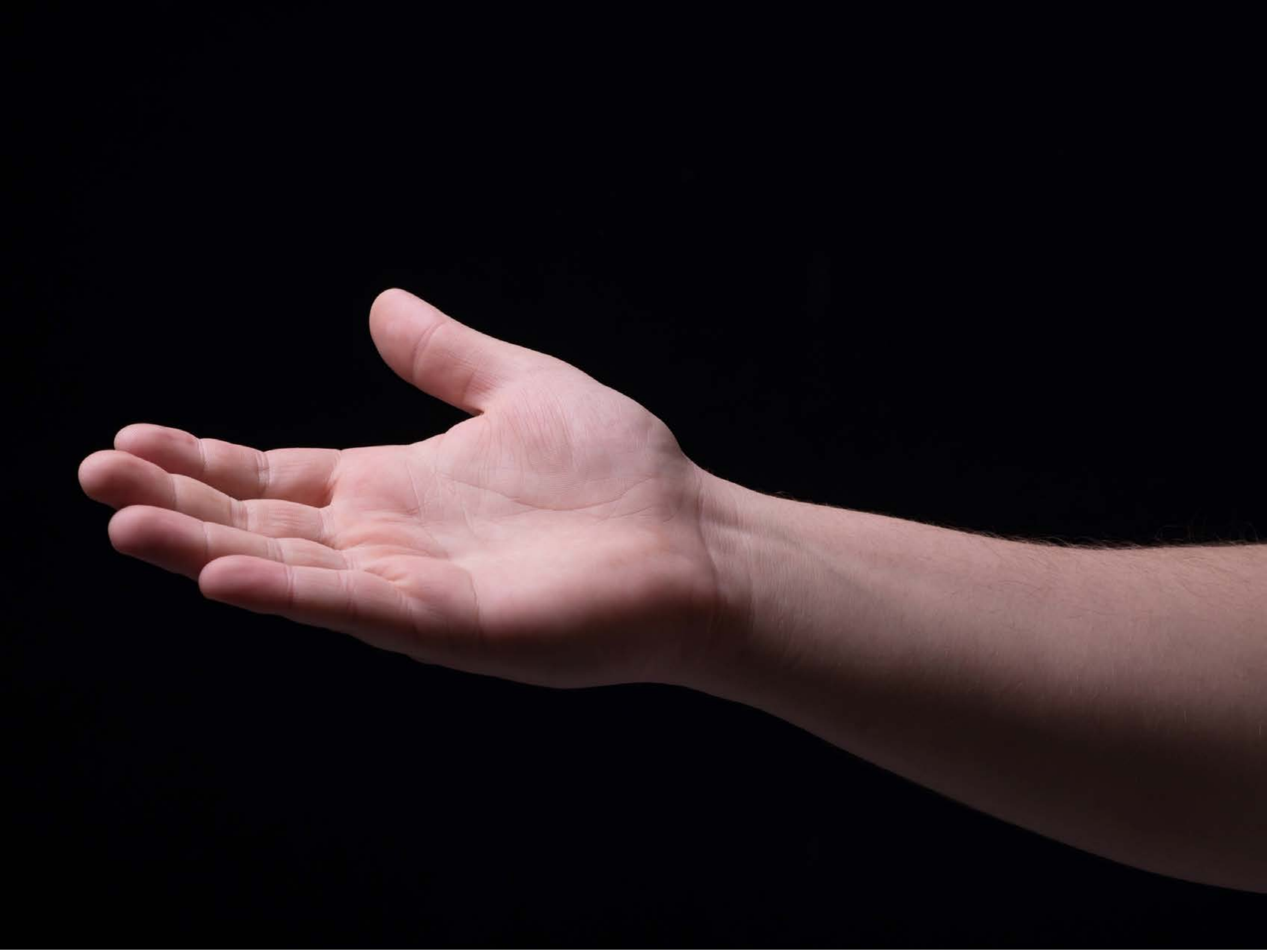

| Category | Craving (0-100) | Valence (1-9) | Arousal (1-9) | Typicality (0-100) | Relatedness | | | | | HSV | | |
|---|---|---|---|---|---|---|---|---|---|---|---|---|
| | | | | | Meth | Opioid | Both | Neither | MethToOpioid | Hue | Saturation | Value |
| Neutral tool hand simple | 13.15 (22.99) | 4.93 (1.207) | 2.74 (2.1) | 19.3 (28.86) | 0 | 0.03704 | 0.03704 | 0.92593 | 0.03704 | 0.567 (0.312) | 0.608 (0.288) | 0.142 (0.25) |

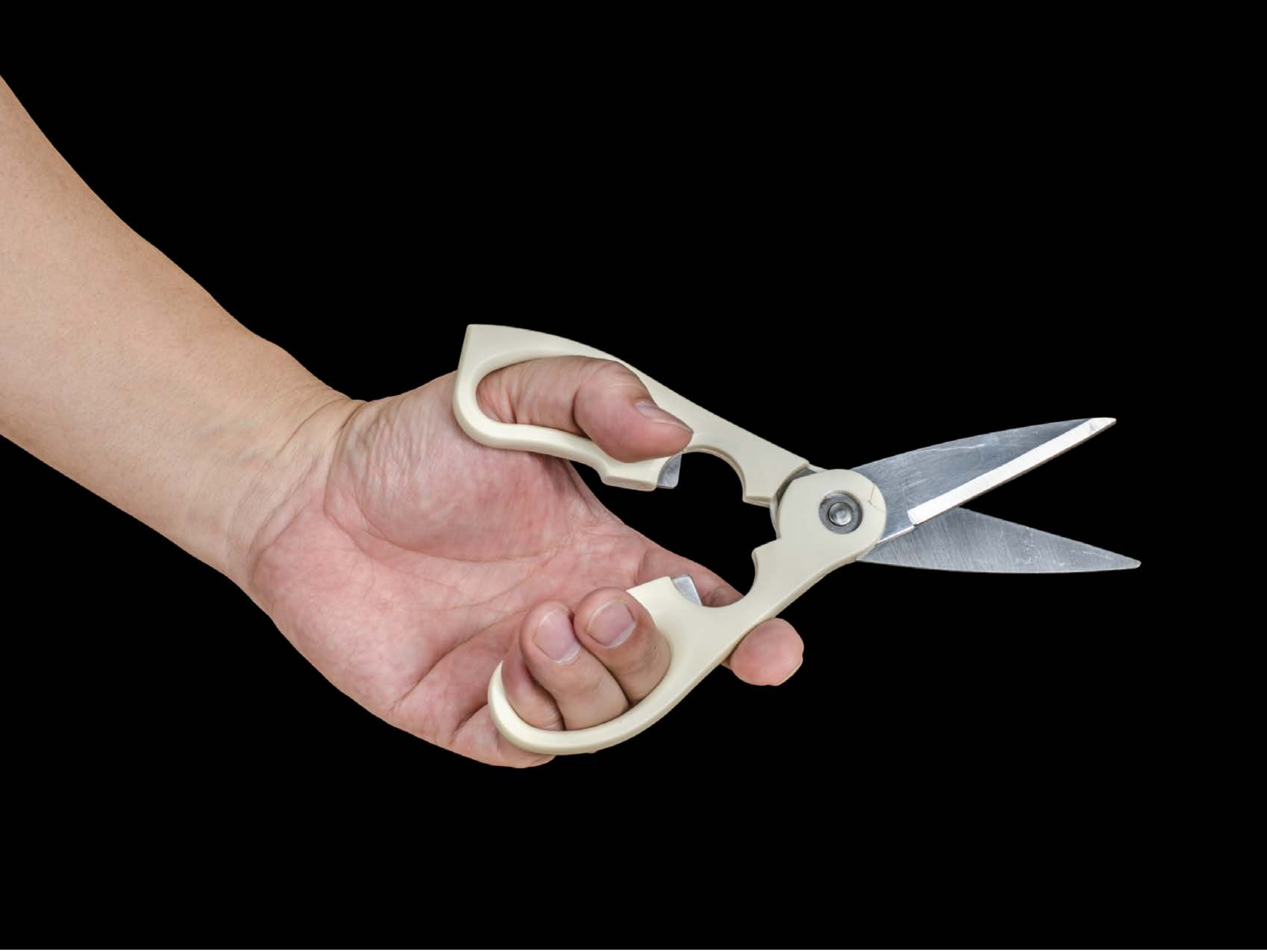

| Category | Craving (0-100) | Valence (1-9) | Arousal (1-9) | Typicality (0-100) | Relatedness | | | | | HSV | | |
|---|---|---|---|---|---|---|---|---|---|---|---|---|
| | | | | | Meth | Opioid | Both | Neither | MethToOpioid | Hue | Saturation | Value |
| Neutral tool hand simple | 7.43 (18.67) | 4.61 (1.75) | 2.21 (1.95) | 9.57 (19.52 | 0 | 0 | 0.03571 | 0.96429 | 0 | 0.029 (0.106) | 0.062 (0.137) | 0.174 (0.321) |

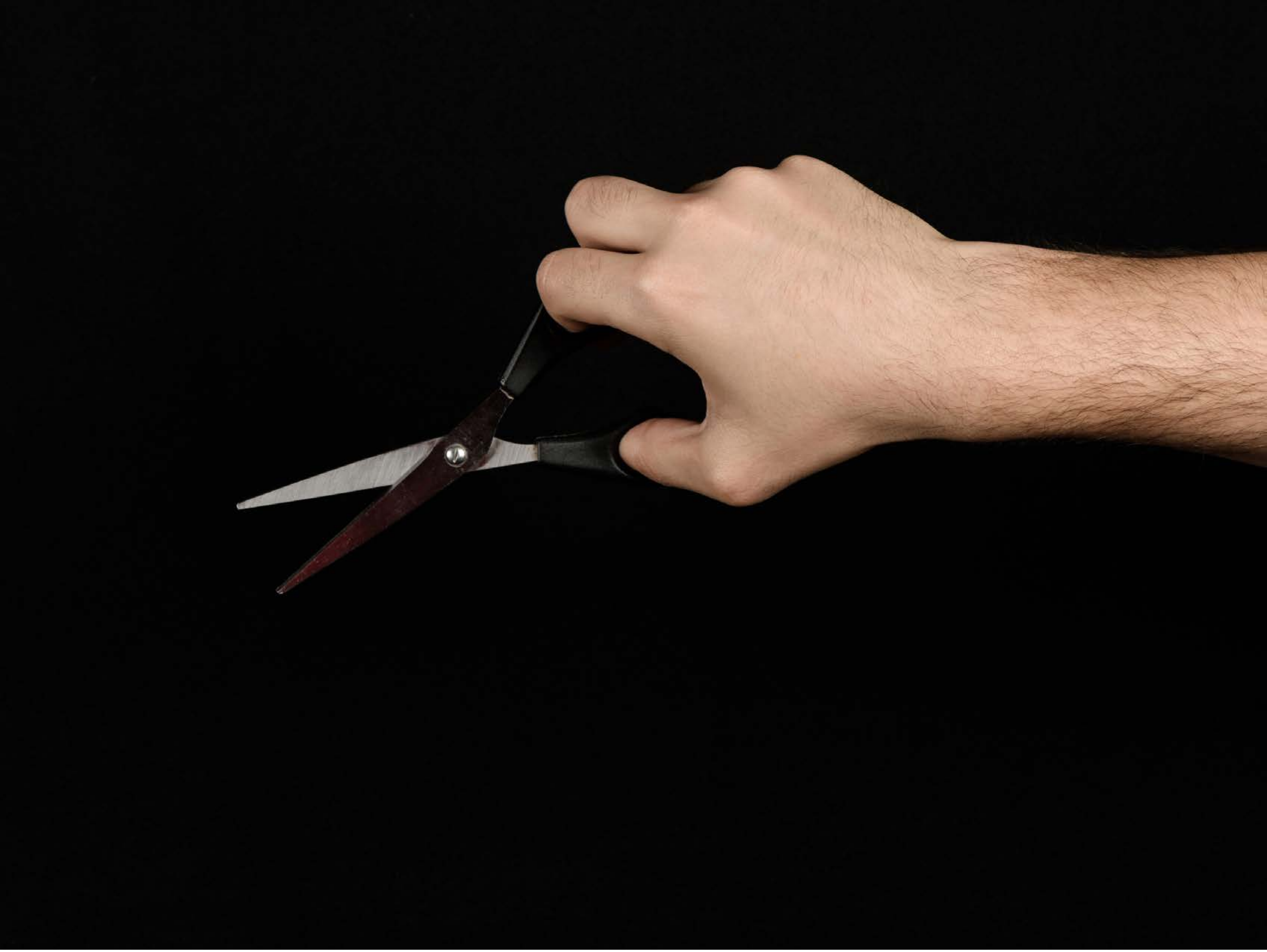

| Category | Craving (0-100) | Valence (1-9) | Arousal (1-9) | Typicality (0-100) | Relatedness | | | | | HSV | | |
|---|---|---|---|---|---|---|---|---|---|---|---|---|
| | | | | | Meth | Opioid | Both | Neither | MethToOpioid | Hue | Saturation | Value |
| Neutral tool hand simple | 9.68 (17.81) | 4.79 (1.371) | 2.46 (1.88) | 9.07 (17.36) | 0 | 0 | 0 | 1 | 0 | 0.018 (0.085 | 0.061 (0.147) | 0.118 (0.258) |

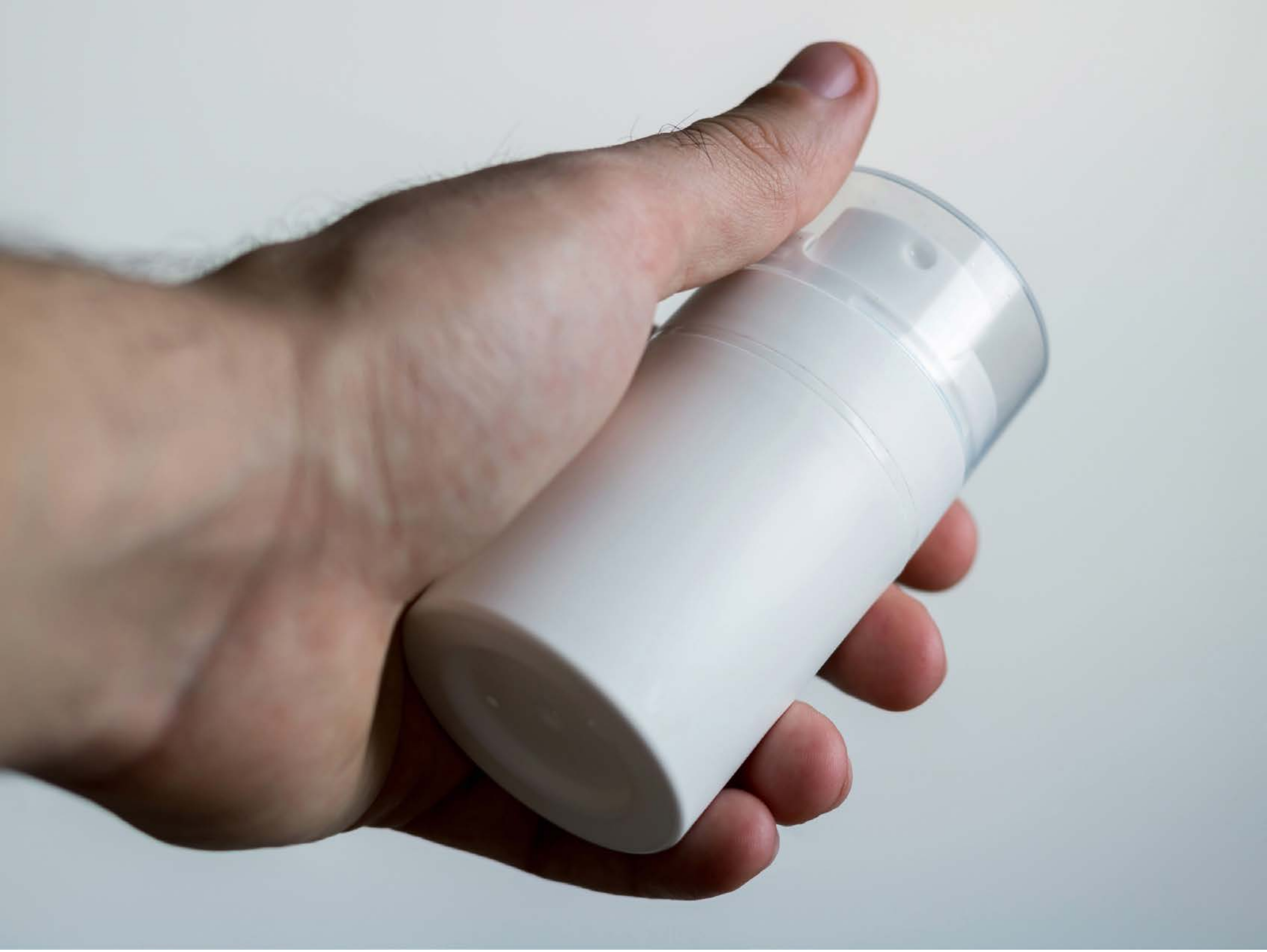

| Category | Craving (0-100) | Valence (1-9) | Arousal (1-9) | Typicality (0-100) | Relatedness | | | | | HSV | | |
|---|---|---|---|---|---|---|---|---|---|---|---|---|
| | | | | | Meth | Opioid | Both | Neither | MethToOpioid | Hue | Saturation | Value |
| Neutral tool hand simple | 16.96 (28.1) | 5.21 (1.228) | 2.75 (2.2) | 16.57 (27.46) | 0 | 0.036 | 0 | 0.964 | 0.036 | 0.367 (0.259) | 0.157 (0.174) | 0.699 (0.194) |

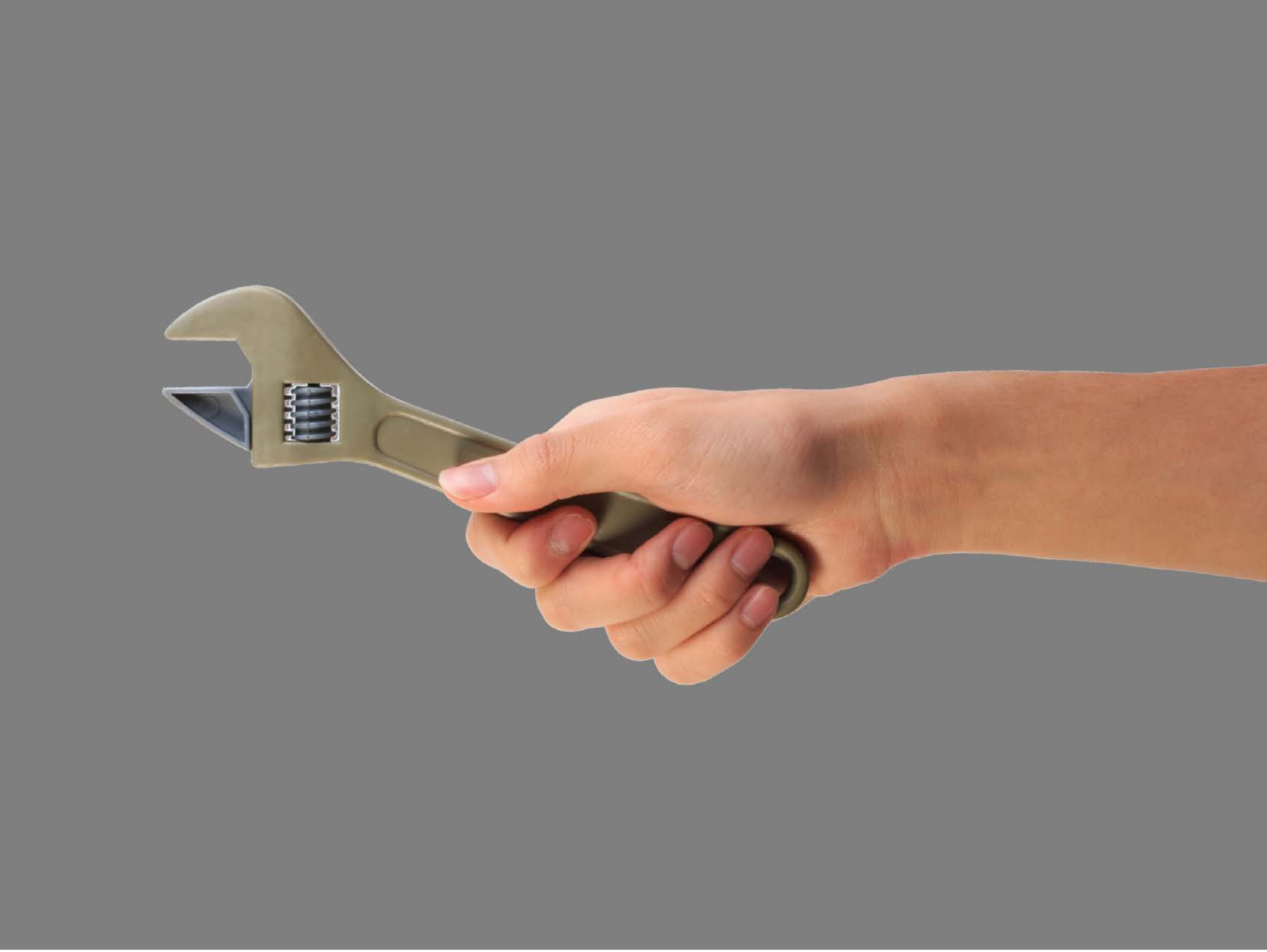

| Category | Craving (0-100) | Valence (1-9) | Arousal (1-9) | Typicality (0-100) | Relatedness | | | | | HSV | | |
|---|---|---|---|---|---|---|---|---|---|---|---|---|
| | | | | | Meth | Opioid | Both | Neither | MethToOpioid | Hue | Saturation | Value |
| Neutral tool hand simple | 10.59 (19.45) | 5.07 (0.385) | 2.3 (1.84) | 12.7 (20.6) | 0 | 0 | 0 | 1 | 0 | 0.016 (0.066) | 0.072 (0.168) | 0.533 (0.102) |

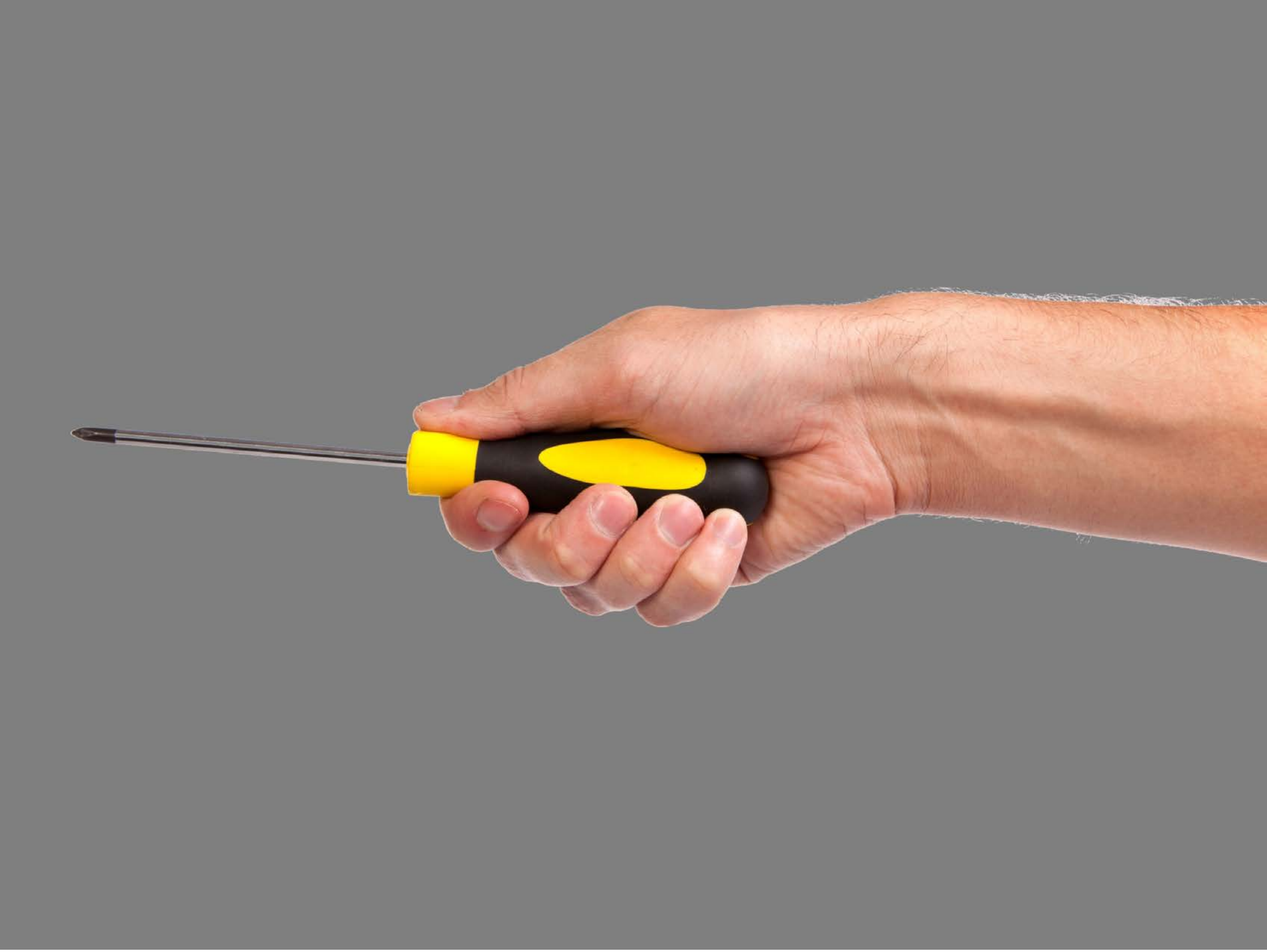

| Category | Craving (0-100) | Valence (1-9) | Arousal (1-9) | Typicality (0-100) | Relatedness | | | | | HSV | | |
|---|---|---|---|---|---|---|---|---|---|---|---|---|
| | | | | | Meth | Opioid | Both | Neither | MethToOpioid | Hue | Saturation | Value |
| Neutral tool hand simple | 8.96 (18.58) | 4.44 (1.251) | 2.44 (1.87) | 9.56 (18.15) | 0 | 0 | 0 | 1 | 0 | 0.015 (0.075) | 0.075 (0.18) | 0.55 (0.14) |

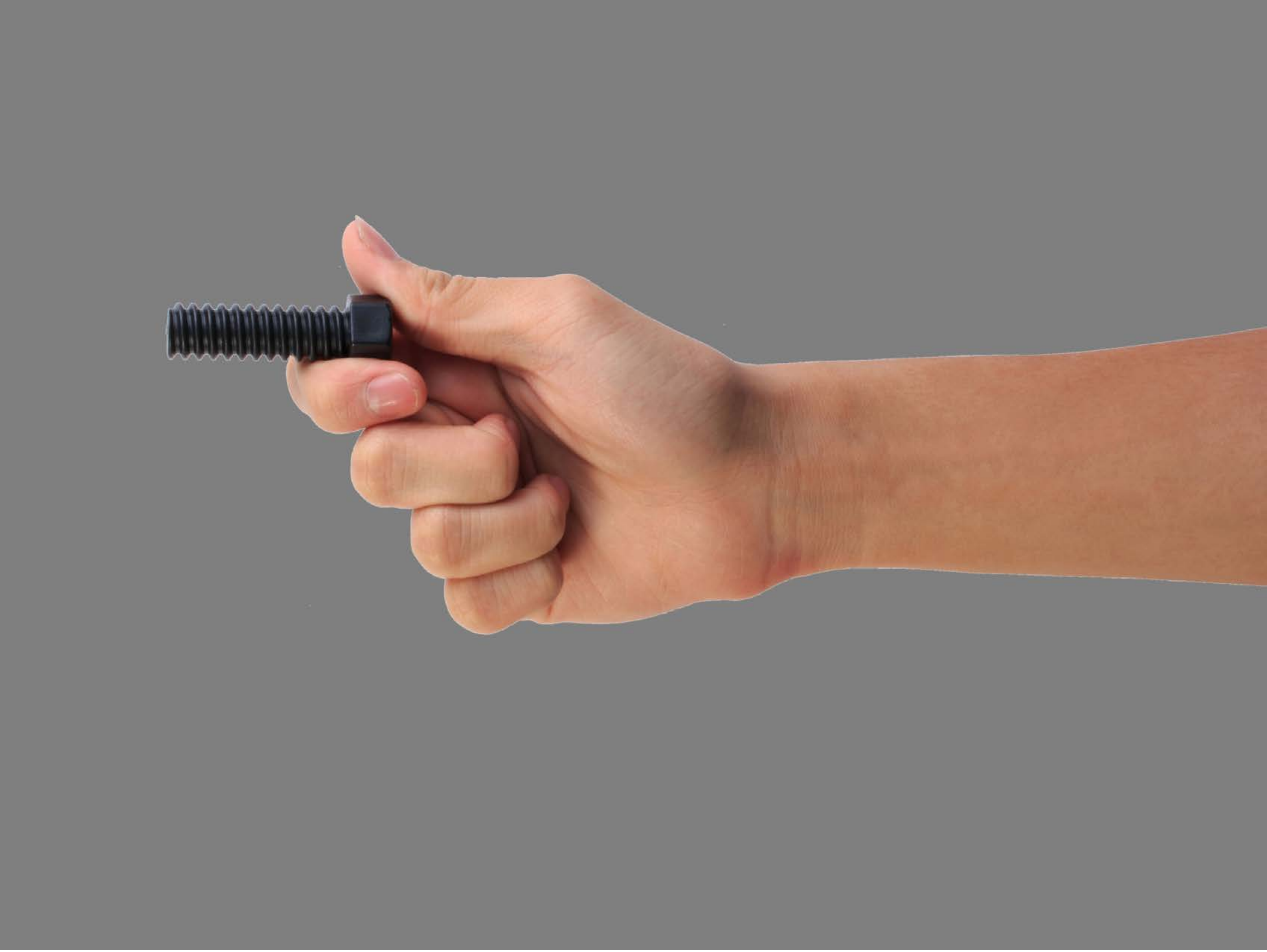

| Category | Craving (0-100) | Valence (1-9) | Arousal (1-9) | Typicality (0-100) | Relatedness | | | | | HSV | | |
|---|---|---|---|---|---|---|---|---|---|---|---|---|
| | | | | | Meth | Opioid | Both | Neither | MethToOpioid | Hue | Saturation | Value |
| Neutral tool hand simple | 11.11 (23.31) | 5.07 (1.562) | 2.36 (2.04) | 13.5 (24.35) | 0 | 0 | 0 | 1 | 0 | 0.019 (0.079) | 0.092 (0.184) | 0.538 (0.106) |

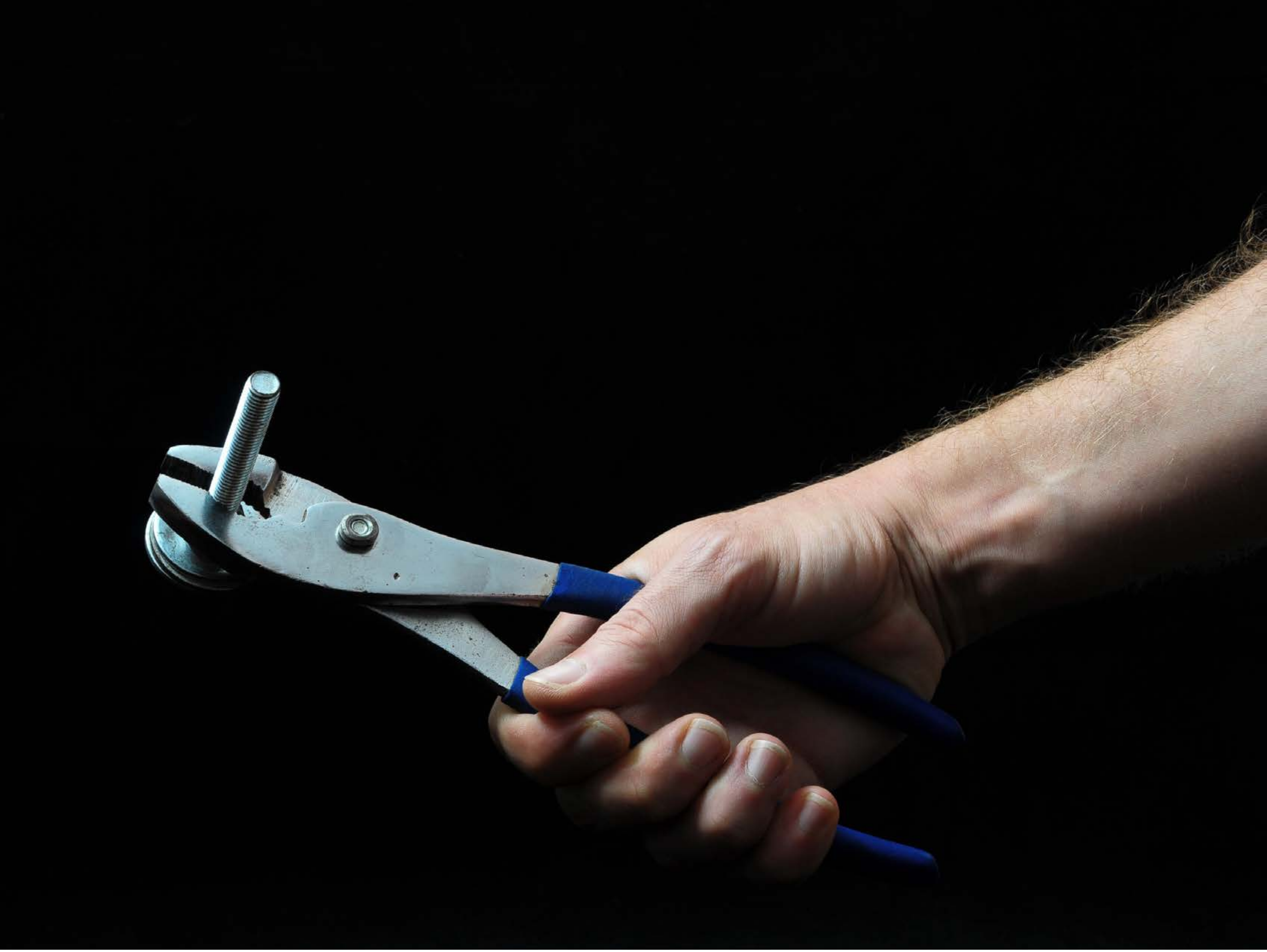

| Category | Craving (0-100) | Valence (1-9) | Arousal (1-9) | Typicality (0-100) | Relatedness | | | | | HSV | | |
|---|---|---|---|---|---|---|---|---|---|---|---|---|
| | | | | | Meth | Opioid | Both | Neither | MethToOpioid | Hue | Saturation | Value |
| Neutral tool hand simple | 12.11 (20.28) | 4.96 (1.453) | 2.25 (1.88) | 13.21 (20.08) | 0 | 0 | 0 | 1 | 0 | 0.061 (0.17) | 0.088 (0.179) | 0.101 (0.228) |

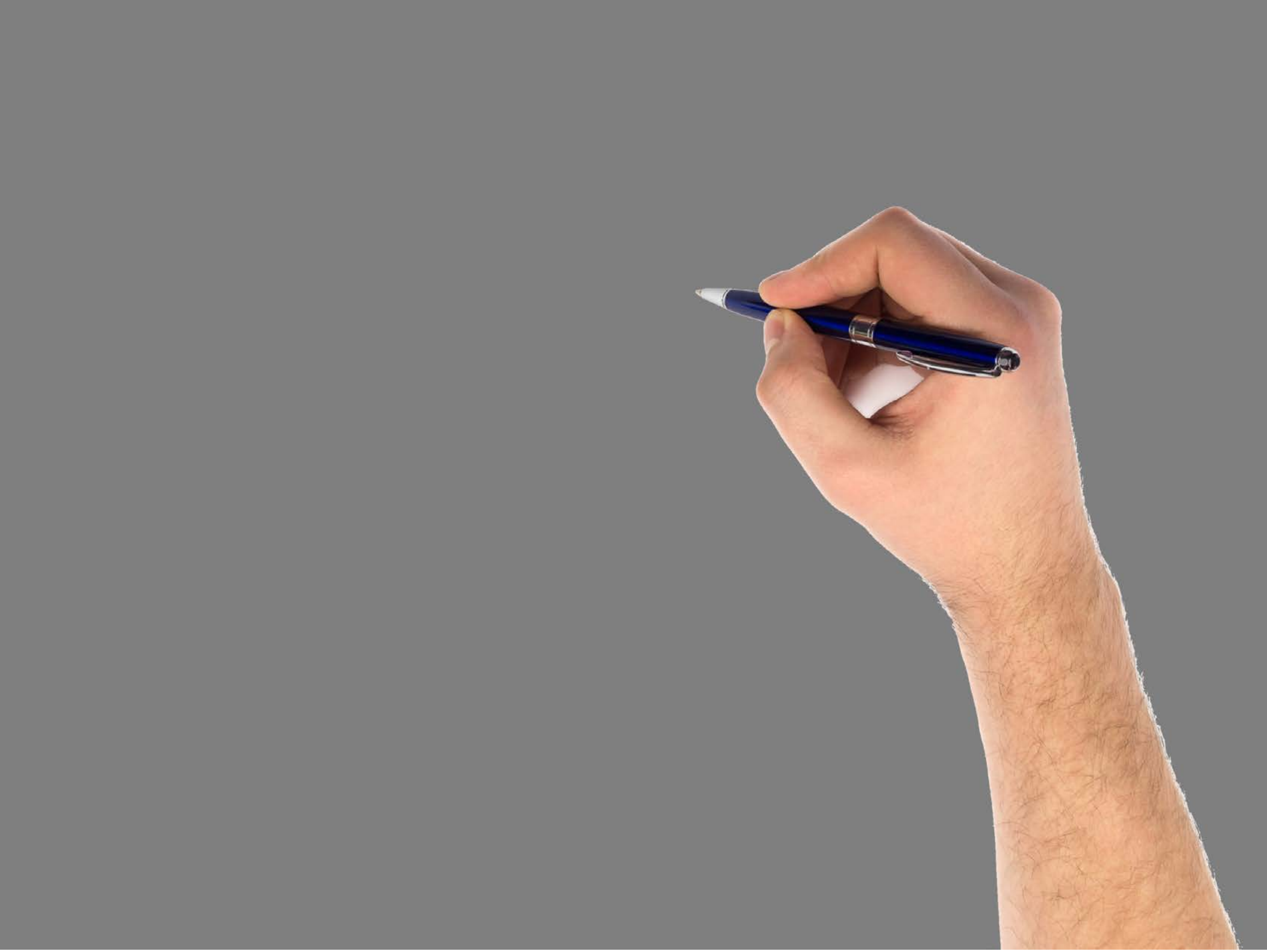

| Category | Craving (0-100) | Valence (1-9) | Arousal (1-9) | Typicality (0-100) | Relatedness | | | | | HSV | | |
|---|---|---|---|---|---|---|---|---|---|---|---|---|
| | | | | | Meth | Opioid | Both | Neither | MethToOpioid | Hue | Saturation | Value |
| Neutral tool hand simple | 8.64 (16.63) | 4.82 (1.634) | 2.14 (1.8) | 10.04 (18.04) | 0 | 0 | 0.036 | 0.964 | 0 | 0.014 (0.072) | 0.054 (0.143) | 0.542 (0.13) |

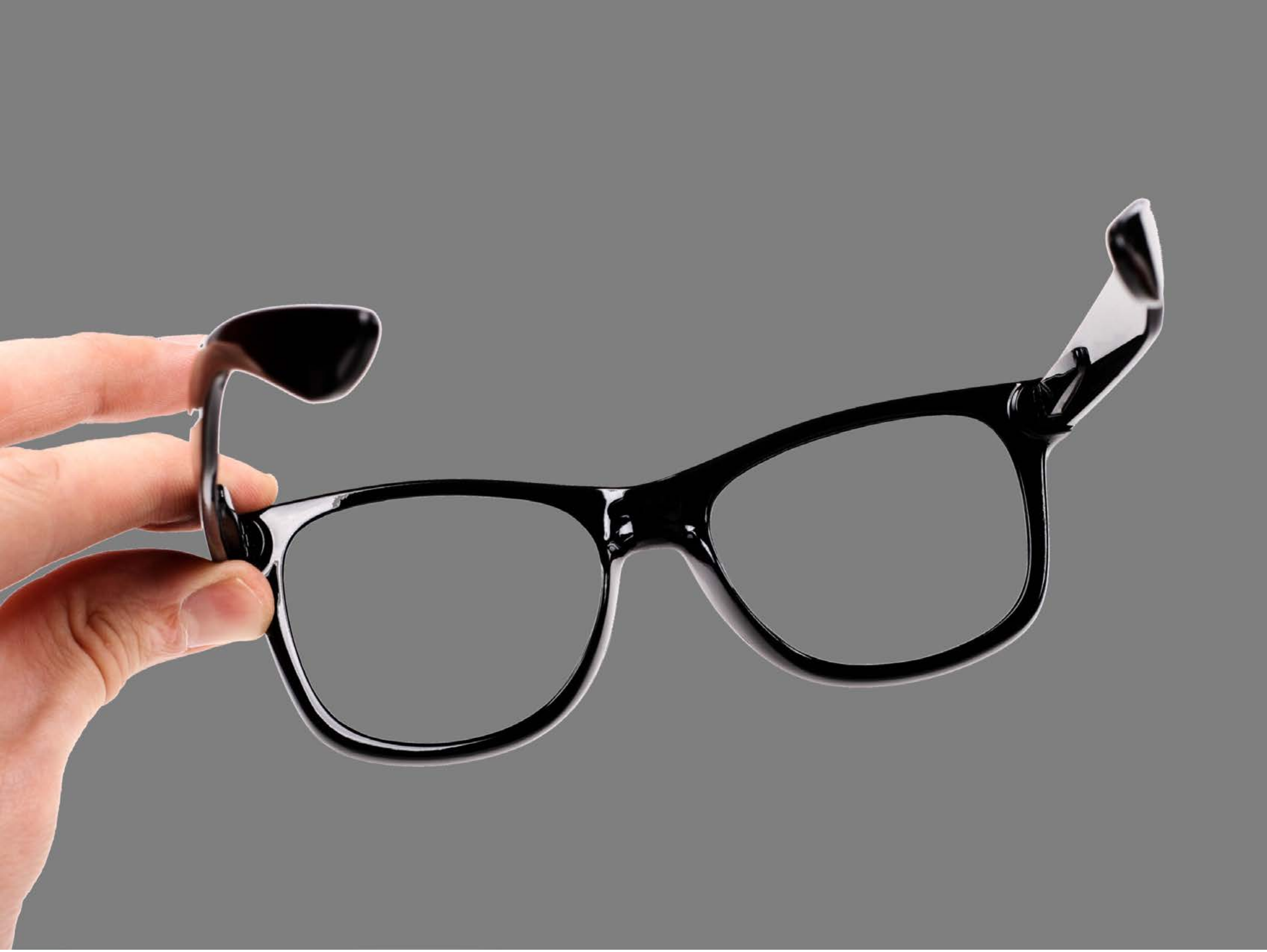

| Category | Craving (0-100) | Valence (1-9) | Arousal (1-9) | Typicality (0-100) | Relatedness | | | | | HSV | | |
|---|---|---|---|---|---|---|---|---|---|---|---|---|
| | | | | | Meth | Opioid | Both | Neither | MethToOpioid | Hue | Saturation | Value |
| Neutral tool hand simple | 7.85 (16.21) | 4.85 (1.35) | 2.56 (1.93) | 11.26 (19.27) | 0 | 0 | 0 | 1 | 0 | 0.038 (0.169) | 0.043 (0.146) | 0.5 (0.146) |

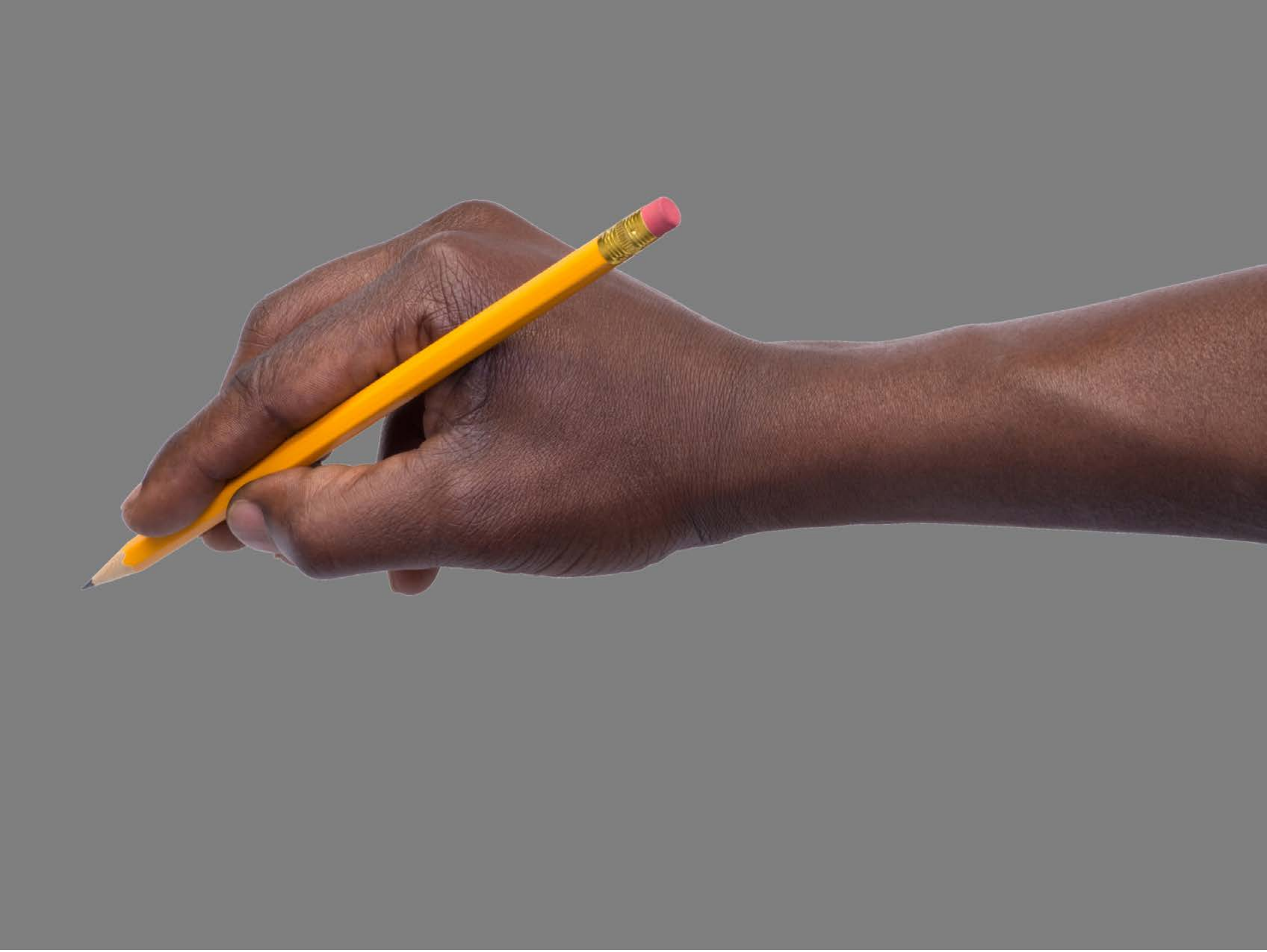

| Category | Craving (0-100) | Valence (1-9) | Arousal (1-9) | Typicality (0-100) | Relatedness | | | | | HSV | | |
|---|---|---|---|---|---|---|---|---|---|---|---|---|
| | | | | | Meth | Opioid | Both | Neither | MethToOpioid | Hue | Saturation | Value |
| Neutral tool hand simple | 16.41 (29.76) | 4.56 (1.368) | 2.63 (2.06) | 15.04 (25.44 | 0 | 0 | 0 | 1 | 0 | 0.052 (0.202 | 0.094 (0.188) | 0.488 (0.081) |

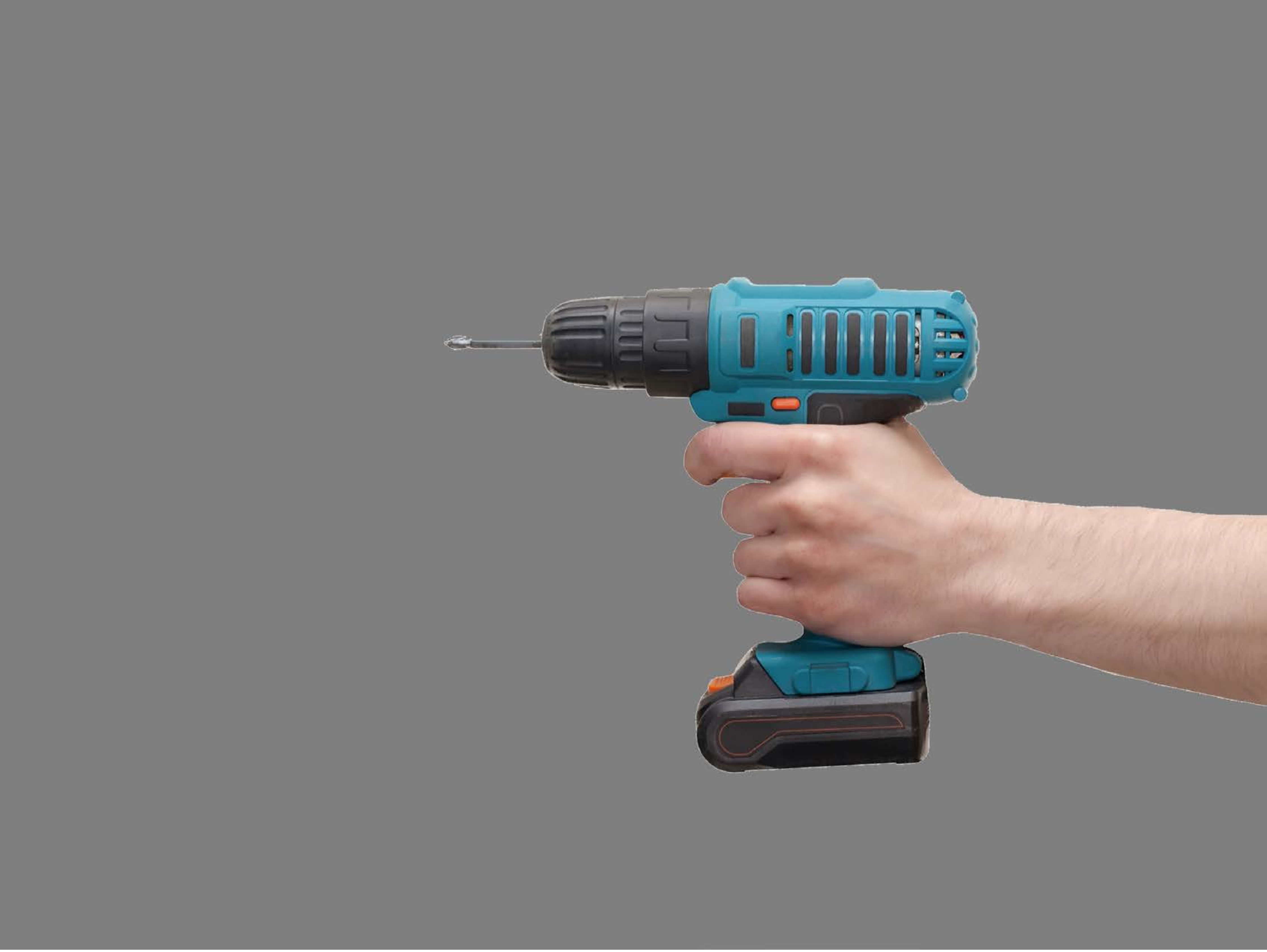

| Category | Craving (0-100) | Valence (1-9) | Arousal (1-9) | Typicality (0-100) | Relatedness | | | | | HSV | | |
|---|---|---|---|---|---|---|---|---|---|---|---|---|
| | | | | | Meth | Opioid | Both | Neither | MethToOpioid | Hue | Saturation | Value |
| Neutral tool hand simple | 6.79 (15.44) | 4.64 (1.545) | 2.39 (1.87) | 7.71 (15.23) | 0 | 0 | 0 | 1 | 0 | 0.042 (0.156) | 0.045 (0.134) | 0.512 (0.098) |

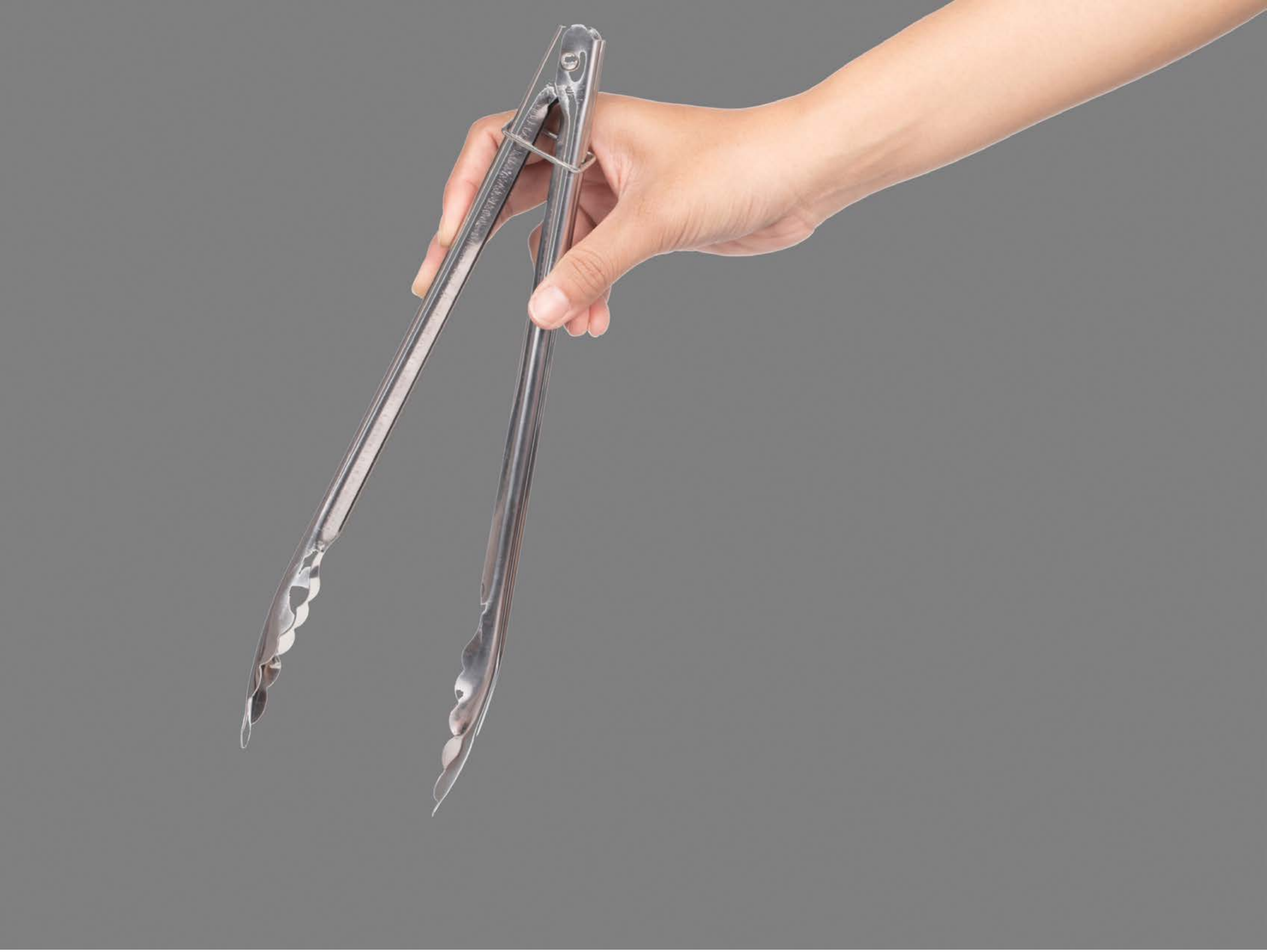

| Category | Craving (0-100) | Valence (1-9) | Arousal (1-9) | Typicality (0-100) | Relatedness | | | | | HSV | | |
|---|---|---|---|---|---|---|---|---|---|---|---|---|
| | | | | | Meth | Opioid | Both | Neither | MethToOpioid | Hue | Saturation | Value |
| Neutral tool hand simple | 13.68 (25.41) | 4.93 (1.386) | 2.29 (1.9) | 16.64 (25.77) | 0 | 0 | 0 | 1 | 0 | 0.03 (0.139) | 0.032 (0.089) | 0.541 (0.129) |

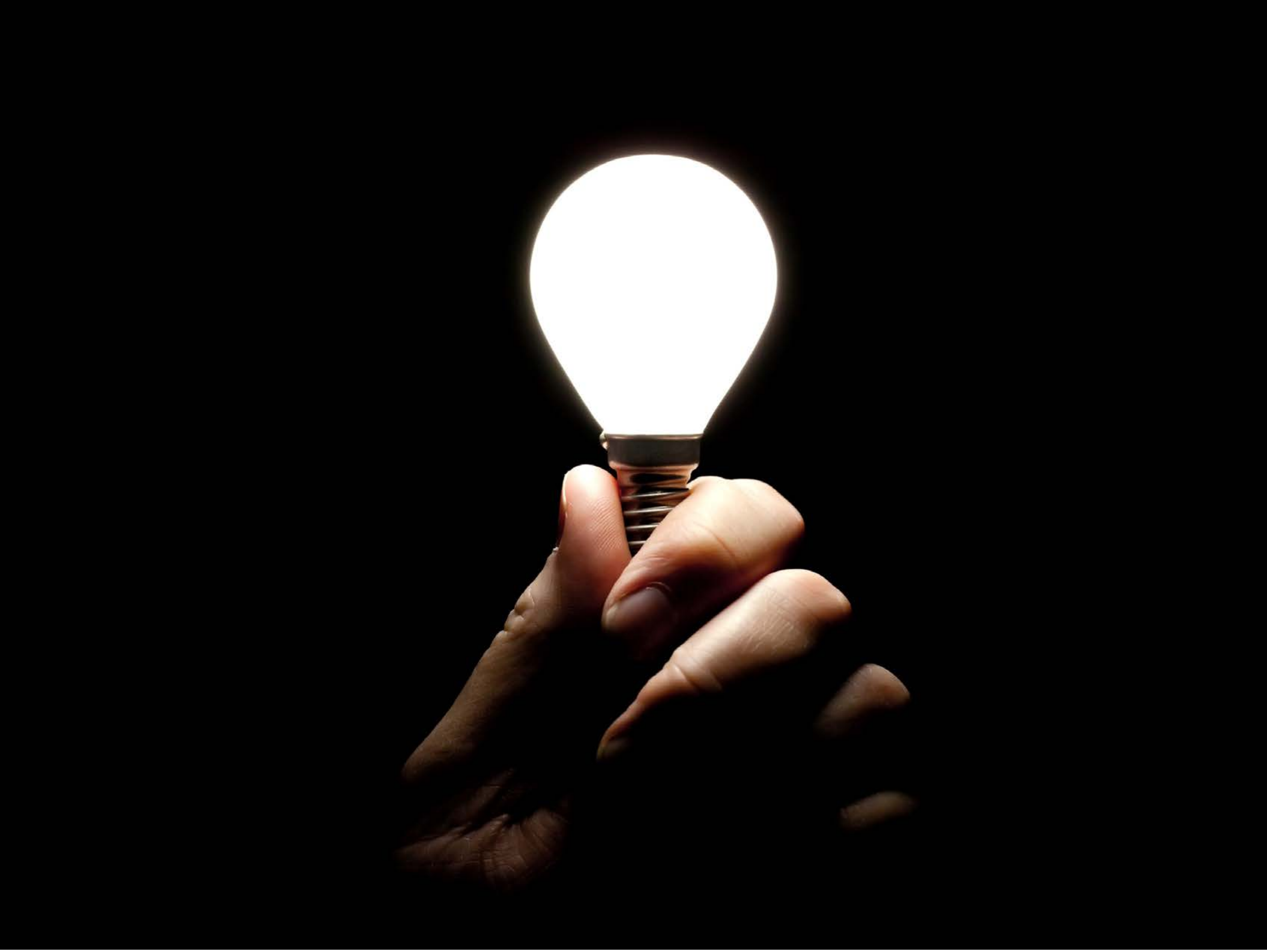

| Category | Craving (0-100) | Valence (1-9) | Arousal (1-9) | Typicality (0-100) | Relatedness | | | | | HSV | | |
|---|---|---|---|---|---|---|---|---|---|---|---|---|
| | | | | | Meth | Opioid | Both | Neither | MethToOpioid | Hue | Saturation | Value |
| Neutral tool hand simple | 41.81 (31.41) | 4.96 (0.759) | 2.93 (2.15) | 42 (30.71) | 0.333 | | 0.037 | 0.630 | -0.333 | 0.015 (0.097) | 0.061 (0.203) | 0.075 (0.241) |

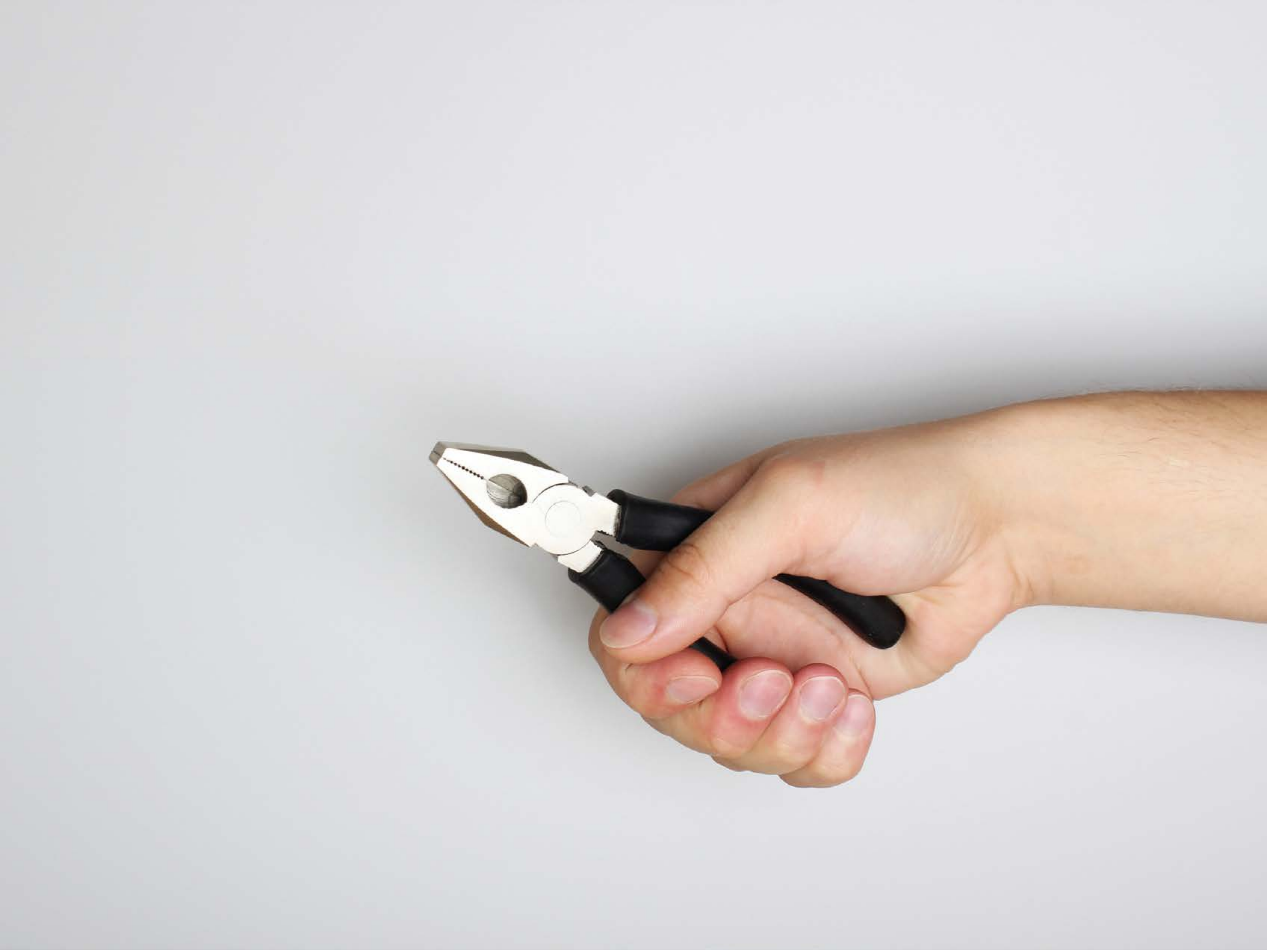

| Category | Craving (0-100) | Valence (1-9) | Arousal (1-9) | Typicality (0-100) | Relatedness | | | | | HSV | | |
|---|---|---|---|---|---|---|---|---|---|---|---|---|
| | | | | | Meth | Opioid | Both | Neither | MethToOpioid | Hue | Saturation | Value |
| Neutral tool hand simple | 8.85 (17.64) | 4.74 (0.944) | 2.44 (1.87) | 9.78 (17.65) | 0 | 0 | 0 | 1 | 0 | 0.507 (0.236) | 0.056 (0.108) | 0.859 (0.099) |

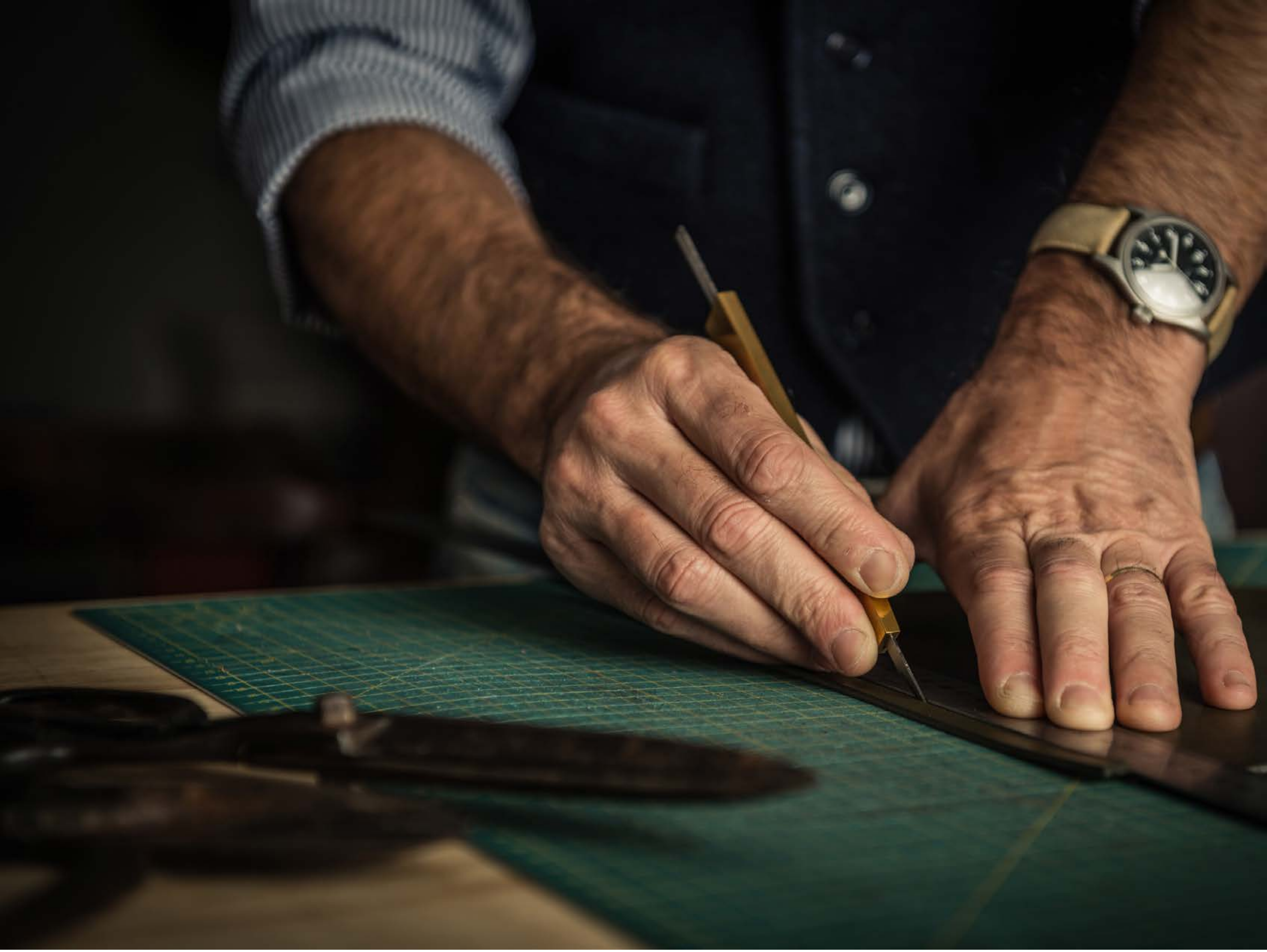

| Category | Craving (0-100) | Valence (1-9) | Arousal (1-9) | Typicality (0-100) | Relatedness | | | | | HSV | | |
|---|---|---|---|---|---|---|---|---|---|---|---|---|
| | | | | | Meth | Opioid | Both | Neither | MethToOpioid | Hue | Saturation | Value |
| Neutral tool hand complex | 7.7 (15.86\|) | 4.85 (0.818) | 2.33 (1.82) | 7.19 (15.68) | 0 | 0 | 0 | 1 | 0 | 0.244 (0.244) | 0.359 (0.169) | 0.226 (0.206) |

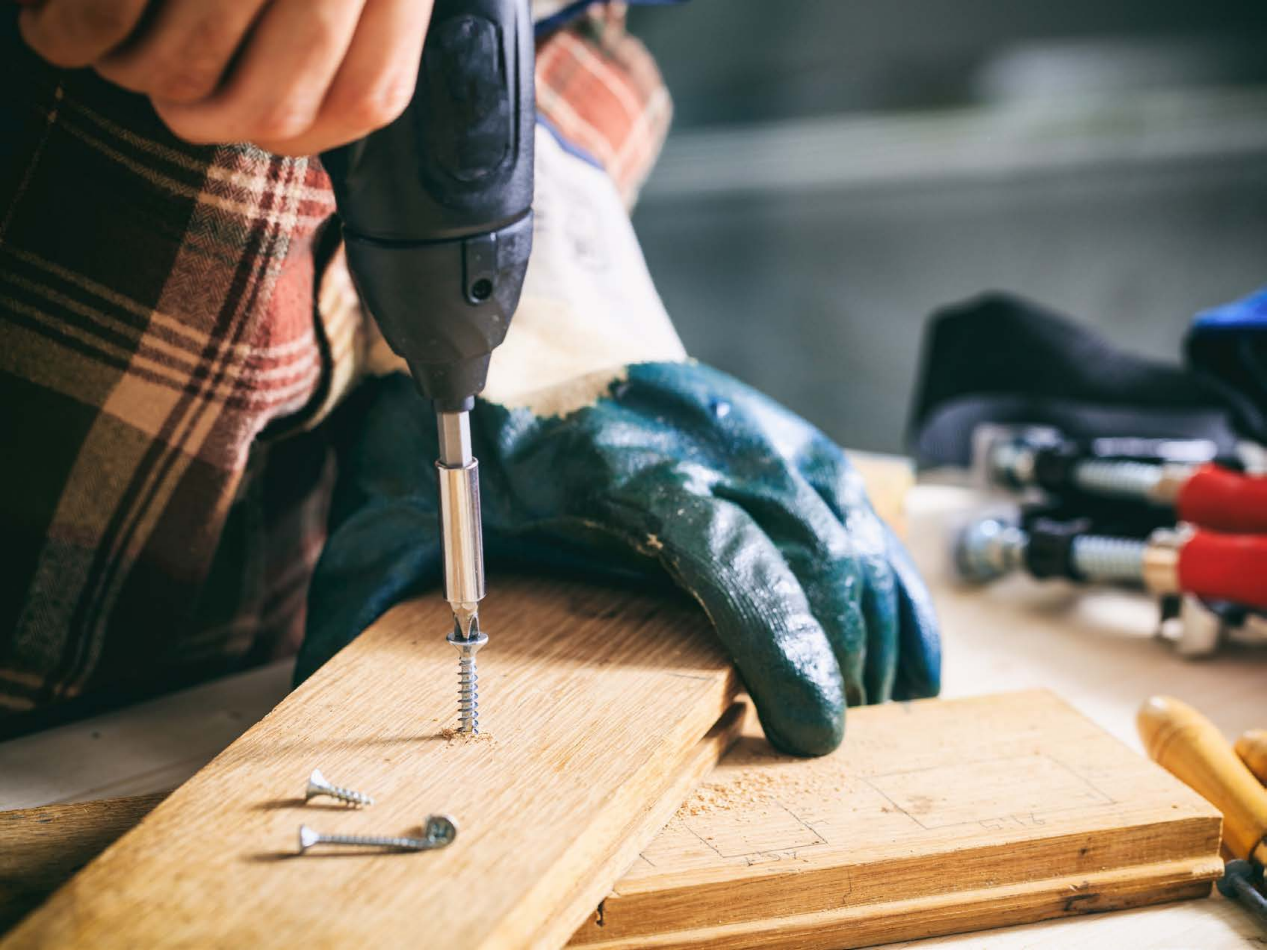

| Category | Craving (0-100) | Valence (1-9) | Arousal (1-9) | Typicality (0-100) | Relatedness | | | | | HSV | | |
|---|---|---|---|---|---|---|---|---|---|---|---|---|
| | | | | | Meth | Opioid | Both | Neither | MethToOpioid | Hue | Saturation | Value |
| Neutral tool hand complex | 10.07 (18.05) | 4.89 (1.05) | 2.52 (1.95) | 10.74 (17.92) | 0 | 0 | 0 | 1 | 0 | 0.298 (0.225) | 0.338 (0.203) | 0.532 (0.316) |

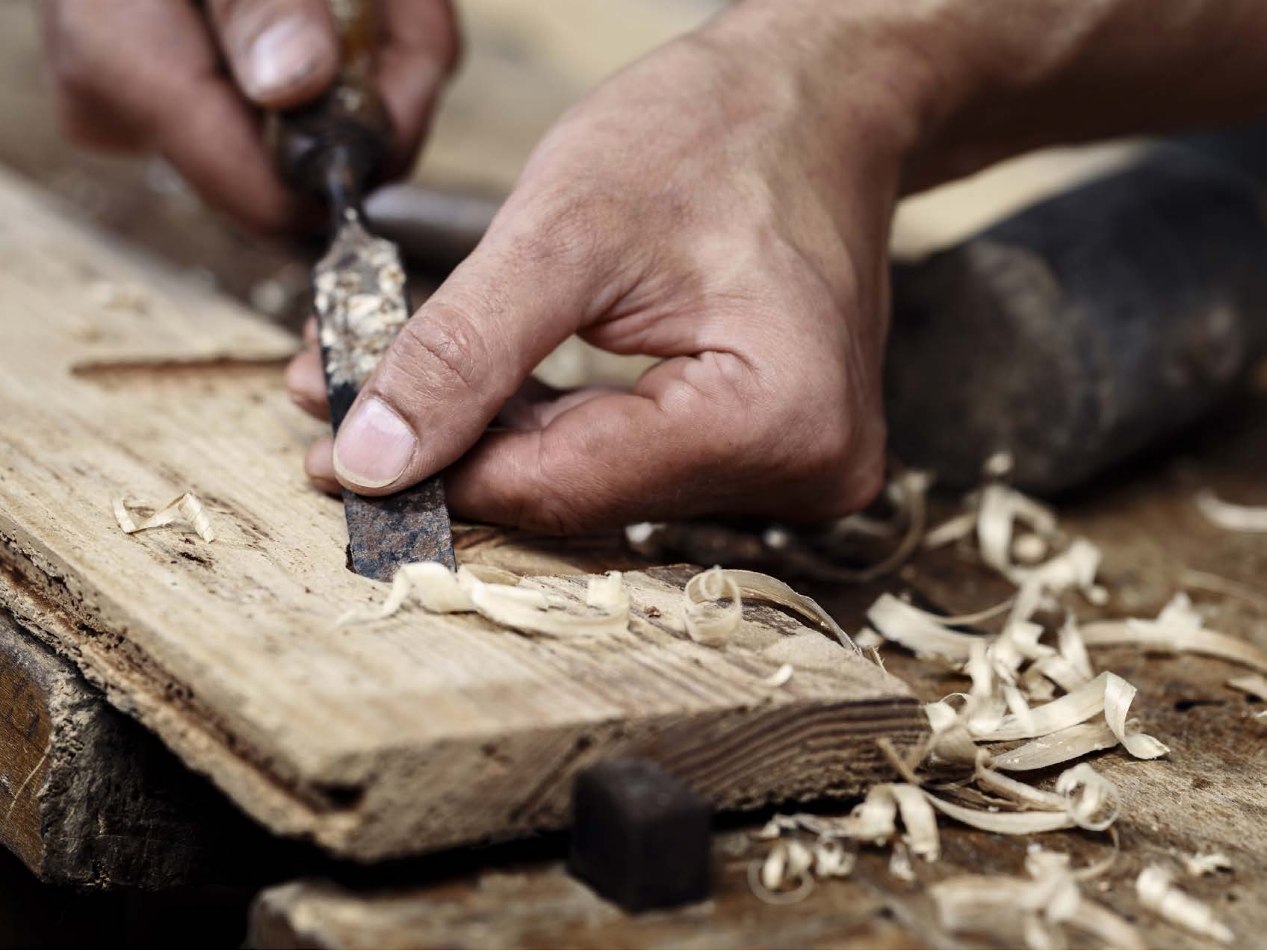

| Category | Craving (0-100) | Valence (1-9) | Arousal (1-9) | Typicality (0-100) | Relatedness | | | | | HSV | | |
|---|---|---|---|---|---|---|---|---|---|---|---|---|
| | | | | | Meth | Opioid | Both | Neither | MethToOpioid | Hue | Saturation | Value |
| Neutral tool hand complex | 5.5 (11.81) | 5.11 (1.771) | 2.14 (1.8) | 7.96 (15.56) | 0 | 0 | 0 | 1 | 0 | 0.14 (0.232) | 0.298 (0.163) | 0.485 (0.269) |

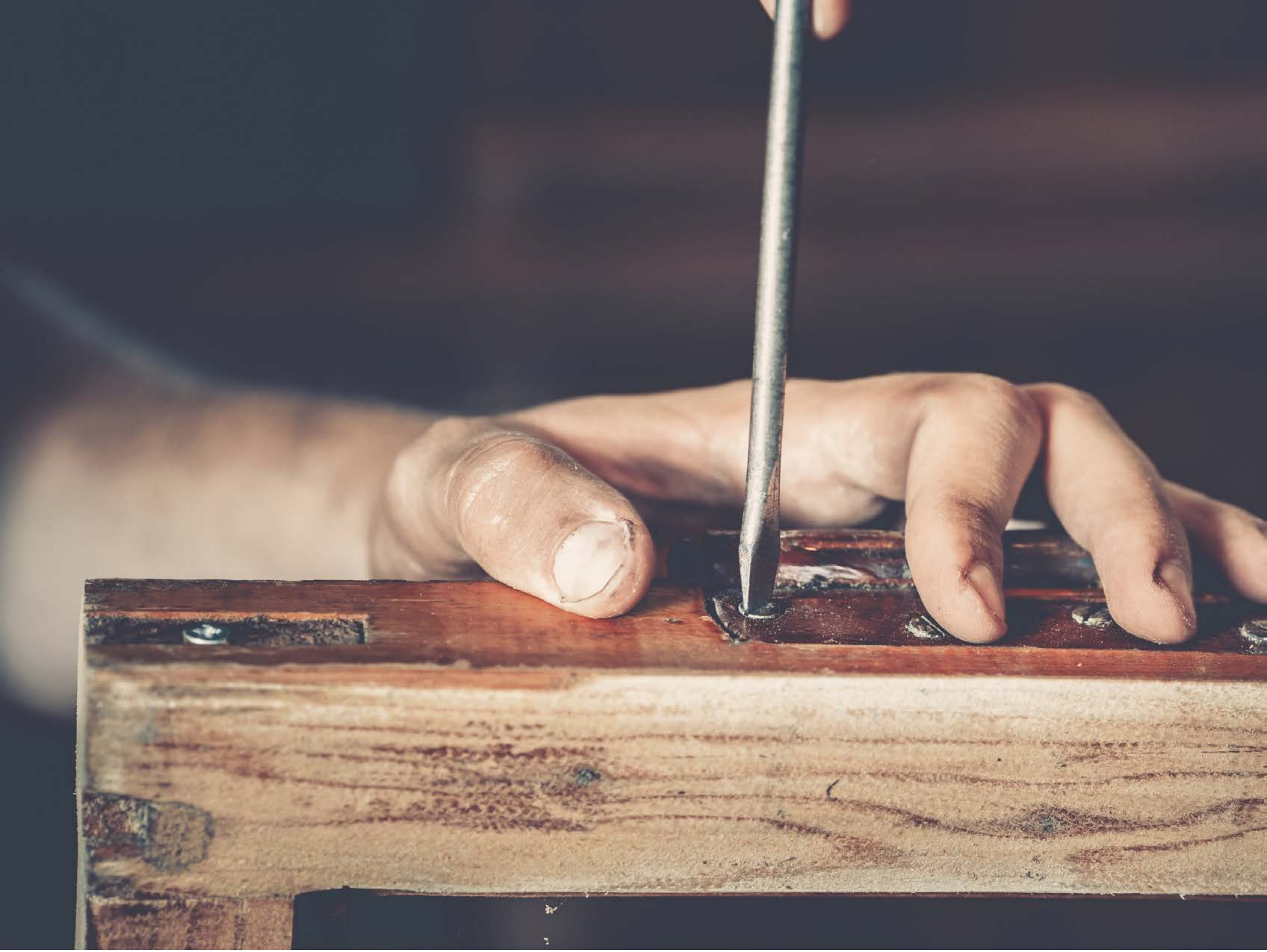

| Category | Craving (0-100) | Valence (1-9) | Arousal (1-9) | Typicality (0-100) | Relatedness | | | | | HSV | | |
|---|---|---|---|---|---|---|---|---|---|---|---|---|
| | | | | | Meth | Opioid | Both | Neither | MethToOpioid | Hue | Saturation | Value |
| Neutral tool hand complex | 7.56 (15.89 | 4.48 (1.312) | 2.37 (1.9) | 8.74 (15.87) | 0 | 0 | 0 | 1 | 0 | 0.424 (0.332) | 0.221 (0.09) | 0.498 (0.244) |

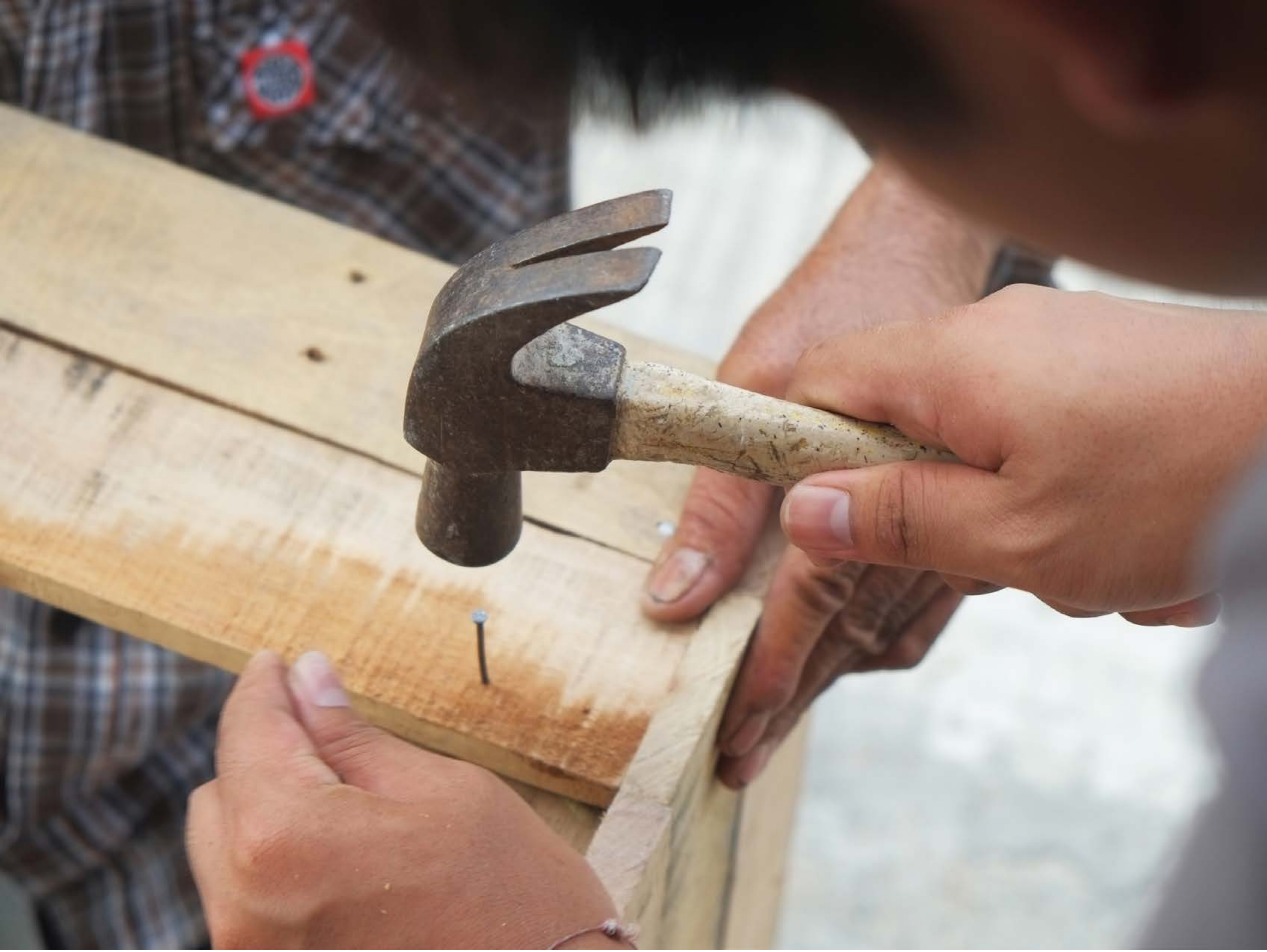

| Category | Craving (0-100) | Valence (1-9) | Arousal (1-9) | Typicality (0-100) | Relatedness | | | | | HSV | | |
|---|---|---|---|---|---|---|---|---|---|---|---|---|
| | | | | | Meth | Opioid | Both | Neither | MethToOpioid | Hue | Saturation | Value |
| Neutral tool hand complex | 20.29 (33.43) | 4.93 (1.514) | 2.32 (1.81) | 10.96 (19.09) | 0 | 0 | 0 | 1 | 0 | 0.209 (0.256) | 0.242 (0.175) | 0.606 (0.262) |

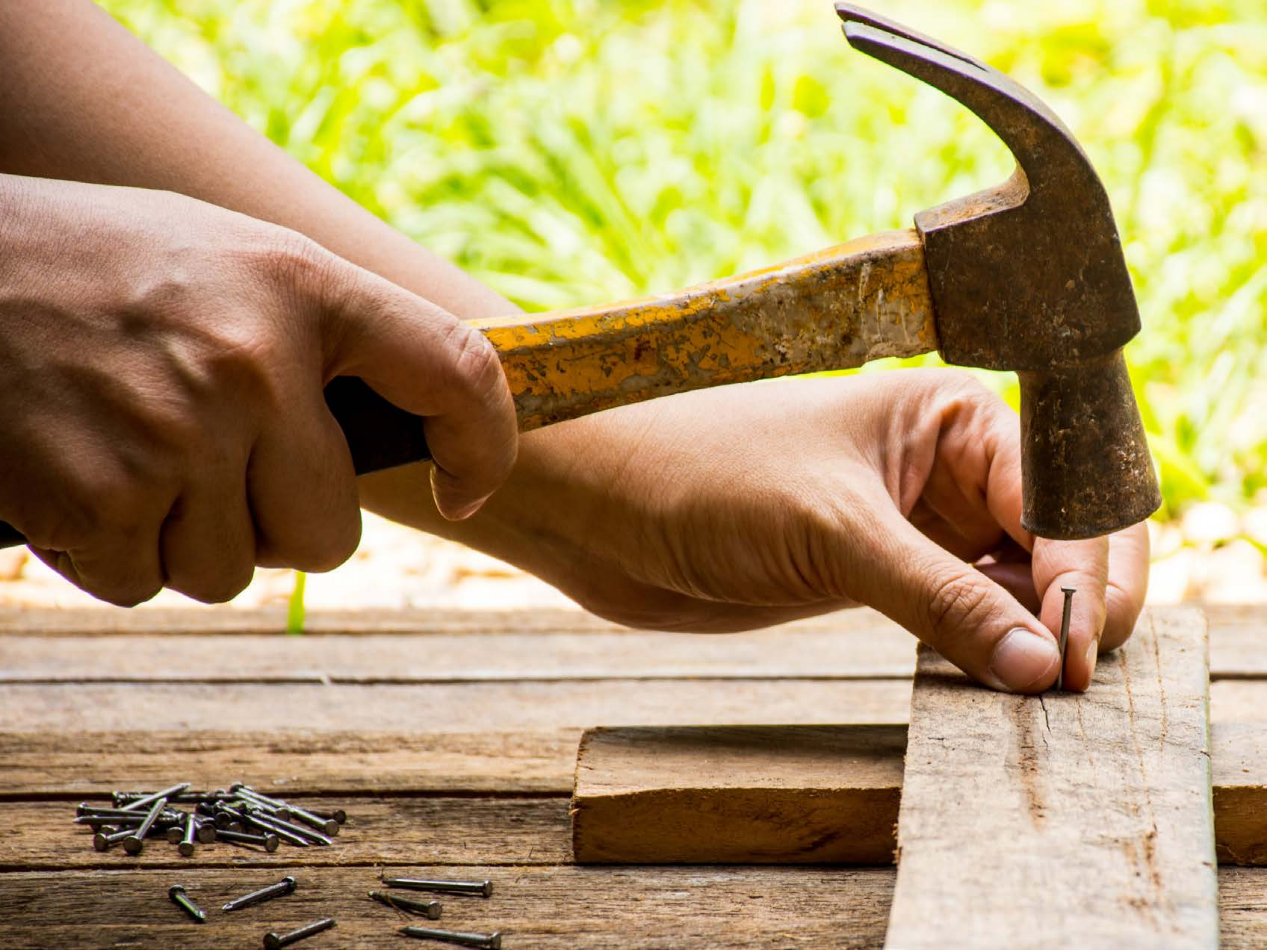

| Category | Craving (0-100) | Valence (1-9) | Arousal (1-9) | Typicality (0-100) | Relatedness | | | | | HSV | | |
|---|---|---|---|---|---|---|---|---|---|---|---|---|
| | | | | | Meth | Opioid | Both | Neither | MethToOpioid | Hue | Saturation | Value |
| Neutral tool hand complex | 10.5 (21.78) | 5.14 (1.976) | 2.39 (2.25) | 11.82 (21.87) | 0.036 | 0 | 0.036 | 0.929 | -0.036 | 0.107 (0.088) | 0.479 (0.301) | 0.69 (0.278) |

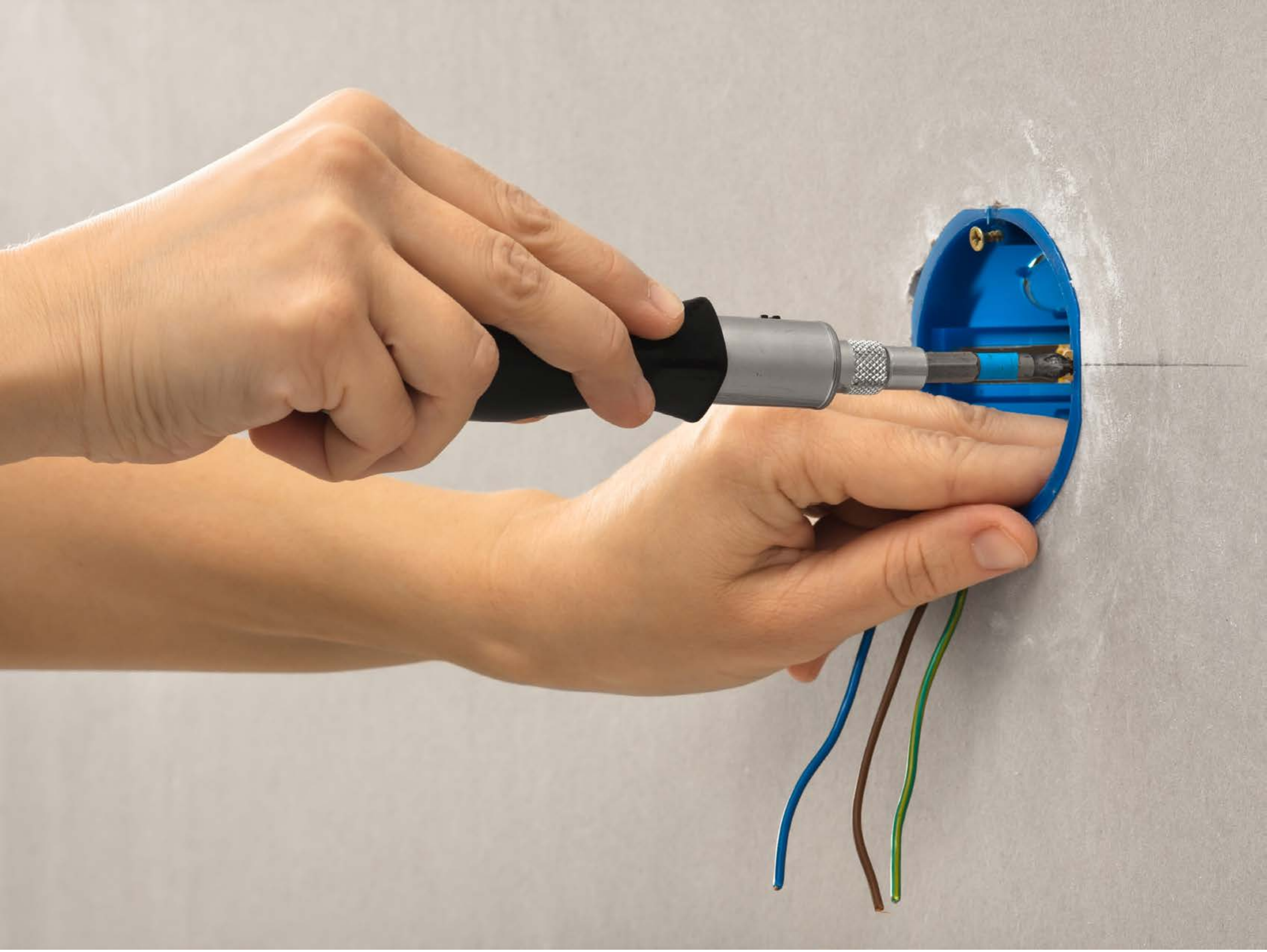

| Category | Craving (0-100) | Valence (1-9) | Arousal (1-9) | Typicality (0-100) | Relatedness | | | | | HSV | | |
|---|---|---|---|---|---|---|---|---|---|---|---|---|
| | | | | | Meth | Opioid | Both | Neither | MethToOpioid | Hue | Saturation | Value |
| Neutral tool hand complex | 15.81 (26.56) | 4.85 (0.77) | 2.81 (2) | 12.11 (18.99) | 0 | 0 | 0 | 1 | 0 | 0.095 (0.086) | 0.238 (0.209) | 0.743 (0.136) |

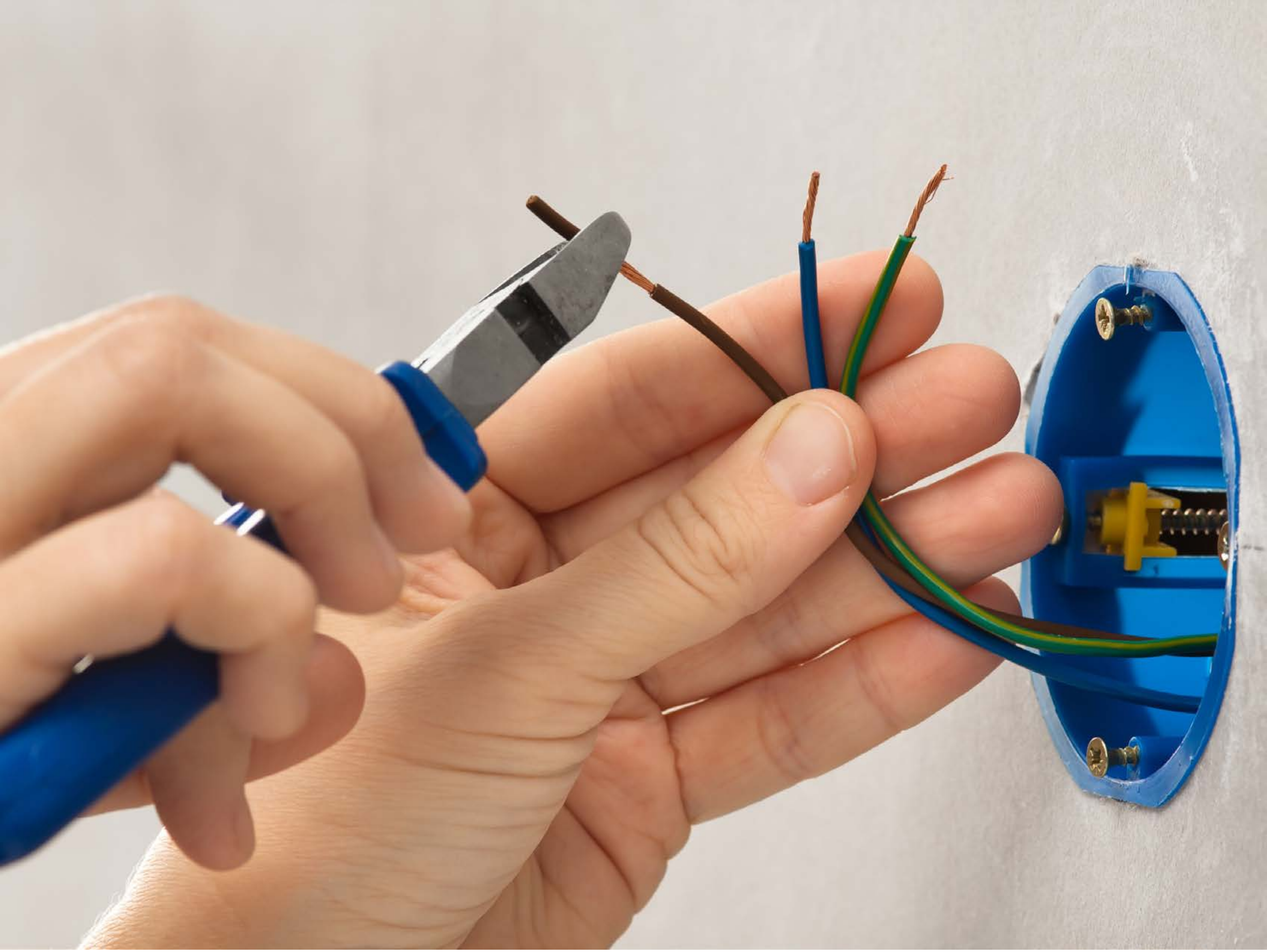

| Category | Craving (0-100) | Valence (1-9) | Arousal (1-9) | Typicality (0-100) | Relatedness | | | | | HSV | | |
|---|---|---|---|---|---|---|---|---|---|---|---|---|
| | | | | | Meth | Opioid | Both | Neither | MethToOpioid | Hue | Saturation | Value |
| Neutral tool hand complex | 13.21 (24.78) | 4.96 (1.453) | 2.57 (1.89) | 12.61 (19.45) | 0 | 0 | 0 | 1 | 0 | 0.128 (0.162) | 0.321 (0.28) | 0.738 (0.17) |

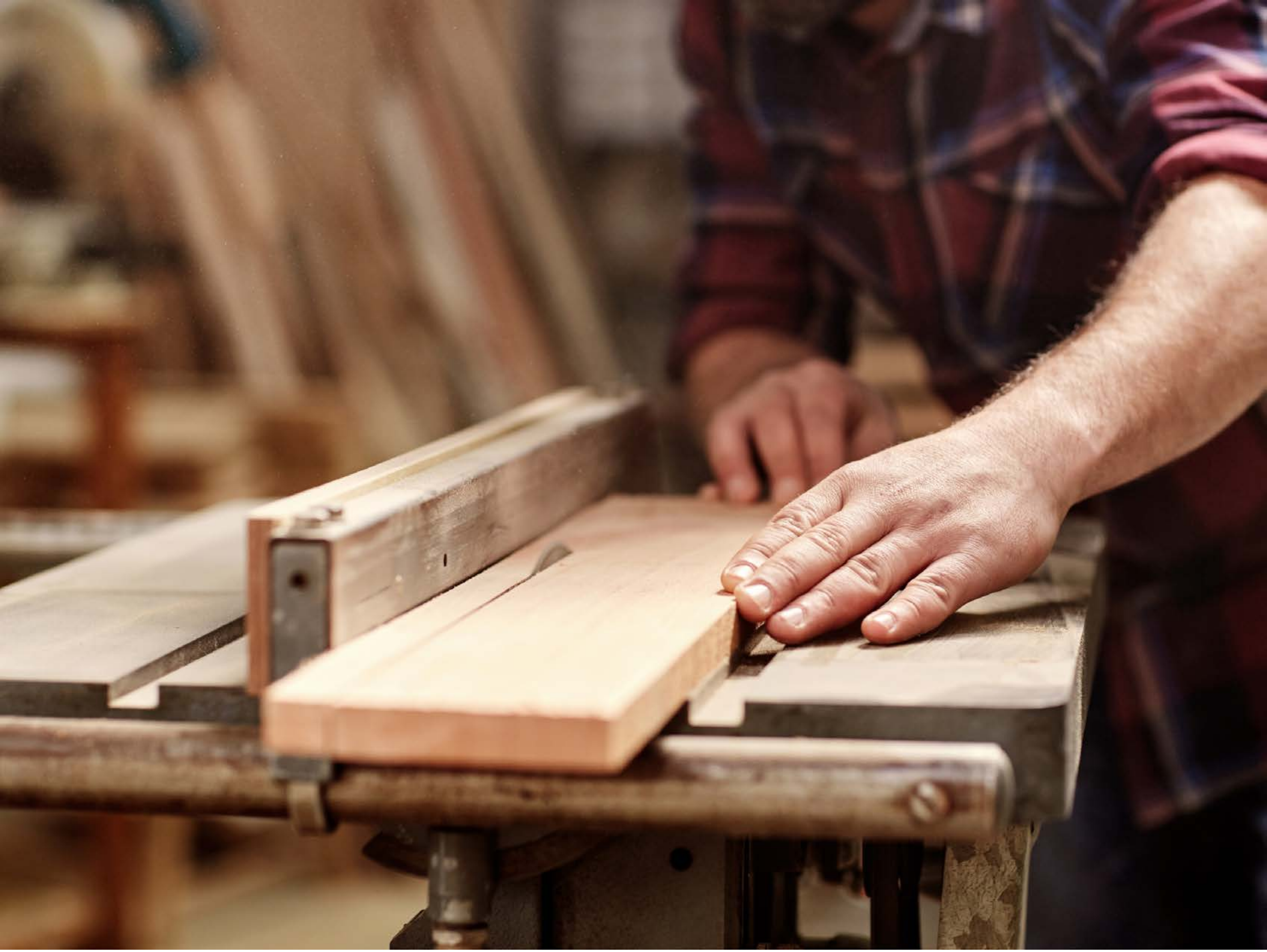

| Category | Craving (0-100) | Valence (1-9) | Arousal (1-9) | Typicality (0-100) | Relatedness | | | | | HSV | | |
|---|---|---|---|---|---|---|---|---|---|---|---|---|
| | | | | | Meth | Opioid | Both | Neither | MethToOpioid | Hue | Saturation | Value |
| Neutral tool hand complex | 6.89 (15.39) | 4.64 (1.521) | 2.11 (1.73) | 8.61 (15.55) | 0 | 0 | 0 | 1 | 0 | 0.152 (0.268) | 0.455 (0.197) | 0.479 (0.277) |

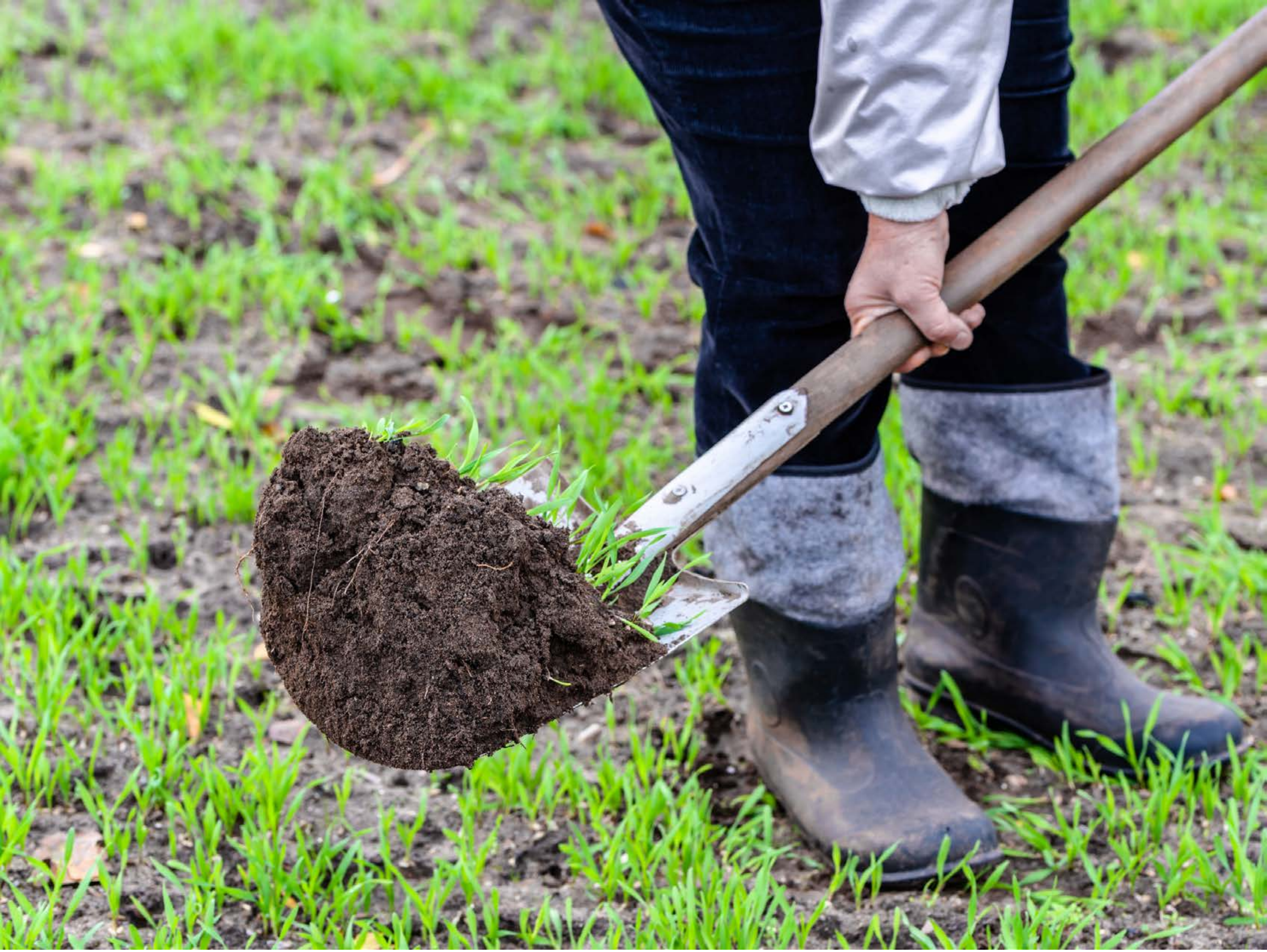

| Category | Craving (0-100) | Valence (1-9) | Arousal (1-9) | Typicality (0-100) | Relatedness | | | | | HSV | | |
|---|---|---|---|---|---|---|---|---|---|---|---|---|
| | | | | | Meth | Opioid | Both | Neither | MethToOpioid | Hue | Saturation | Value |
| Neutral tool hand complex | 5.54 (13.07) | 4.79 (1.792) | 2.21 (1.75) | 6.61 (13.24) | 0 | 0 | 0 | 1 | 0 | 0.355 (0.282) | 0.349 (0.248) | 0.578 (0.252) |

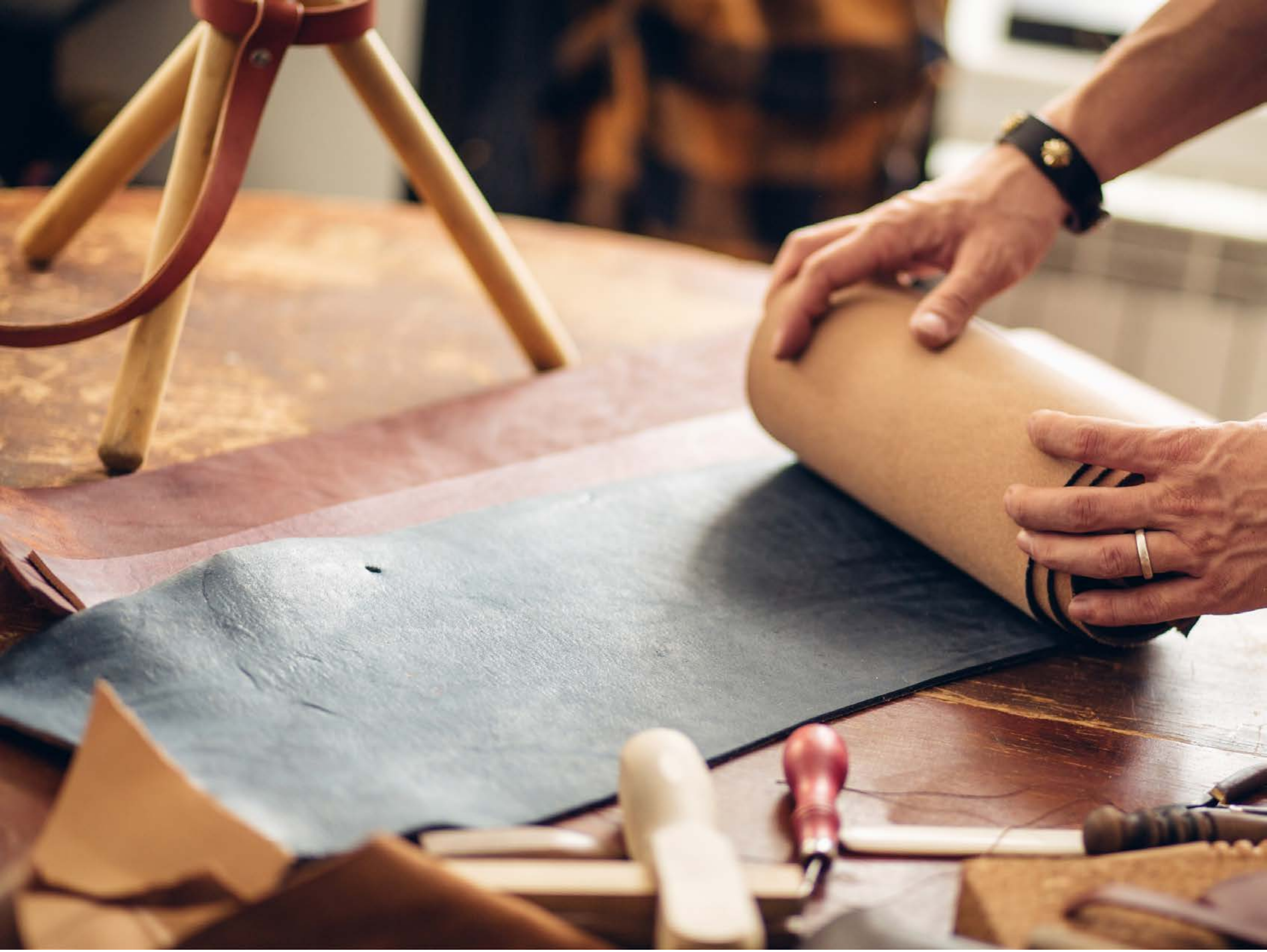

| Category | Craving (0-100) | Valence (1-9) | Arousal (1-9) | Typicality (0-100) | Relatedness | | | | | HSV | | |
|---|---|---|---|---|---|---|---|---|---|---|---|---|
| | | | | | Meth | Opioid | Both | Neither | MethToOpioid | Hue | Saturation | Value |
| Neutral tool hand complex | 9.93 (19.89) | 4.78 (1.251) | 2.7 (2.13) | 9.37 (17.55) | 0 | 0 | 0 | 1 | 0 | 0.202 (0.253) | 0.337 (0.226) | 0.641 (0.261) |

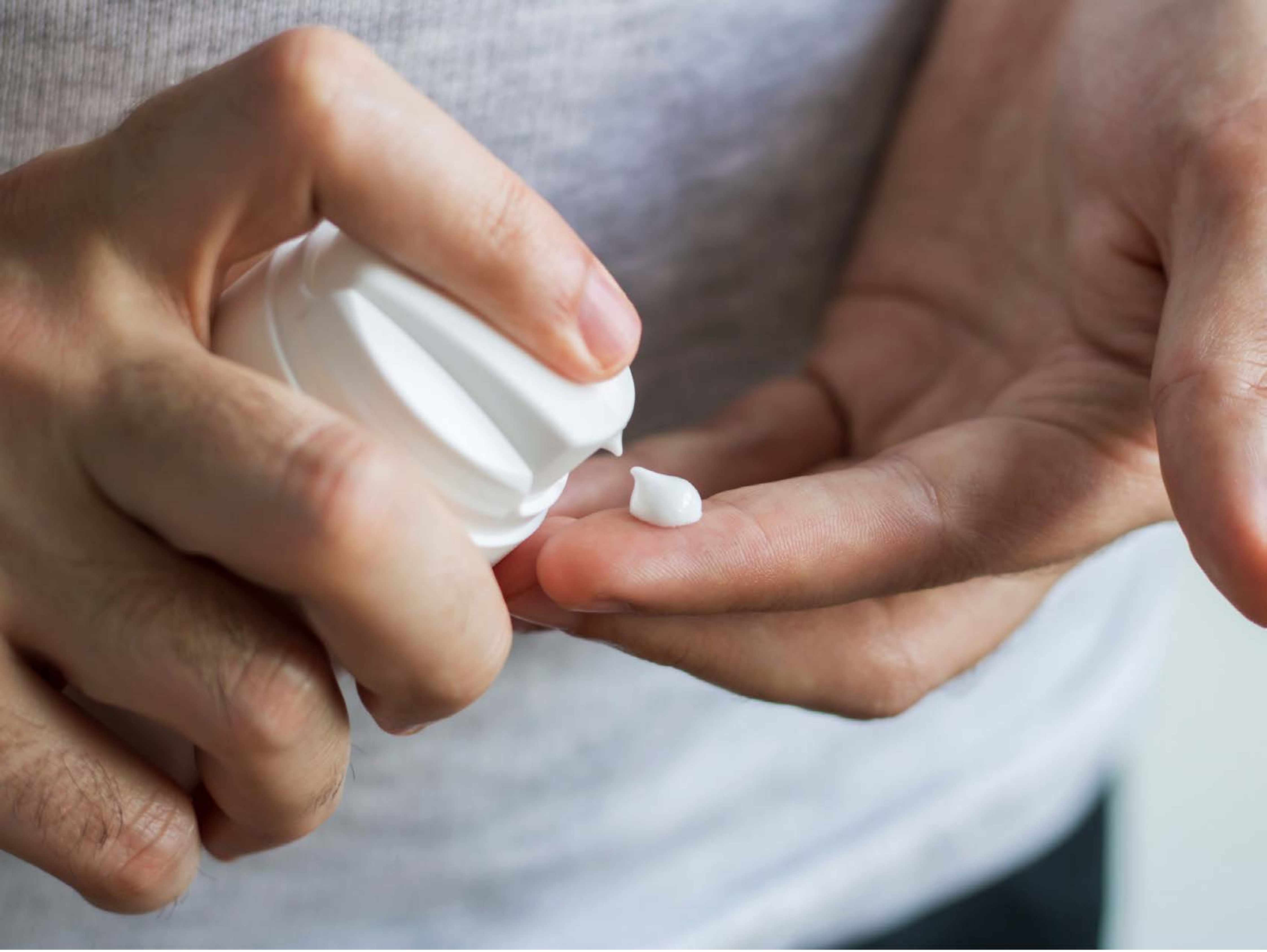

| Category | Craving (0-100) | Valence (1-9) | Arousal (1-9) | Typicality (0-100) | Relatedness | | | | | HSV | | |
|---|---|---|---|---|---|---|---|---|---|---|---|---|
| | | | | | Meth | Opioid | Both | Neither | MethToOpioid | Hue | Saturation | Value |
| Neutral tool hand complex | 8.85 (17.64) | 4.78 (1.155) | 2.33 (1.92) | 16.37 (24.97) | 0 | 0 | 0 | 1 | 0 | 0.257 (0.275) | 0.274 (0.178) | 0.601 (0.212) |

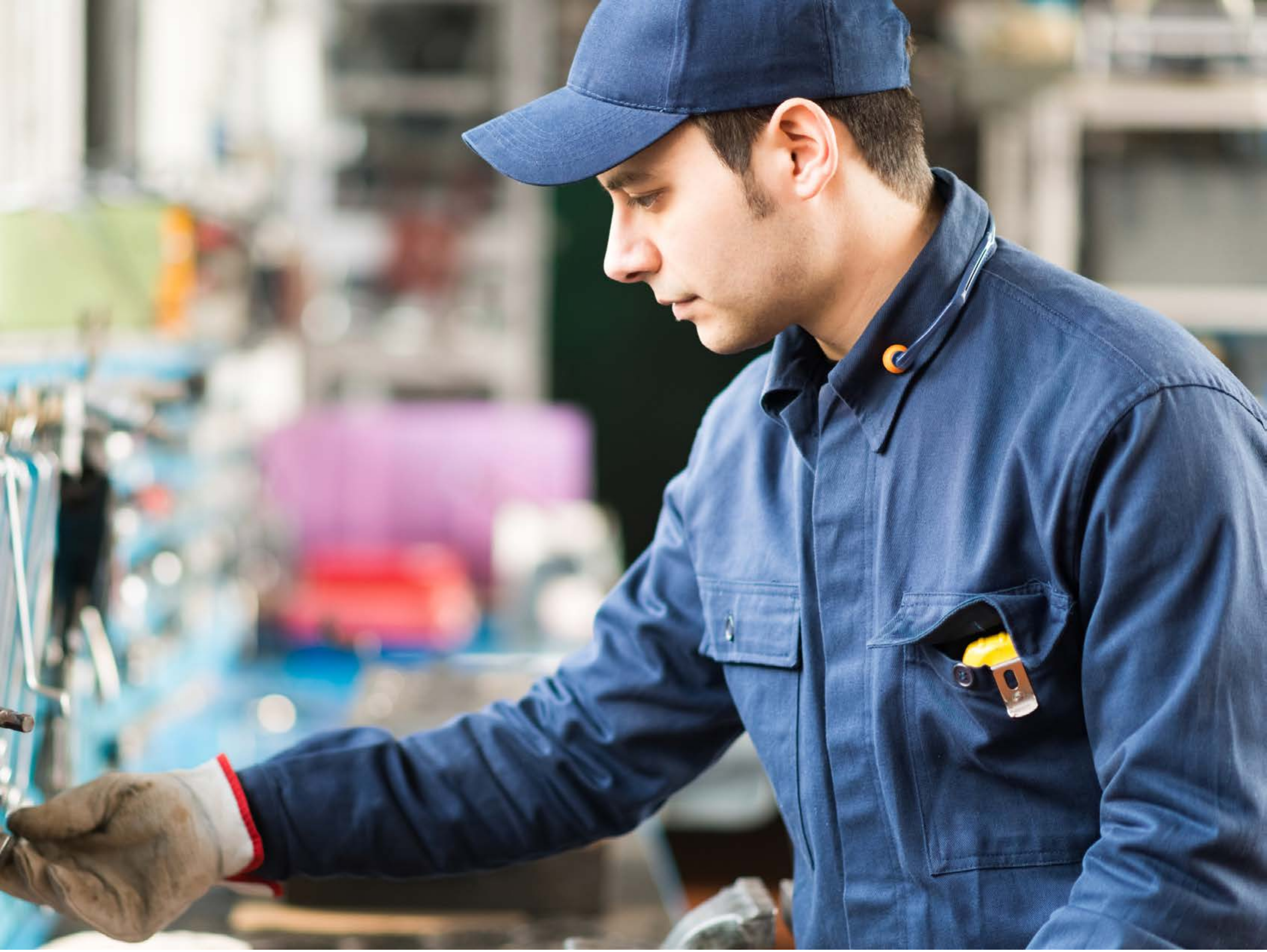

| Category | Craving (0-100) | Valence (1-9) | Arousal (1-9) | Typicality (0-100) | Relatedness | | | | | HSV | | |
|---|---|---|---|---|---|---|---|---|---|---|---|---|
| | | | | | Meth | Opioid | Both | Neither | MethToOpioid | Hue | Saturation | Value |
| Neutral tool face | 14.32 (25.2) | 4.93 (1.386) | 2.46 (1.99) | 14.89 (24.89) | 0 | 0 | 0 | 1 | 0 | 0.393 (0.265) | 0.305 (0.195) | 0.591 (0.259) |

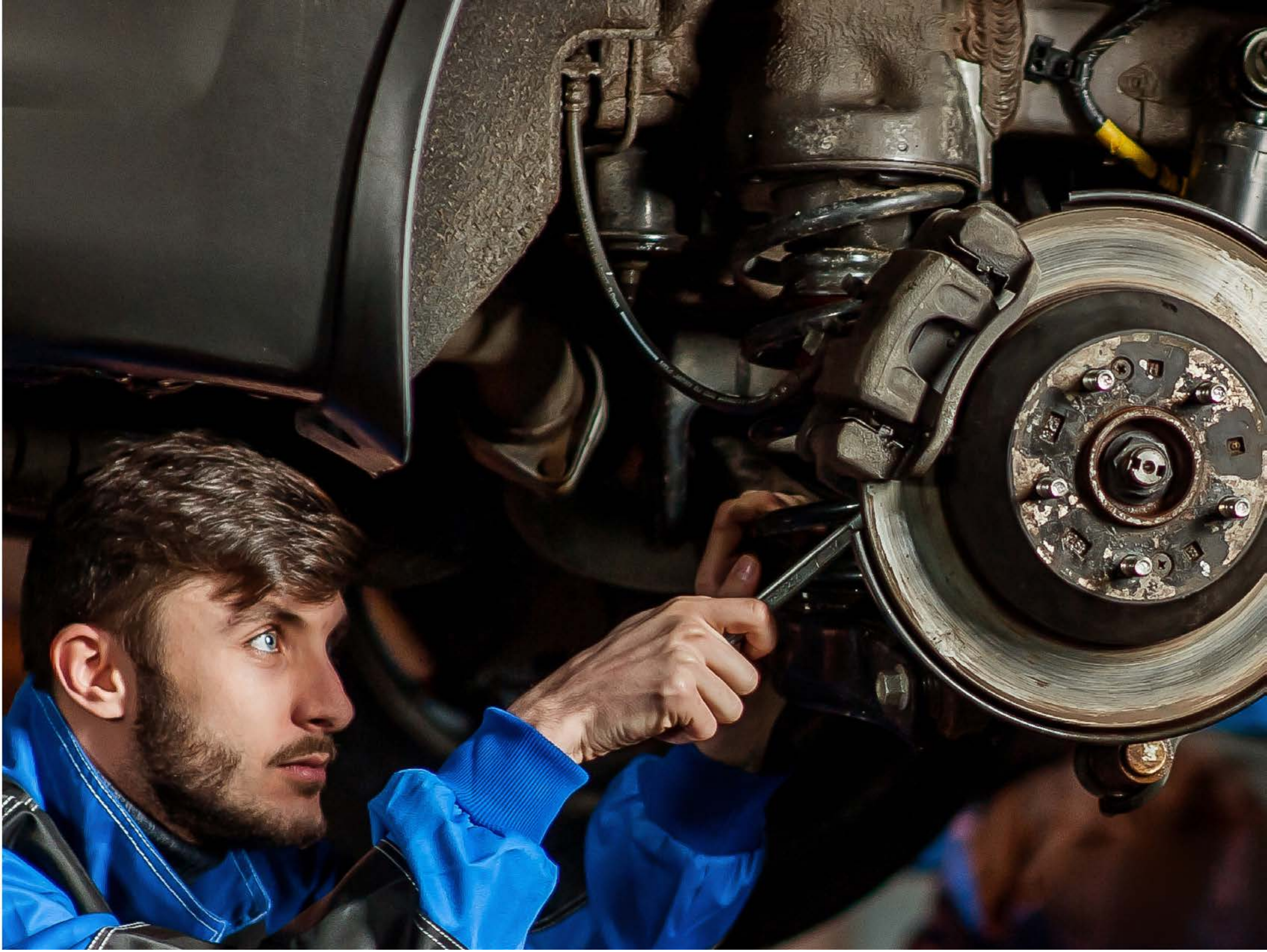

| Category | Craving (0-100) | Valence (1-9) | Arousal (1-9) | Typicality (0-100) | Relatedness | | | | | HSV | | |
|---|---|---|---|---|---|---|---|---|---|---|---|---|
| | | | | | Meth | Opioid | Both | Neither | MethToOpioid | Hue | Saturation | Value |
| Neutral tool face | 11.96 (19.53) | 4.78 (1.155) | 2.89 (2.24) | 15.07 (21.05) | 0.037 | 0 | 0 | 0.963 | -0.037 | 0.197 (0.264) | 0.412 (0.306) | 0.263 (0.241) |

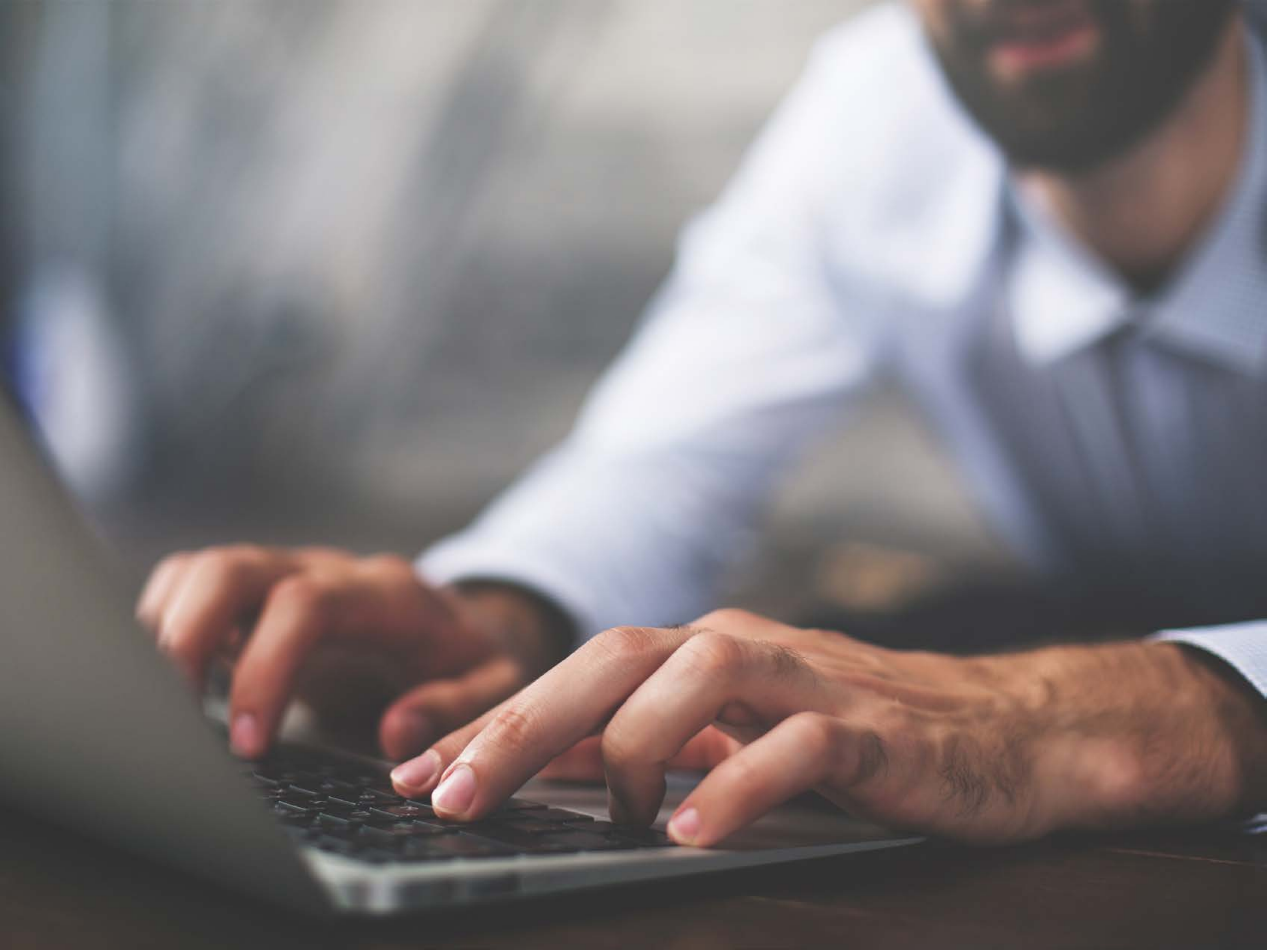

| Category | Craving (0-100) | Valence (1-9) | Arousal (1-9) | Typicality (0-100) | Relatedness | | | | | HSV | | |
|---|---|---|---|---|---|---|---|---|---|---|---|---|
| | | | | | Meth | Opioid | Both | Neither | MethToOpioid | Hue | Saturation | Value |
| Neutral tool face | 8.71 (17.49) | 5.04 (1.835) | 2.54 (2.06) | 10.82 (19.02) | 0.036 | 0 | 0 | 0.964 | -0.036 | 0.356 (0.313) | 0.127 (0.123) | 0.528 (0.222) |

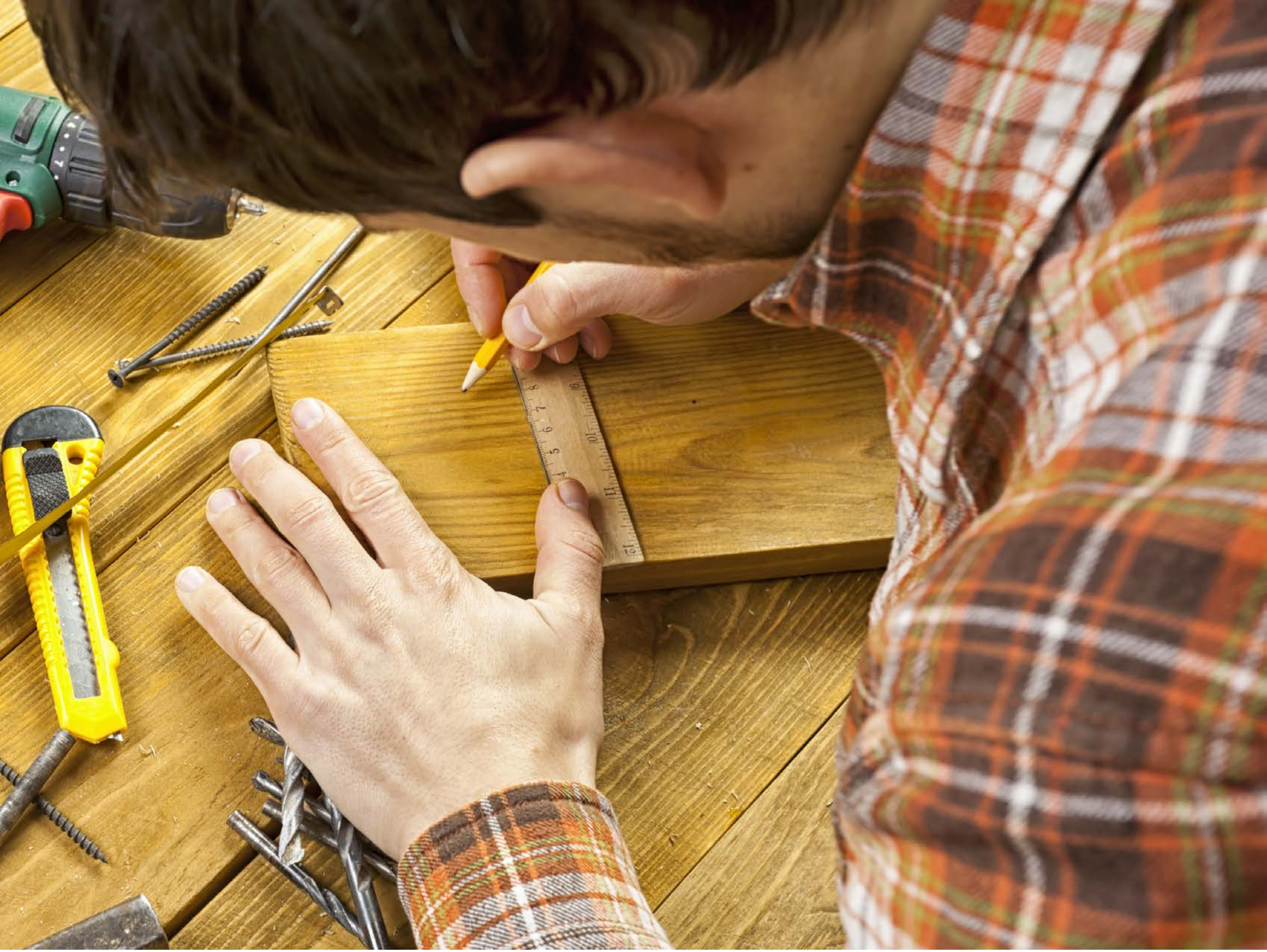

| Category | Craving (0-100) | Valence (1-9) | Arousal (1-9) | Typicality (0-100) | Relatedness | | | | | HSV | | |
|---|---|---|---|---|---|---|---|---|---|---|---|---|
| | | | | | Meth | Opioid | Both | Neither | MethToOpioid | Hue | Saturation | Value |
| Neutral tool face | 9.33 (17.69) | 4.85 (1.231) | 2.52 (1.95) | 11.26 (17.49) | 0 | 0 | 0 | 1 | 0 | 0.088 (0.065) | 0.522 (0.222) | 0.621 (0.241) |

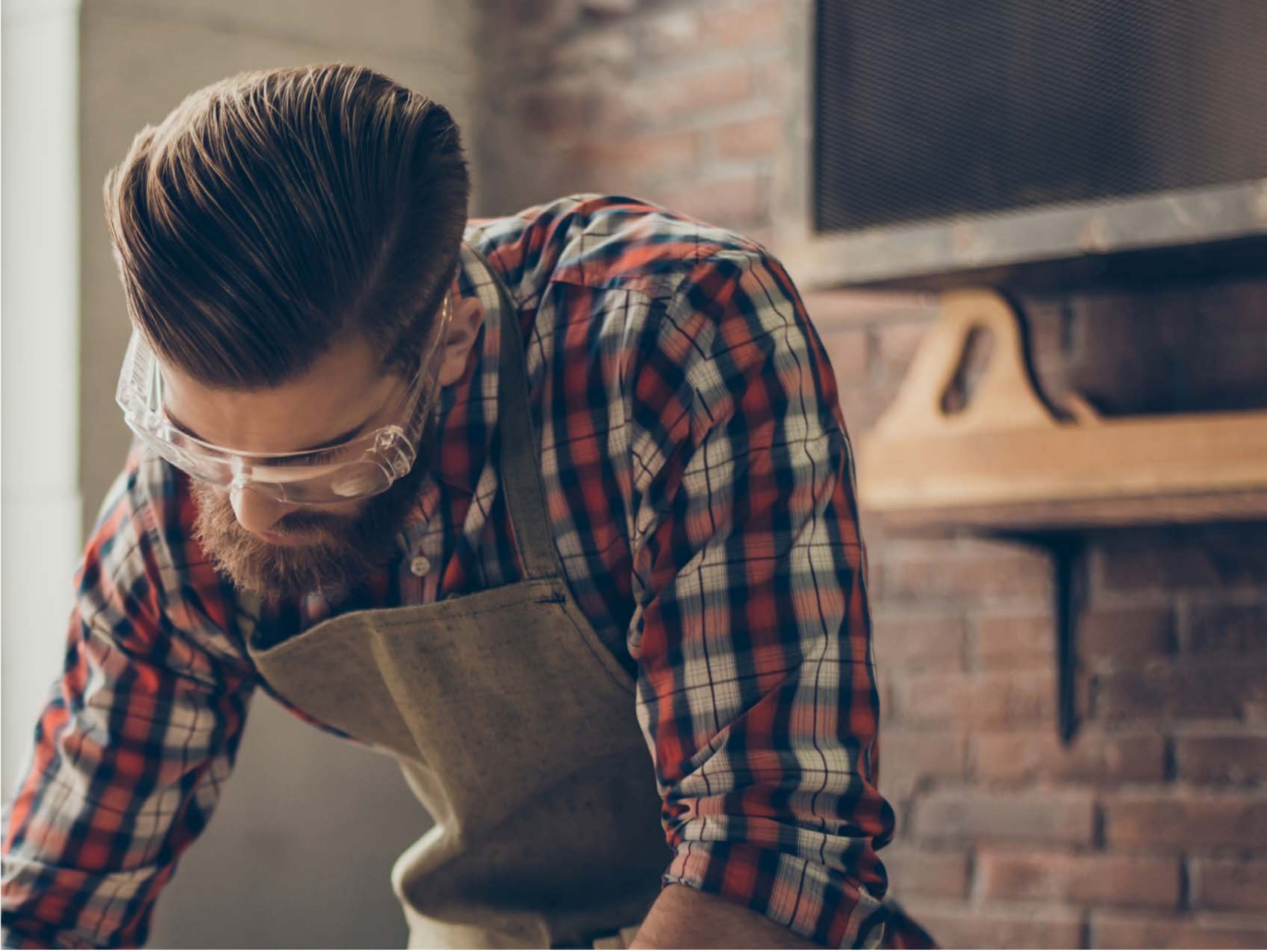

| Category | Craving (0-100) | Valence (1-9) | Arousal (1-9) | Typicality (0-100) | Relatedness | | | | | HSV | | |
|---|---|---|---|---|---|---|---|---|---|---|---|---|
| | | | | | Meth | Opioid | Both | Neither | MethToOpioid | Hue | Saturation | Value |
| Neutral tool face | 8.32 (20.67) | 4.86 (1.671) | 2.14 (1.78) | 10.86 (22.29) | 0 | 0 | 0 | 1 | 0 | 0.343 (0.326) | 0.291 (0.195) | 0.432 (0.203) |

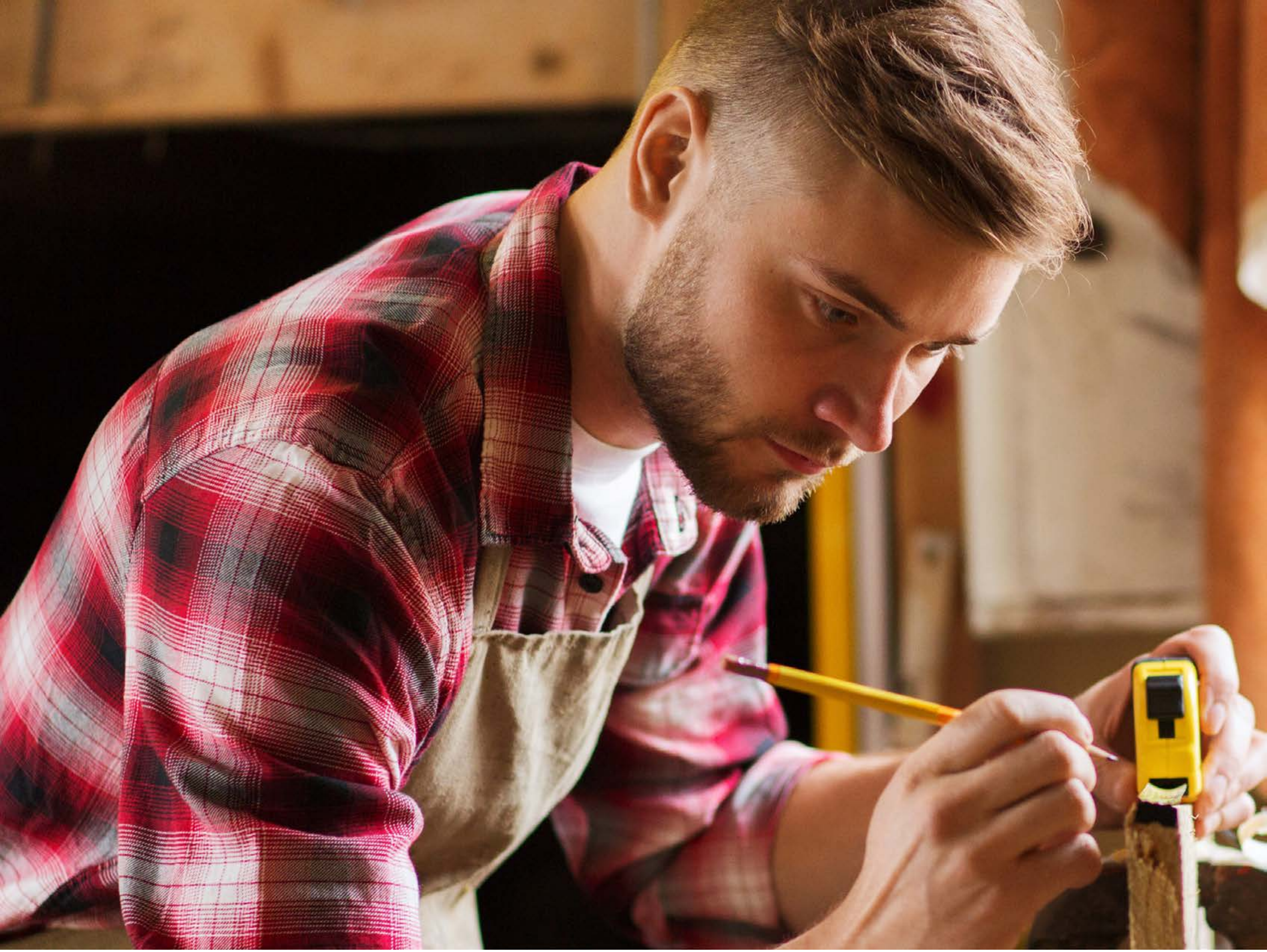

| Category | Craving (0-100) | Valence (1-9) | Arousal (1-9) | Typicality (0-100) | Relatedness | | | | | HSV | | |
|---|---|---|---|---|---|---|---|---|---|---|---|---|
| | | | | | Meth | Opioid | Both | Neither | MethToOpioid | Hue | Saturation | Value |
| Neutral tool face | 8.85 (17.73 | 4.85 (1.231 | 2.67 (1.98 | 9.44 (17.56 | 0 | 0 | 0 | 1 | 0 | 0.254 (0.374 | 0.559 (0.233 | 0.512 (0.272 |

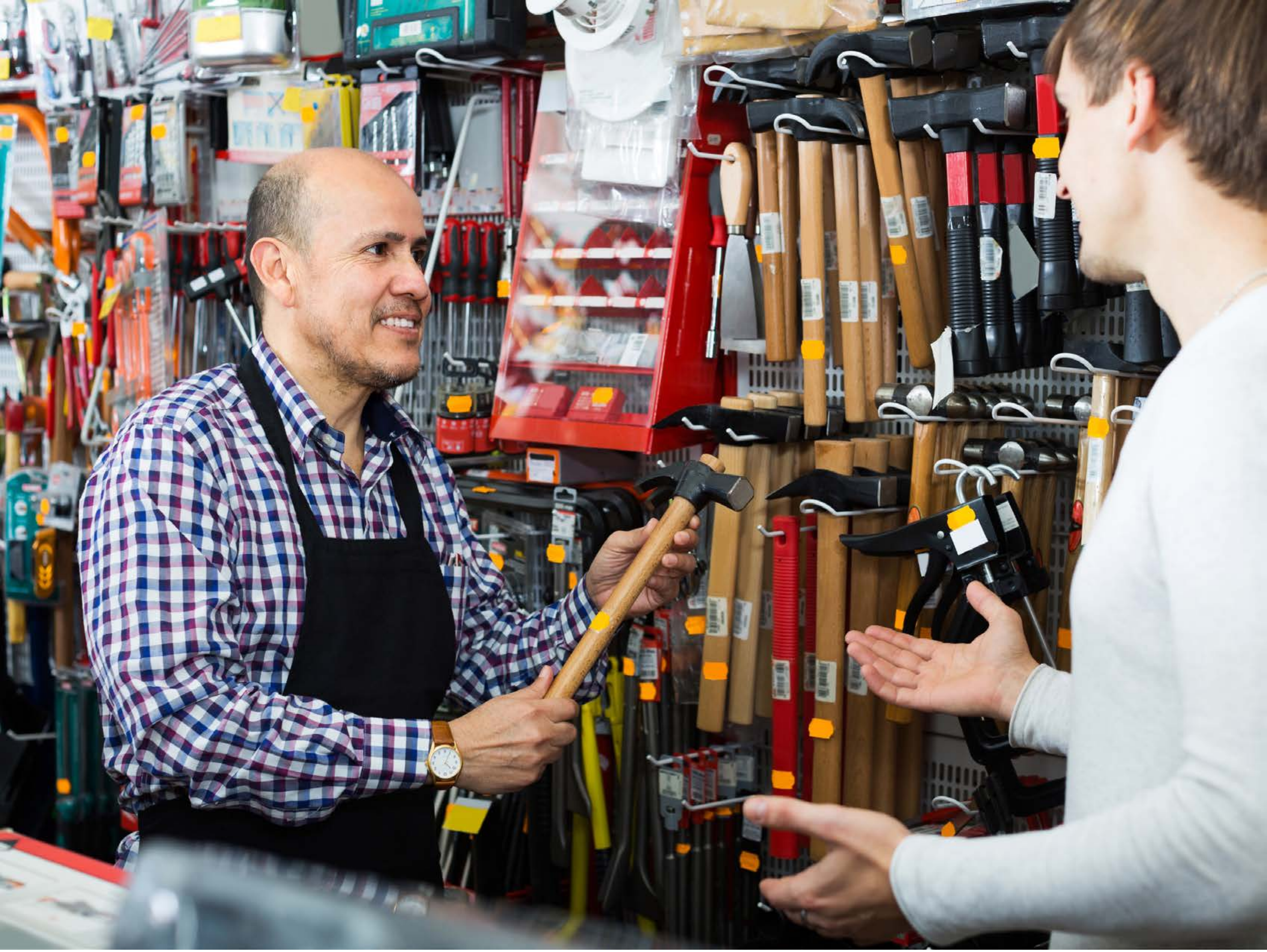

| Category | Craving (0-100) | Valence (1-9) | Arousal (1-9) | Typicality (0-100) | Relatedness | | | | | HSV | | |
|---|---|---|---|---|---|---|---|---|---|---|---|---|
| | | | | | Meth | Opioid | Both | Neither | MethToOpioid | Hue | Saturation | Value |
| Neutral tool face | 12.52 (19.21) | 5.04 (1.055) | 2.74 (1.95) | 13.11 (19.2) | 0 | 0 | 0 | 1 | 0 | 0.404 (0.304) | 0.294 (0.258) | 0.53 (0.313) |

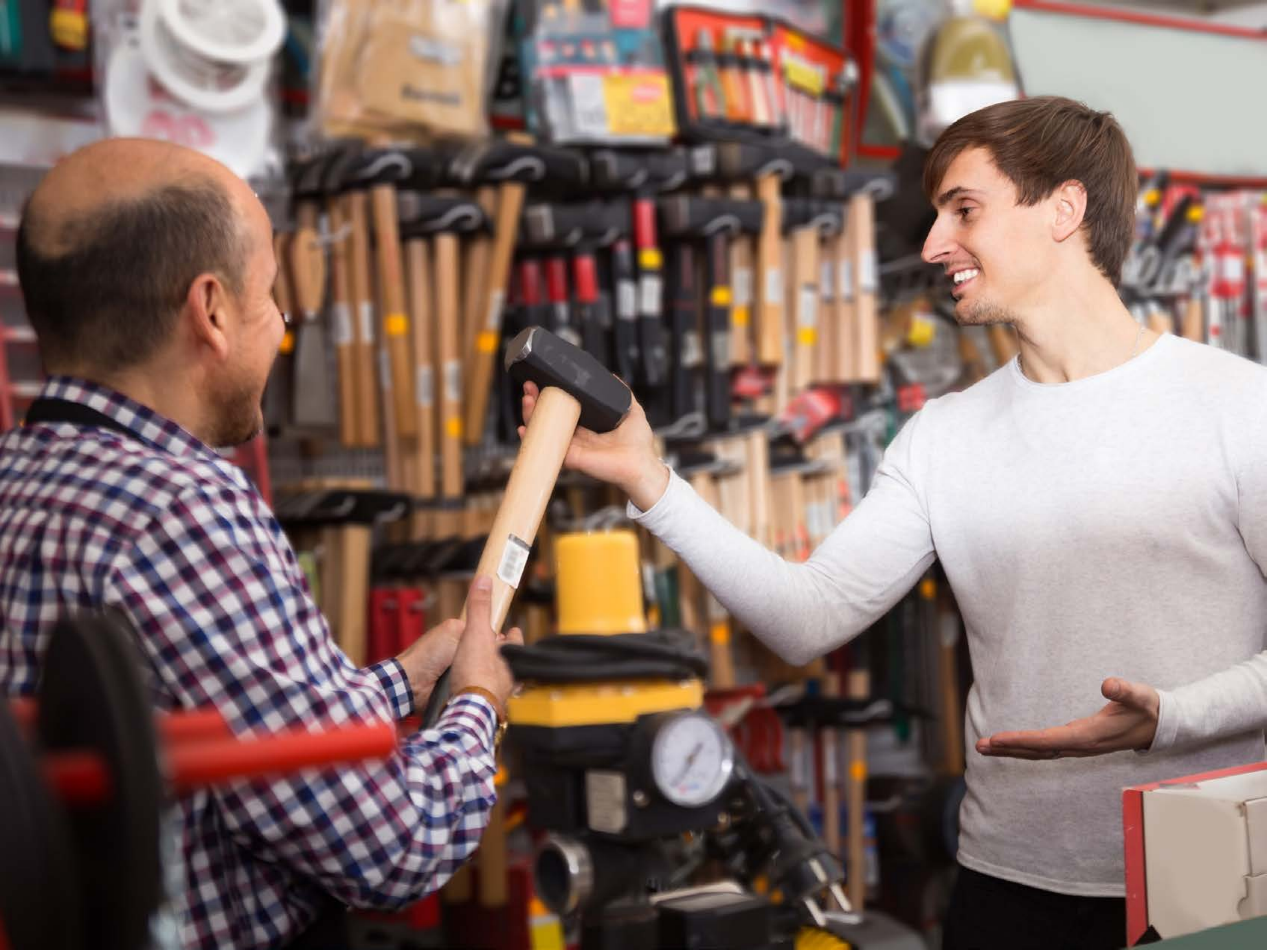

| Category | Craving (0-100) | Valence (1-9) | Arousal (1-9) | Typicality (0-100) | Relatedness | | | | | HSV | | |
|---|---|---|---|---|---|---|---|---|---|---|---|---|
| | | | | | Meth | Opioid | Both | Neither | MethToOpioid | Hue | Saturation | Value |
| Neutral tool face | 9.11 (17.85) | 4.81 (1.178) | 2.63 (2) | 9.89 (17.55) | 0 | 0 | 0 | 1 | 0 | 0.389 (0.402) | 0.279 (0.22) | 0.516 (0.287) |

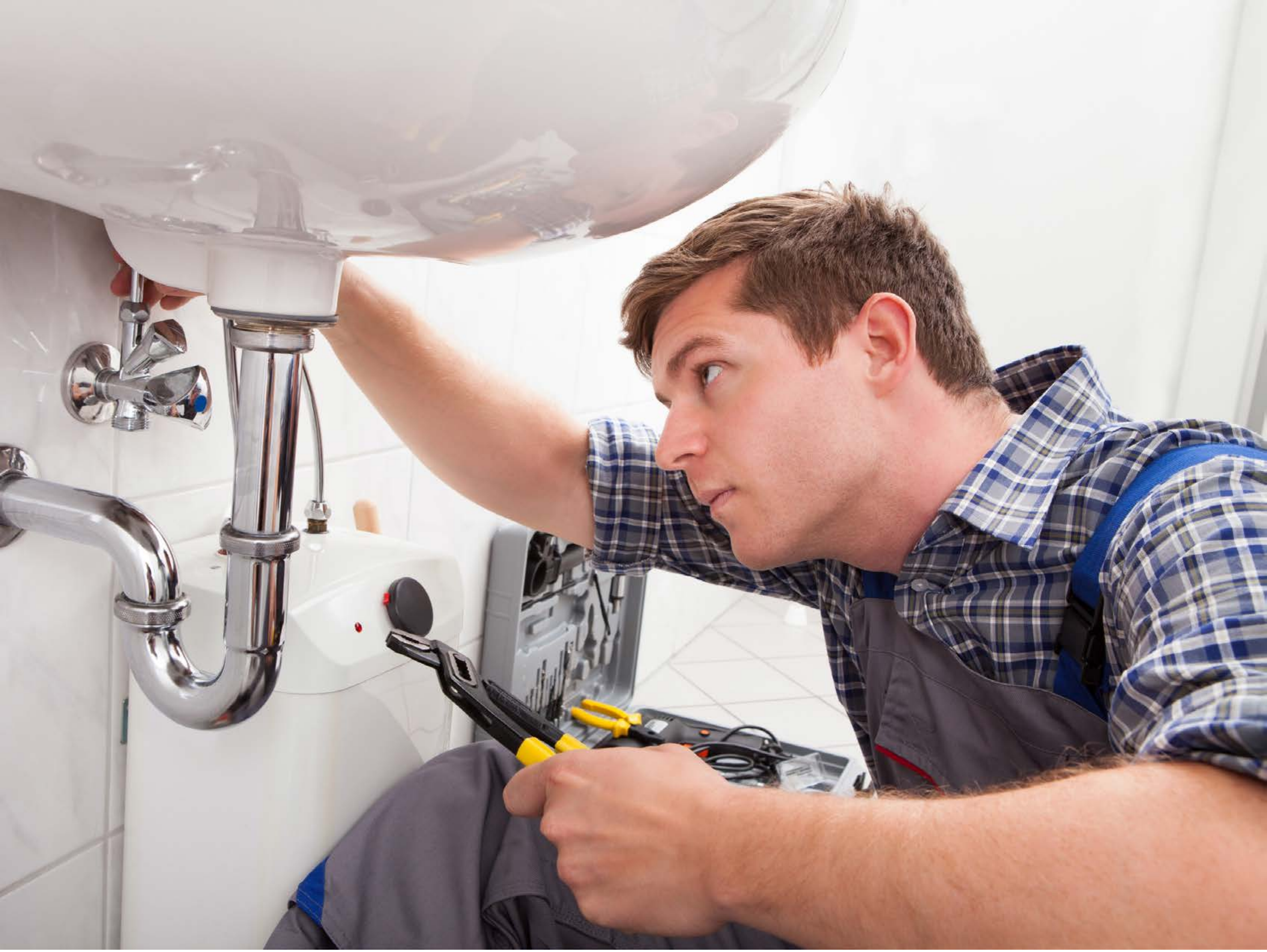

| Category | Craving (0-100) | Valence (1-9) | Arousal (1-9) | Typicality (0-100) | Relatedness | | | | | HSV | | |
|---|---|---|---|---|---|---|---|---|---|---|---|---|
| | | | | | Meth | Opioid | Both | Neither | MethToOpioid | Hue | Saturation | Value |
| Neutral tool face | 12.5 (27.28) | 4.93 (1.741) | 2.29 (1.86) | 13.71 (25.55) | 0 | 0 | 0 | 1 | 0 | 0.248 (0.345) | 0.152 (0.176) | 0.726 (0.229) |

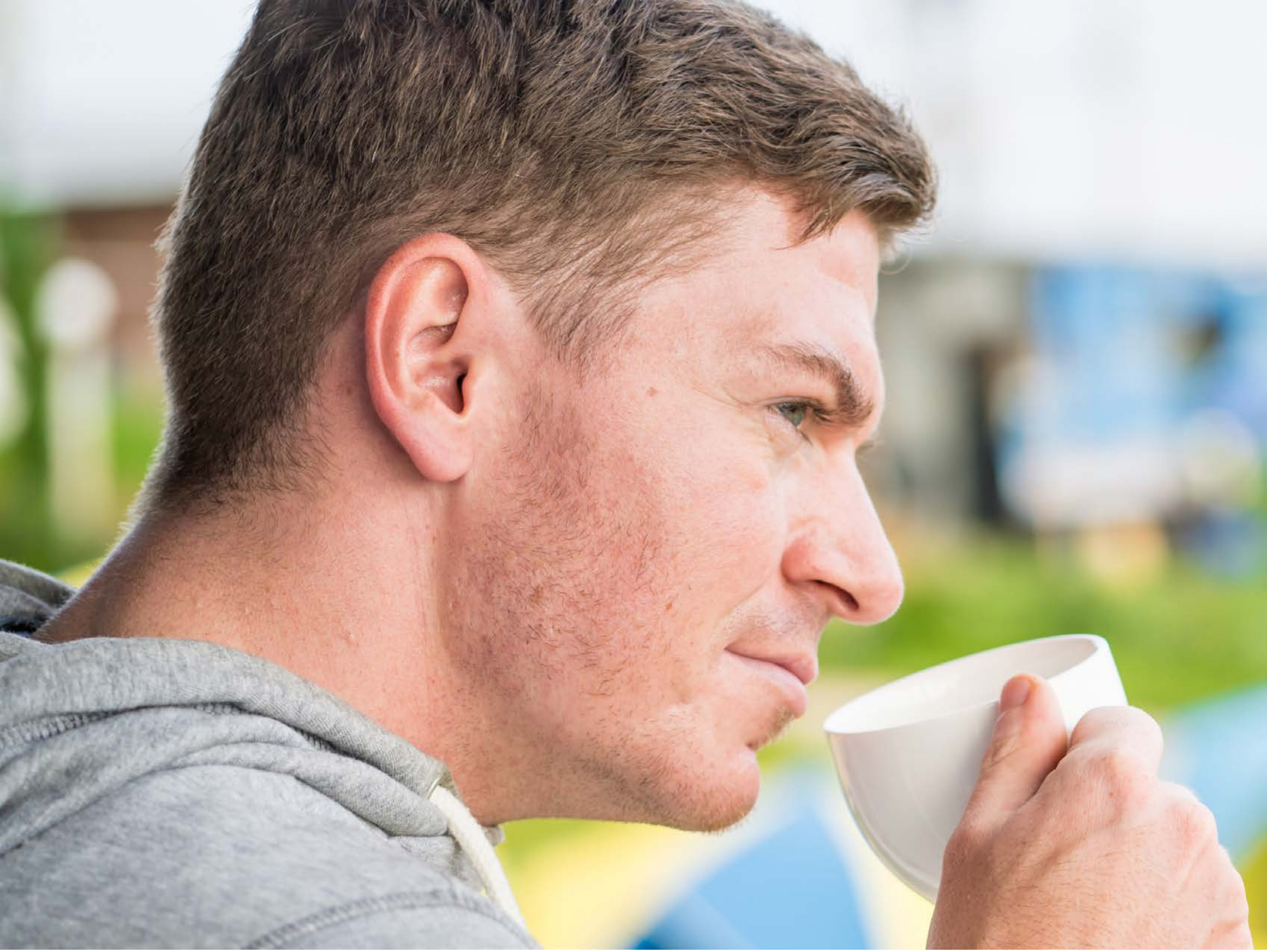

| Category | Craving (0-100) | Valence (1-9) | Arousal (1-9) | Typicality (0-100) | Relatedness | | | | | HSV | | |
|---|---|---|---|---|---|---|---|---|---|---|---|---|
| | | | | | Meth | Opioid | Both | Neither | MethToOpioid | Hue | Saturation | Value |
| Neutral tool face | 10.61 (23.34) | 4.79 (1.729) | 2.25 (1.82) | 11.25 (23.15) | 0 | 0 | 0 | 1 | 0 | 0.238 (0.249) | 0.244 (0.172) | 0.741 (0.222) |

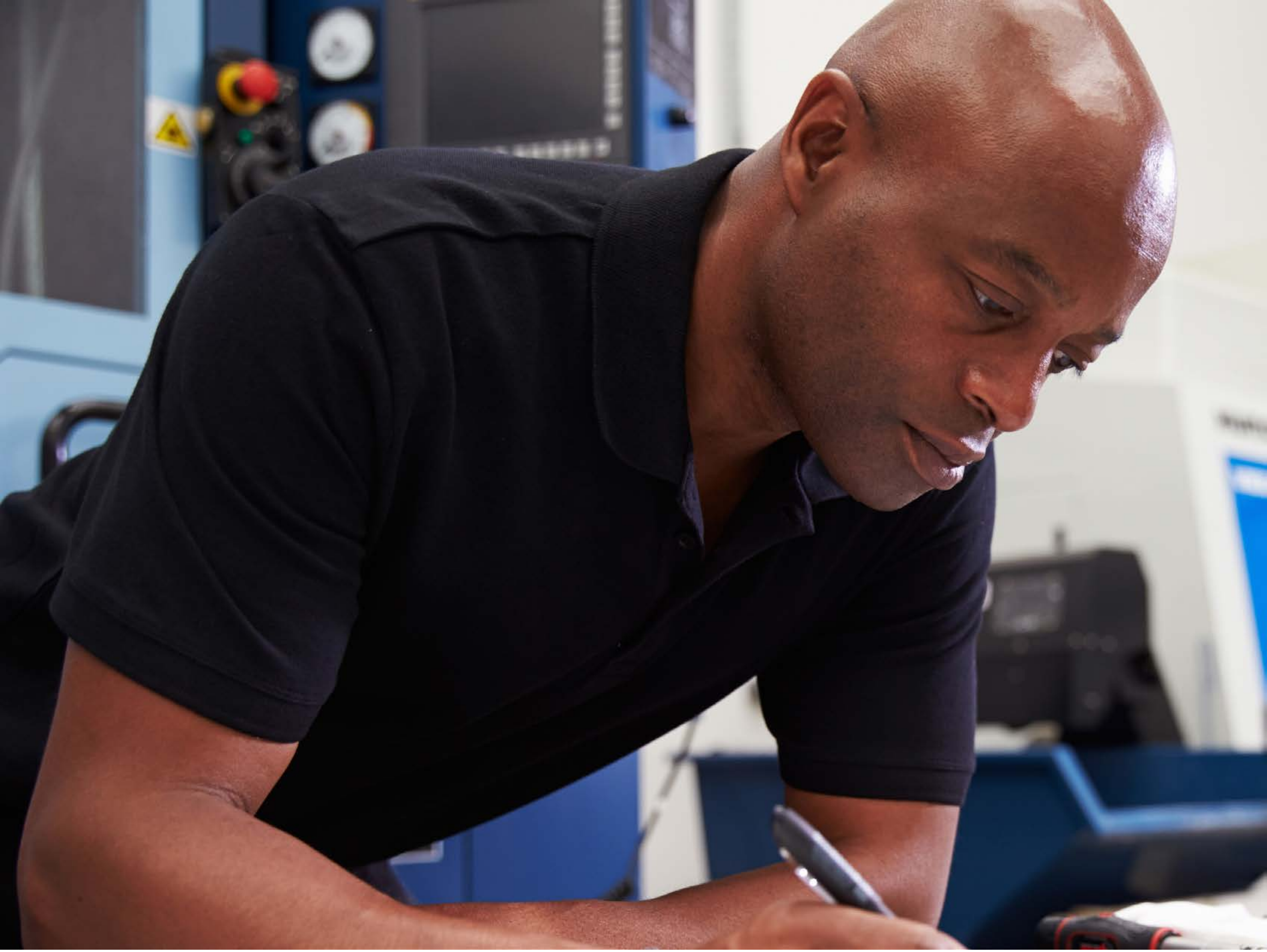

| Category | Craving (0-100) | Valence (1-9) | Arousal (1-9) | Typicality (0-100) | Relatedness | | | | | HSV | | |
|---|---|---|---|---|---|---|---|---|---|---|---|---|
| | | | | | Meth | Opioid | Both | Neither | MethToOpioid | Hue | Saturation | Value |
| Neutral tool face | 12.59 (22.71) | 4.93 (0.874) | 2.7 (2.13) | 11.37 (22.41) | 0 | 0.037 | 0 | 0.963 | 0.037 | 0.456 (0.342) | 0.307 (0.224) | 0.357 (0.296) |

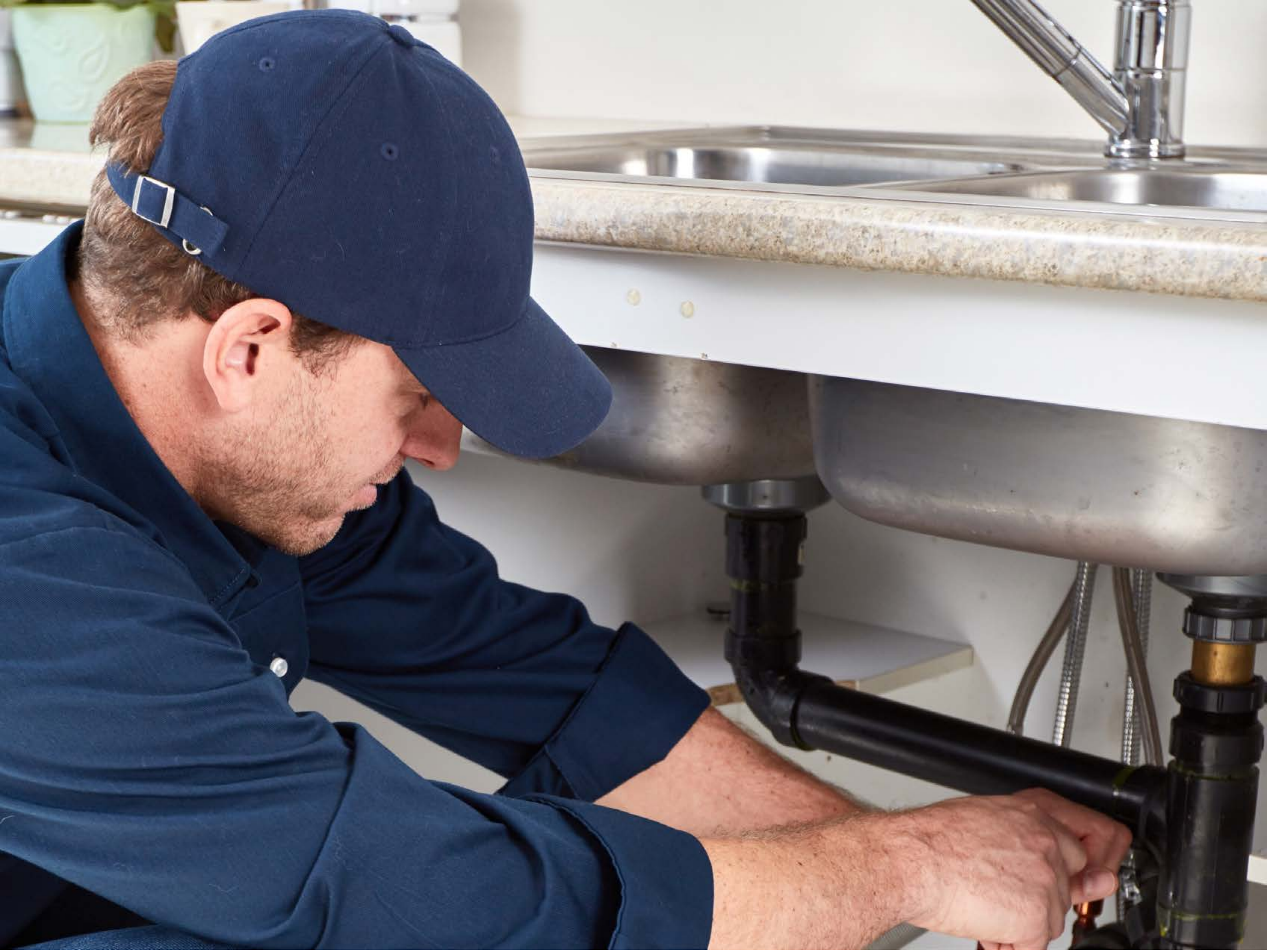

| Category | Craving (0-100) | Valence (1-9) | Arousal (1-9) | Typicality (0-100) | Relatedness | | | | | HSV | | |
|---|---|---|---|---|---|---|---|---|---|---|---|---|
| | | | | | Meth | Opioid | Both | Neither | MethToOpioid | Hue | Saturation | Value |
| Neutral tool face | 7.56 (15.85) | 4.7 (1.068) | 2.56 (1.93) | 11.19 (17.57) | 0 | 0 | 0 | 1 | 0 | 0.374 (0.306) | 0.286 (0.232) | 0.528 (0.274) |

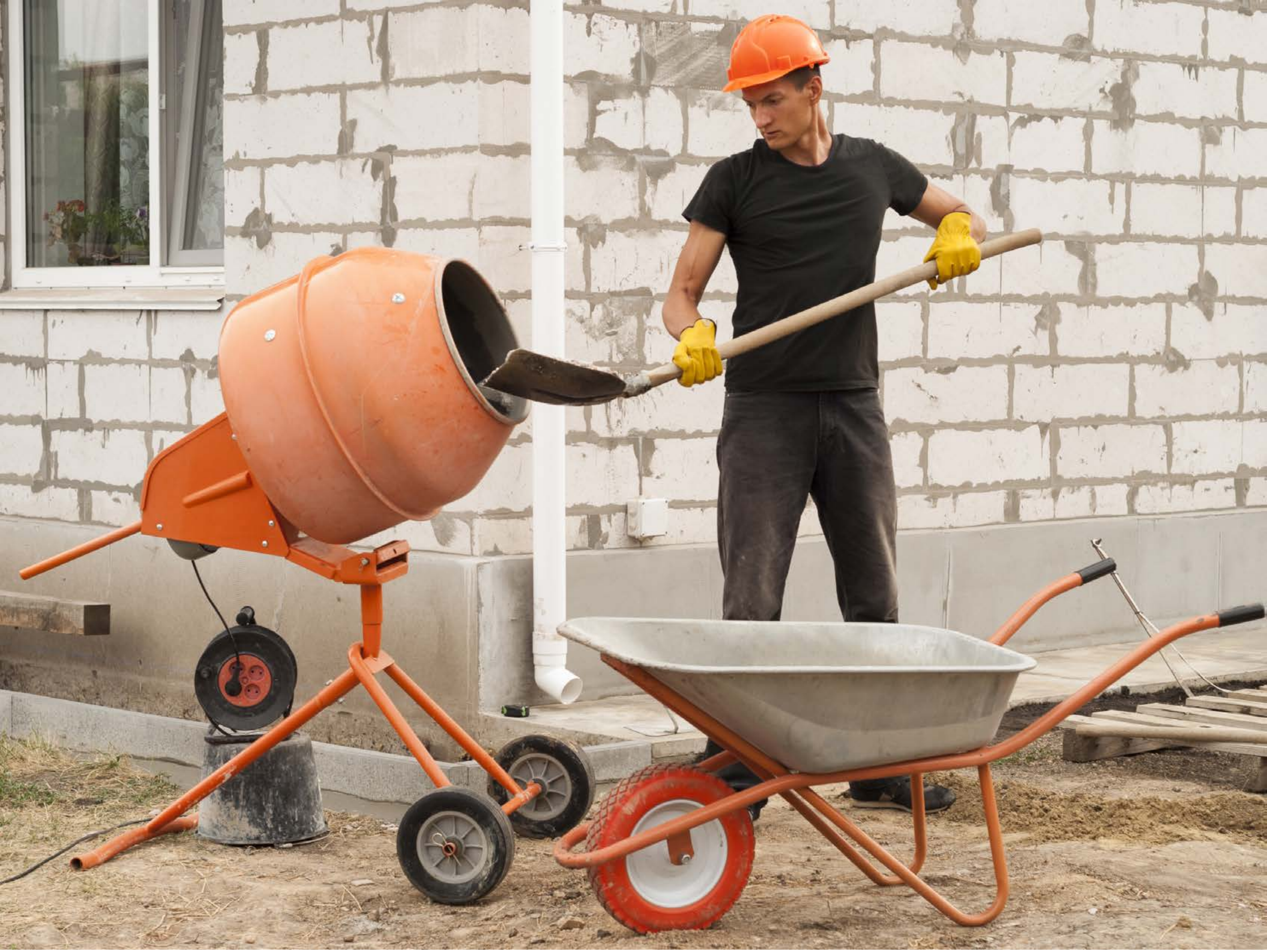

| Category | Craving (0-100) | Valence (1-9) | Arousal (1-9) | Typicality (0-100) | Relatedness | | | | | HSV | | |
|---|---|---|---|---|---|---|---|---|---|---|---|---|
| | | | | | Meth | Opioid | Both | Neither | MethToOpioid | Hue | Saturation | Value |
| Neutral tool face | 10 (19.09) | 4.96 (1.551) | 2.79 (2.27) | 9.93 (19.33) | 0.071 | 0 | 0 | 0.929 | -0.071 | 0.087 (0.048) | 0.224 (0.197) | 0.698 (0.222) |

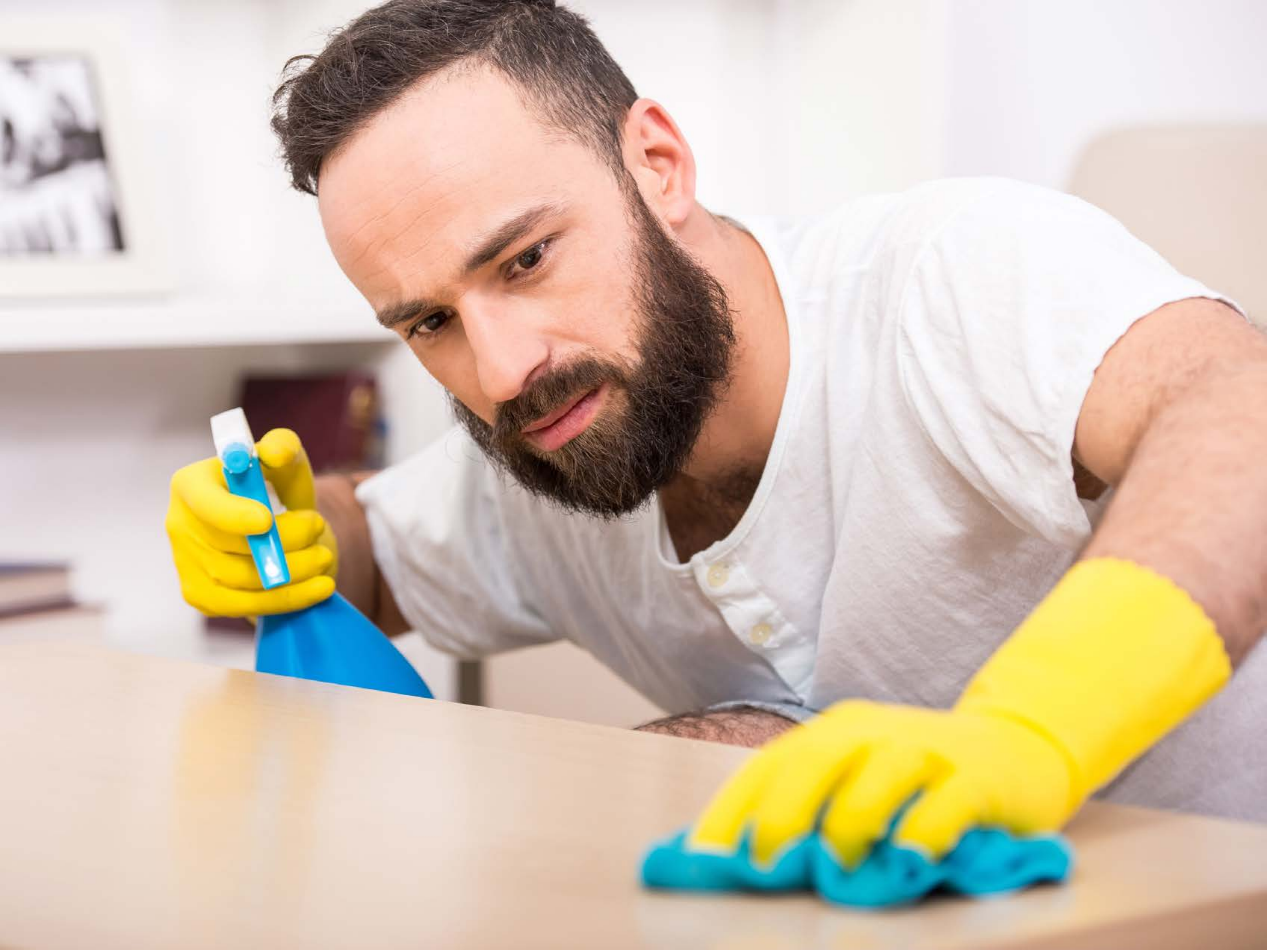

| Category | Craving (0-100) | Valence (1-9) | Arousal (1-9) | Typicality (0-100) | Relatedness | | | | | HSV | | |
|---|---|---|---|---|---|---|---|---|---|---|---|---|
| | | | | | Meth | Opioid | Both | Neither | MethToOpioid | Hue | Saturation | Value |
| Neutral tool face | 14.36 (24.63) | 4.75 (1.713) | 2.18 (2) | 16.07 (23.69) | 0.071 | 0 | 0.036 | 0.893 | -0.071 | 0.32 (0.379) | 0.229 (0.254) | 0.84 (0.187) |

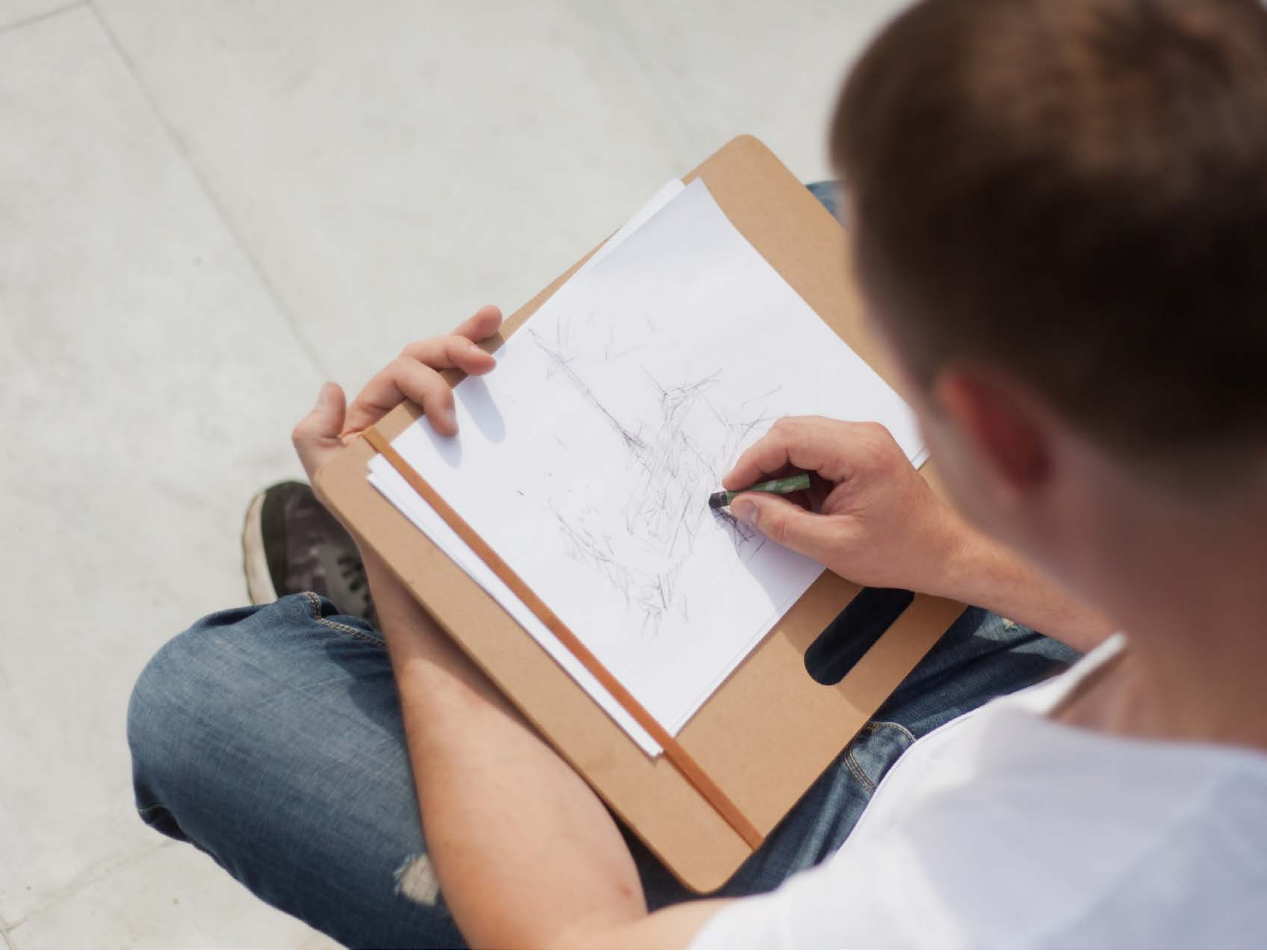

| Category | Craving (0-100) | Valence (1-9) | Arousal (1-9) | Typicality (0-100) | Relatedness | | | | | HSV | | |
|---|---|---|---|---|---|---|---|---|---|---|---|---|
| | | | | | Meth | Opioid | Both | Neither | MethToOpioid | Hue | Saturation | Value |
| Neutral tool face | 17.93 (27.47) | 4.78 (0.847) | 2.52 (1.95) | 20.56 (27.33) | 0.074 | 0 | 0 | 0.926 | -0.074 | 0.261 (0.268) | 0.199 (0.188) | 0.651 (0.262) |

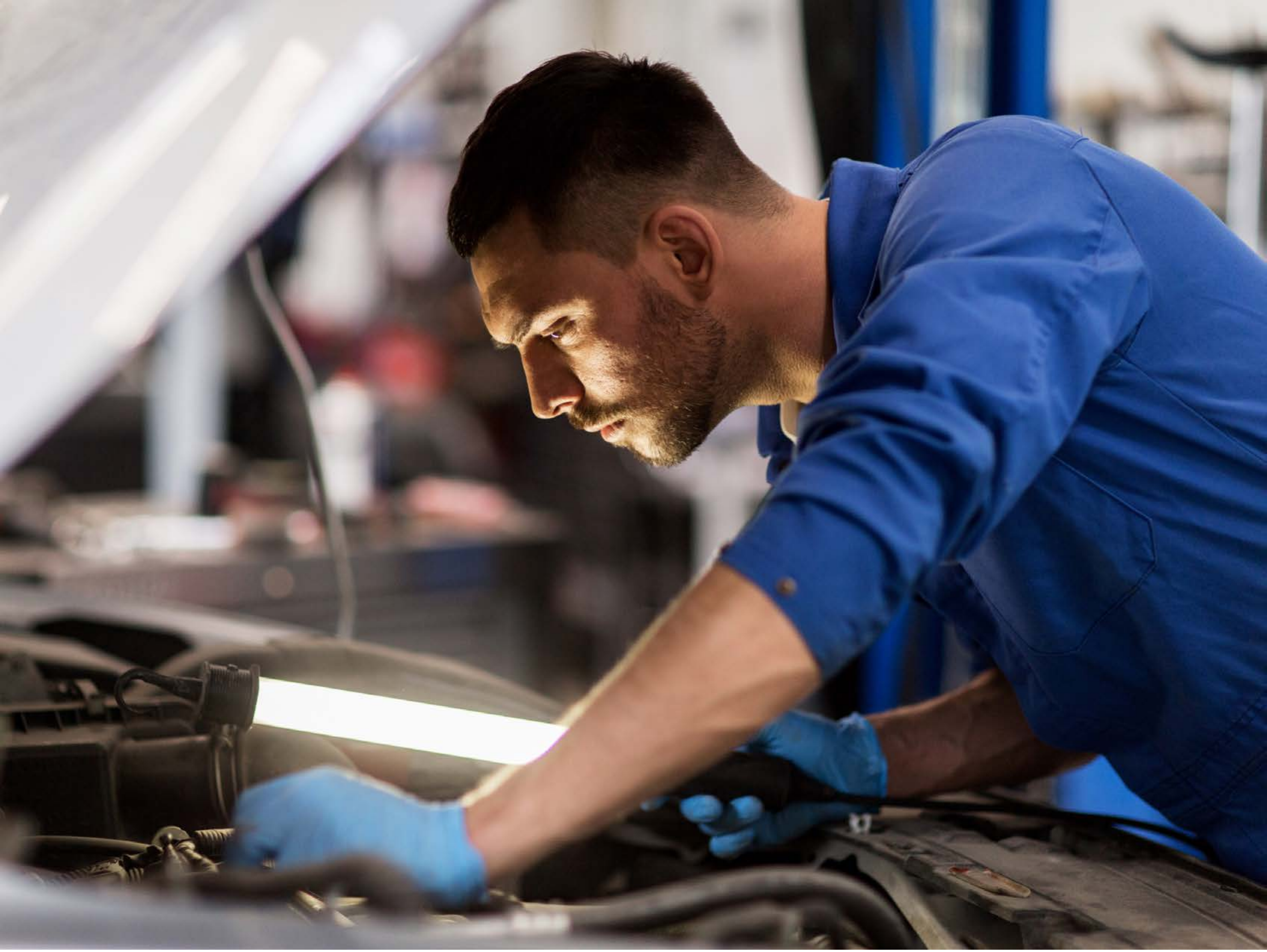

| Category | Craving (0-100) | Valence (1-9) | Arousal (1-9) | Typicality (0-100) | Relatedness | | | | | HSV | | |
|---|---|---|---|---|---|---|---|---|---|---|---|---|
| | | | | | Meth | Opioid | Both | Neither | MethToOpioid | Hue | Saturation | Value |
| Neutral tool face | 16.56 (28.49) | 4.89 (1.281) | 2.93 (2.11) | 20.26 (26.36) | 0.037 | 0 | 0 | 0.963 | -0.037 | 0.331 (0.323) | 0.393 (0.308) | 0.47 (0.271) |

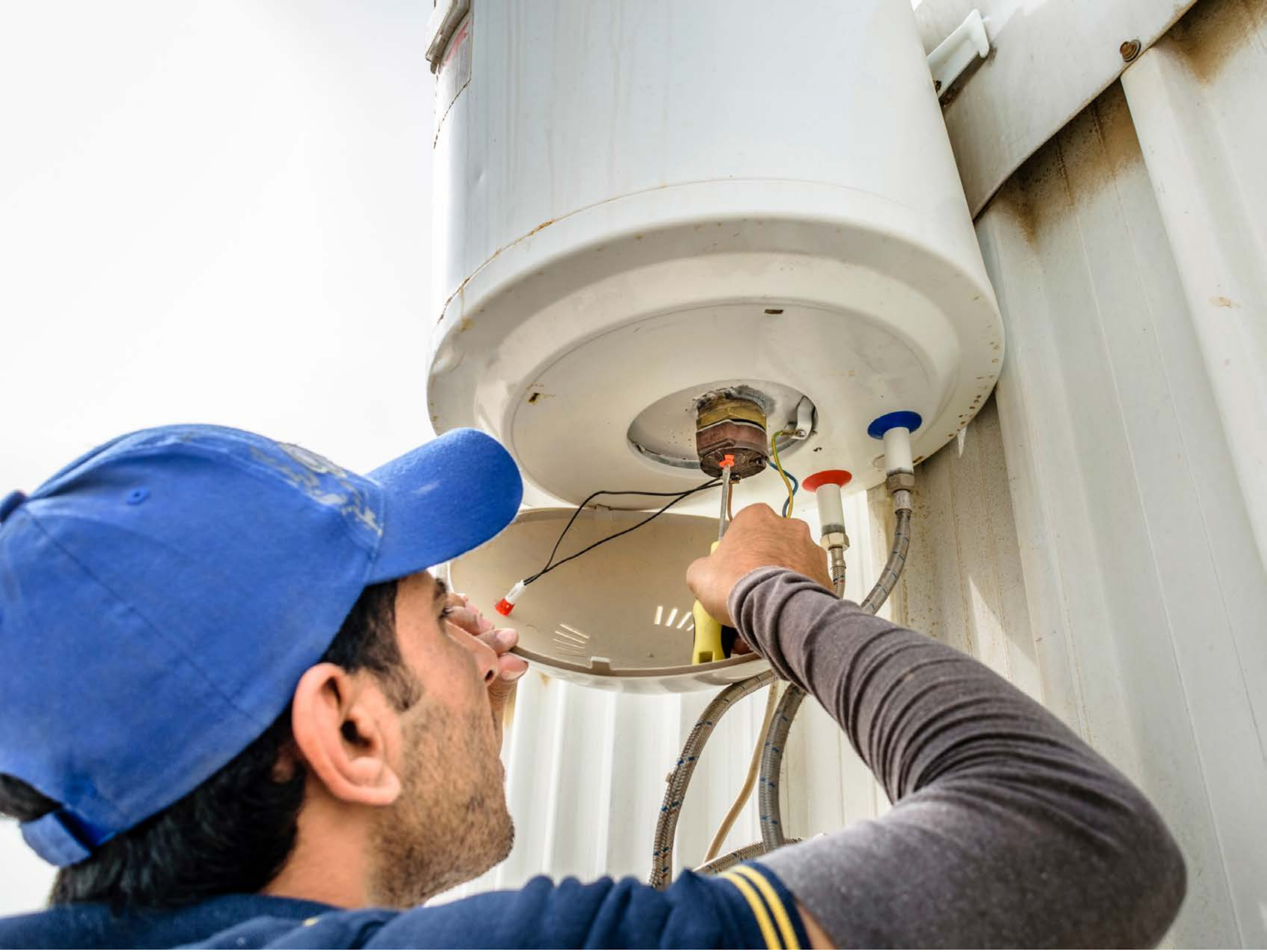

| Category | Craving (0-100) | Valence (1-9) | Arousal (1-9) | Typicality (0-100) | Relatedness | | | | | HSV | | |
|---|---|---|---|---|---|---|---|---|---|---|---|---|
| | | | | | Meth | Opioid | Both | Neither | MethToOpioid | Hue | Saturation | Value |
| Neutral tool face | 9.81 (18.02) | 4.89 (0.577) | 2.56 (1.93) | 10.81 (17.78) | 0 | 0 | 0 | 1 | 0 | 0.24 (0.244) | 0.213 (0.251) | 0.716 (0.227) |

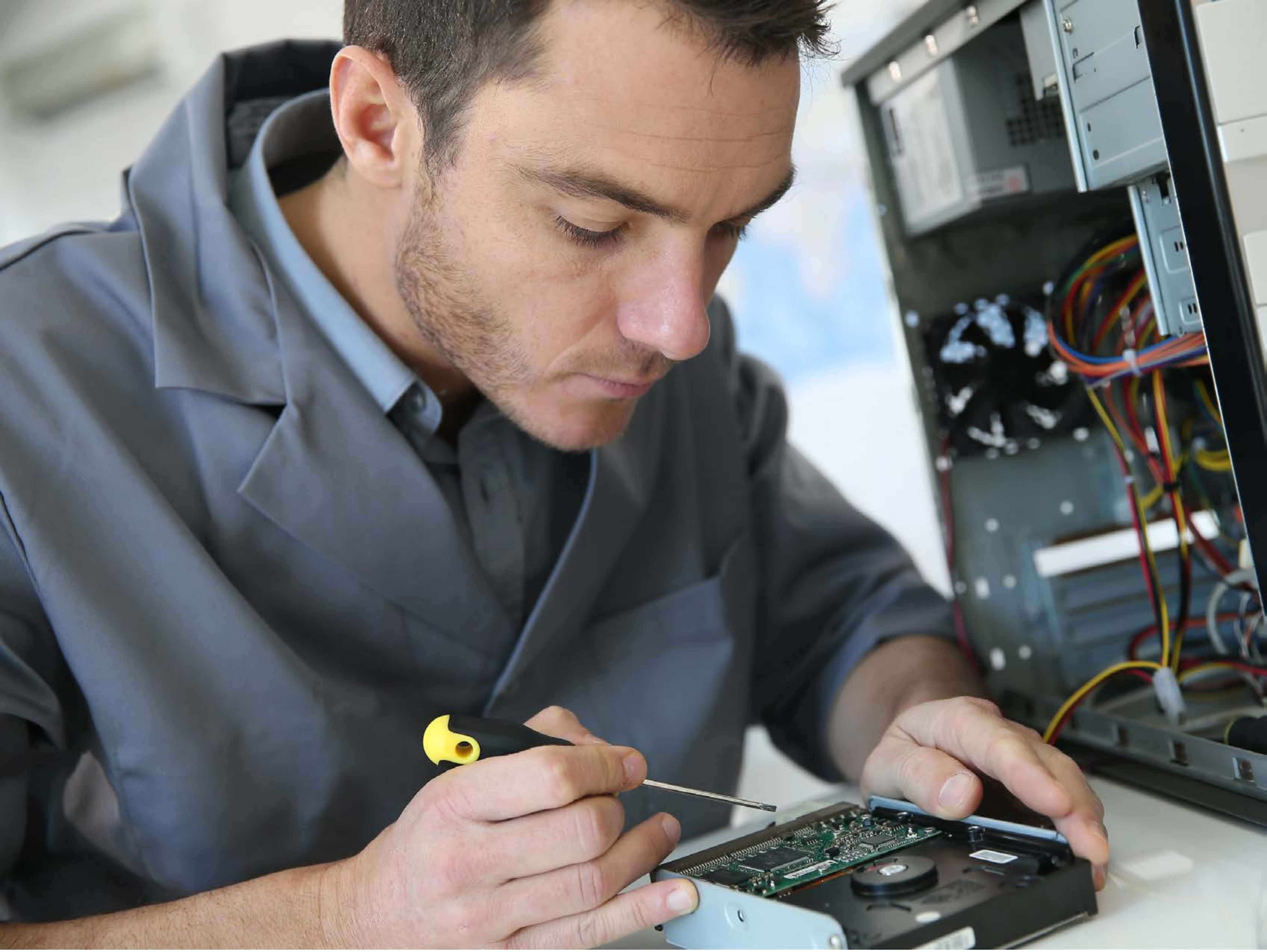

| Category | Craving (0-100) | Valence (1-9) | Arousal (1-9) | Typicality (0-100) | Relatedness | | | | | HSV | | |
|---|---|---|---|---|---|---|---|---|---|---|---|---|
| | | | | | Meth | Opioid | Both | Neither | MethToOpioid | Hue | Saturation | Value |
| Neutral tool face | 9.57 (17.27) | 4.89 (1.641) | 2.32 (1.89) | 11.96 (17.95) | 0 | 0 | 0 | 1 | 0 | 0.401 (0.234) | 0.201 (0.123) | 0.48 (0.263) |

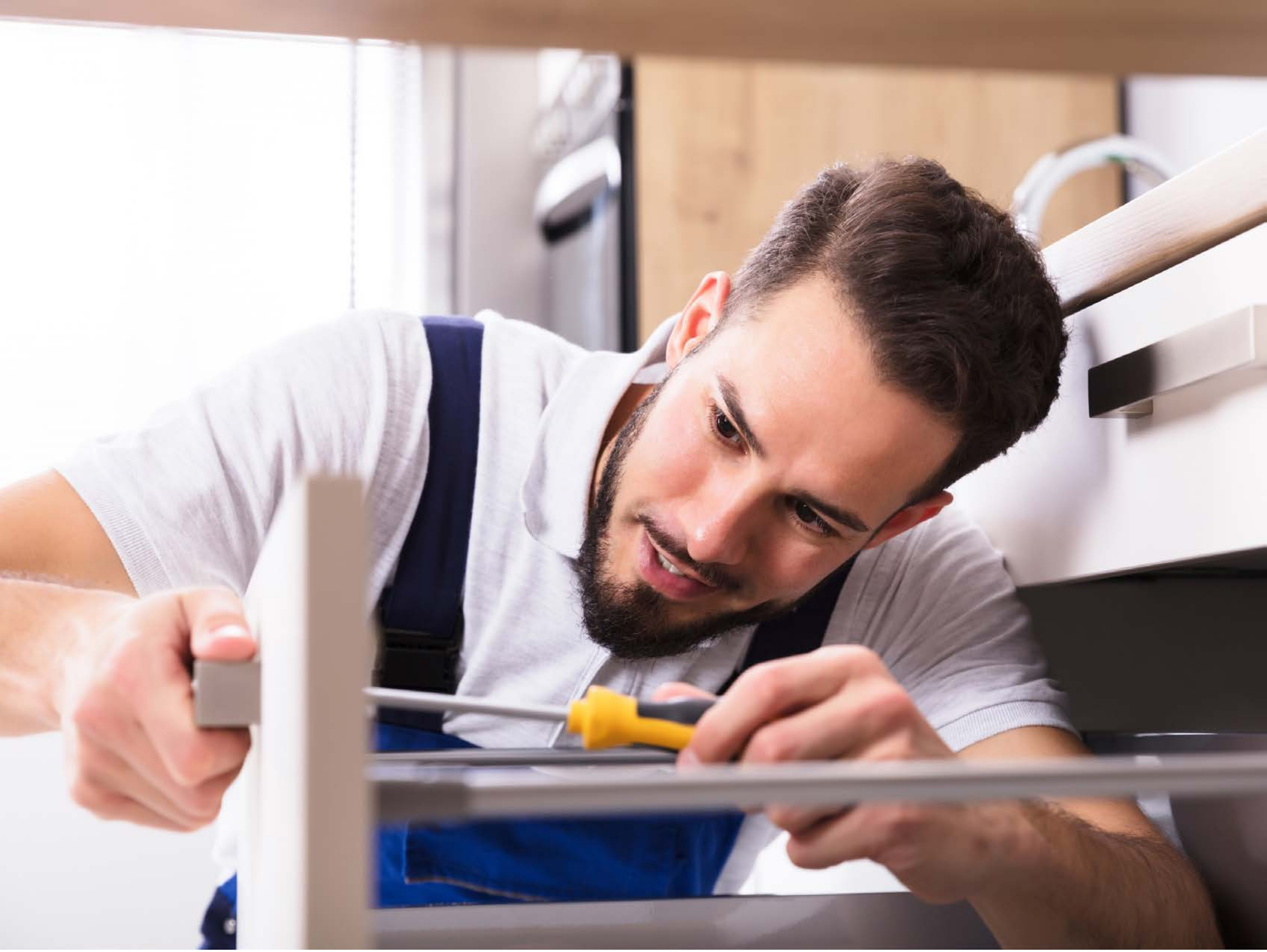

| Category | Craving (0-100) | Valence (1-9) | Arousal (1-9) | Typicality (0-100) | Relatedness | | | | | HSV | | |
|---|---|---|---|---|---|---|---|---|---|---|---|---|
| | | | | | Meth | Opioid | Both | Neither | MethToOpioid | Hue | Saturation | Value |
| Neutral tool face | 8.32 (21.29) | 4.79 (1.572) | 2.14 (2.01) | 10.29 (22.22) | 0 | 0 | 0.036 | 0.964 | 0 | 0.373 (0.395) | 0.201 (0.203) | 0.709 (0.281) |

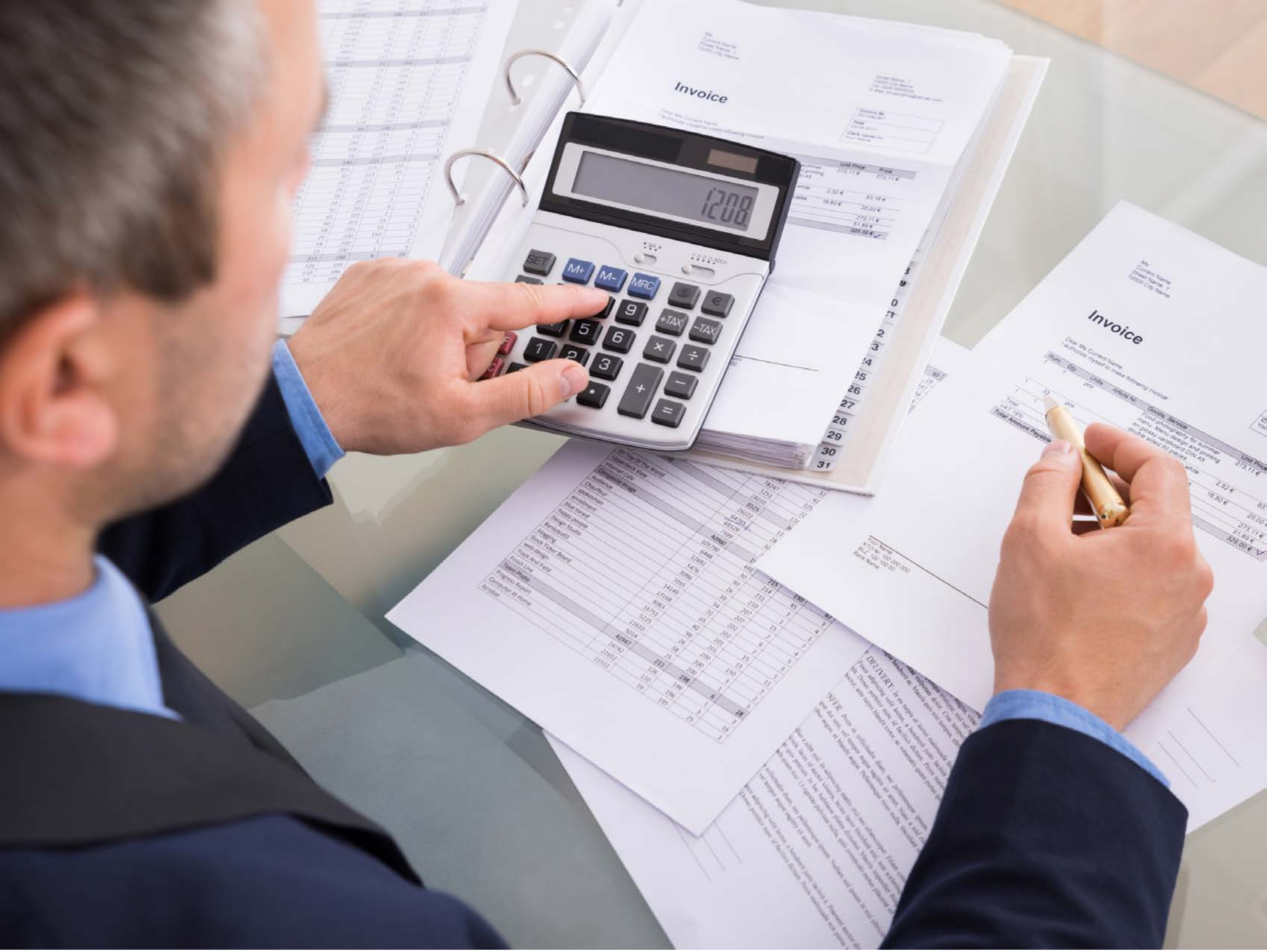


| Category | Craving (0-100) | Valence (1-9) | Arousal (1-9) | Typicality (0-100) | Relatedness | | | | | HSV | | |
|---|---|---|---|---|---|---|---|---|---|---|---|---|
| | | | | | Meth | Opioid | Both | Neither | MethToOpioid | Hue | Saturation | Value |
| Neutral tool face | 6.89 (15.5) | 4.82 (1.786) | 2.54 (1.9) | 7.64 (15.35) | 0 | 0 | 0 | 1 | 0 | 0.496 (0.303) | 0.158 (0.151) | 0.695 (0.271) |
