## Supplementary Material 3 for "LIBR Methamphetamine and Opioid Cue Database (LIBR MOCD): Development and Validation"

### Meth and Opioid Cue Database

#### Meth Image

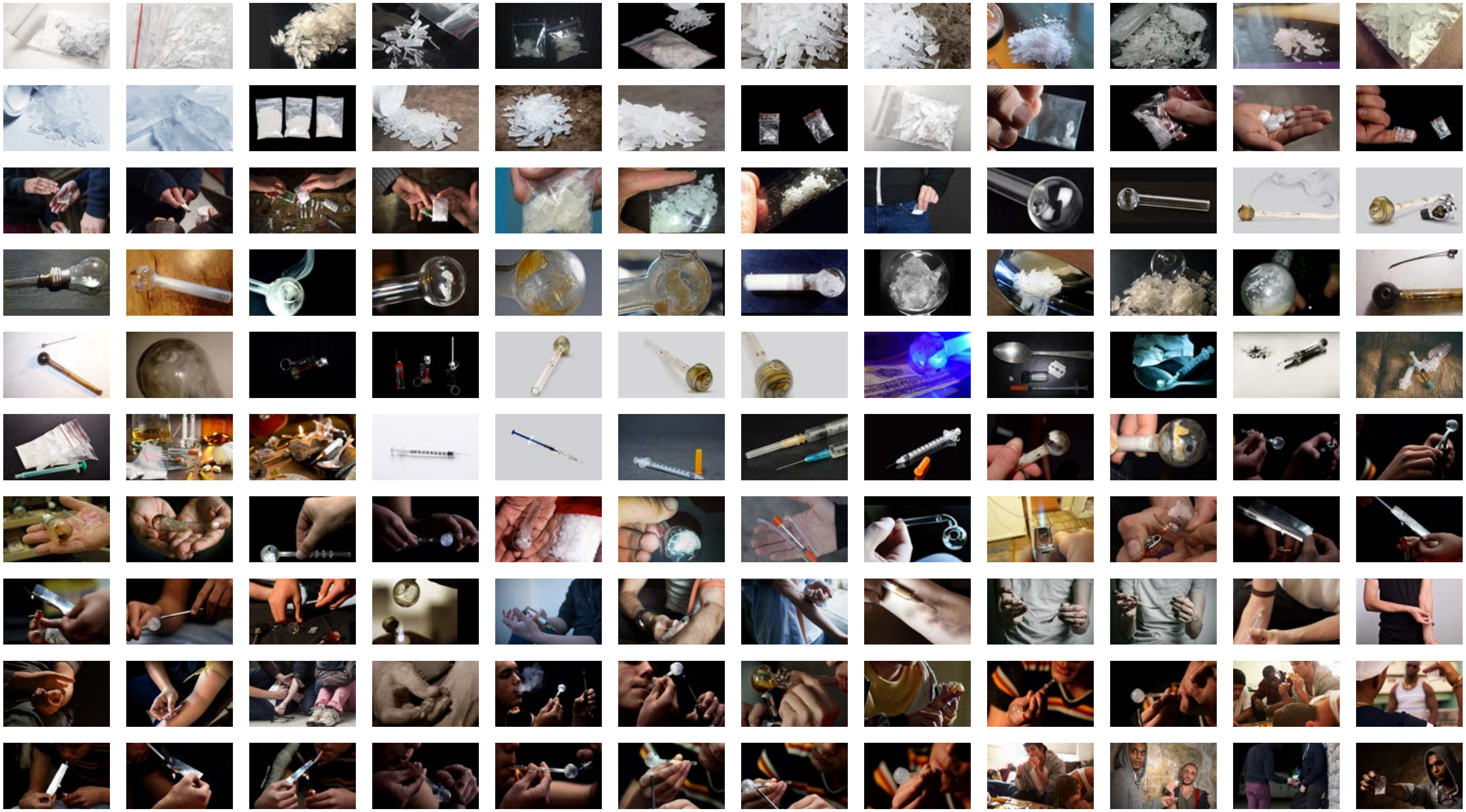

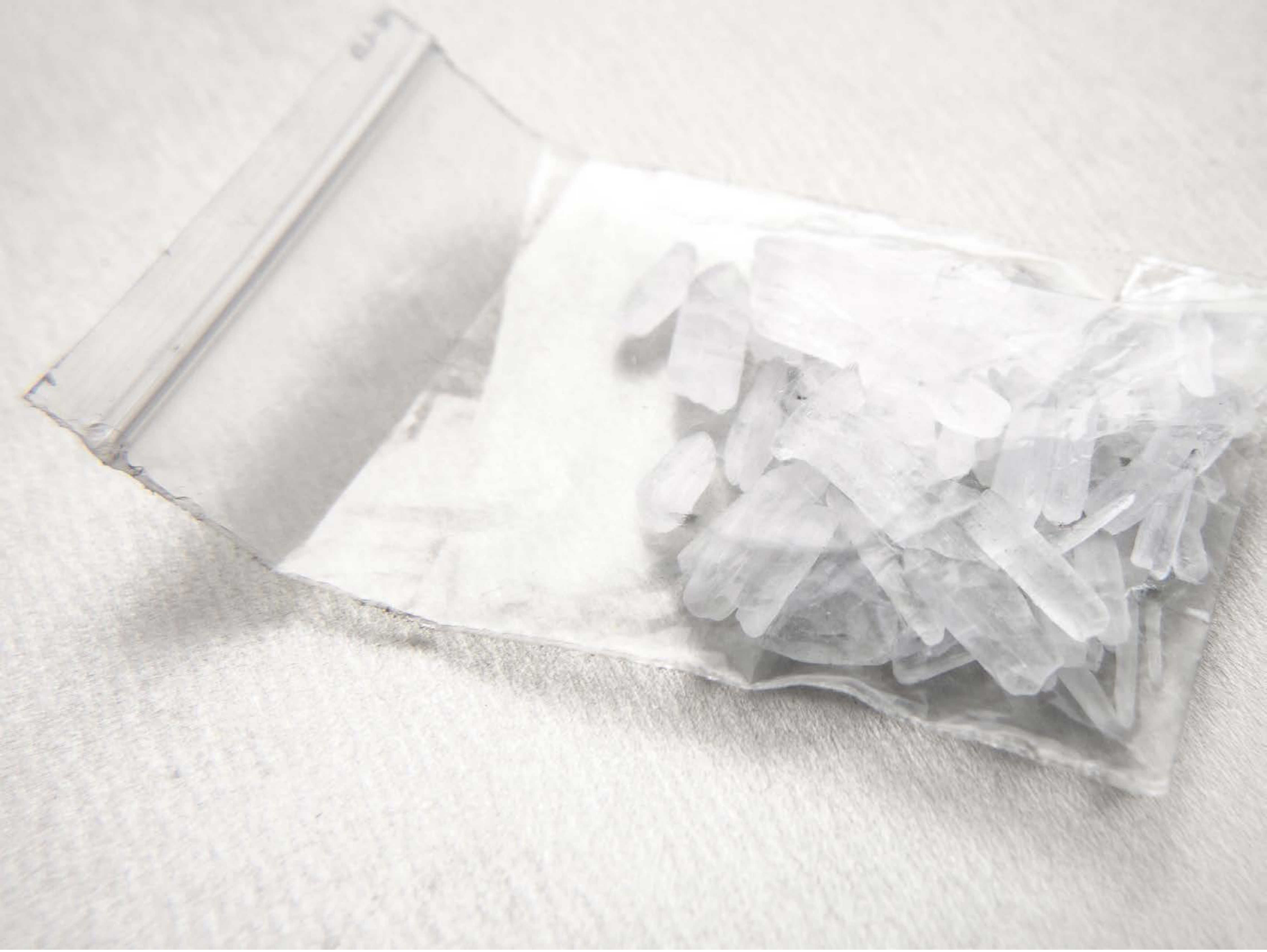

| Category | Craving (0-100) | Valence (1-9) | Arousal (1-9) | Typicality (0-100) | Relatedness | | | | | HSV | | |
|---|---|---|---|---|---|---|---|---|---|---|---|---|
| | | | | | Meth | Opioid | Both | Neither | MethToOpioid | Hue | Saturation | Value |
| Meth | 92.07 (12.9) | 4.5 (1.953) | 4.5 (2.8) | 93.32 (11.12) | 1 | 0 | 0 | 0 | -1 | 0.22 (0.299) | 0.024 (0.015) | 0.882 (0.093) |

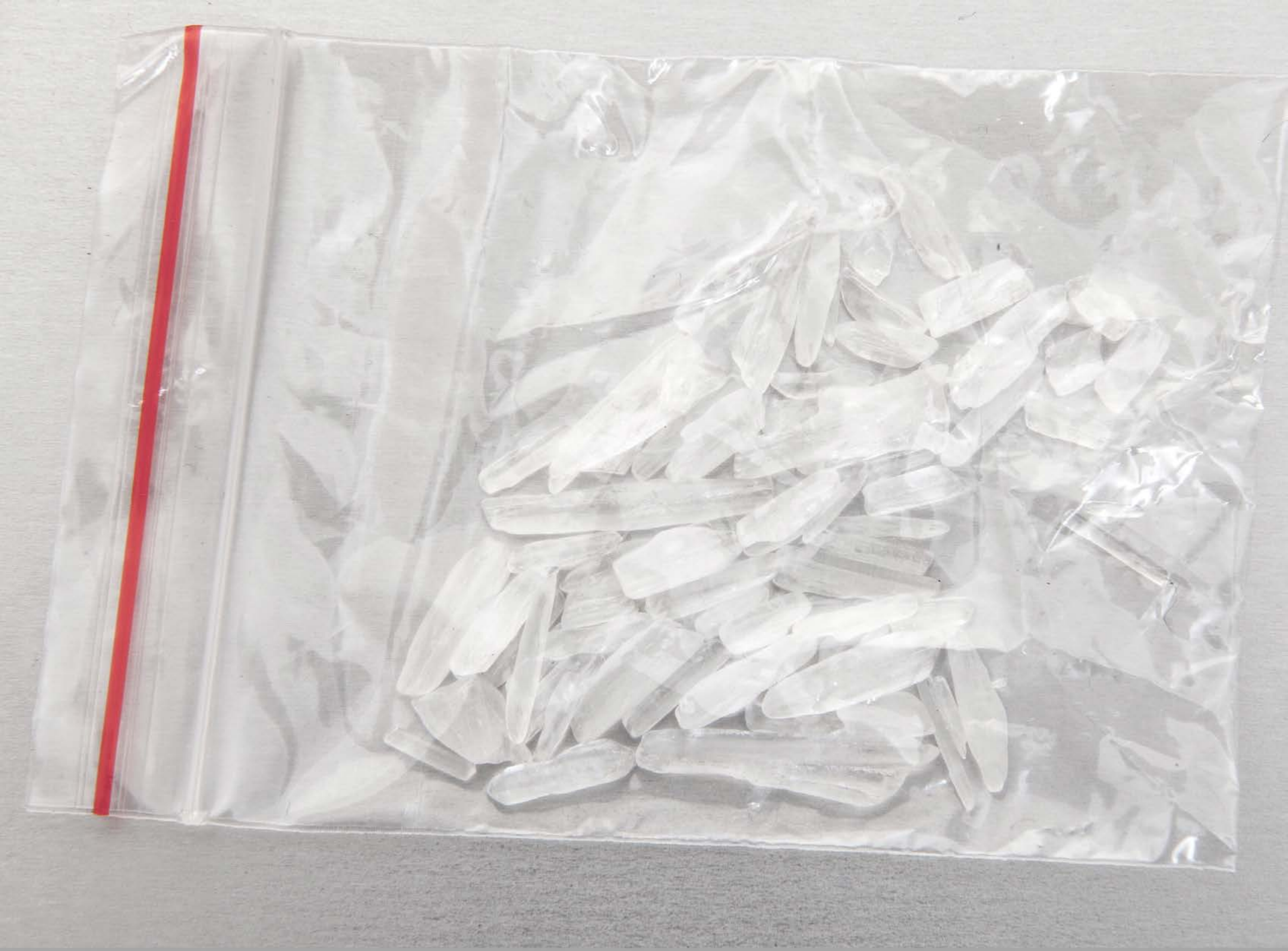

| Category | Craving (0-100) | Valence (1-9) | Arousal (1-9) | Typicality (0-100) | Relatedness | | | | | HSV | | |
|---|---|---|---|---|---|---|---|---|---|---|---|---|
| | | | | | Meth | Opioid | Both | Neither | MethToOpioid | Hue | Saturation | Value |
| Meth | 91.26 (13.03) | 4.7 (2.053) | 4.7 (2.87) | 89.93 (15.26) | 1 | 0 | 0 | 0 | -1 | 0.314 (0.373) | 0.022 (0.062) | 0.822 (0.081) |

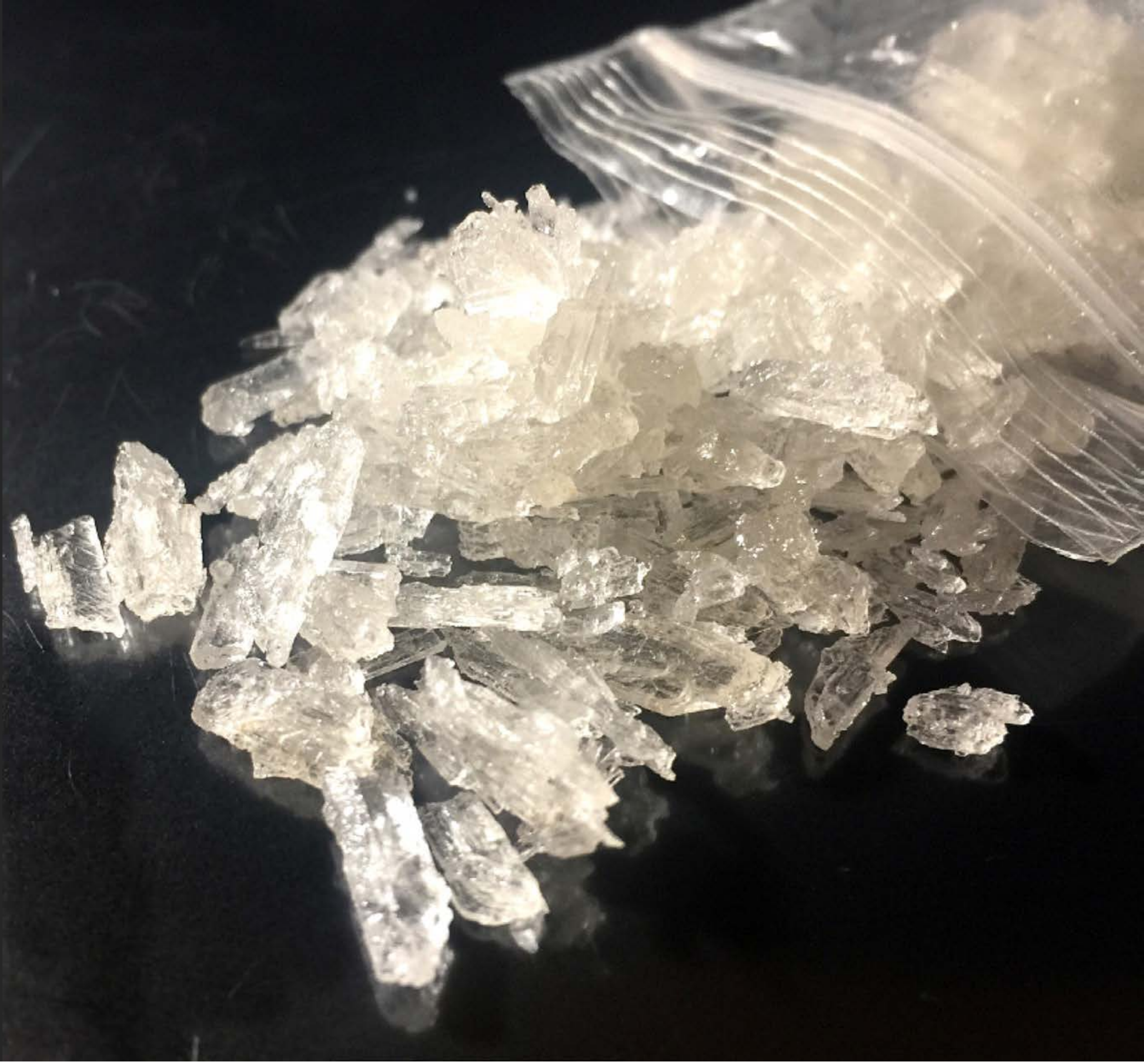

| Category | Craving (0-100) | Valence (1-9) | Arousal (1-9) | Typicality (0-100) | Relatedness | | | | | HSV | | |
|---|---|---|---|---|---|---|---|---|---|---|---|---|
| | | | | | Meth | Opioid | Both | Neither | MethToOpioid | Hue | Saturation | Value |
| Meth | 88.75 (15.22) | 4.64 (2.181) | 4.89 (2.86) | 91.36 (14.08) | 1 | 0 | 0 | 0 | -1 | 0.195 (0.245) | 0.131 (0.127) | 0.387 (0.333) |

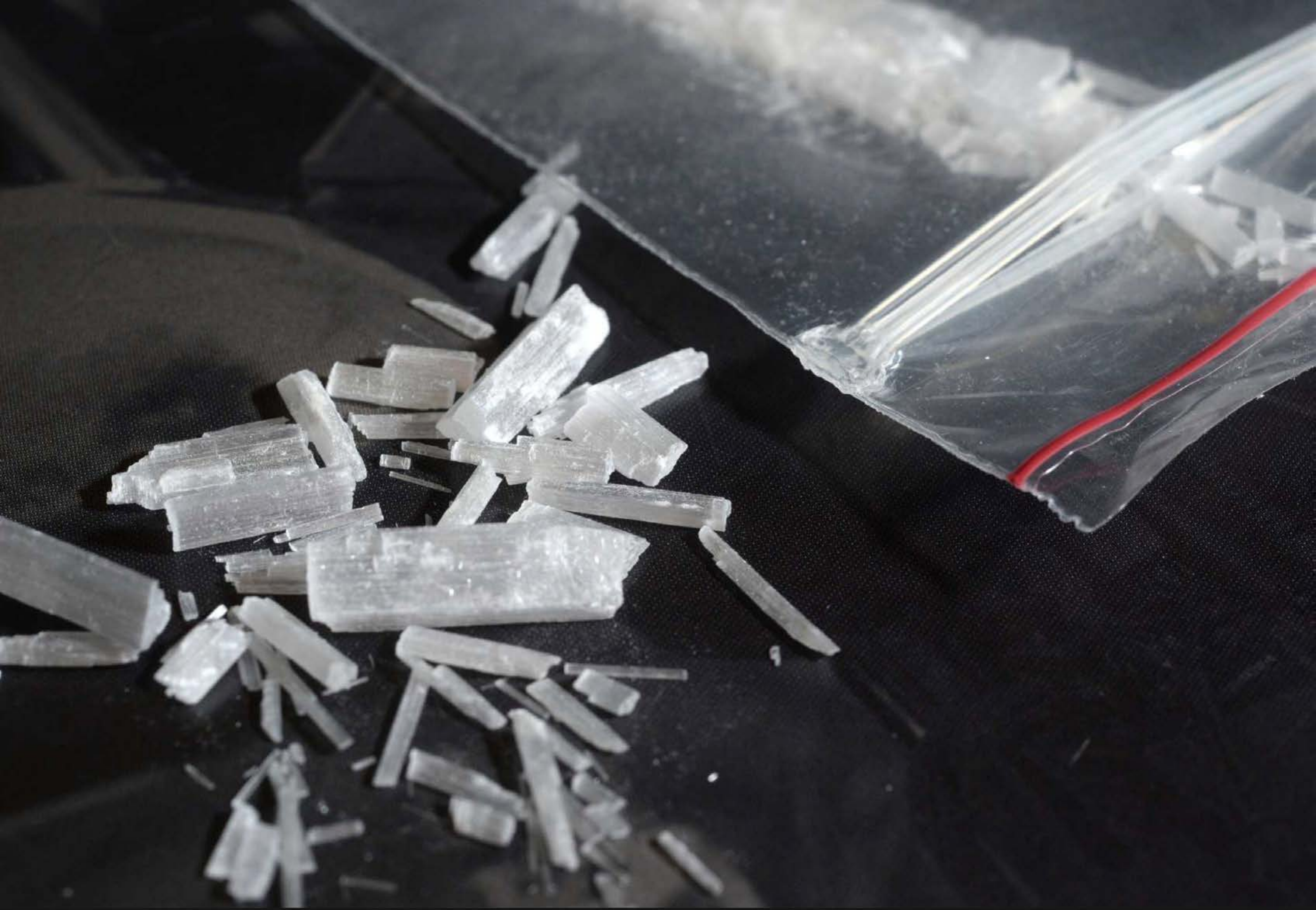

| Category | Craving (0-100) | Valence (1-9) | Arousal (1-9) | Typicality (0-100) | Relatedness | | | | | HSV | | |
|---|---|---|---|---|---|---|---|---|---|---|---|---|
| | | | | | Meth | Opioid | Both | Neither | MethToOpioid | Hue | Saturation | Value |
| Meth | 85.22 (22.52) | 4.52 (1.827) | 4.63 (2.6) | 84.59 (23.82) | 1 | 0 | 0 | 0 | -1 | 0.419 (0.324) | 0.085 (0.088) | 0.307 (0.255) |

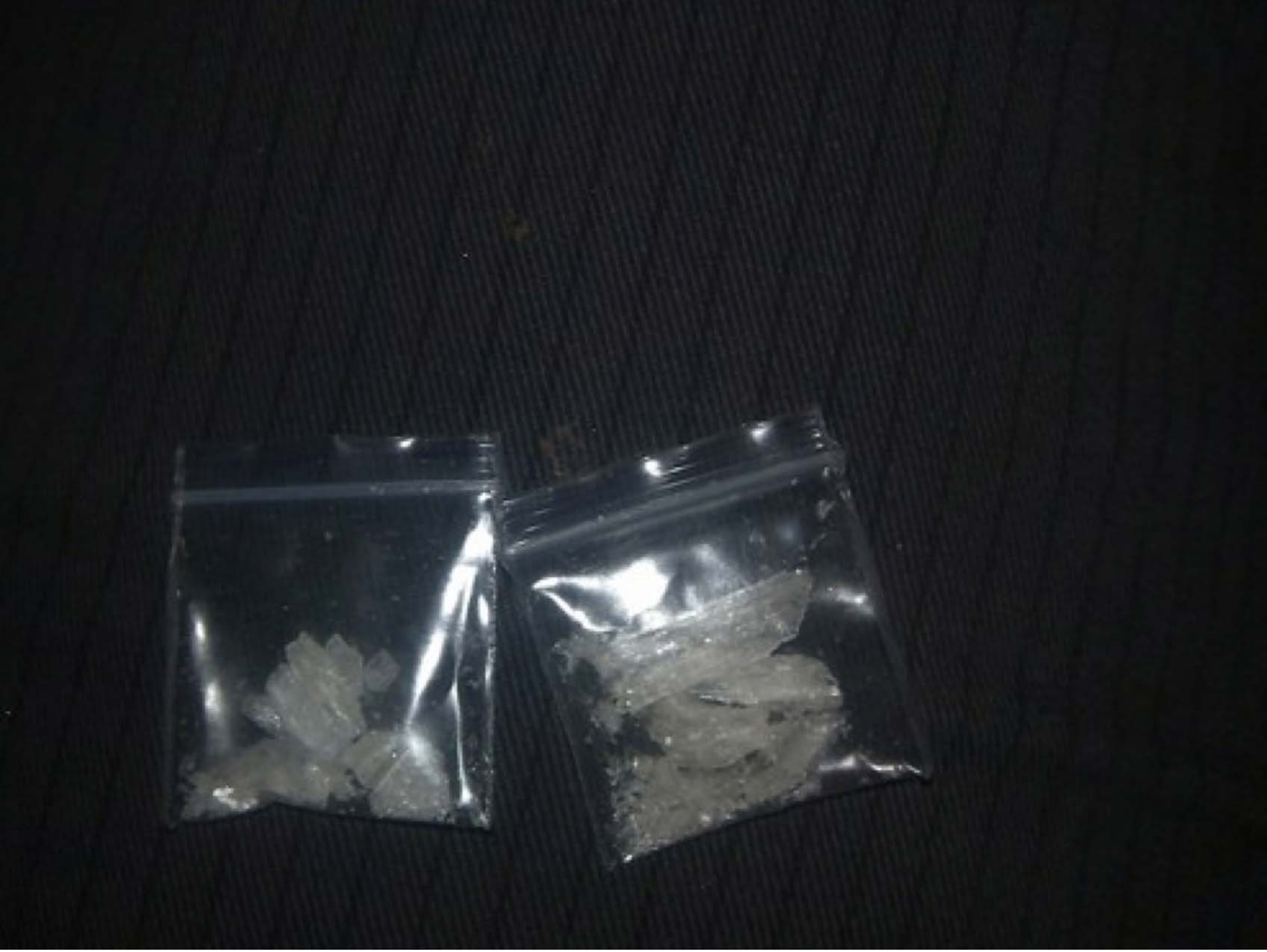

| Category | Craving (0-100) | Valence (1-9) | Arousal (1-9) | Typicality (0-100) | Relatedness | | | | | HSV | | |
|---|---|---|---|---|---|---|---|---|---|---|---|---|
| | | | | | Meth | Opioid | Both | Neither | MethToOpioid | Hue | Saturation | Value |
| Meth | 91.43 (13.8) | 4.5 (2.134) | 4.96 (2.87) | 93.11 (13.25) | 1 | 0 | 0 | 0 | -1 | 0.504 (0.199) | 0.141 (0.08) | 0.145 (0.129) |

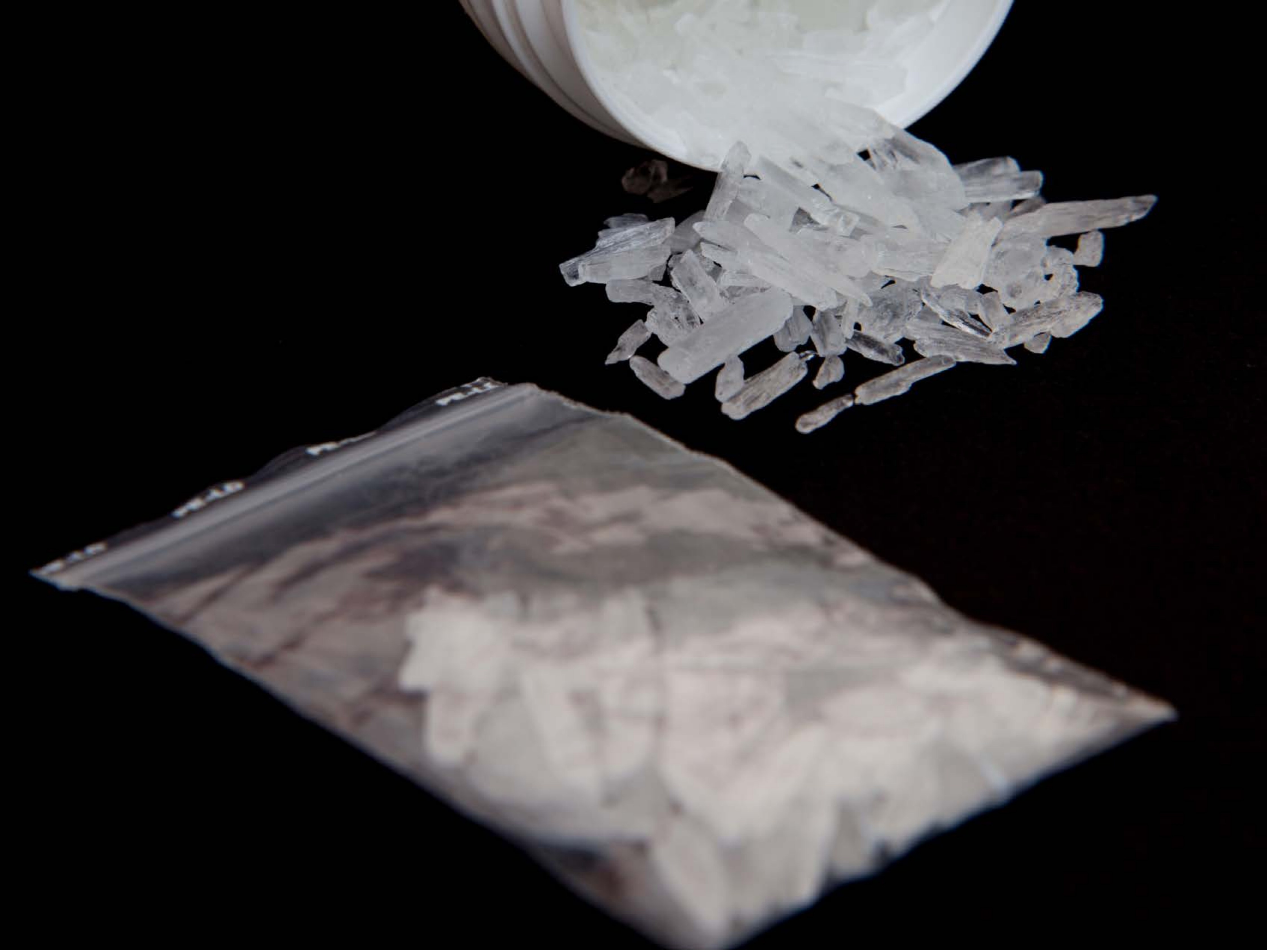

| Category | Craving (0-100) | Valence (1-9) | Arousal (1-9) | Typicality (0-100) | Relatedness | | | | | HSV | | |
|---|---|---|---|---|---|---|---|---|---|---|---|---|
| | | | | | Meth | Opioid | Both | Neither | MethToOpioid | Hue | Saturation | Value |
| Meth | 91.79 (12.59) | 4.39 (2.266) | 4.68 (2.91) | 94.32 (10.15) | 1 | 0 | 0 | 0 | -1 | 0.247 (0.363) | 0.152 (0.309) | 0.232 (0.294) |

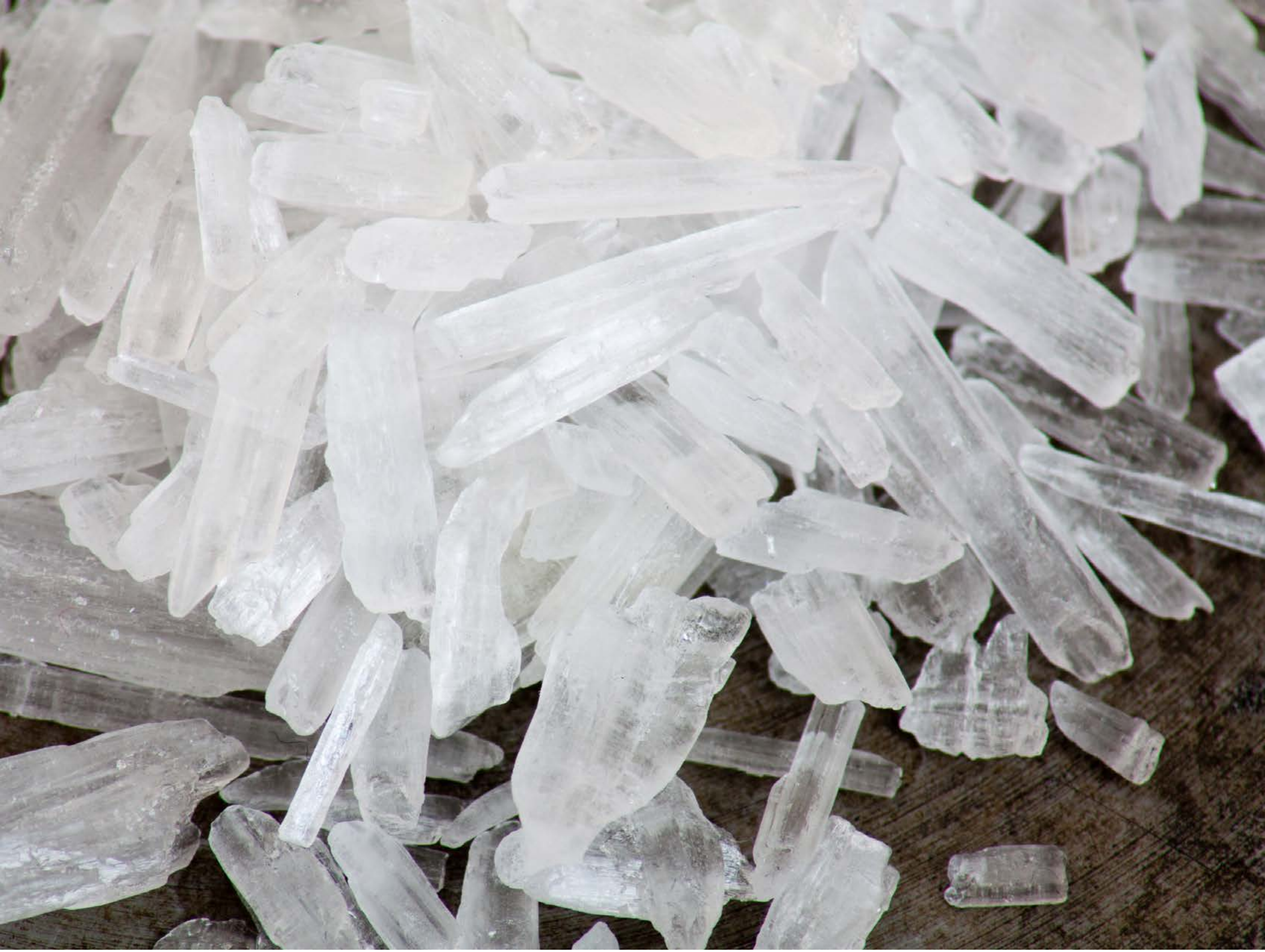

| Category | Craving (0-100) | Valence (1-9) | Arousal (1-9) | Typicality (0-100) | Relatedness | | | | | HSV | | |
|---|---|---|---|---|---|---|---|---|---|---|---|---|
| | | | | | Meth | Opioid | Both | Neither | MethToOpioid | Hue | Saturation | Value |
| Meth | 91.41 (12.29) | 4.74 (1.992) | 4.67 (2.9) | 92.67 (12.4) | 1 | 0 | 0 | 0 | -1 | 0.438 (0.311) | 0.085 (0.154) | 0.678 (0.218) |

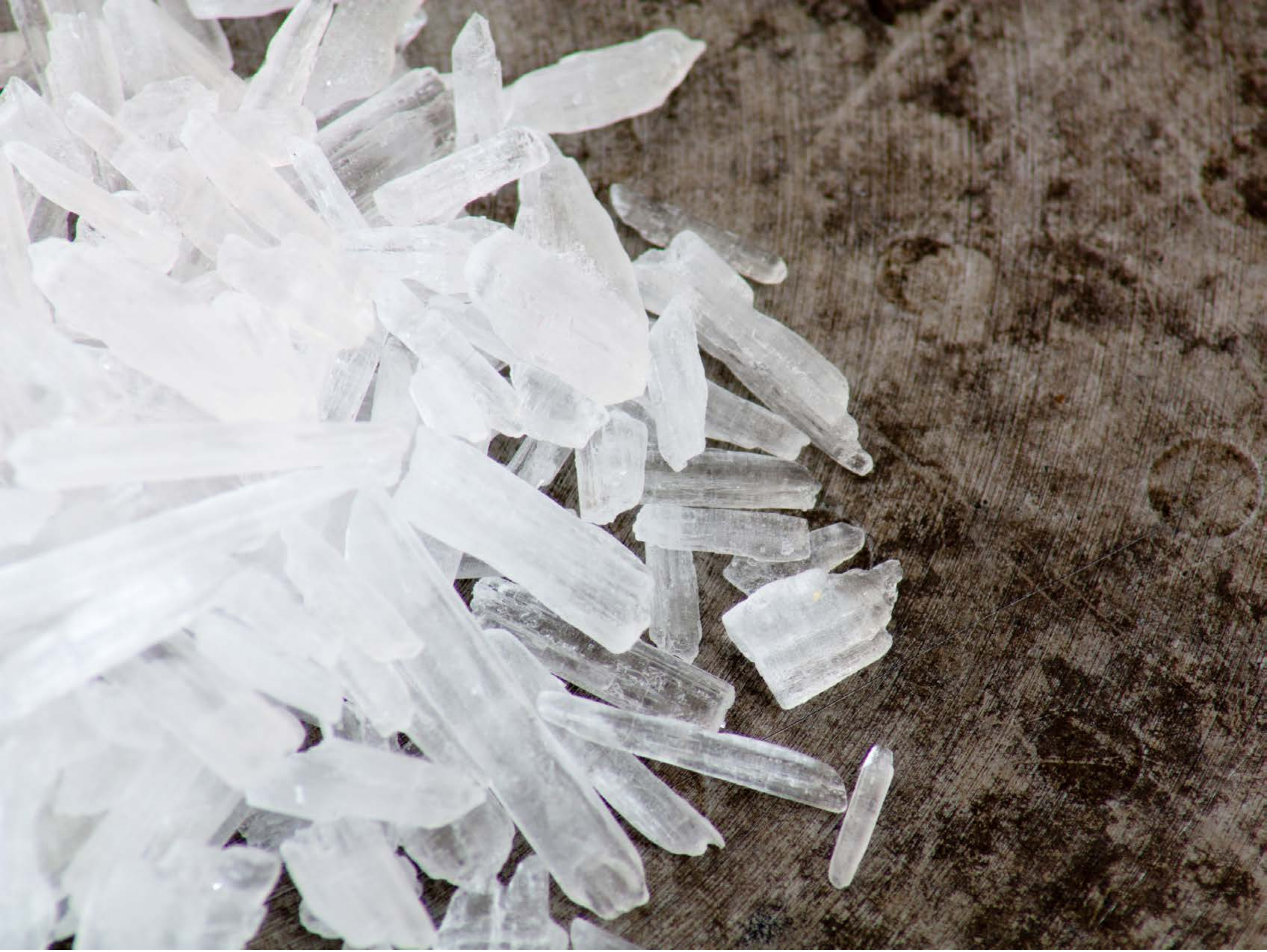

| Category | Craving (0-100) | Valence (1-9) | Arousal (1-9) | Typicality (0-100) | Relatedness | | | | | HSV | | |
|---|---|---|---|---|---|---|---|---|---|---|---|---|
| | | | | | Meth | Opioid | Both | Neither | MethToOpioid | Hue | Saturation | Value |
| Meth | 85 (23.29) | 4.48 (1.968) | 4.59 (2.71) | 84.89 (24.56) | 1 | 0 | 0 | 0 | -1 | 0.375 (0.296) | 0.161 (0.186) | 0.627 (0.26) |

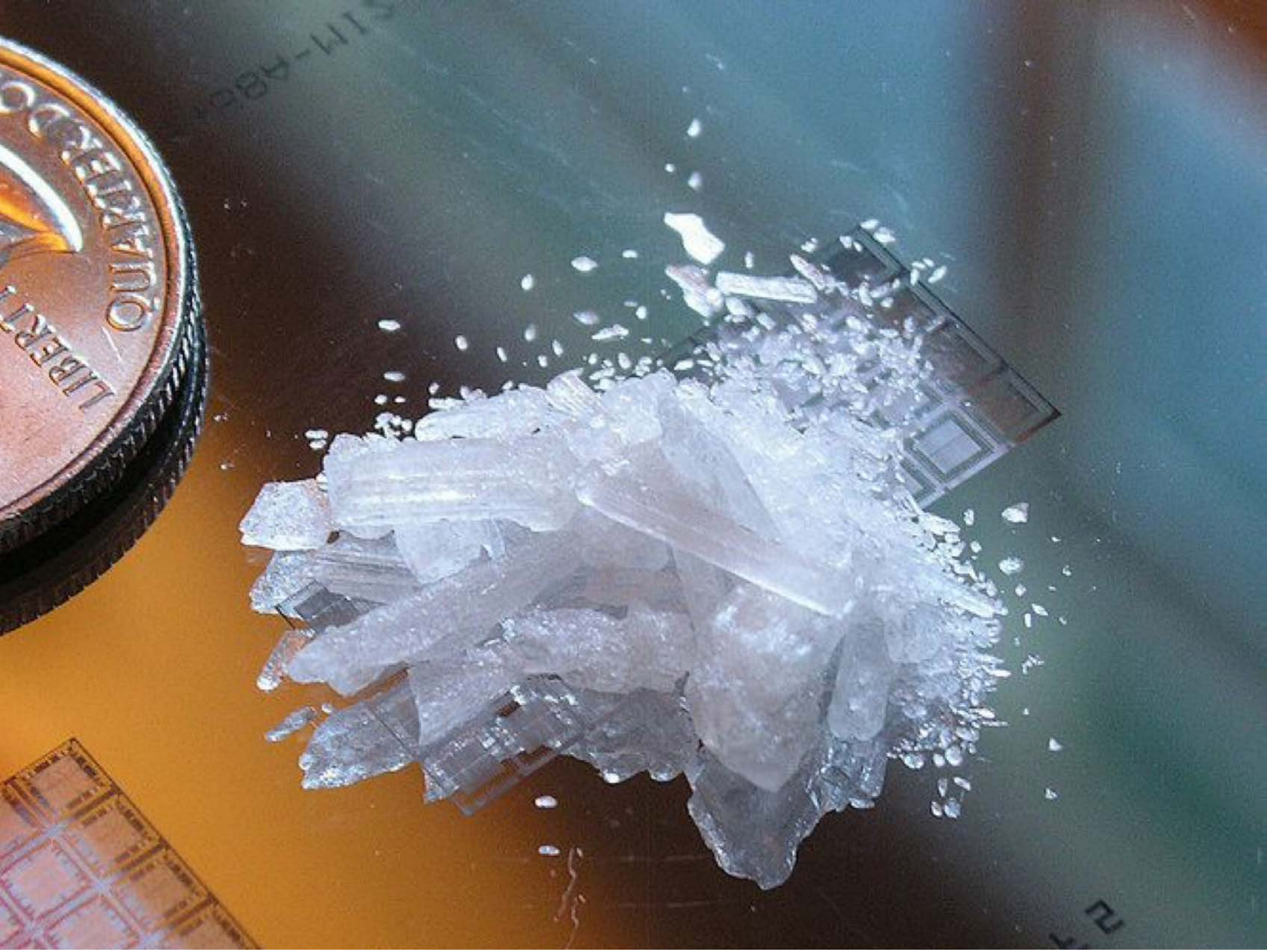

| Category | Craving (0-100) | Valence (1-9) | Arousal (1-9) | Typicality (0-100) | Relatedness | | | | | HSV | | |
|---|---|---|---|---|---|---|---|---|---|---|---|---|
| | | | | | Meth | Opioid | Both | Neither | MethToOpioid | Hue | Saturation | Value |
| Meth | 90.41 (15.33) | 4.56 (2.276) | 4.67 (2.94) | 91.44 (16.23) | 1 | 0 | 0 | 0 | -1 | 0.458 (0.257) | 0.312 (0.206) | 0.548 (0.22) |

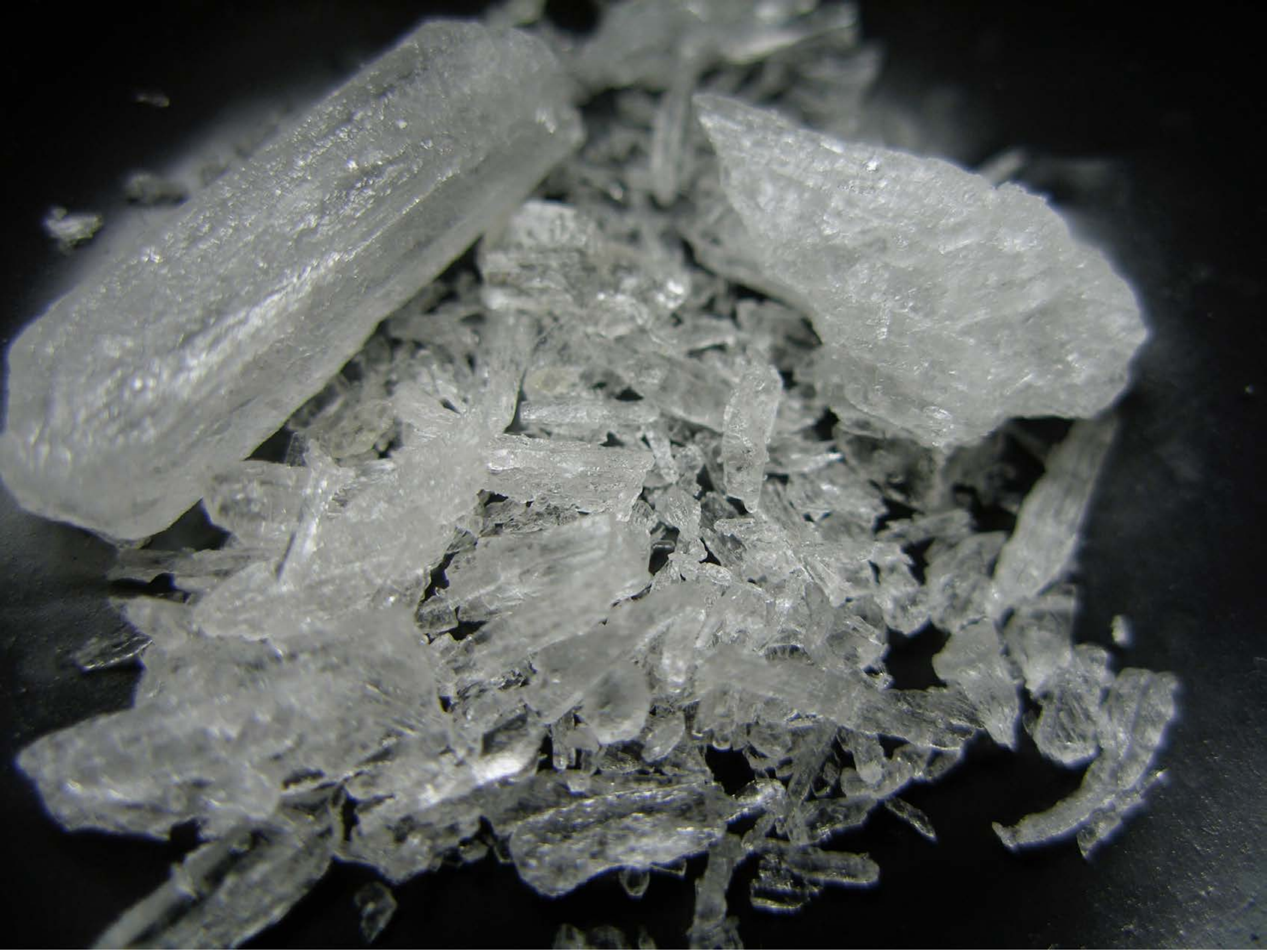

| Category | Craving (0-100) | Valence (1-9) | Arousal (1-9) | Typicality (0-100) | Relatedness | | | | | HSV | | |
|---|---|---|---|---|---|---|---|---|---|---|---|---|
| | | | | | Meth | Opioid | Both | Neither | MethToOpioid | Hue | Saturation | Value |
| Meth | 90.19 (14.79) | 4.3 (1.938) | 4.81 (2.66) | 90.26 (16.03) | 1 | 0 | 0 | 0 | -1 | 0.338 (0.199) | 0.094 (0.107) | 0.369 (0.229) |

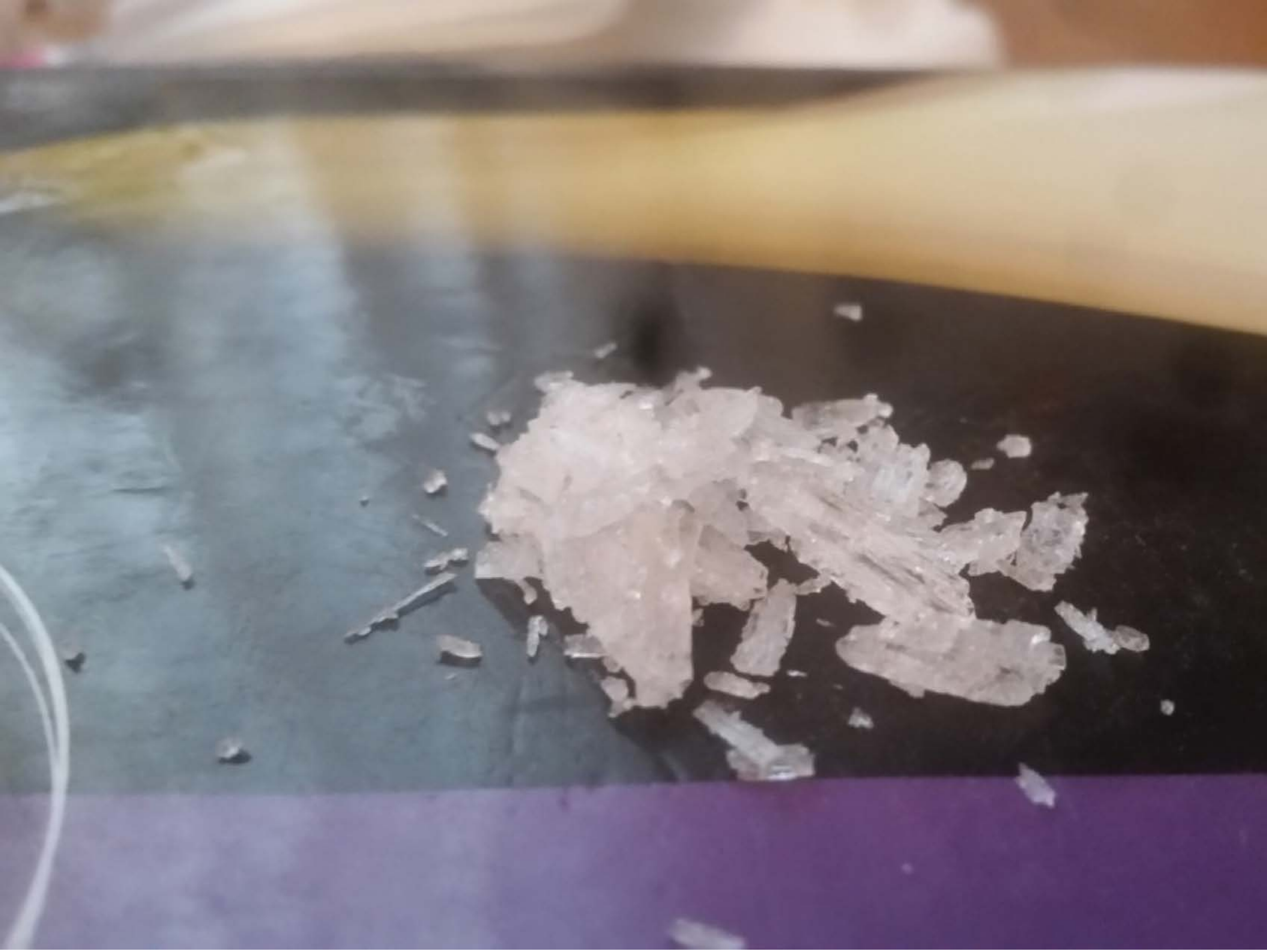

| Category | Craving (0-100) | Valence (1-9) | Arousal (1-9) | Typicality (0-100) | Relatedness | | | | | HSV | | |
|---|---|---|---|---|---|---|---|---|---|---|---|---|
| | | | | | Meth | Opioid | Both | Neither | MethToOpioid | Hue | Saturation | Value |
| Meth | 94 (11.24) | 4.25 (2.066) | 4.82 (2.97) | 94.57 (10.72) | 1 | 0 | 0 | 0 | -1 | 0.391 (0.332) | 0.209 (0.123) | 0.559 (0.198) |

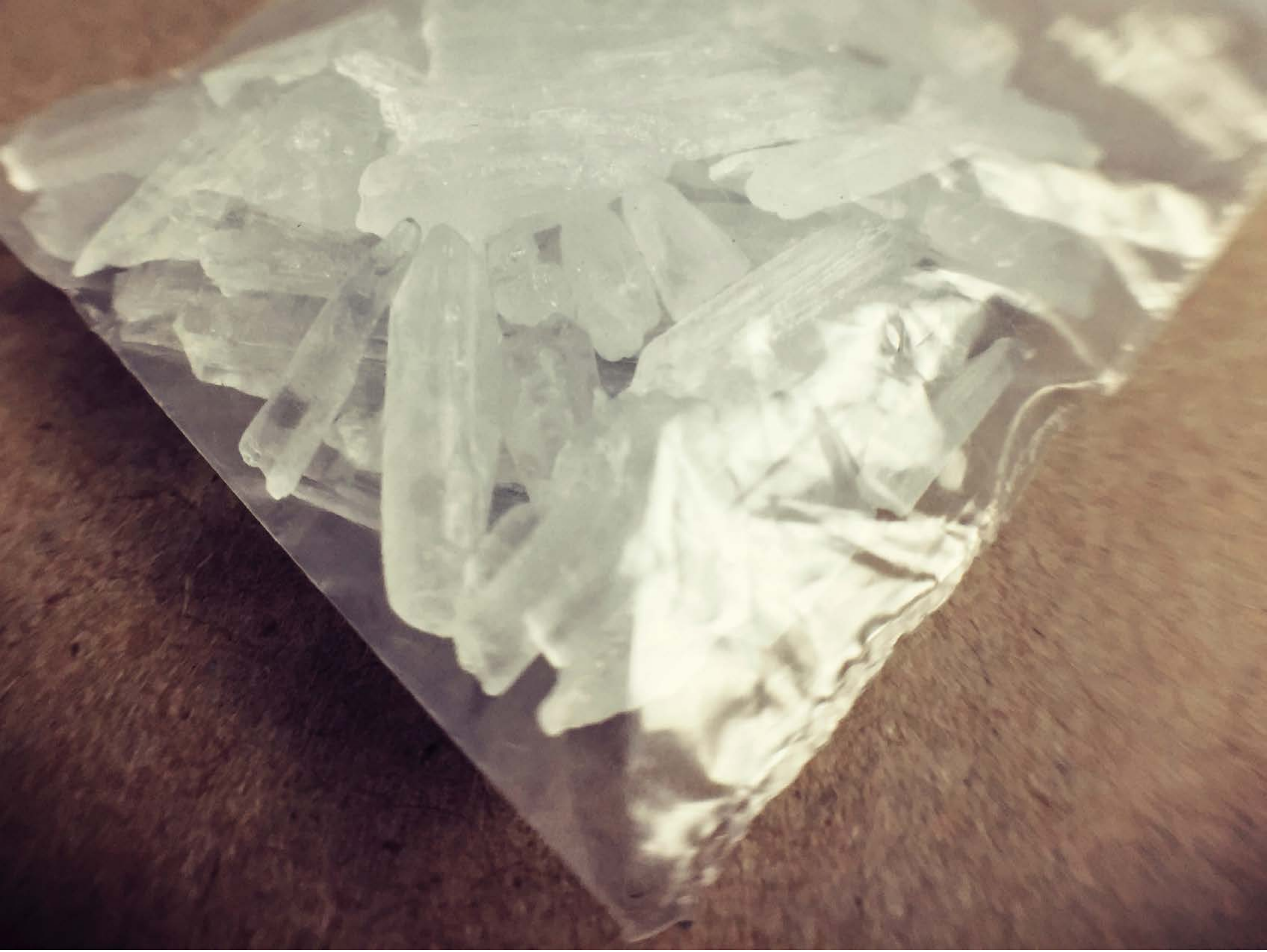

| Category | Craving (0-100) | Valence (1-9) | Arousal (1-9) | Typicality (0-100) | Relatedness | | | | | HSV | | |
|---|---|---|---|---|---|---|---|---|---|---|---|---|
| | | | | | Meth | Opioid | Both | Neither | MethToOpioid | Hue | Saturation | Value |
| Meth | 91.79 (14.57) | 4.46 (2.027) | 4.57 (3.04) | 93.36 (13.5) | 1 | 0 | 0 | 0 | -1 | 0.267 (0.318) | 0.235 (0.133) | 0.608 (0.224) |

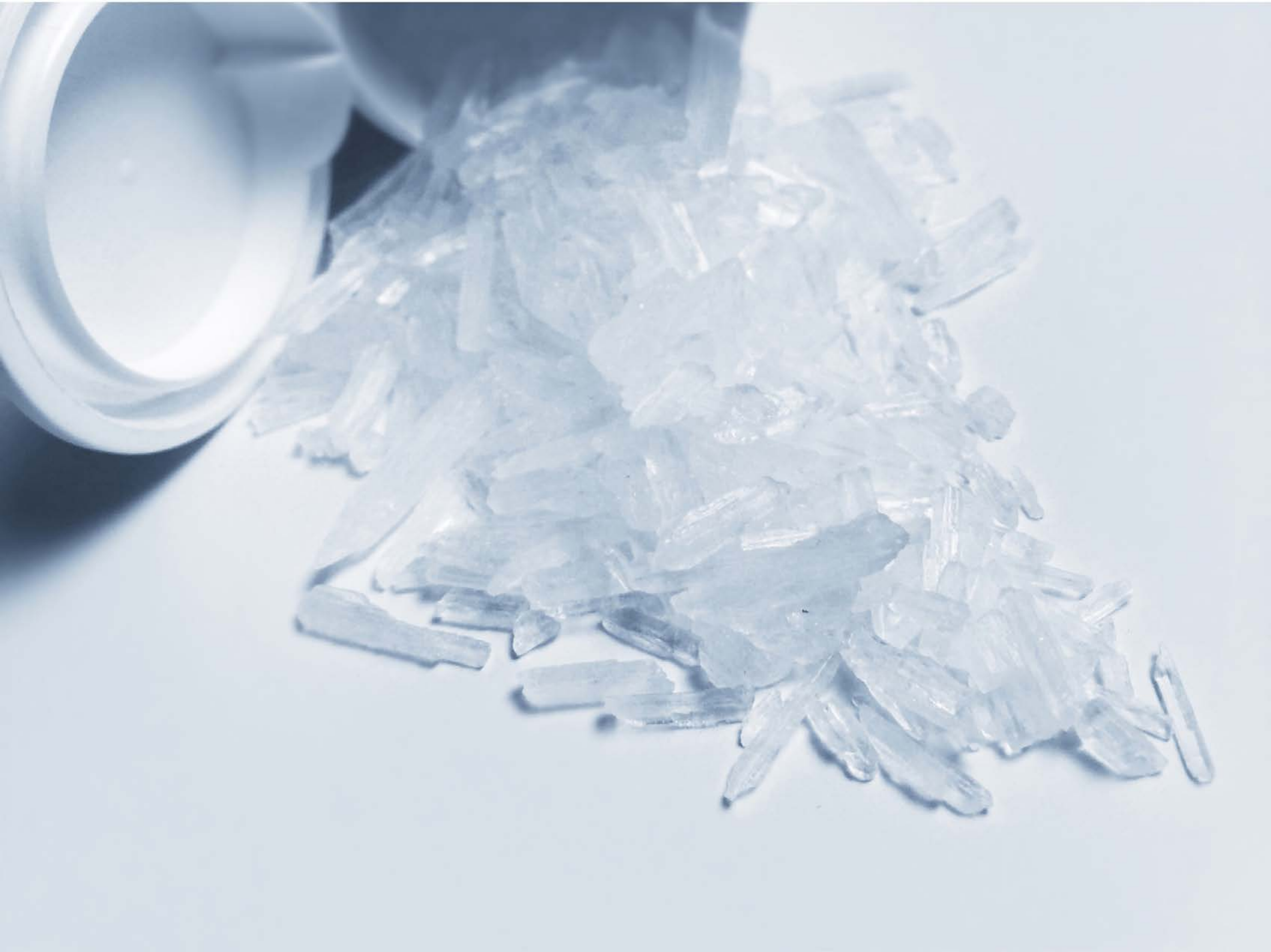

| Category | Craving (0-100) | Valence (1-9) | Arousal (1-9) | Typicality (0-100) | Relatedness | | | | | HSV | | |
|---|---|---|---|---|---|---|---|---|---|---|---|---|
| | | | | | Meth | Opioid | Both | Neither | MethToOpioid | Hue | Saturation | Value |
| Meth | 94.14 (11.19) | 4.29 (2.275) | 4.93 (2.87) | 94.89 (10.35) | 1 | 0 | 0 | 0 | -1 | 0.587 (0.022) | 0.101 (0.076) | 0.869 (0.105) |

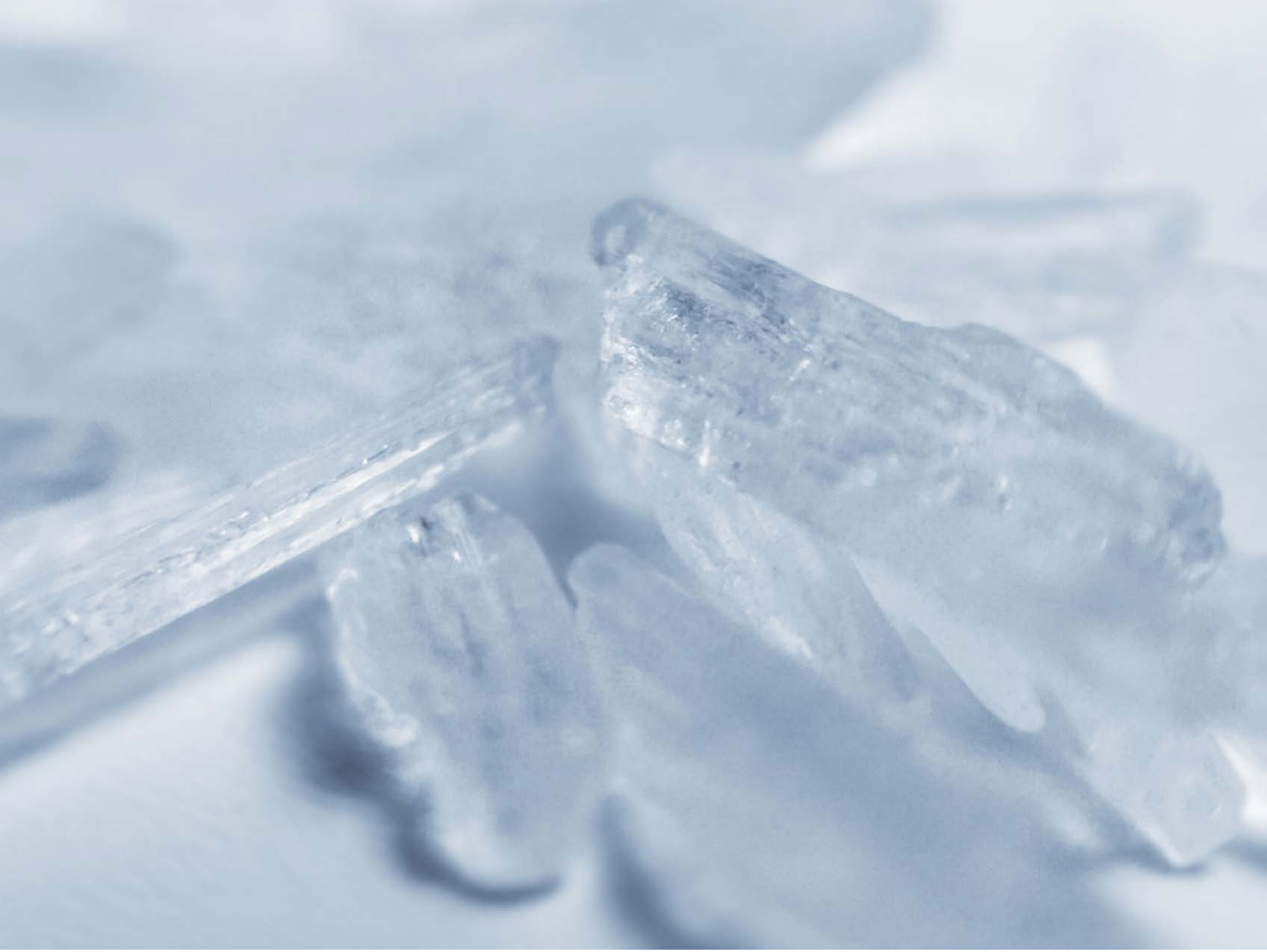

| Category | Craving (0-100) | Valence (1-9) | Arousal (1-9) | Typicality (0-100) | Relatedness | | | | | HSV | | |
|---|---|---|---|---|---|---|---|---|---|---|---|---|
| | | | | | Meth | Opioid | Both | Neither | MethToOpioid | Hue | Saturation | Value |
| Meth | 89.96 (20.59) | 4.29 (2.209) | 4.64 (2.86) | 91.71 (20.62) | 1 | 0 | 0 | 0 | -1 | 0.584 (0.006) | 0.148 (0.054) | 0.832 (0.057) |

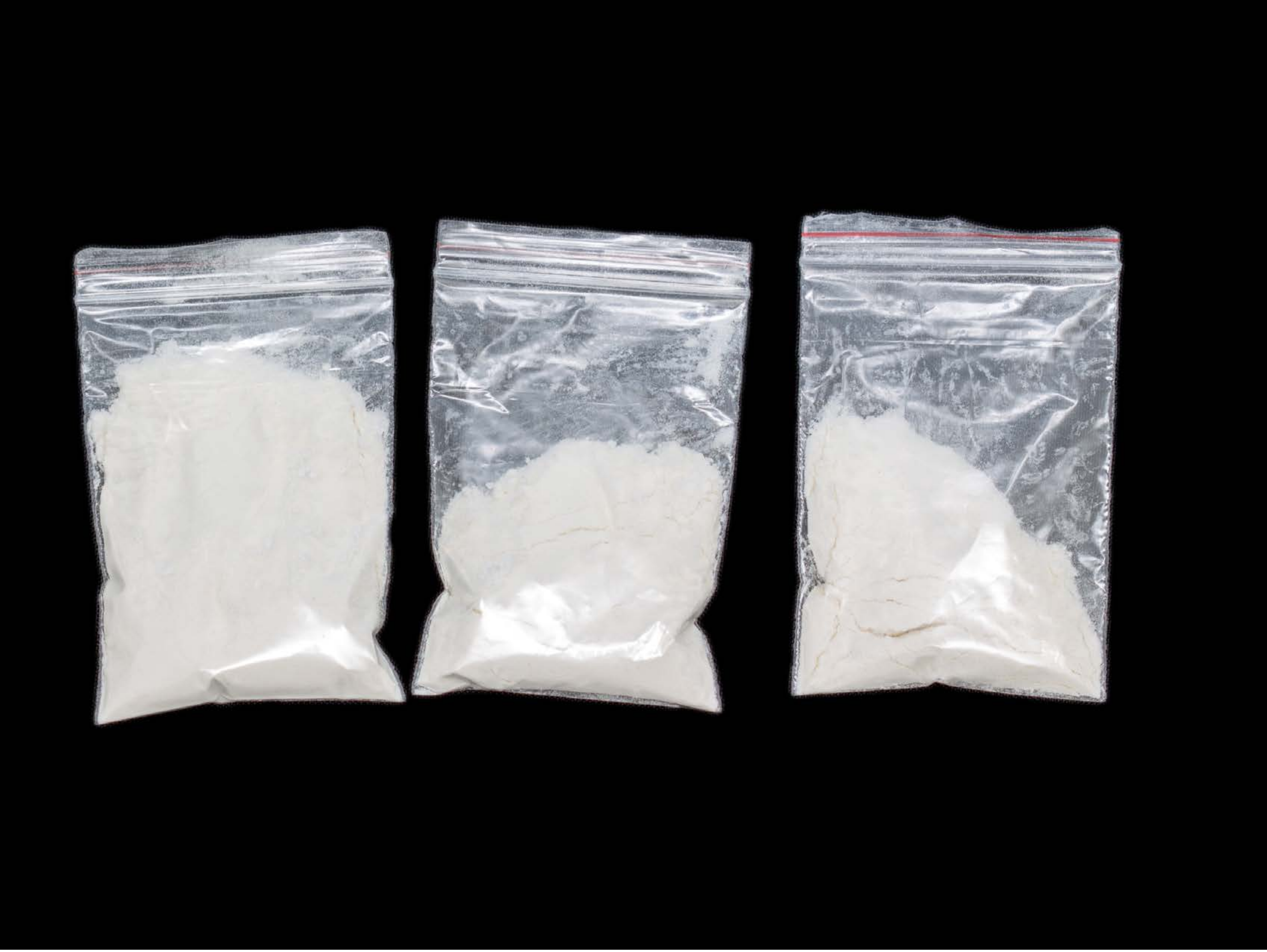

| Category | Craving (0-100) | Valence (1-9) | Arousal (1-9) | Typicality (0-100) | Relatedness | | | | | HSV | | |
|---|---|---|---|---|---|---|---|---|---|---|---|---|
| | | | | | Meth | Opioid | Both | Neither | MethToOpioid | Hue | Saturation | Value |
| Meth | 88.11 (18.25) | 4.41 (2.08) | 4.67 (2.79) | 88.15 (19.19) | 0.370 | 0.222 | 0.407 | 0 | -0.148 | 0.148 (0.261) | 0.024 (0.07) | 0.283 (0.383) |

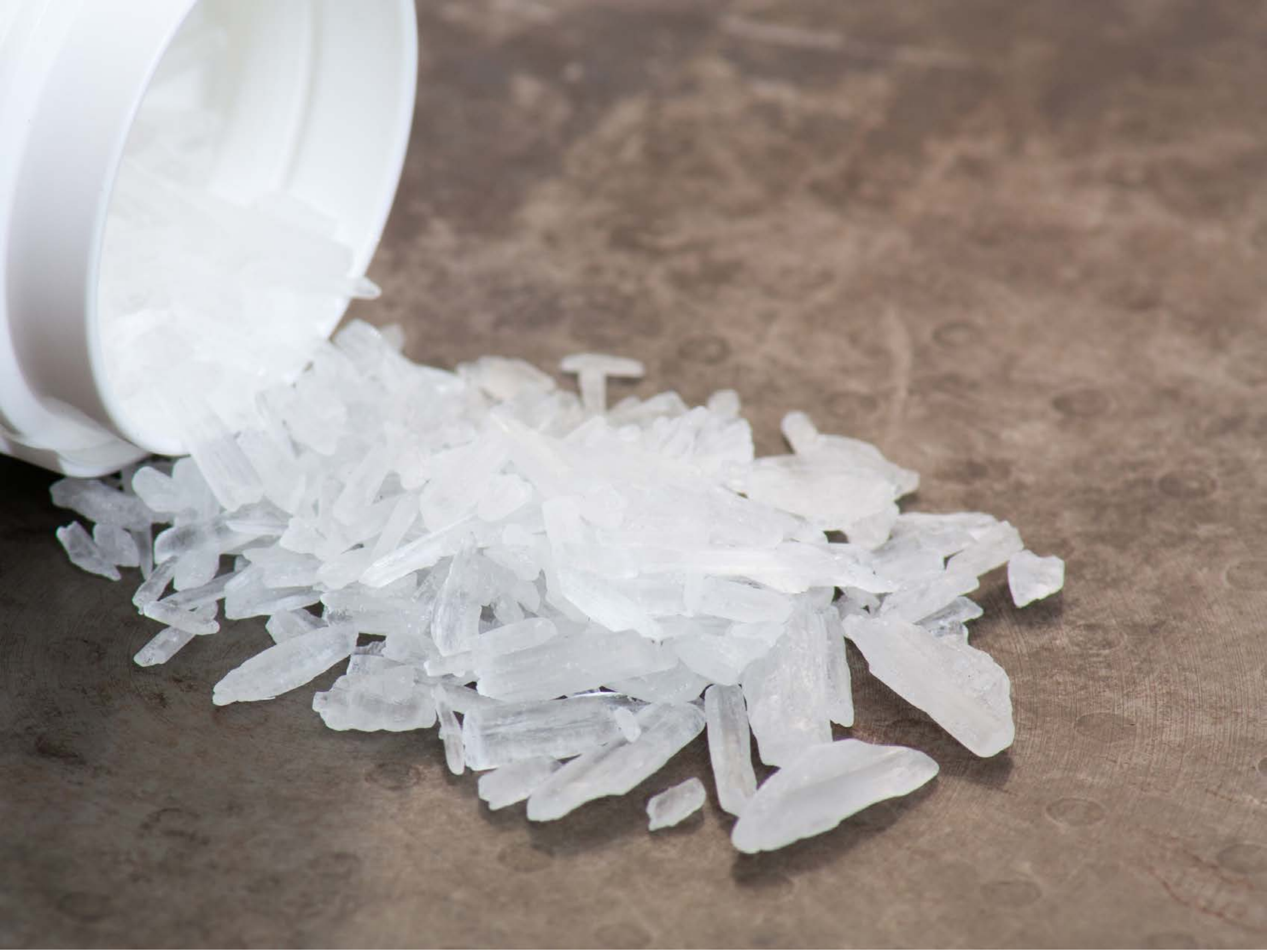

| Category | Craving (0-100) | Valence (1-9) | Arousal (1-9) | Typicality (0-100) | Relatedness | | | | | HSV | | |
|---|---|---|---|---|---|---|---|---|---|---|---|---|
| | | | | | Meth | Opioid | Both | Neither | MethToOpioid | Hue | Saturation | Value |
| Meth | 87.89 (16.86) | 4.41 (1.947) | 4.67 (2.83) | 87.3 (18.22) | 1 | 0 | 0 | 0 | -1 | 0.269 (0.276) | 0.152 (0.099) | 0.631 (0.195) |

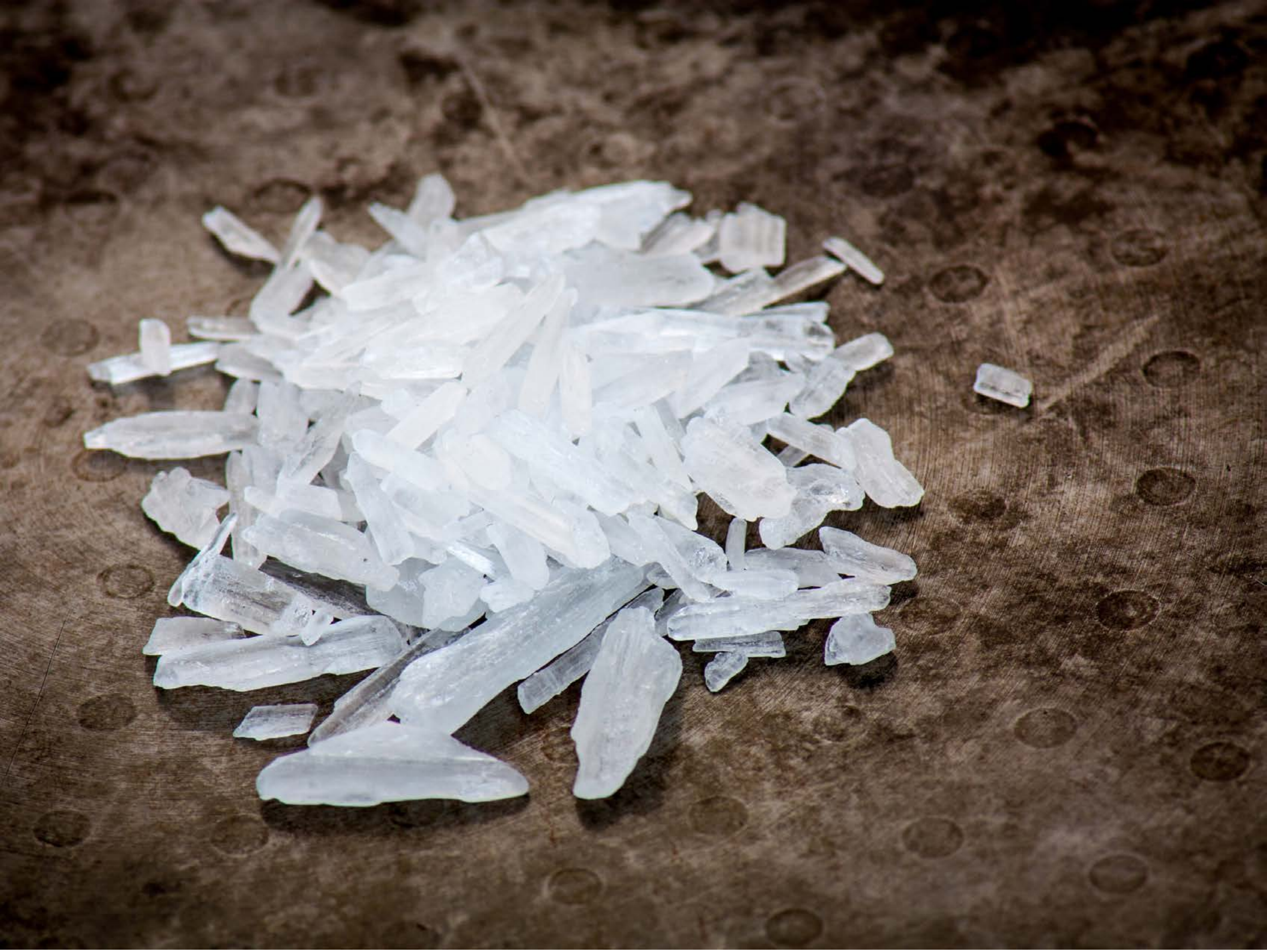

| Category | Craving (0-100) | Valence (1-9) | Arousal (1-9) | Typicality (0-100) | Relatedness | | | | | HSV | | |
|---|---|---|---|---|---|---|---|---|---|---|---|---|
| | | | | | Meth | Opioid | Both | Neither | MethToOpioid | Hue | Saturation | Value |
| Meth | 88.41 (16.64) | 4.48 (2.007) | 4.93 (2.8) | 88.96 (17.53) | 1 | 0 | 0 | 0 | -1 | 0.228 (0.255) | 0.332 (0.227) | 0.462 (0.251) |

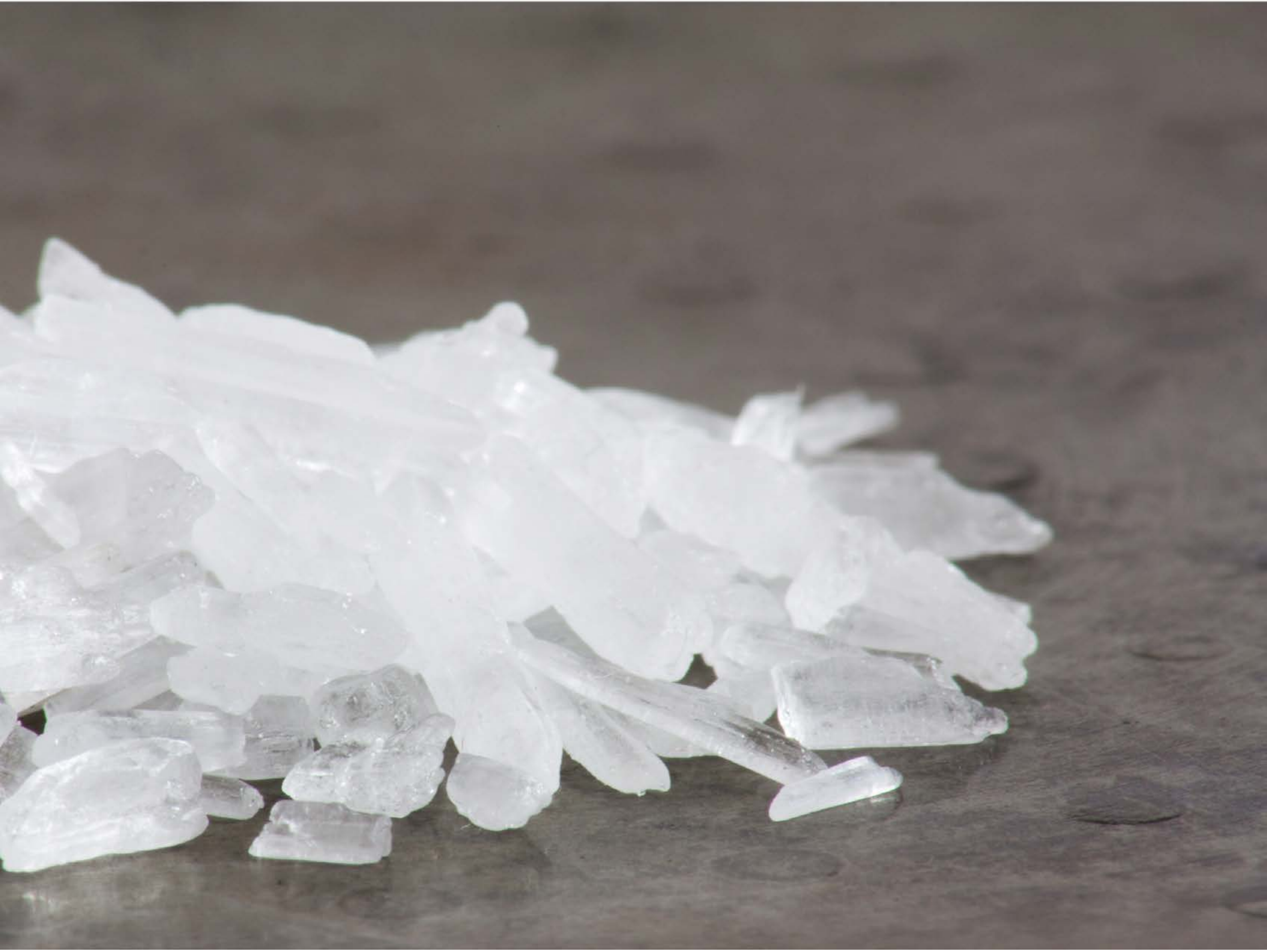

| Category | Craving (0-100) | Valence (1-9) | Arousal (1-9) | Typicality (0-100) | Relatedness | | | | | HSV | | |
|---|---|---|---|---|---|---|---|---|---|---|---|---|
| | | | | | Meth | Opioid | Both | Neither | MethToOpioid | Hue | Saturation | Value |
| Meth | 88.32 (23.6) | 4.25 (2.048) | 4.82 (2.93) | 90.11 (20.42) | 0.964 | 0 | 0 | 0.036 | -0.964 | 0.301 (0.301) | 0.078 (0.049) | 0.581 (0.209) |

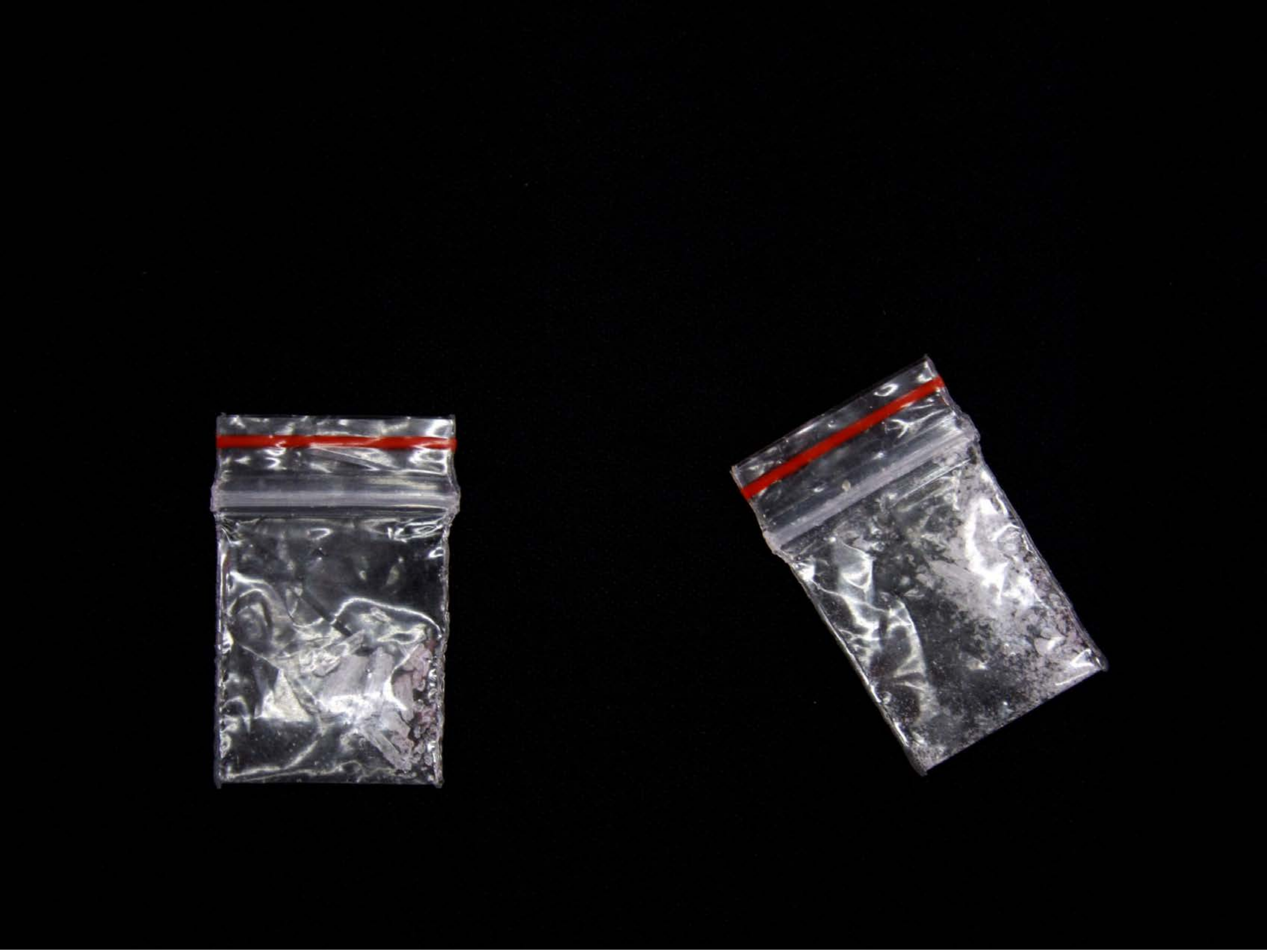

| Category | Craving (0-100) | Valence (1-9) | Arousal (1-9) | Typicality (0-100) | Relatedness | | | | | HSV | | |
|---|---|---|---|---|---|---|---|---|---|---|---|---|
| | | | | | Meth | Opioid | Both | Neither | MethToOpioid | Hue | Saturation | Value |
| Meth | 88.89 (16.67) | 4.68 (2.038) | 4.68 (2.87) | 92 (14.01) | 0.75 | 0.071 | 0.179 | 0 | -0.679 | 0.278 (0.352) | 0.264 (0.39) | 0.069 (0.177) |

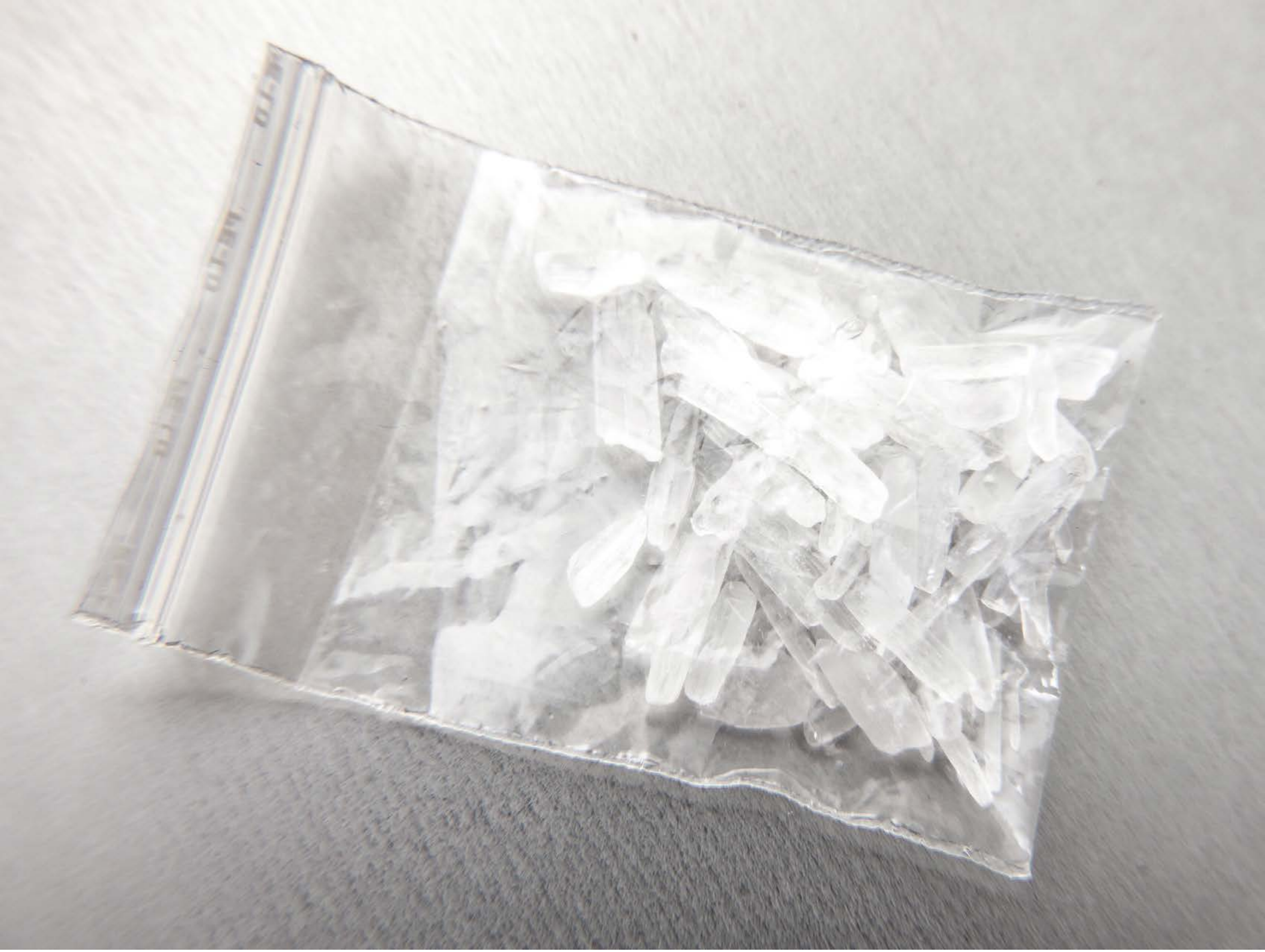

| Category | Craving (0-100) | Valence (1-9) | Arousal (1-9) | Typicality (0-100) | Relatedness | | | | | HSV | | |
|---|---|---|---|---|---|---|---|---|---|---|---|---|
| | | | | | Meth | Opioid | Both | Neither | MethToOpioid | Hue | Saturation | Value |
| Meth | 87.19 (18.32) | 4.37 (1.801) | 4.67 (2.63) | 84.81 (19.42) | 1 | 0 | 0 | 0 | -1 | 0.243 (0.31) | 0.022 (0.016) | 0.792 (0.146) |

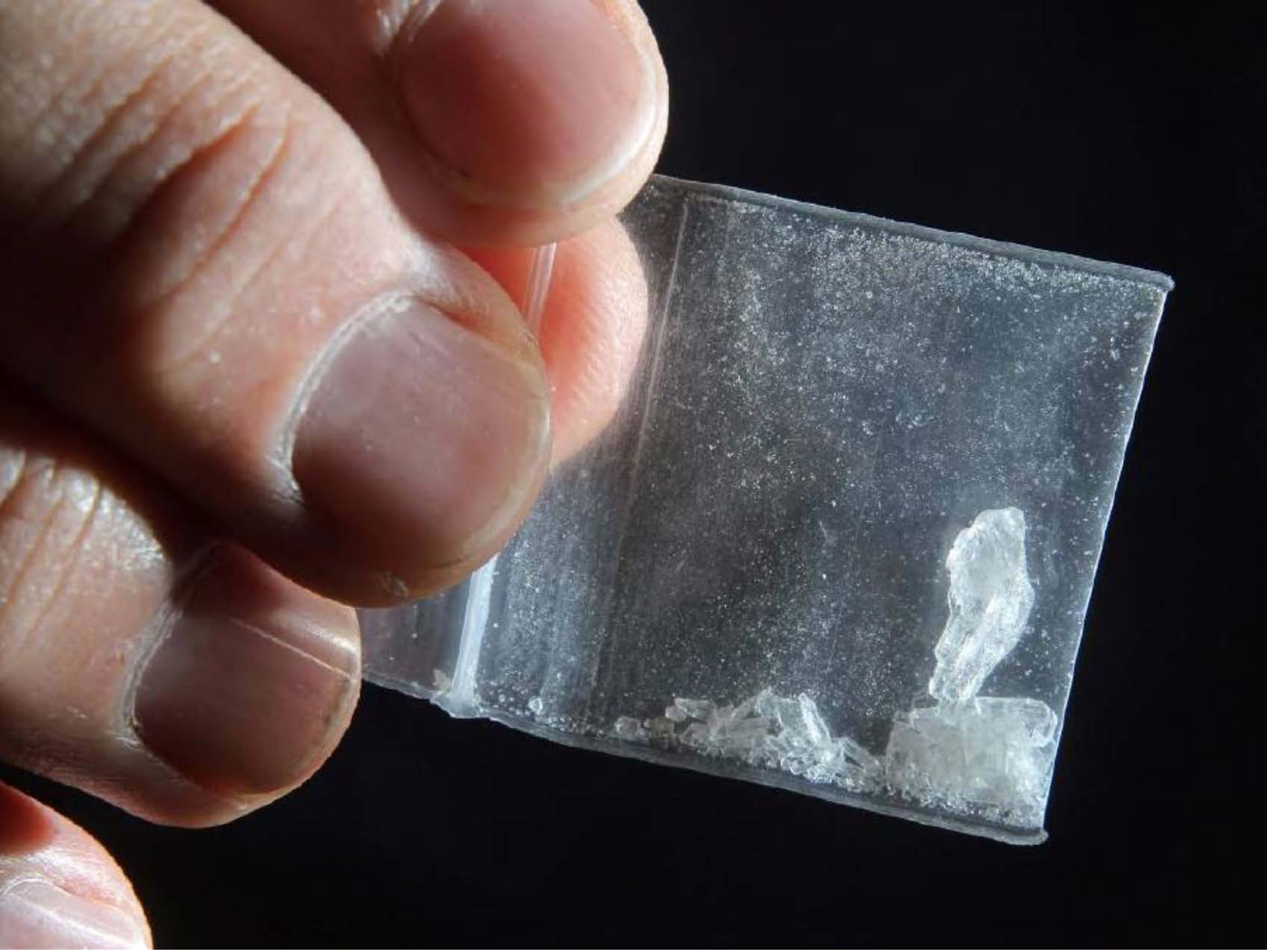

| Category | Craving (0-100) | Valence (1-9) | Arousal (1-9) | Typicality (0-100) | Relatedness | | | | | HSV | | |
|---|---|---|---|---|---|---|---|---|---|---|---|---|
| | | | | | Meth | Opioid | Both | Neither | MethToOpioid | Hue | Saturation | Value |
| Meth Hand | 90.5 (15.68) | 4.57 (1.971) | 4.64 (2.77) | 95.07 (10.18) | 1 | 0 | 0 | 0 | -1 | 0.411 (0.32) | 0.302 (0.197) | 0.345 (0.273) |

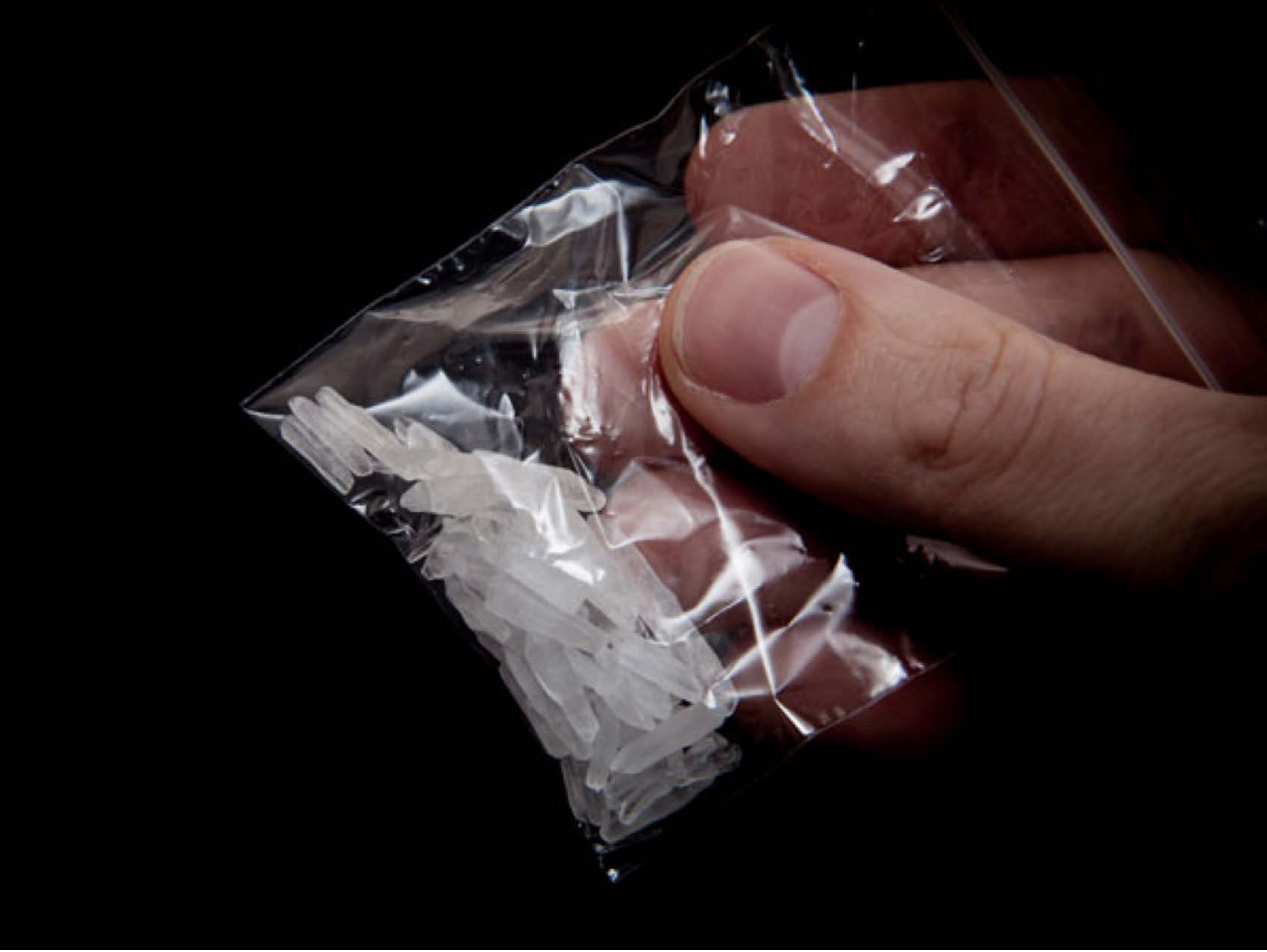

| Category | Craving (0-100) | Valence (1-9) | Arousal (1-9) | Typicality (0-100) | Relatedness | | | | | HSV | | |
|---|---|---|---|---|---|---|---|---|---|---|---|---|
| | | | | | Meth | Opioid | Both | Neither | MethToOpioid | Hue | Saturation | Value |
| Meth Hand | 89.19 (15.56) | 4.59 (2.206) | 4.74 (2.84) | 89.11 (17.86) | 1 | 0 | 0 | 0 | -1 | 0.188 (0.355) | 0.164 (0.274) | 0.158 (0.236) |

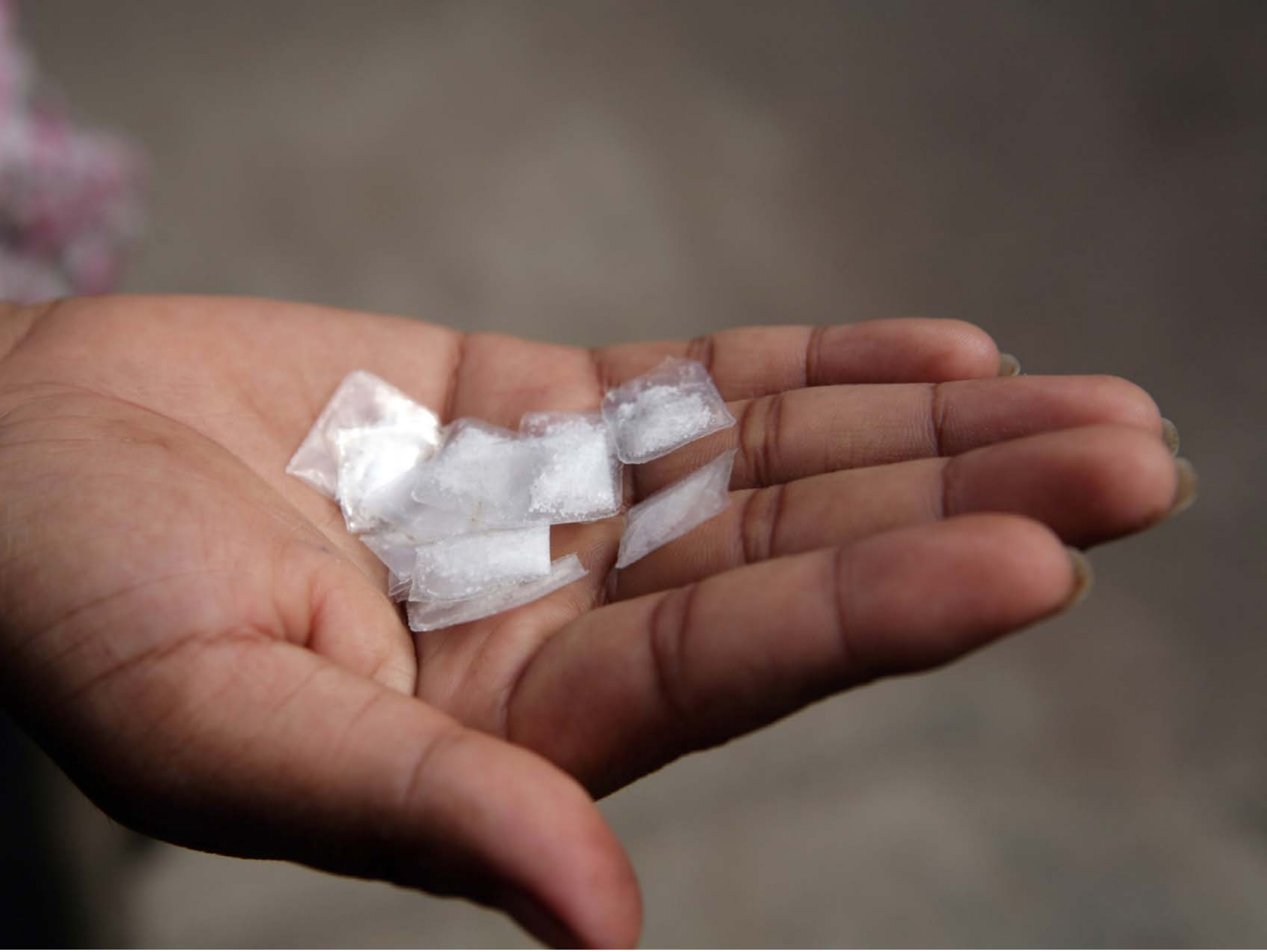

| Category | Craving (0-100) | Valence (1-9) | Arousal (1-9) | Typicality (0-100) | Relatedness | | | | | HSV | | |
|---|---|---|---|---|---|---|---|---|---|---|---|---|
| | | | | | Meth | Opioid | Both | Neither | MethToOpioid | Hue | Saturation | Value |
| Meth Hand | 93.93 (10.94) | 4.43 (2.098) | 4.79 (2.87) | 94.14 (10.12) | 0.964 | 0 | 0.036 | 0 | -0.964 | 0.118 (0.215) | 0.283 (0.169) | 0.442 (0.176) |

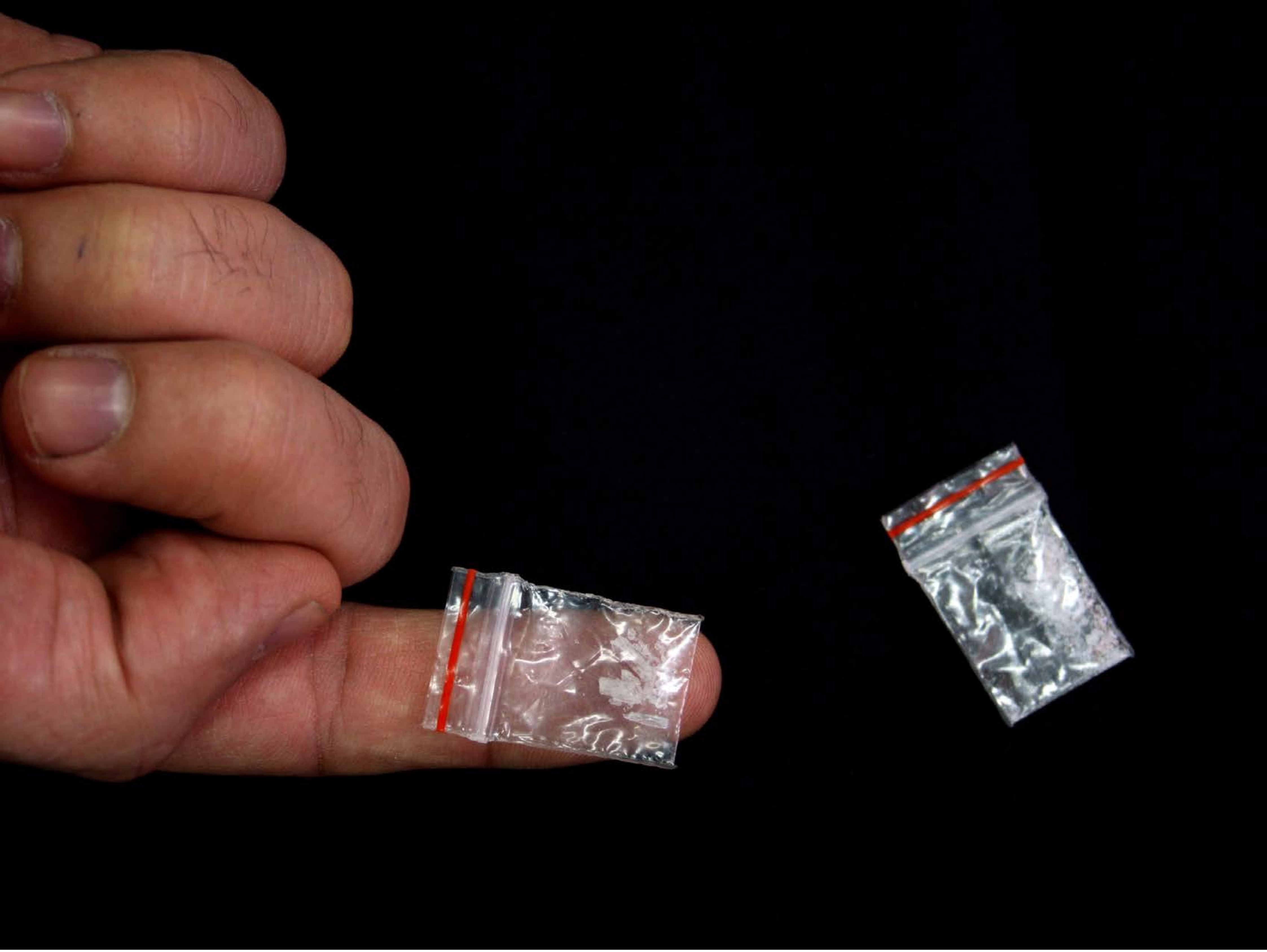

| Category | Craving (0-100) | Valence (1-9) | Arousal (1-9) | Typicality (0-100) | Relatedness | | | | | HSV | | |
|---|---|---|---|---|---|---|---|---|---|---|---|---|
| | | | | | Meth | Opioid | Both | Neither | MethToOpioid | Hue | Saturation | Value |
| Meth Hand | 87.93 (16.71) | 4.63 (1.904) | 4.63 (2.84) | 84.52 (20.22) | 0.889 | 0 | 0.111 | 0 | -0.889 | 0.238 (0.329) | 0.393 (0.39) | 0.155 (0.251) |

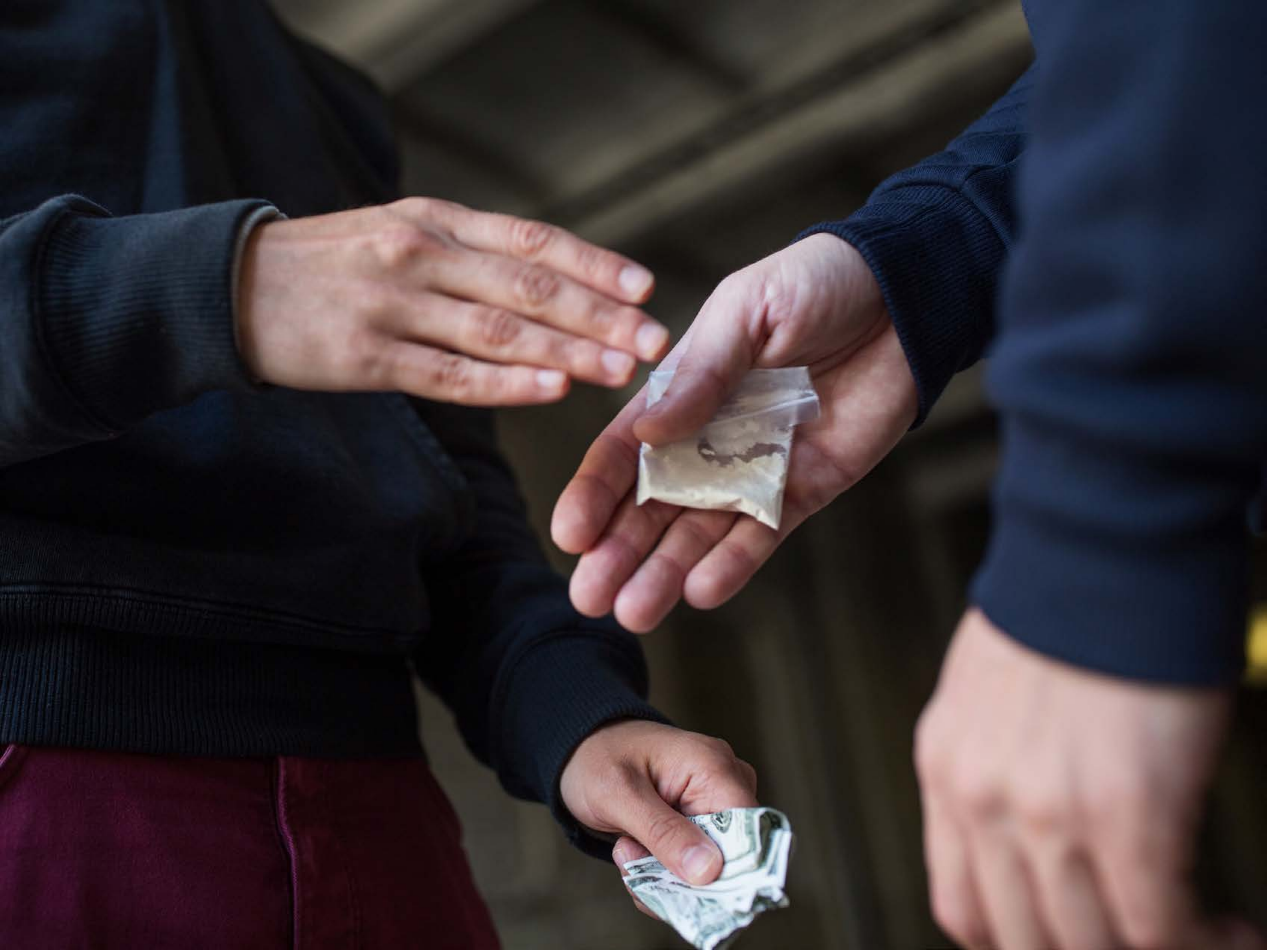

| Category | Craving (0-100) | Valence (1-9) | Arousal (1-9) | Typicality (0-100) | Relatedness | | | | | HSV | | |
|---|---|---|---|---|---|---|---|---|---|---|---|---|
| | | | | | Meth | Opioid | Both | Neither | MethToOpioid | Hue | Saturation | Value |
| Meth Hand | 85.15 (20.99) | 4.41 (1.947) | 4.56 (2.76) | 88.26 (16.68) | 0.259 | 0.296 | 0.444 | 0 | 0.037 | 0.432 (0.338) | 0.372 (0.202) | 0.257 (0.237) |

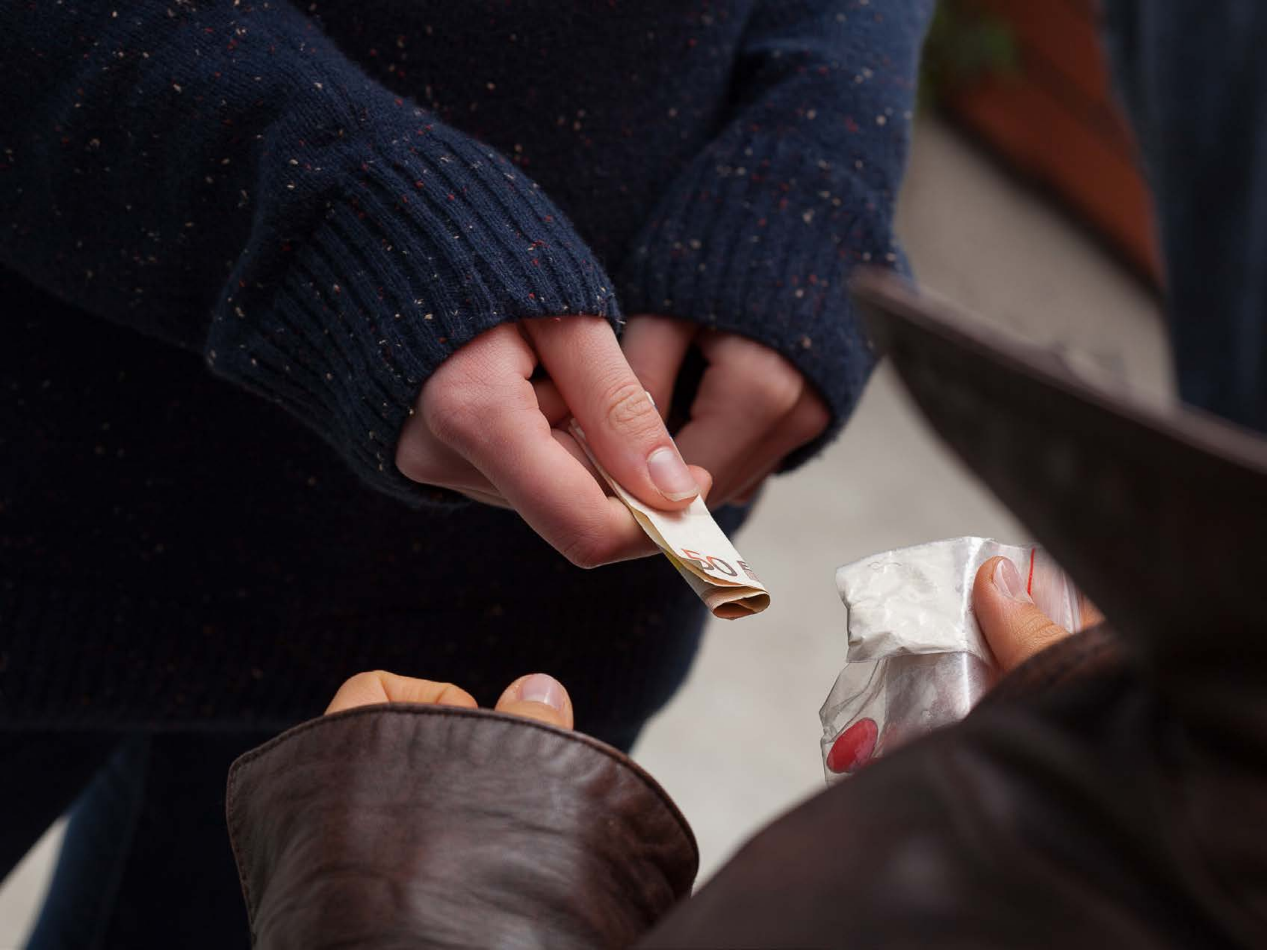


| Category | Craving (0-100) | Valence (1-9) | Arousal (1-9) | Typicality (0-100) | Relatedness | | | | | HSV | | |
|---|---|---|---|---|---|---|---|---|---|---|---|---|
| | | | | | Meth | Opioid | Both | Neither | MethToOpioid | Hue | Saturation | Value |
| Meth Hand | 81.82 (19.29) | 4.46 (1.551) | 4.54 (2.67) | 88.86 (15.99) | 0.107 | 0.036 | 0.857 | 0 | -0.071 | 0.462 (0.357) | 0.368 (0.169) | 0.223 (0.226) |

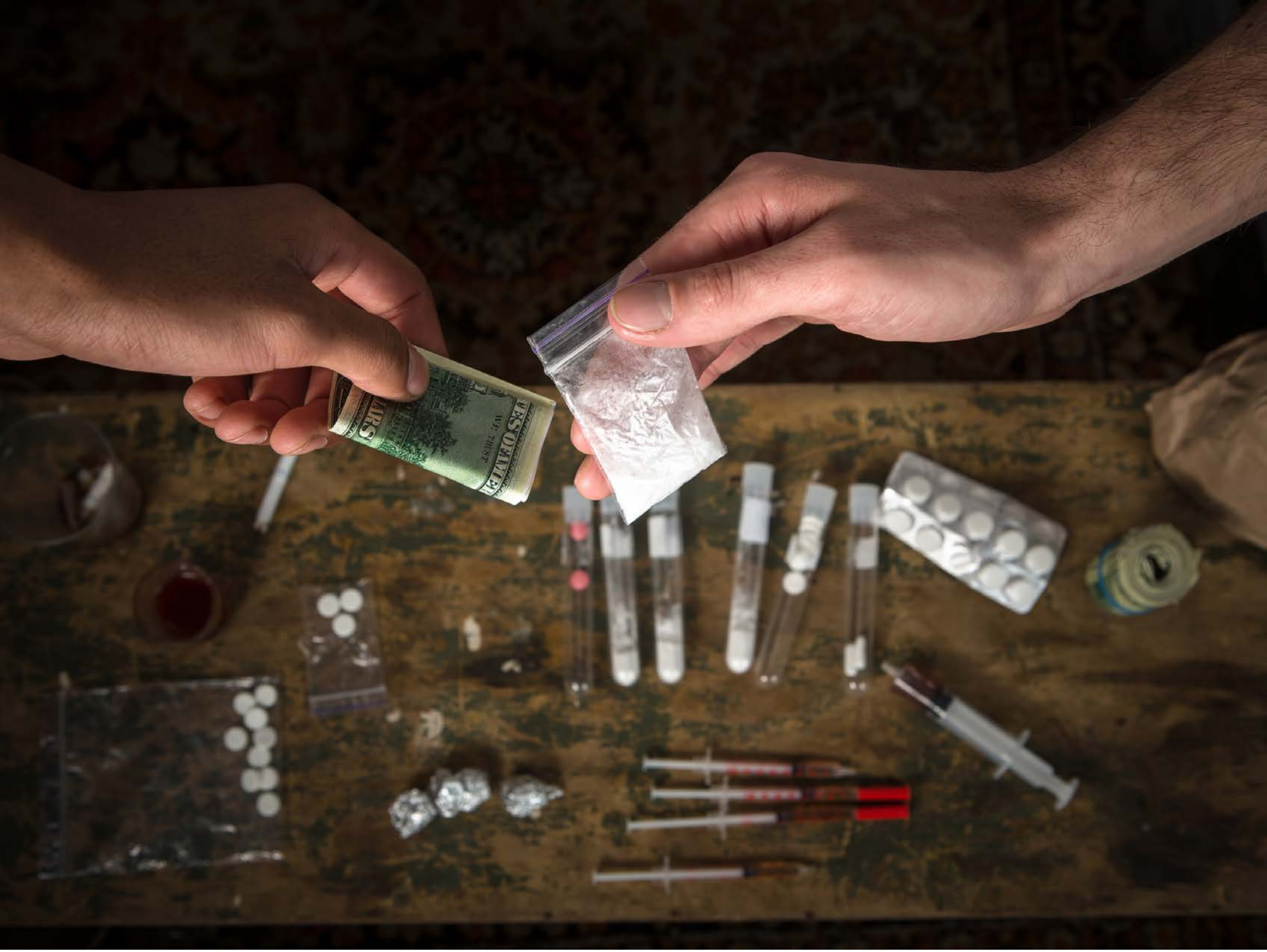

| Category | Craving (0-100) | Valence (1-9) | Arousal (1-9) | Typicality (0-100) | Relatedness | | | | | HSV | | |
|---|---|---|---|---|---|---|---|---|---|---|---|---|
| | | | | | Meth | Opioid | Both | Neither | MethToOpioid | Hue | Saturation | Value |
| Meth Hand | 88.96 (16.49) | 4.89 (1.888) | 4.96 (2.7) | 87.56 (18.67) | 0.074 | 0.259 | 0.667 | 0 | 0.185 | 0.092 (0.111) | 0.511 (0.212) | 0.268 (0.214) |

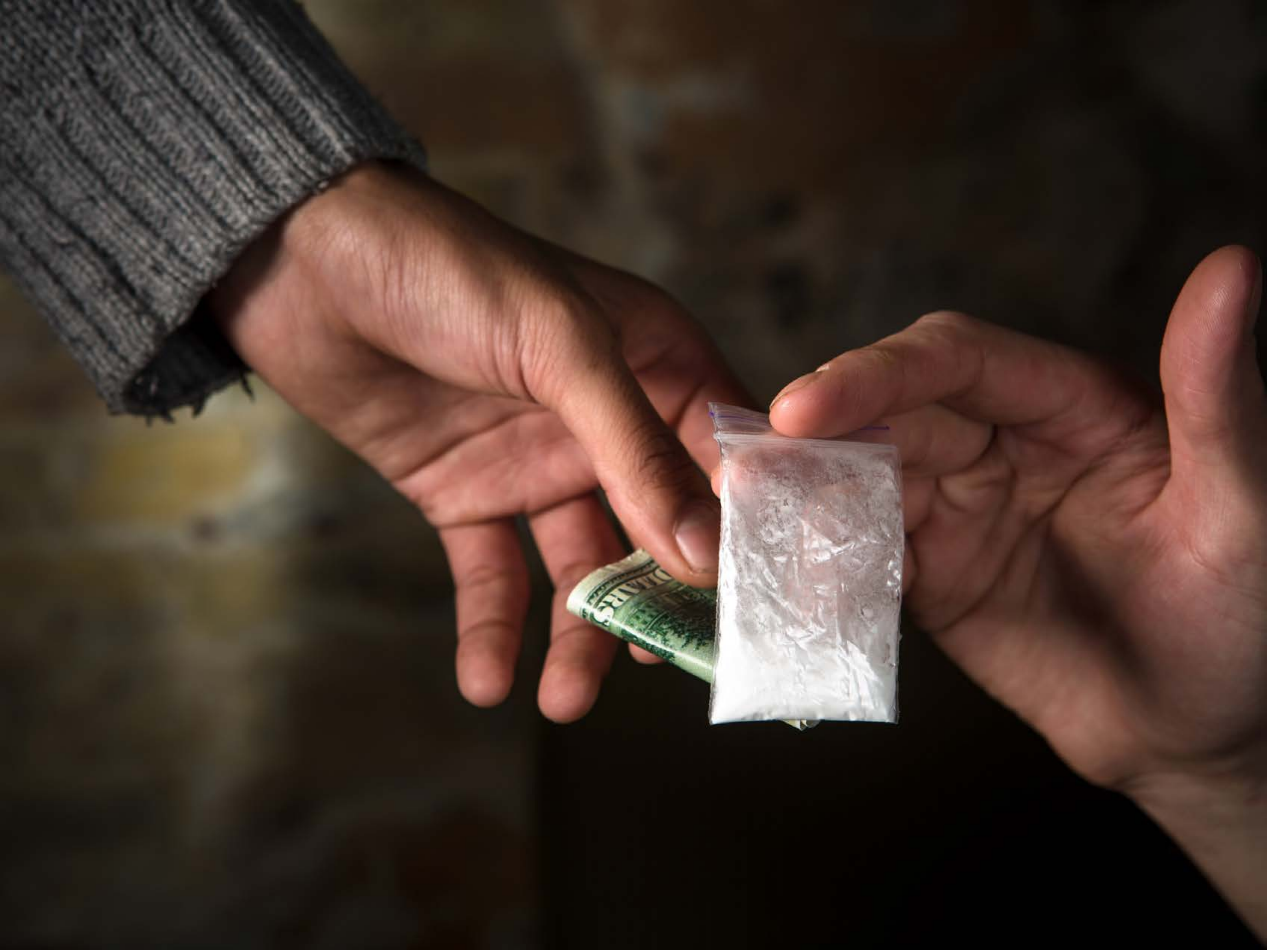

| | | | | | Relatedness | | | | | HSV | | |
|---|---|---|---|---|---|---|---|---|---|---|---|---|
| **Category** | **Craving (0-100)** | **Valence (1-9)** | **Arousal (1-9)** | **Typicality (0-100)** | Meth | Opioid | Both | Neither | MethToOpioid | Hue | Saturation | Value |
| Meth Hand | 82.63 (20.33) | 4.56 (1.717) | 4.26 (2.54) | 85.93 (19.59) | 0.333 | 0.185 | 0.481 | 0 | -0.148 | 0.106 (0.161) | 0.439 (0.234) | 0.27 (0.227) |

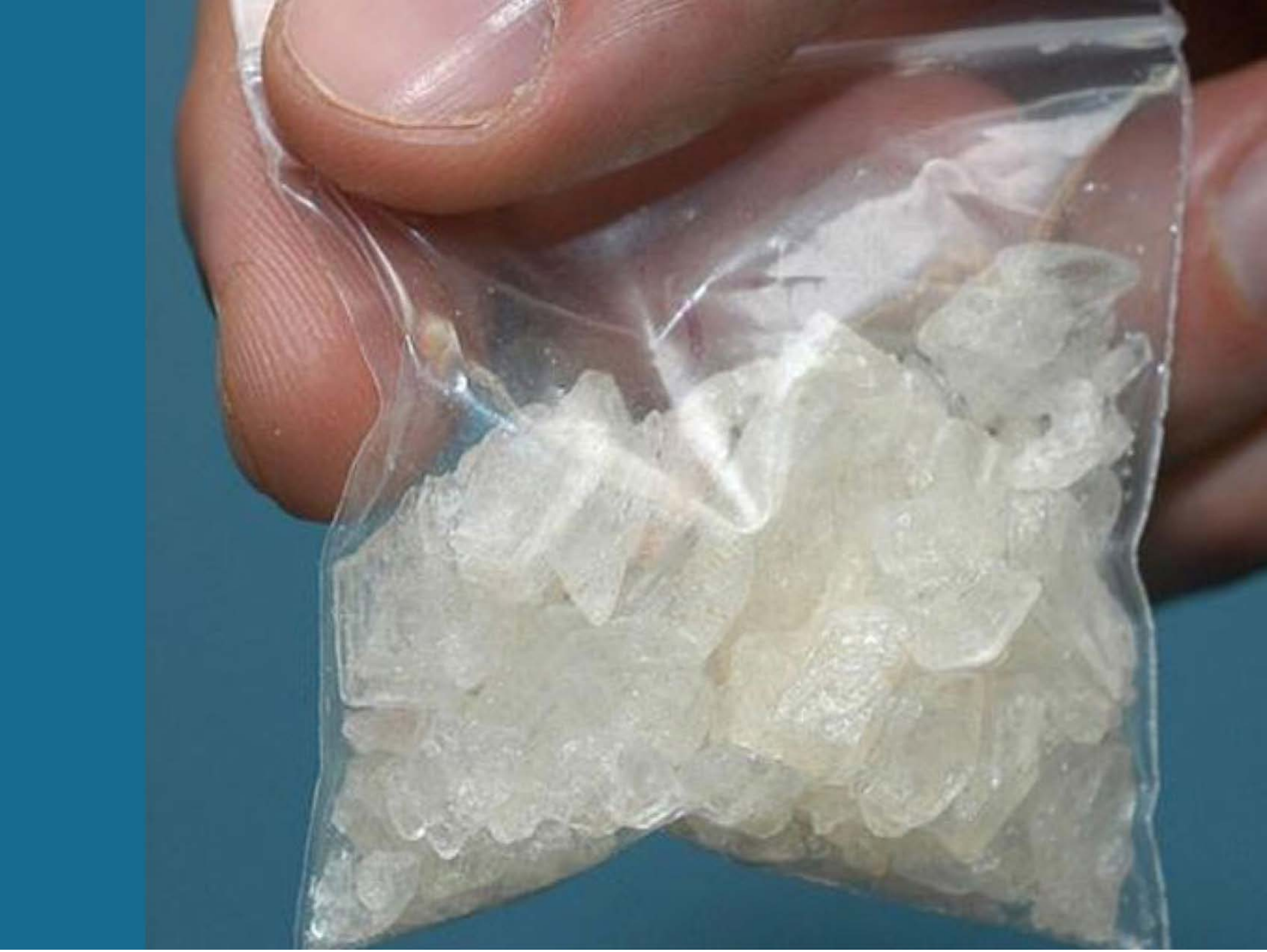

| Category | Craving (0-100) | Valence (1-9) | Arousal (1-9) | Typicality (0-100) | Relatedness | | | | | HSV | | |
|---|---|---|---|---|---|---|---|---|---|---|---|---|
| | | | | | Meth | Opioid | Both | Neither | MethToOpioid | Hue | Saturation | Value |
| Meth Hand | 88.21 (22.57) | 4.46 (1.934) | 4.64 (2.95) | 89.39 (22.11) | 1 | 0 | 0 | 0 | -1 | 0.321 (0.275) | 0.304 (0.273) | 0.58 (0.156) |

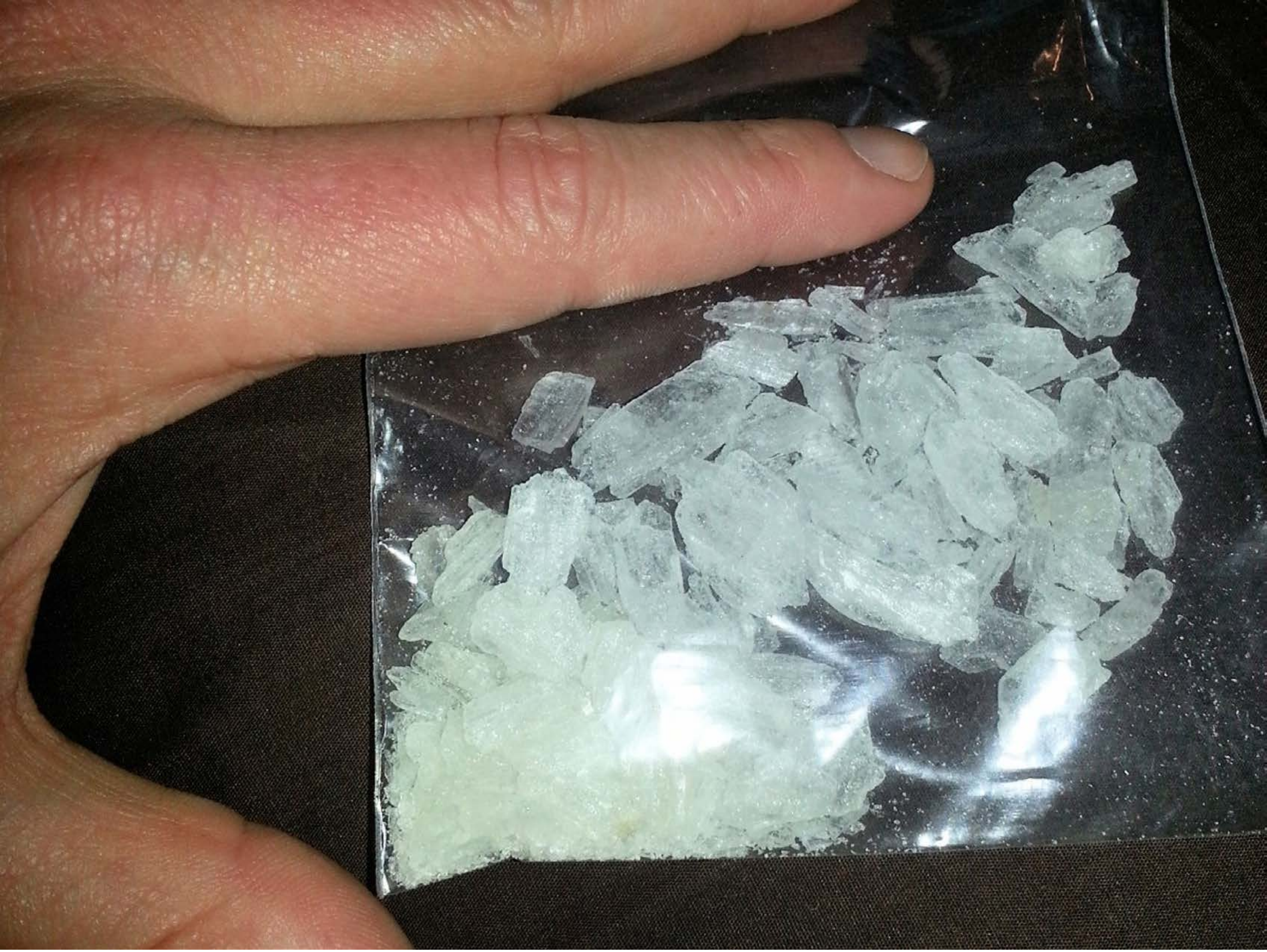

| Category | Craving (0-100) | Valence (1-9) | Arousal (1-9) | Typicality (0-100) | Relatedness | | | | | HSV | | |
|---|---|---|---|---|---|---|---|---|---|---|---|---|
| | | | | | Meth | Opioid | Both | Neither | MethToOpioid | Hue | Saturation | Value |
| Meth Hand | 90.61 (14.99) | 4.68 (1.765) | 5.04 (2.65) | 93.5 (13.82) | 1 | 0 | 0 | 0 | -1 | 0.263 (0.221) | 0.277 (0.2) | 0.475 (0.285) |

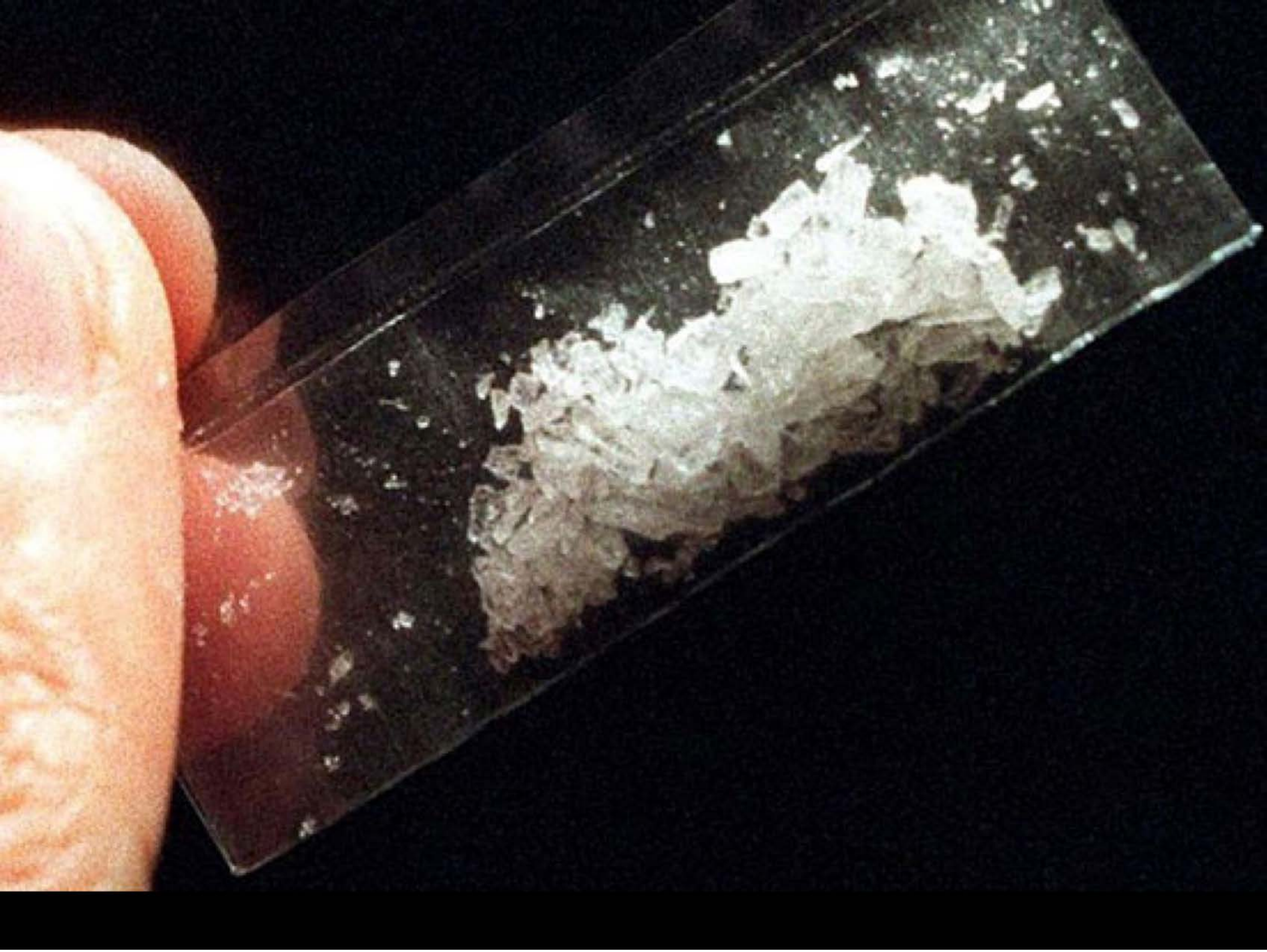

| Category | Craving (0-100) | Valence (1-9) | Arousal (1-9) | Typicality (0-100) | Relatedness | | | | | HSV | | |
|---|---|---|---|---|---|---|---|---|---|---|---|---|
| | | | | | Meth | Opioid | Both | Neither | MethToOpioid | Hue | Saturation | Value |
| Meth Hand | 91.71 (12.98) | 4.41 (2.062) | 4.89 (2.81) | 93.29 (11.22) | 1 | 0 | 0 | 0 | -1 | 0.404 (0.316) | 0.392 (0.279) | 0.291 (0.344) |

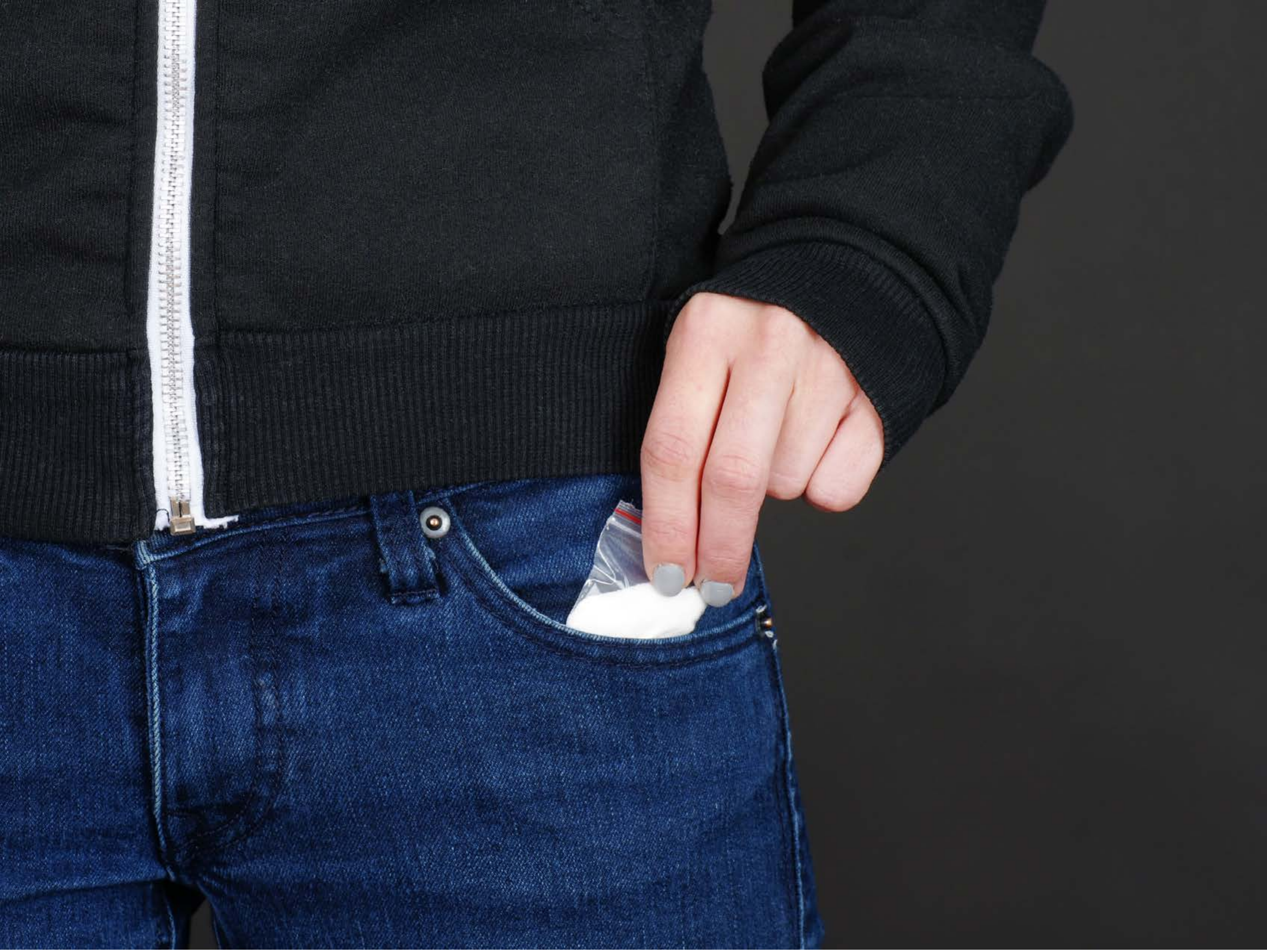

| Category | Craving (0-100) | Valence (1-9) | Arousal (1-9) | Typicality (0-100) | Relatedness | | | | | HSV | | |
|---|---|---|---|---|---|---|---|---|---|---|---|---|
| | | | | | Meth | Opioid | Both | Neither | MethToOpioid | Hue | Saturation | Value |
| Meth Hand | 82.56 (19.41) | 4.41 (1.647) | 4.19 (2.63) | 83.89 (19.41) | 0.296 | 0.222 | 0.481 | 0 | -0.074 | 0.505 (0.275) | 0.29 (0.288) | 0.249 (0.193) |

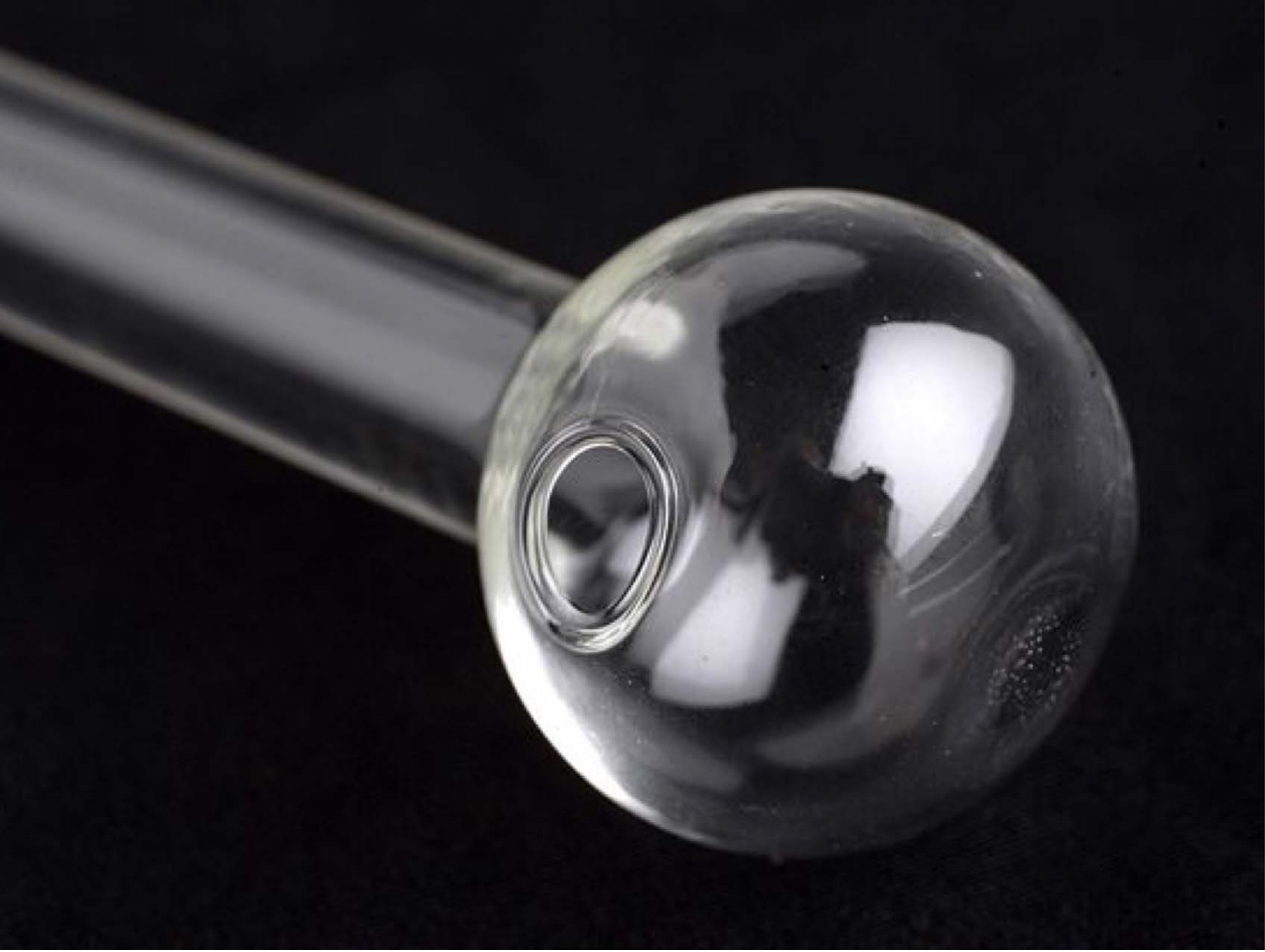

| Category | Craving (0-100) | Valence (1-9) | Arousal (1-9) | Typicality (0-100) | Relatedness | | | | | HSV | | |
|---|---|---|---|---|---|---|---|---|---|---|---|---|
| | | | | | Meth | Opioid | Both | Neither | MethToOpioid | Hue | Saturation | Value |
| Meth Instrument | 84.89 (18.71) | 4.59 (1.782) | 4.22 (2.5) | 88.48 (17.27) | 0.889 | 0 | 0.111 | 0 | -0.889 | 0.709 (0.182) | 0.225 (0.178) | 0.192 (0.224) |

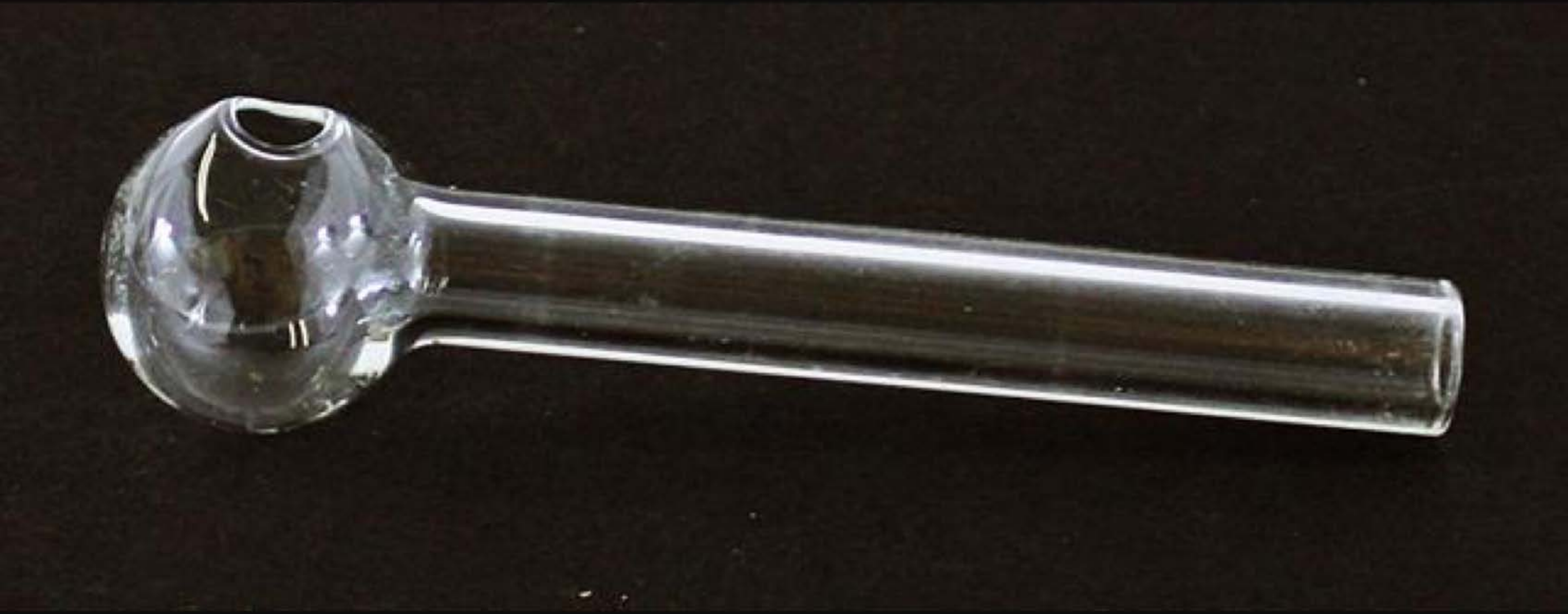

| Category | Craving (0-100) | Valence (1-9) | Arousal (1-9) | Typicality (0-100) | Relatedness | | | | | HSV | | |
|---|---|---|---|---|---|---|---|---|---|---|---|---|
| | | | | | Meth | Opioid | Both | Neither | MethToOpioid | Hue | Saturation | Value |
| Meth Instrument | 87.14 (22.33) | 4.11 (1.75) | 4.11 (2.71) | 91.64 (19.86) | 0.929 | 0 | 0.071 | 0 | -0.929 | 0.07 (0.142)) | 0.158 (0.174) | 0.122 (0.148) |

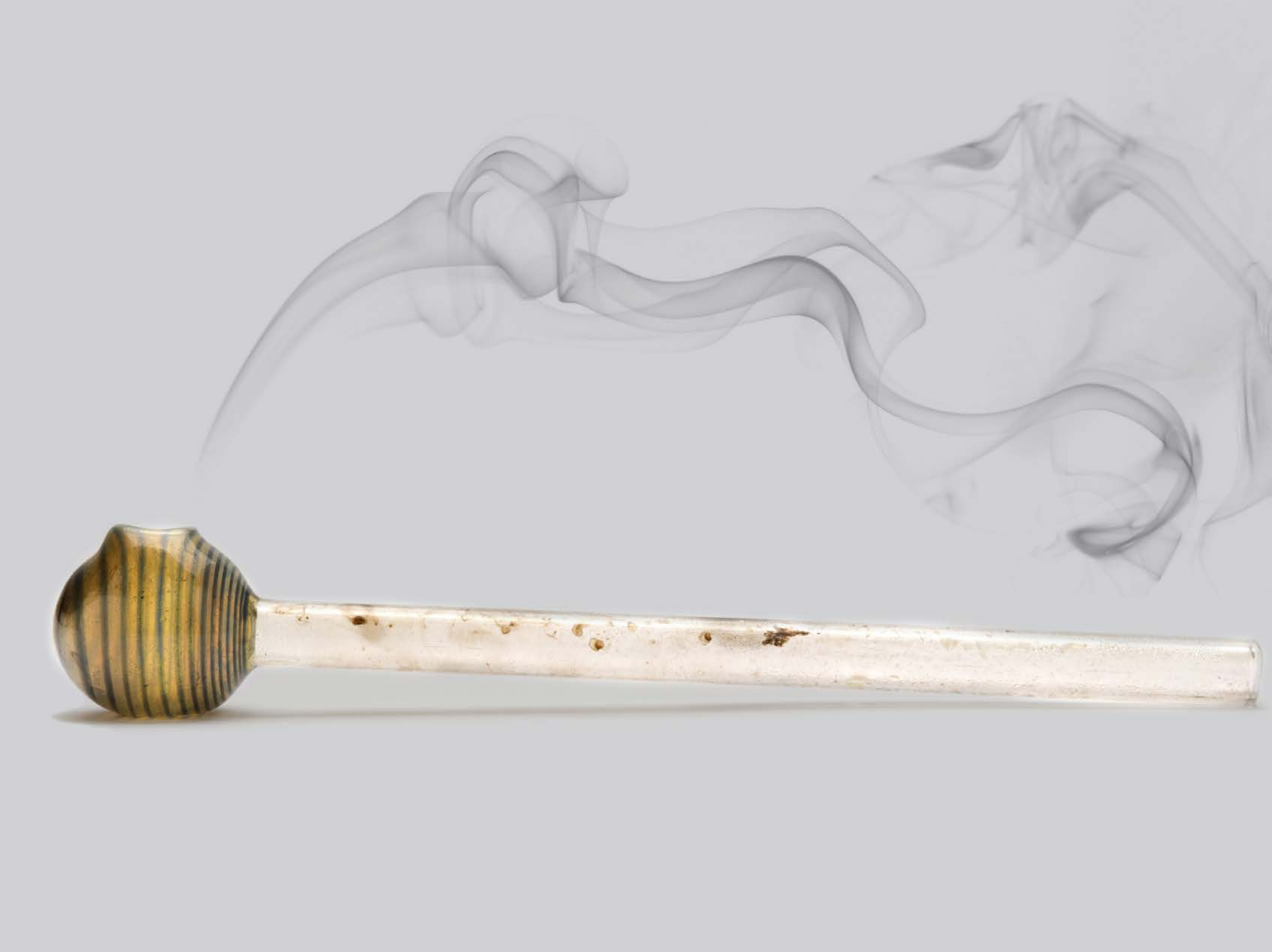

| Category | Craving (0-100) | Valence (1-9) | Arousal (1-9) | Typicality (0-100) | Relatedness | | | | | HSV | | |
|---|---|---|---|---|---|---|---|---|---|---|---|---|
| | | | | | Meth | Opioid | Both | Neither | MethToOpioid | Hue | Saturation | Value |
| Meth Instrument | 91.68 (13.61) | 4.29 (1.979) | 4.61 (2.87) | 92.64 (13.64) | 0.929 | 0 | 0.071 | 0 | -0.929 | 0.011 (0.04) | 0.019 (0.081) | 0.974 (0.081) |

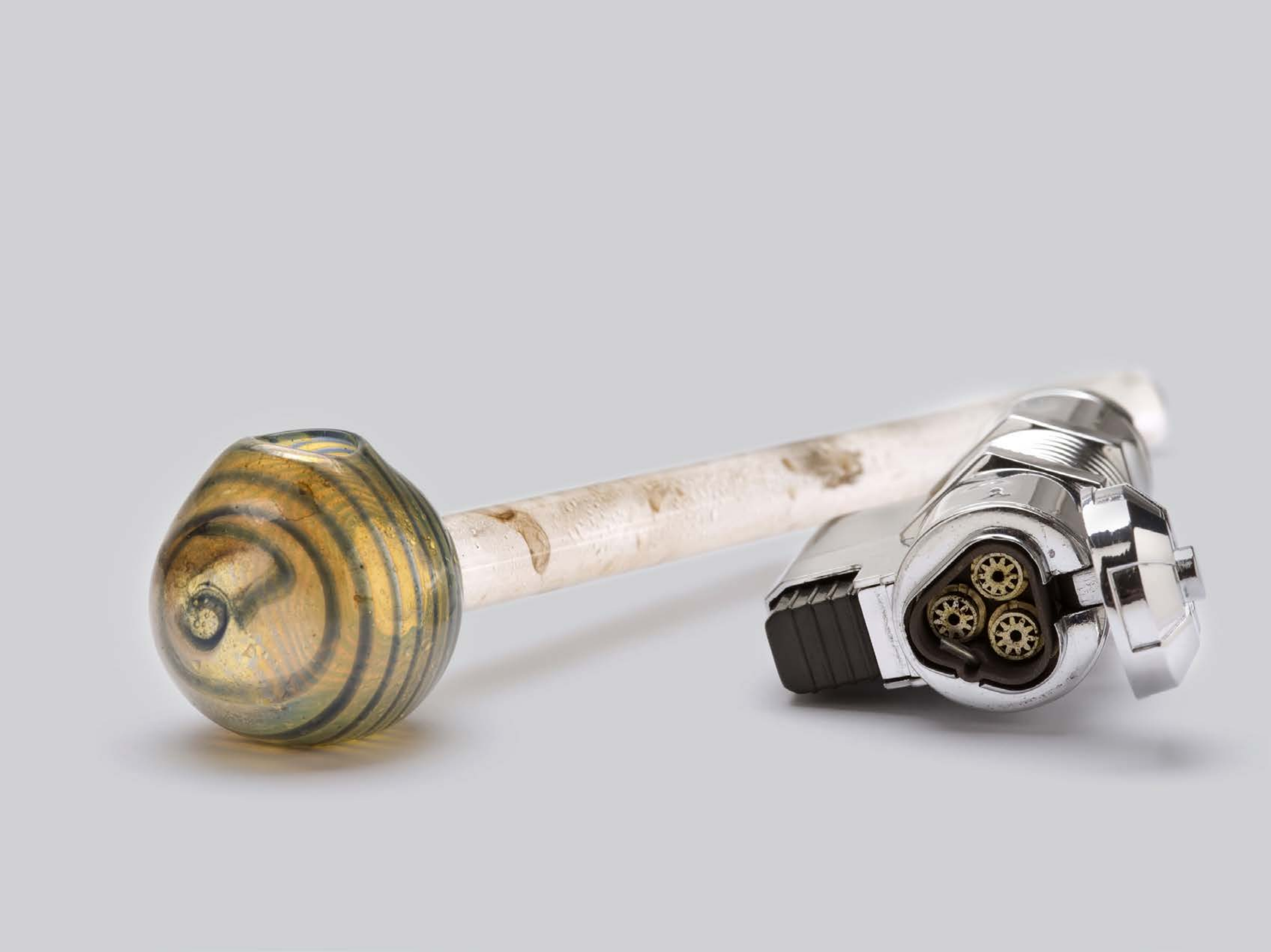

| Category | Craving (0-100) | Valence (1-9) | Arousal (1-9) | Typicality (0-100) | Relatedness | | | | | HSV | | |
|---|---|---|---|---|---|---|---|---|---|---|---|---|
| | | | | | Meth | Opioid | Both | Neither | MethToOpioid | Hue | Saturation | Value |
| Meth Instrument | 84.56 (19.19) | 4.3 (1.793) | 4.74 (2.65) | 86.67 (17.46) | 0.926 | 0 | 0.074 | 0 | -0.926 | 0.074 (0.214) | 0.042 (0.117) | 0.928 (0.177) |

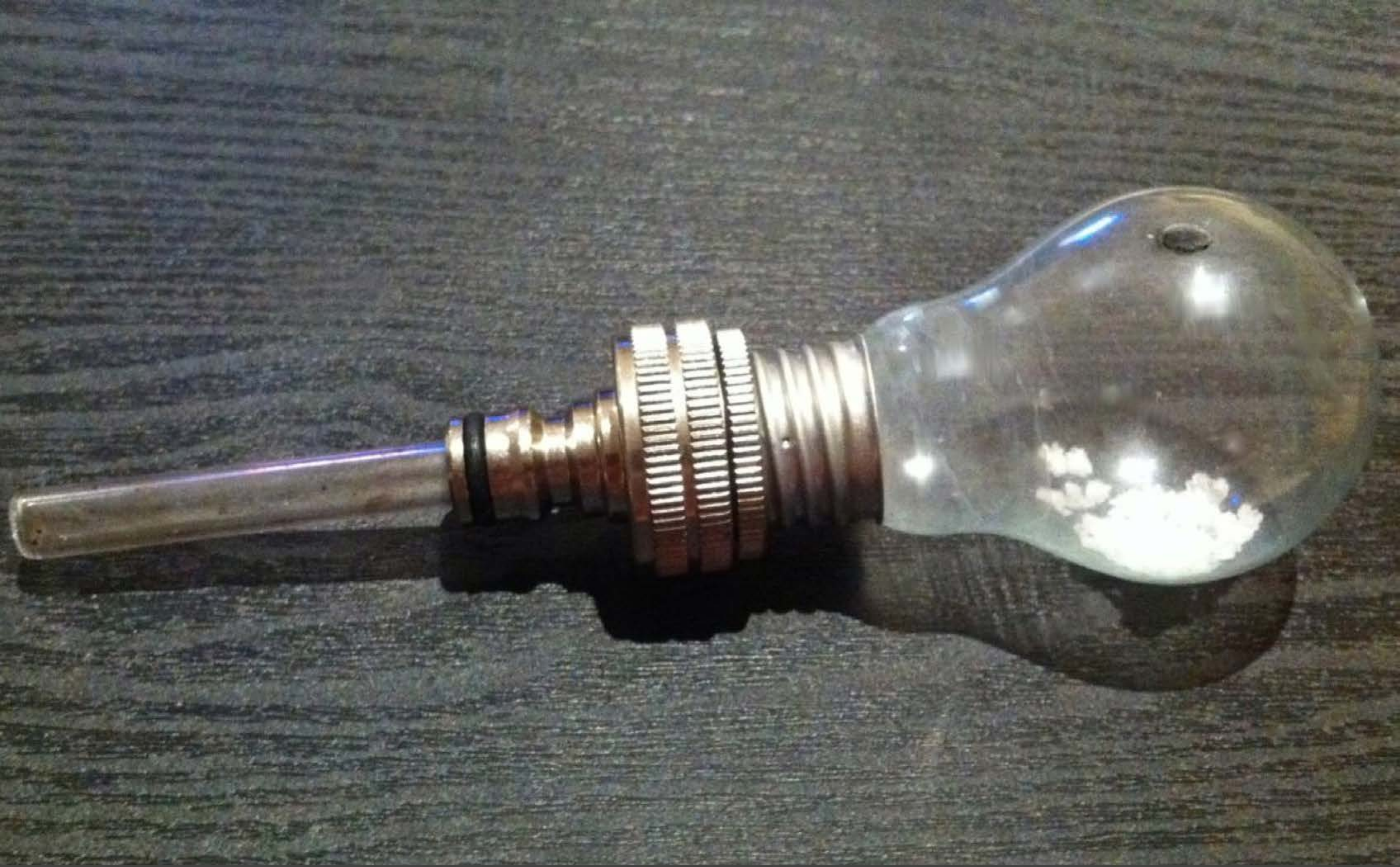

| Category | Craving (0-100) | Valence (1-9) | Arousal (1-9) | Typicality (0-100) | Relatedness | | | | | HSV | | |
|---|---|---|---|---|---|---|---|---|---|---|---|---|
| | | | | | Meth | Opioid | Both | Neither | MethToOpioid | Hue | Saturation | Value |
| Meth Instrument | 87.54 (17.66) | 4.71 (2.016) | 4.54 (2.83) | 89.21 (15.54) | 0.929 | 0 | 0.036 | 0.036 | -0.929 | 0.256 (0.23) | 0.091 (0.092) | 0.365 (0.152) |

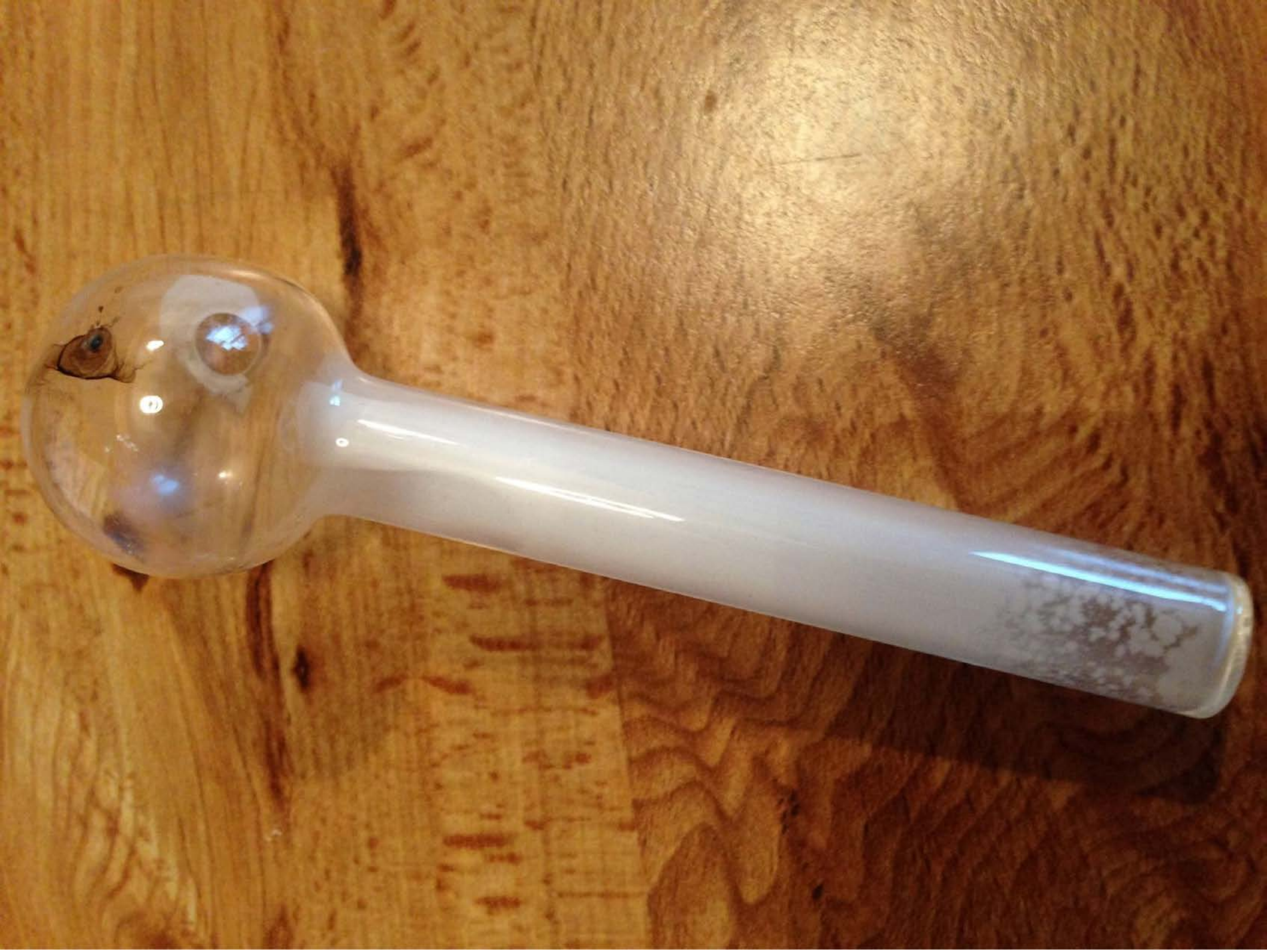

| Category | Craving (0-100) | Valence (1-9) | Arousal (1-9) | Typicality (0-100) | Relatedness | | | | | HSV | | |
|---|---|---|---|---|---|---|---|---|---|---|---|---|
| | | | | | Meth | Opioid | Both | Neither | MethToOpioid | Hue | Saturation | Value |
| Meth Instrument | 89.57 (20.86) | 4.43 (1.814) | 4.71 (2.59) | 91.46 (20.52) | 0.893 | 0 | 0.107 | 0 | -0.893 | 0.093 (0.097) | 0.646 (0.237) | 0.611 (0.169) |

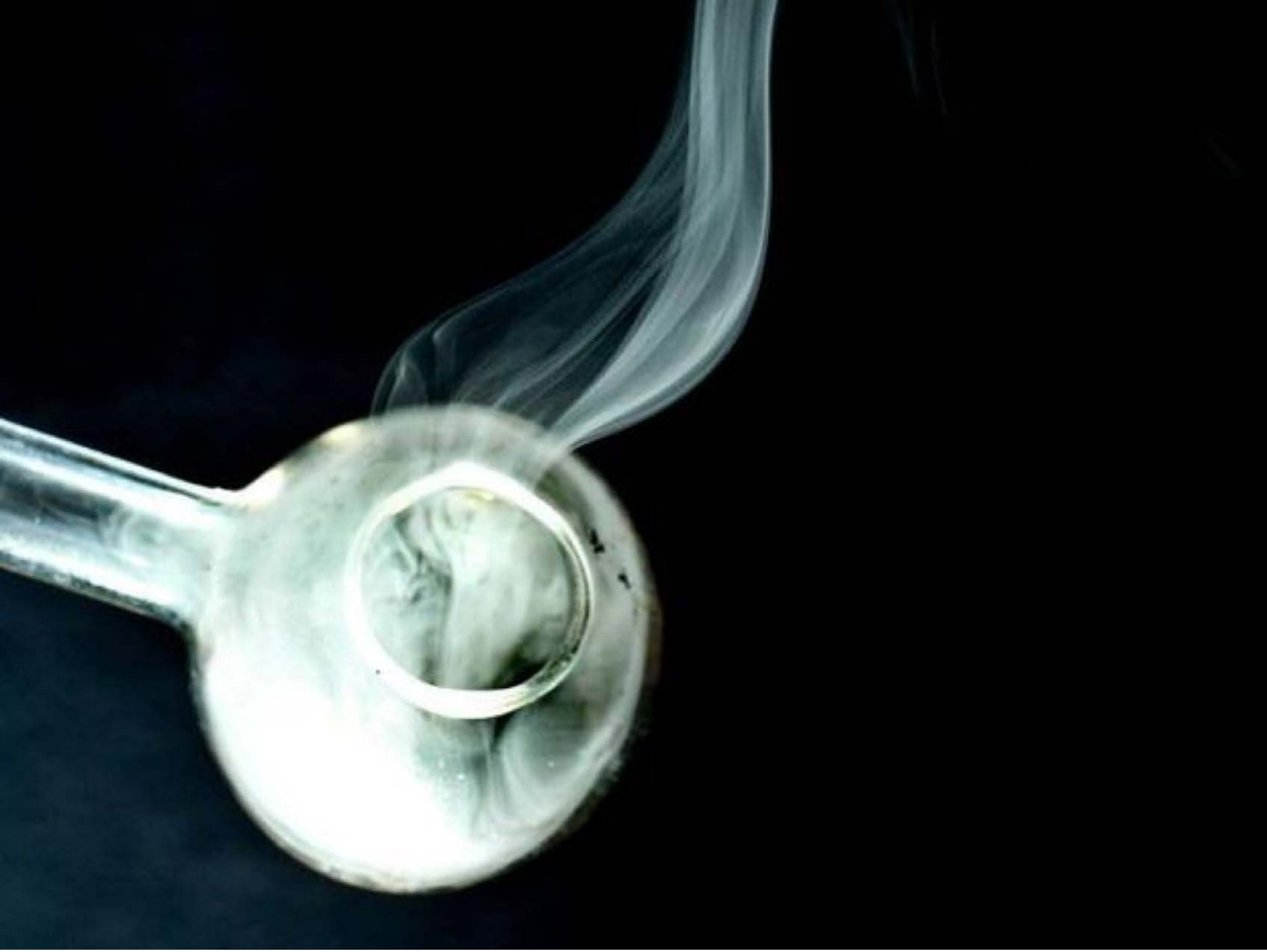

| Category | Craving (0-100) | Valence (1-9) | Arousal (1-9) | Typicality (0-100) | Relatedness | | | | | HSV | | |
|---|---|---|---|---|---|---|---|---|---|---|---|---|
| | | | | | Meth | Opioid | Both | Neither | MethToOpioid | Hue | Saturation | Value |
| Meth Instrument | 88 (17.47) | 4.48 (1.988) | 4.59 (2.83) | 89.41 (18.05) | 0.963 | 0 | 0.037 | 0 | -0.963 | 0.219 (0.27) | 0.21 (0.342) | 0.177 (0.315) |

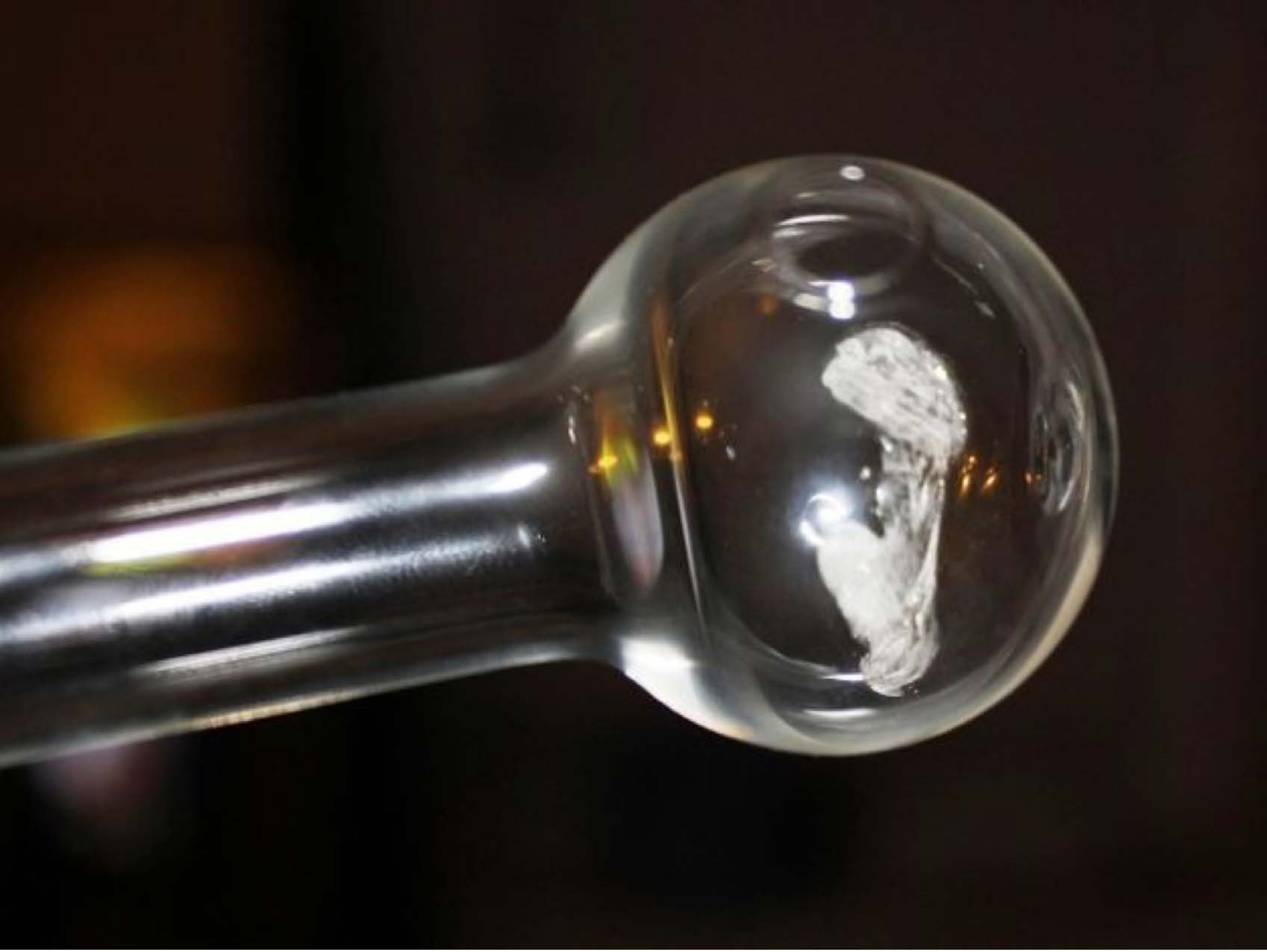

| Category | Craving (0-100) | Valence (1-9) | Arousal (1-9) | Typicality (0-100) | Relatedness | | | | | HSV | | |
|---|---|---|---|---|---|---|---|---|---|---|---|---|
| | | | | | Meth | Opioid | Both | Neither | MethToOpioid | Hue | Saturation | Value |
| Meth Instrument | 87.26 (18.64) | 4.59 (1.986) | 4.67 (2.7) | 88.04 (18.74) | 1 | 0 | 0 | 0 | -1 | 0.127 (0.18) | 0.397 (0.263) | 0.203 (0.202) |

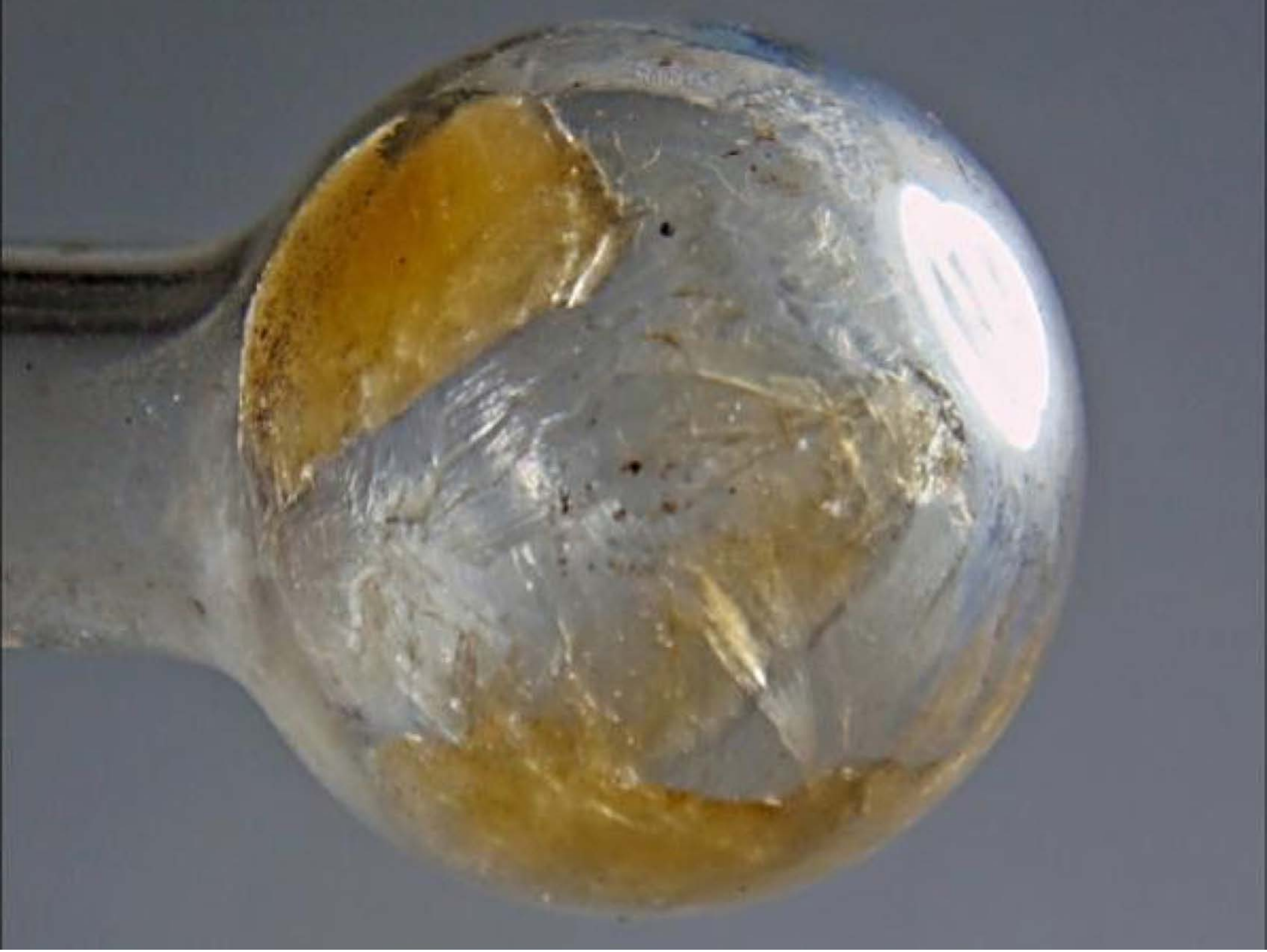

| Category | Craving (0-100) | Valence (1-9) | Arousal (1-9) | Typicality (0-100) | Relatedness | | | | | HSV | | |
|---|---|---|---|---|---|---|---|---|---|---|---|---|
| | | | | | Meth | Opioid | Both | Neither | MethToOpioid | Hue | Saturation | Value |
| Meth Instrument | 87.3 (18.96) | 4.44 (1.695) | 4.93 (2.6) | 87.67 (17.95) | 1 | 0 | 0 | 0 | -1 | 0.376 (0.264) | 0.189 (0.207) | 0.49 (0.118) |

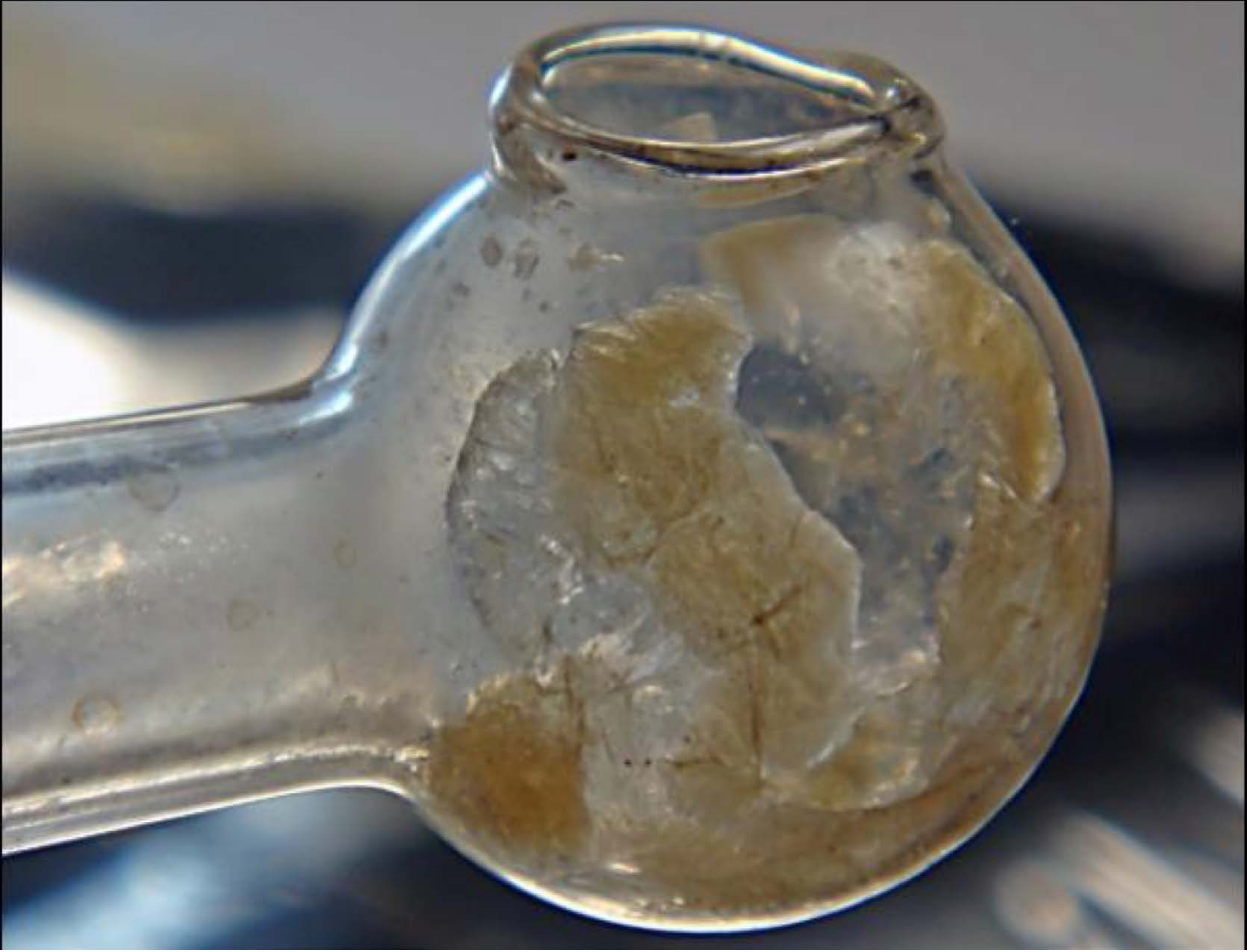

| Category | Craving (0-100) | Valence (1-9) | Arousal (1-9) | Typicality (0-100) | Relatedness | | | | | HSV | | |
|---|---|---|---|---|---|---|---|---|---|---|---|---|
| | | | | | Meth | Opioid | Both | Neither | MethToOpioid | Hue | Saturation | Value |
| Meth Instrument | 87.85 (16.91) | 4.41 (2.024) | 4.81 (2.65) | 88.78 (17.03) | 1 | 0 | 0 | 0 | -1 | 0.31 (0.268) | 0.187 (0.165) | 0.461 (0.17) |

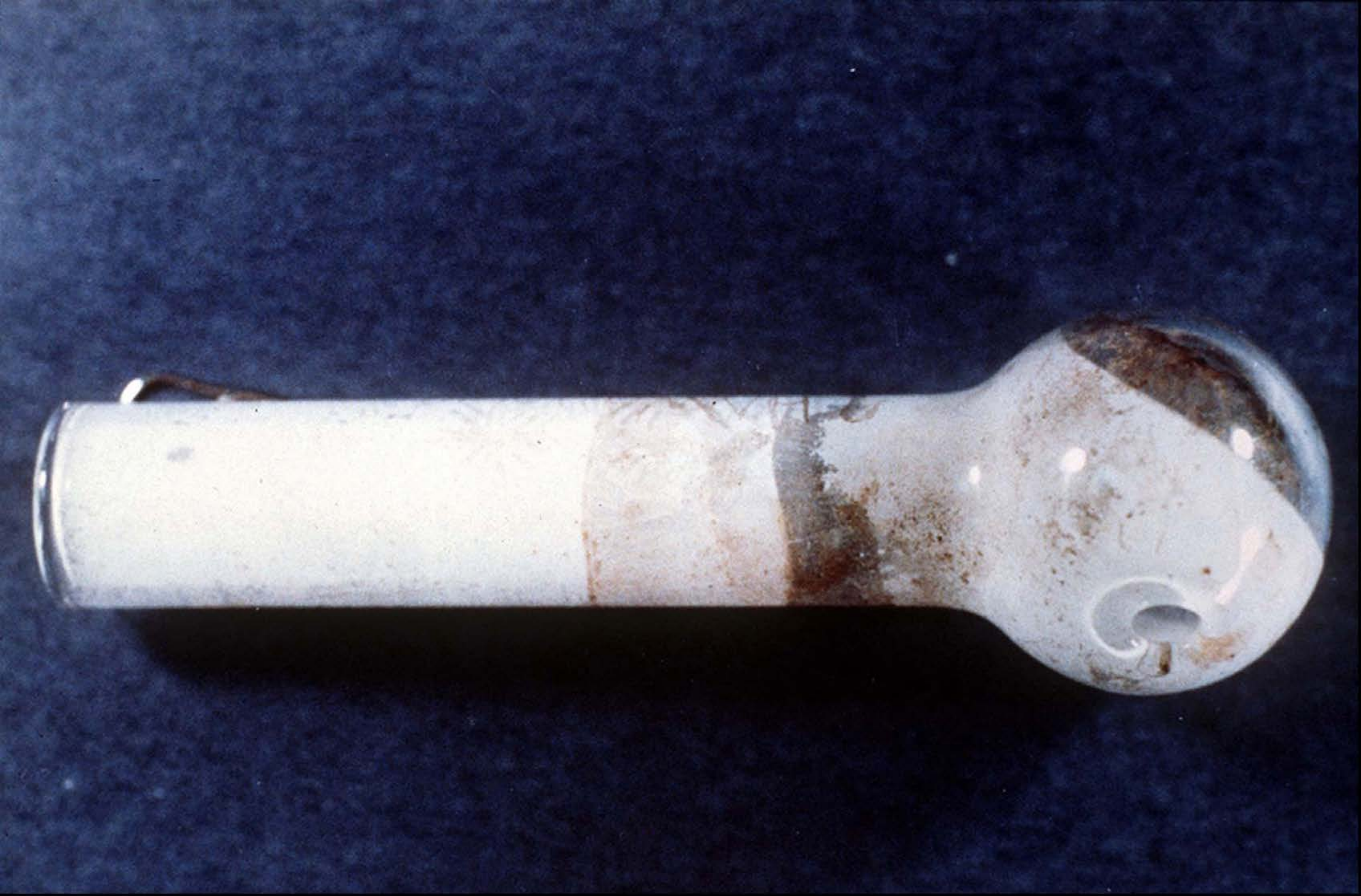

| Category | Craving (0-100) | Valence (1-9) | Arousal (1-9) | Typicality (0-100) | Relatedness | | | | | HSV | | |
|---|---|---|---|---|---|---|---|---|---|---|---|---|
| | | | | | Meth | Opioid | Both | Neither | MethToOpioid | Hue | Saturation | Value |
| Meth Instrument | 93.79 (10.8) | 4.59 (2.024) | 4.43 (2.81) | 94.54 (10.15) | 0.926 | 0 | 0.074 | 0 | -0.926 | 0.537 (0.276) | 0.407 (0.285) | 0.323 (0.294) |

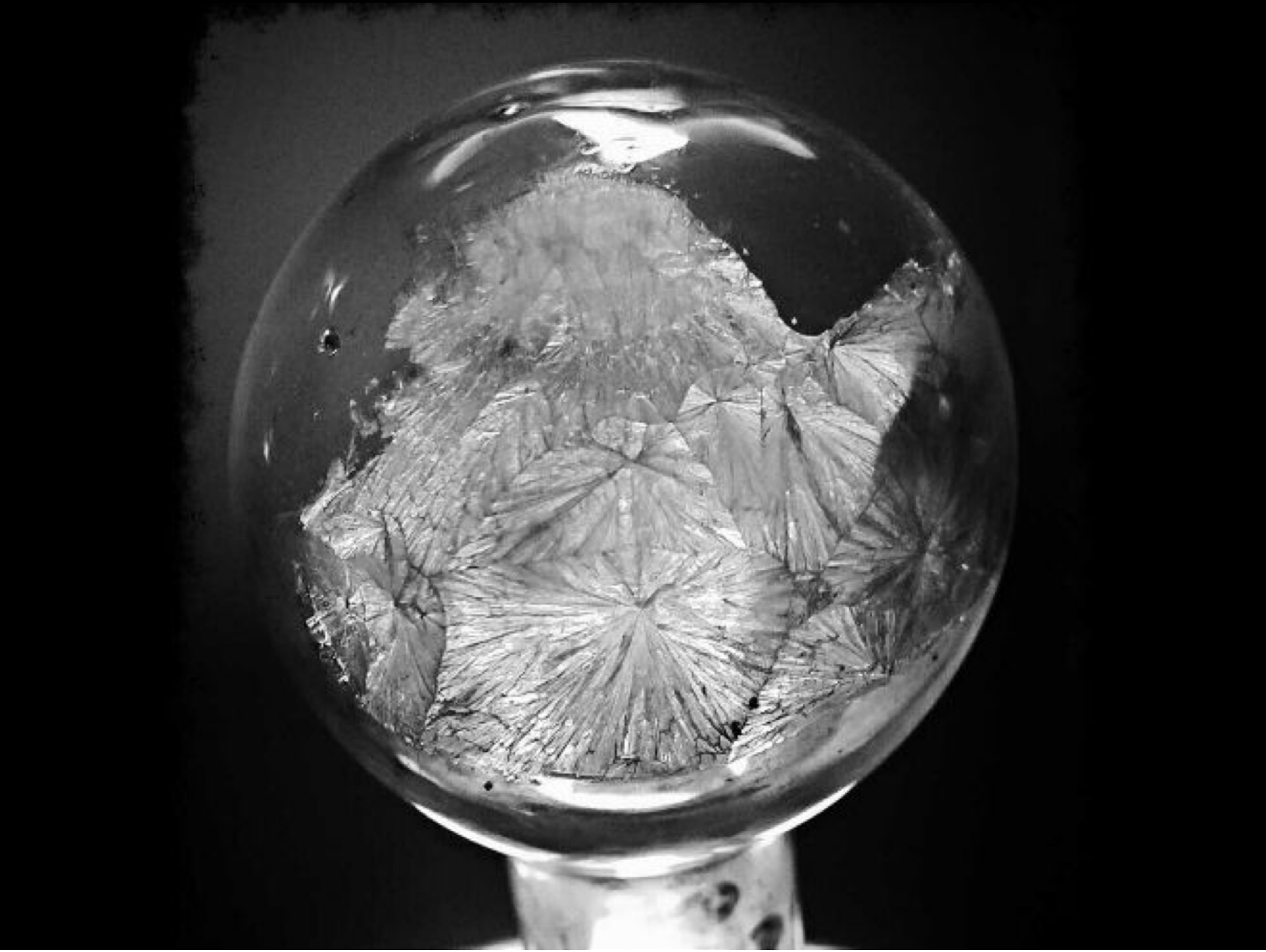

| Category | Craving (0-100) | Valence (1-9) | Arousal (1-9) | Typicality (0-100) | Relatedness | | | | | HSV | | |
|---|---|---|---|---|---|---|---|---|---|---|---|---|
| | | | | | Meth | Opioid | Both | Neither | MethToOpioid | Hue | Saturation | Value |
| Meth Instrument | 80.22 (25.73) | 4.11 (2.044) | 4.74 (2.73) | 79.81 (26.79) | 0.963 | 0 | 0 | 0.037 | -0.963 | 0 (0) | 0 (0) | 0.235 (0.281) |

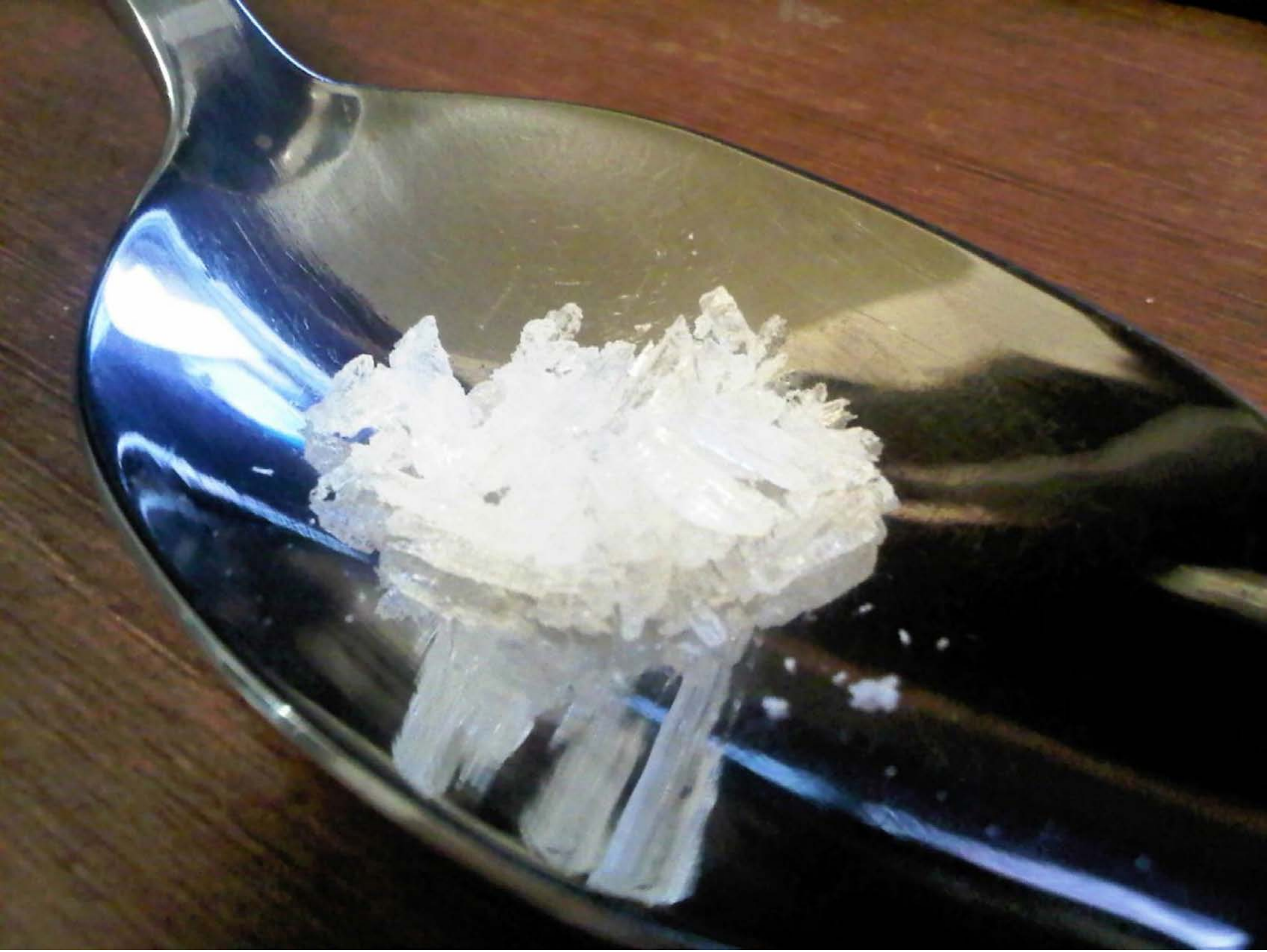

| Category | Craving (0-100) | Valence (1-9) | Arousal (1-9) | Typicality (0-100) | Relatedness | | | | | HSV | | |
|---|---|---|---|---|---|---|---|---|---|---|---|---|
| | | | | | Meth | Opioid | Both | Neither | MethToOpioid | Hue | Saturation | Value |
| Meth Instrument | 90.63 (14.98) | 4.33 (2.184) | 4.89 (2.87) | 90.78 (14.45) | 1 | 0 | 0 | 0 | -1 | 0.221 (0.237) | 0.292 (0.23) | 0.421 (0.283) |

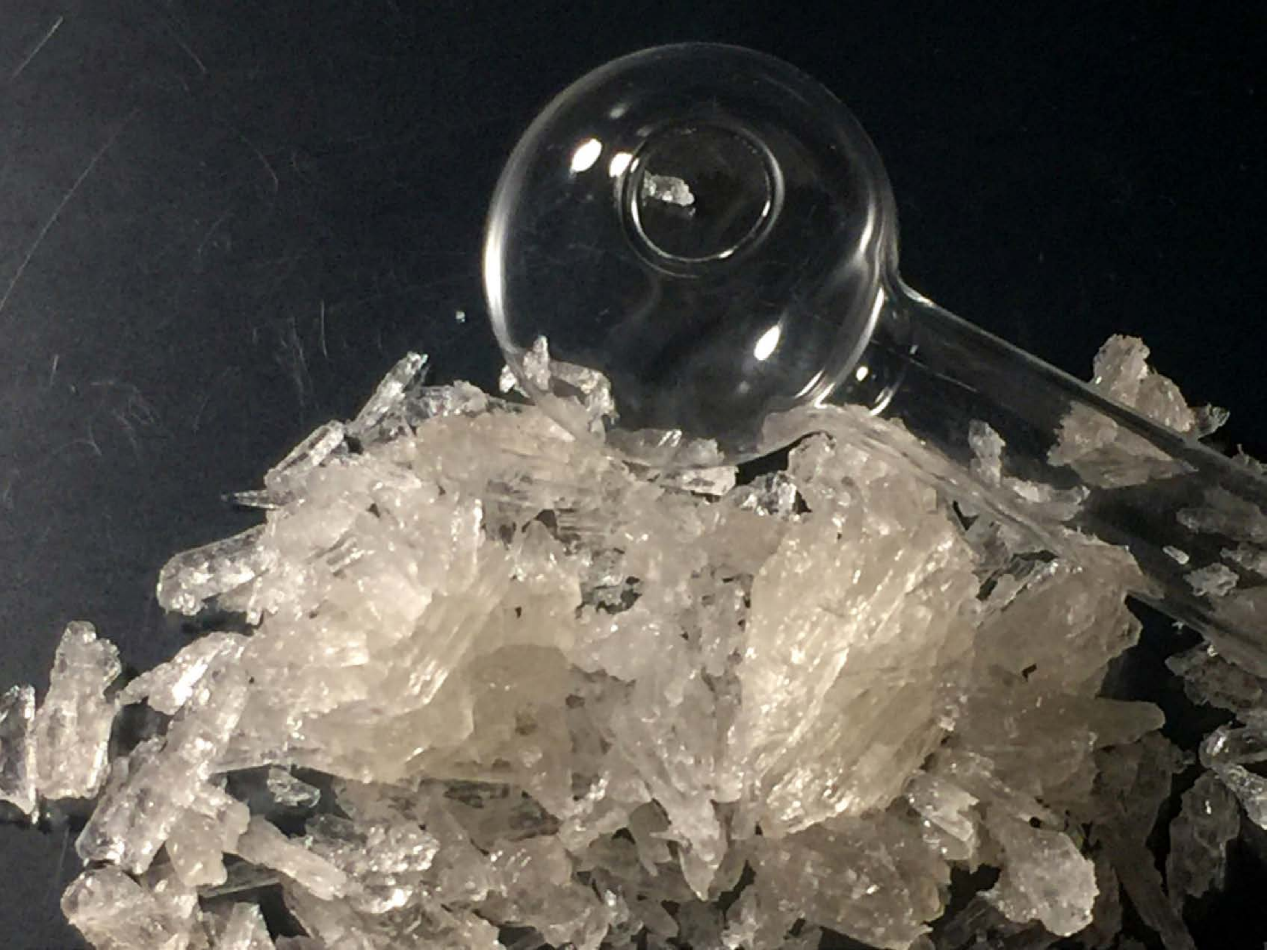

| Category | Craving (0-100) | Valence (1-9) | Arousal (1-9) | Typicality (0-100) | Relatedness | | | | | HSV | | |
|---|---|---|---|---|---|---|---|---|---|---|---|---|
| | | | | | Meth | Opioid | Both | Neither | MethToOpioid | Hue | Saturation | Value |
| Meth Instrument | 92.43 (12.51) | 4.54 (2.186) | 5 (2.82) | 93.21 (11.99) | 1 | 0 | 0 | 0 | -1 | 0.309 (0.24) | 0.231 (0.179) | 0.36 (0.263) |

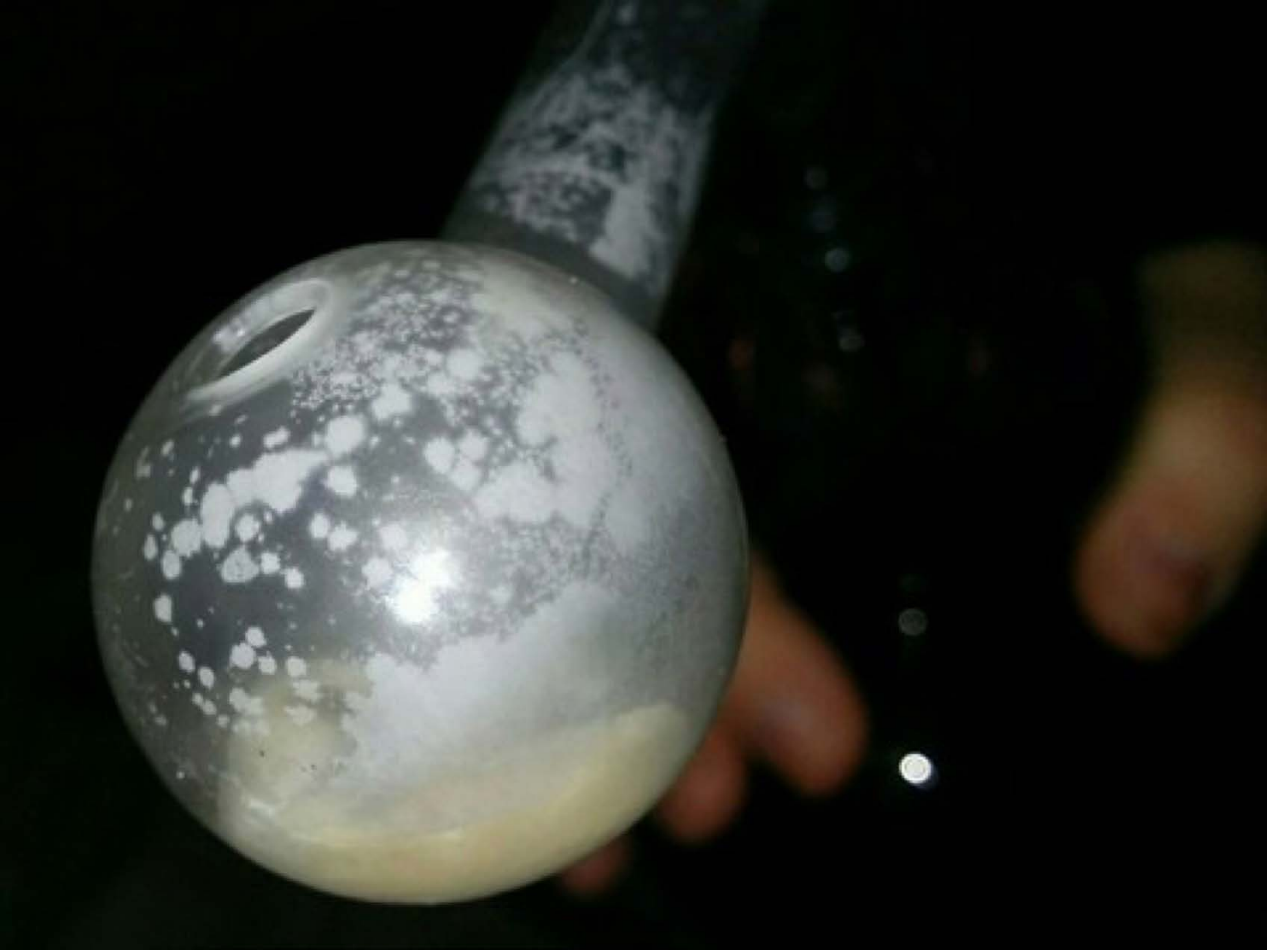

| Category | Craving (0-100) | Valence (1-9) | Arousal (1-9) | Typicality (0-100) | Relatedness | | | | | HSV | | |
|---|---|---|---|---|---|---|---|---|---|---|---|---|
| | | | | | Meth | Opioid | Both | Neither | MethToOpioid | Hue | Saturation | Value |
| Meth Instrument | 93 (12.46) | 4.57 (1.687) | 4.5 (2.73) | 93.04 (11.81) | 1 | 0 | 0 | 0 | -1 | 0.231 (0.205) | 0.324 (0.395) | 0.194 (0.26) |

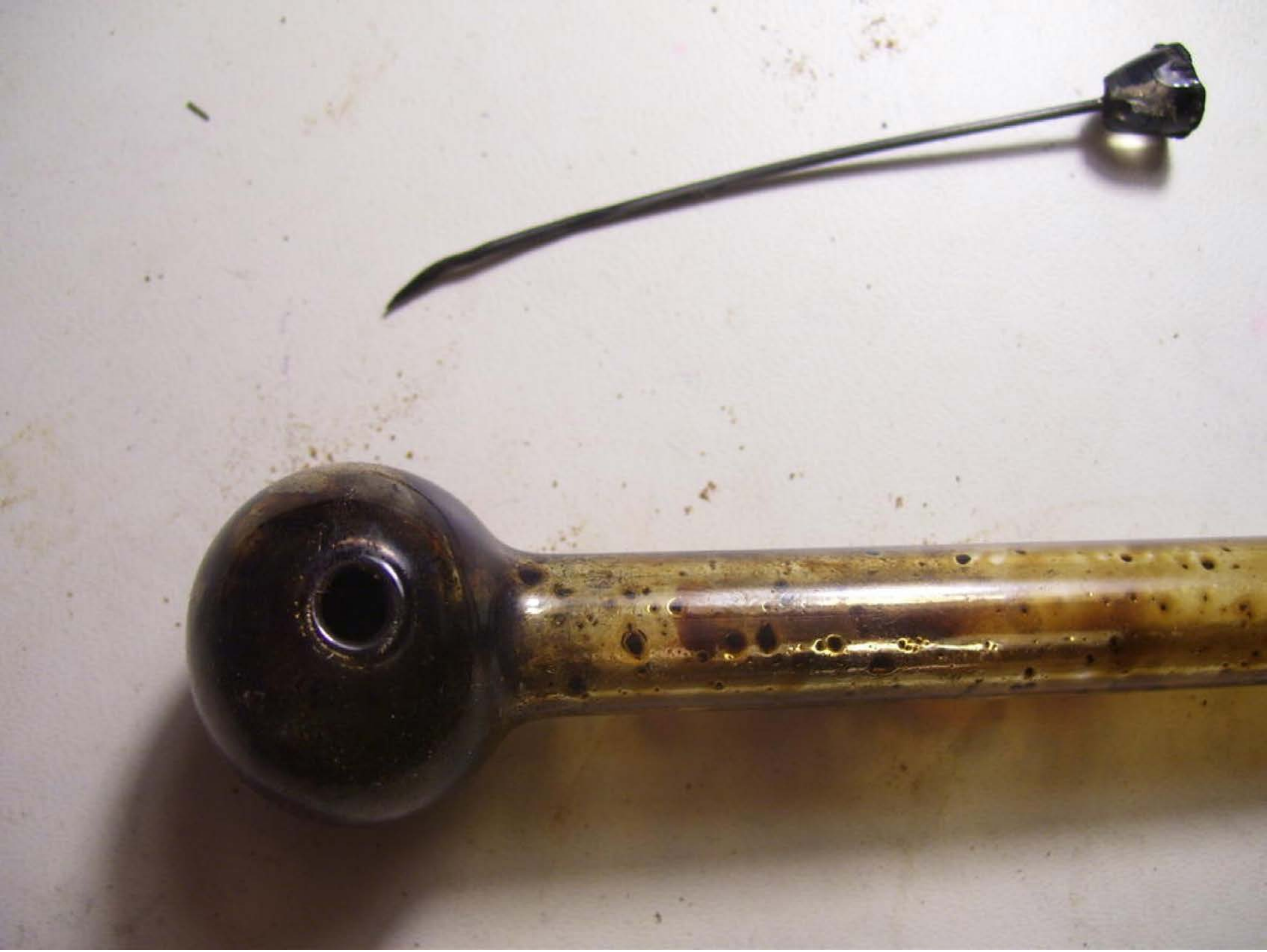

| Category | Craving (0-100) | Valence (1-9) | Arousal (1-9) | Typicality (0-100) | Relatedness | | | | | HSV | | |
|---|---|---|---|---|---|---|---|---|---|---|---|---|
| | | | | | Meth | Opioid | Both | Neither | MethToOpioid | Hue | Saturation | Value |
| Meth Instrument | 92.54 (13.65) | 4.43 (1.933) | 4.61 (2.66) | 91.04 (14.48) | 0.857 | 0 | 0.143 | 0 | -0.857 | 0.323 (0.404) | 0.18 (0.217) | 0.66 (0.263) |

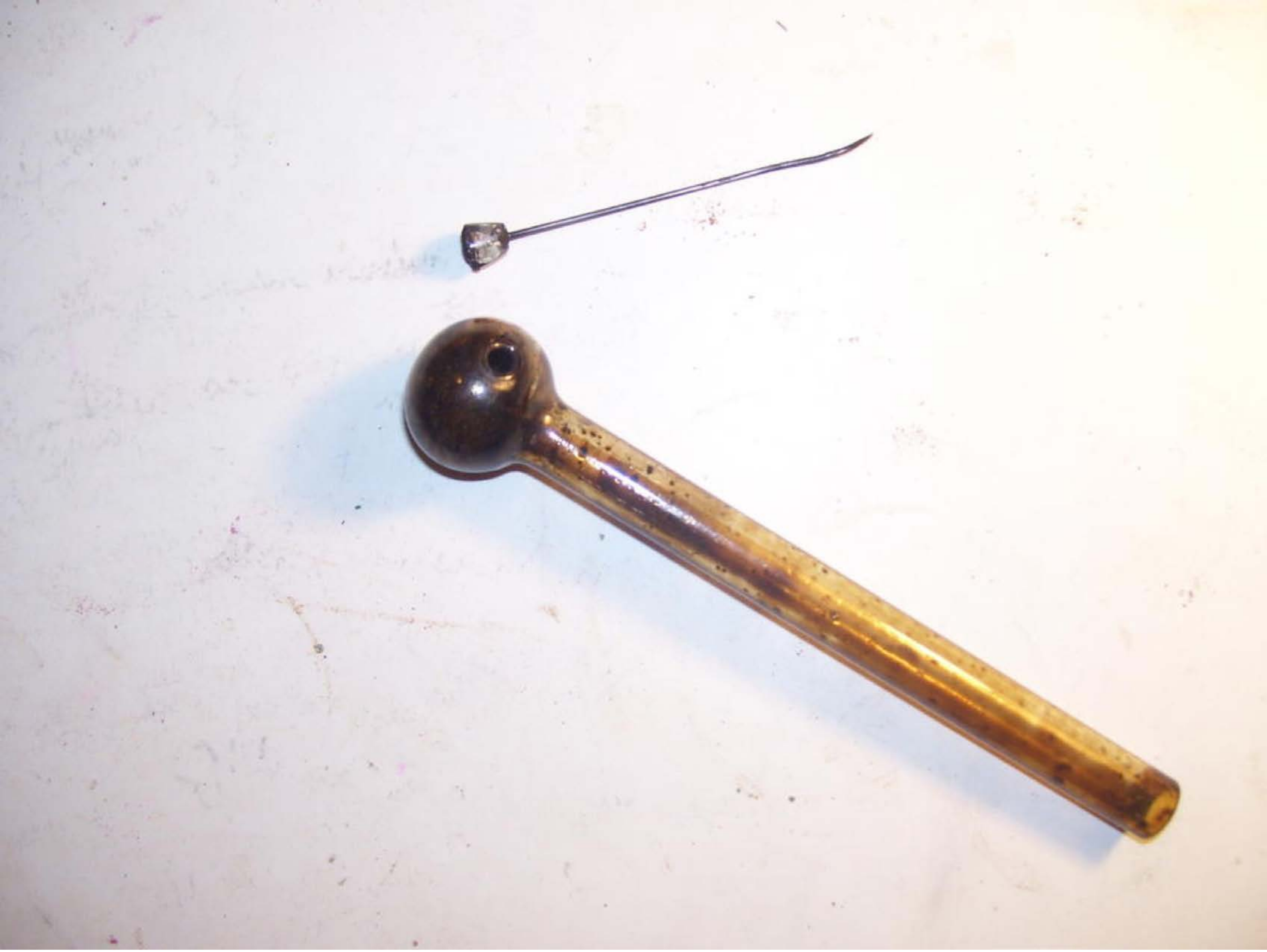

| Category | Craving (0-100) | Valence (1-9) | Arousal (1-9) | Typicality (0-100) | Relatedness | | | | | HSV | | |
|---|---|---|---|---|---|---|---|---|---|---|---|---|
| | | | | | Meth | Opioid | Both | Neither | MethToOpioid | Hue | Saturation | Value |
| Meth Instrument | 91.39 (19.59) | 3.93 (2.017) | 4.71 (2.84) | 94.79 (10.02) | 0.893 | 0 | 0.071 | 0.036 | -0.893 | 0.501 (0.38) | 0.06 (0.104) | 0.912 (0.122) |

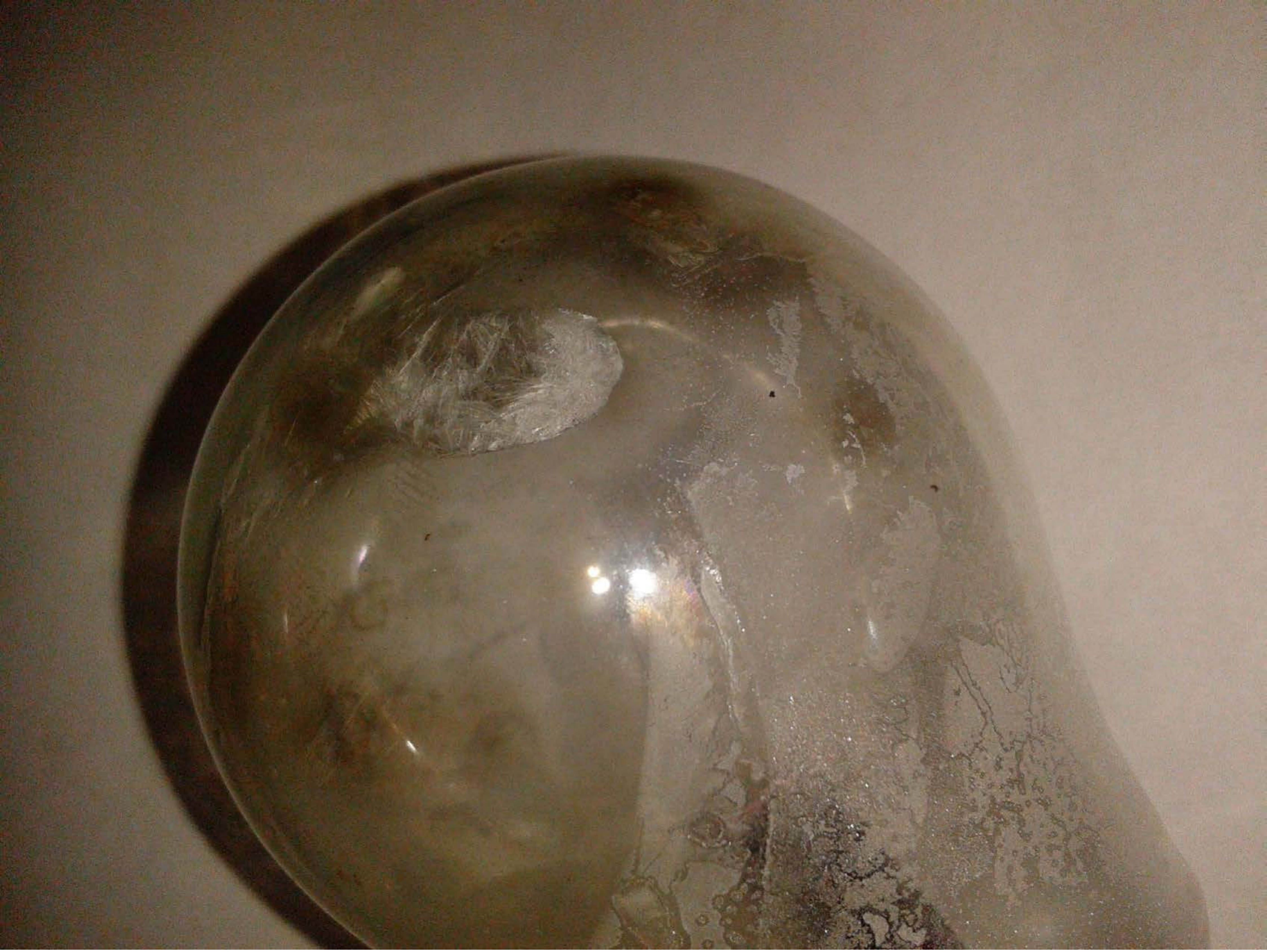

| Category | Craving (0-100) | Valence (1-9) | Arousal (1-9) | Typicality (0-100) | Relatedness | | | | | HSV | | |
|---|---|---|---|---|---|---|---|---|---|---|---|---|
| | | | | | Meth | Opioid | Both | Neither | MethToOpioid | Hue | Saturation | Value |
| Meth Instrument | 86.44 (17.41) | 4.59 (2.08) | 5 (2.76) | 87.44 (17.57) | 1 | 0 | 0 | 0 | -1 | 0.09 (0.039) | 0.344 (0.162) | 0.457 (0.156) |

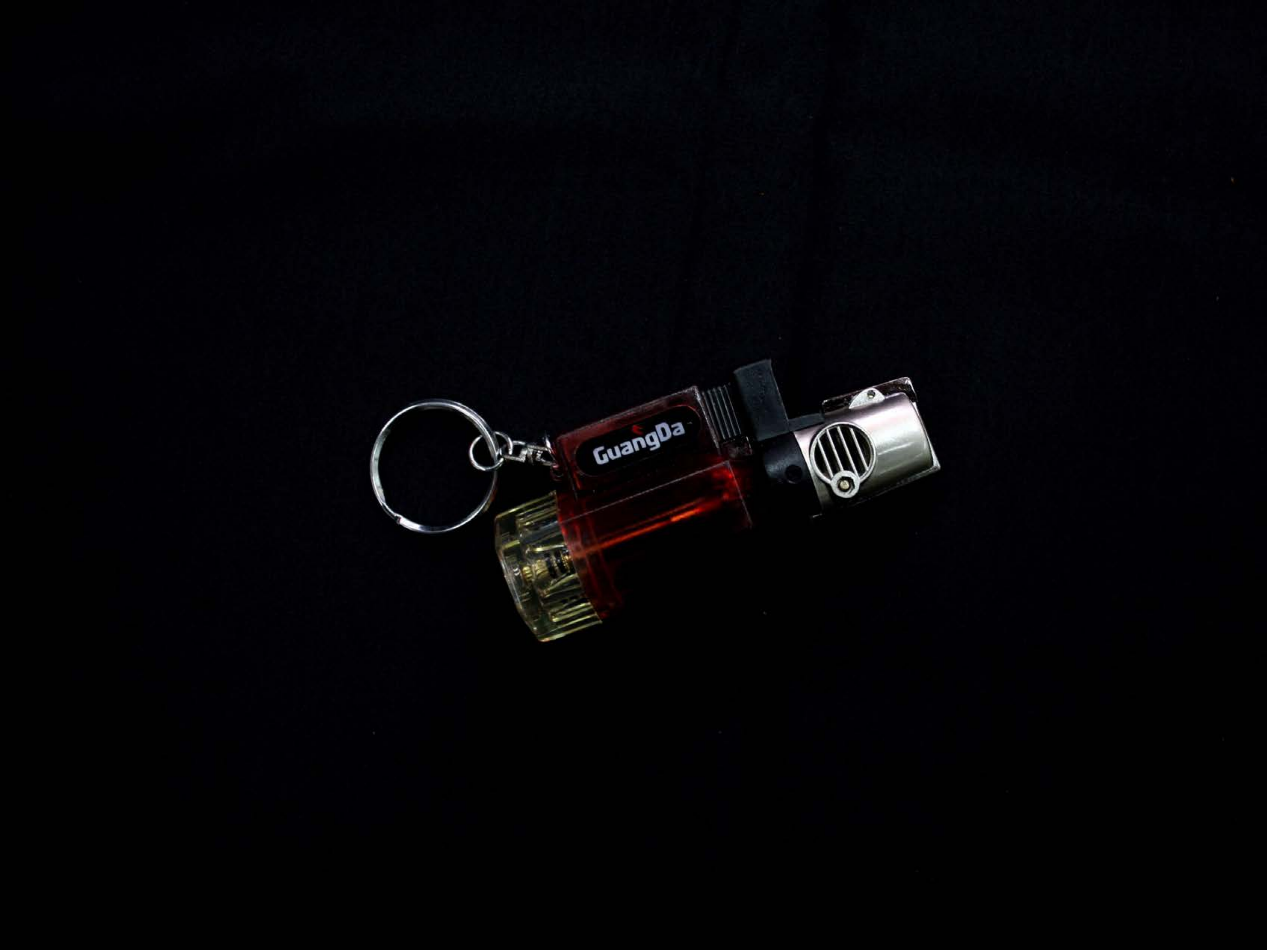


| Category | Craving (0-100) | Valence (1-9) | Arousal (1-9) | Typicality (0-100) | Relatedness | | | | | HSV | | |
|---|---|---|---|---|---|---|---|---|---|---|---|---|
| | | | | | Meth | Opioid | Both | Neither | MethToOpioid | Hue | Saturation | Value |
| Meth Instrument | 62.39 (30.91) | 4.43 (1.814) | 3.5 (2.38) | 73.39 (31.57) | 0.25 | 0 | 0.464 | 0.286 | -0.25 | 0.458 (0.31) | 0.559 (0.409) | 0.027 (0.076) |

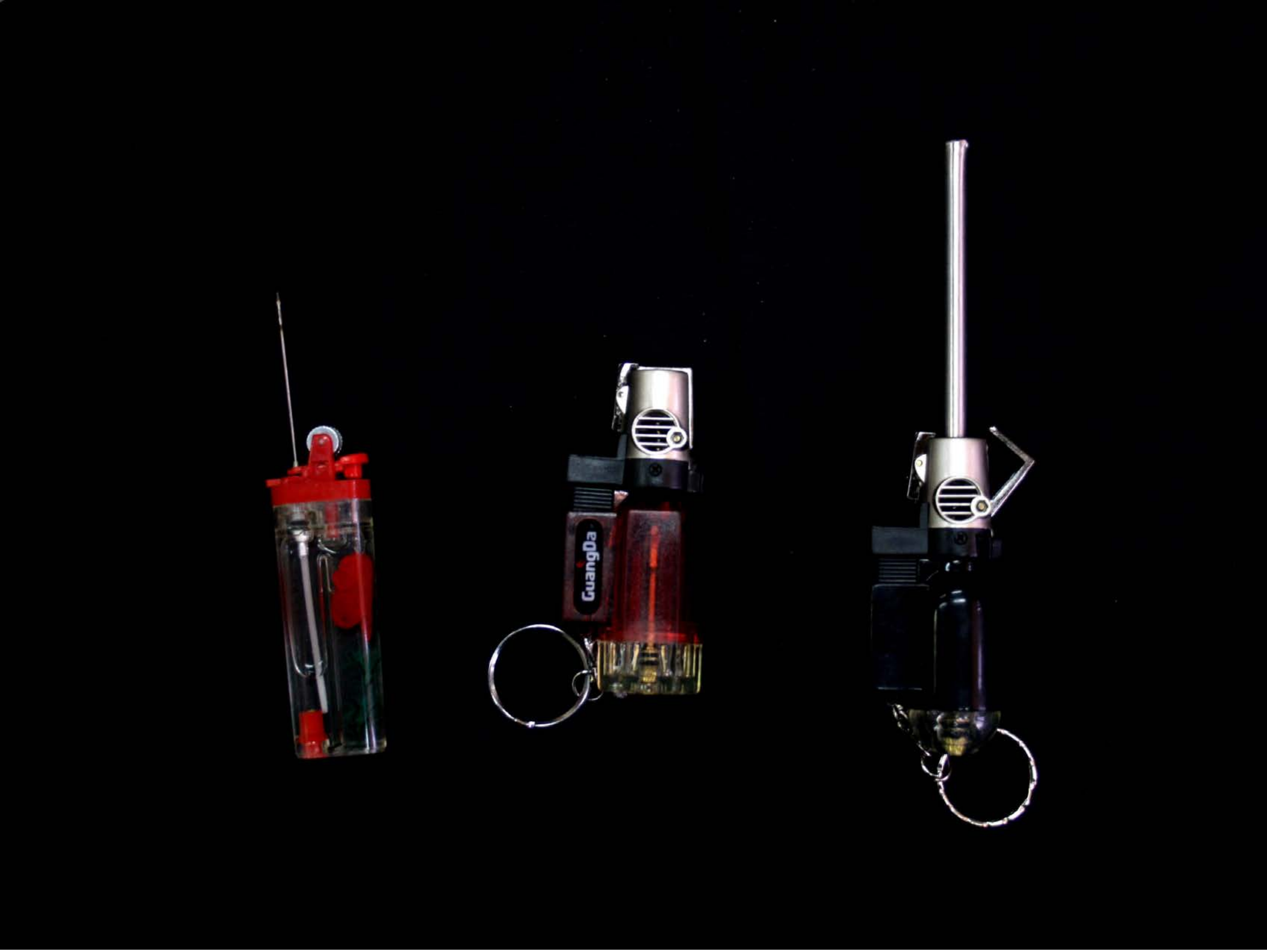


| Category | Craving (0-100) | Valence (1-9) | Arousal (1-9) | Typicality (0-100) | Relatedness | | | | | HSV | | |
|---|---|---|---|---|---|---|---|---|---|---|---|---|
| | | | | | Meth | Opioid | Both | Neither | MethToOpioid | Hue | Saturation | Value |
| Meth Instrument | 54.7 (31.81) | 4.56 (1.05) | 3.41 (2.12) | 64.48 (33.79) | 0.296 | 0 | 0.444 | 0.259 | -0.296 | 0.173 (0.3) | 0.226 (0.399) | 0.028 (0.118) |

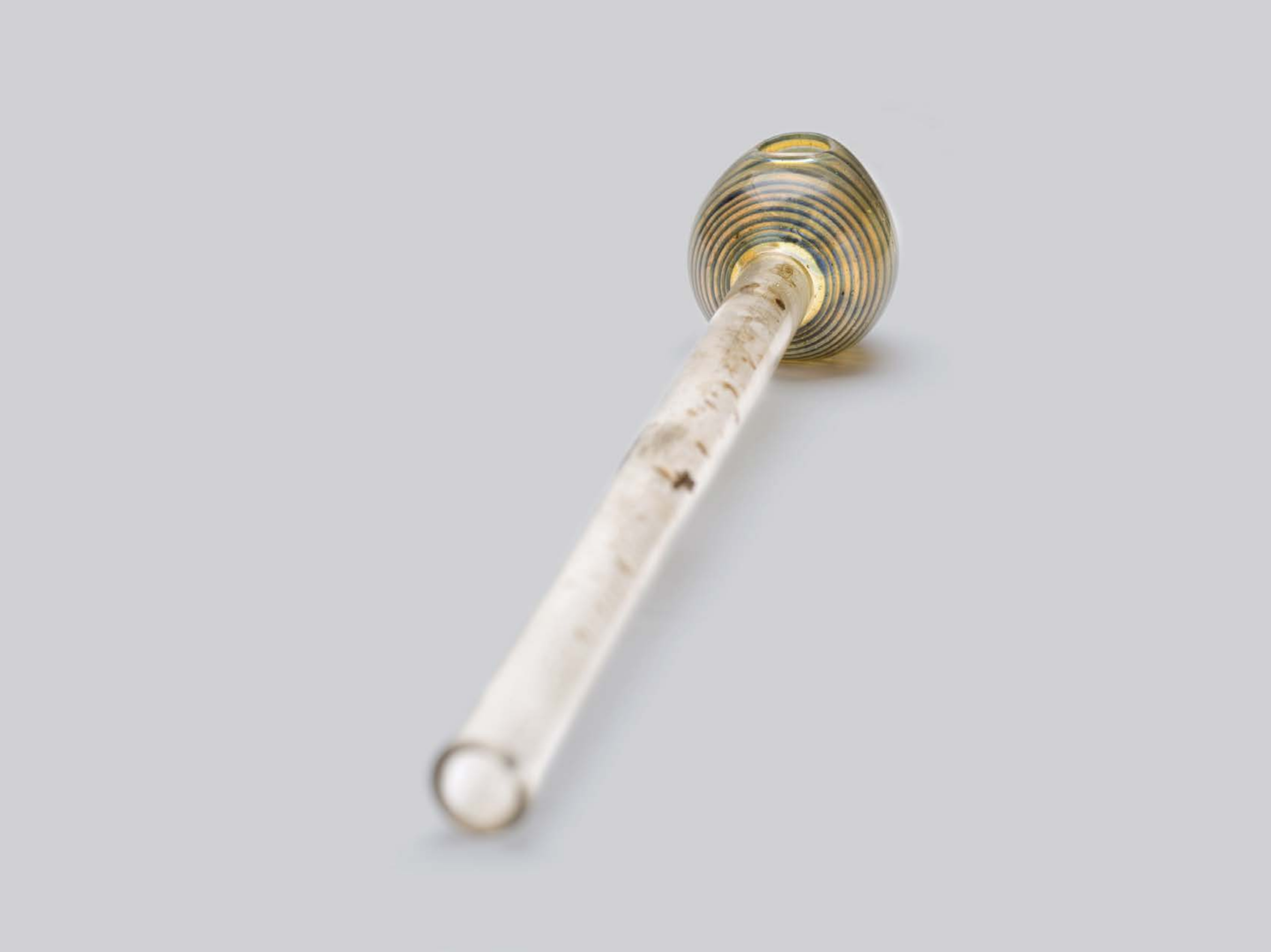

| Category | Craving (0-100) | Valence (1-9) | Arousal (1-9) | Typicality (0-100) | Relatedness | | | | | HSV | | |
|---|---|---|---|---|---|---|---|---|---|---|---|---|
| | | | | | Meth | Opioid | Both | Neither | MethToOpioid | Hue | Saturation | Value |
| Meth Instrument | 83.71 (27.19) | 4.25 (1.974) | 4.21 (2.79) | 85.89 (26.21) | 0.857 | 0 | 0.107 | 0.036 | -0.857 | 0.032 (0.137) | 0.012 (0.056) | 0.982 (0.072) |

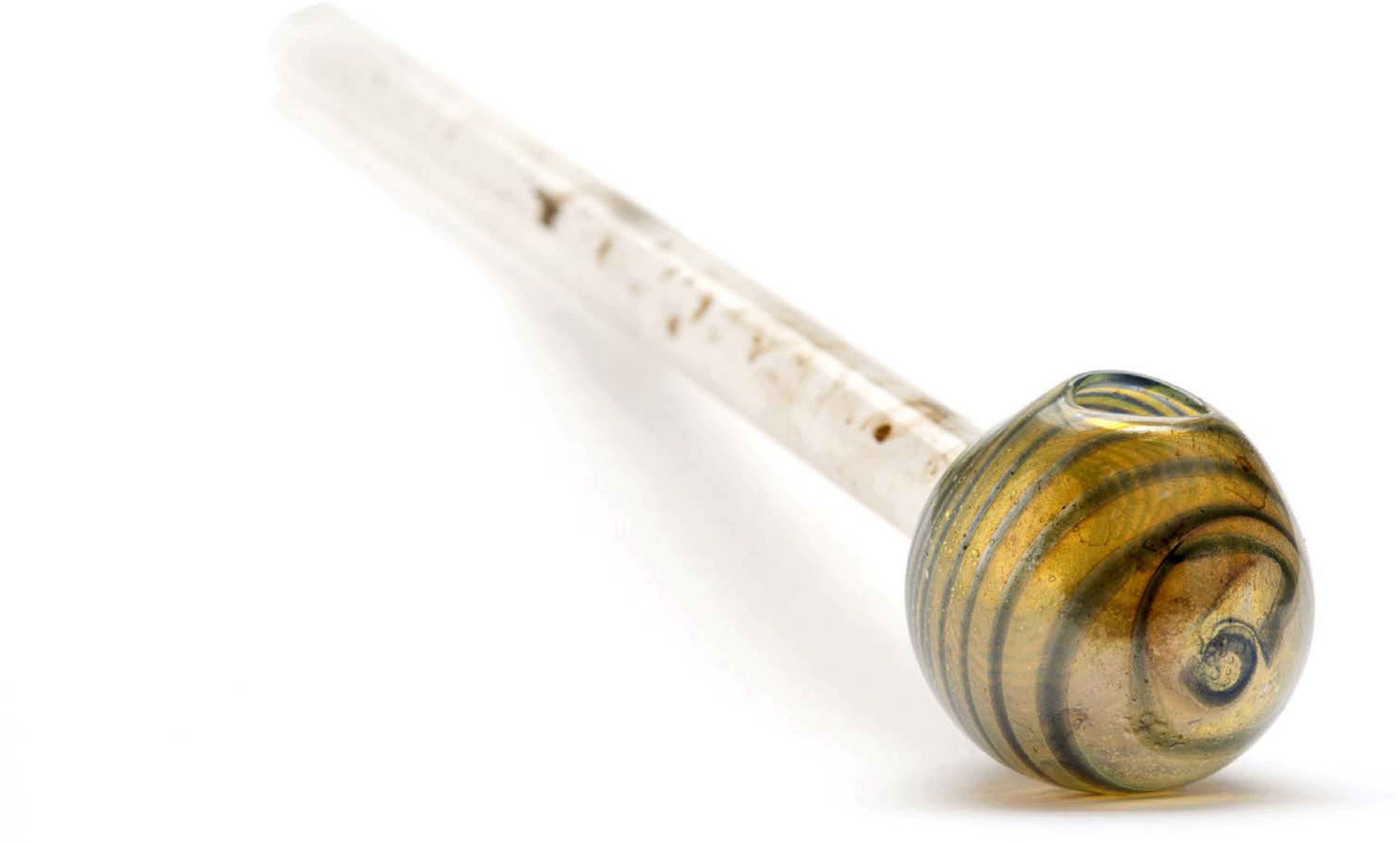

| Category | Craving (0-100) | Valence (1-9) | Arousal (1-9) | Typicality (0-100) | Relatedness | | | | | HSV | | |
|---|---|---|---|---|---|---|---|---|---|---|---|---|
| | | | | | Meth | Opioid | Both | Neither | MethToOpioid | Hue | Saturation | Value |
| Meth Instrument | 89.18 (16.73) | 4.82 (1.806) | 4.5 (2.8) | 92.75 (13.66) | 0.929 | 0 | 0.071 | 0 | -0.929 | 0.029 (0.095) | 0.04 (0.142) | 0.964 (0.115) |

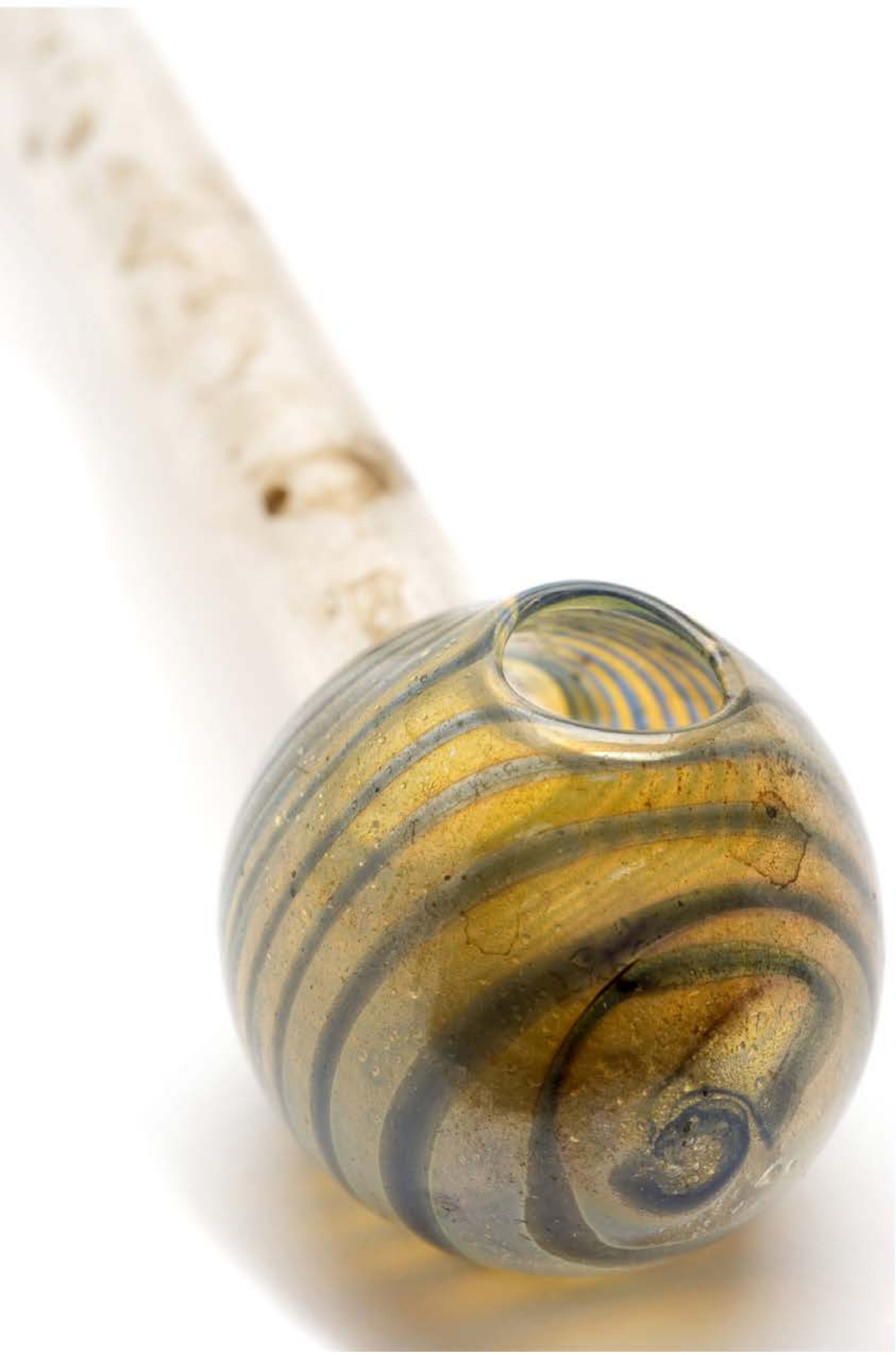

| Category | Craving (0-100) | Valence (1-9) | Arousal (1-9) | Typicality (0-100) | Relatedness | | | | | HSV | | |
|---|---|---|---|---|---|---|---|---|---|---|---|---|
| | | | | | Meth | Opioid | Both | Neither | MethToOpioid | Hue | Saturation | Value |
| Meth Instrument | 88.11 (16.38) | 4.48 (1.868) | 4.59 (2.8) | 86.3 (19.86) | 0.889 | 0.037 | 0.074 | 0 | -0.852 | 0.032 (0.067) | 0.073 (0.173) | 0.929 (0.157) |

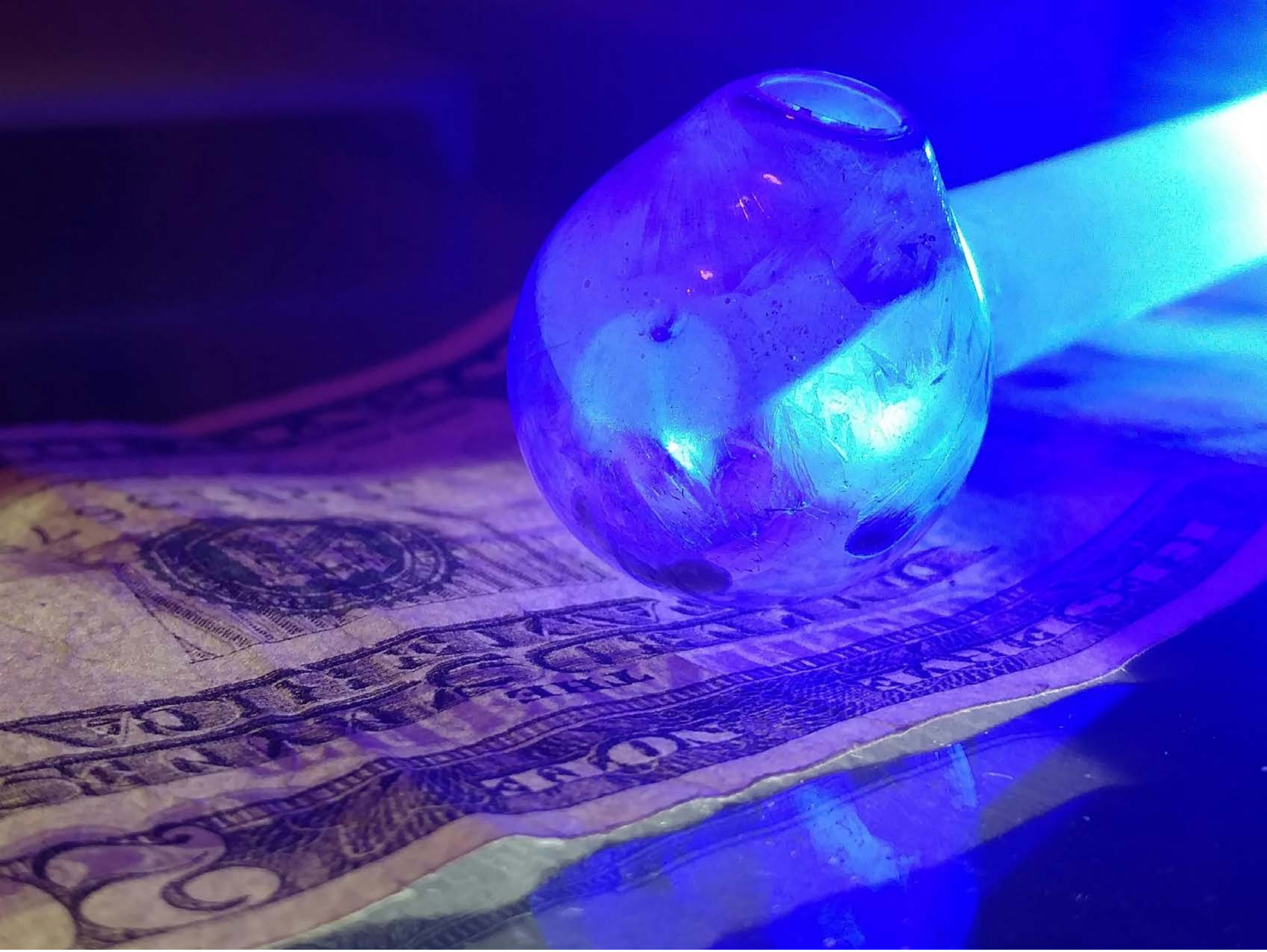

| Category | Craving (0-100) | Valence (1-9) | Arousal (1-9) | Typicality (0-100) | Relatedness | | | | | HSV | | |
|---|---|---|---|---|---|---|---|---|---|---|---|---|
| | | | | | Meth | Opioid | Both | Neither | MethToOpioid | Hue | Saturation | Value |
| Meth Instrument | 86.63 (18.62) | 4.41 (1.866) | 4.52 (2.61) | 84.78 (20.52) | 0.926 | 0 | 0.074 | 0 | -0.926 | 0.678 (0.125) | 0.751 (0.259) | 0.69 (0.267) |

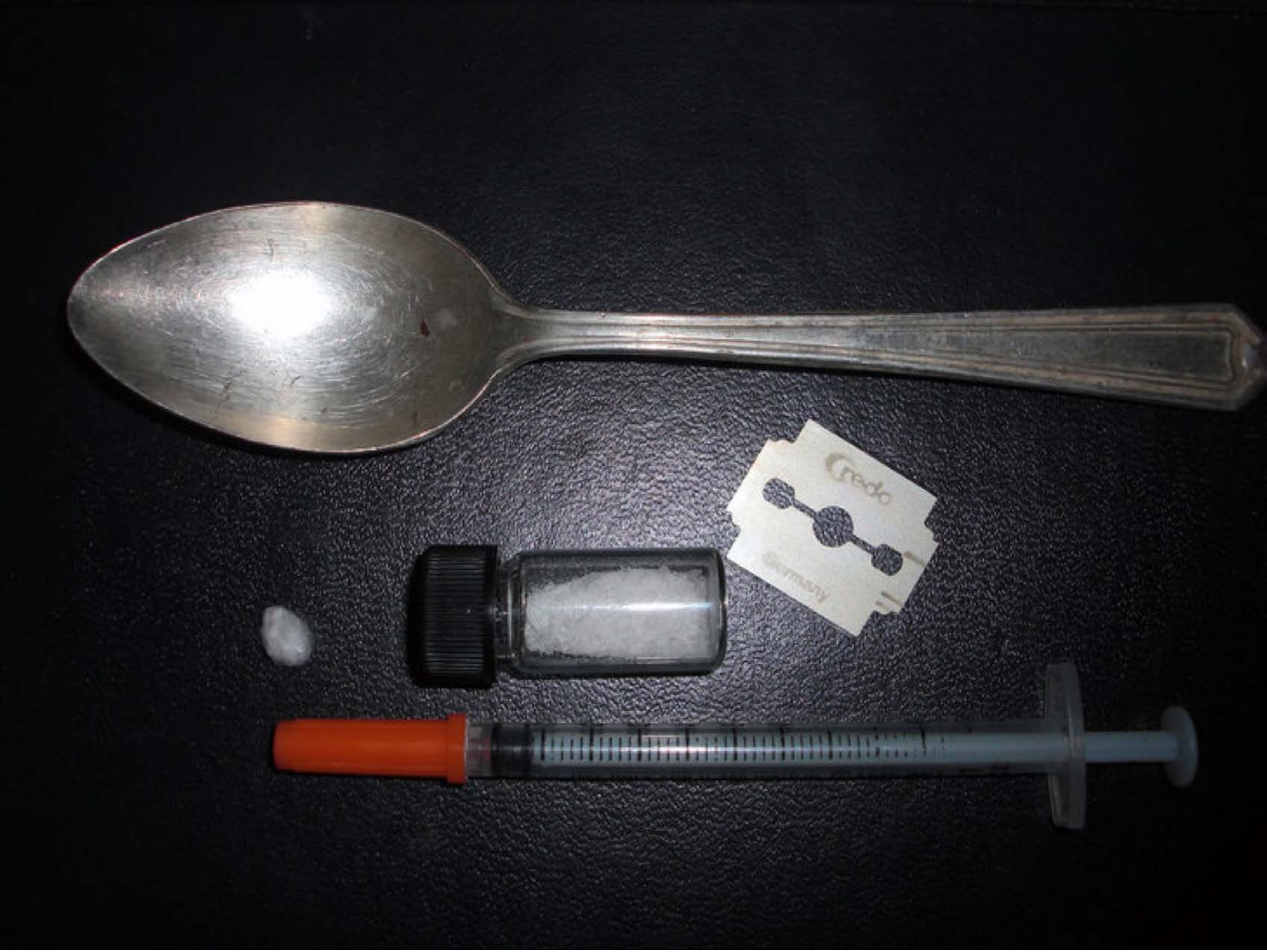


| Category | Craving (0-100) | Valence (1-9) | Arousal (1-9) | Typicality (0-100) | Relatedness | | | | | HSV | | |
|---|---|---|---|---|---|---|---|---|---|---|---|---|
| | | | | | Meth | Opioid | Both | Neither | MethToOpioid | Hue | Saturation | Value |
| Meth Instrument | 91.18 (15.8) | 4.32 (2.056) | 5.04 (2.9) | 90.57 (16.19) | 0.607 | 0.036 | 0.357 | 0 | -0.571 | 0.455 (0.299) | 0.123 (0.134) | 0.179 (0.212) |

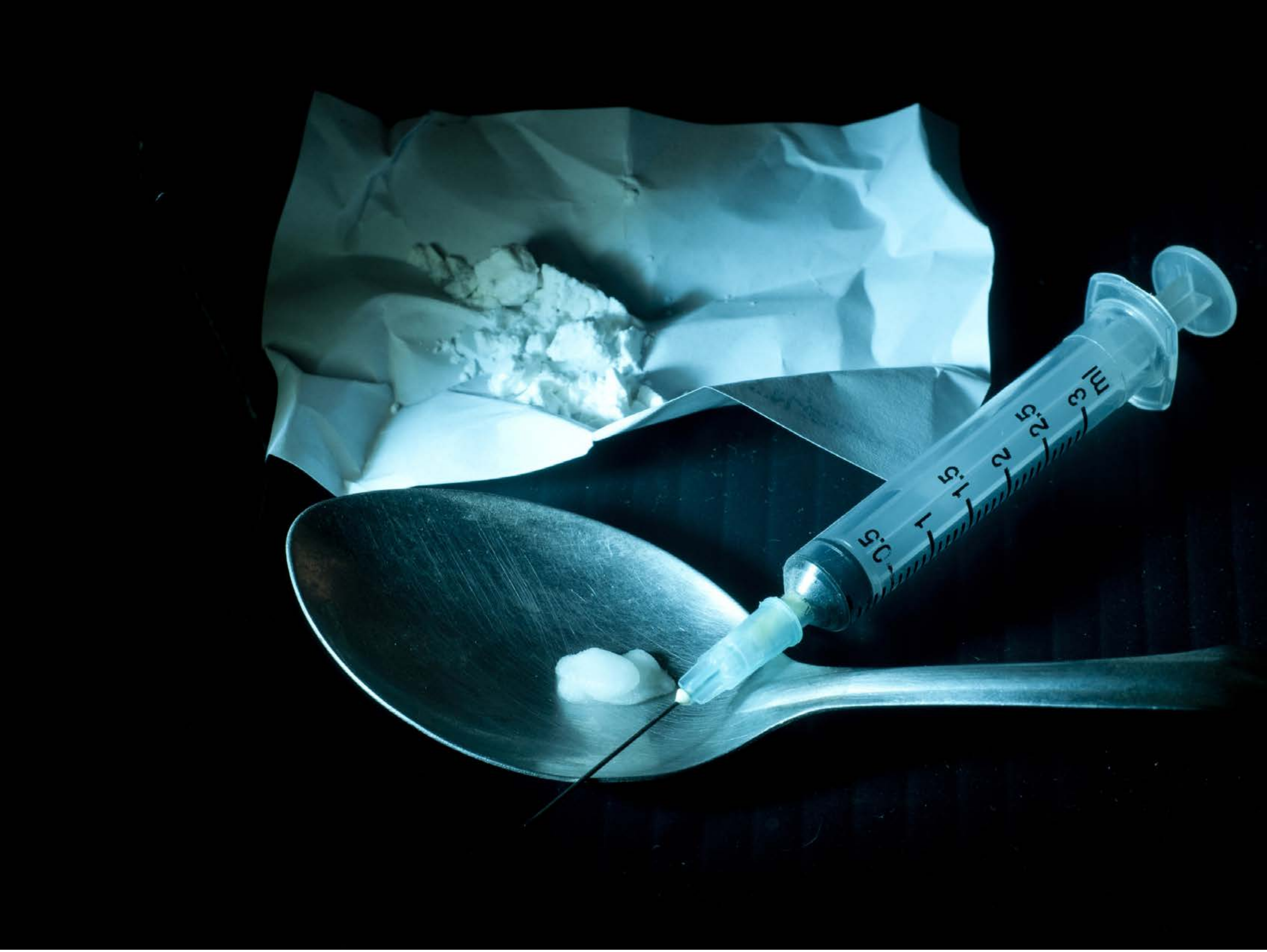

| Category | Craving (0-100) | Valence (1-9) | Arousal (1-9) | Typicality (0-100) | Relatedness | | | | | HSV | | |
|---|---|---|---|---|---|---|---|---|---|---|---|---|
| | | | | | Meth | Opioid | Both | Neither | MethToOpioid | Hue | Saturation | Value |
| Meth Instrument | 90.29 (19.86) | 4.46 (2.081) | 4.82 (3.03) | 91.79 (19.28) | 0.321 | 0.143 | 0.536 | 0 | -0.179 | 0.274 (0.279) | 0.331 (0.376) | 0.204 (0.312) |

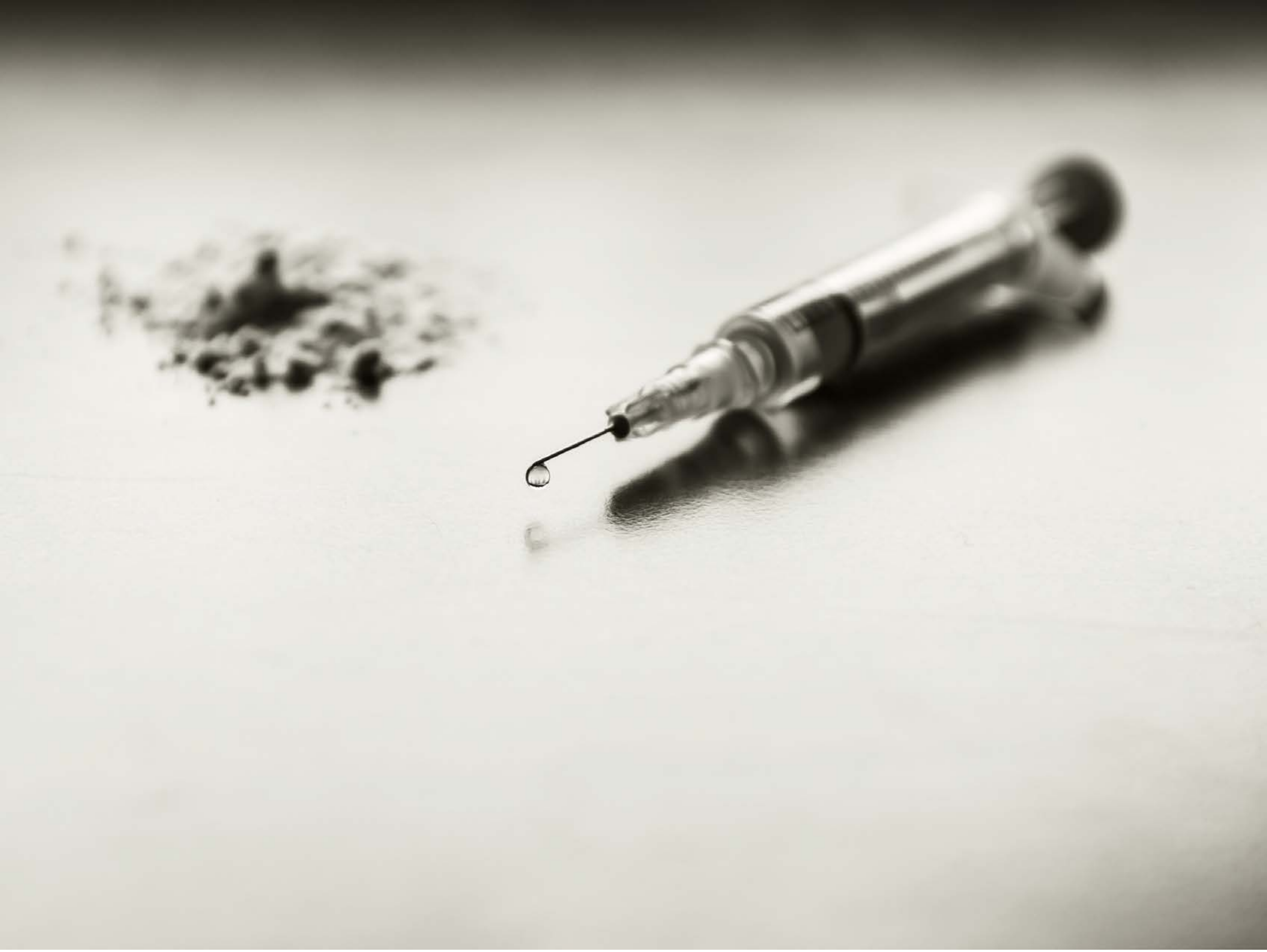

| Category | Craving (0-100) | Valence (1-9) | Arousal (1-9) | Typicality (0-100) | Relatedness | | | | | HSV | | |
|---|---|---|---|---|---|---|---|---|---|---|---|---|
| | | | | | Meth | Opioid | Both | Neither | MethToOpioid | Hue | Saturation | Value |
| Meth Instrument | 86.3 (18.7) | 4.48 (1.968) | 4.59 (2.68) | 86.44 (18.87) | 0.185 | 0.259 | 0.556 | 0 | 0.074 | 0.138 (0.019) | 0.067 (0.063) | 0.787 (0.217) |

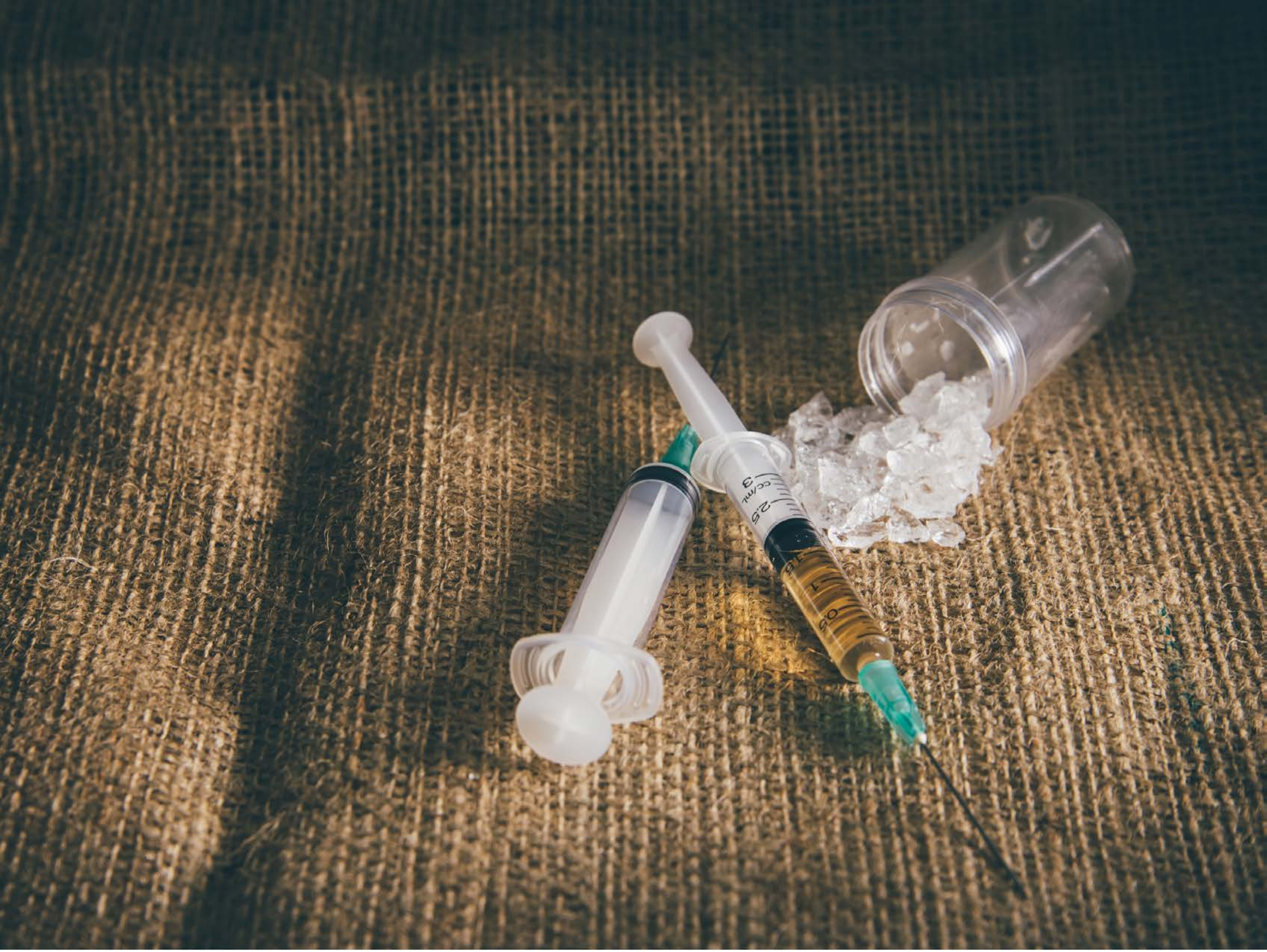


| Category | Craving (0-100) | Valence (1-9) | Arousal (1-9) | Typicality (0-100) | Relatedness | | | | | HSV | | |
|---|---|---|---|---|---|---|---|---|---|---|---|---|
| | | | | | Meth | Opioid | Both | Neither | MethToOpioid | Hue | Saturation | Value |
| Meth Instrument | 91.96 (13.69) | 4.61 (2.025) | 4.93 (2.8) | 93 (13.29) | 0.714 | 0 | 0.286 | 0 | -0.714 | 0.223 (0.187) | 0.278 (0.142) | 0.386 (0.198) |

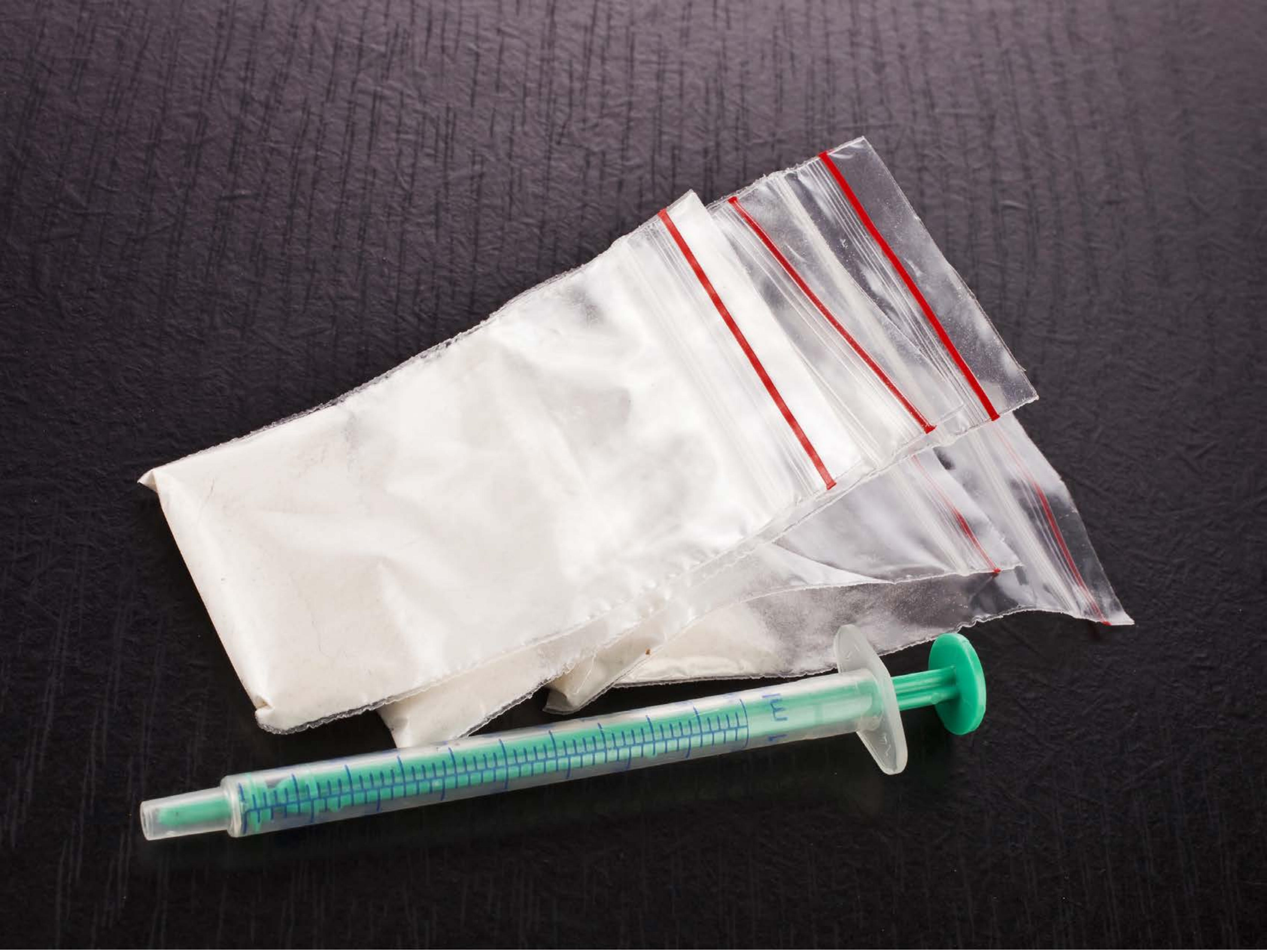

| Category | Craving (0-100) | Valence (1-9) | Arousal (1-9) | Typicality (0-100) | Relatedness | | | | | HSV | | |
|---|---|---|---|---|---|---|---|---|---|---|---|---|
| | | | | | Meth | Opioid | Both | Neither | MethToOpioid | Hue | Saturation | Value |
| Meth Instrument | 81.44 (19.38) | 4.26 (1.678) | 4.44 (2.74) | 84.89 (19.57) | 0.185 | 0.296 | 0.481 | 0.037 | 0.111 | 0.659 (0.354) | 0.116 (0.102) | 0.41 (0.309) |

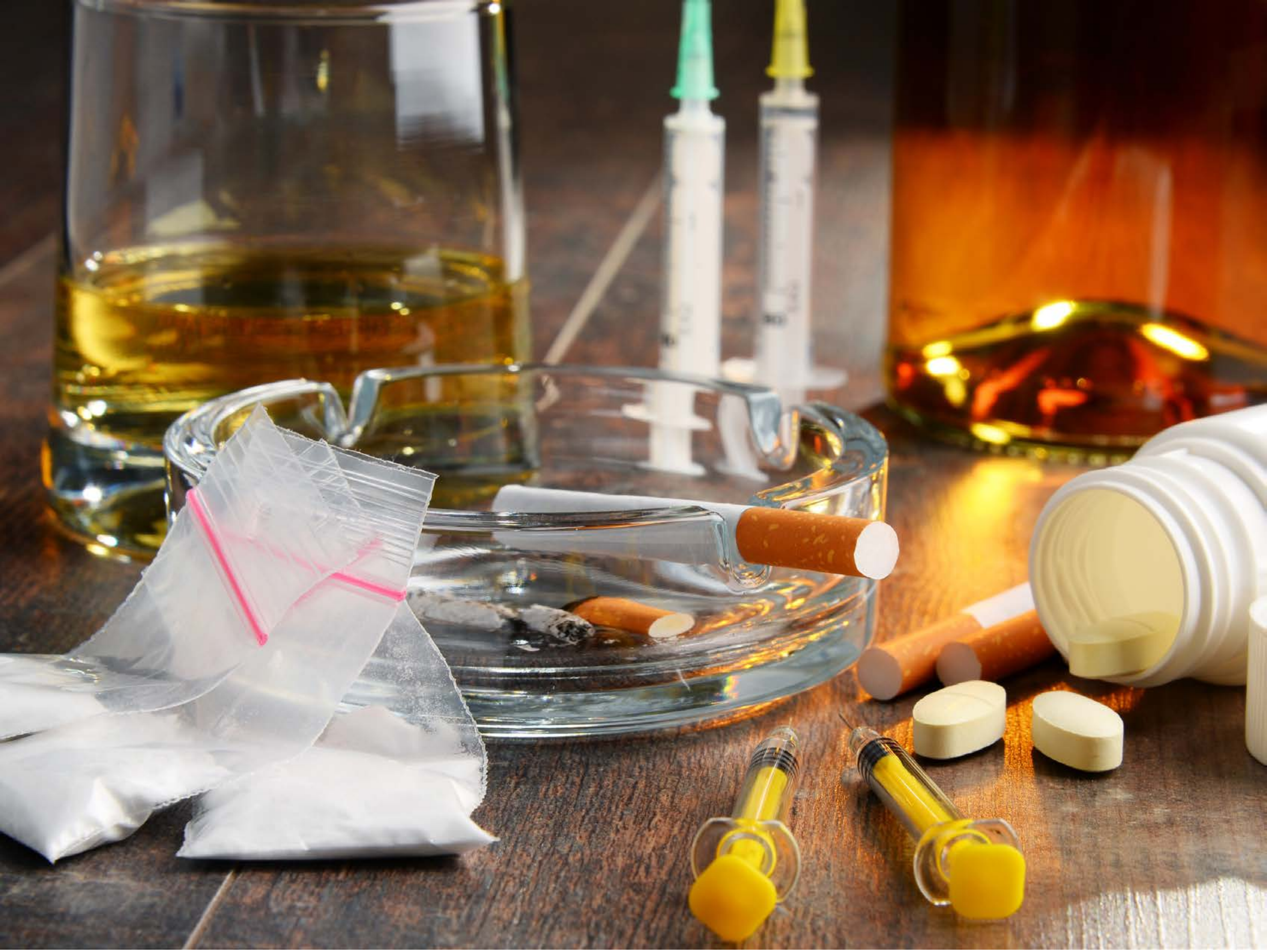

| Category | Craving (0-100) | Valence (1-9) | Arousal (1-9) | Typicality (0-100) | Relatedness | | | | | HSV | | |
|---|---|---|---|---|---|---|---|---|---|---|---|---|
| | | | | | Meth | Opioid | Both | Neither | MethToOpioid | Hue | Saturation | Value |
| Meth Instrument | 93.11 (14.23) | 4.5 (1.836) | 4.64 (2.83) | 93.54 (13.11) | 0 | 0.071 | 0.929 | 0 | 0.071 | 0.156 (0.22) | 0.419 (0.34) | 0.547 (0.259) |

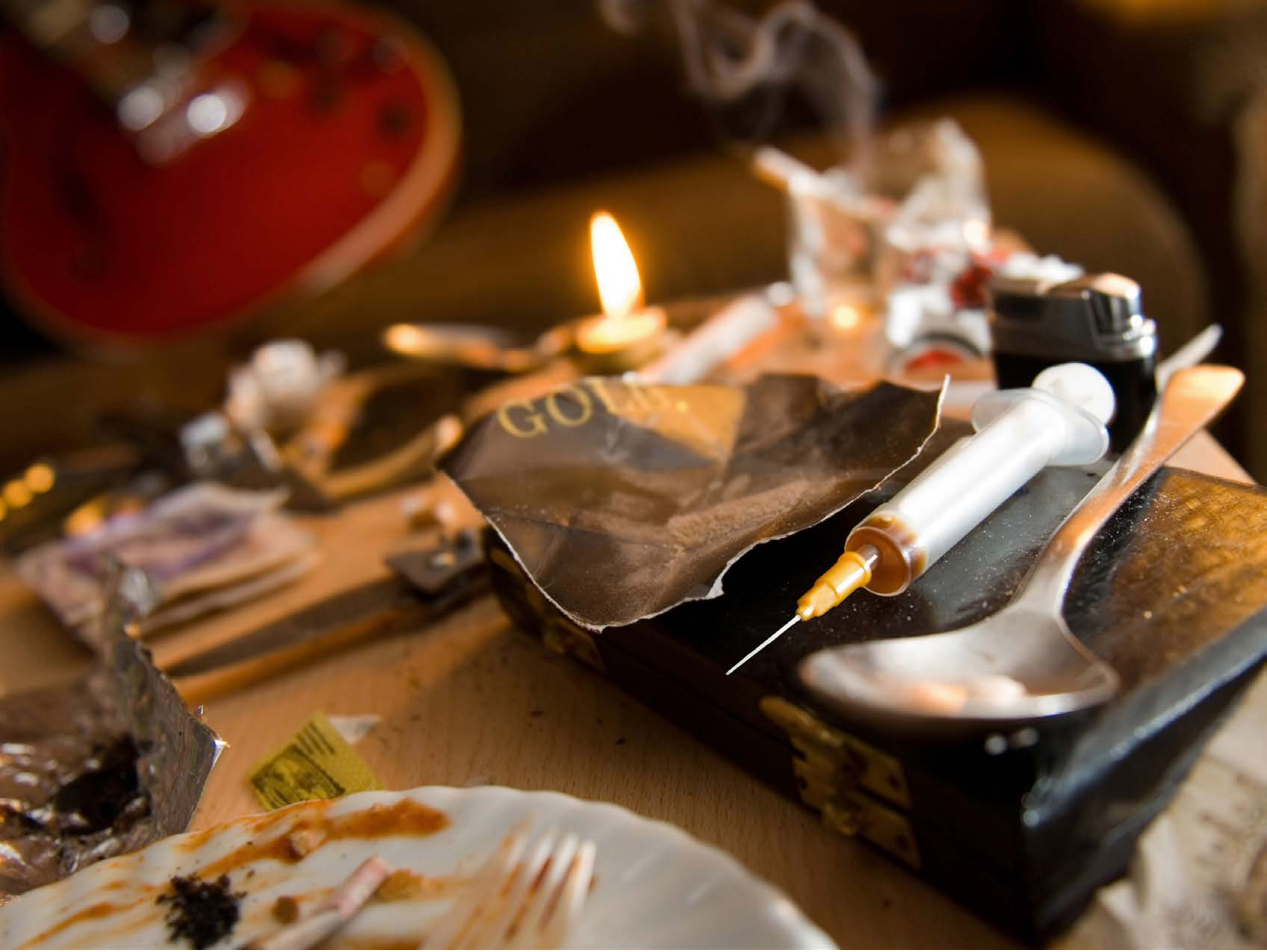


| Category | Craving (0-100) | Valence (1-9) | Arousal (1-9) | Typicality (0-100) | Relatedness | | | | | HSV | | |
|---|---|---|---|---|---|---|---|---|---|---|---|---|
| | | | | | Meth | Opioid | Both | Neither | MethToOpioid | Hue | Saturation | Value |
| Meth Instrument | 88.33 (16) | 4.3 (2.181) | 4.67 (2.88) | 89.96 (16.39) | 0.074 | 0.370 | 0.519 | 0.037 | 0.296 | 0.097 (0.152) | 0.63 (0.302) | 0.42 (0.256) |

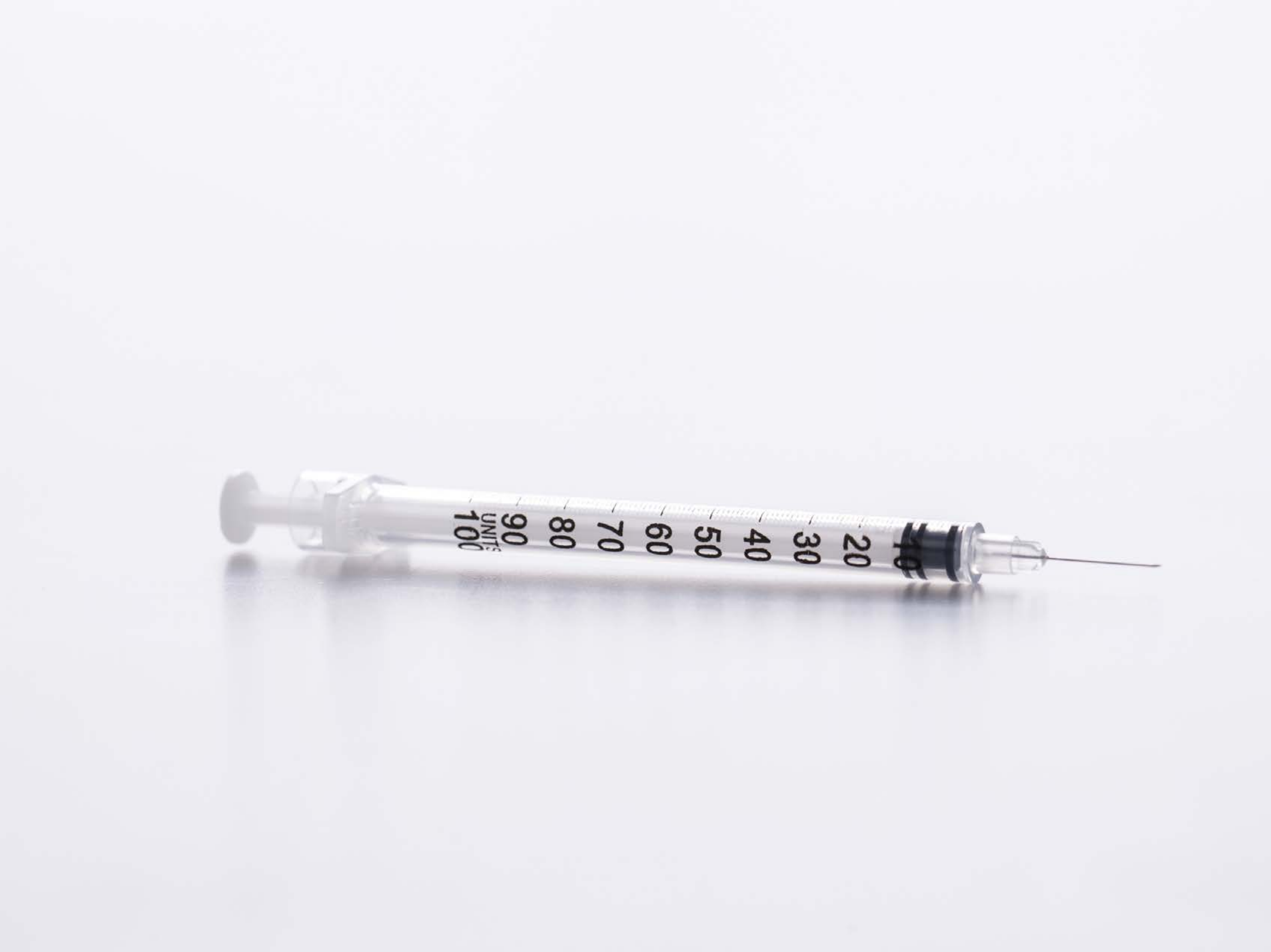


| | | | | | Relatedness | | | | | HSV | | |
|---|---|---|---|---|---|---|---|---|---|---|---|---|
| **Category** | **Craving (0-100)** | **Valence (1-9)** | **Arousal (1-9)** | **Typicality (0-100)** | Meth | Opioid | Both | Neither | MethToOpioid | Hue | Saturation | Value |
| Meth Instrument | 83.48 (20.07) | 4.63 (1.597) | 4.89 (2.5) | 82 (21.36) | 0.037 | 0.037 | 0.852 | 0.074 | 0 | 0.697 (0.055) | 0.023 (0.014) | 0.95 (0.066) |

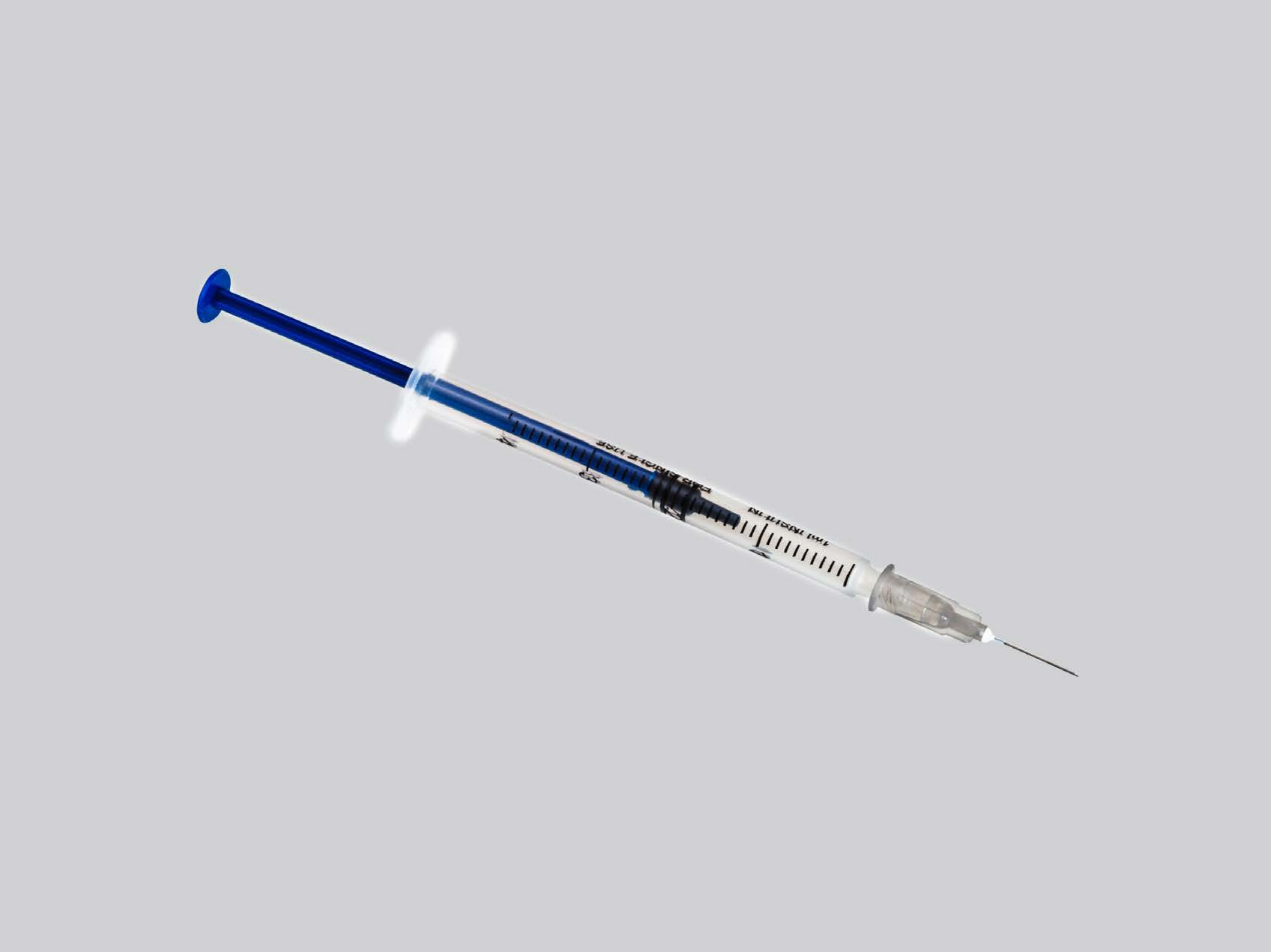

| Category | Craving (0-100) | Valence (1-9) | Arousal (1-9) | Typicality (0-100) | Relatedness | | | | | HSV | | |
|---|---|---|---|---|---|---|---|---|---|---|---|---|
| | | | | | Meth | Opioid | Both | Neither | MethToOpioid | Hue | Saturation | Value |
| Meth Instrument | 81.63 (23.8) | 4.56 (1.553) | 4.63 (2.59) | 83.33 (25.29) | 0.074 | 0 | 0.815 | 0.111 | -0.074 | 0.012 (0.085) | 0.009 (0.084) | 0.99 (0.072) |

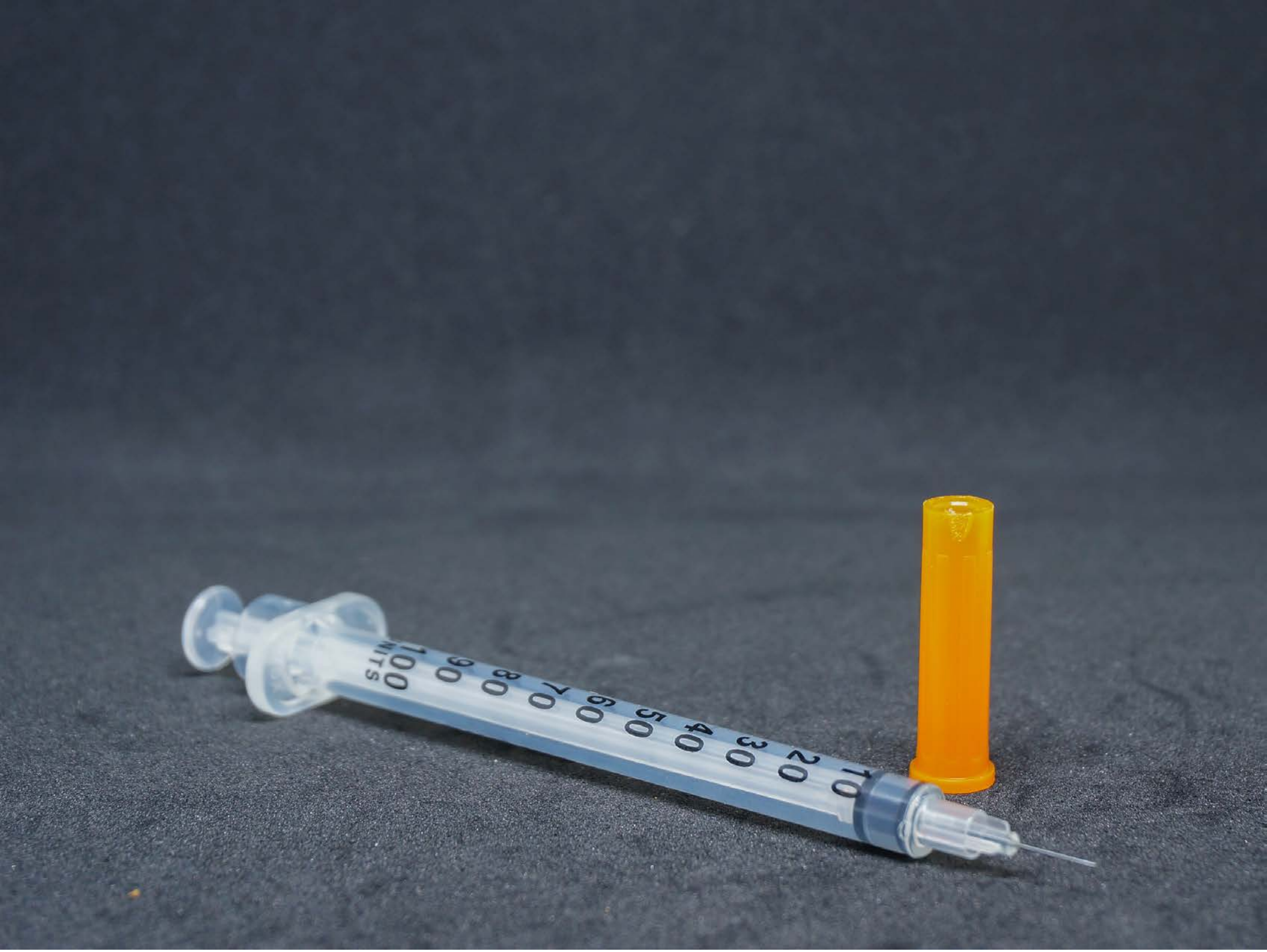

| Category | Craving (0-100) | Valence (1-9) | Arousal (1-9) | Typicality (0-100) | Relatedness | | | | | HSV | | |
|---|---|---|---|---|---|---|---|---|---|---|---|---|
| | | | | | Meth | Opioid | Both | Neither | MethToOpioid | Hue | Saturation | Value |
| Meth Instrument | 92.04 (11.67) | 4.46 (1.915) | 4.68 (2.74) | 94.39 (10.26) | 0.071 | 0 | 0.857 | 0.071 | -0.071 | 0.589 (0.07) | 0.242 (0.114) | 0.363 (0.131) |

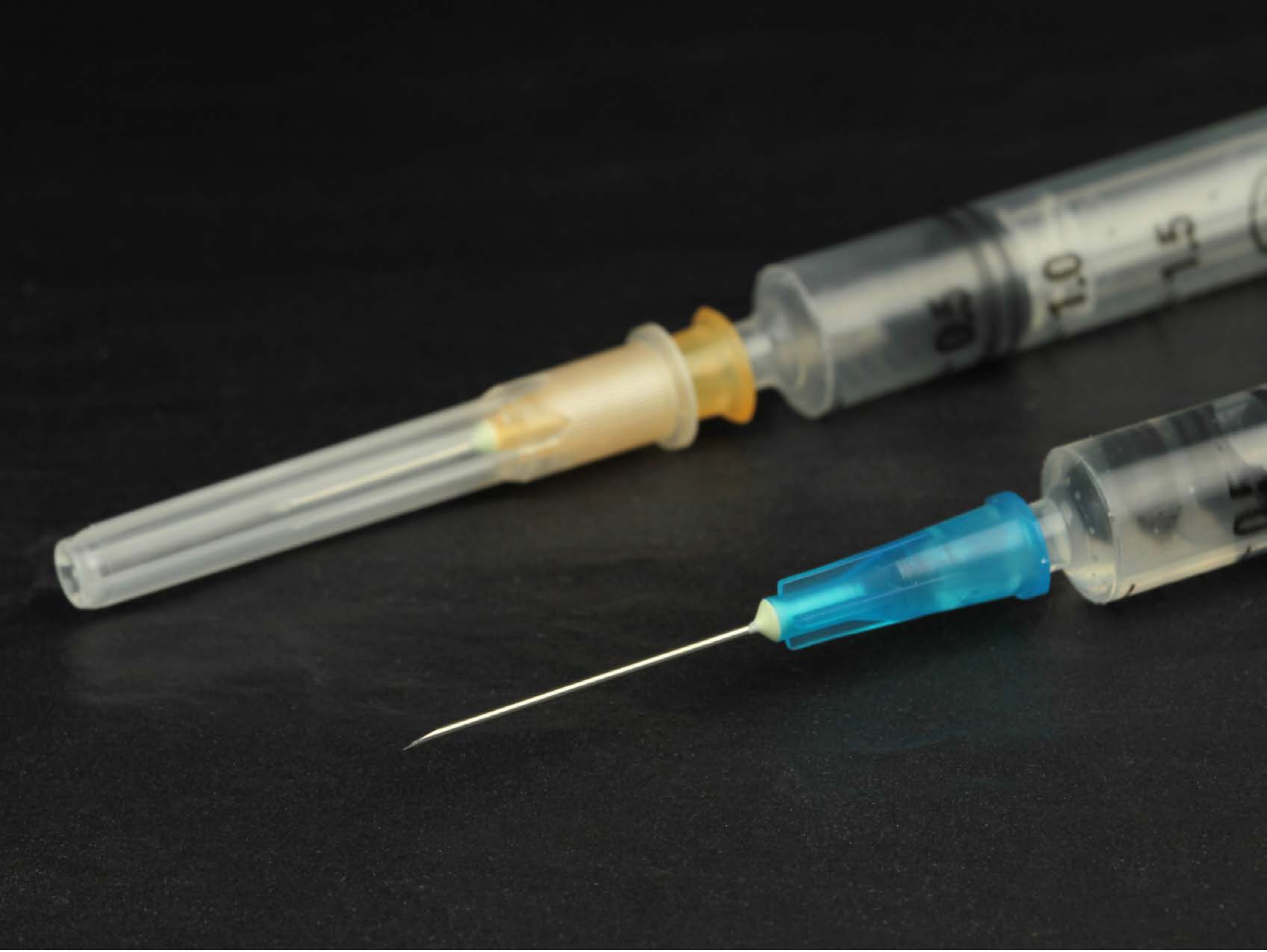


| Category | Craving (0-100) | Valence (1-9) | Arousal (1-9) | Typicality (0-100) | Relatedness | | | | | HSV | | |
|---|---|---|---|---|---|---|---|---|---|---|---|---|
| | | | | | Meth | Opioid | Both | Neither | MethToOpioid | Hue | Saturation | Value |
| Meth Instrument | 80.67 (25.5) | 4.67 (1.664) | 4.52 (2.58) | 84.44 (24.41) | 0.111 | 0 | 0.815 | 0.074 | -0.111 | 0.105 (0.158) | 0.056 (0.127) | 0.192 (0.186) |

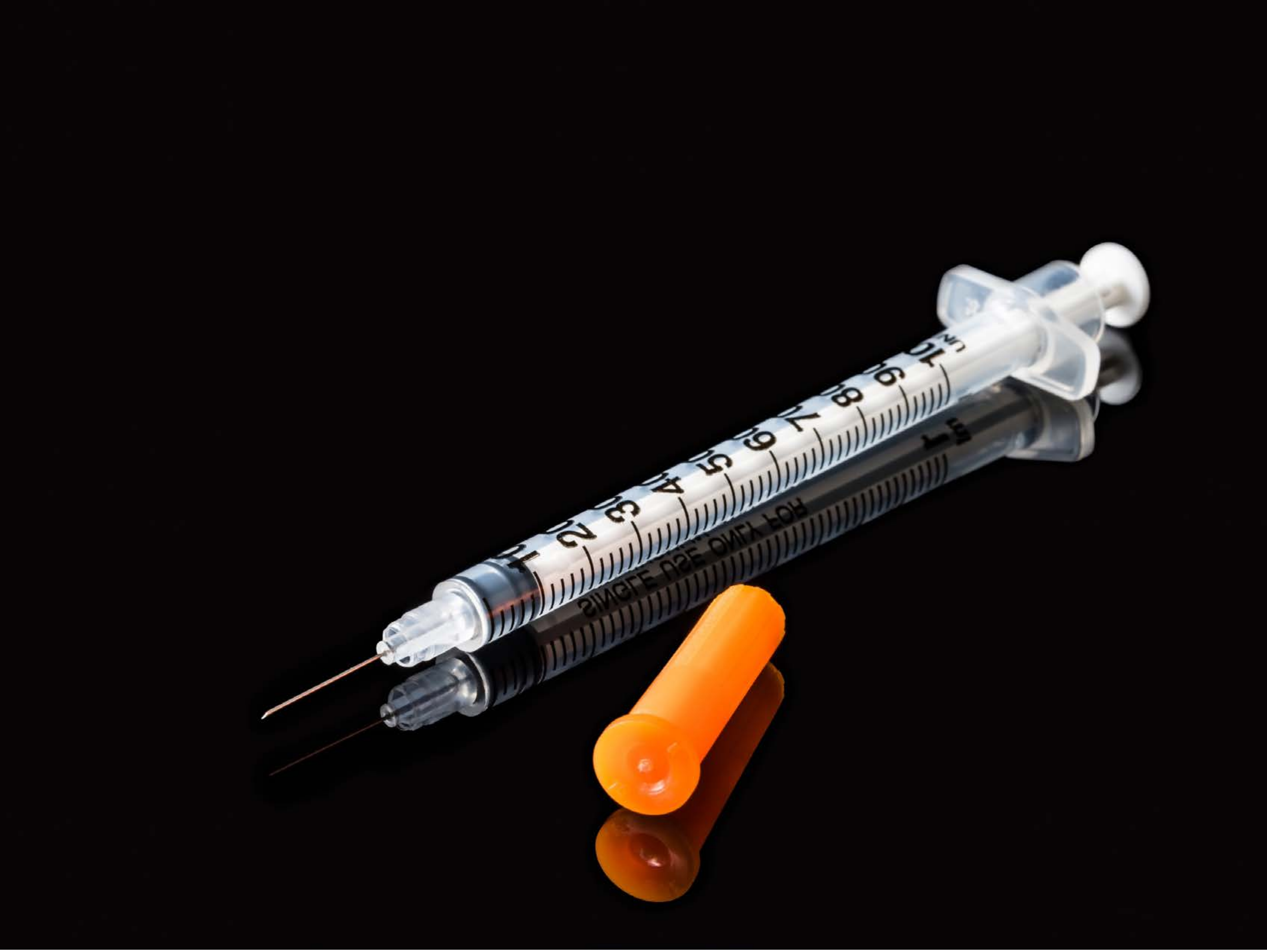


| Category | Craving (0-100) | Valence (1-9) | Arousal (1-9) | Typicality (0-100) | Relatedness | | | | | HSV | | |
|---|---|---|---|---|---|---|---|---|---|---|---|---|
| | | | | | Meth | Opioid | Both | Neither | MethToOpioid | Hue | Saturation | Value |
| Meth Instrument | 92.46 (14) | 4.32 (2.109) | 4.82 (2.67) | 94 (13.12) | 0.071 | 0 | 0.893 | 0.036 | -0.071 | 0.816 (0.261) | 0.291 (0.147) | 0.097 (0.219) |

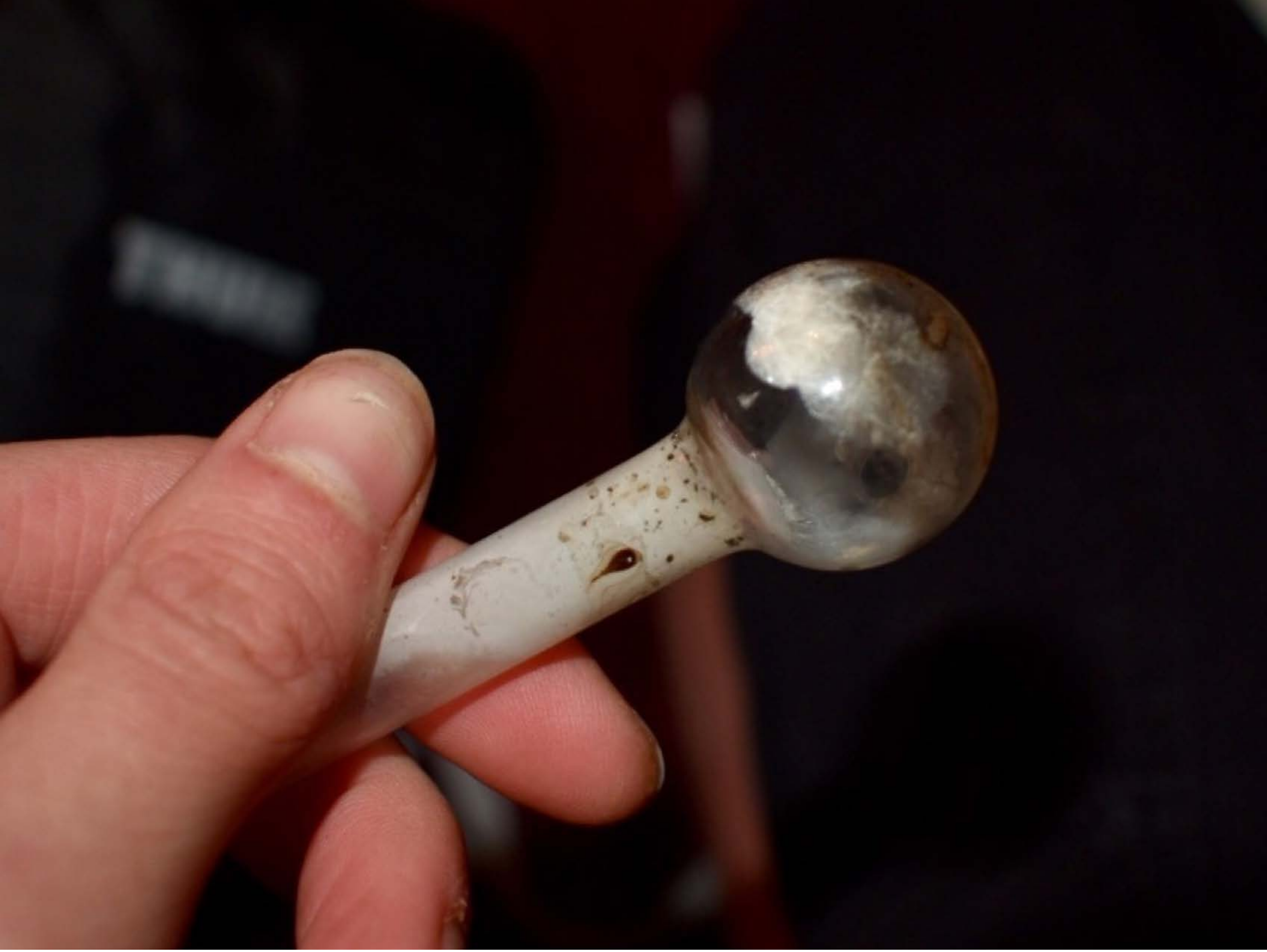

| Category | Craving (0-100) | Valence (1-9) | Arousal (1-9) | Typicality (0-100) | Relatedness | | | | | HSV | | |
|---|---|---|---|---|---|---|---|---|---|---|---|---|
| | | | | | Meth | Opioid | Both | Neither | MethToOpioid | Hue | Saturation | Value |
| Meth Instrument Hand | 92.18 (12.15) | 4.36 (1.89) | 5.04 (2.71) | 95 (10.67) | 1 | 0 | 0 | 0 | -1 | 0.497 (0.428) | 0.592 (0.286) | 0.225 (0.278) |

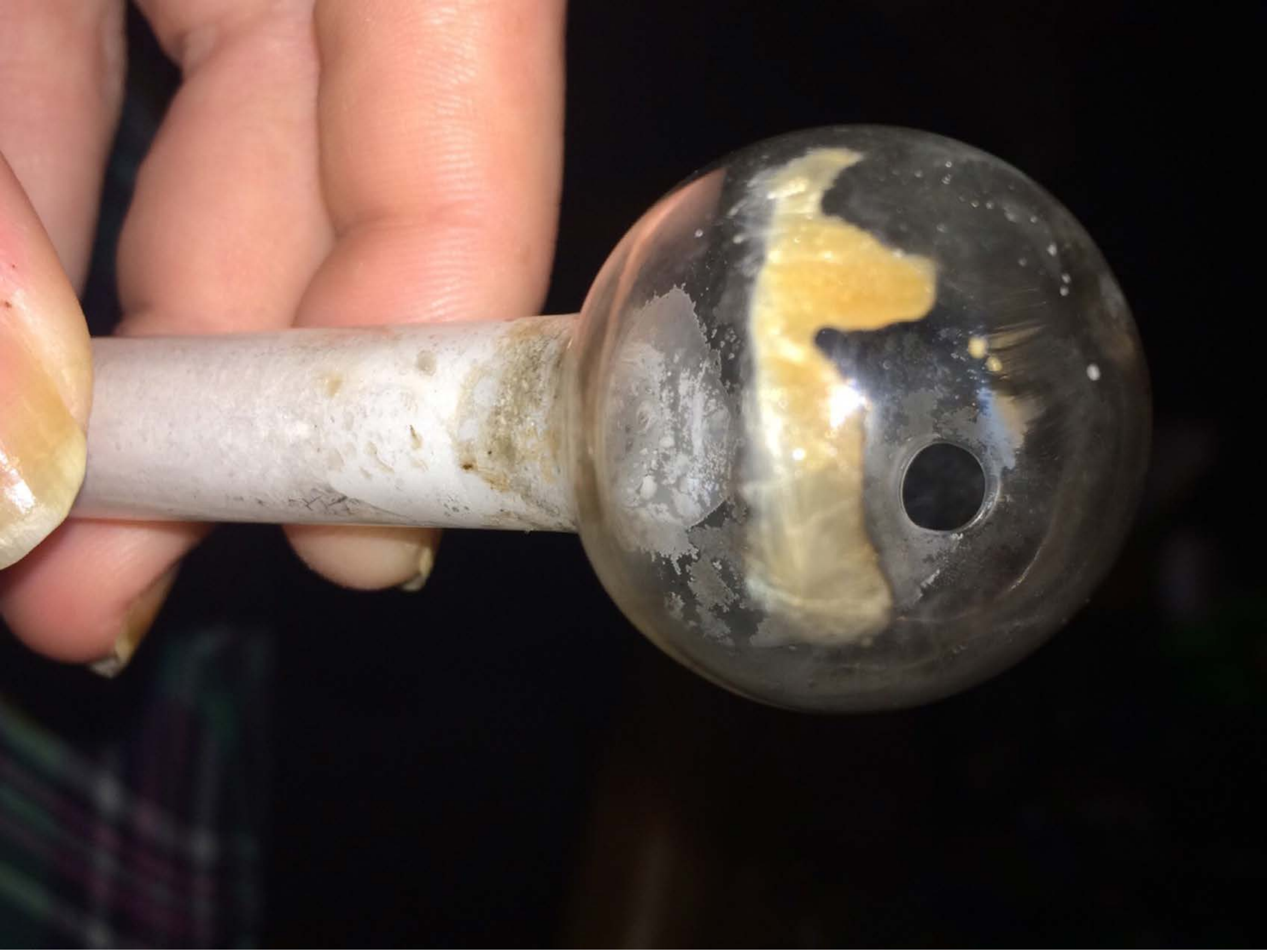

| Category | Craving (0-100) | Valence (1-9) | Arousal (1-9) | Typicality (0-100) | Relatedness | | | | | HSV | | |
|---|---|---|---|---|---|---|---|---|---|---|---|---|
| | | | | | Meth | Opioid | Both | Neither | MethToOpioid | Hue | Saturation | Value |
| Meth Instrument Hand | 89.04 (15.17) | 4.3 (1.958) | 5.07 (2.67) | 88.07 (17.12) | 1 | 0 | 0 | 0 | -1 | 0.31 (0.356) | 0.26 (0.168) | 0.337 (0.352) |

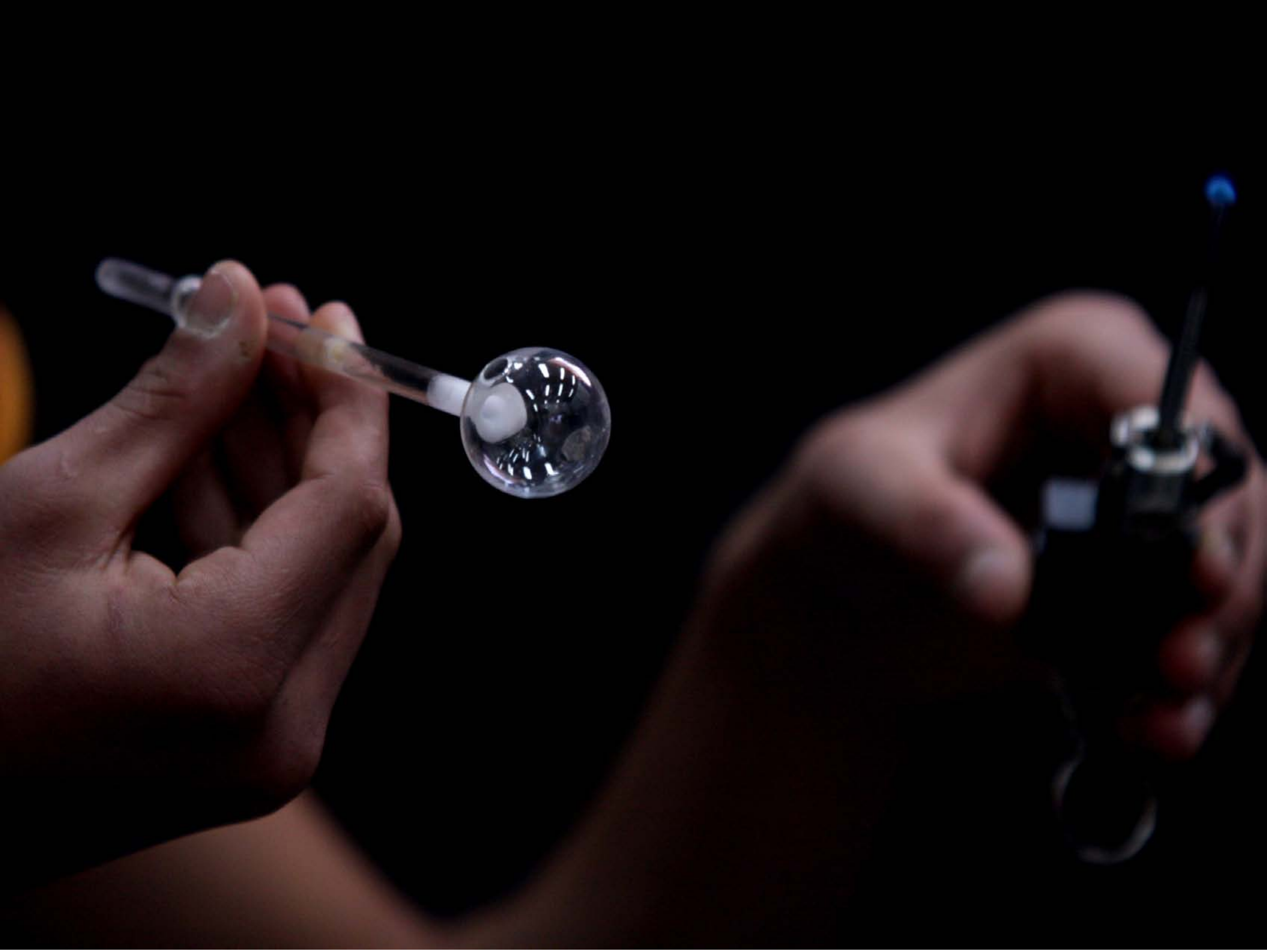

| Category | Craving (0-100) | Valence (1-9) | Arousal (1-9) | Typicality (0-100) | Relatedness | | | | | HSV | | |
|---|---|---|---|---|---|---|---|---|---|---|---|---|
| | | | | | Meth | Opioid | Both | Neither | MethToOpioid | Hue | Saturation | Value |
| Meth Instrument Hand | 86 (19.23) | 4.52 (1.988) | 4.78 (2.74) | 87.85 (19.28) | 0.926 | 0 | 0.074 | 0 | -0.926 | 0.196 (0.37) | 0.259 (0.36) | 0.083 (0.152) |

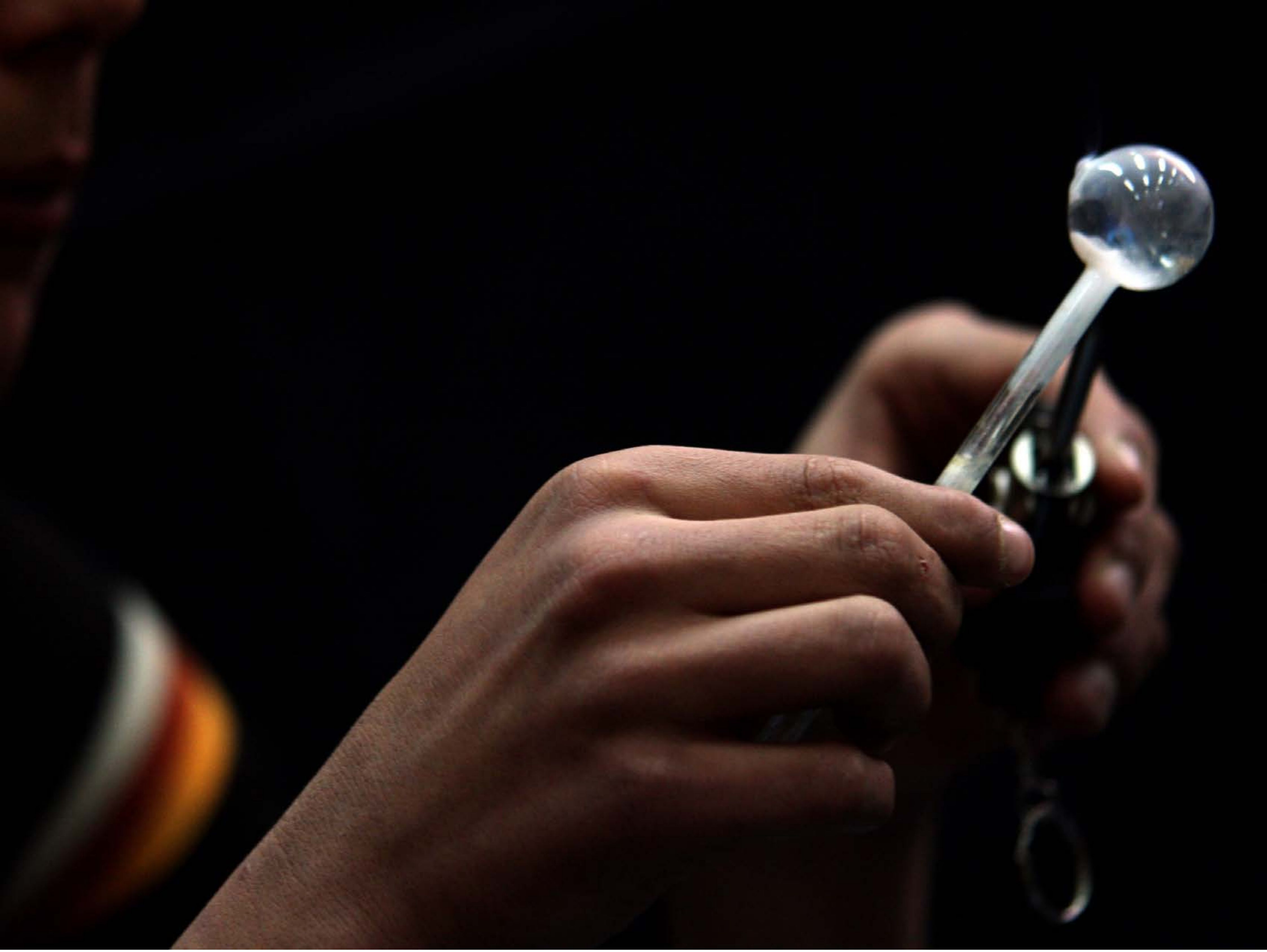

| Category | Craving (0-100) | Valence (1-9) | Arousal (1-9) | Typicality (0-100) | Relatedness | | | | | HSV | | |
|---|---|---|---|---|---|---|---|---|---|---|---|---|
| | | | | | Meth | Opioid | Both | Neither | MethToOpioid | Hue | Saturation | Value |
| Meth Instrument Hand | 83.26 (23.52) | 4.48 (1.889) | 4.48 (2.69) | 83.81 (24.9) | 0.889 | 0 | 0.111 | 0 | -0.889 | 0.299 (0.341) | 0.5 (0.354) | 0.11 (0.189) |

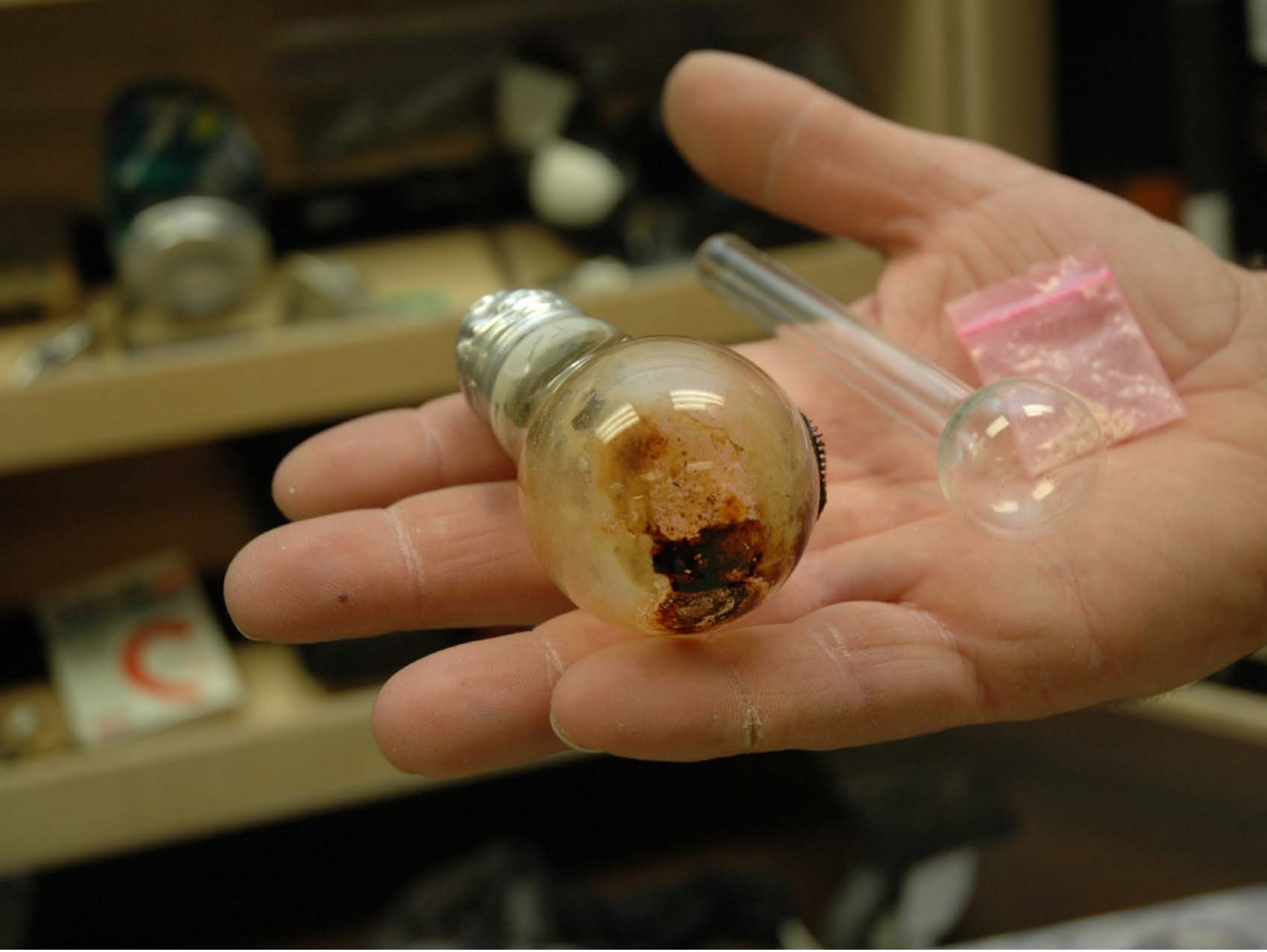

| Category | Craving (0-100) | Valence (1-9) | Arousal (1-9) | Typicality (0-100) | Relatedness | | | | | HSV | | |
|---|---|---|---|---|---|---|---|---|---|---|---|---|
| | | | | | Meth | Opioid | Both | Neither | MethToOpioid | Hue | Saturation | Value |
| Meth Instrument Hand | 92.21 (12.09) | 4.32 (1.945) | 4.79 (2.86) | 93.18 (12.36) | 0.893 | 0 | 0.107 | 0 | -0.893 | 0.109 (0.078) | 0.468 (0.203) | 0.377 (0.254) |

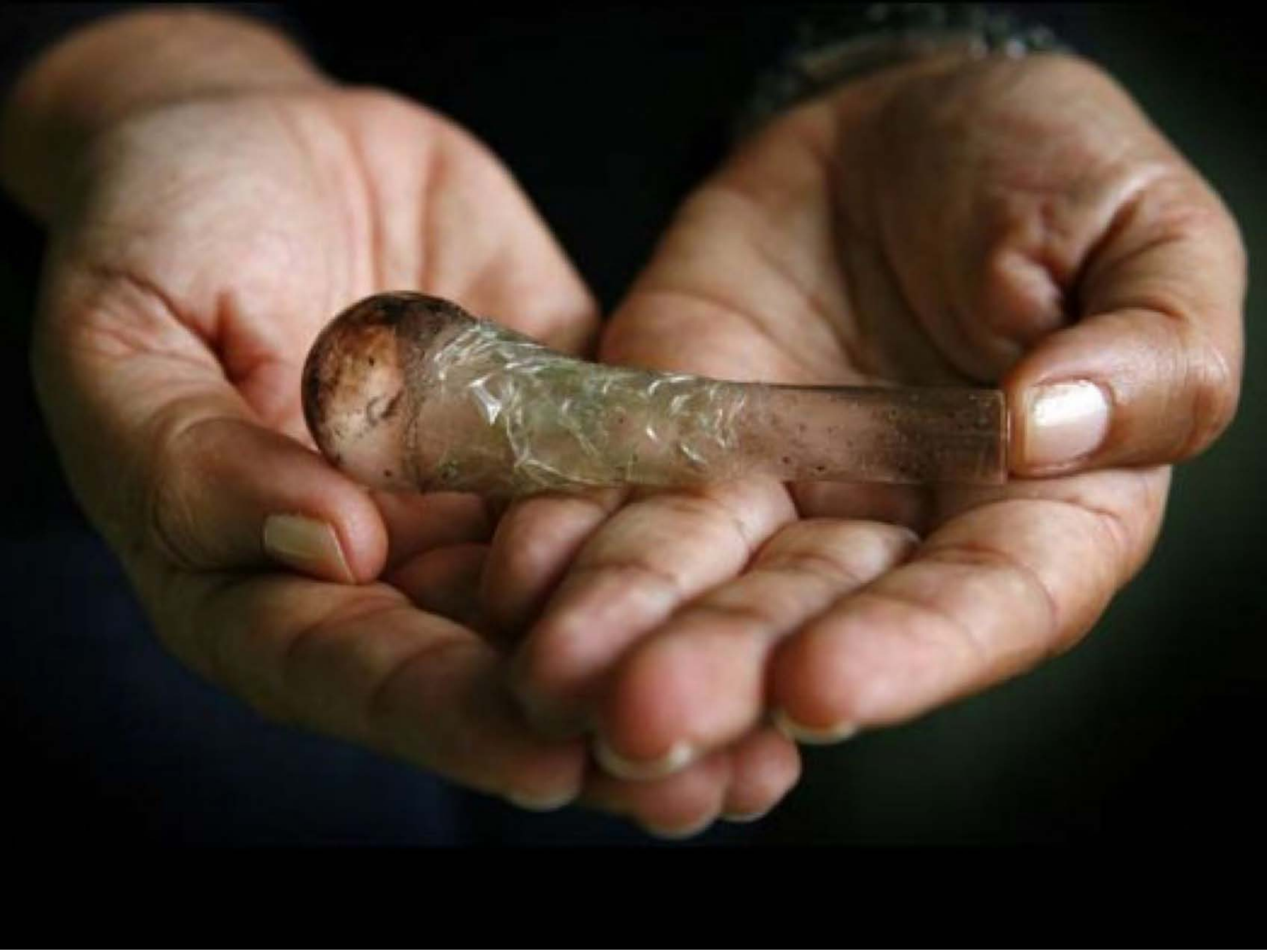

| Category | Craving (0-100) | Valence (1-9) | Arousal (1-9) | Typicality (0-100) | Relatedness | | | | | HSV | | |
|---|---|---|---|---|---|---|---|---|---|---|---|---|
| | | | | | Meth | Opioid | Both | Neither | MethToOpioid | Hue | Saturation | Value |
| Meth Instrument Hand | 81.67 (24.5) | 4.3 (1.75) | 4.63 (2.56) | 81.3 (25.23) | 0.889 | 0 | 0.074 | 0.037 | -0.889 | 0.139 (0.174) | 0.419 (0.258) | 0.311 (0.288) |

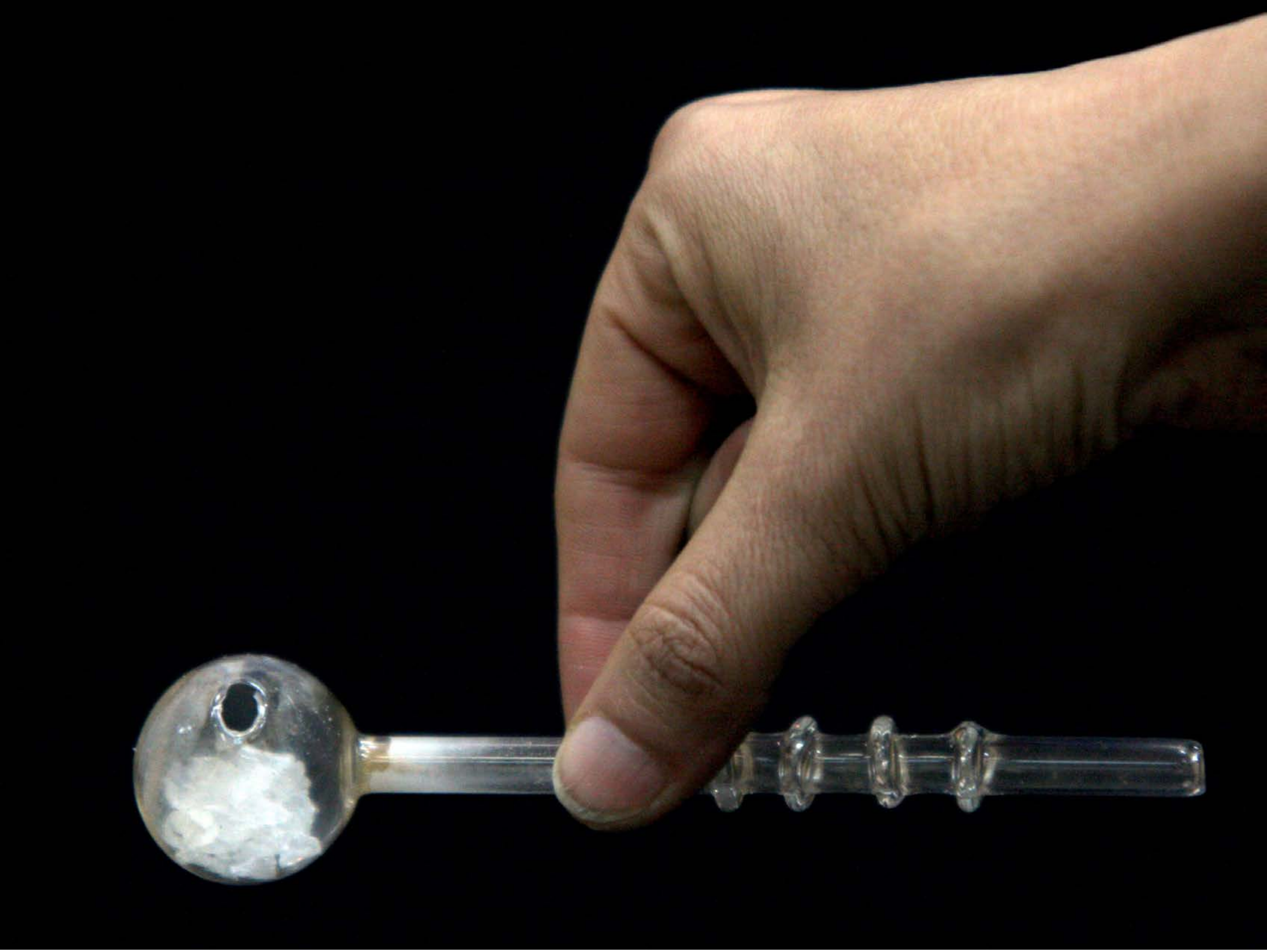

| Category | Craving (0-100) | Valence (1-9) | Arousal (1-9) | Typicality (0-100) | Relatedness | | | | | HSV | | |
|---|---|---|---|---|---|---|---|---|---|---|---|---|
| | | | | | Meth | Opioid | Both | Neither | MethToOpioid | Hue | Saturation | Value |
| Meth Instrument Hand | 90.14 (20.9) | 4.25 (1.898) | 4.75 (2.66) | 91.43 (20.45) | 1 | 0 | 0 | 0 | -1 | 0.226 (0.295) | 0.388 (0.402) | 0.18 (0.281) |

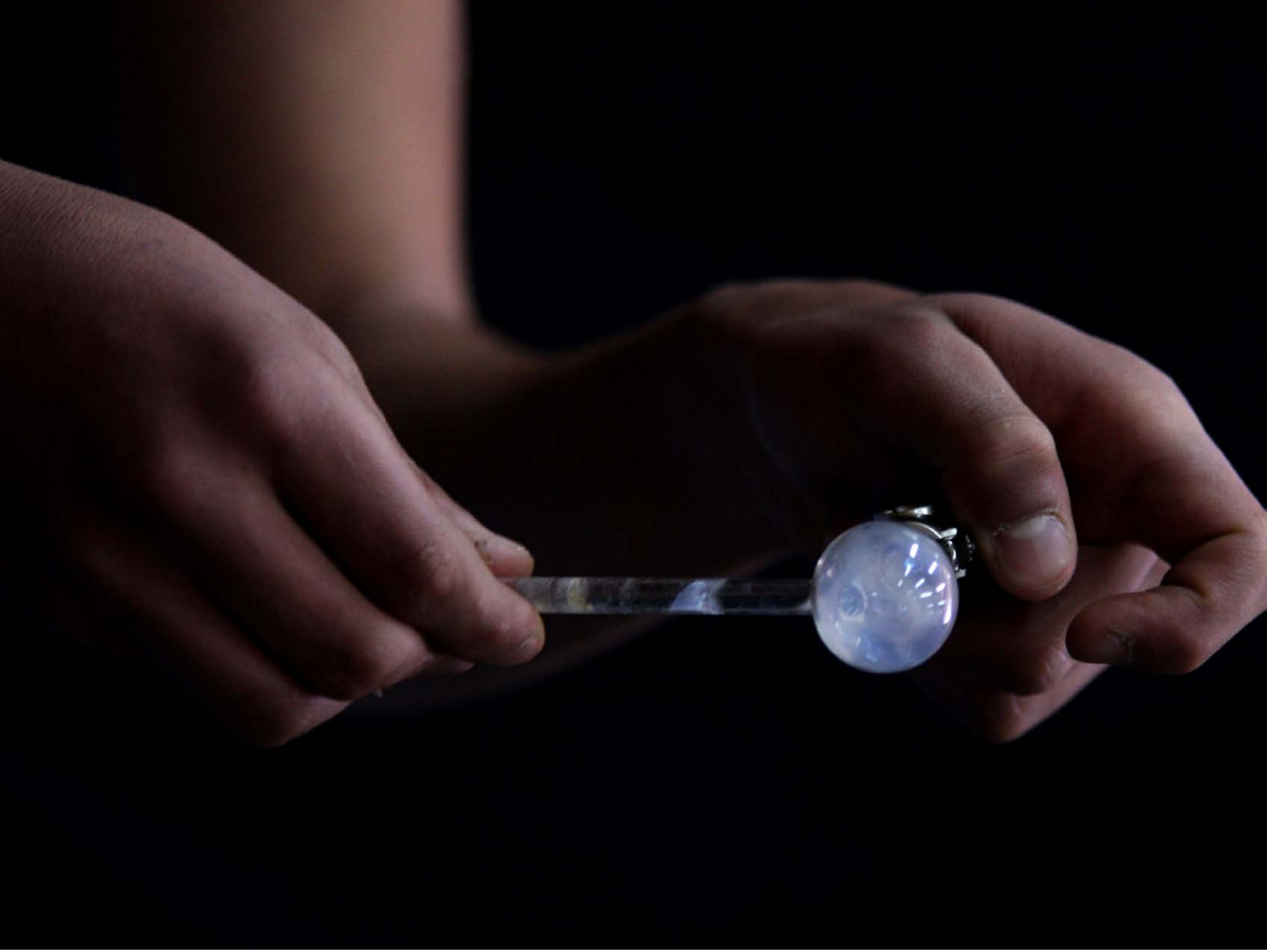

| Category | Craving (0-100) | Valence (1-9) | Arousal (1-9) | Typicality (0-100) | Relatedness | | | | | HSV | | |
|---|---|---|---|---|---|---|---|---|---|---|---|---|
| | | | | | Meth | Opioid | Both | Neither | MethToOpioid | Hue | Saturation | Value |
| Meth Instrument Hand | 89.29 (21.13) | 4.07 (2.089) | 4.61 (2.87) | 90.5 (20.93) | 0.893 | 0 | 0.107 | 0 | -0.893 | 0.377 (0.398) | 0.498 (0.399) | 0.1 (0.166) |

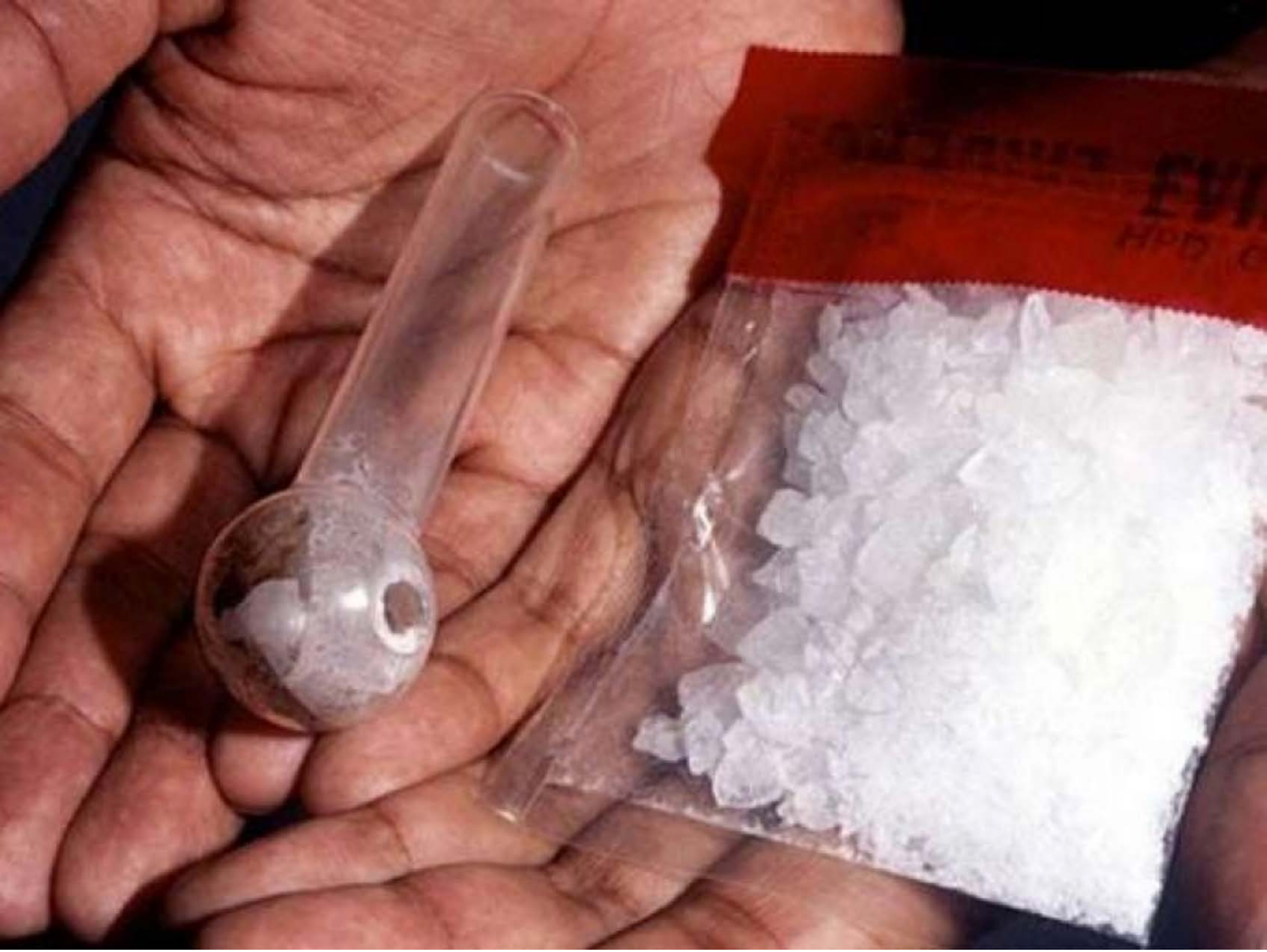

| Category | Craving (0-100) | Valence (1-9) | Arousal (1-9) | Typicality (0-100) | Relatedness | | | | | HSV | | |
| --- | --- | --- | --- | --- | --- | --- | --- | --- | --- | --- | --- | --- |
| | | | | | Meth | Opioid | Both | Neither | MethToOpioid | Hue | Saturation | Value |
| Meth Instrument Hand | 93.64 (12.24) | 4.25 (2.137) | 5.11 (2.87) | 95.43 (10.87) | 1 | 0 | 0 | 0 | -1 | 0.264 (0.389) | 0.427 (0.297) | 0.62 (0.243) |

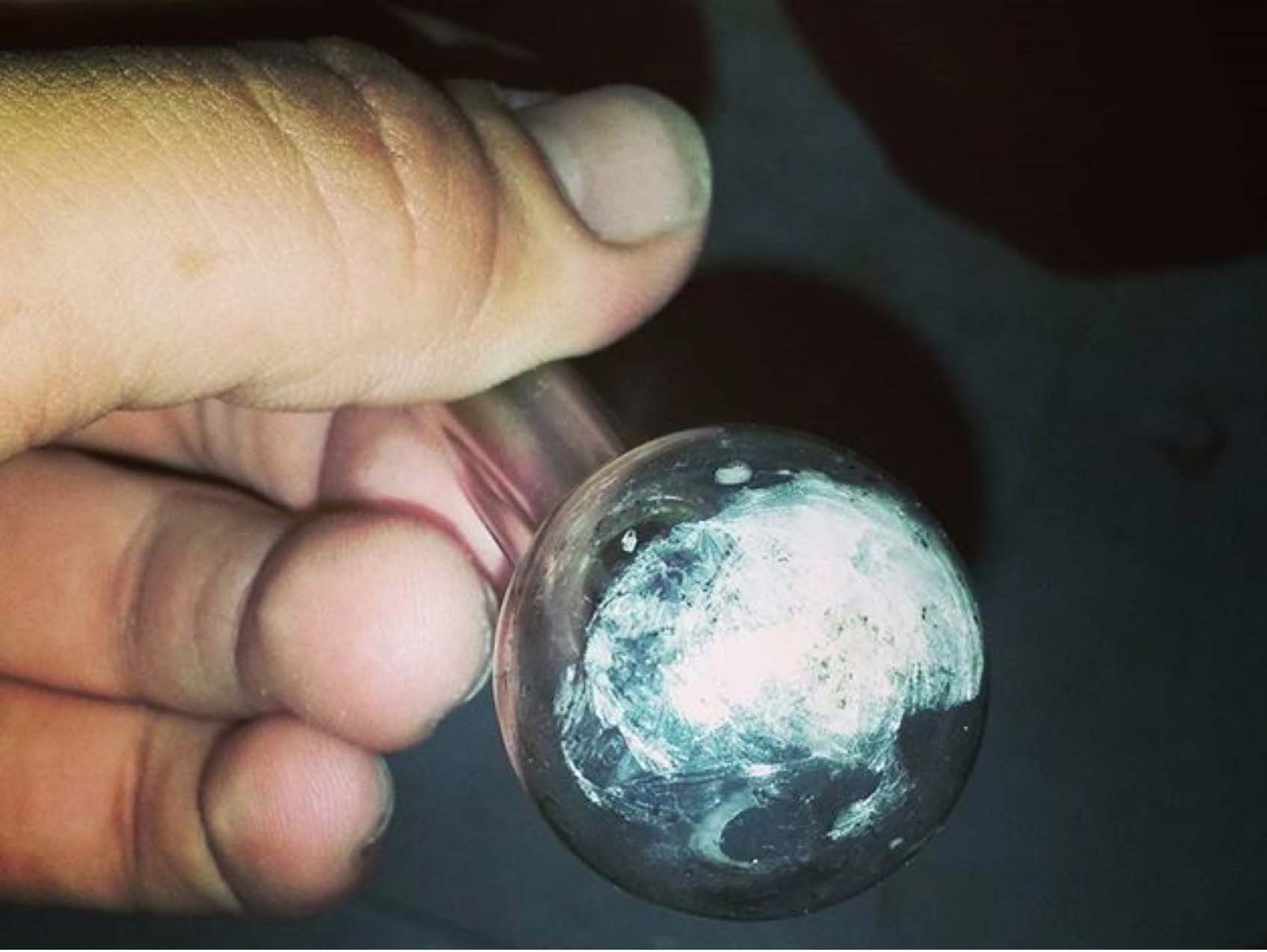

| Category | Craving (0-100) | Valence (1-9) | Arousal (1-9) | Typicality (0-100) | Relatedness | | | | | HSV | | |
|---|---|---|---|---|---|---|---|---|---|---|---|---|
| | | | | | Meth | Opioid | Both | Neither | MethToOpioid | Hue | Saturation | Value |
| Meth Instrument Hand | 85.41 (19.17) | 4.07 (1.94) | 4.3 (2.78) | 87.89 (18.57) | 0.963 | 0 | 0.037 | 0 | -0.963 | 0.272 (0.254) | 0.31 (0.142) | 0.421 (0.298) |

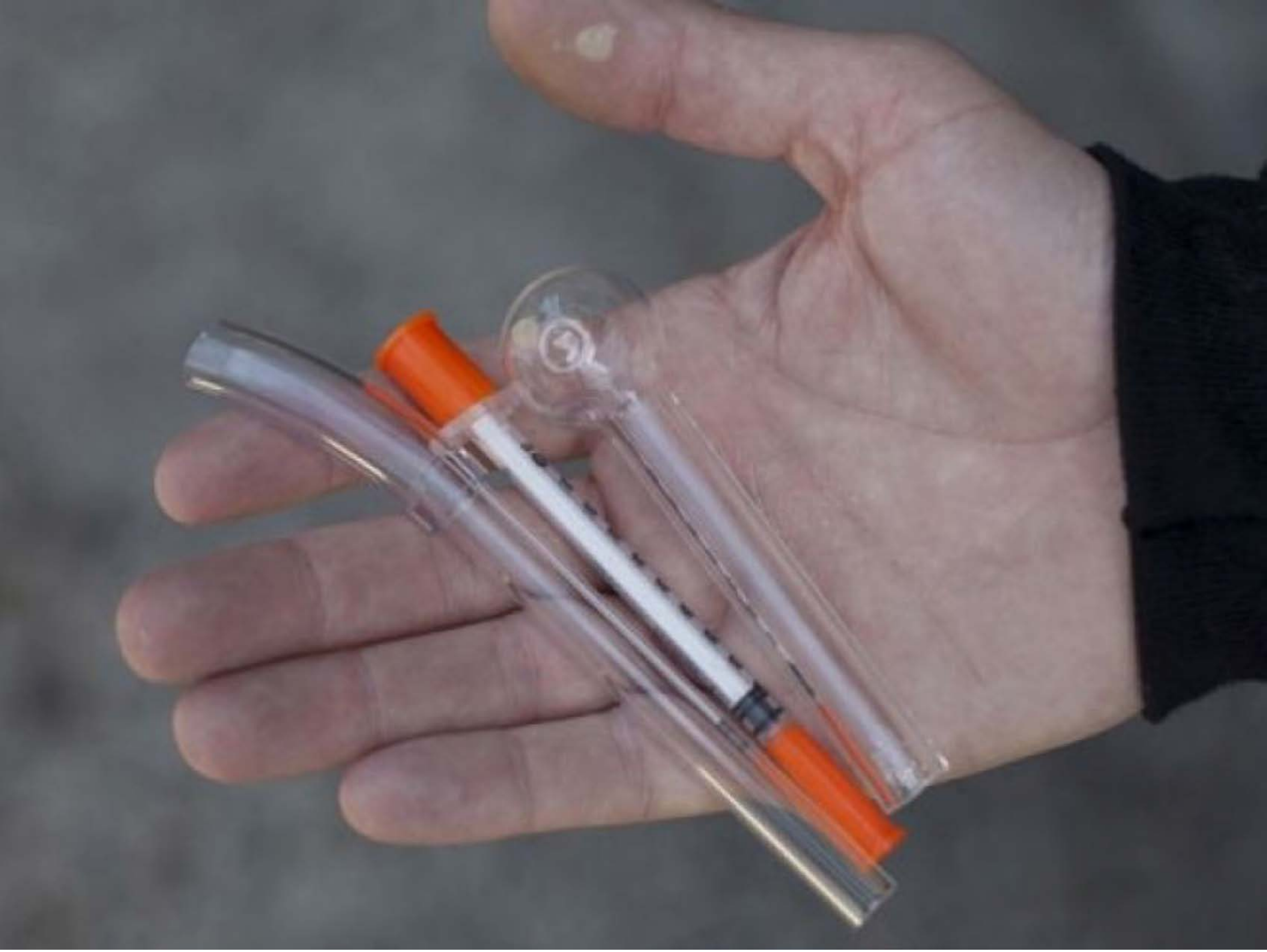

| Category | Craving (0-100) | Valence (1-9) | Arousal (1-9) | Typicality (0-100) | Relatedness | | | | | HSV | | |
|---|---|---|---|---|---|---|---|---|---|---|---|---|
| | | | | | Meth | Opioid | Both | Neither | MethToOpioid | Hue | Saturation | Value |
| Meth Instrument Hand | 82.85 (20) | 4.63 (1.904) | 4.48 (2.65) | 86.48 (18.72) | 0.556 | 0 | 0.444 | 0 | -0.556 | 0.641 (0.302) | 0.154 (0.126) | 0.444 (0.152) |

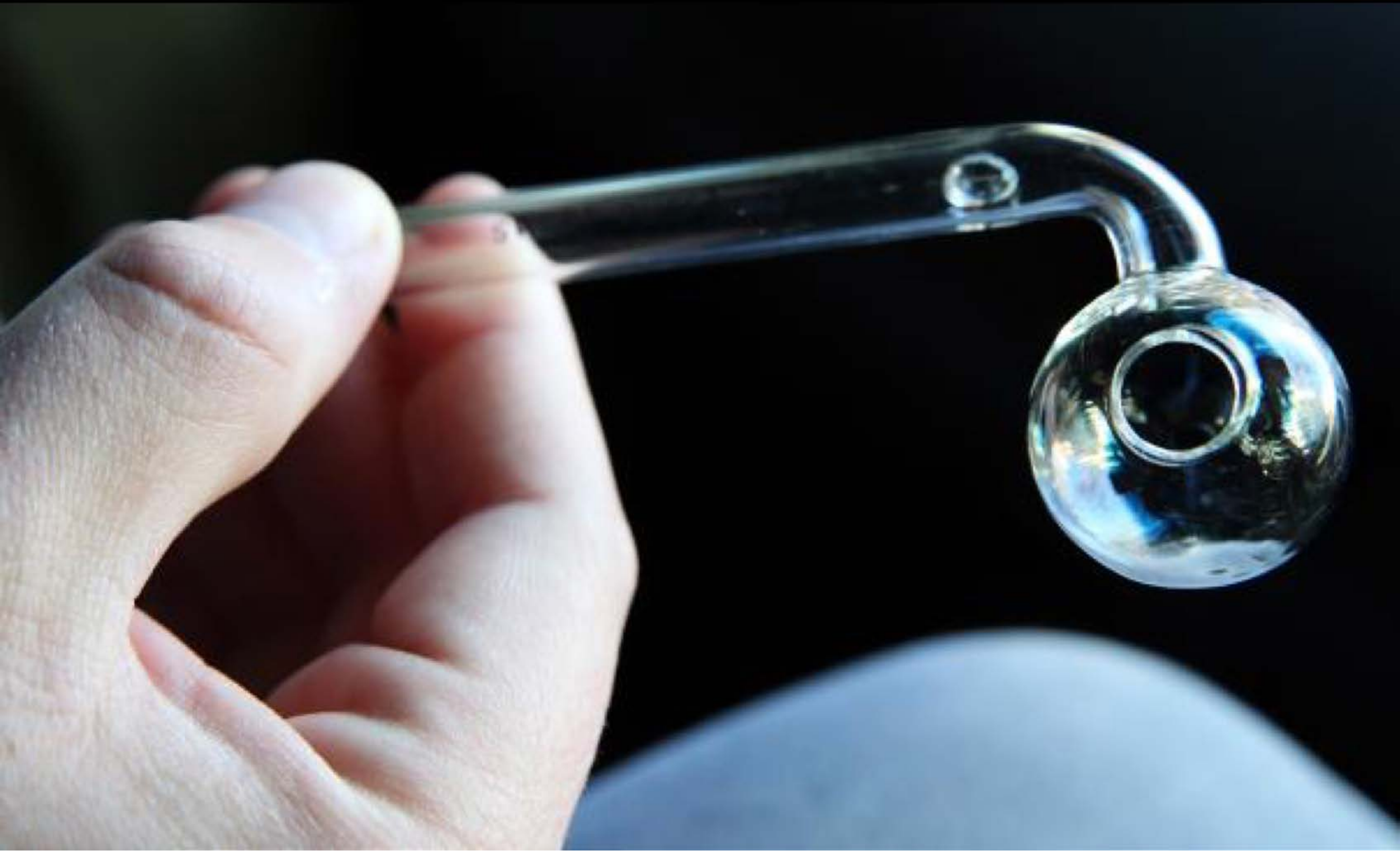

| Category | Craving (0-100) | Valence (1-9) | Arousal (1-9) | Typicality (0-100) | Relatedness | | | | | HSV | | |
|---|---|---|---|---|---|---|---|---|---|---|---|---|
| | | | | | Meth | Opioid | Both | Neither | MethToOpioid | Hue | Saturation | Value |
| Meth Instrument Hand | 87.32 (18.57) | 4.61 (1.853) | 4.46 (2.62) | 84.64 (26.56) | 0.857 | 0 | 0.107 | 0.036 | -0.857 | 0.324 (0.33) | 0.2 (0.242) | 0.296 (0.358) |

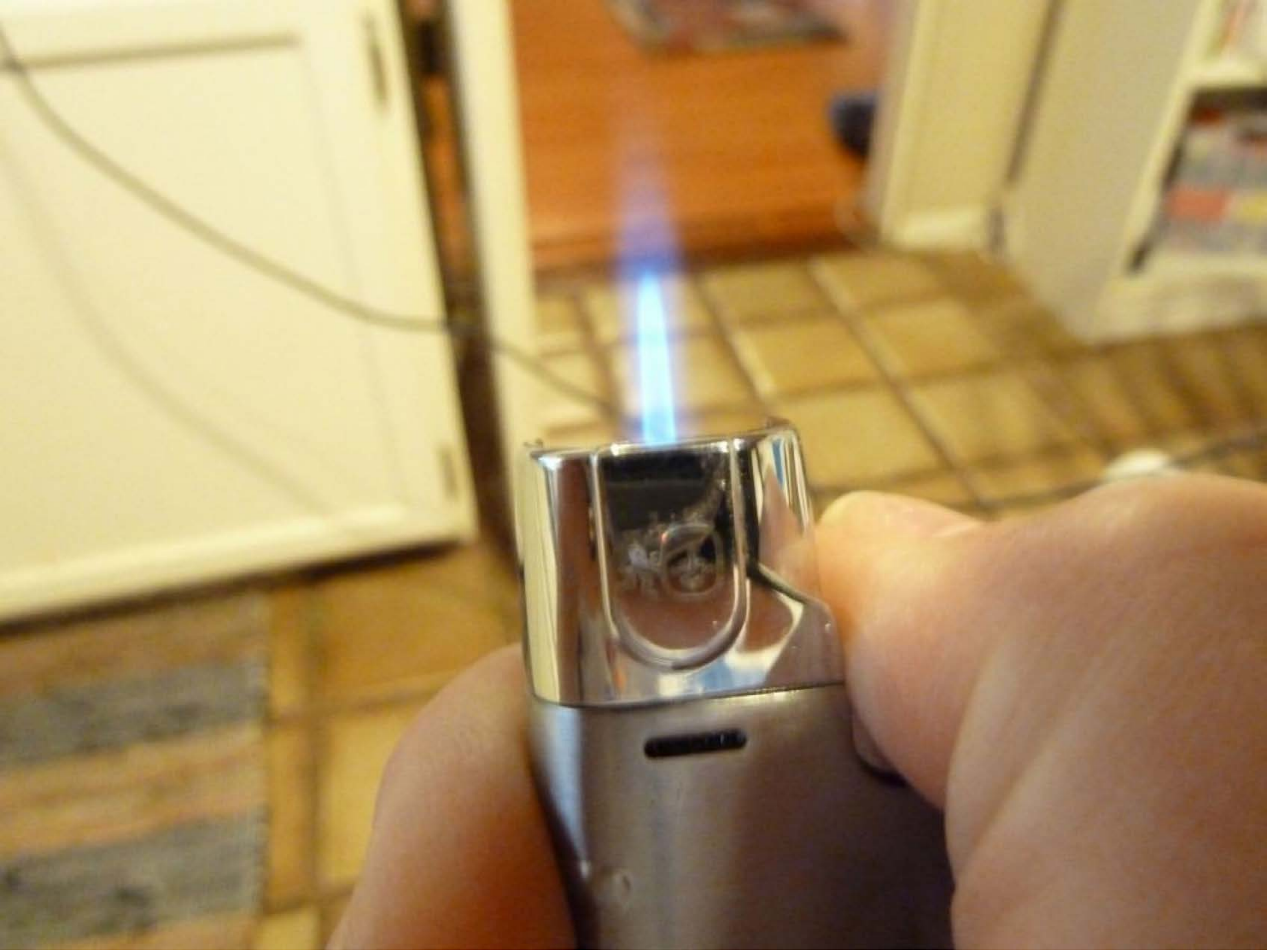

| Category | Craving (0-100) | Valence (1-9) | Arousal (1-9) | Typicality (0-100) | Relatedness | | | | | HSV | | |
|---|---|---|---|---|---|---|---|---|---|---|---|---|
| | | | | | Meth | Opioid | Both | Neither | MethToOpioid | Hue | Saturation | Value |
| Meth Instrument Hand | 65.39 (30.01) | 4.93 (1.538) | 3.71 (2.32) | 77.46 (21.45) | 0.357 | 0.036 | 0.464 | 0.143 | -0.321 | 0.117 (0.103) | 0.462 (0.192) | 0.68 (0.207) |

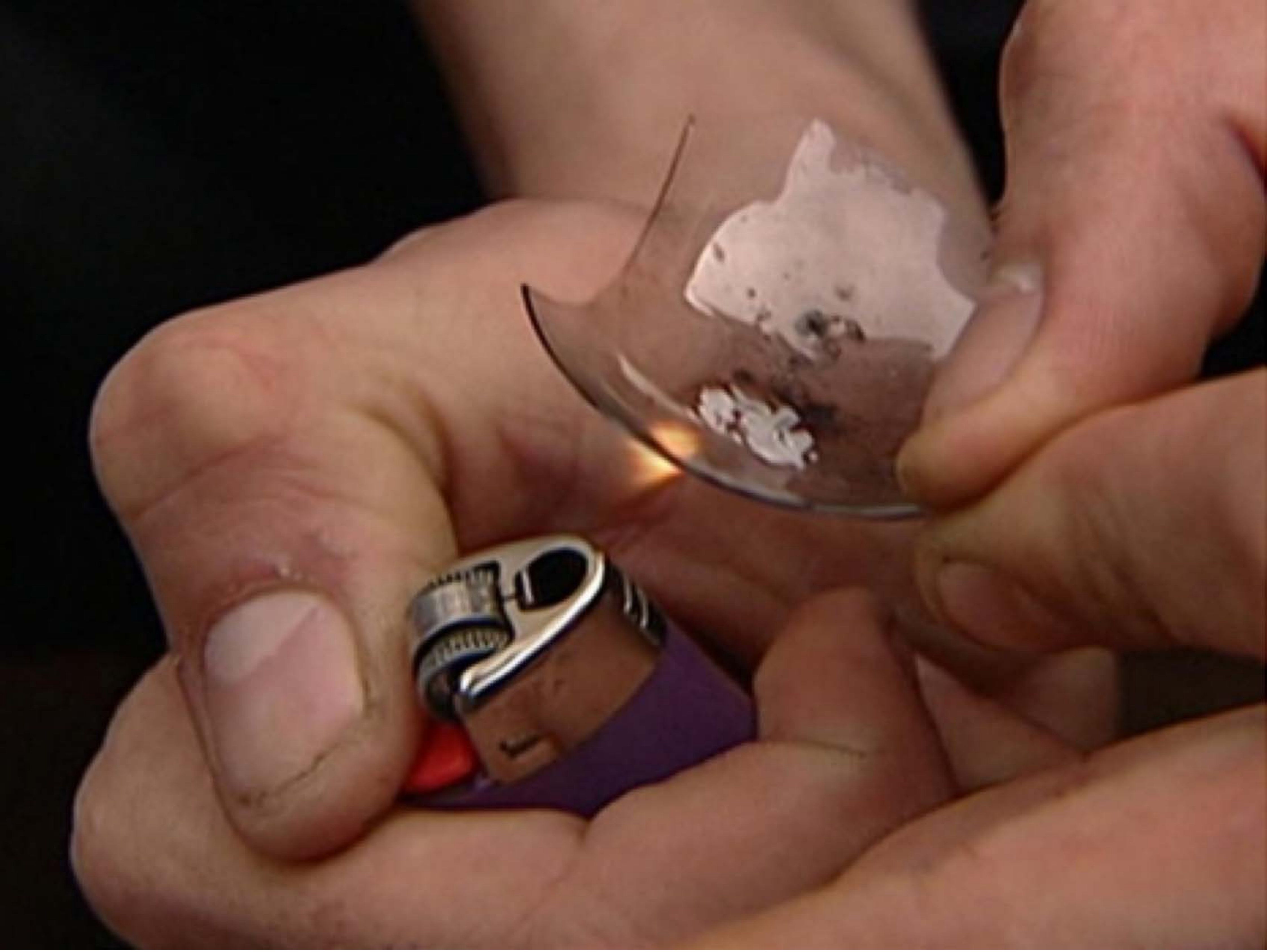

| Category | Craving (0-100) | Valence (1-9) | Arousal (1-9) | Typicality (0-100) | Relatedness | | | | | HSV | | |
|---|---|---|---|---|---|---|---|---|---|---|---|---|
| | | | | | Meth | Opioid | Both | Neither | MethToOpioid | Hue | Saturation | Value |
| Meth Instrument Hand | 90.57 (13.87) | 4.54 (1.753) | 4.5 (2.83) | 89.39 (14.18) | 0.893 | 0.036 | 0.071 | 0 | -0.857 | 0.089 (0.175) | 0.446 (0.199) | 0.417 (0.232) |

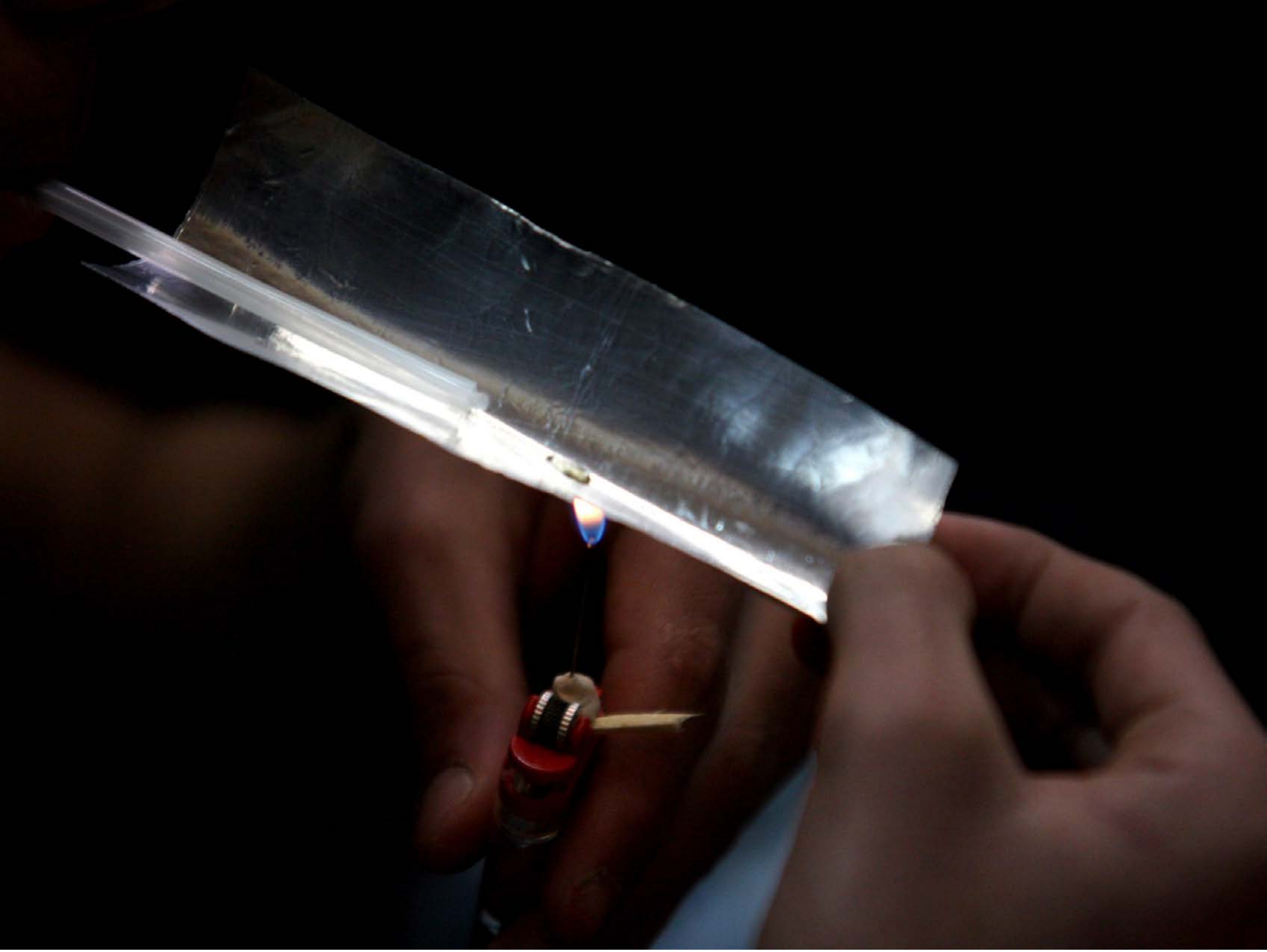

| Category | Craving (0-100) | Valence (1-9) | Arousal (1-9) | Typicality (0-100) | Relatedness | | | | | HSV | | |
|---|---|---|---|---|---|---|---|---|---|---|---|---|
| | | | | | Meth | Opioid | Both | Neither | MethToOpioid | Hue | Saturation | Value |
| Meth Instrument Hand | 85.89 (18.59) | 4.59 (2.005) | 4.96 (2.68) | 86.85 (17.12) | 0.444 | 0.148 | 0.407 | 0 | -0.296 | 0.145 (0.276) | 0.289 (0.355) | 0.129 (0.199) |

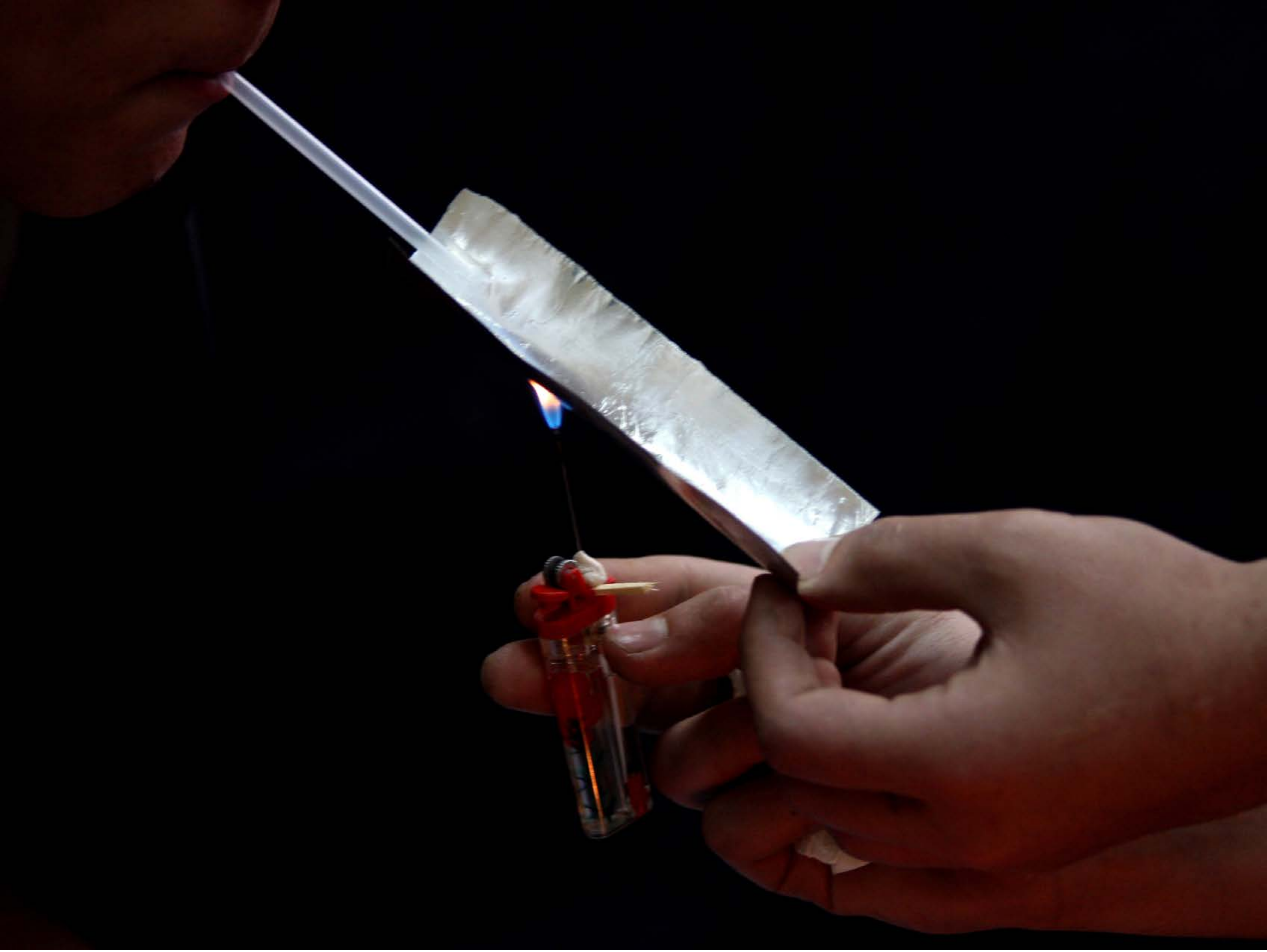

| Category | Craving (0-100) | Valence (1-9) | Arousal (1-9) | Typicality (0-100) | Relatedness | | | | | HSV | | |
|---|---|---|---|---|---|---|---|---|---|---|---|---|
| | | | | | Meth | Opioid | Both | Neither | MethToOpioid | Hue | Saturation | Value |
| Meth Instrument Hand | 82.56 (19.91) | 4.78 (1.783) | 4.41 (2.65) | 82.89 (19.84) | 0.407 | 0.185 | 0.407 | 0 | -0.222 | 0.219 (0.314) | 0.432 (0.446) | 0.112 (0.206) |

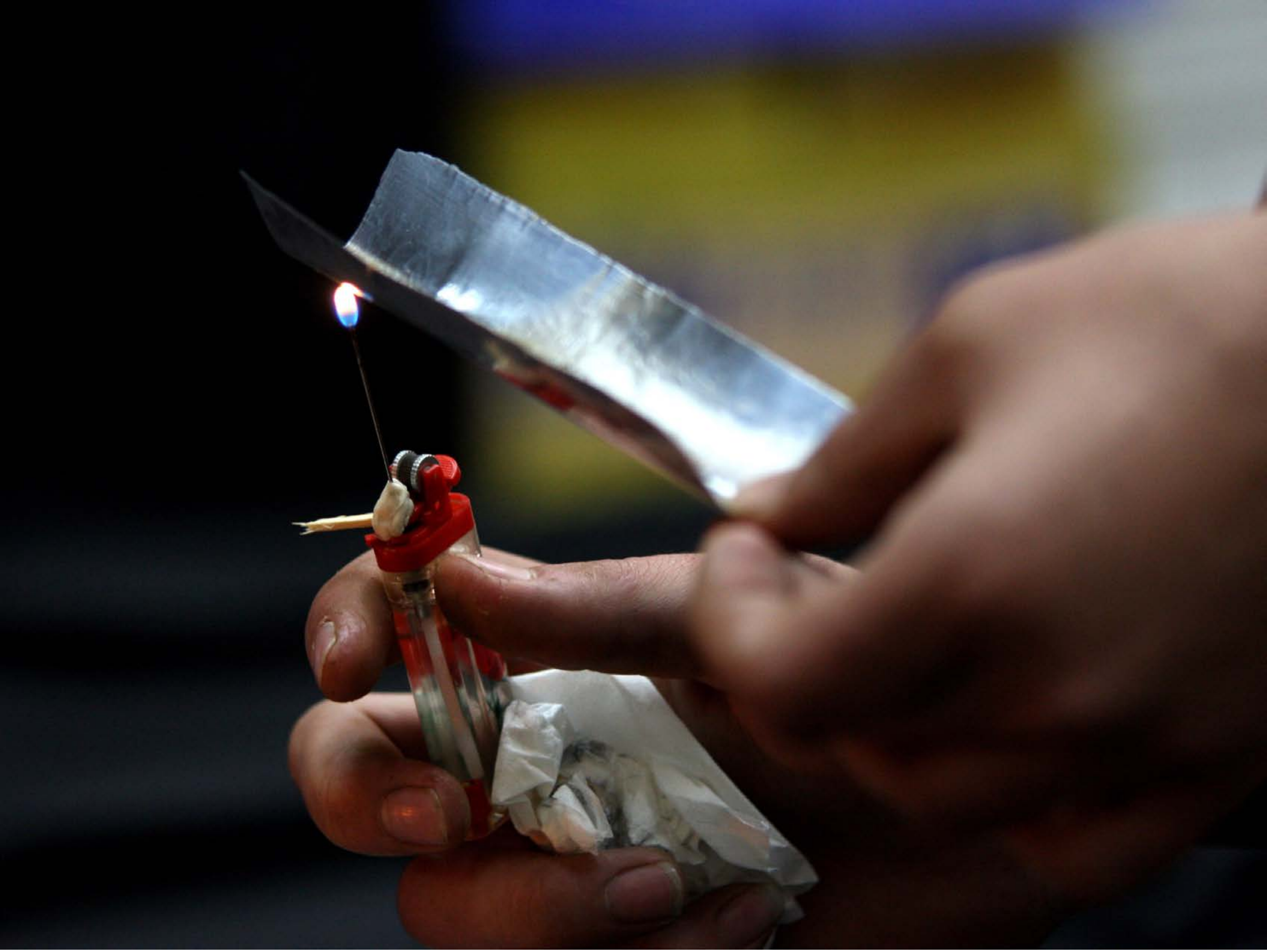

| Category | Craving (0-100) | Valence (1-9) | Arousal (1-9) | Typicality (0-100) | Relatedness | | | | | HSV | | |
|---|---|---|---|---|---|---|---|---|---|---|---|---|
| | | | | | Meth | Opioid | Both | Neither | MethToOpioid | Hue | Saturation | Value |
| Meth Instrument Hand | 88.21 (20.02) | 4.46 (1.915) | 4.64 (2.84) | 91.68 (11.22) | 0.357 | 0.143 | 0.464 | 0.036 | -0.214 | 0.237 (0.279) | 0.442 (0.316) | 0.262 (0.244) |

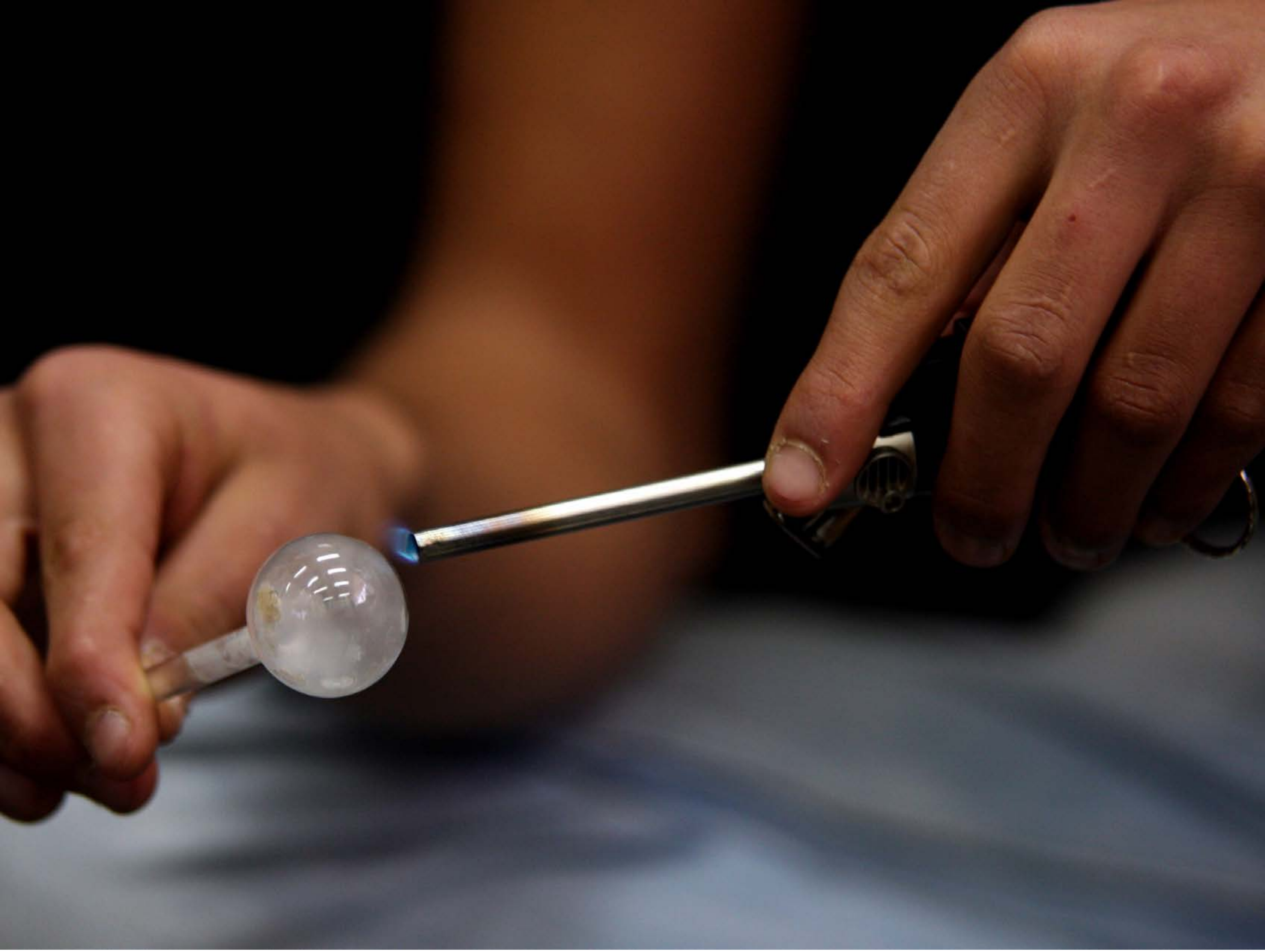

| Category | Craving (0-100) | Valence (1-9) | Arousal (1-9) | Typicality (0-100) | Relatedness | | | | | HSV | | |
|---|---|---|---|---|---|---|---|---|---|---|---|---|
| | | | | | Meth | Opioid | Both | Neither | MethToOpioid | Hue | Saturation | Value |
| Meth Instrument Hand | 90.18 (19.59) | 4.54 (1.895) | 4.68 (2.74) | 91.68 (19.44) | 0.929 | 0 | 0.071 | 0 | -0.929 | 0.221 (0.287) | 0.397 (0.335) | 0.296 (0.213) |

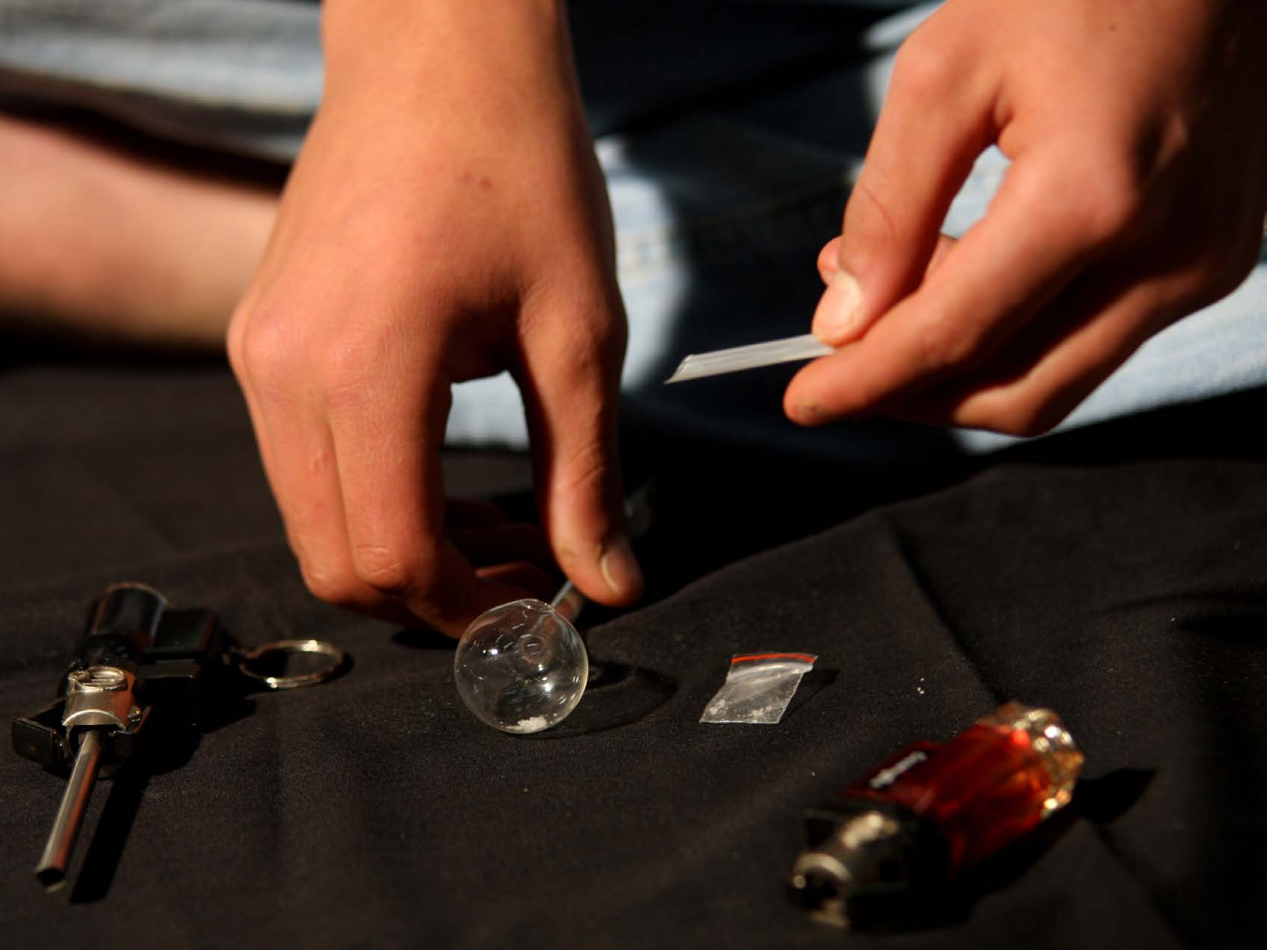

| Category | Craving (0-100) | Valence (1-9) | Arousal (1-9) | Typicality (0-100) | Relatedness | | | | | HSV | | |
|---|---|---|---|---|---|---|---|---|---|---|---|---|
| | | | | | Meth | Opioid | Both | Neither | MethToOpioid | Hue | Saturation | Value |
| Meth Instrument Hand | 86.15 (17.61) | 4.26 (1.767) | 4.44 (2.68) | 86.59 (18.16) | 0.852 | 0 | 0.148 | 0 | -0.852 | 0.129 (0.201) | 0.469 (0.265) | 0.312 (0.319) |

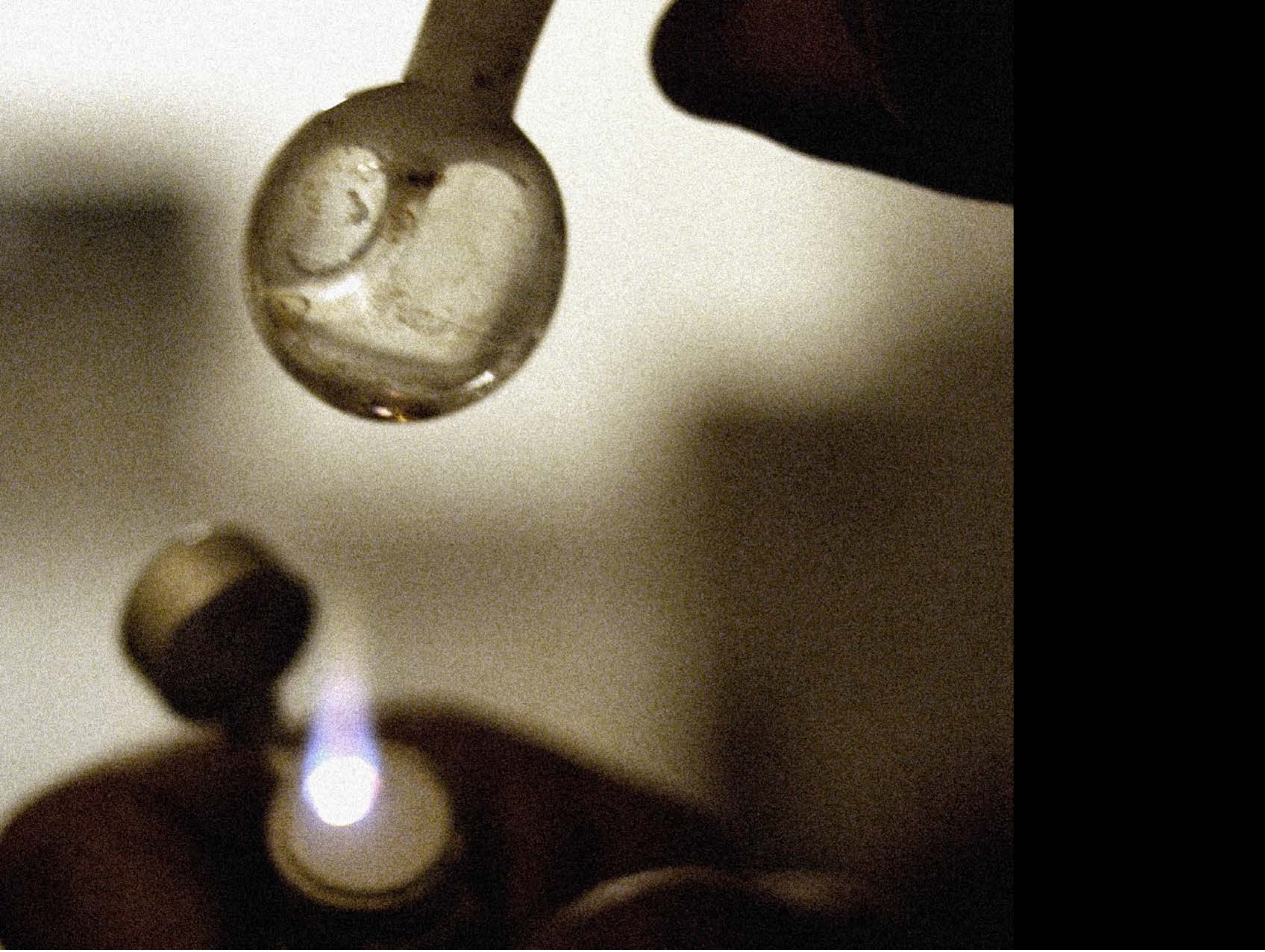

| Category | Craving (0-100) | Valence (1-9) | Arousal (1-9) | Typicality (0-100) | Relatedness | | | | | HSV | | |
|---|---|---|---|---|---|---|---|---|---|---|---|---|
| | | | | | Meth | Opioid | Both | Neither | MethToOpioid | Hue | Saturation | Value |
| Meth Instrument Hand | 87.41 (22.09) | 4.37 (1.964) | 4.81 (2.76) | 90.63 (16.54) | 0.889 | 0 | 0.111 | 0 | -0.889 | 0.124 (0.209) | 0.278 (0.267) | 0.359 (0.325) |

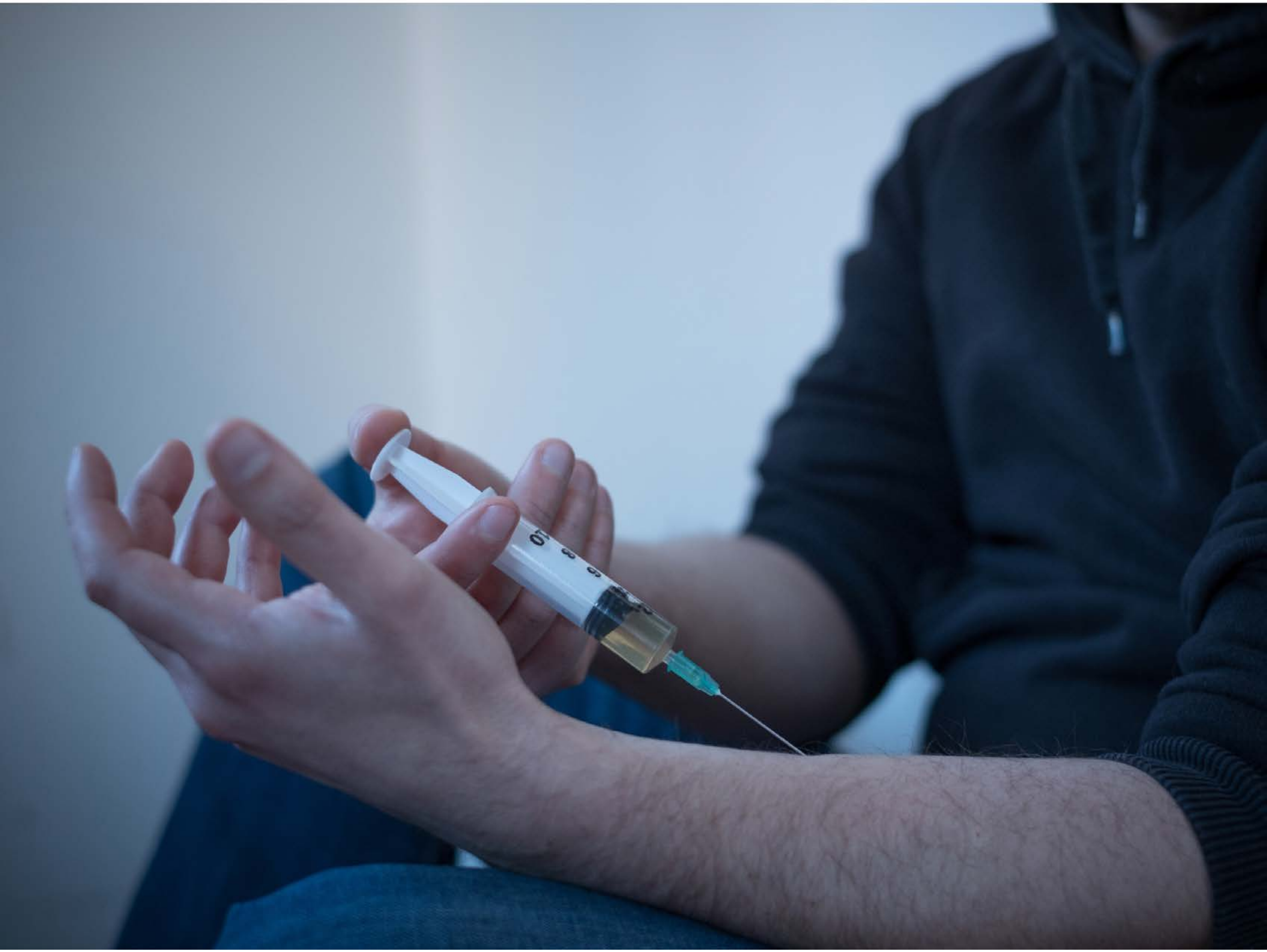

| Category | Craving (0-100) | Valence (1-9) | Arousal (1-9) | Typicality (0-100) | Relatedness | | | | | HSV | | |
|---|---|---|---|---|---|---|---|---|---|---|---|---|
| | | | | | Meth | Opioid | Both | Neither | MethToOpioid | Hue | Saturation | Value |
| Meth Injection Hand | 84.56 (20.69) | 4.44 (1.761) | 4.78 (2.62) | 86.56 (20.3) | 0.148 | 0.037 | 0.815 | 0 | -0.111 | 0.622 (0.077) | 0.384 (0.267) | 0.468 (0.269) |

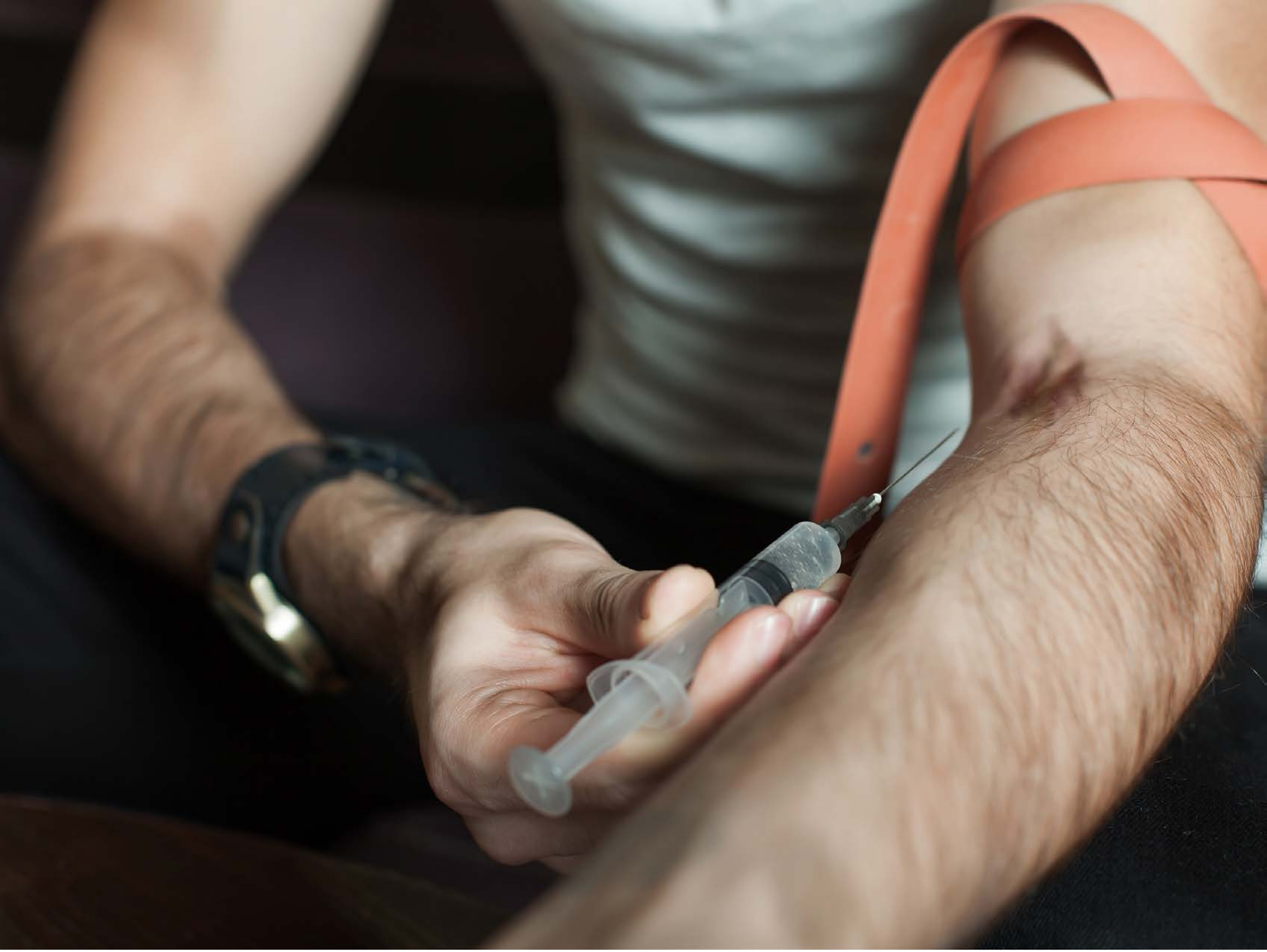

| Category | Craving (0-100) | Valence (1-9) | Arousal (1-9) | Typicality (0-100) | Relatedness | | | | | HSV | | |
|---|---|---|---|---|---|---|---|---|---|---|---|---|
| | | | | | Meth | Opioid | Both | Neither | MethToOpioid | Hue | Saturation | Value |
| Meth Injection Hand | 84.74 (23.23) | 3.96 (1.85) | 4.59 (2.74) | 85.41 (23.76) | 0.185 | 0 | 0.815 | 0 | -0.185 | 0.192 (0.258) | 0.282 (0.161) | 0.417 (0.298) |

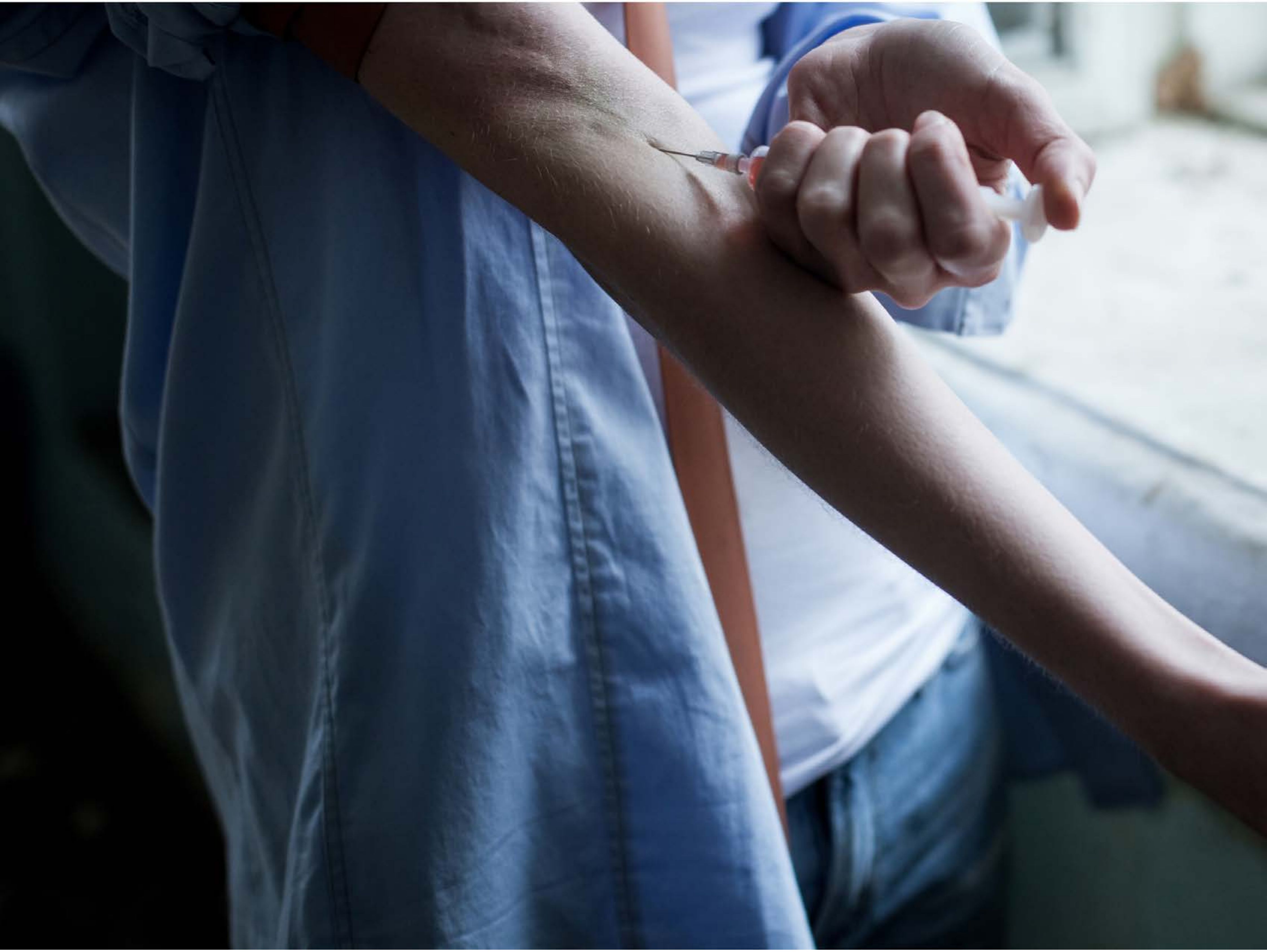

| Category | Craving (0-100) | Valence (1-9) | Arousal (1-9) | Typicality (0-100) | Relatedness | | | | | HSV | | |
|---|---|---|---|---|---|---|---|---|---|---|---|---|
| | | | | | Meth | Opioid | Both | Neither | MethToOpioid | Hue | Saturation | Value |
| Meth Injection Hand | 91.18 (14.09) | 4.14 (2.206) | 5.07 (2.89) | 92.79 (13.67) | 0.107 | 0 | 0.893 | 0 | -0.107 | 0.484 (0.253) | 0.357 (0.226) | 0.402 (0.301) |

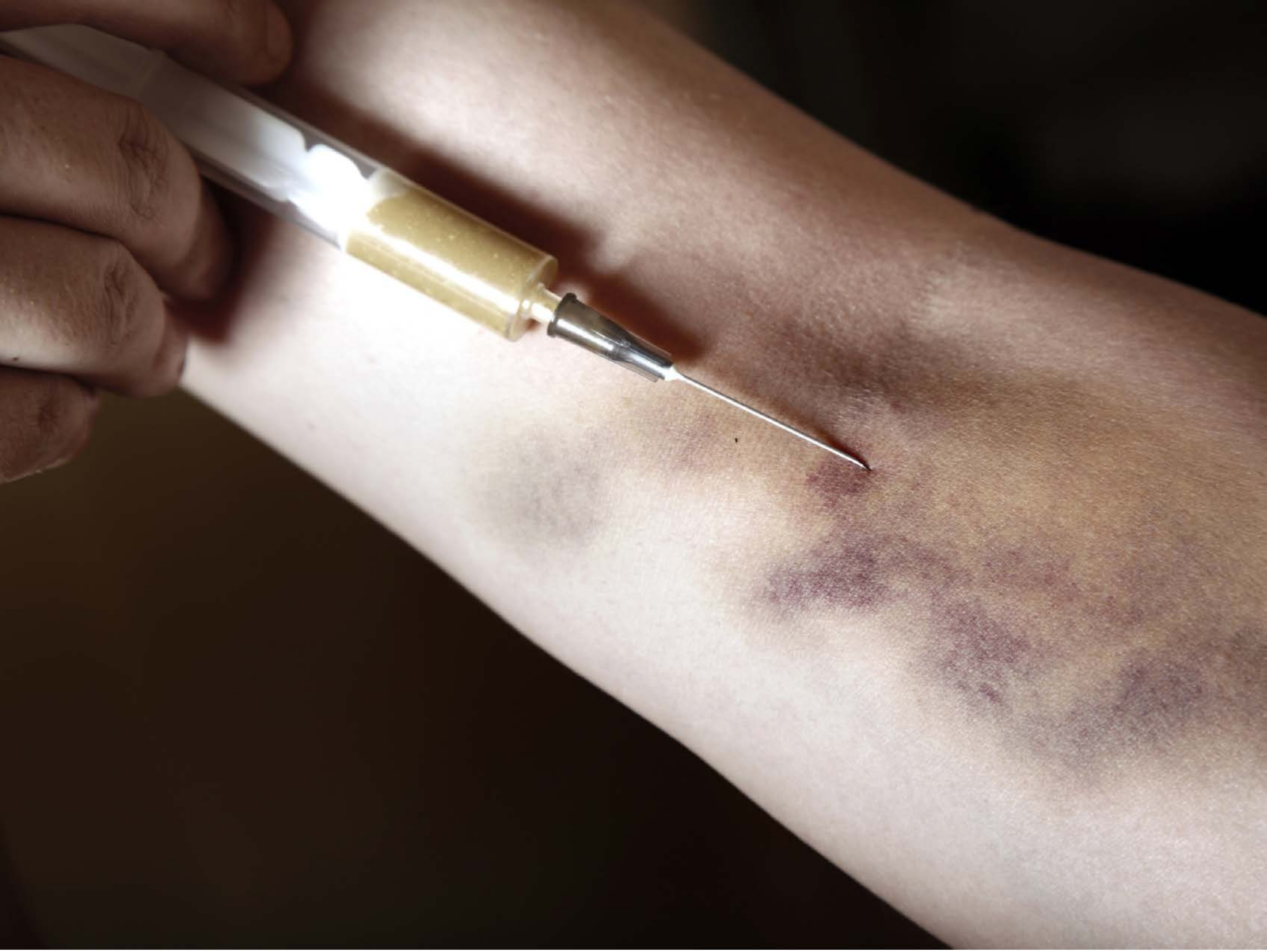

| Category | Craving (0-100) | Valence (1-9) | Arousal (1-9) | Typicality (0-100) | Relatedness | | | | | HSV | | |
|---|---|---|---|---|---|---|---|---|---|---|---|---|
| | | | | | Meth | Opioid | Both | Neither | MethToOpioid | Hue | Saturation | Value |
| Meth Injection Hand | 84.89 (18.96) | 3.85 (1.812) | 4.85 (2.66) | 86.52 (18.45) | 0.111 | 0.111 | 0.778 | 0 | 0 | 0.08 (0.141) | 0.355 (0.222) | 0.456 (0.315) |

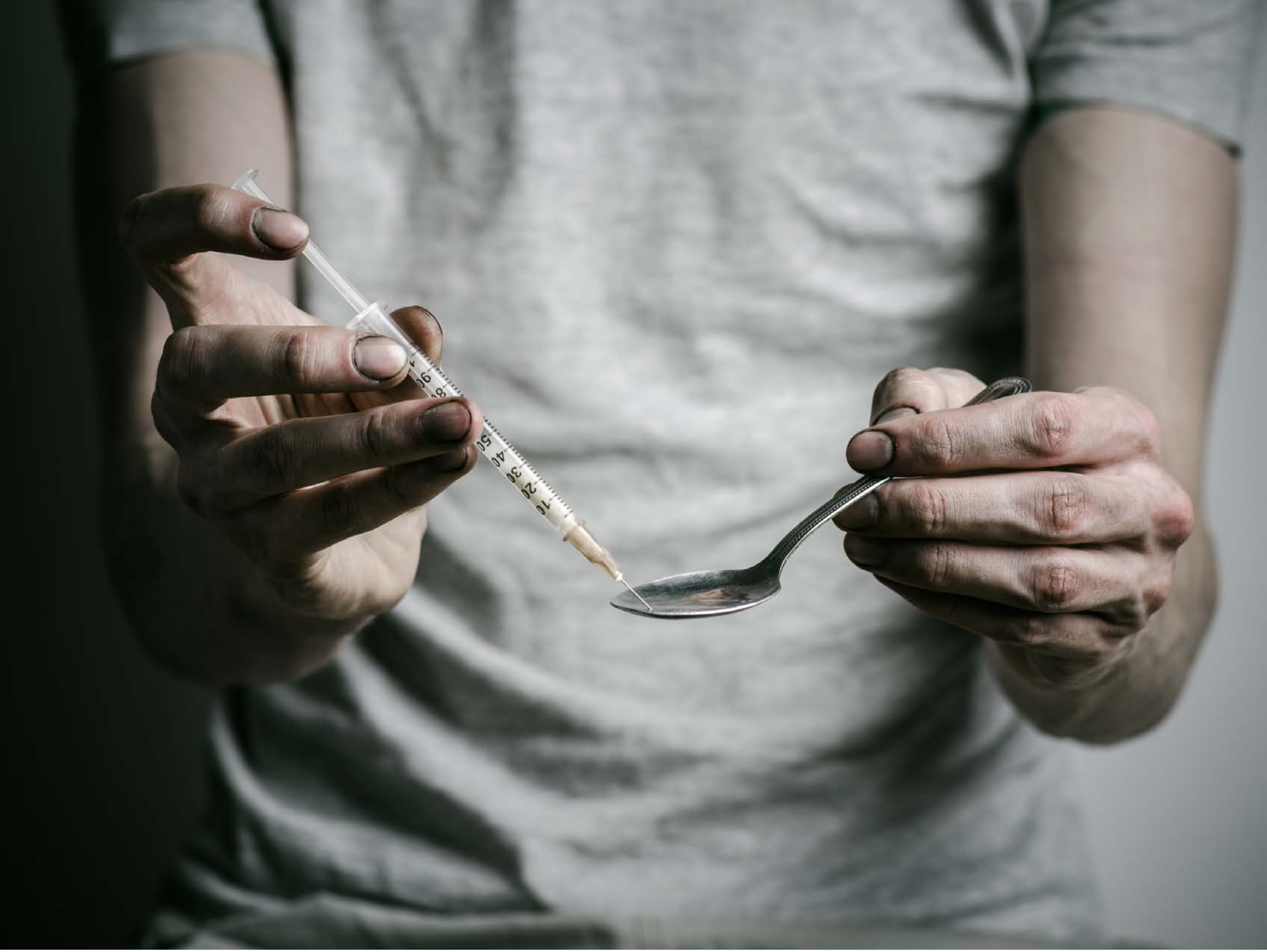

| Category | Craving (0-100) | Valence (1-9) | Arousal (1-9) | Typicality (0-100) | Relatedness: Meth | Opioid | Both | Neither | MethToOpioid | HSV: Hue | Saturation | Value |
|---|---|---|---|---|---|---|---|---|---|---|---|---|
| Meth Injection Hand | 90.75 (15.72) | 4.14 (1.82) | 4.79 (2.57) | 92.75 (13.59) | 0.107 | 0.071 | 0.821 | 0 | -0.036 | 0.344 (0.168) | 0.149 (0.162) | 0.467 (0.26) |

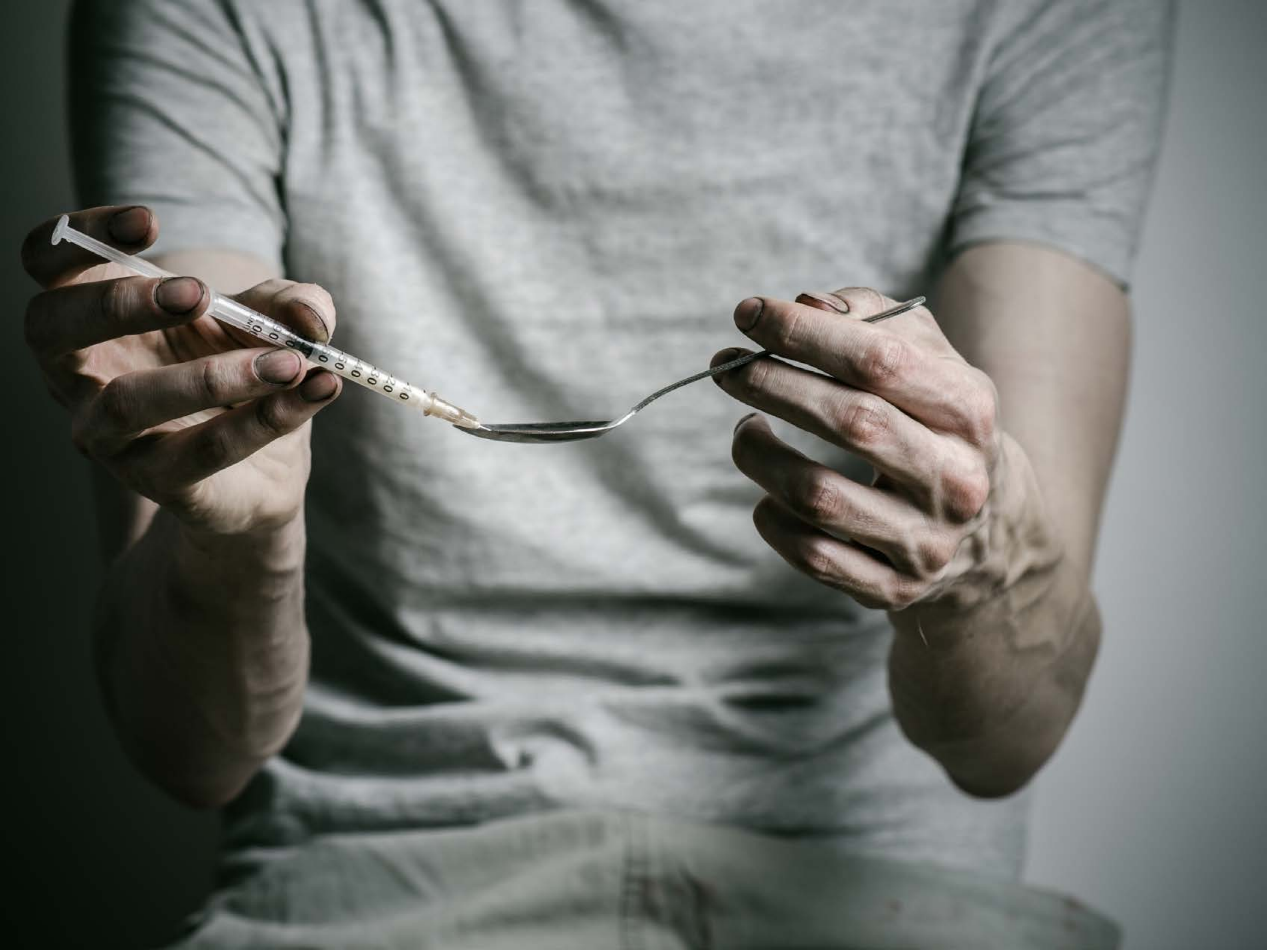

| Category | Craving (0-100) | Valence (1-9) | Arousal (1-9) | Typicality (0-100) | Relatedness | | | | | HSV | | |
|---|---|---|---|---|---|---|---|---|---|---|---|---|
| | | | | | Meth | Opioid | Both | Neither | MethToOpioid | Hue | Saturation | Value |
| Meth Injection Hand | 92.21 (12.63) | 4.36 (2.022) | 4.89 (2.73) | 94.25 (11.25) | 0.107 | 0 | 0.893 | 0 | -0.107 | 0.372 (0.163) | 0.126 (0.132) | 0.472 (0.23) |

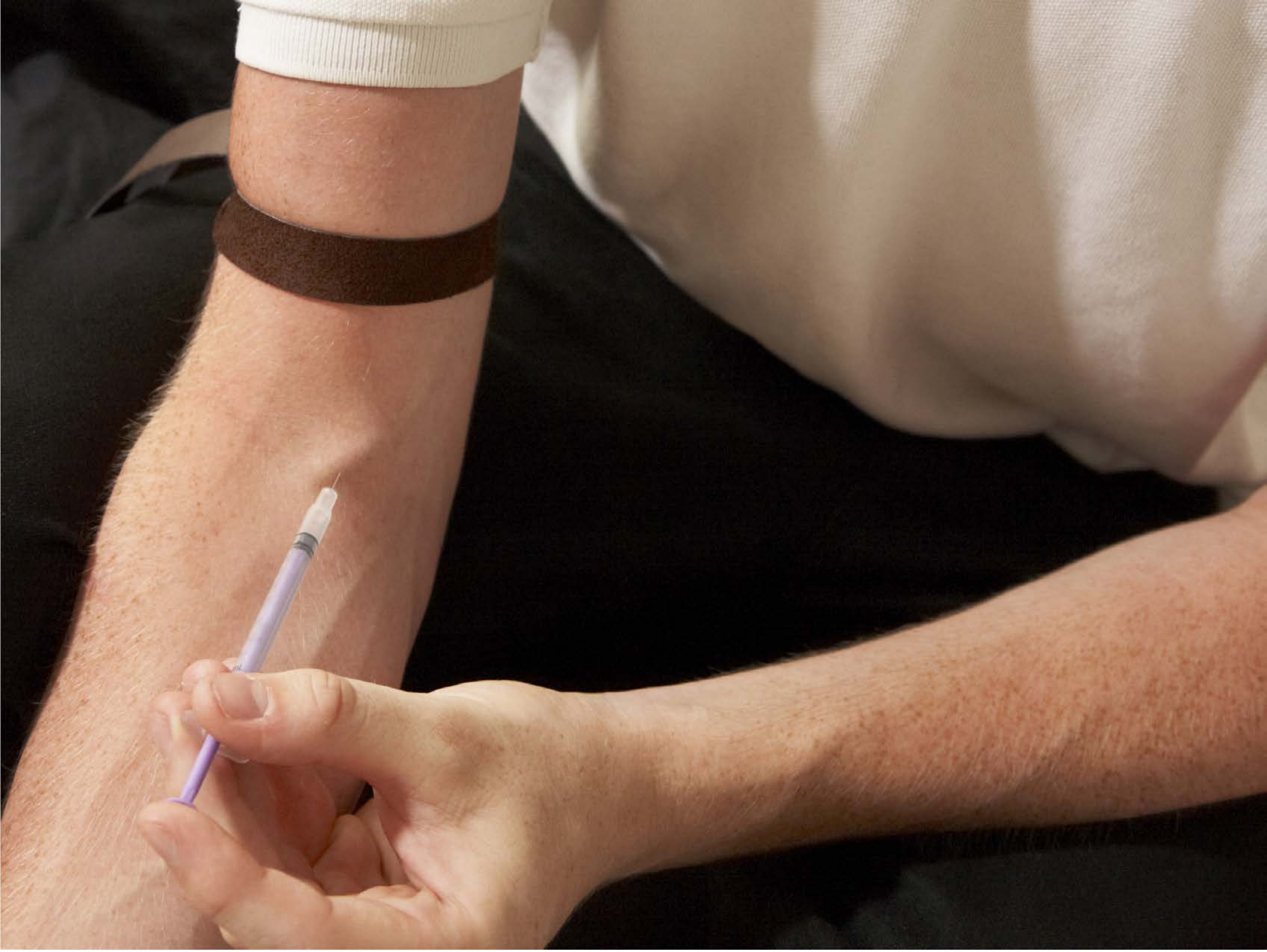

| Category | Craving (0-100) | Valence (1-9) | Arousal (1-9) | Typicality (0-100) | Relatedness | | | | | HSV | | |
|---|---|---|---|---|---|---|---|---|---|---|---|---|
| | | | | | Meth | Opioid | Both | Neither | MethToOpioid | Hue | Saturation | Value |
| Meth Injection Hand | 88.19 (17.34) | 4.19 (1.861) | 4.7 (2.66) | 89.19 (17.8) | 0.111 | 0 | 0.889 | 0 | -0.111 | 0.157 (0.266) | 0.384 (0.191) | 0.459 (0.311) |

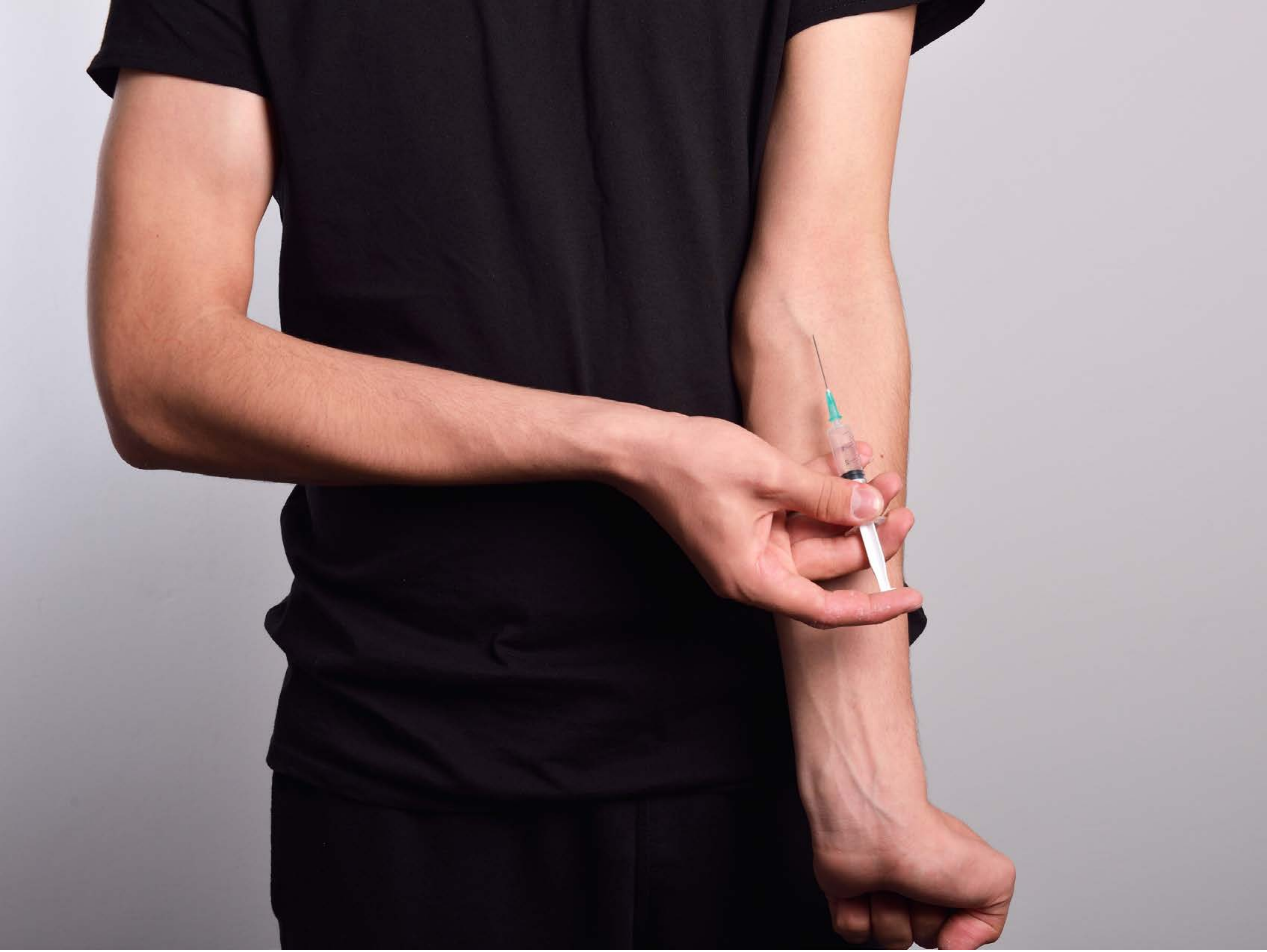

| Category | Craving (0-100) | Valence (1-9) | Arousal (1-9) | Typicality (0-100) | Relatedness | | | | | HSV | | |
|---|---|---|---|---|---|---|---|---|---|---|---|---|
| | | | | | Meth | Opioid | Both | Neither | MethToOpioid | Hue | Saturation | Value |
| Meth Injection Hand | 91.68 (13.64) | 4.21 (2.331) | 4.86 (3.11) | 93.18 (13.16) | 0.143 | 0 | 0.857 | 0 | -0.143 | 0.659 (0.334) | 0.232 (0.19) | 0.505 (0.344) |

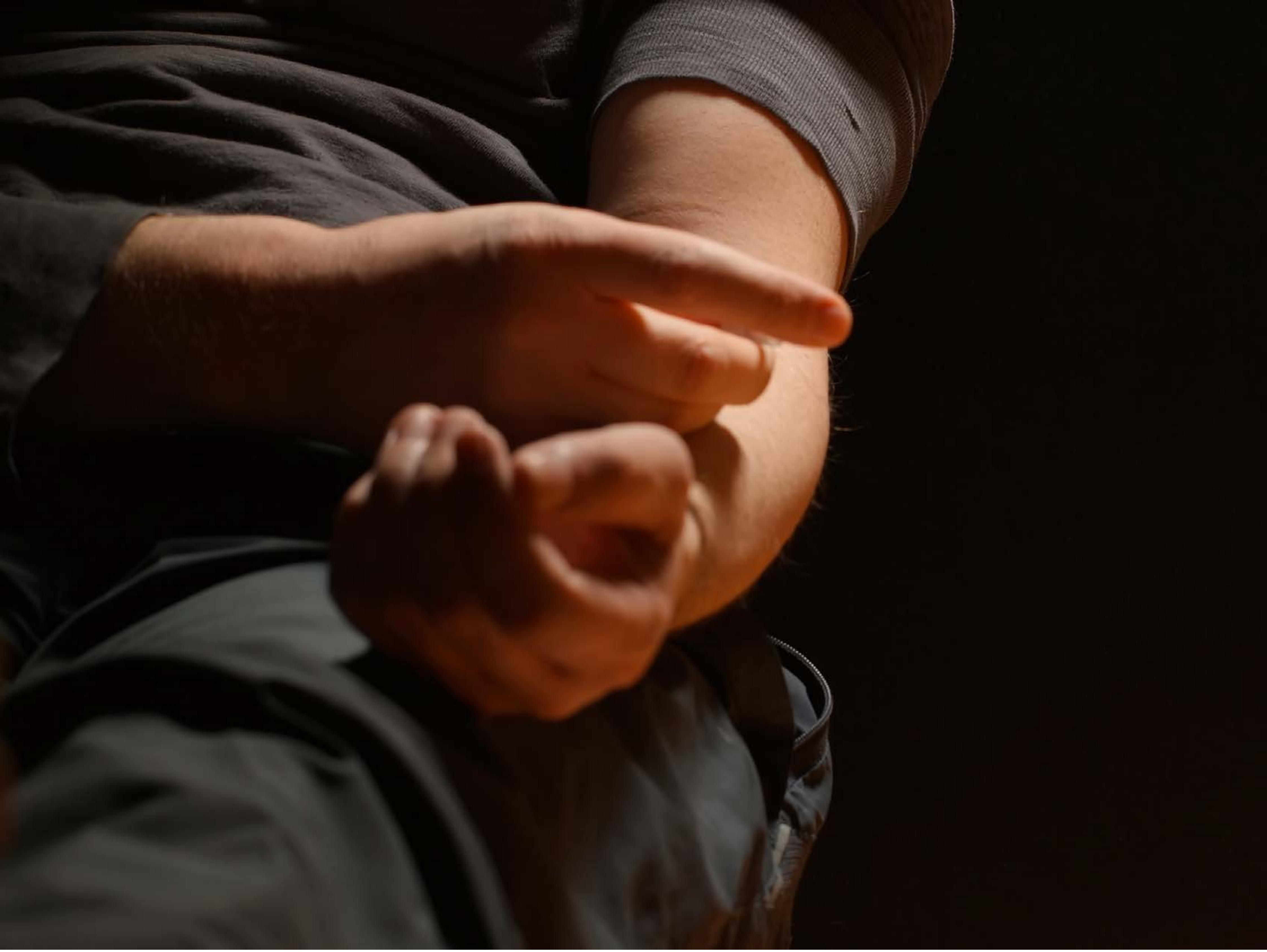

| Category | Craving (0-100) | Valence (1-9) | Arousal (1-9) | Typicality (0-100) | Relatedness | | | | | HSV | | |
|---|---|---|---|---|---|---|---|---|---|---|---|---|
| | | | | | Meth | Opioid | Both | Neither | MethToOpioid | Hue | Saturation | Value |
| Meth Injection Hand | 84.86 (22.82) | 4.11 (2.006) | 4.64 (2.64) | 85.75 (23.28) | 0.148 | 0 | 0.815 | 0.037 | -0.148 | 0.113 (0.18) | 0.569 (0.261) | 0.168 (0.195) |

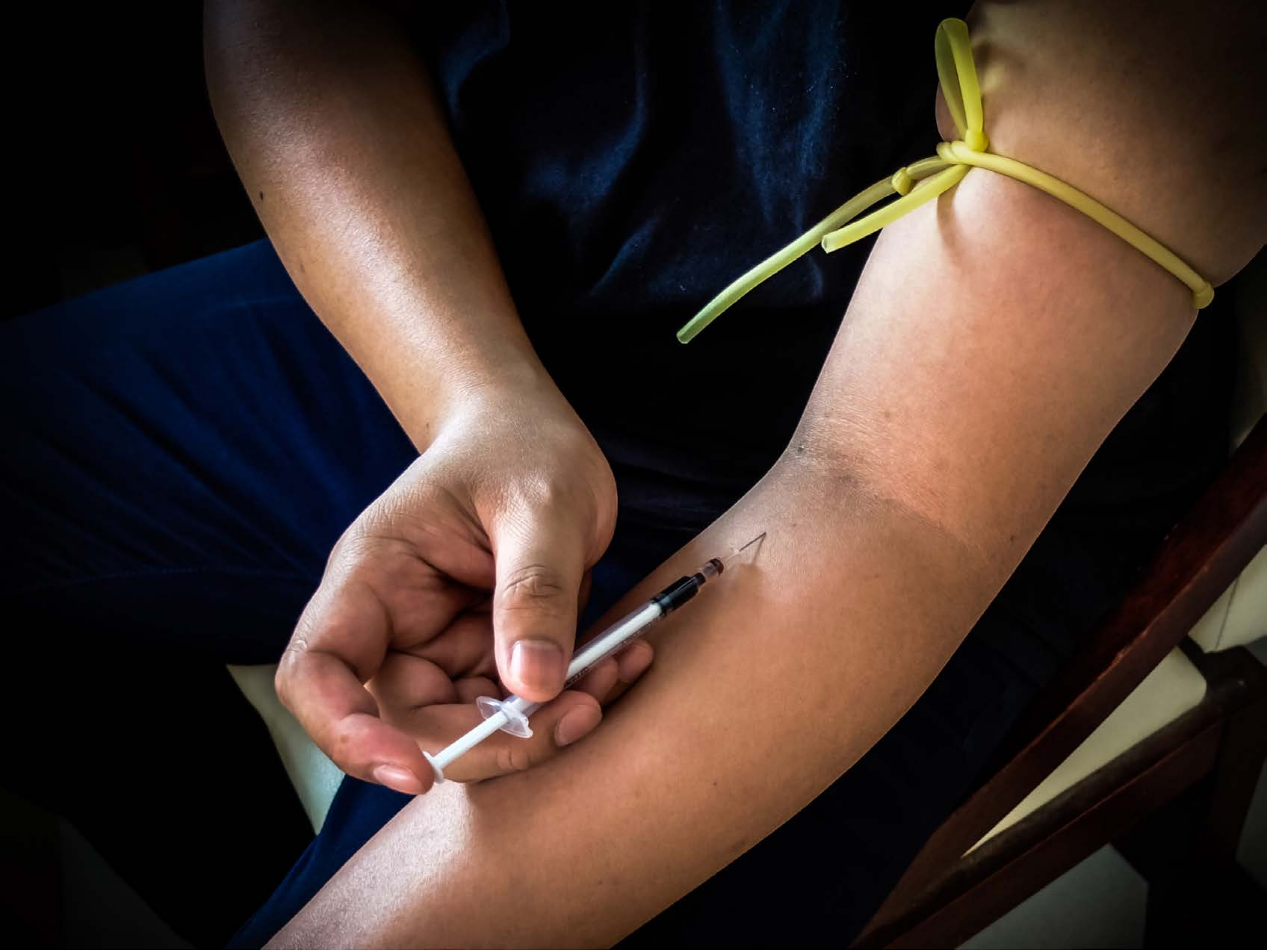

| Category | Craving (0-100) | Valence (1-9) | Arousal (1-9) | Typicality (0-100) | Relatedness | | | | | HSV | | |
|---|---|---|---|---|---|---|---|---|---|---|---|---|
| | | | | | Meth | Opioid | Both | Neither | MethToOpioid | Hue | Saturation | Value |
| Meth Injection Hand | 93.21 (11.52) | 3.93 (2.193) | 4.96 (2.82) | 92.21 (13.63) | 0.107 | 0.143 | 0.75 | 0 | 0.036 | 0.221 (0.291) | 0.442 (0.307) | 0.28 (0.303) |

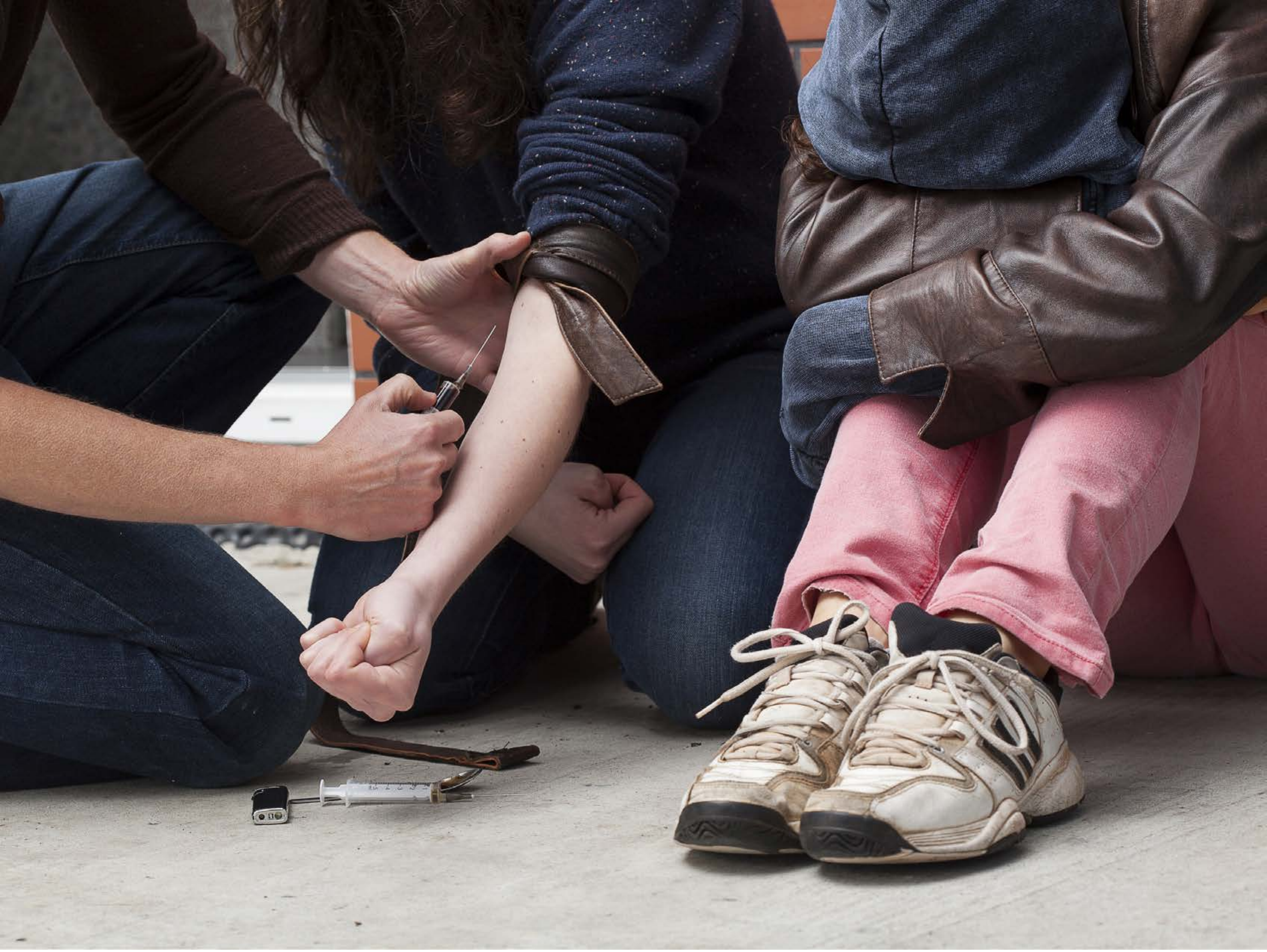

| Category | Craving (0-100) | Valence (1-9) | Arousal (1-9) | Typicality (0-100) | Relatedness | | | | | HSV | | |
|---|---|---|---|---|---|---|---|---|---|---|---|---|
| | | | | | Meth | Opioid | Both | Neither | MethToOpioid | Hue | Saturation | Value |
| Meth Injection Hand | 90.89 (14.88) | 4.3 (2.091) | 4.89 (2.75) | 88.81 (17.43) | 0.111 | 0.185 | 0.667 | 0.037 | 0.074 | 0.411 (0.367) | 0.245 (0.16) | 0.395 (0.291) |

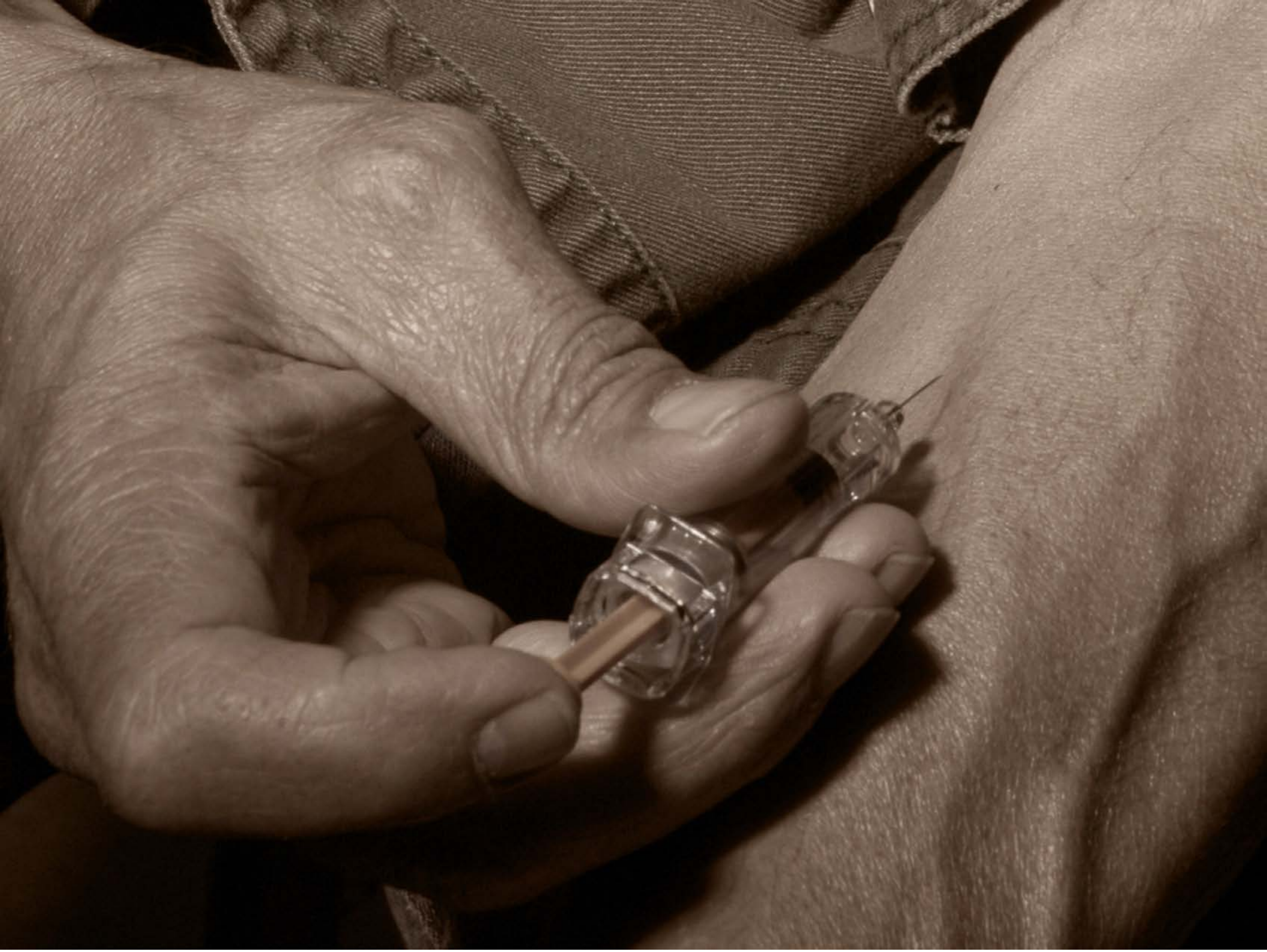

| Category | Craving (0-100) | Valence (1-9) | Arousal (1-9) | Typicality (0-100) | Relatedness | | | | | HSV | | |
|---|---|---|---|---|---|---|---|---|---|---|---|---|
| | | | | | Meth | Opioid | Both | Neither | MethToOpioid | Hue | Saturation | Value |
| Meth Injection Hand | 88.78 (15.91) | 4.33 (2.019) | 4.7 (2.96) | 90.41 (15.7) | 0.074 | 0 | 0.889 | 0.037 | -0.074 | 0.078 (0.083) | 0.422 (0.245) | 0.39 (0.203) |

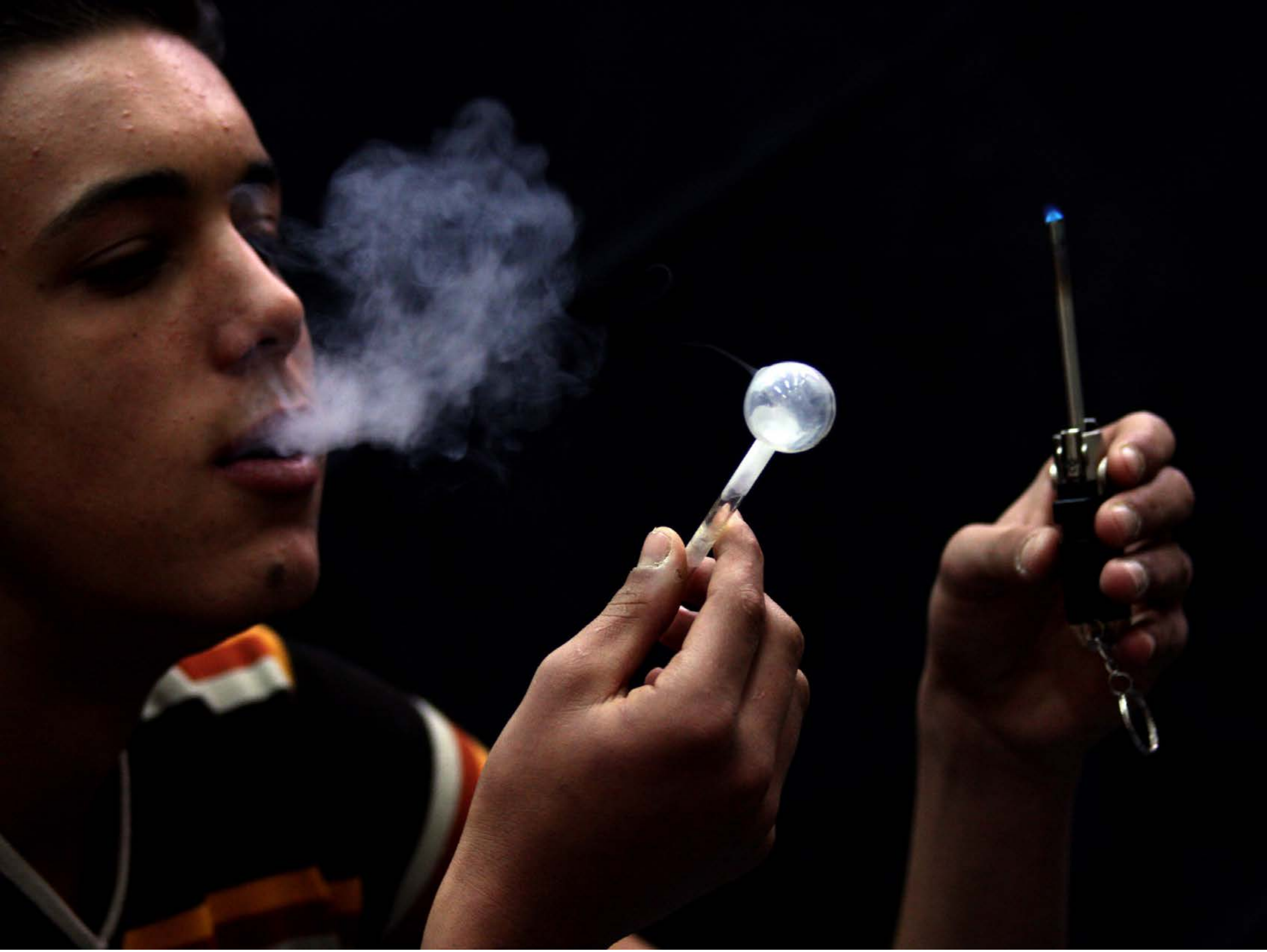

| Category | Craving (0-100) | Valence (1-9) | Arousal (1-9) | Typicality (0-100) | Relatedness | | | | | HSV | | |
|---|---|---|---|---|---|---|---|---|---|---|---|---|
| | | | | | Meth | Opioid | Both | Neither | MethToOpioid | Hue | Saturation | Value |
| Meth Face Activities | 92.5 (11.74) | 4.5 (1.953) | 4.57 (2.97) | 95.25 (9.85) | 0.821 | 0 | 0.179 | 0 | -0.821 | 0.371 (0.343) | 0.613 (0.354) | 0.128 (0.199) |

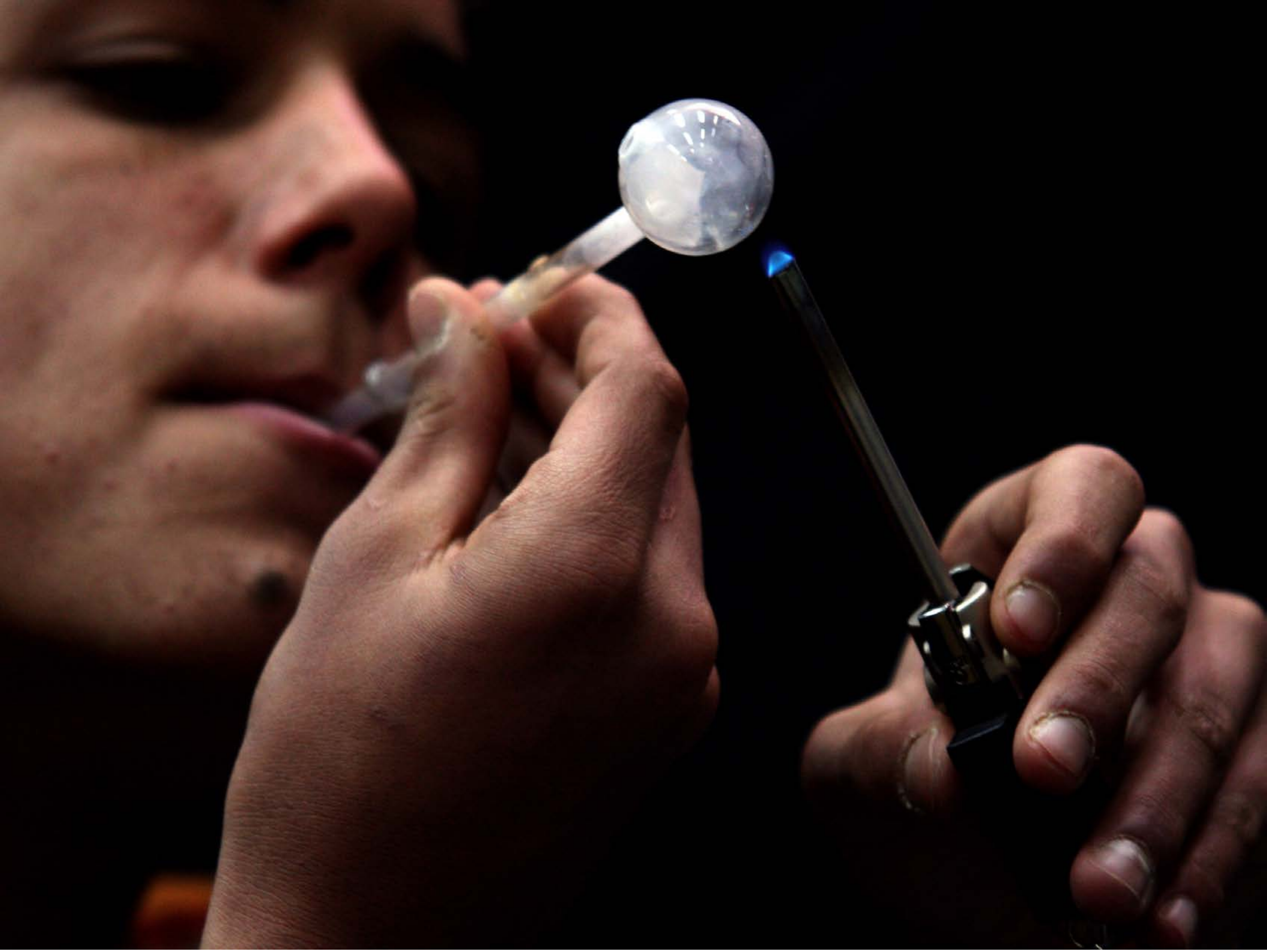

| Category | Craving (0-100) | Valence (1-9) | Arousal (1-9) | Typicality (0-100) | Relatedness | | | | | HSV | | |
|---|---|---|---|---|---|---|---|---|---|---|---|---|
| | | | | | Meth | Opioid | Both | Neither | MethToOpioid | Hue | Saturation | Value |
| Meth Face Activities | 87.81 (15.82) | 4.38 (2.002) | 4.67 (2.67) | 90.78 (14.17) | 0.926 | 0 | 0.074 | 0 | -0.926 | 0.166 (0.311) | 0.468 (0.369) | 0.197 (0.249) |

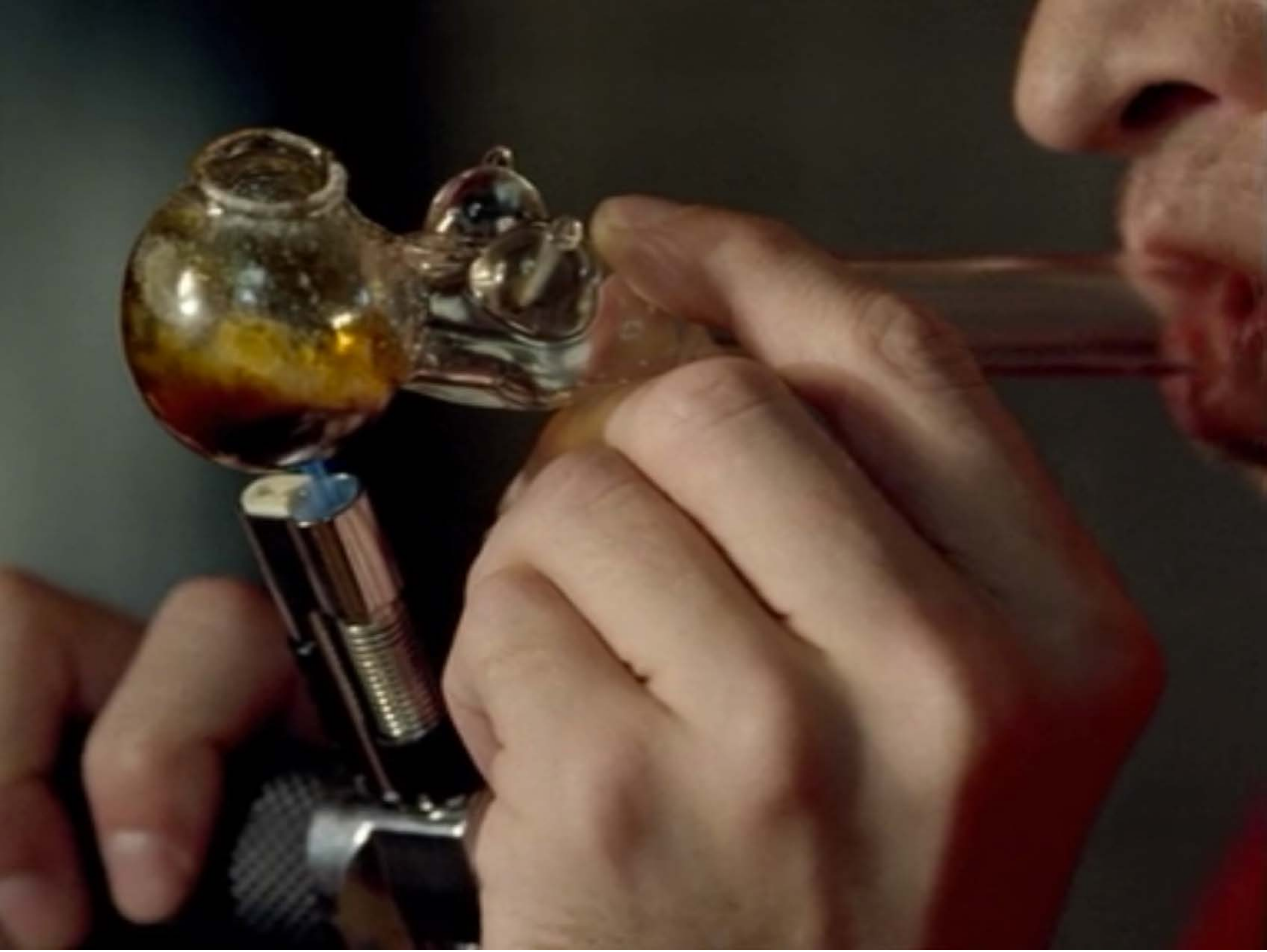

| Category | Craving (0-100) | Valence (1-9) | Arousal (1-9) | Typicality (0-100) | Relatedness | | | | | HSV | | |
|---|---|---|---|---|---|---|---|---|---|---|---|---|
| | | | | | Meth | Opioid | Both | Neither | MethToOpioid | Hue | Saturation | Value |
| Meth Face Activities | 91.14 (15.65) | 4.75 (1.993) | 4.89 (2.79) | 92.61 (13.25) | 0.75 | 0 | 0.25 | 0 | -0.75 | 0.165 (0.243) | 0.434 (0.245) | 0.304 (0.199) |

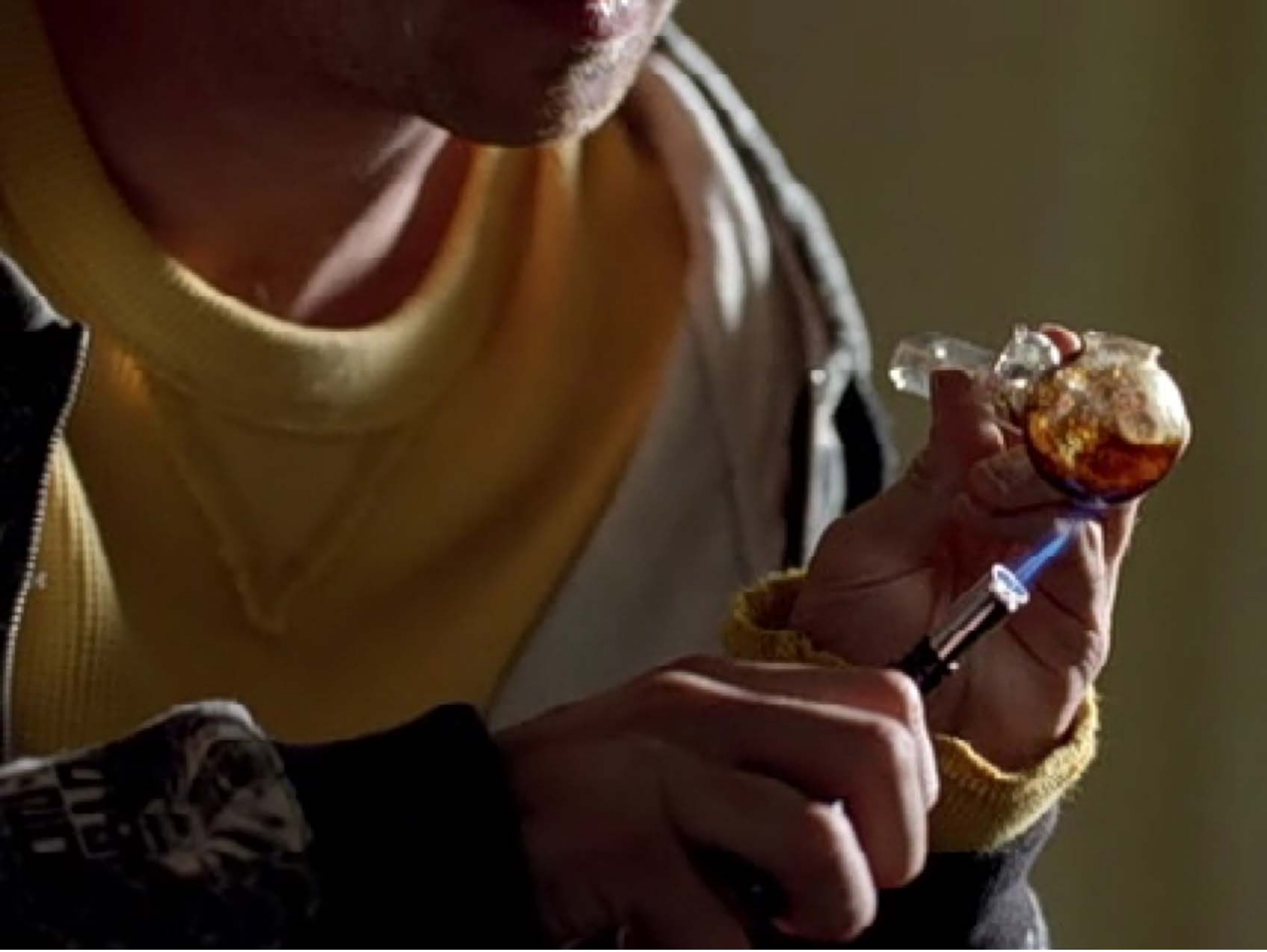

| Category | Craving (0-100) | Valence (1-9) | Arousal (1-9) | Typicality (0-100) | Relatedness | | | | | HSV | | |
|---|---|---|---|---|---|---|---|---|---|---|---|---|
| | | | | | Meth | Opioid | Both | Neither | MethToOpioid | Hue | Saturation | Value |
| Meth Face Activities | 89.85 (15.2) | 4.3 (2.091) | 4.74 (2.85) | 90.78 (14.36) | 0.926 | 0 | 0.074 | 0 | -0.926 | 0.155 (0.243) | 0.568 (0.264) | 0.269 (0.16) |

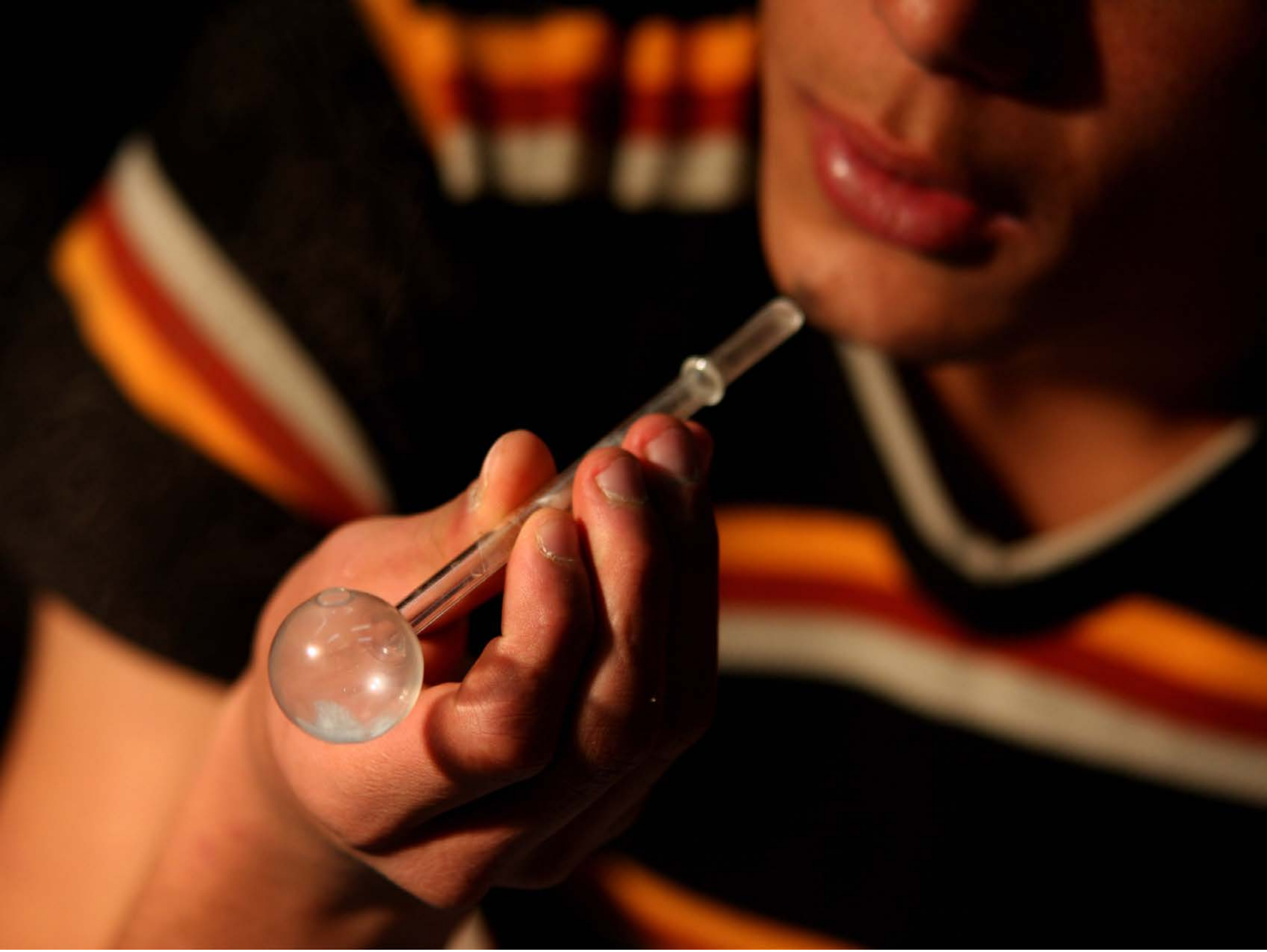

| Category | Craving (0-100) | Valence (1-9) | Arousal (1-9) | Typicality (0-100) | Relatedness | | | | | HSV | | |
|---|---|---|---|---|---|---|---|---|---|---|---|---|
| | | | | | Meth | Opioid | Both | Neither | MethToOpioid | Hue | Saturation | Value |
| Meth Face Activities | 89.18 (21.17) | 4.43 (1.952) | 4.89 (2.73) | 94.29 (10.58) | 0.929 | 0 | 0.071 | 0 | -0.929 | 0.132 (0.274) | 0.662 (0.338) | 0.301 (0.313) |

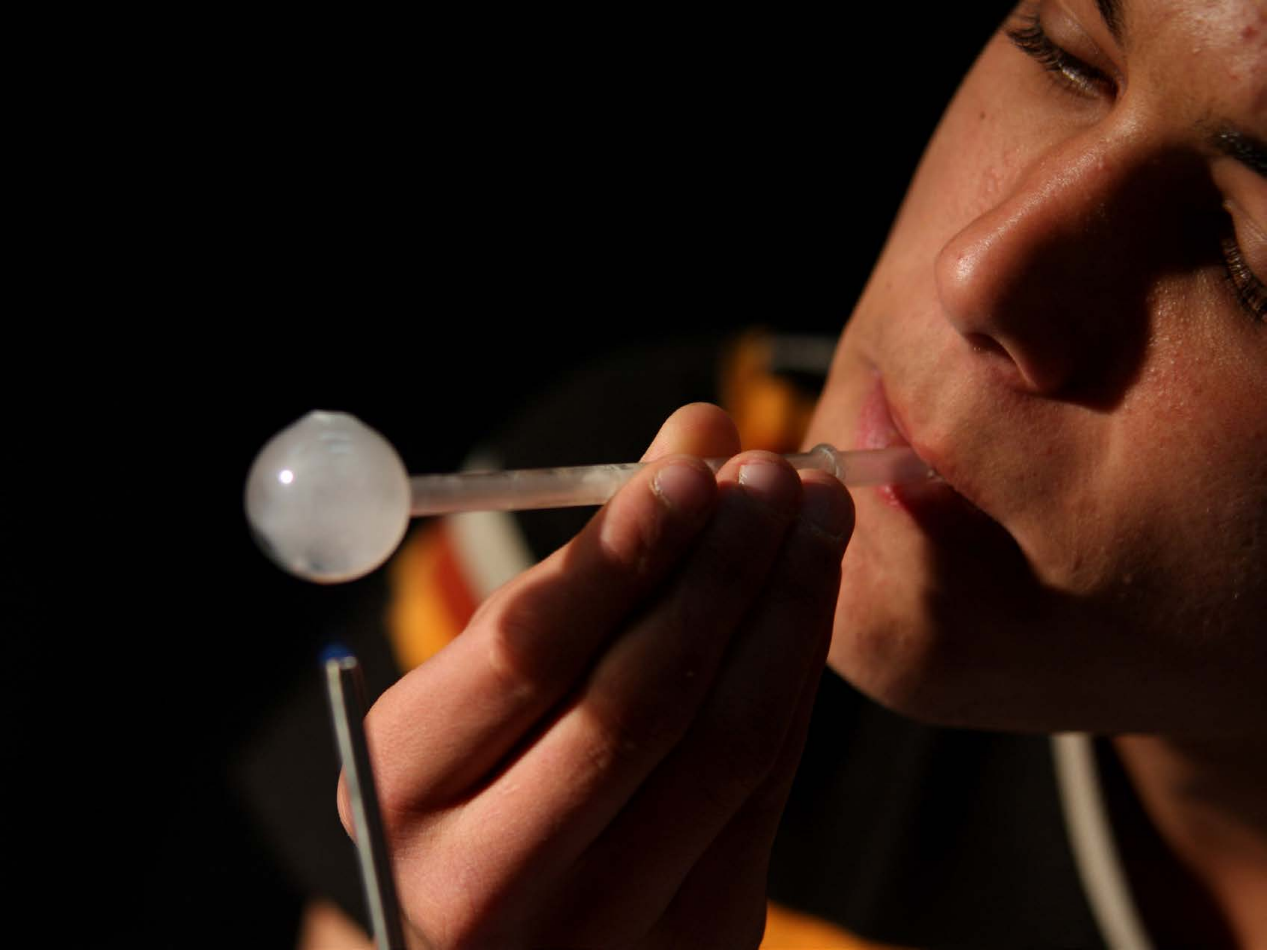

| Category | Craving (0-100) | Valence (1-9) | Arousal (1-9) | Typicality (0-100) | Relatedness | | | | | HSV | | |
|---|---|---|---|---|---|---|---|---|---|---|---|---|
| | | | | | Meth | Opioid | Both | Neither | MethToOpioid | Hue | Saturation | Value |
| Meth Face Activities | 87.63 (16.91) | 4.52 (1.968) | 4.78 (2.74) | 90.85 (14.22) | 0.926 | 0 | 0.074 | 0 | -0.926 | 0.067 (0.183) | 0.433 (0.388) | 0.176 (0.251) |

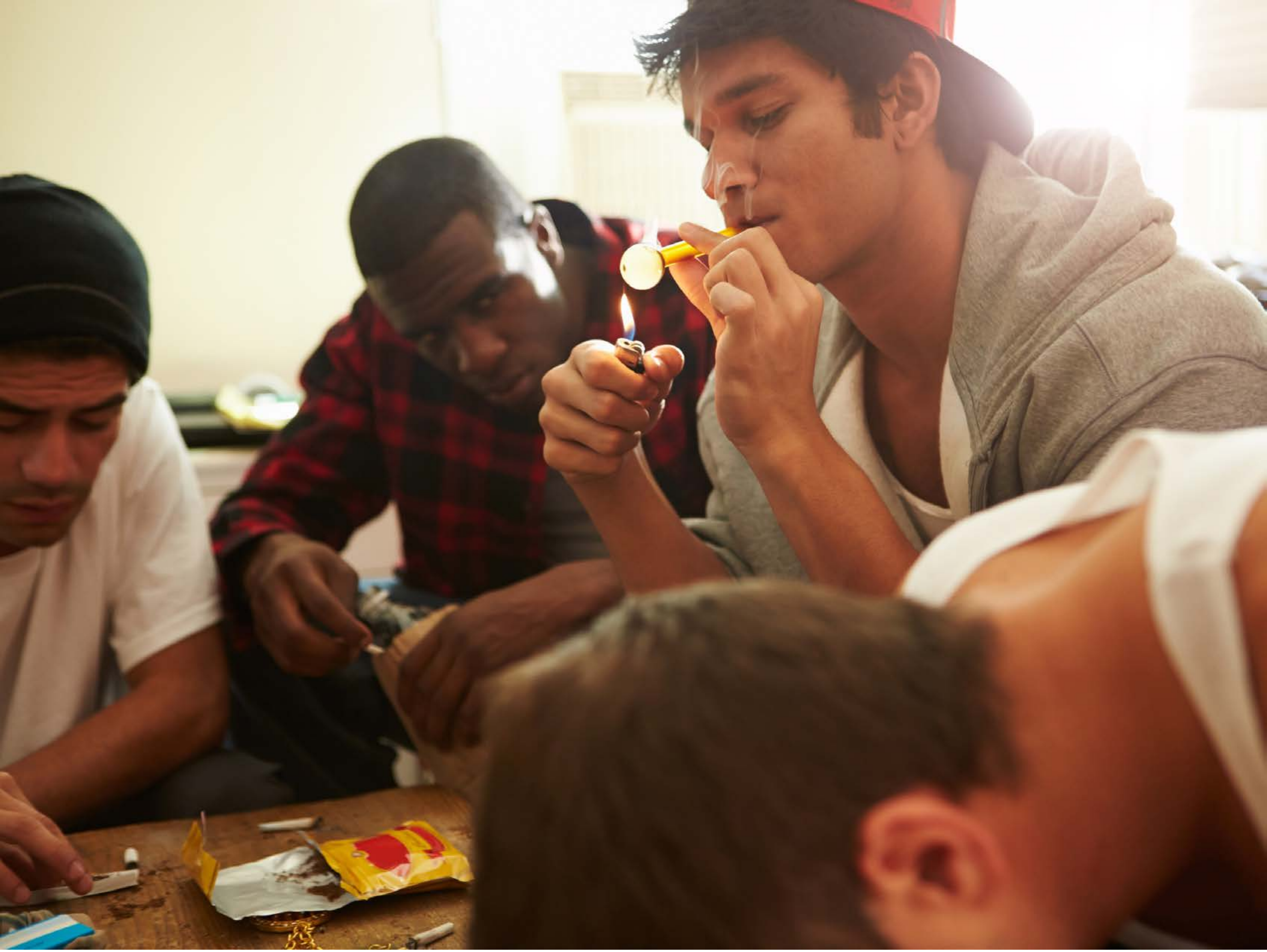

| Category | Craving (0-100) | Valence (1-9) | Arousal (1-9) | Typicality (0-100) | Relatedness | | | | | HSV | | |
|---|---|---|---|---|---|---|---|---|---|---|---|---|
| | | | | | Meth | Opioid | Both | Neither | MethToOpioid | Hue | Saturation | Value |
| Meth Face Activities | 88.19 (16.49) | 4.52 (2.007) | 4.59 (2.9) | 89.7 (16.35) | 0.704 | 0 | 0.259 | 0.037 | -0.704 | 0.099 (0.139) | 0.462 (0.263) | 0.538 (0.293) |

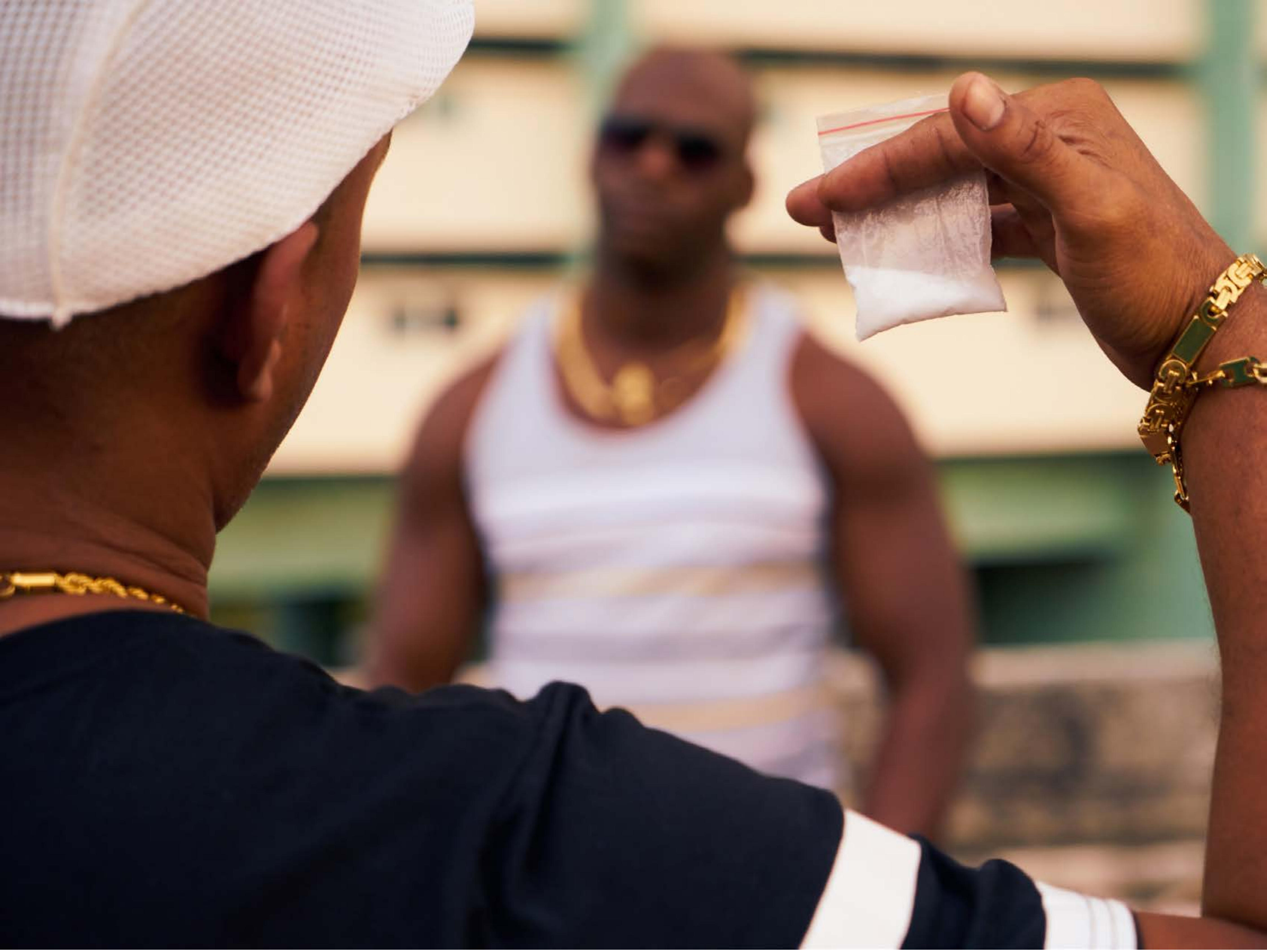

| Category | Craving (0-100) | Valence (1-9) | Arousal (1-9) | Typicality (0-100) | Relatedness | | | | | HSV | | |
|---|---|---|---|---|---|---|---|---|---|---|---|---|
| | | | | | Meth | Opioid | Both | Neither | MethToOpioid | Hue | Saturation | Value |
| Meth Face Activities | 82.44 (19.94) | 4.48 (1.74) | 4.63 (2.63) | 85.78 (18.94) | 0.407 | 0.148 | 0.370 | 0.074 | -0.259 | 0.269 (0.306) | 0.396 (0.228) | 0.52 (0.299) |

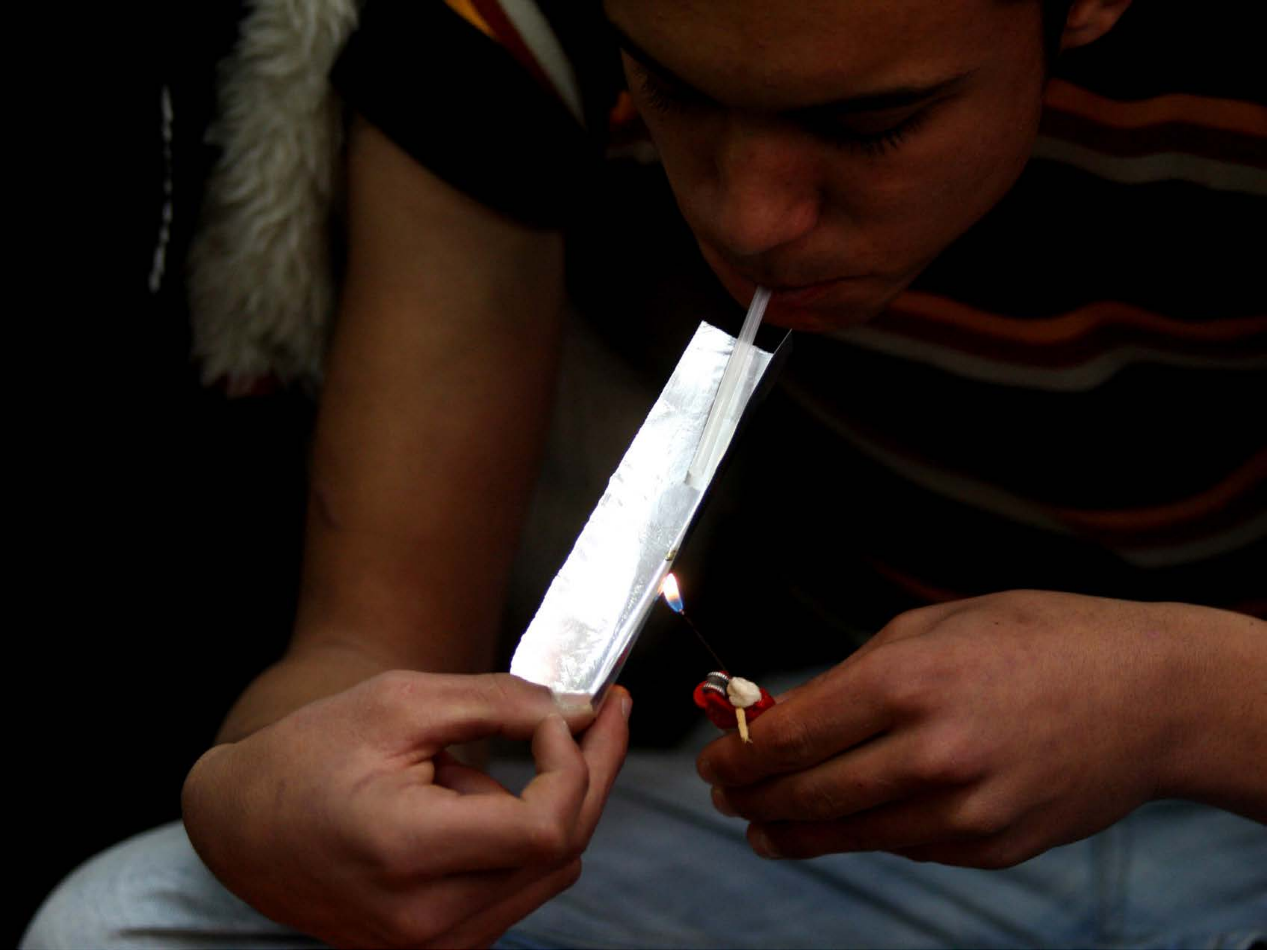

| Category | Craving (0-100) | Valence (1-9) | Arousal (1-9) | Typicality (0-100) | Relatedness | | | | | HSV | | |
|---|---|---|---|---|---|---|---|---|---|---|---|---|
| | | | | | Meth | Opioid | Both | Neither | MethToOpioid | Hue | Saturation | Value |
| Meth Face Activities | 91.89 (13.45) | 4.29 (2.016) | 4.57 (2.91) | 91.89 (13.71) | 0.464 | 0.107 | 0.429 | 0 | -0.357 | 0.11 (0.21) | 0.41 (0.381) | 0.176 (0.211) |

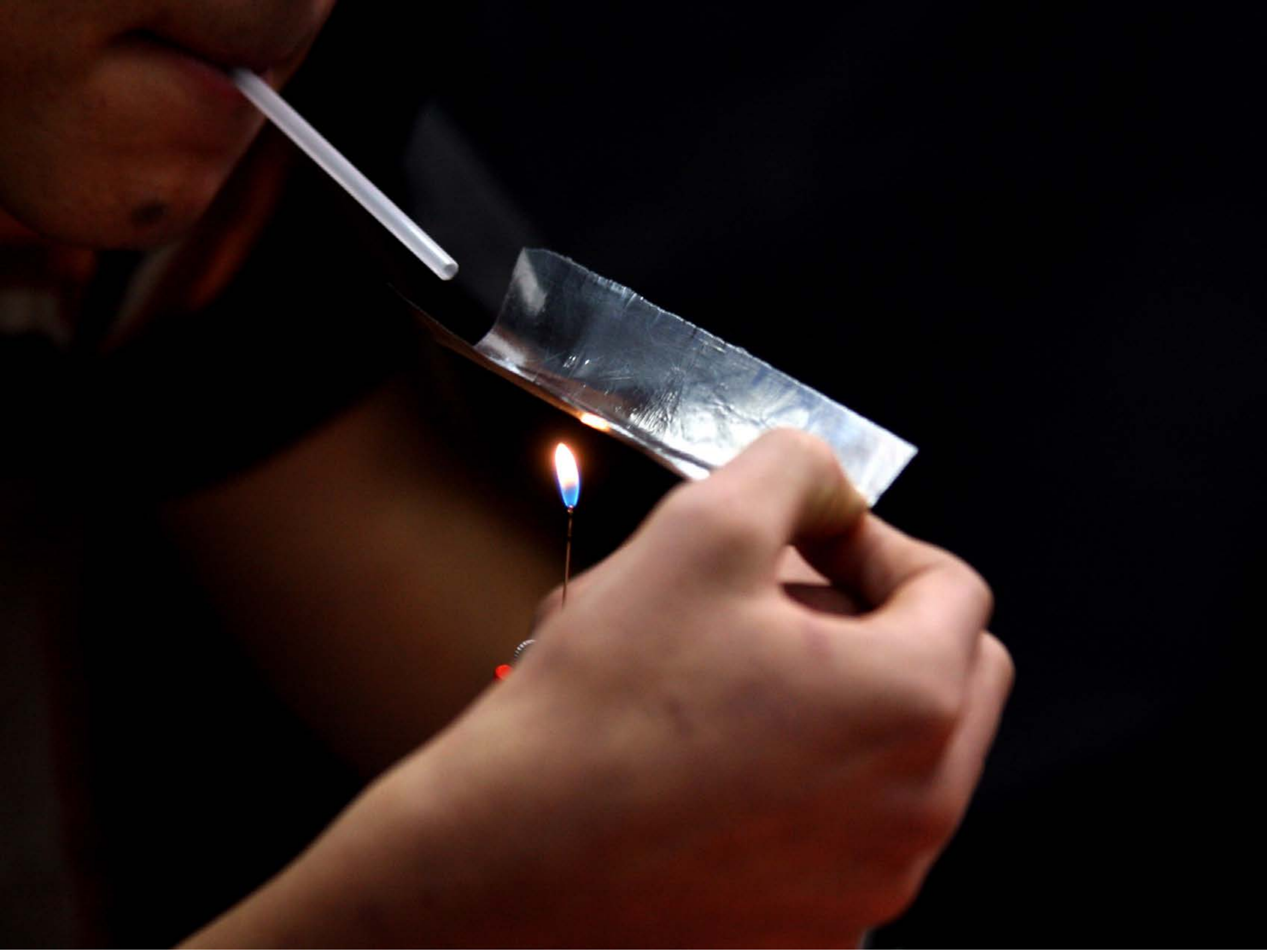

| Category | Craving (0-100) | Valence (1-9) | Arousal (1-9) | Typicality (0-100) | Relatedness | | | | | HSV | | |
|---|---|---|---|---|---|---|---|---|---|---|---|---|
| | | | | | Meth | Opioid | Both | Neither | MethToOpioid | Hue | Saturation | Value |
| Meth Face Activities | 90.18 (15.17) | 4.54 (1.934) | 4.54 (2.74) | 91.07 (13.55) | 0.321 | 0.143 | 0.5 | 0.036 | -0.179 | 0.264 (0.333) | 0.543 (0.386) | 0.153 (0.228) |

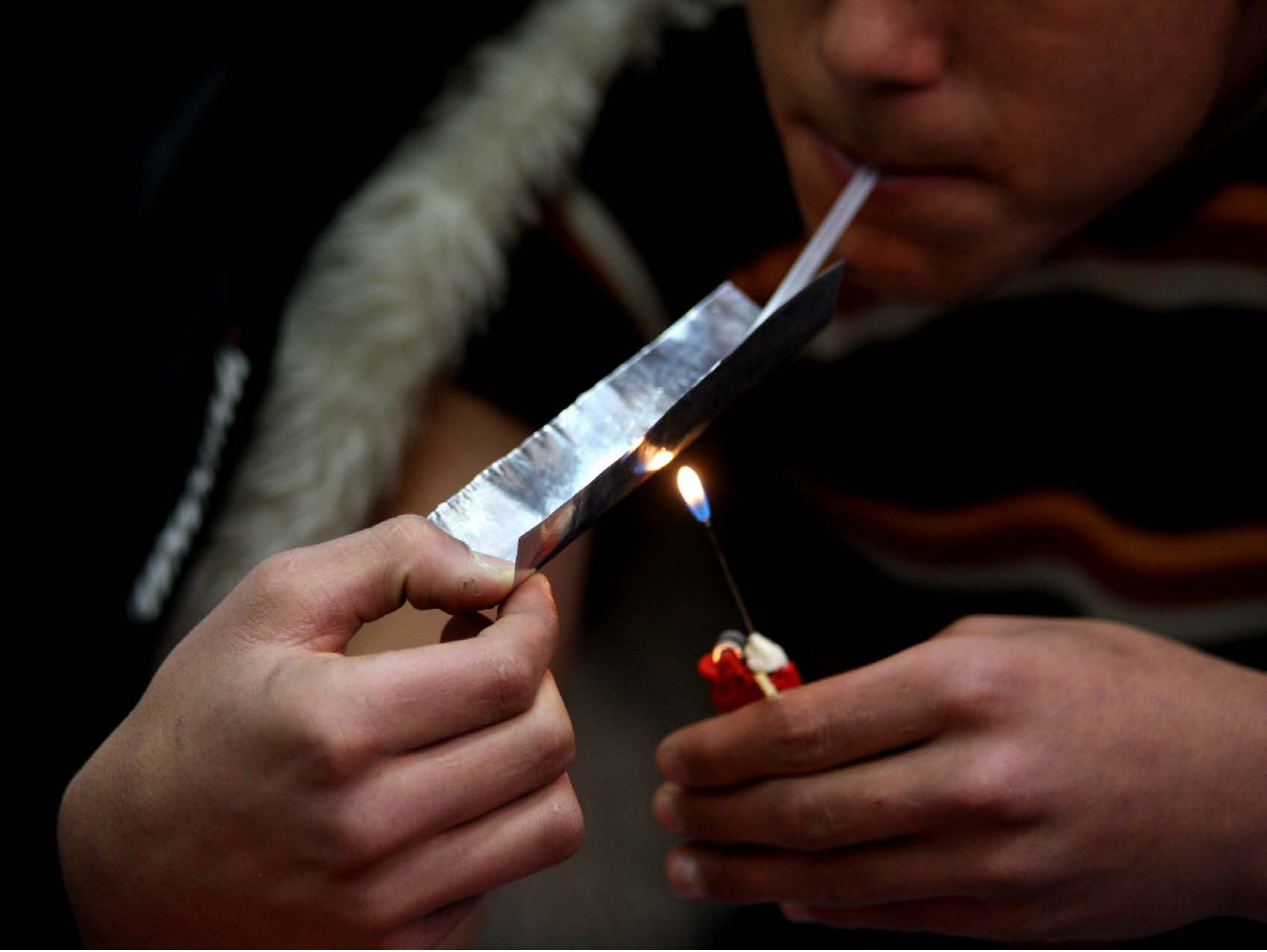

| Category | Craving (0-100) | Valence (1-9) | Arousal (1-9) | Typicality (0-100) | Relatedness | | | | | HSV | | |
|---|---|---|---|---|---|---|---|---|---|---|---|---|
| | | | | | Meth | Opioid | Both | Neither | MethToOpioid | Hue | Saturation | Value |
| Meth Face Activities | 86.85 (18.04) | 4.37 (2.003) | 4.63 (2.87) | 86.11 (18.43) | 0.481 | 0.111 | 0.370 | 0.037 | -0.370 | 0.096 (0.201) | 0.364 (0.342) | 0.206 (0.223) |

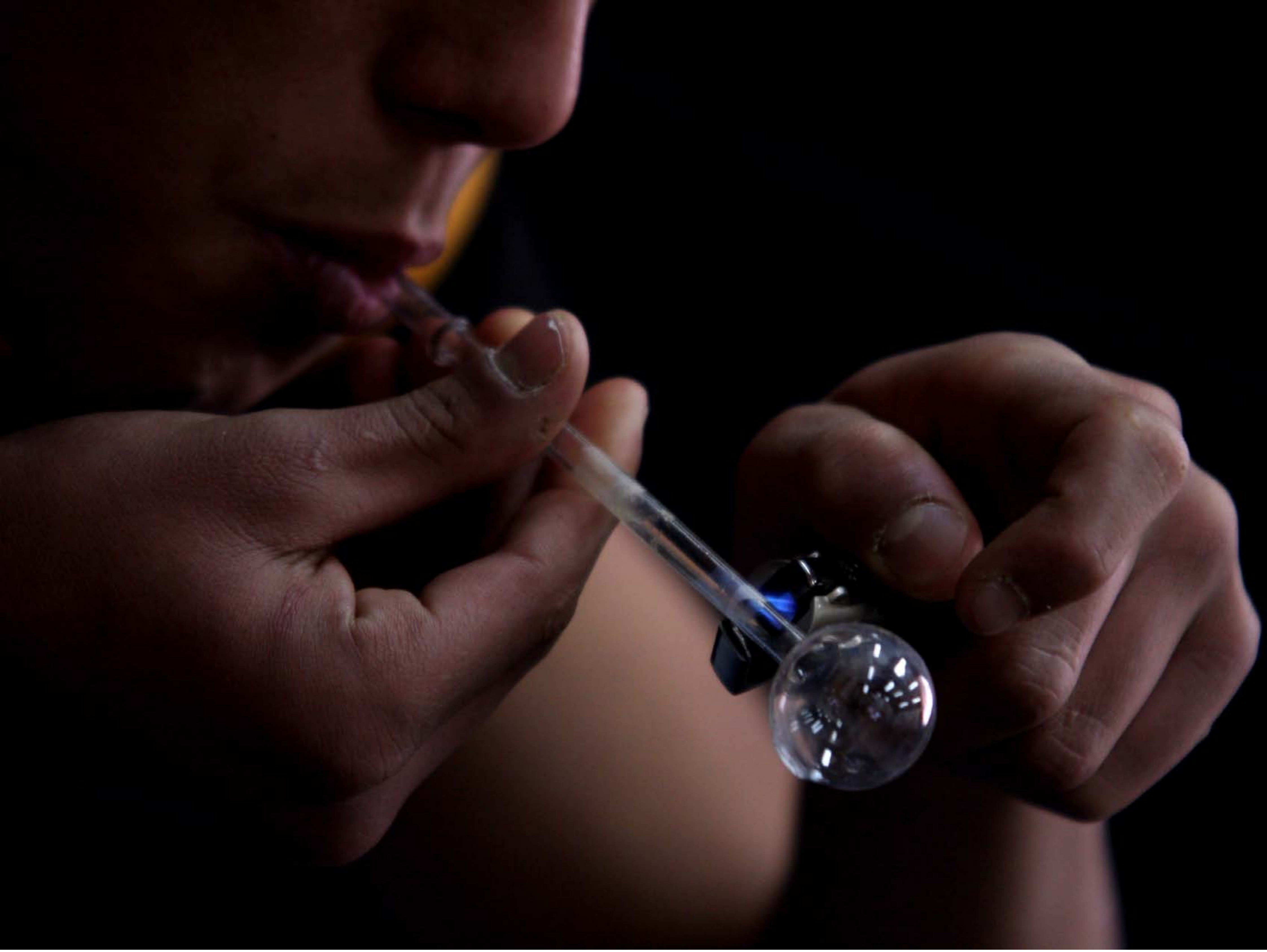

| Category | Craving (0-100) | Valence (1-9) | Arousal (1-9) | Typicality (0-100) | Relatedness | | | | | HSV | | |
|---|---|---|---|---|---|---|---|---|---|---|---|---|
| | | | | | Meth | Opioid | Both | Neither | MethToOpioid | Hue | Saturation | Value |
| Meth Face Activities | 86.63 (17.4) | 4.37 (1.884) | 4.67 (2.6) | 87.7 (18.15) | 0.852 | 0 | 0.148 | 0 | -0.852 | 0.256 (0.4) | 0.449 (0.382) | 0.124 (0.158) |

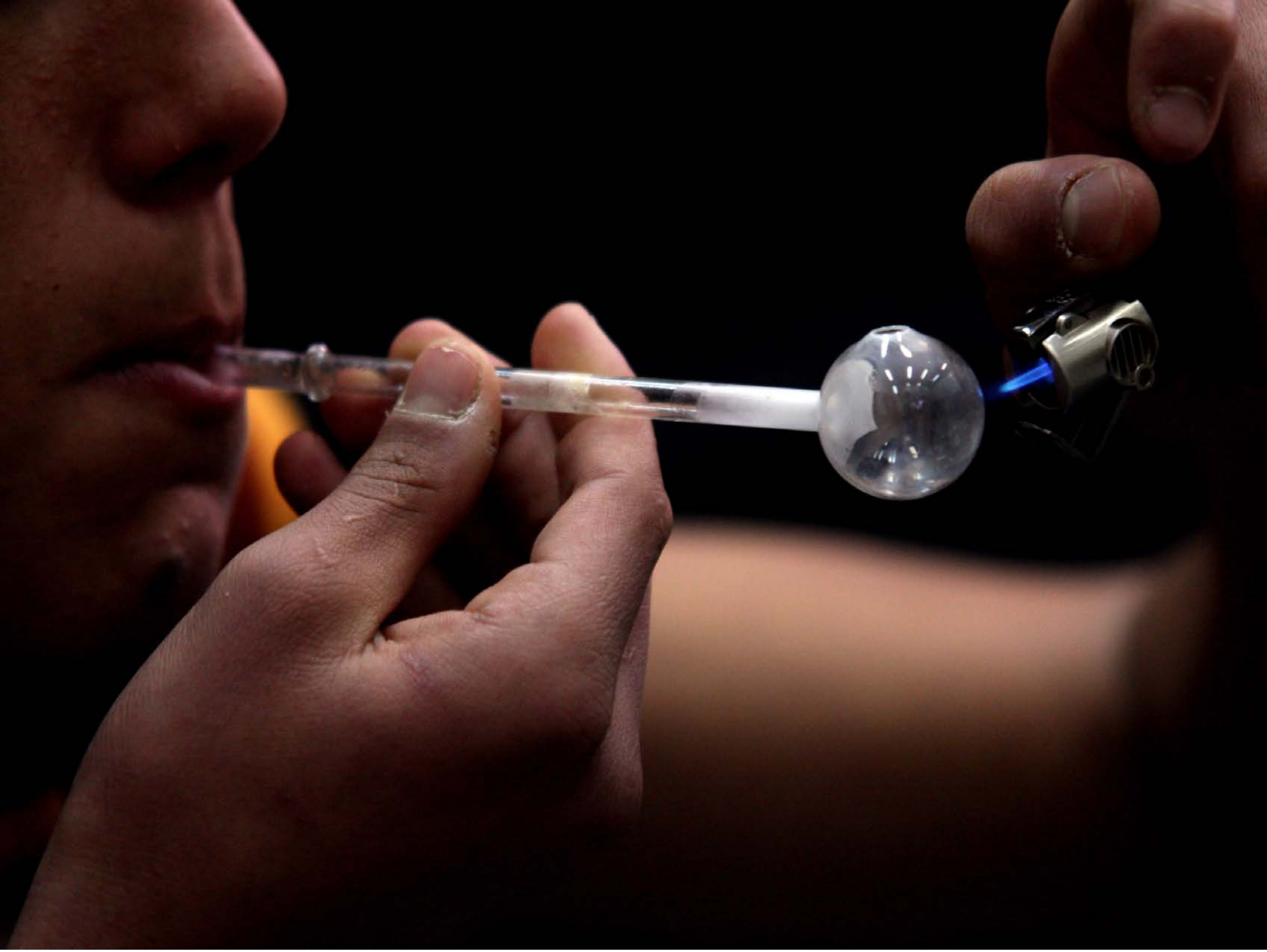

| Category | Craving (0-100) | Valence (1-9) | Arousal (1-9) | Typicality (0-100) | Relatedness | | | | | HSV | | |
|---|---|---|---|---|---|---|---|---|---|---|---|---|
| | | | | | Meth | Opioid | Both | Neither | MethToOpioid | Hue | Saturation | Value |
| Meth Face Activities | 91.93 (11.78) | 4.32 (1.847) | 4.75 (2.86) | 94.68 (10.46) | 0.964 | 0 | 0.036 | 0 | -0.964 | 0.16 (0.321) | 0.492 (0.385) | 0.176 (0.222) |

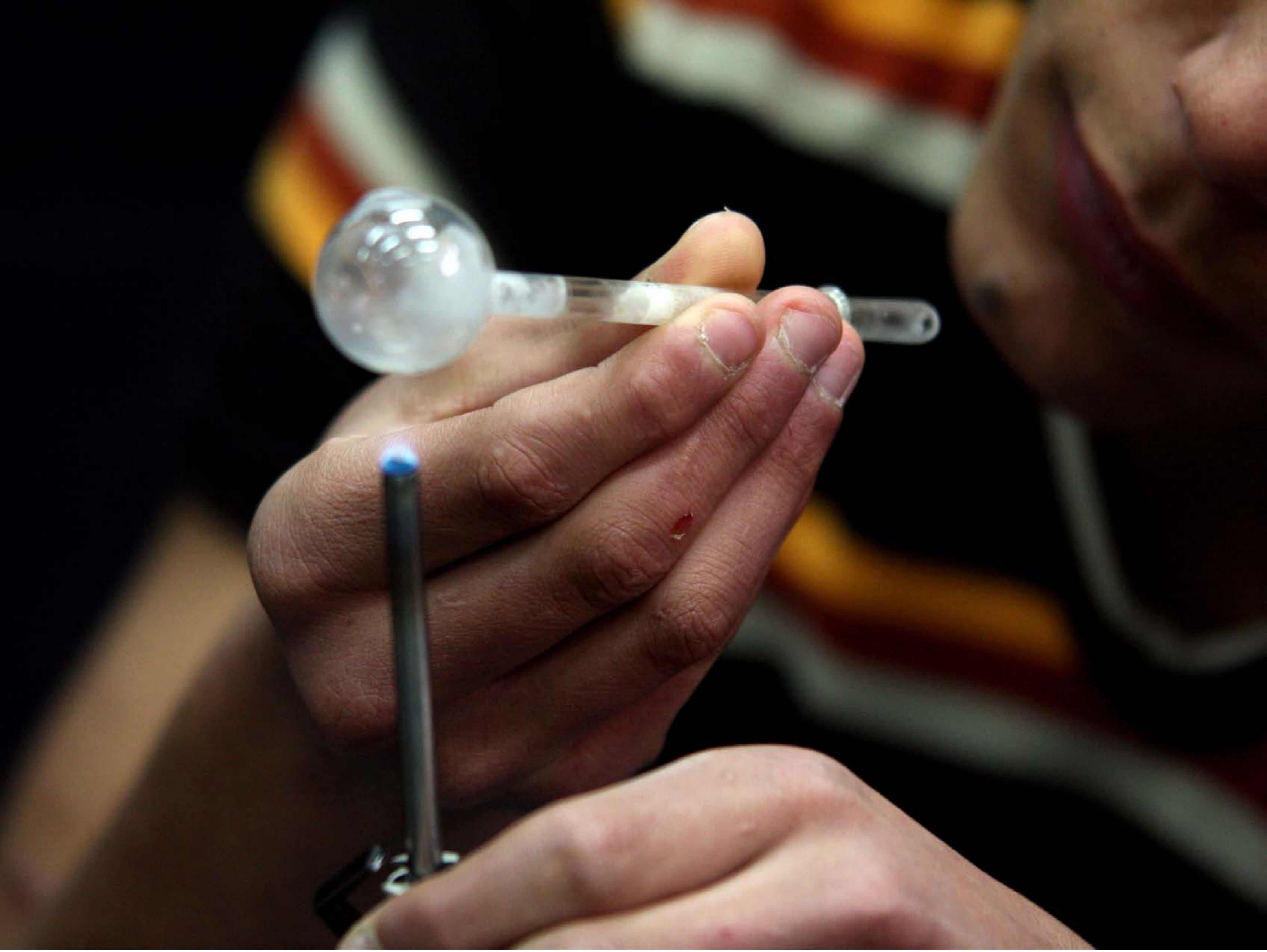

| Category | Craving (0-100) | Valence (1-9) | Arousal (1-9) | Typicality (0-100) | Relatedness | | | | | HSV | | |
|---|---|---|---|---|---|---|---|---|---|---|---|---|
| | | | | | Meth | Opioid | Both | Neither | MethToOpioid | Hue | Saturation | Value |
| Meth Face Activities | 89.57 (22.11) | 4.39 (2.025) | 4.79 (2.74) | 93.68 (13.32) | 0.929 | 0 | 0.071 | 0 | -0.929 | 0.201 (0.29) | 0.469 (0.334) | 0.26 (0.265) |

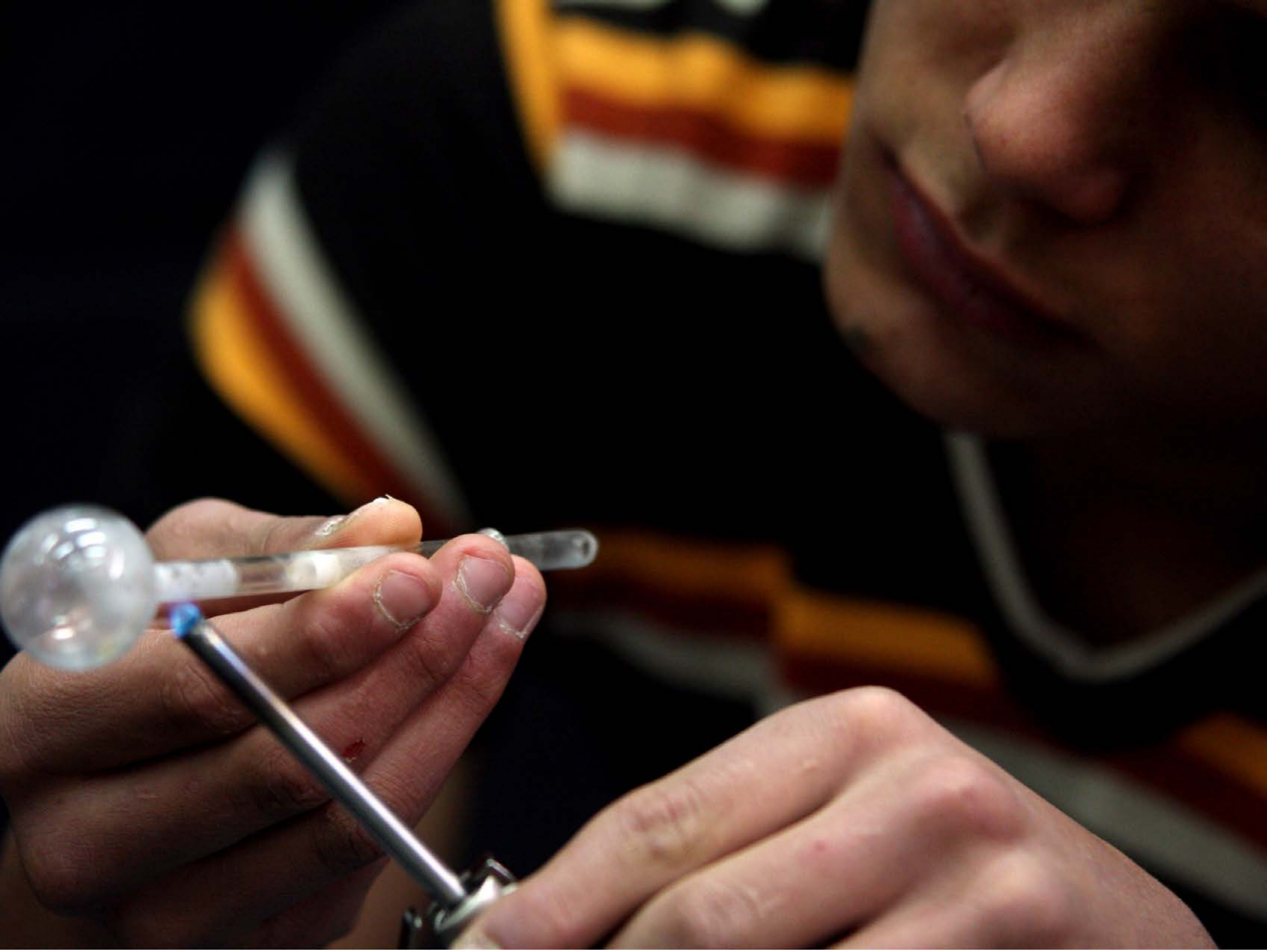

| Category | Craving (0-100) | Valence (1-9) | Arousal (1-9) | Typicality (0-100) | Relatedness | | | | | HSV | | |
|---|---|---|---|---|---|---|---|---|---|---|---|---|
| | | | | | Meth | Opioid | Both | Neither | MethToOpioid | Hue | Saturation | Value |
| Meth Face Activities | 86.11 (18.79) | 4.63 (1.864) | 4.67 (2.77) | 86.56 (19.42) | 0.926 | 0 | 0.074 | 0 | -0.926 | 0.209 (0.313) | 0.508 (0.352) | 0.238 (0.267) |

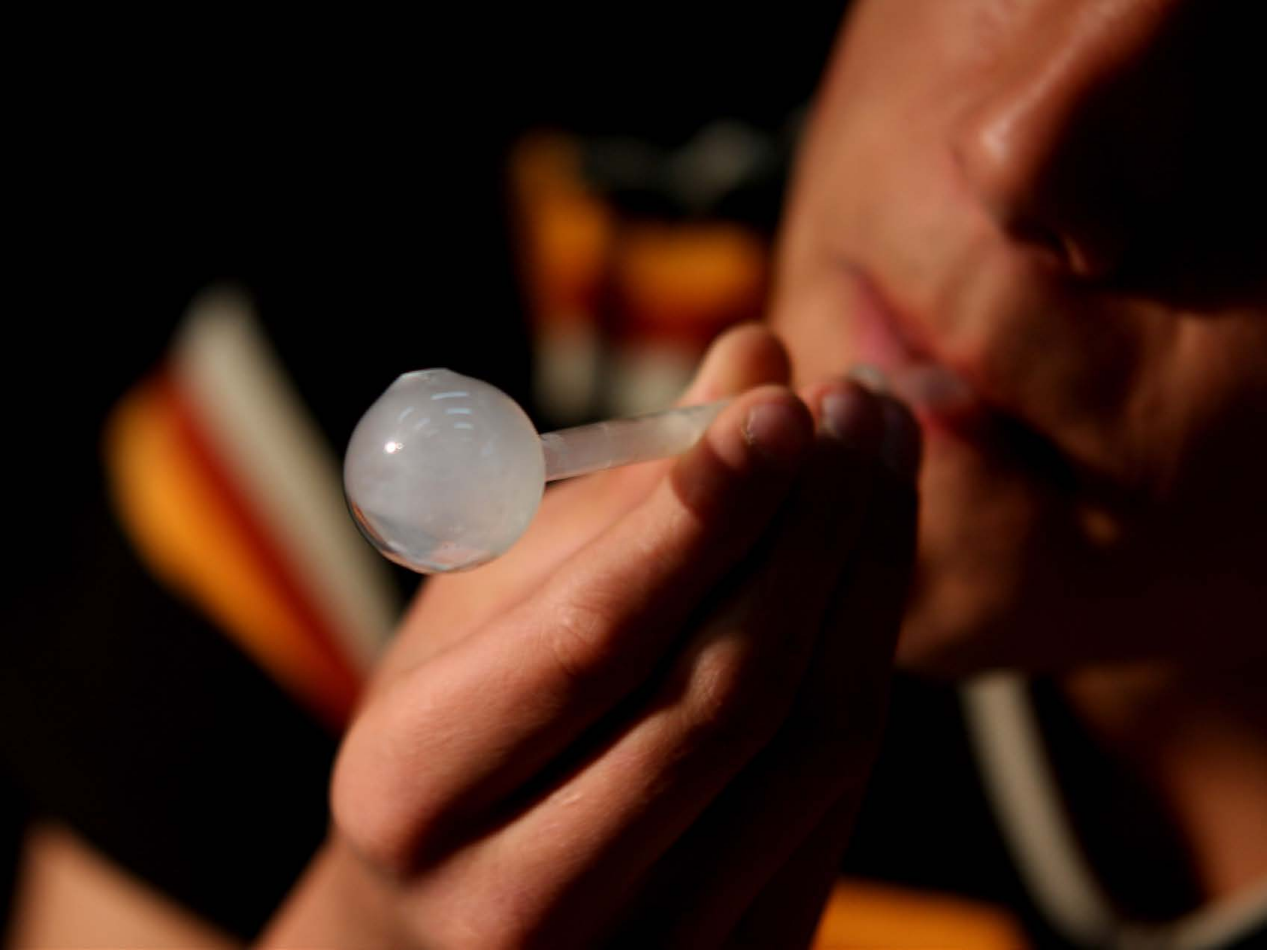

| Category | Craving (0-100) | Valence (1-9) | Arousal (1-9) | Typicality (0-100) | Relatedness | | | | | HSV | | |
|---|---|---|---|---|---|---|---|---|---|---|---|---|
| | | | | | Meth | Opioid | Both | Neither | MethToOpioid | Hue | Saturation | Value |
| Meth Face Activities | 85.78 (18.34) | 4.48 (1.762) | 4.56 (2.62) | 84.3 (20.76) | 0.963 | 0 | 0.037 | 0 | -0.963 | 0.134 (0.295) | 0.579 (0.408) | 0.245 (0.285) |

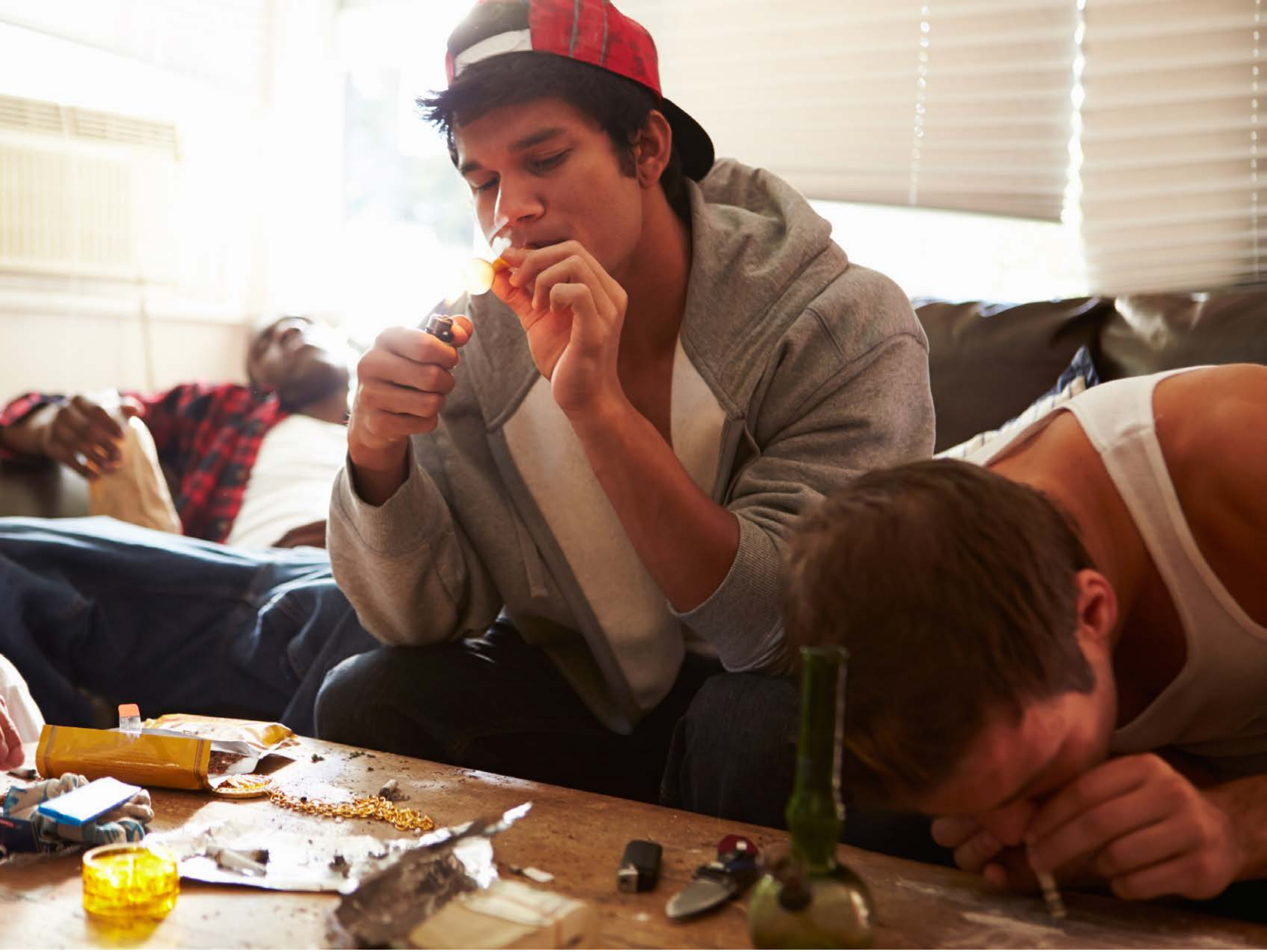

| Category | Craving (0-100) | Valence (1-9) | Arousal (1-9) | Typicality (0-100) | Relatedness | | | | | HSV | | |
|---|---|---|---|---|---|---|---|---|---|---|---|---|
| | | | | | Meth | Opioid | Both | Neither | MethToOpioid | Hue | Saturation | Value |
| Meth Face Activities | 88.04 (16.58) | 4.33 (2.13) | 4.67 (2.84) | 87.37 (18.37) | 0.519 | 0 | 0.481 | 0 | -0.519 | 0.139 (0.227) | 0.395 (0.29) | 0.553 (0.318) |

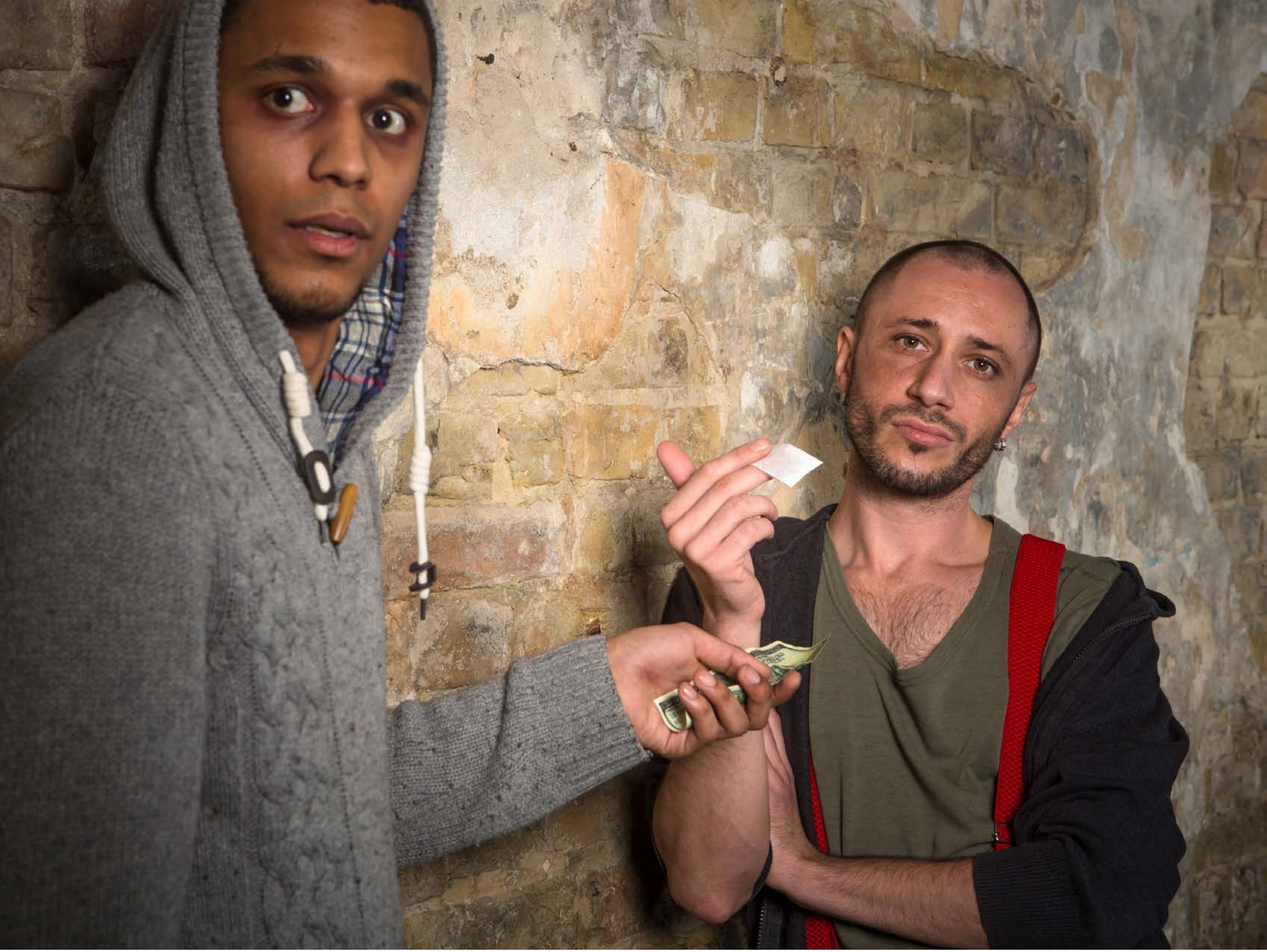

| Category | Craving (0-100) | Valence (1-9) | Arousal (1-9) | Typicality (0-100) | Relatedness | | | | | HSV | | |
|---|---|---|---|---|---|---|---|---|---|---|---|---|
| | | | | | Meth | Opioid | Both | Neither | MethToOpioid | Hue | Saturation | Value |
| Meth Face Activities | 86.29 (17.92) | 4.75 (1.506) | 4.5 (2.62) | 90.29 (16.58) | 0.25 | 0.036 | 0.714 | 0 | -0.214 | 0.153 (0.236) | 0.264 (0.185) | 0.473 (0.23) |

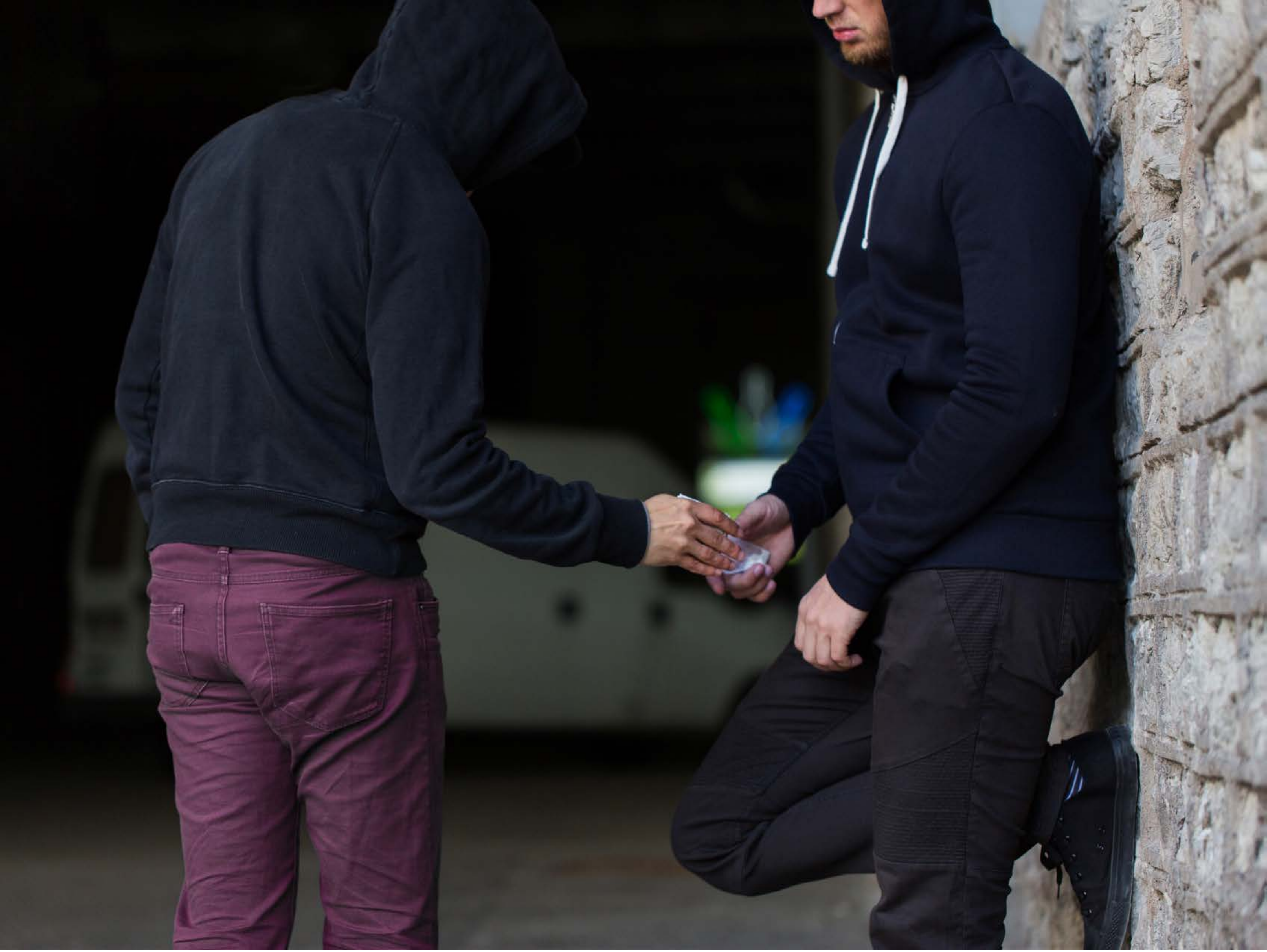

| Category | Craving (0-100) | Valence (1-9) | Arousal (1-9) | Typicality (0-100) | Relatedness | | | | | HSV | | |
|---|---|---|---|---|---|---|---|---|---|---|---|---|
| | | | | | Meth | Opioid | Both | Neither | MethToOpioid | Hue | Saturation | Value |
| Meth Face Activities | 86.25 (16.09) | 4.43 (1.933) | 4.71 (3) | 90.14 (12.66) | 0 | 0.036 | 0.964 | 0 | 0.036 | 0.49 (0.313) | 0.322 (0.231) | 0.207 (0.203) |

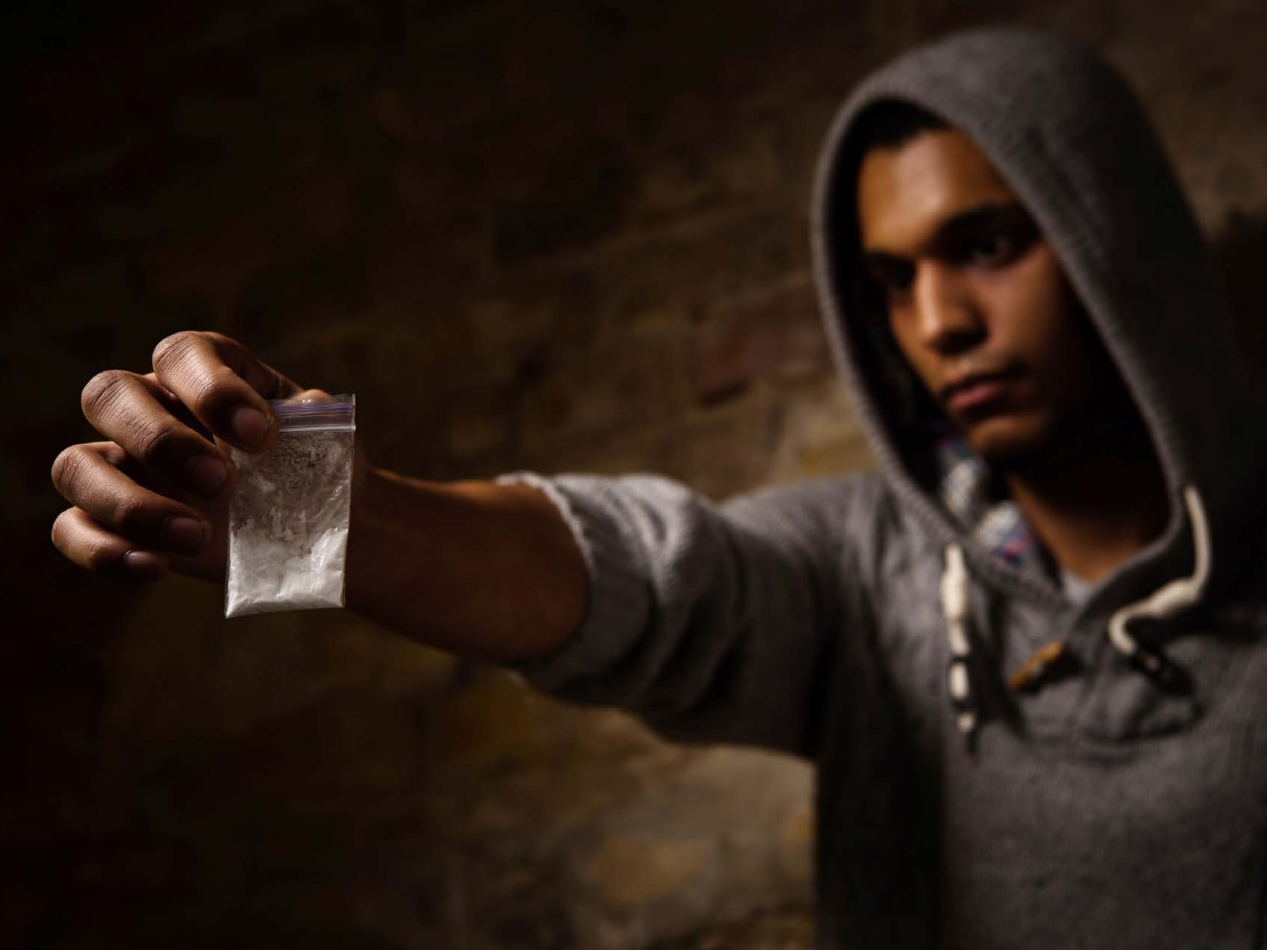

| Category | Craving (0-100) | Valence (1-9) | Arousal (1-9) | Typicality (0-100) | Relatedness | | | | | HSV | | |
|---|---|---|---|---|---|---|---|---|---|---|---|---|
| | | | | | Meth | Opioid | Both | Neither | MethToOpioid | Hue | Saturation | Value |
| Meth Face Activities | 87.71 (14.35) | 4.54 (1.934) | 4.75 (2.78) | 93.29 (12.22) | 0.393 | 0.107 | 0.5 | 0 | -0.286 | 0.092 (0.151) | 0.609 (0.242) | 0.162 (0.143) |
