## Supplementary Material 4 for "LIBR Methamphetamine and Opioid Cue Database (LIBR MOCD): Development and Validation"

### Meth and Opioid Cue Database

#### Opioid Image

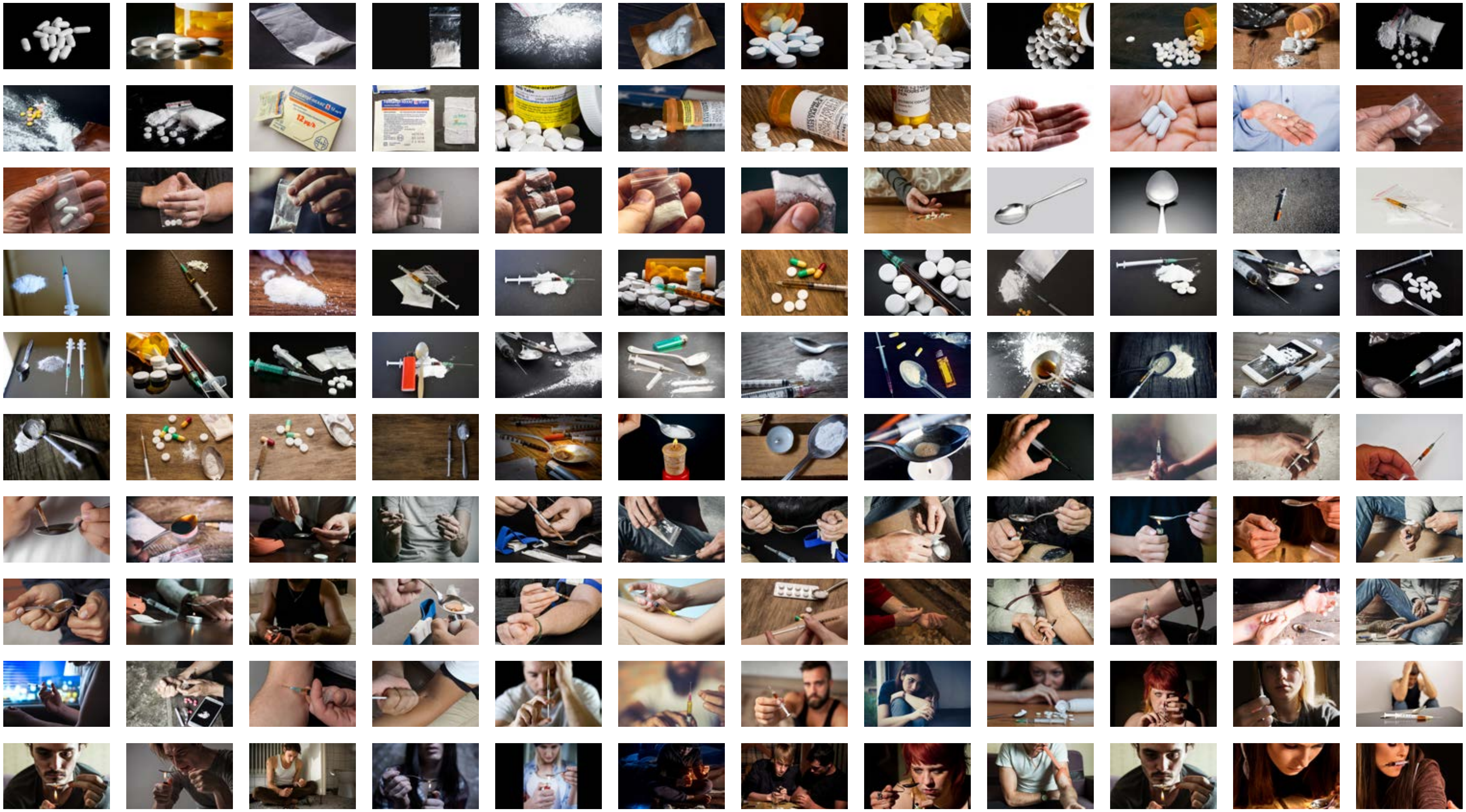

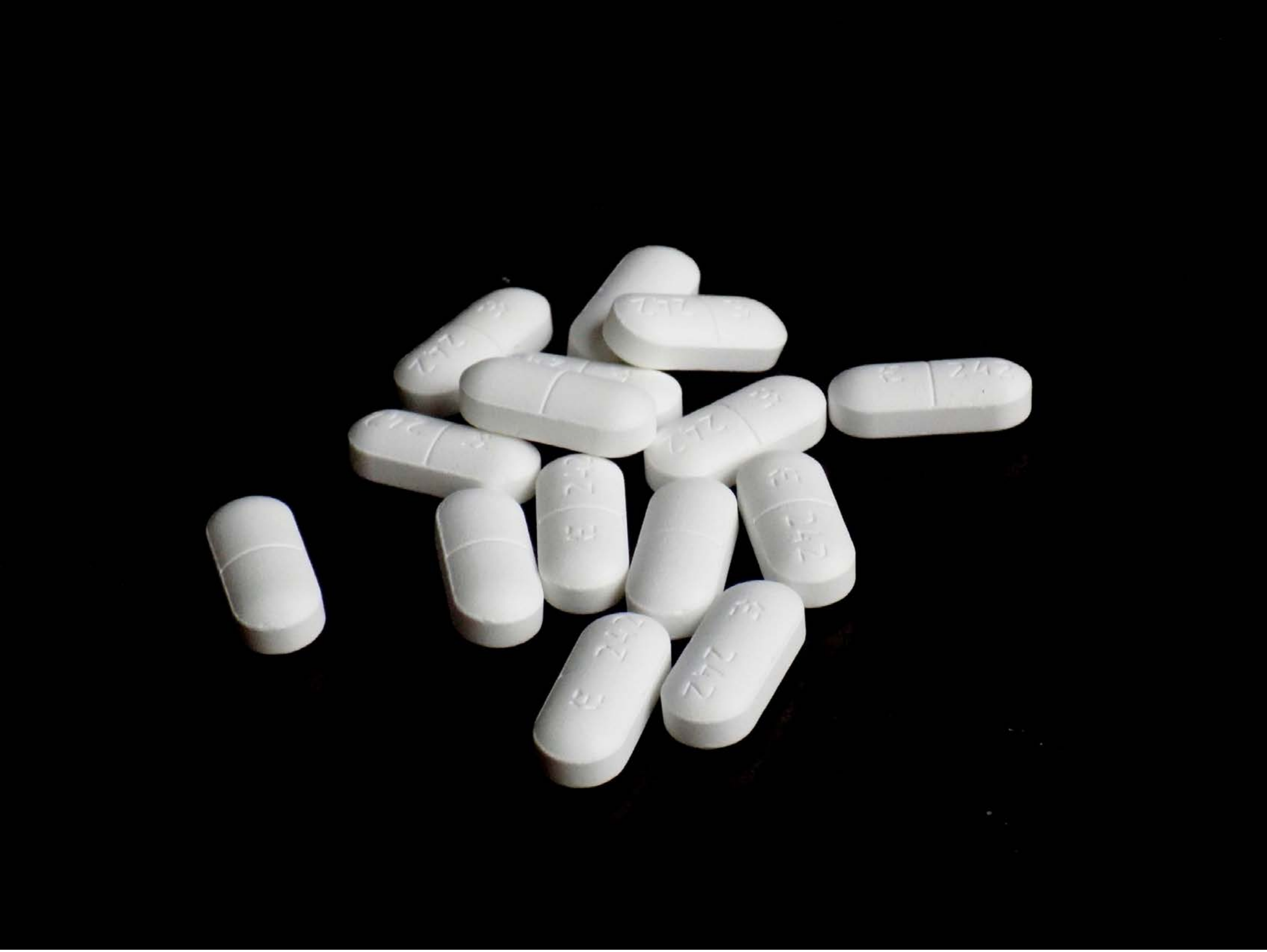

| Category | Craving (0-100) | Valence (1-9) | Arousal (1-9) | Typicality (0-100) | Relatedness | | | | | HSV | | |
|---|---|---|---|---|---|---|---|---|---|---|---|---|
| | | | | | Meth | Opioid | Both | Neither | MethToOpioid | Hue | Saturation | Value |
| Opioid | 88.04 (14.1) | 4.68 (1.722) | 4.39 (2.75) | 92.14 (11.57) | 0 | 0.929 | 0.071 | 0 | 0.929 | 0.028 (0.114) | 0.013 (0.093) | 0.115 (0.275) |

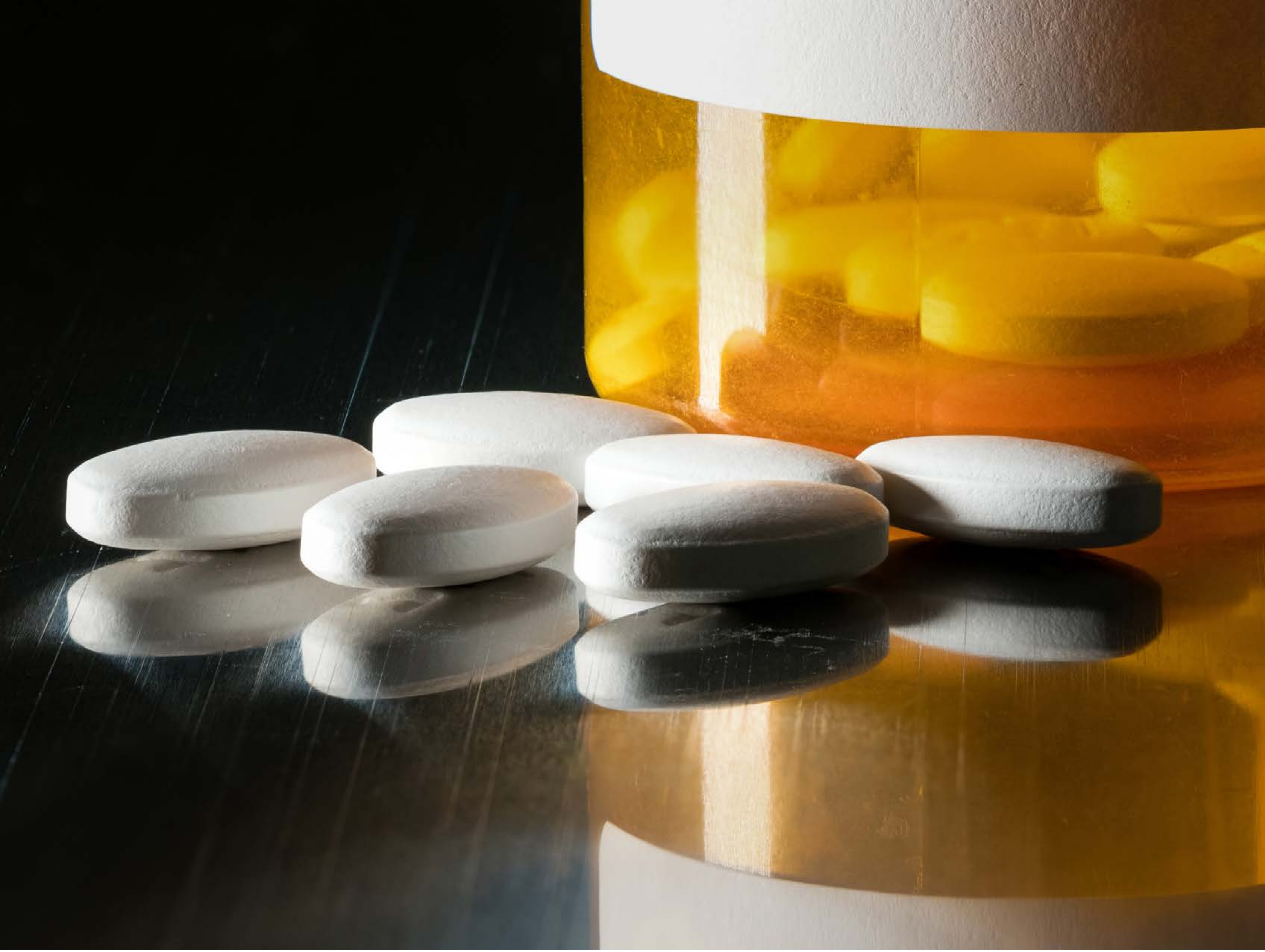

| Category | Craving (0-100) | Valence (1-9) | Arousal (1-9) | Typicality (0-100) | Relatedness | | | | | HSV | | |
|---|---|---|---|---|---|---|---|---|---|---|---|---|
| | | | | | Meth | Opioid | Both | Neither | MethToOpioid | Hue | Saturation | Value |
| Opioid | 81.81 (22.99) | 4.74 (1.655) | 4.56 (2.64) | 83.26 (22.42) | 0 | 0.963 | 0.037 | 0 | 0.963 | 0.164 (0.114) | 0.459 (0.352) | 0.427 (0.327) |

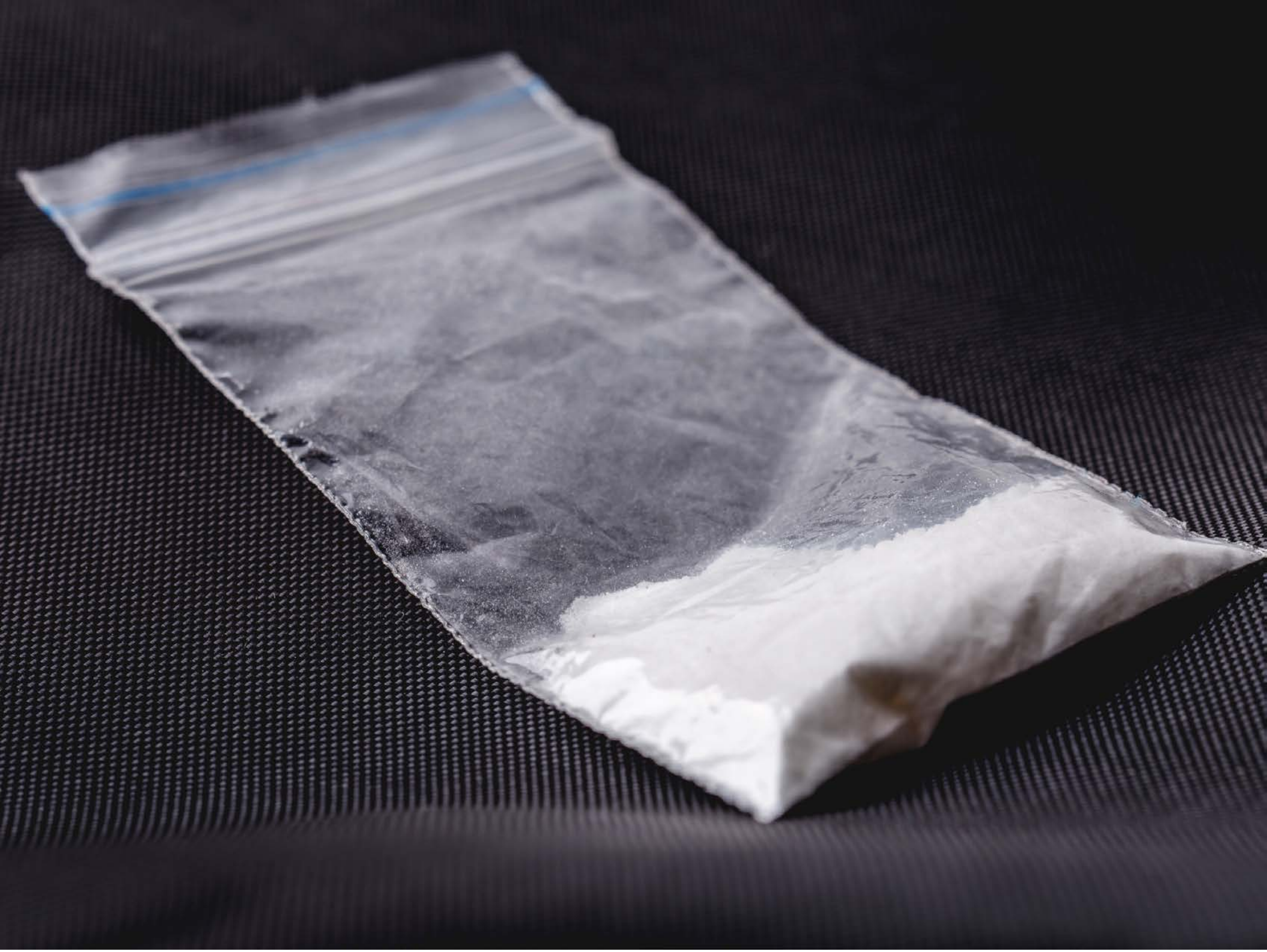

| Category | Craving (0-100) | Valence (1-9) | Arousal (1-9) | Typicality (0-100) | Relatedness | | | | | HSV | | |
|---|---|---|---|---|---|---|---|---|---|---|---|---|
| | | | | | Meth | Opioid | Both | Neither | MethToOpioid | Hue | Saturation | Value |
| Opioid | 89.25 (13.92) | 4.64 (1.81) | 4.46 (2.76) | 94.11 (11.11) | 0.429 | 0.036 | 0.5 | 0.036 | -0.393 | 0.646 (0.236) | 0.123 (0.075) | 0.328 (0.256) |

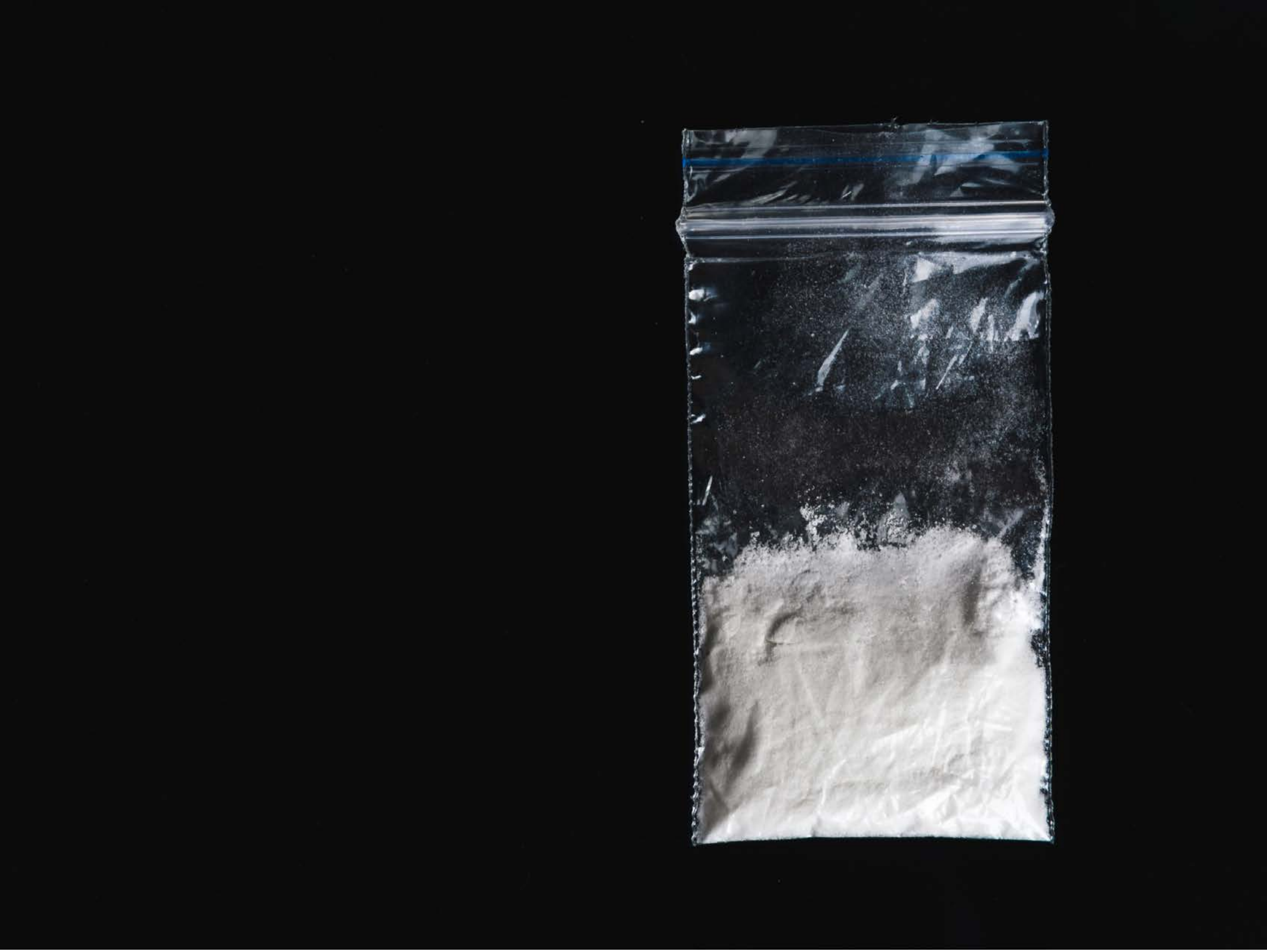

| Category | Craving (0-100) | Valence (1-9) | Arousal (1-9) | Typicality (0-100) | Relatedness | | | | | HSV | | |
|---|---|---|---|---|---|---|---|---|---|---|---|---|
| | | | | | Meth | Opioid | Both | Neither | MethToOpioid | Hue | Saturation | Value |
| Opioid | 89.74 (15.26) | 4.48 (2.007) | 4.59 (2.85) | 90.22 (15.42) | 0.296 | 0.296 | 0.407 | 0 | 0 | 0.118 (0.248) | 0.031 (0.083) | 0.133 (0.218) |

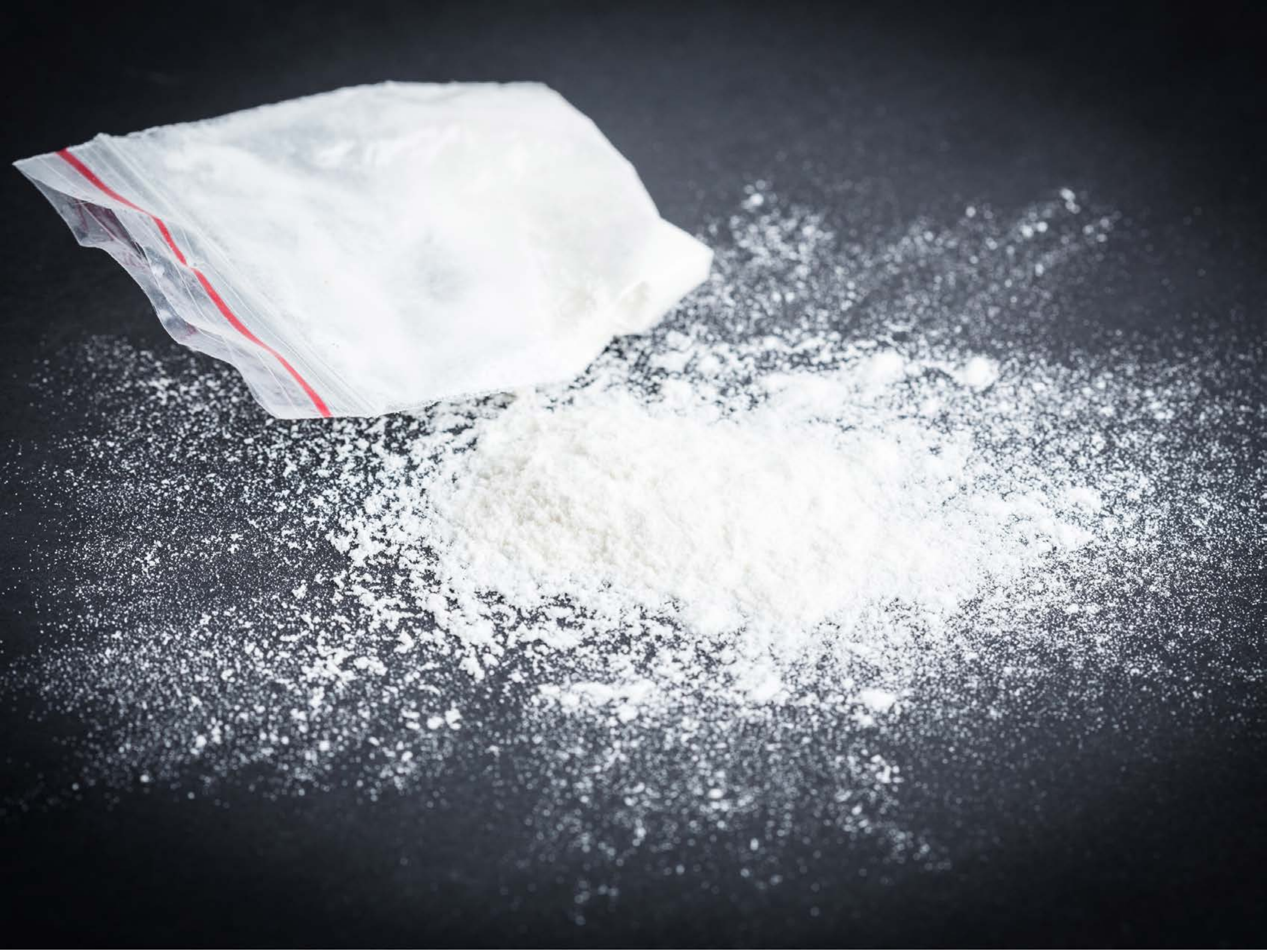

| Category | Craving (0-100) | Valence (1-9) | Arousal (1-9) | Typicality (0-100) | Relatedness | | | | | HSV | | |
|---|---|---|---|---|---|---|---|---|---|---|---|---|
| | | | | | Meth | Opioid | Both | Neither | MethToOpioid | Hue | Saturation | Value |
| Opioid | 87.61 (20.78) | 4.43 (1.814) | 4.89 (2.53) | 90.32 (19.59) | 0.5 | 0.107 | 0.393 | 0 | -0.393 | 0.582 (0.124) | 0.149 (0.098) | 0.495 (0.331) |

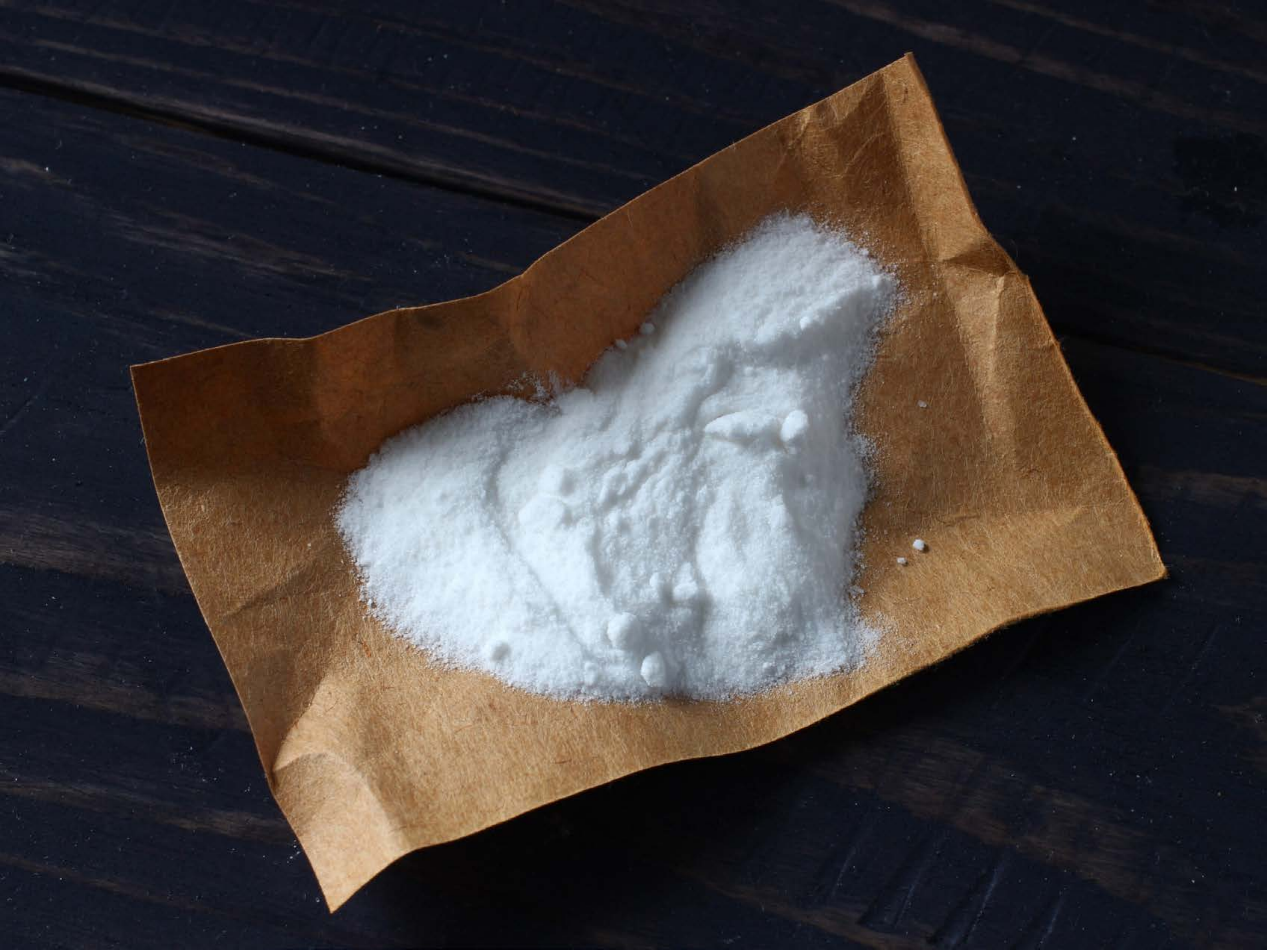

| Category | Craving (0-100) | Valence (1-9) | Arousal (1-9) | Typicality (0-100) | Relatedness | | | | | HSV | | |
|---|---|---|---|---|---|---|---|---|---|---|---|---|
| | | | | | Meth | Opioid | Both | Neither | MethToOpioid | Hue | Saturation | Value |
| Opioid | 84.43 (21.1) | 4.75 (1.456) | 4.46 (2.56) | 88.96 (18.55) | 0.393 | 0.179 | 0.429 | 0 | -0.214 | 0.483 (0.26) | 0.383 (0.184) | 0.296 (0.259) |

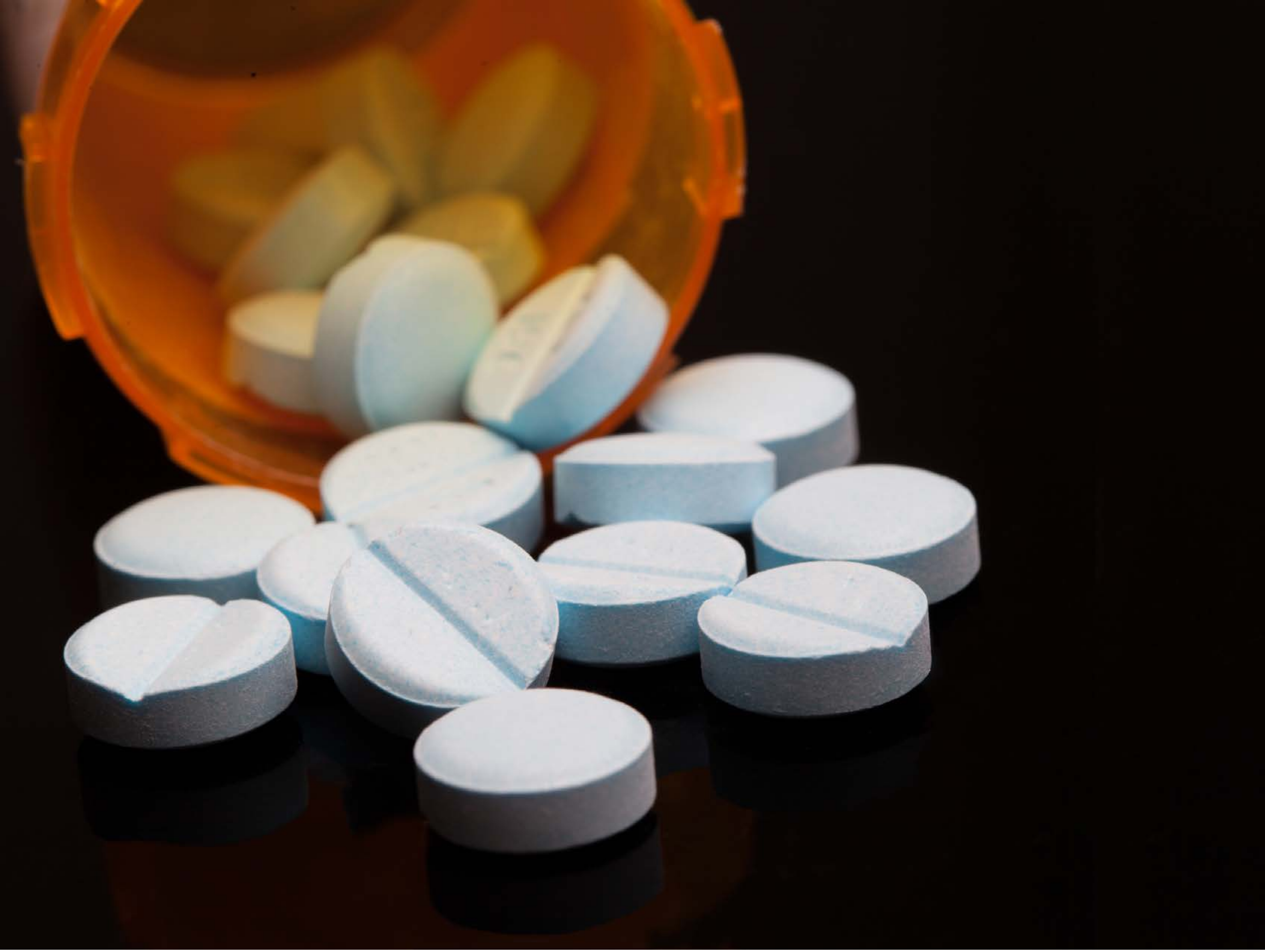

| Category | Craving (0-100) | Valence (1-9) | Arousal (1-9) | Typicality (0-100) | Relatedness | | | | | HSV | | |
|---|---|---|---|---|---|---|---|---|---|---|---|---|
| | | | | | Meth | Opioid | Both | Neither | MethToOpioid | Hue | Saturation | Value |
| Opioid | 84.41 (19.57) | 4.89 (2.006) | 4.44 (2.68) | 85.81 (19.27) | 0 | 0.963 | 0.037 | 0 | 0.963 | 0.205 (0.286) | 0.538 (0.337) | 0.313 (0.321) |

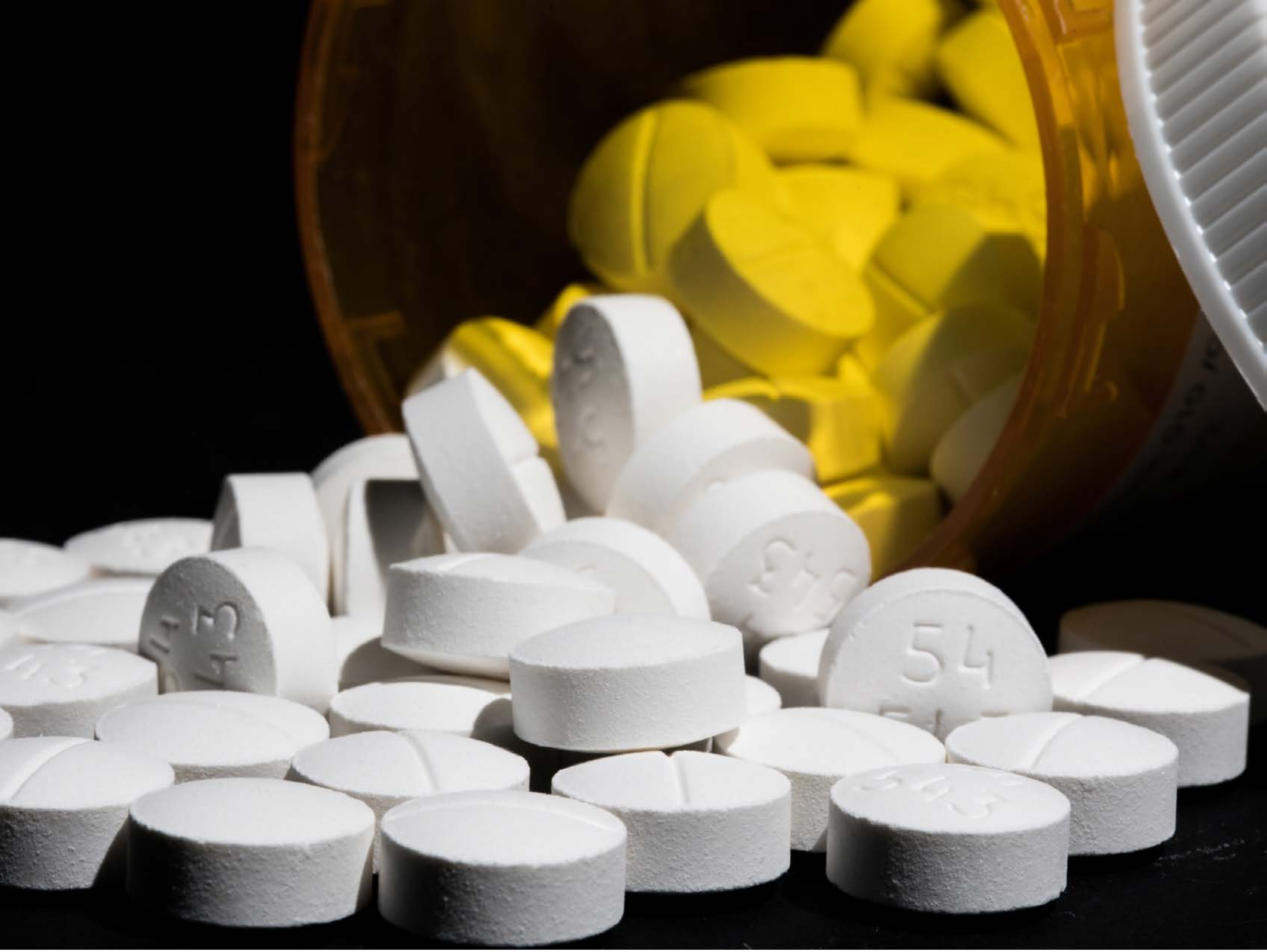

| Category | Craving (0-100) | Valence (1-9) | Arousal (1-9) | Typicality (0-100) | Relatedness | | | | | HSV | | |
|---|---|---|---|---|---|---|---|---|---|---|---|---|
| | | | | | Meth | Opioid | Both | Neither | MethToOpioid | Hue | Saturation | Value |
| Opioid | 83.44 (19.71) | 4.85 (1.634) | 4.3 (2.77) | 84.48 (19.66) | 0 | 0.963 | 0 | 0.037 | 0.963 | 0.181 (0.261) | 0.318 (0.42) | 0.459 (0.354) |

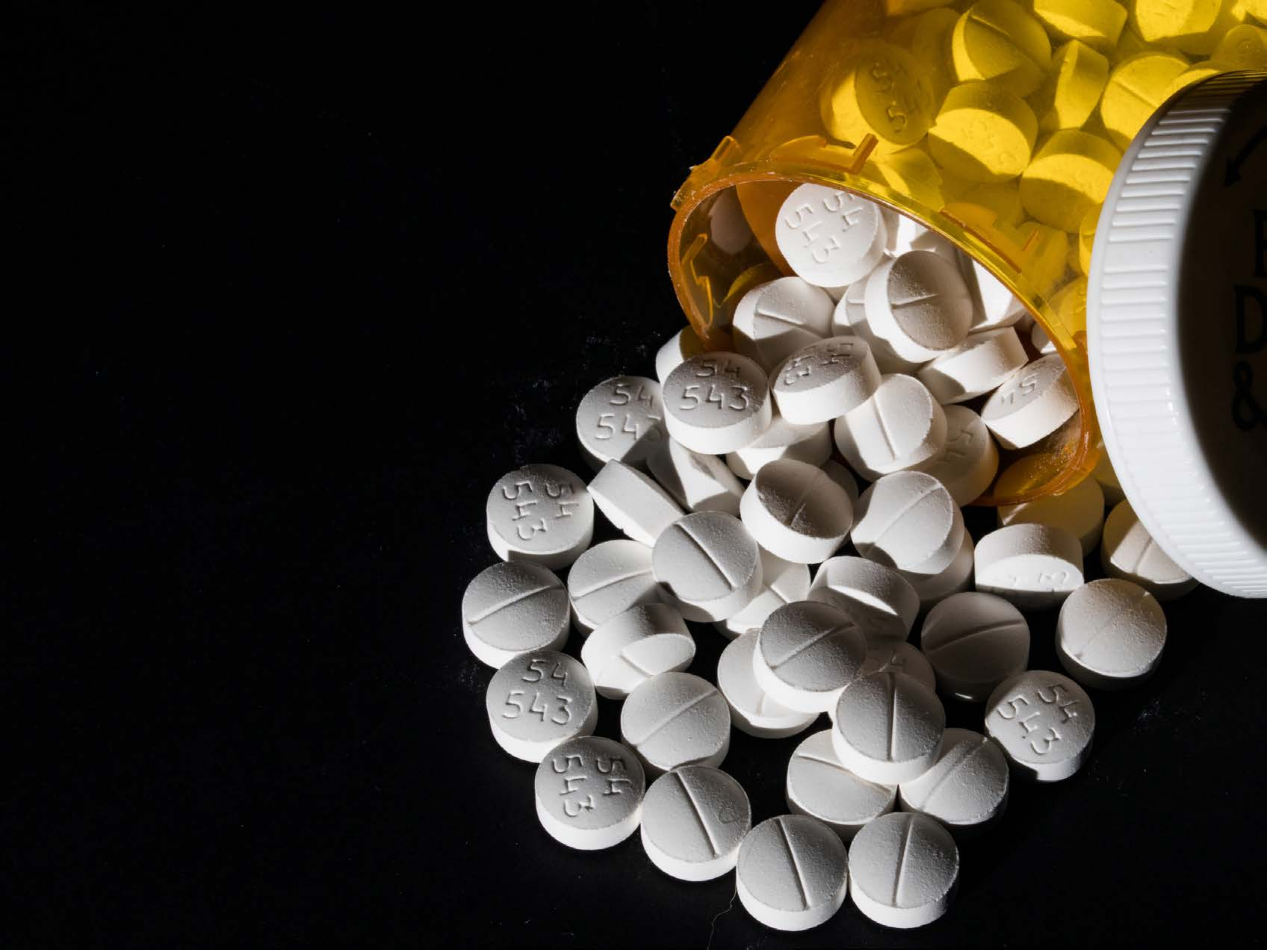


| Category | Craving (0-100) | Valence (1-9) | Arousal (1-9) | Typicality (0-100) | Relatedness | | | | | HSV | | |
|---|---|---|---|---|---|---|---|---|---|---|---|---|
| | | | | | Meth | Opioid | Both | Neither | MethToOpioid | Hue | Saturation | Value |
| Opioid | 85.78 (17.75) | 4.59 (1.866) | 4.41 (2.82) | 87.67 (17.27) | 0 | 0.963 | 0 | 0.037 | 0.963 | 0.305 (0.285) | 0.379 (0.35) | 0.26 (0.328) |

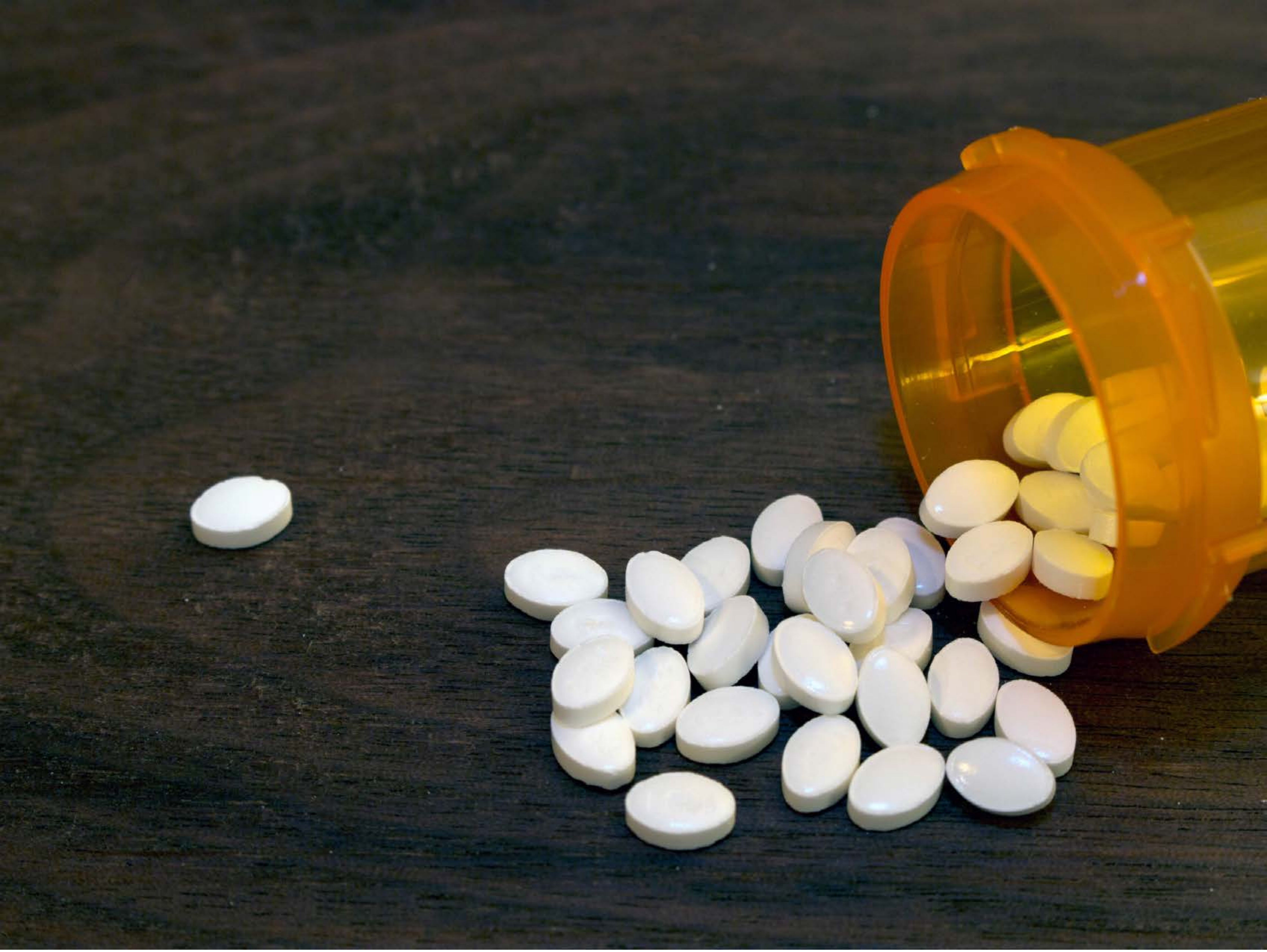

| Category | Craving (0-100) | Valence (1-9) | Arousal (1-9) | Typicality (0-100) | Relatedness | | | | | HSV | | |
|---|---|---|---|---|---|---|---|---|---|---|---|---|
| | | | | | Meth | Opioid | Both | Neither | MethToOpioid | Hue | Saturation | Value |
| Opioid | 82.96 (18.45) | 4.85 (1.657) | 4.15 (2.74) | 84.56 (18.46) | 0 | 0.926 | 0.037 | 0.037 | 0.926 | 0.253 (0.21) | 0.317 (0.305) | 0.313 (0.253) |

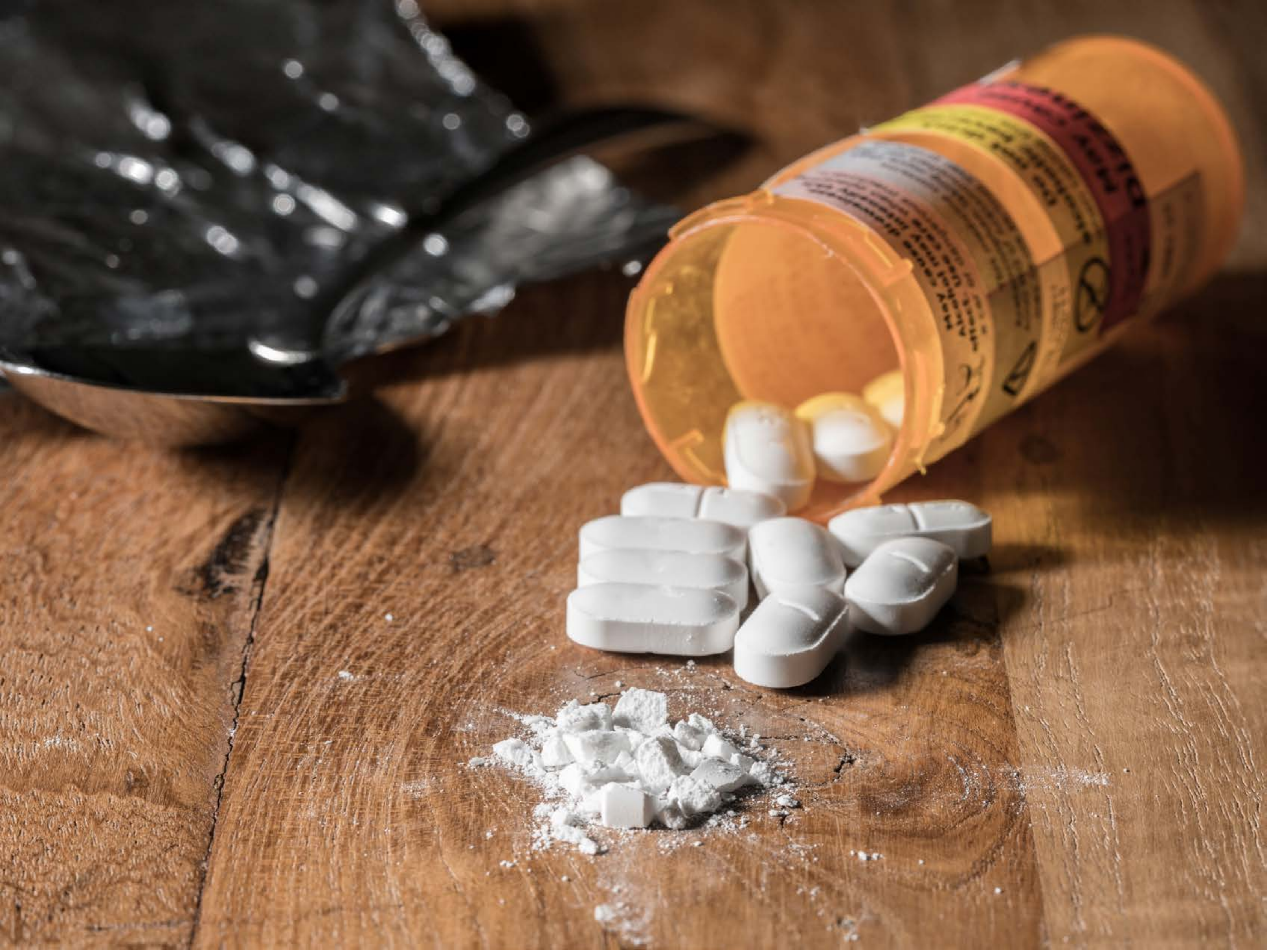

| Category | Craving (0-100) | Valence (1-9) | Arousal (1-9) | Typicality (0-100) | Relatedness | | | | | HSV | | |
|---|---|---|---|---|---|---|---|---|---|---|---|---|
| | | | | | Meth | Opioid | Both | Neither | MethToOpioid | Hue | Saturation | Value |
| Opioid | 89.14 (14.61) | 4.89 (1.663) | 4.61 (2.41) | 89.86 (14.33) | 0 | 0.893 | 0.107 | 0 | 0.893 | 0.091 (0.125) | 0.439 (0.239) | 0.503 (0.255) |

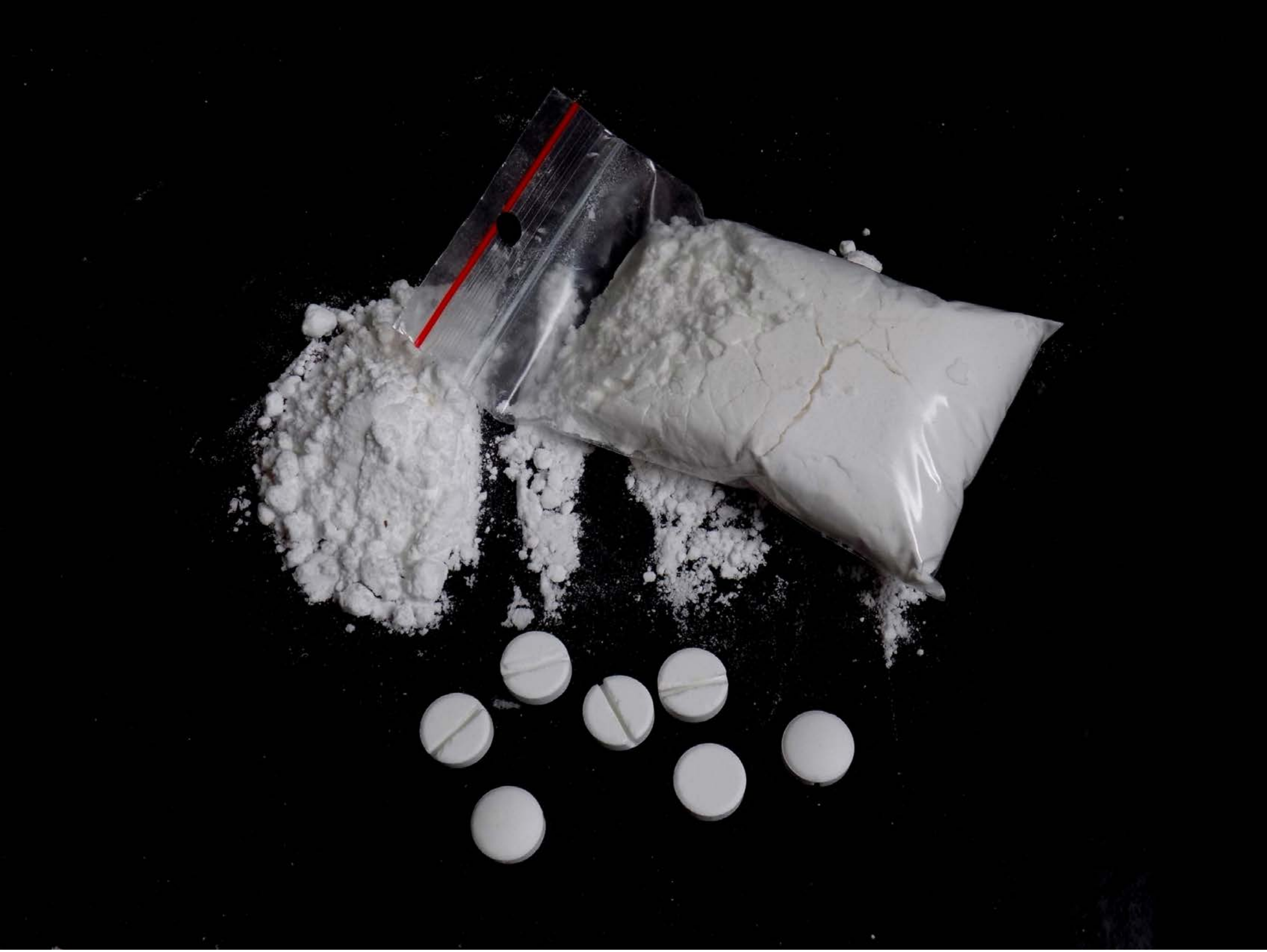

| Category | Craving (0-100) | Valence (1-9) | Arousal (1-9) | Typicality (0-100) | Relatedness | | | | | HSV | | |
|---|---|---|---|---|---|---|---|---|---|---|---|---|
| | | | | | Meth | Opioid | Both | Neither | MethToOpioid | Hue | Saturation | Value |
| Opioid | 85.93 (21.82) | 4.21 (1.969) | 4.46 (2.85) | 89.18 (22.23) | 0 | 0.464 | 0.536 | 0 | 0.464 | 0.166 (0.313) | 0.027 (0.119) | 0.125 (0.243) |

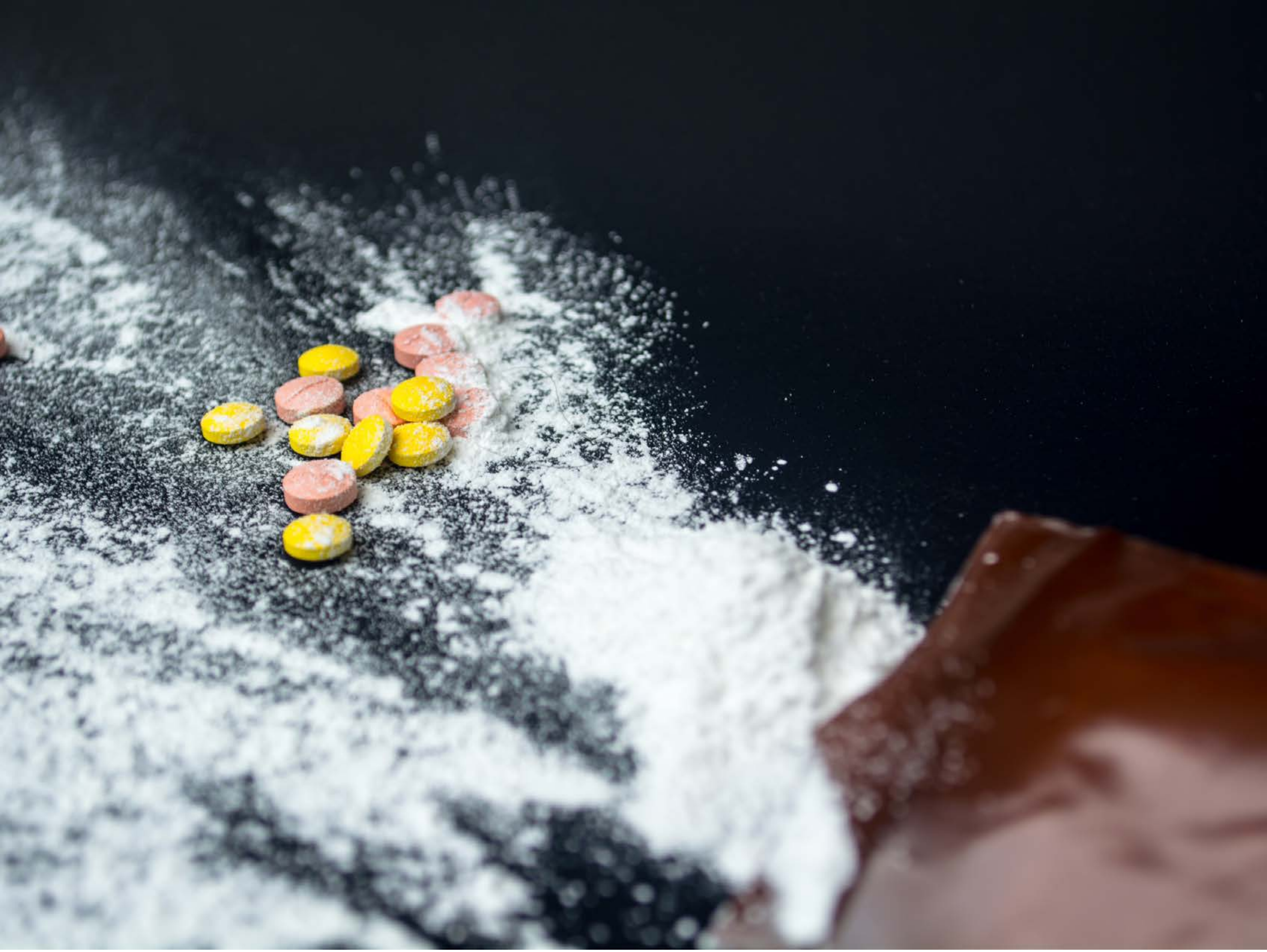

| Category | Craving (0-100) | Valence (1-9) | Arousal (1-9) | Typicality (0-100) | Relatedness | | | | | HSV | | |
|---|---|---|---|---|---|---|---|---|---|---|---|---|
| | | | | | Meth | Opioid | Both | Neither | MethToOpioid | Hue | Saturation | Value |
| Opioid | 85.86 (22.18) | 4.82 (1.806) | 4.36 (2.77) | 86.68 (23.16) | 0 | 0.571 | 0.393 | 0.036 | 0.571 | 0.502 (0.205) | 0.34 (0.273) | 0.433 (0.325) |

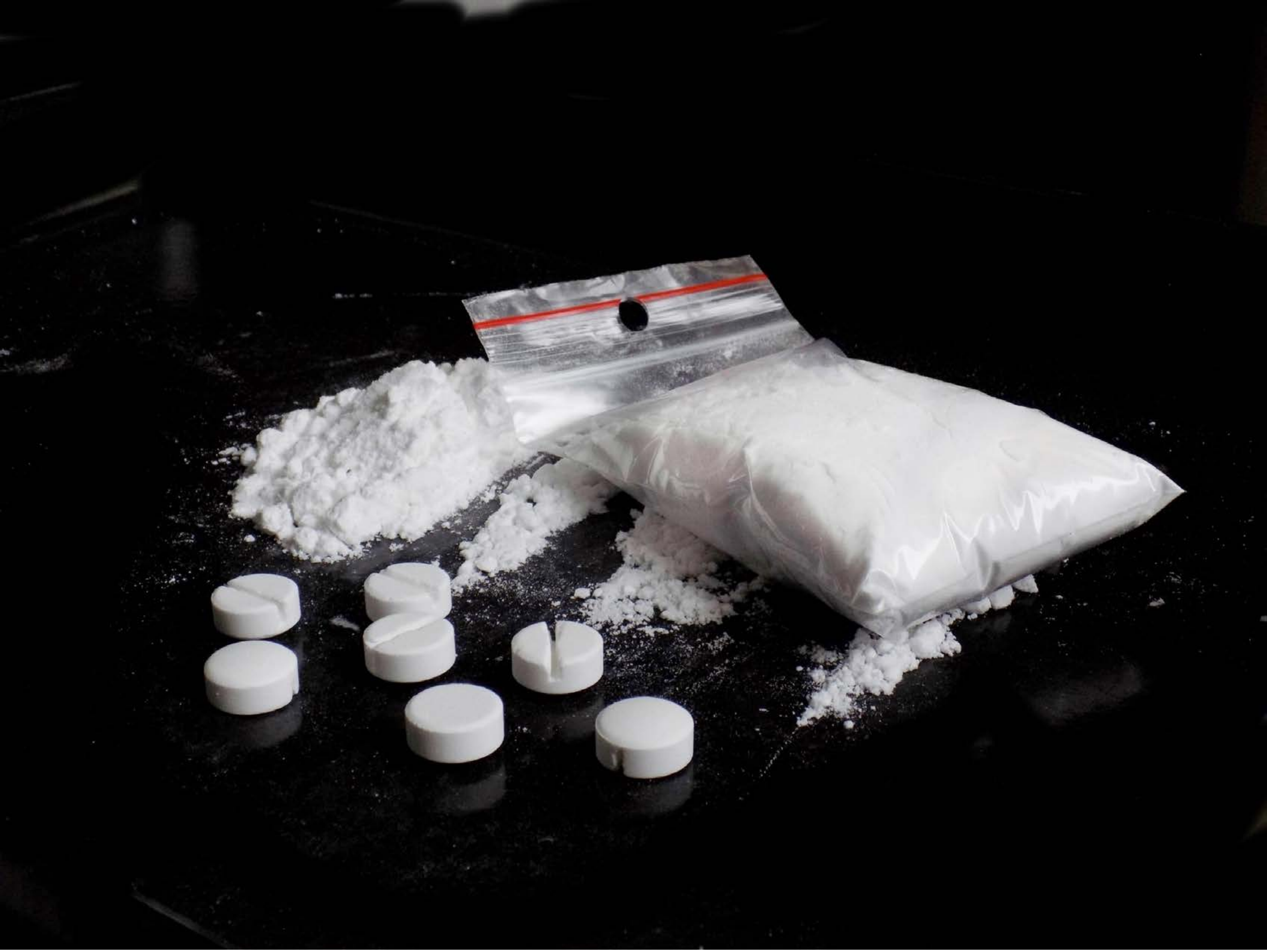

| Category | Craving (0-100) | Valence (1-9) | Arousal (1-9) | Typicality (0-100) | Relatedness | | | | | HSV | | |
|---|---|---|---|---|---|---|---|---|---|---|---|---|
| | | | | | Meth | Opioid | Both | Neither | MethToOpioid | Hue | Saturation | Value |
| Opioid | 89.46 (15.02) | 4.46 (1.972) | 4.57 (2.95) | 90.11 (15.05) | 0 | 0.536 | 0.464 | 0 | 0.536 | 0.171 (0.321) | 0.043 (0.151) | 0.157 (0.293) |

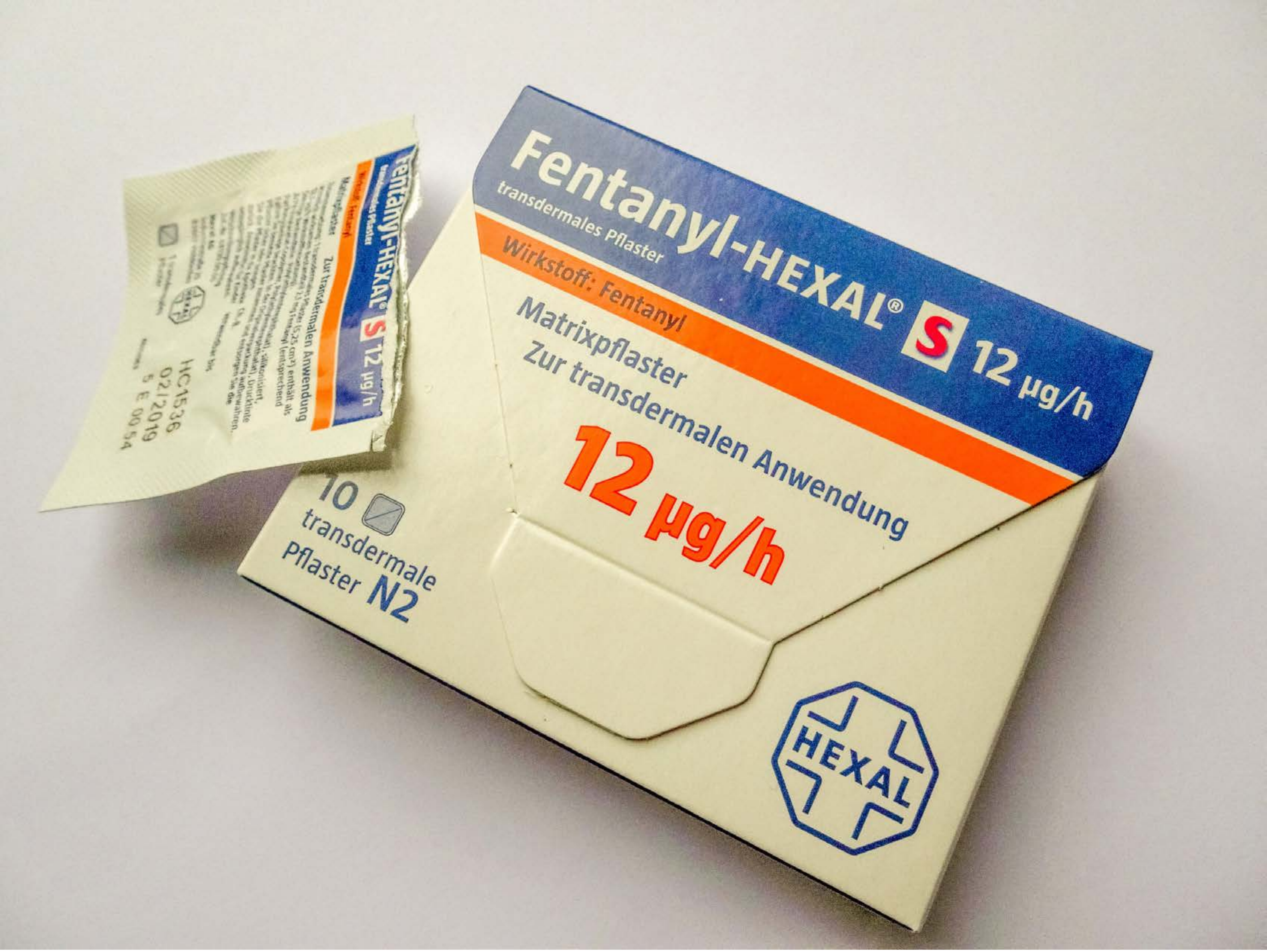


| Category | Craving (0-100) | Valence (1-9) | Arousal (1-9) | Typicality (0-100) | Relatedness | | | | | HSV | | |
|---|---|---|---|---|---|---|---|---|---|---|---|---|
| | | | | | Meth | Opioid | Both | Neither | MethToOpioid | Hue | Saturation | Value |
| Opioid | 86.15 (17.01) | 4.59 (1.6) | 4.37 (2.66) | 84.7 (17.2) | 0 | 0.926 | 0.074 | 0 | 0.926 | 0.153 (0.148) | 0.175 (0.215) | 0.795 (0.129) |

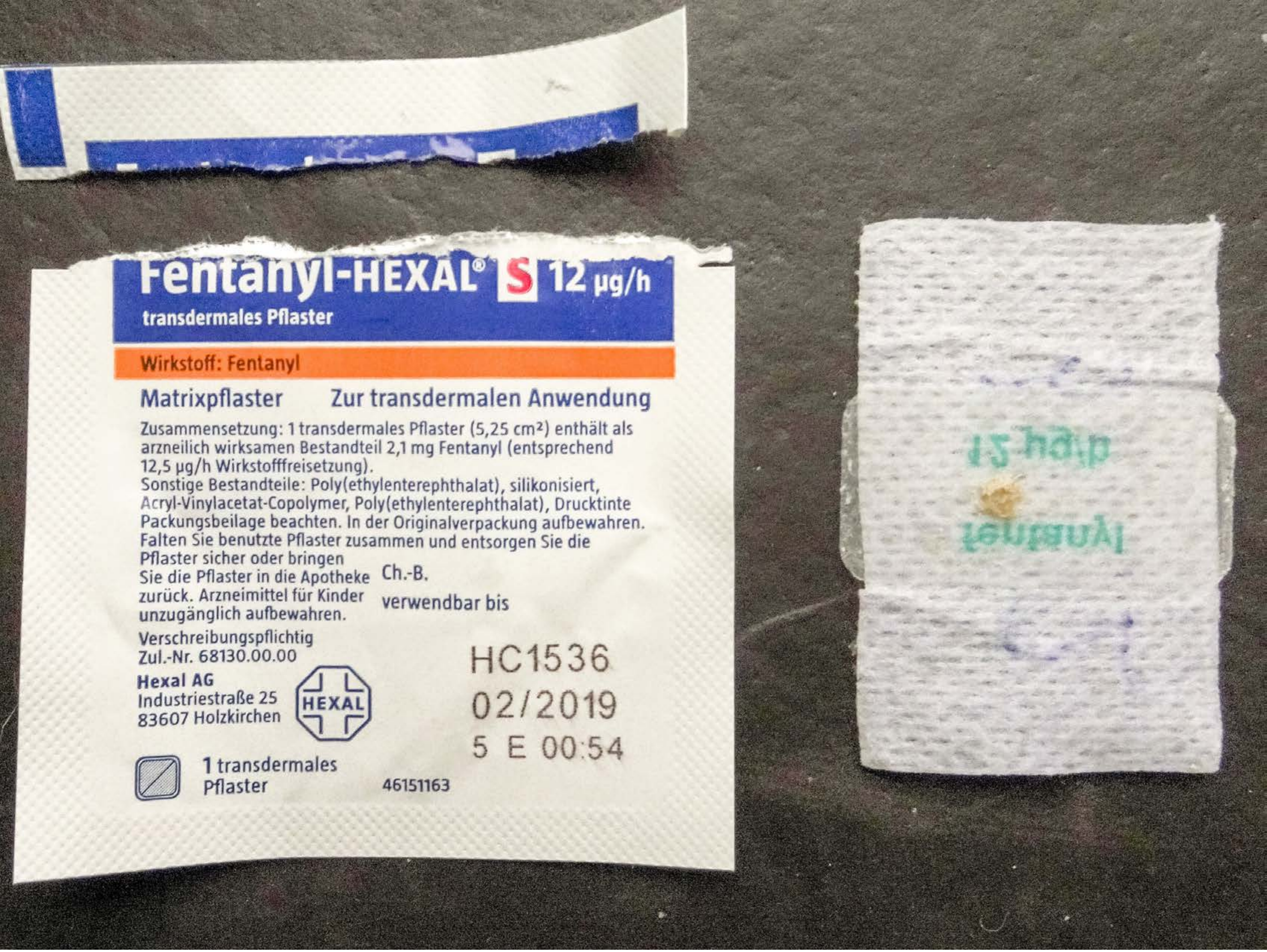


| Category | Craving (0-100) | Valence (1-9) | Arousal (1-9) | Typicality (0-100) | Relatedness | | | | | HSV | | |
|---|---|---|---|---|---|---|---|---|---|---|---|---|
| | | | | | Meth | Opioid | Both | Neither | MethToOpioid | Hue | Saturation | Value |
| Opioid | 84.85 (19.45) | 4.59 (2.024) | 4.37 (2.82) | 83.96 (17.78) | 0 | 0.926 | 0.074 | 0 | 0.926 | 0.181 (0.198) | 0.143 (0.155) | 0.651 (0.288) |

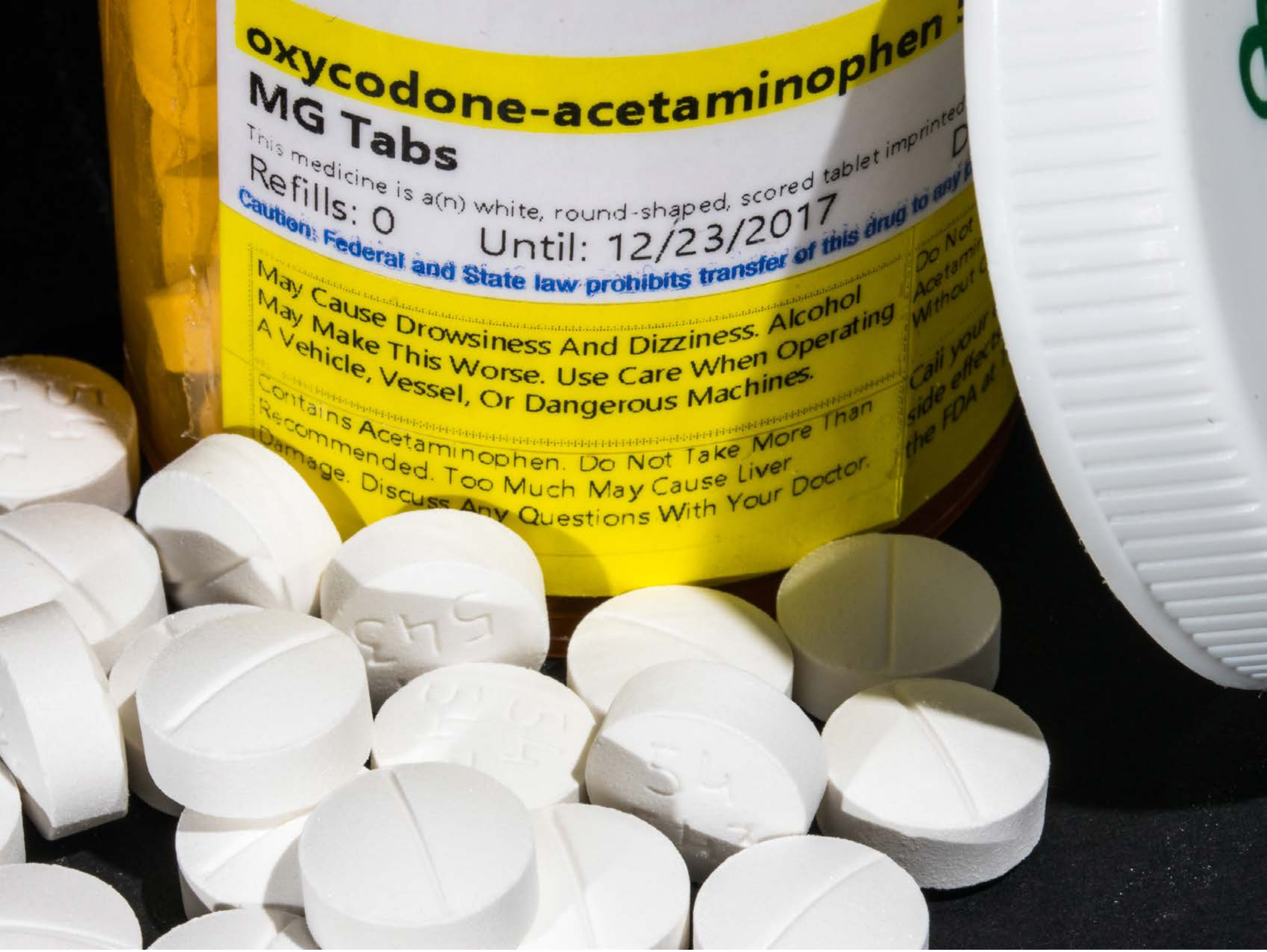


| Category | Craving (0-100) | Valence (1-9) | Arousal (1-9) | Typicality (0-100) | Relatedness | | | | | HSV | | |
|---|---|---|---|---|---|---|---|---|---|---|---|---|
| | | | | | Meth | Opioid | Both | Neither | MethToOpioid | Hue | Saturation | Value |
| Opioid | 89.37 (14.85) | 4.7 (1.898) | 4.78 (2.42) | 90.78 (13.58) | 0 | 0.963 | 0.037 | 0 | 0.963 | 0.277 (0.272) | 0.312 (0.408) | 0.67 (0.309) |

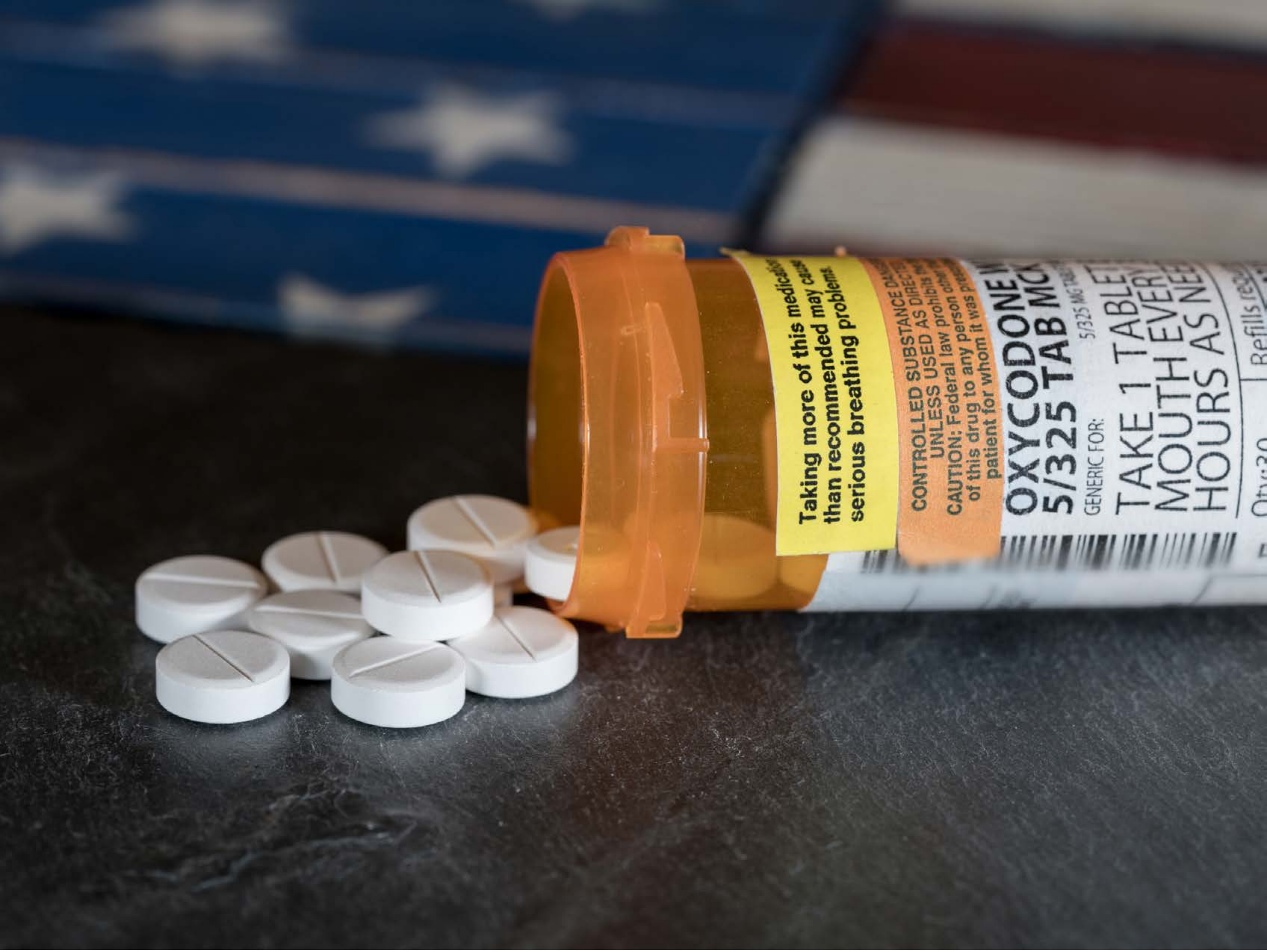


| Category | Craving (0-100) | Valence (1-9) | Arousal (1-9) | Typicality (0-100) | Relatedness | | | | | HSV | | |
|---|---|---|---|---|---|---|---|---|---|---|---|---|
| | | | | | Meth | Opioid | Both | Neither | MethToOpioid | Hue | Saturation | Value |
| Opioid | 84.75 (18.4) | 4.82 (1.744) | 4.61 (2.79) | 88.89 (15.72) | 0 | 1 | 0 | 0 | 1 | 0.341 (0.285) | 0.317 (0.288) | 0.375 (0.251) |

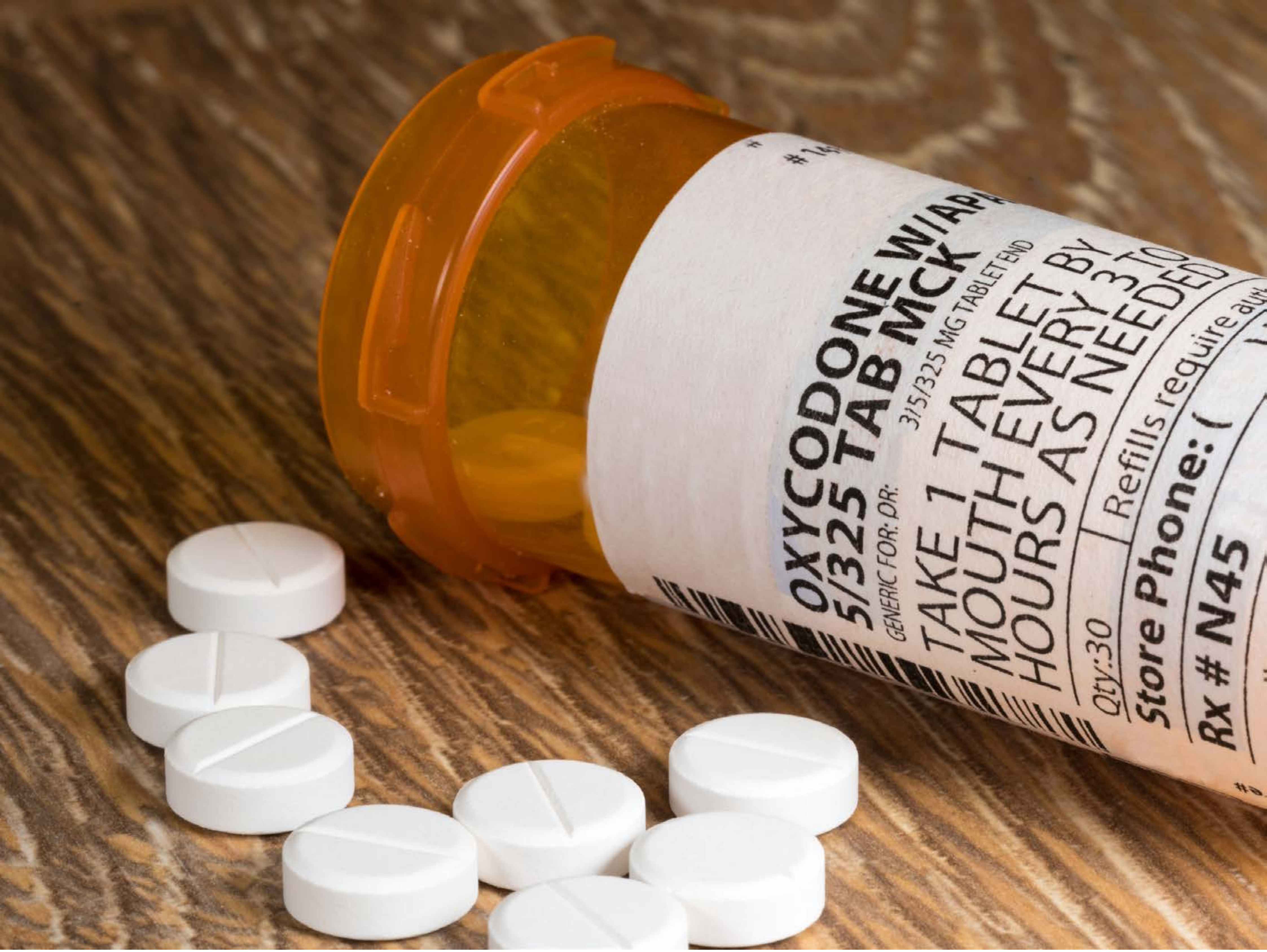


| Category | Craving (0-100) | Valence (1-9) | Arousal (1-9) | Typicality (0-100) | Relatedness | | | | | HSV | | |
|---|---|---|---|---|---|---|---|---|---|---|---|---|
| | | | | | Meth | Opioid | Both | Neither | MethToOpioid | Hue | Saturation | Value |
| Opioid | 92.21 (11.42) | 4.43 (1.933) | 4.57 (2.77) | 94.25 (10.59) | | 1 | | | 1 | 0.087 (0.128) | 0.448 (0.33) | 0.59 (0.235) |

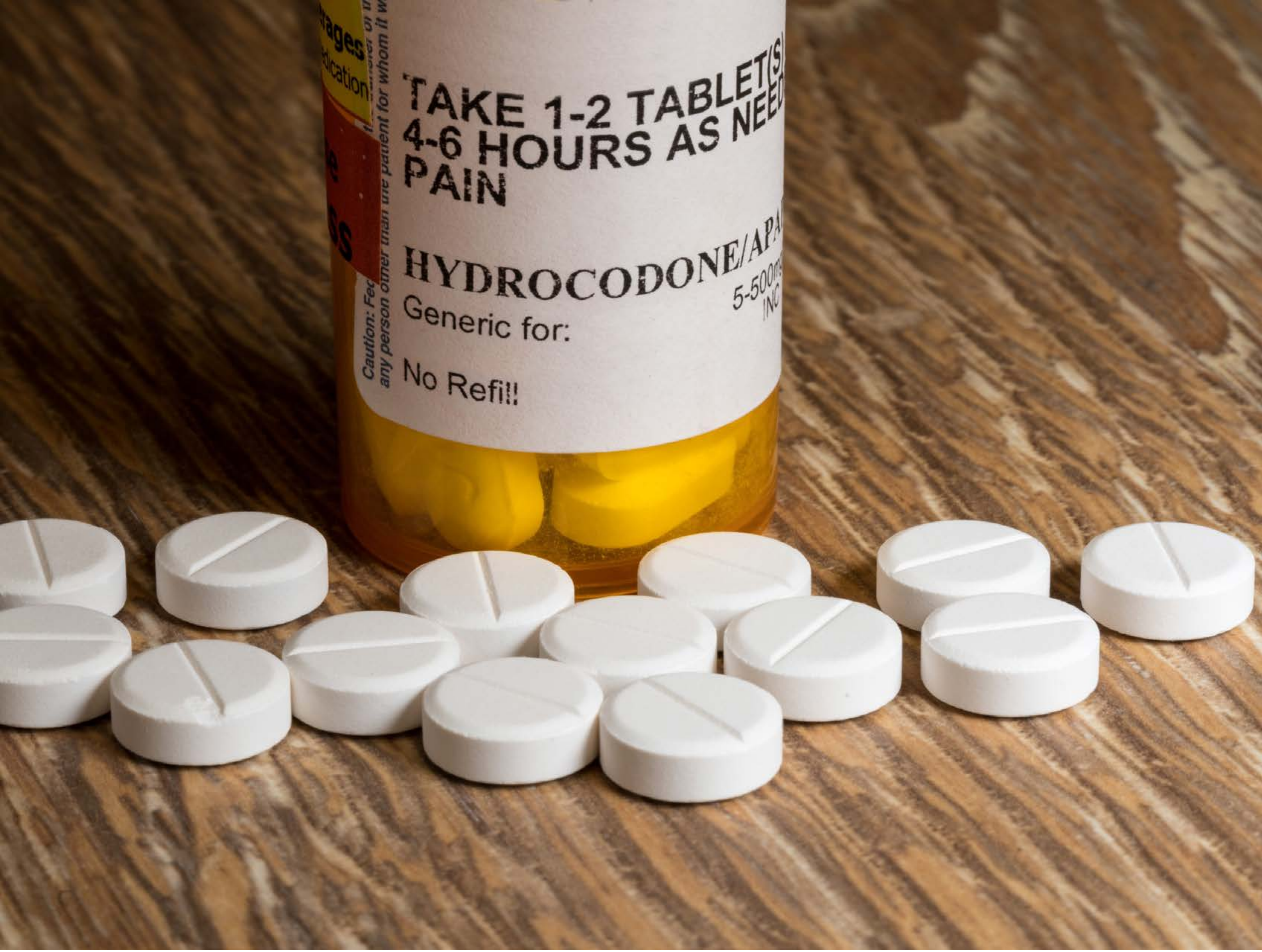


| Category | Craving (0-100) | Valence (1-9) | Arousal (1-9) | Typicality (0-100) | Relatedness | | | | | HSV | | |
|---|---|---|---|---|---|---|---|---|---|---|---|---|
| | | | | | Meth | Opioid | Both | Neither | MethToOpioid | Hue | Saturation | Value |
| Opioid | 83.3 (24.2) | 4.63 (1.779) | 4.48 (2.55) | 88.33 (17.85) | 0 | 0.963 | 0.037 | 0 | 0.963 | 0.079 (0.071) | 0.426 (0.291) | 0.561 (0.234) |

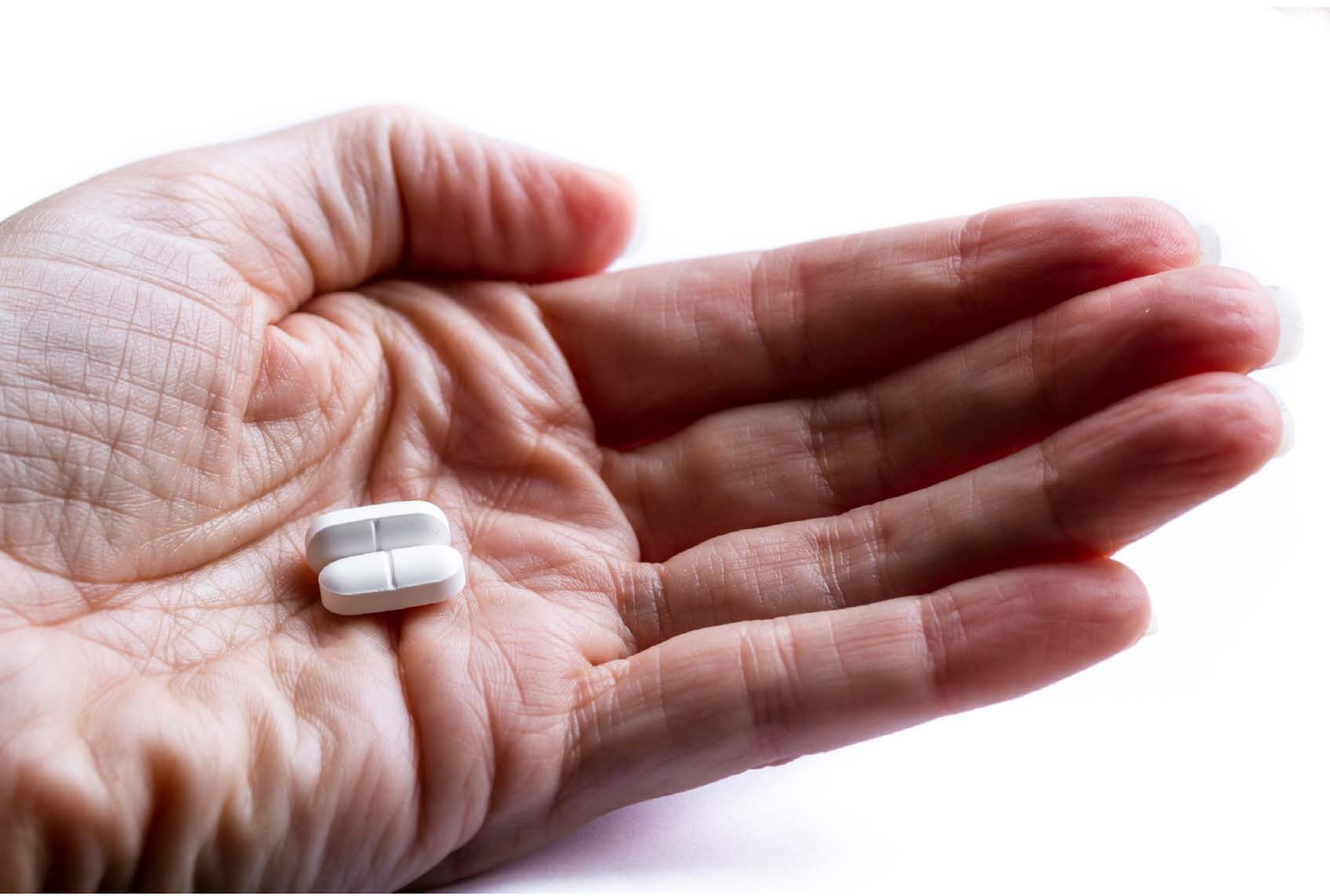

| Category | Craving (0-100) | Valence (1-9) | Arousal (1-9) | Typicality (0-100) | Relatedness | | | | | HSV | | |
|---|---|---|---|---|---|---|---|---|---|---|---|---|
| | | | | | Meth | Opioid | Both | Neither | MethToOpioid | Hue | Saturation | Value |
| Opioid Hand | 84.75 (18.32) | 4.82 (1.906) | 4.64 (2.72) | 88.75 (17.04) | 0 | 0.964 | 0.036 | 0 | 0.964 | 0.241 (0.4) | 0.216 (0.268 | 0.789 (0.245) |

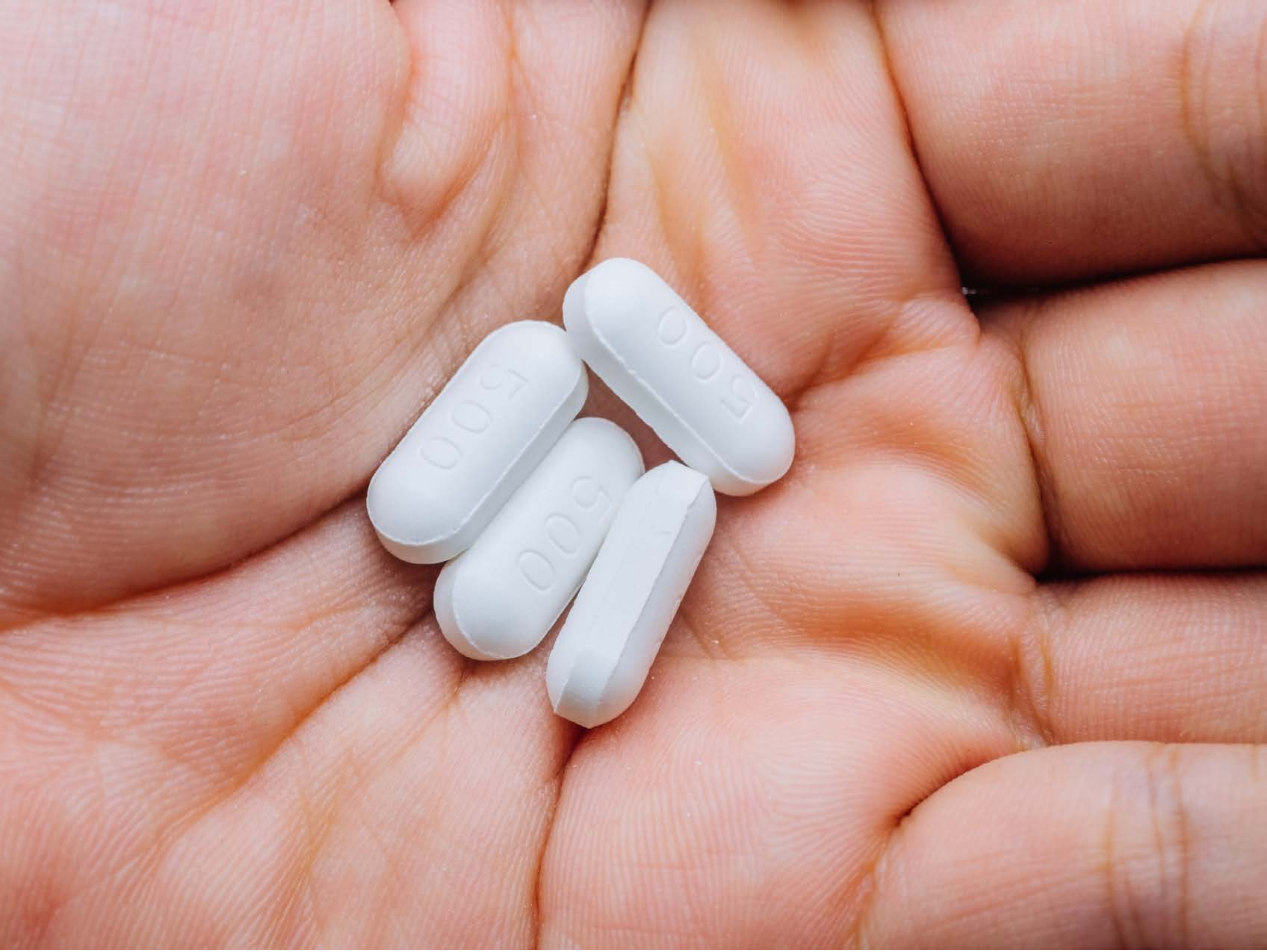


| Category | Craving (0-100) | Valence (1-9) | Arousal (1-9) | Typicality (0-100) | Relatedness | | | | | HSV | | |
|---|---|---|---|---|---|---|---|---|---|---|---|---|
| | | | | | Meth | Opioid | Both | Neither | MethToOpioid | Hue | Saturation | Value |
| Opioid Hand | 77.11 (27.45 | 4.85 (1.725 | 4.37 (2.79 | 74.74 (29.03) | 0 | 0.778 | 0.074 | 0.148 | 0.778 | 0.132 (0.263) | 0.321 (0.171) | 0.776 (0.125) |

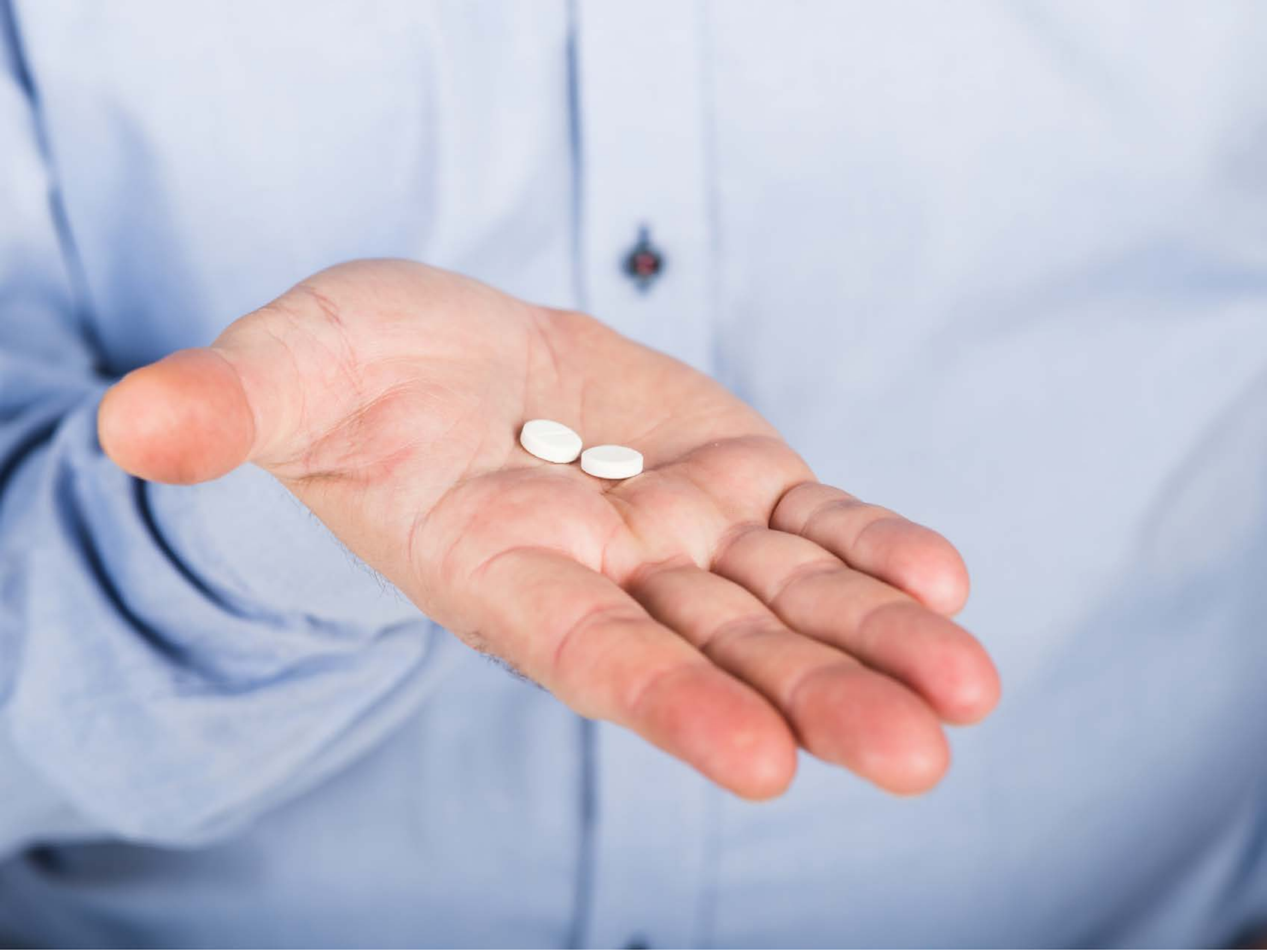

| Category | Craving (0-100) | Valence (1-9) | Arousal (1-9) | Typicality (0-100) | Relatedness | | | | | HSV | | |
|---|---|---|---|---|---|---|---|---|---|---|---|---|
| | | | | | Meth | Opioid | Both | Neither | MethToOpioid | Hue | Saturation | Value |
| Opioid Hand | 87.68 (16.4) | 4.93 (1.654) | 4.32 (2.71) | 86.21 (17.46) | 0 | 0.964 | 0.036 | 0 | 0.964 | 0.493 (0.236) | 0.2 (0.116) | 0.853 (0.102) |

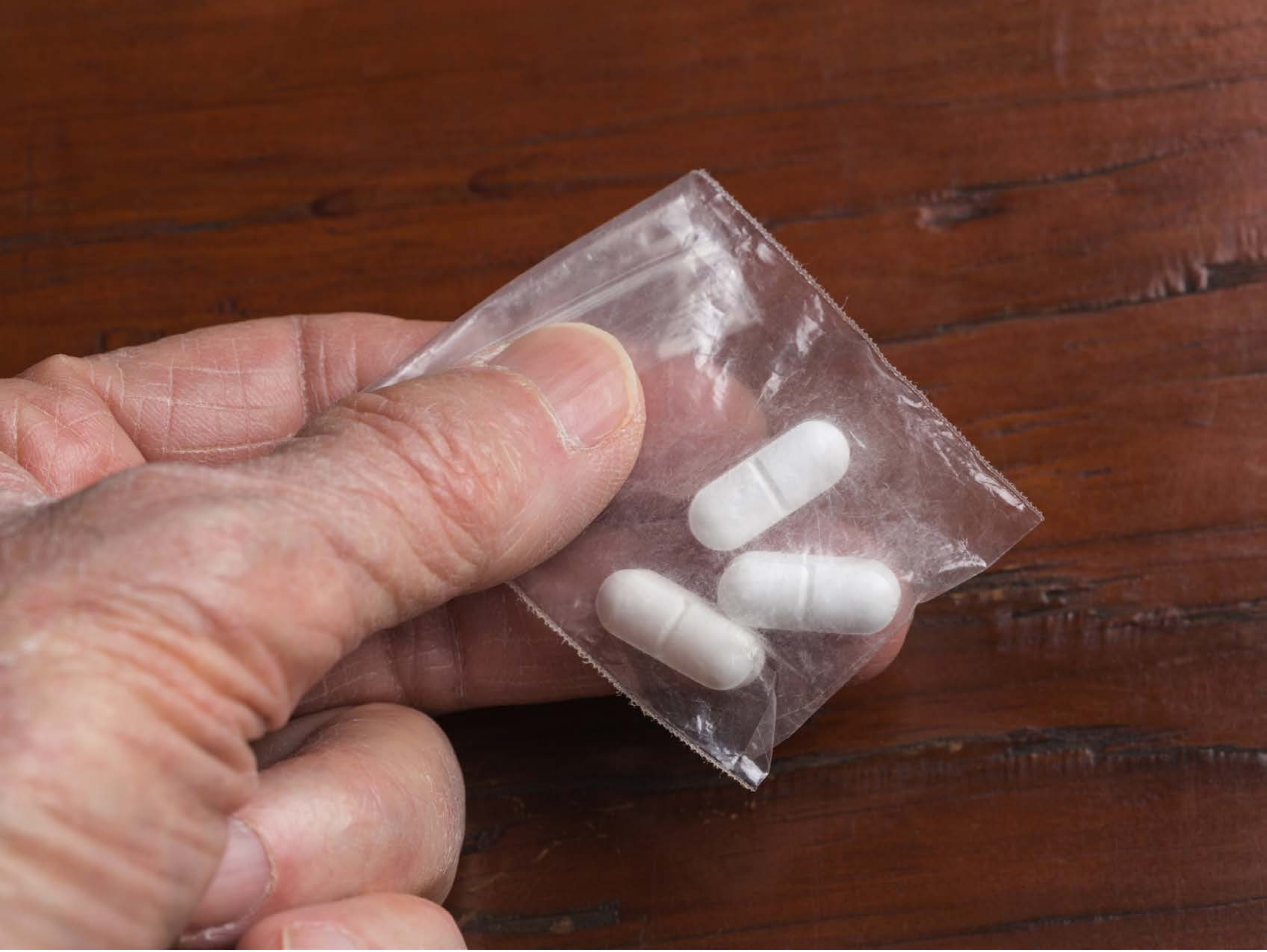

| Category | Craving (0-100) | Valence (1-9) | Arousal (1-9) | Typicality (0-100) | Relatedness | | | | | HSV | | |
|---|---|---|---|---|---|---|---|---|---|---|---|---|
| | | | | | Meth | Opioid | Both | Neither | MethToOpioid | Hue | Saturation | Value |
| Opioid Hand | 87.3 (16.87) | 4.7 (1.772) | 4.52 (2.76) | 88.26 (16.19) | 0 | 0.963 | 0.037 | 0 | 0.963 | 0.108 (0.249) | 0.503 (0.223) | 0.467 (0.207) |

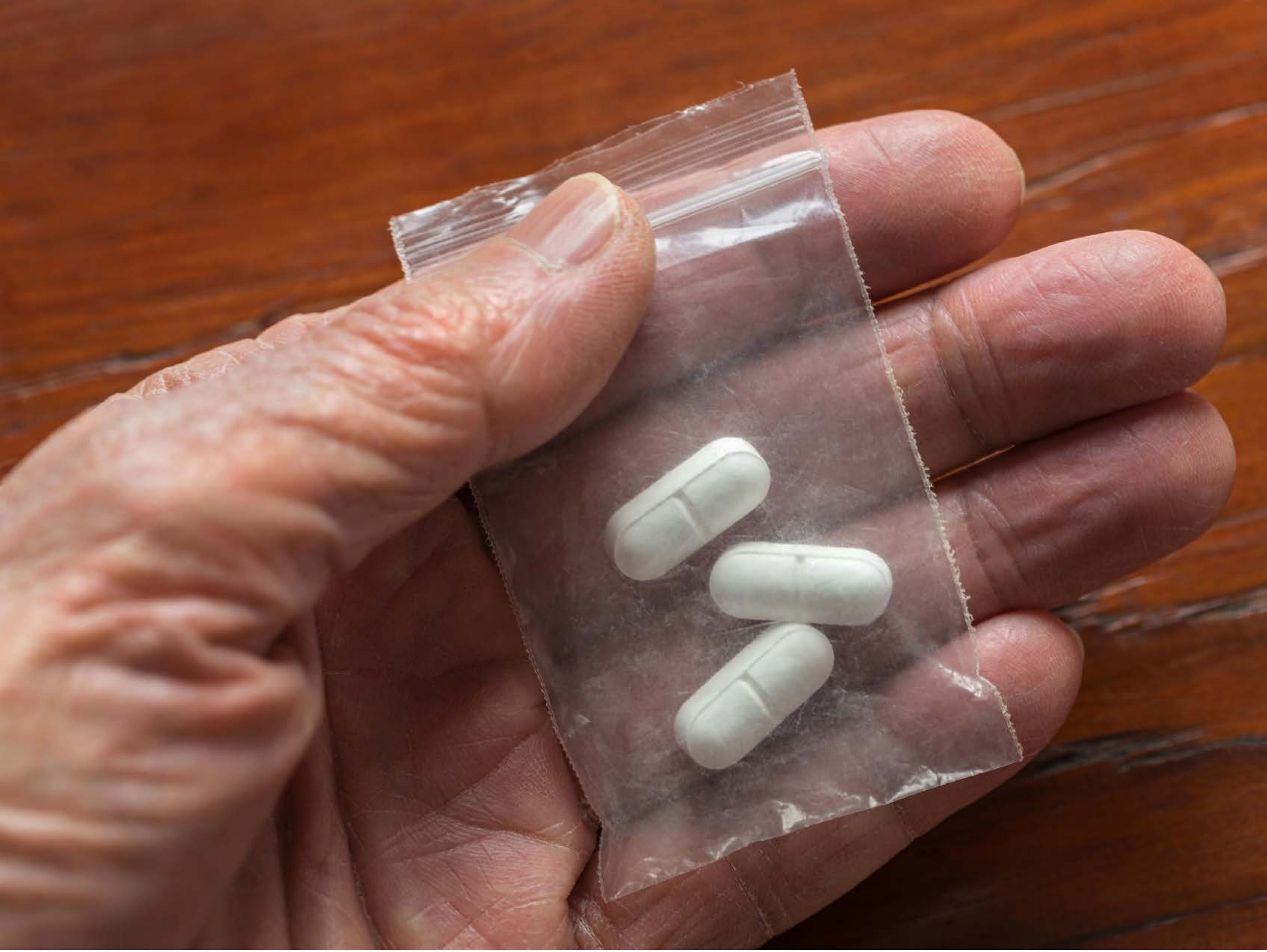

| Category | Craving (0-100) | Valence (1-9) | Arousal (1-9) | Typicality (0-100) | Relatedness | | | | | HSV | | |
|---|---|---|---|---|---|---|---|---|---|---|---|---|
| | | | | | Meth | Opioid | Both | Neither | MethToOpioid | Hue | Saturation | Value |
| Opioid Hand | 82.22 (19.35) | 4.52 (1.718) | 4.59 (2.61) | 84.63 (18.77) | 0 | 0.963 | 0 | 0.037 | 0.963 | 0.061 (0.135) | 0.51 (0.24) | 0.504 (0.184) |

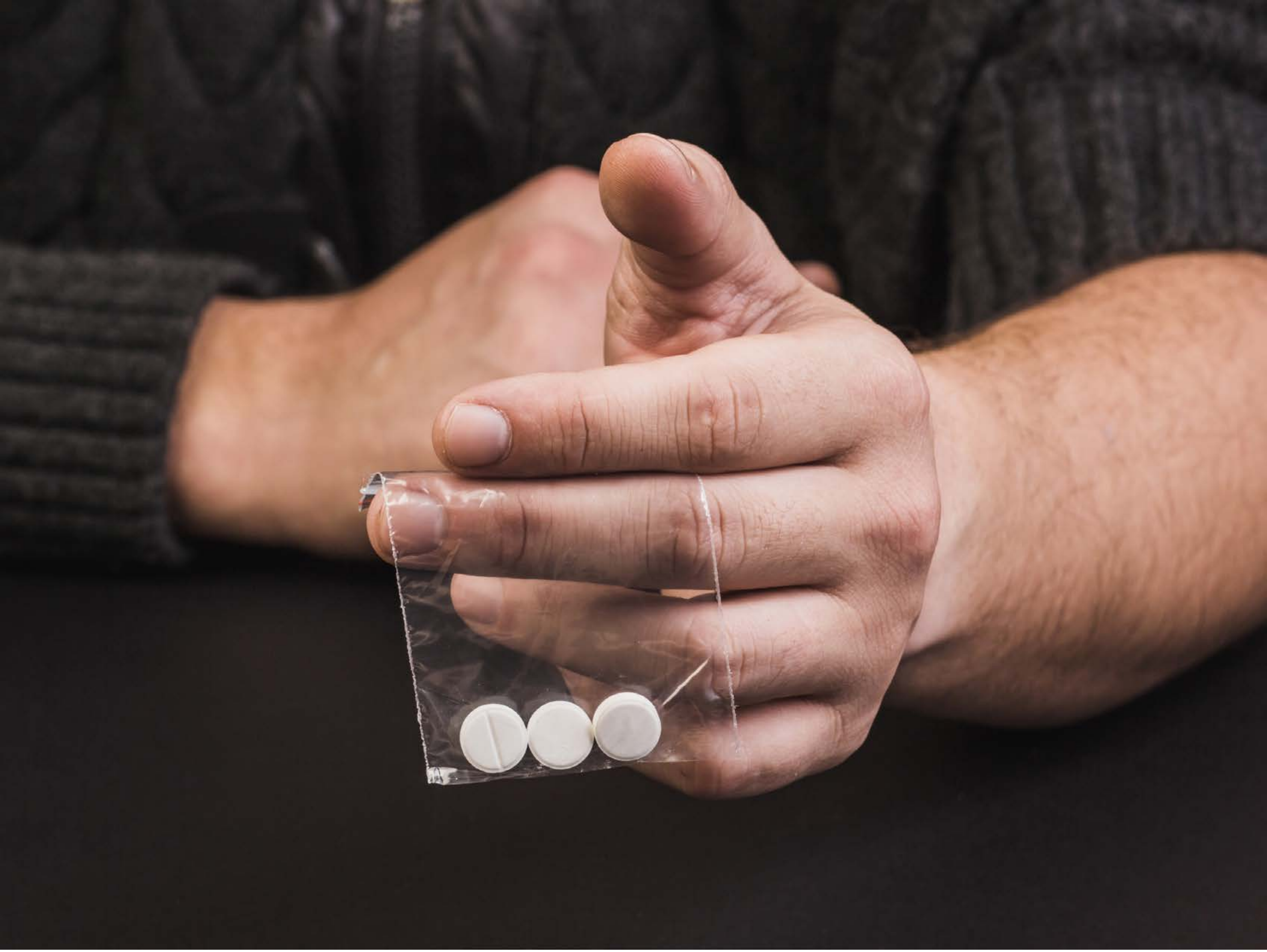

| Category | Craving (0-100) | Valence (1-9) | Arousal (1-9) | Typicality (0-100) | Relatedness | | | | | HSV | | |
|---|---|---|---|---|---|---|---|---|---|---|---|---|
| | | | | | Meth | Opioid | Both | Neither | MethToOpioid | Hue | Saturation | Value |
| Opioid Hand | 87.46 (14.46) | 4.79 (1.75) | 4.57 (2.7) | 91.82 (11.48) | 0.036 | 0.964 | 0 | 0 | 0.929 | 0.299 (0.405) | 0.214 (0.133) | 0.352 (0.297) |

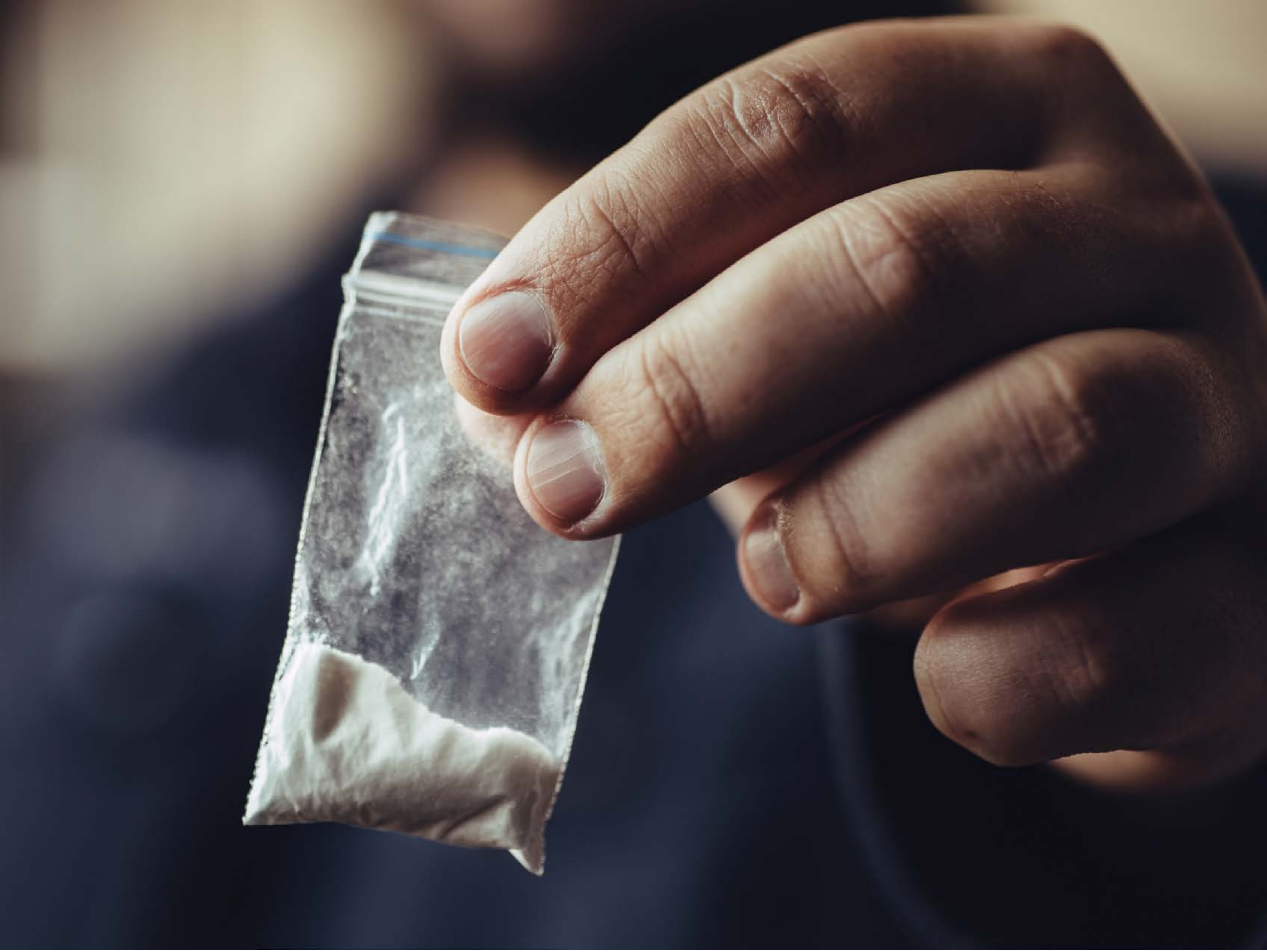

| Category | Craving (0-100) | Valence (1-9) | Arousal (1-9) | Typicality (0-100) | Relatedness | | | | | HSV | | |
|---|---|---|---|---|---|---|---|---|---|---|---|---|
| | | | | | Meth | Opioid | Both | Neither | MethToOpioid | Hue | Saturation | Value |
| Opioid Hand | 85.85 (17.04) | 4.41 (1.623) | 4.41 (2.61) | 85.85 (19.07) | 0.370 | 0.296 | 0.333 | 0 | -0.074 | 0.468 (0.328) | 0.275 (0.129) | 0.352 (0.235) |

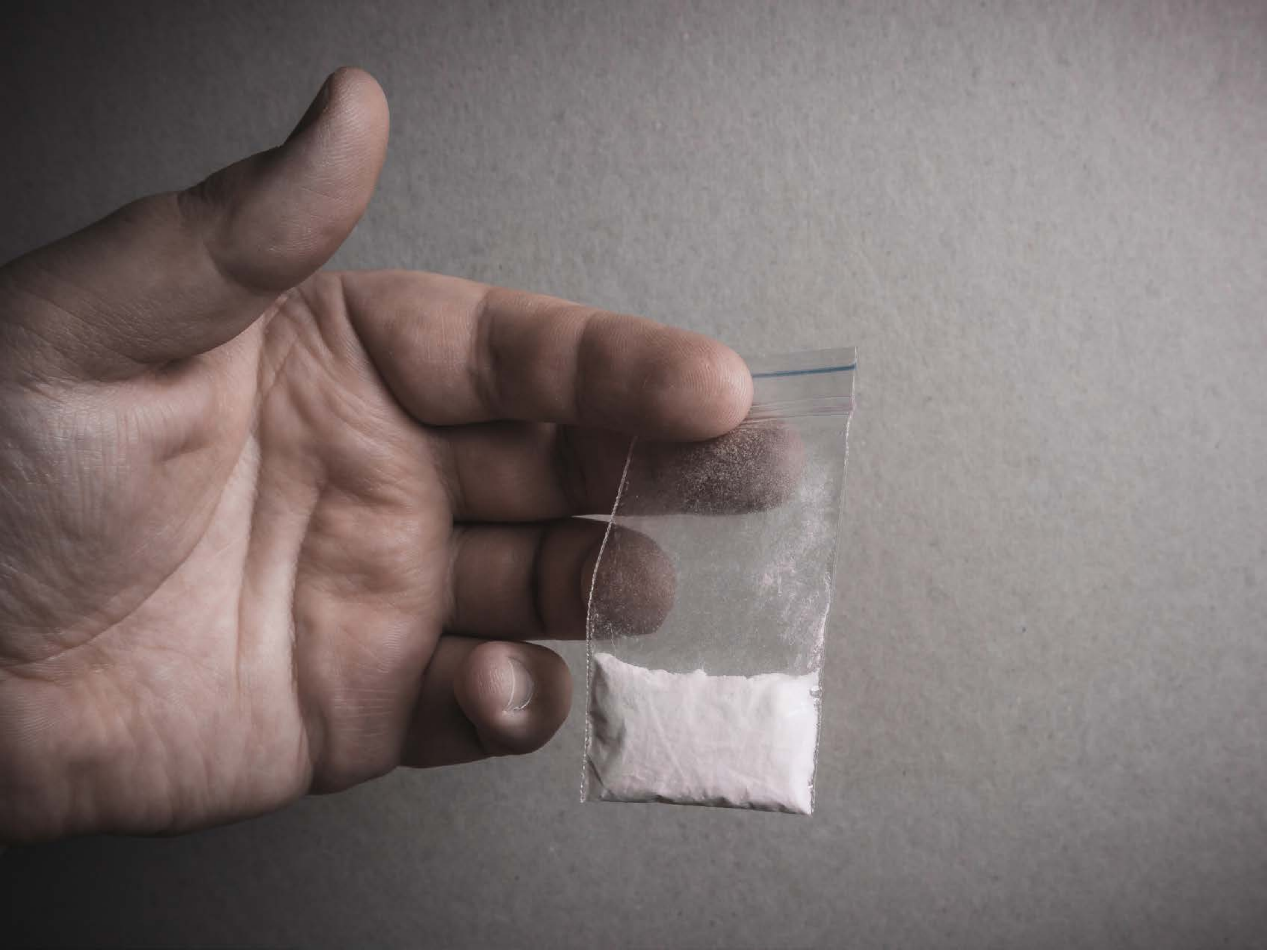

| Category | Craving (0-100) | Valence (1-9) | Arousal (1-9) | Typicality (0-100) | Relatedness | | | | | HSV | | |
|---|---|---|---|---|---|---|---|---|---|---|---|---|
| | | | | | Meth | Opioid | Both | Neither | MethToOpioid | Hue | Saturation | Value |
| Opioid Hand | 80.81 (21.16) | 4.44 (1.368) | 4.07 (2.37) | 83.07 (21.29) | 0.444 | 0.259 | 0.296 | 0 | -0.185 | 0.433 (0.456) | 0.106 (0.105) | 0.492 (0.168) |

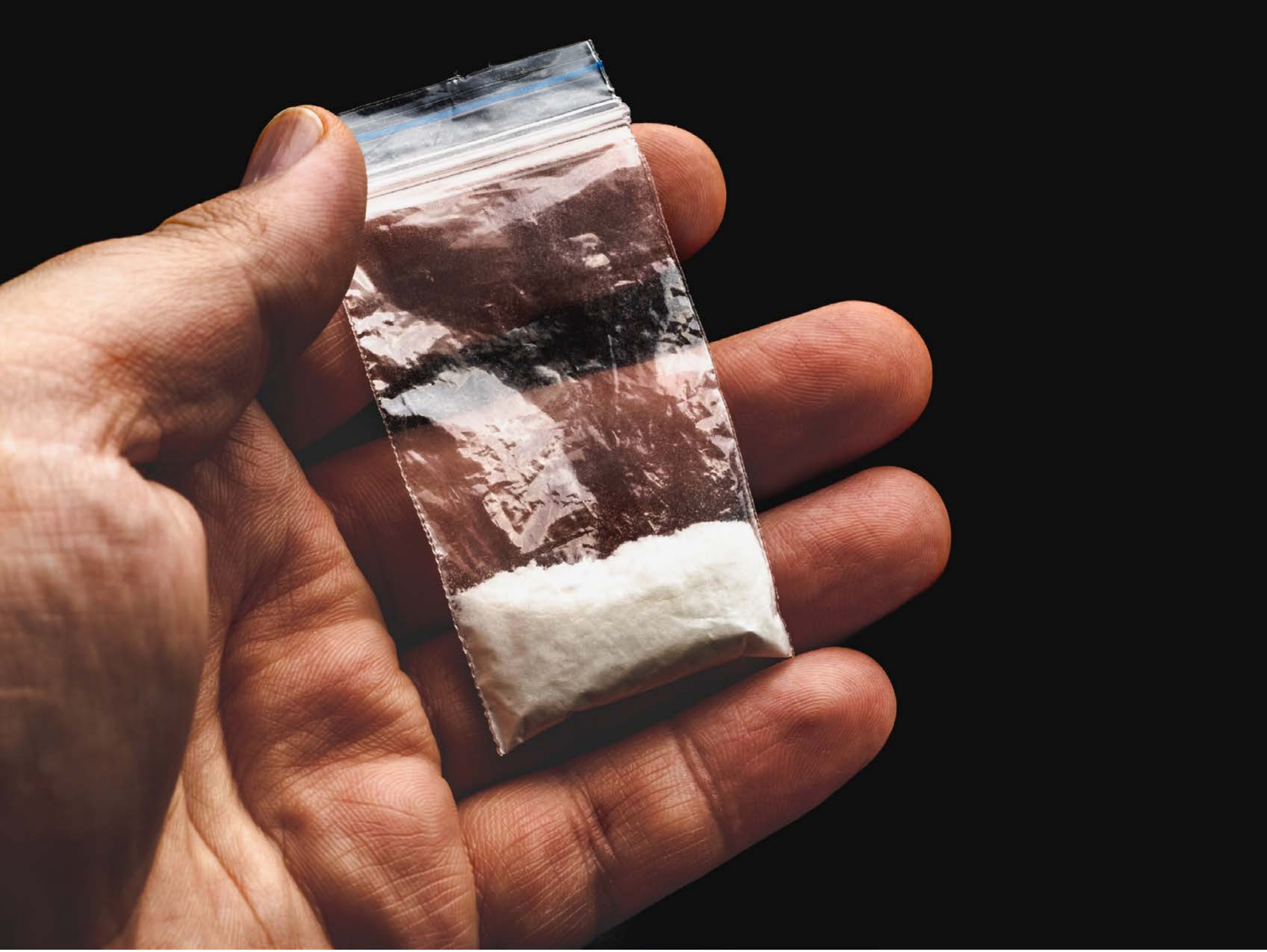

| Category | Craving (0-100) | Valence (1-9) | Arousal (1-9) | Typicality (0-100) | Relatedness | | | | | HSV | | |
|---|---|---|---|---|---|---|---|---|---|---|---|---|
| | | | | | Meth | Opioid | Both | Neither | MethToOpioid | Hue | Saturation | Value |
| Opioid Hand | 90.57 (16.41) | 4.71 (1.823) | 4.75 (2.78) | 91.04 (16.1) | 0.571 | 0.143 | 0.25 | 0.036 | -0.429 | 0.065 (0.194) | 0.213 (0.229) | 0.336 (0.312) |

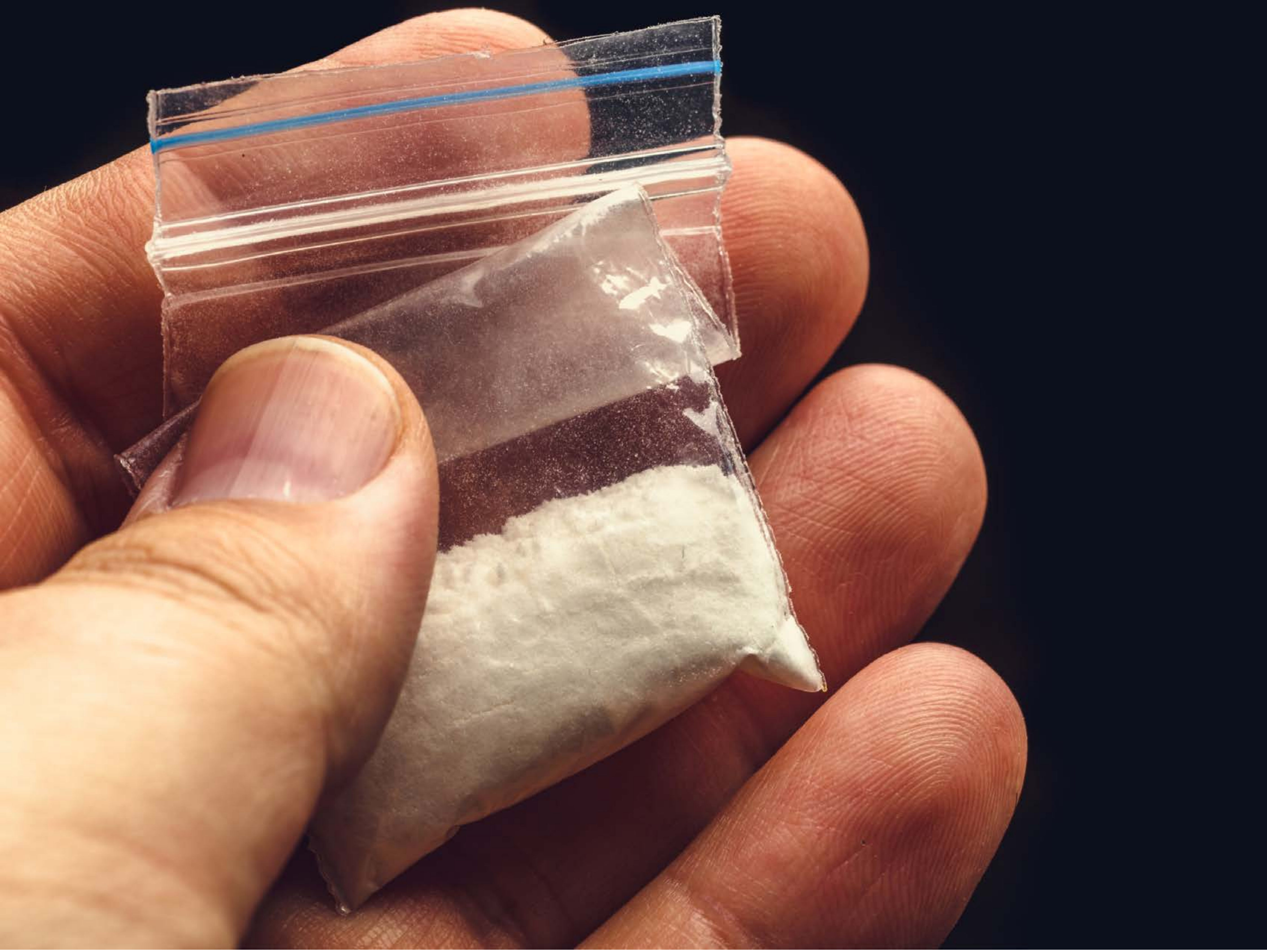

| Category | Craving (0-100) | Valence (1-9) | Arousal (1-9) | Typicality (0-100) | Relatedness | | | | | HSV | | |
|---|---|---|---|---|---|---|---|---|---|---|---|---|
| | | | | | Meth | Opioid | Both | Neither | MethToOpioid | Hue | Saturation | Value |
| Opioid Hand | 85.96 (21.98) | 4.89 (1.499) | 4.57 (2.78) | 89.25 (21.96) | 0.464 | 0.071 | 0.429 | 0.036 | -0.393 | 0.324 (0.333) | 0.468 (0.139) | 0.481 (0.327) |

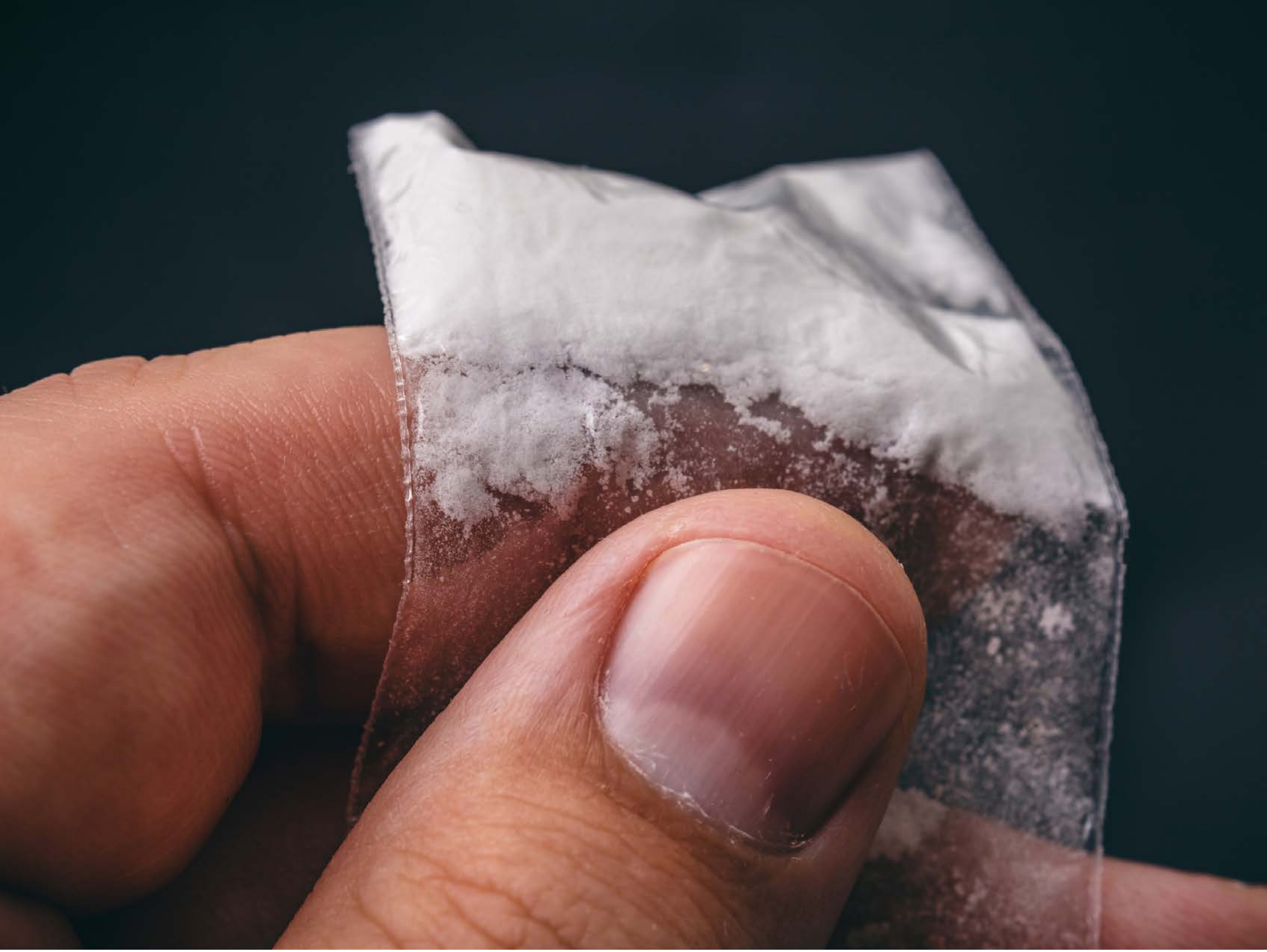

| Category | Craving (0-100) | Valence (1-9) | Arousal (1-9) | Typicality (0-100) | Relatedness | | | | | HSV | | |
|---|---|---|---|---|---|---|---|---|---|---|---|---|
| | | | | | Meth | Opioid | Both | Neither | MethToOpioid | Hue | Saturation | Value |
| Opioid Hand | 91.32 (12.72) | 4.64 (1.87) | 4.61 (2.69) | 94.14 (10.14) | 0.5 | 0.036 | 0.393 | 0.071 | -0.464 | 0.405 (0.353) | 0.392 (0.209) | 0.413 (0.264) |

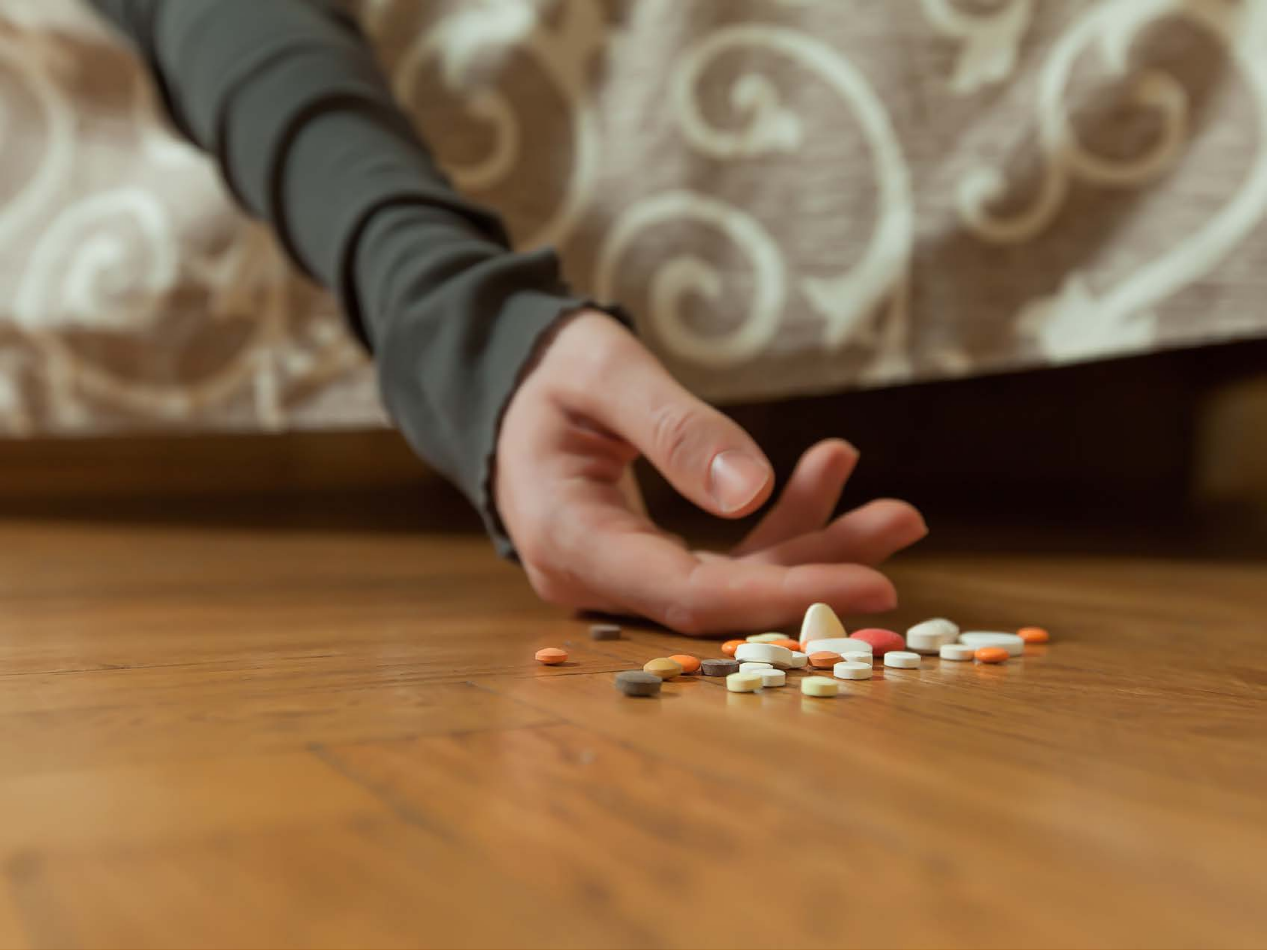

| Category | Craving (0-100) | Valence (1-9) | Arousal (1-9) | Typicality (0-100) | Relatedness | | | | | HSV | | |
|---|---|---|---|---|---|---|---|---|---|---|---|---|
| | | | | | Meth | Opioid | Both | Neither | MethToOpioid | Hue | Saturation | Value |
| Opioid Hand | 83.33 (21.79) | 4.63 (1.735) | 4.37 (2.32) | 85.48 (19.22) | 0 | 0.889 | 0.074 | 0.037 | 0.889 | 0.099 (0.053) | 0.483 (0.241) | 0.528 (0.198) |

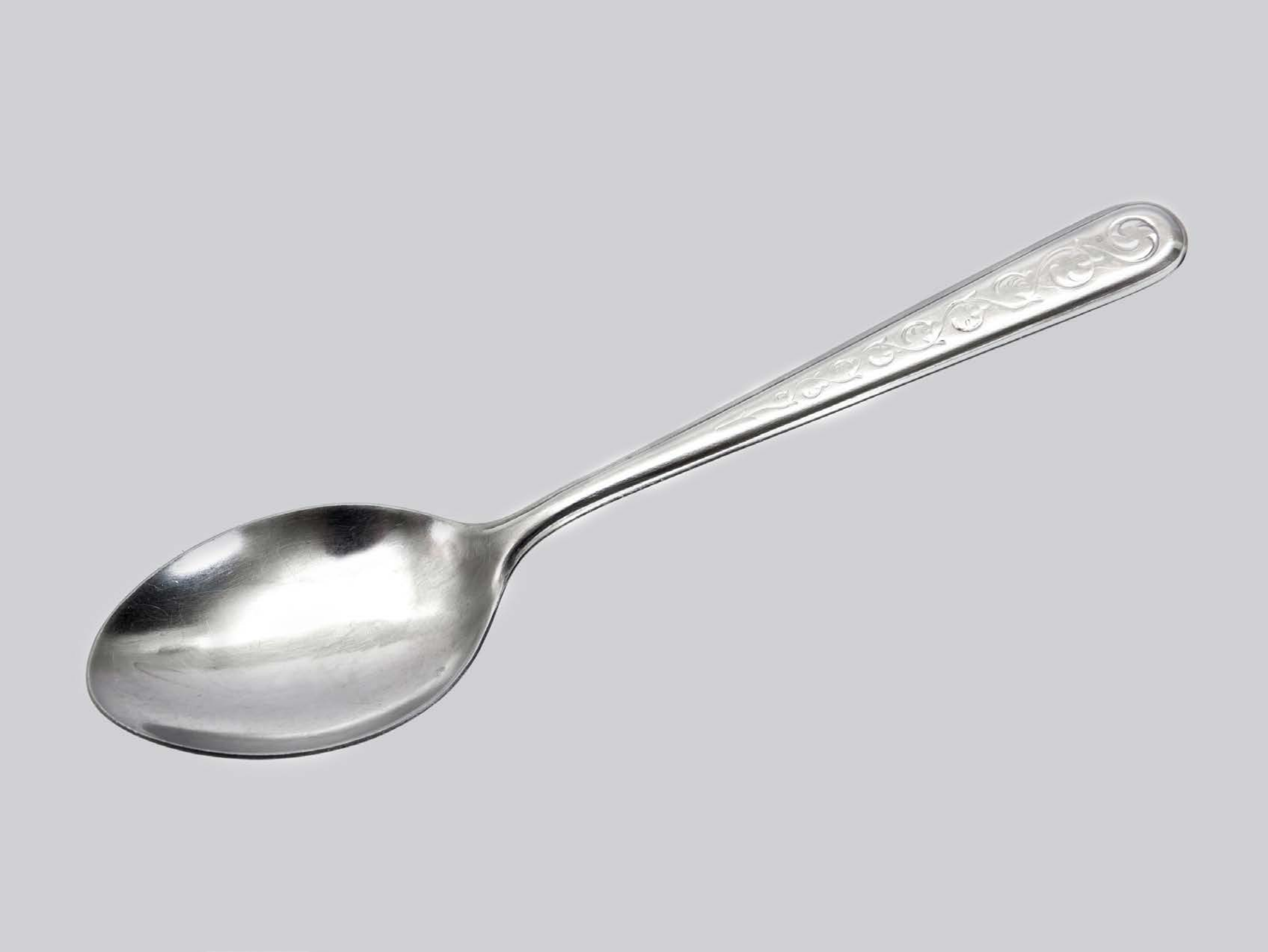

| Category | Craving (0-100) | Valence (1-9) | Arousal (1-9) | Typicality (0-100) | Relatedness | | | | | HSV | | |
|---|---|---|---|---|---|---|---|---|---|---|---|---|
| | | | | | Meth | Opioid | Both | Neither | MethToOpioid | Hue | Saturation | Value |
| Opioid Injection Instrument | 59.19 (31.64) | 4.85 (1.433) | 4.04 (2.26) | 75.96 (25.62) | 0.074 | 0.037 | 0.630 | 0.259 | -0.037 | 0.03 (0.135) | 0.002 (0.009) | 0.971 (0.112) |

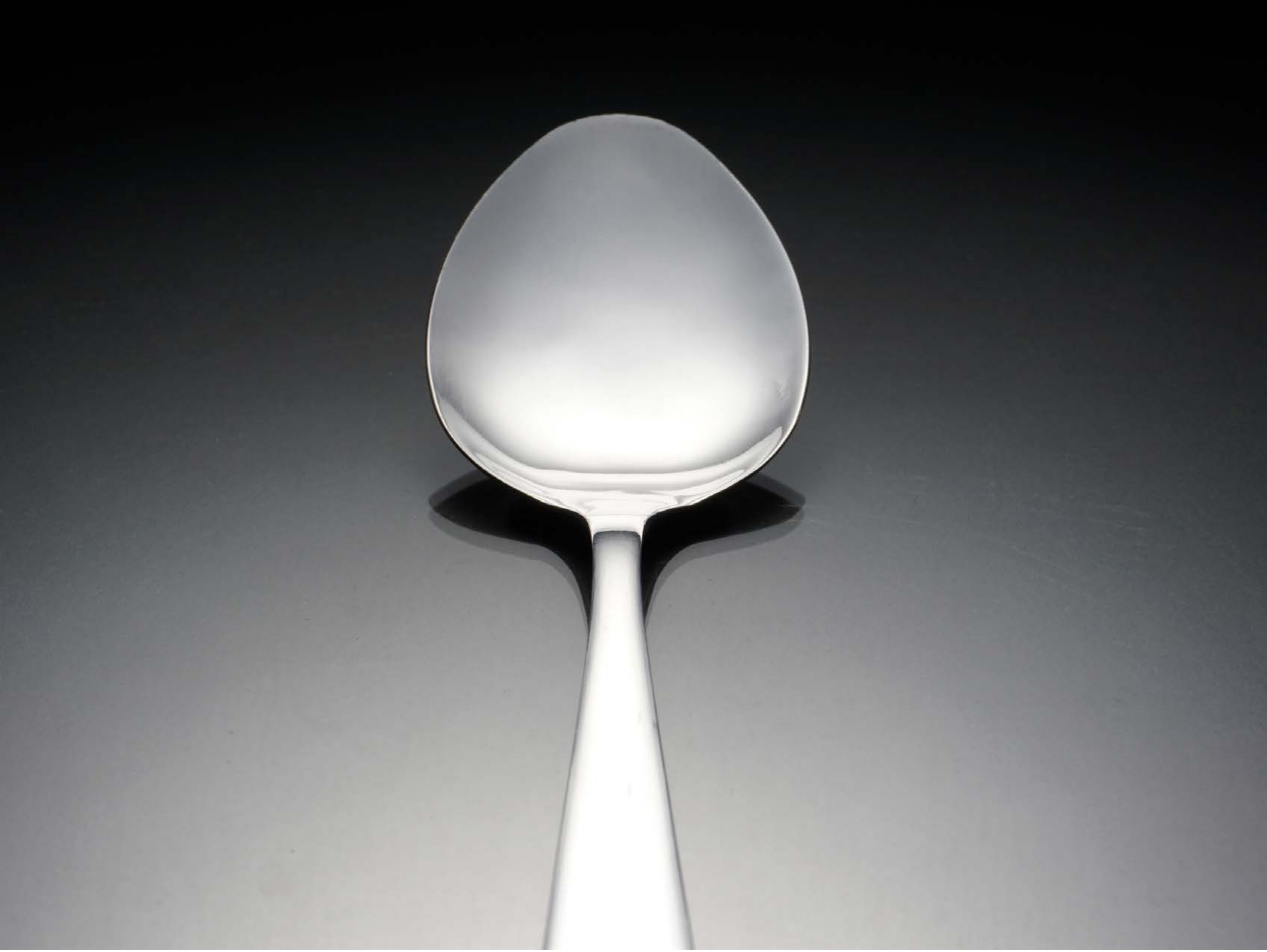

| Category | Craving (0-100) | Valence (1-9) | Arousal (1-9) | Typicality (0-100) | Relatedness | | | | | HSV | | |
|---|---|---|---|---|---|---|---|---|---|---|---|---|
| | | | | | Meth | Opioid | Both | Neither | MethToOpioid | Hue | Saturation | Value |
| Opioid Injection Instrument | 70.18 (26.98) | 4.57 (1.597) | 3.96 (2.63) | 79.36 (27.94) | 0.036 | 0 | 0.786 | 0.179 | -0.036 | 0.383 (0.284) | 0.062 (0.1) | 0.405 (0.289) |

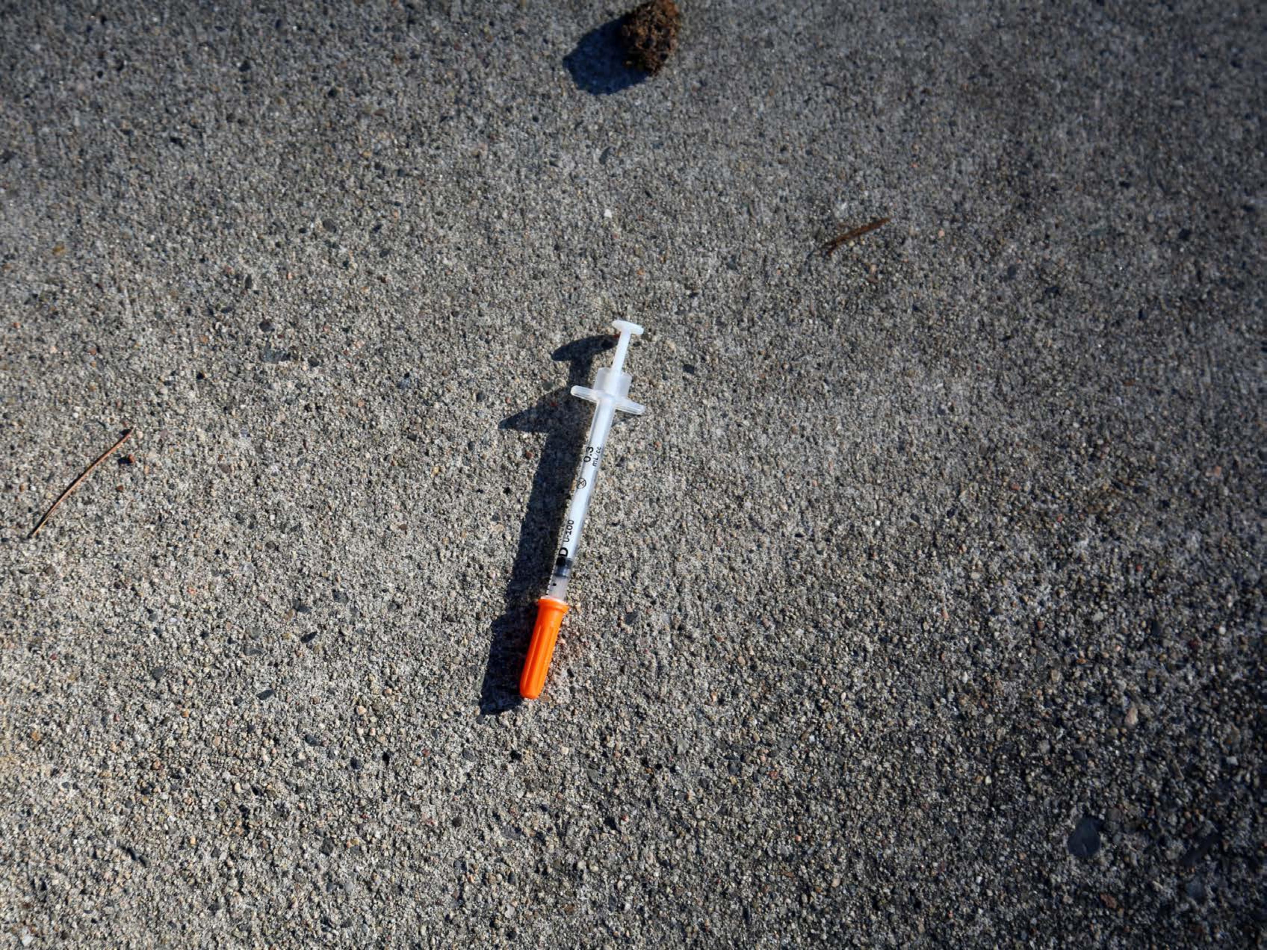

| Category | Craving (0-100) | Valence (1-9) | Arousal (1-9) | Typicality (0-100) | Relatedness | | | | | HSV | | |
|---|---|---|---|---|---|---|---|---|---|---|---|---|
| | | | | | Meth | Opioid | Both | Neither | MethToOpioid | Hue | Saturation | Value |
| Opioid Injection Instrument | 89.79 (20.63) | 4.18 (2.056) | 4.71 (2.89) | 86.5 (26.51) | 0.071 | 0.036 | 0.893 | 0 | -0.036 | 0.366 (0.281) | 0.104 (0.123) | 0.416 (0.188) |

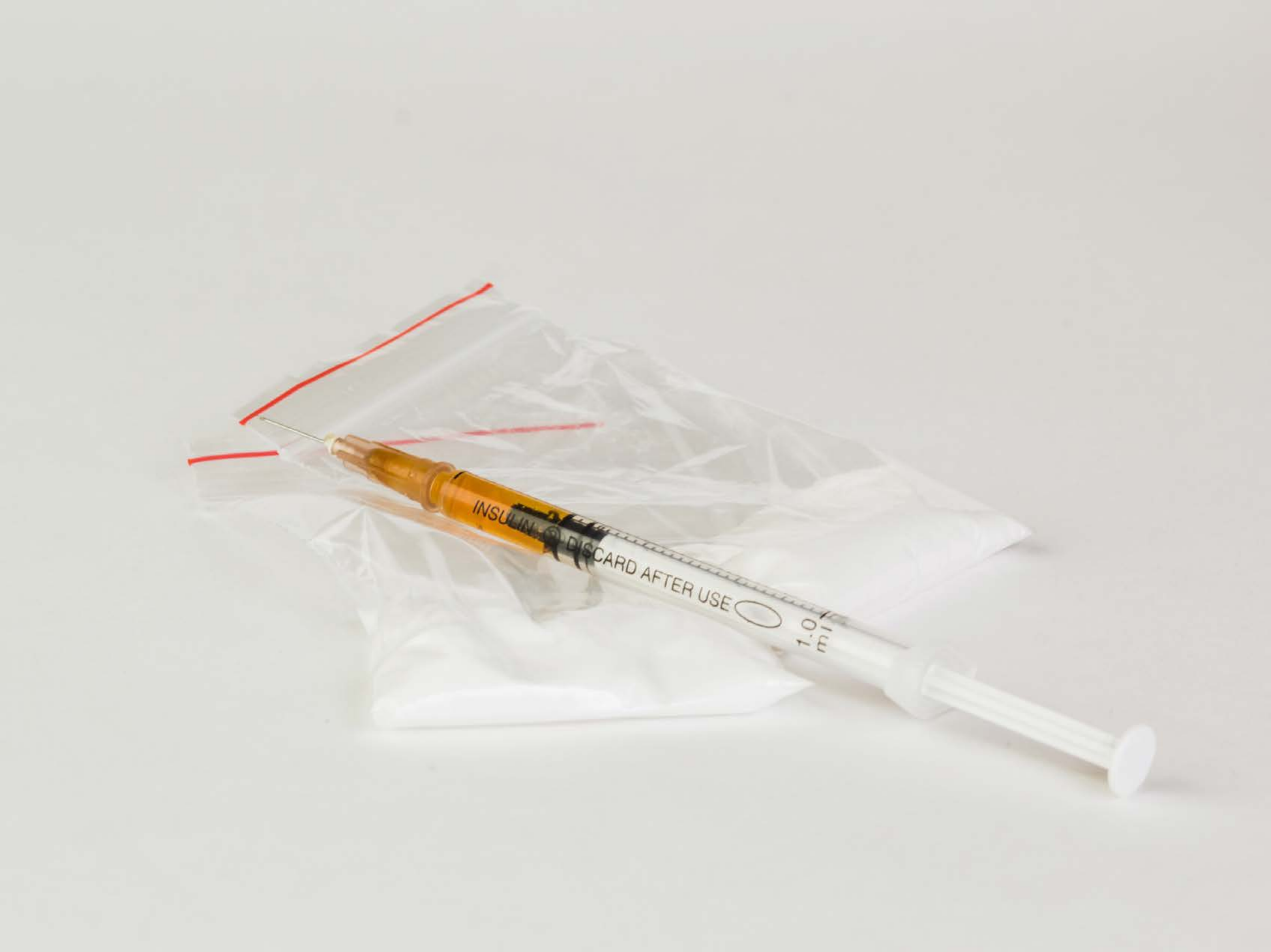


| Category | Craving (0-100) | Valence (1-9) | Arousal (1-9) | Typicality (0-100) | Relatedness | | | | | HSV | | |
|---|---|---|---|---|---|---|---|---|---|---|---|---|
| | | | | | Meth | Opioid | Both | Neither | MethToOpioid | Hue | Saturation | Value |
| Opioid Injection Instrument | 89.04 (16.39) | 4.33 (1.754) | 4.81 (2.69) | 90.07 (16.05) | 0.185 | 0.370 | 0.444 | 0 | 0.185 | 0.145 (0.079) | 0.037 (0.075) | 0.856 (0.039) |

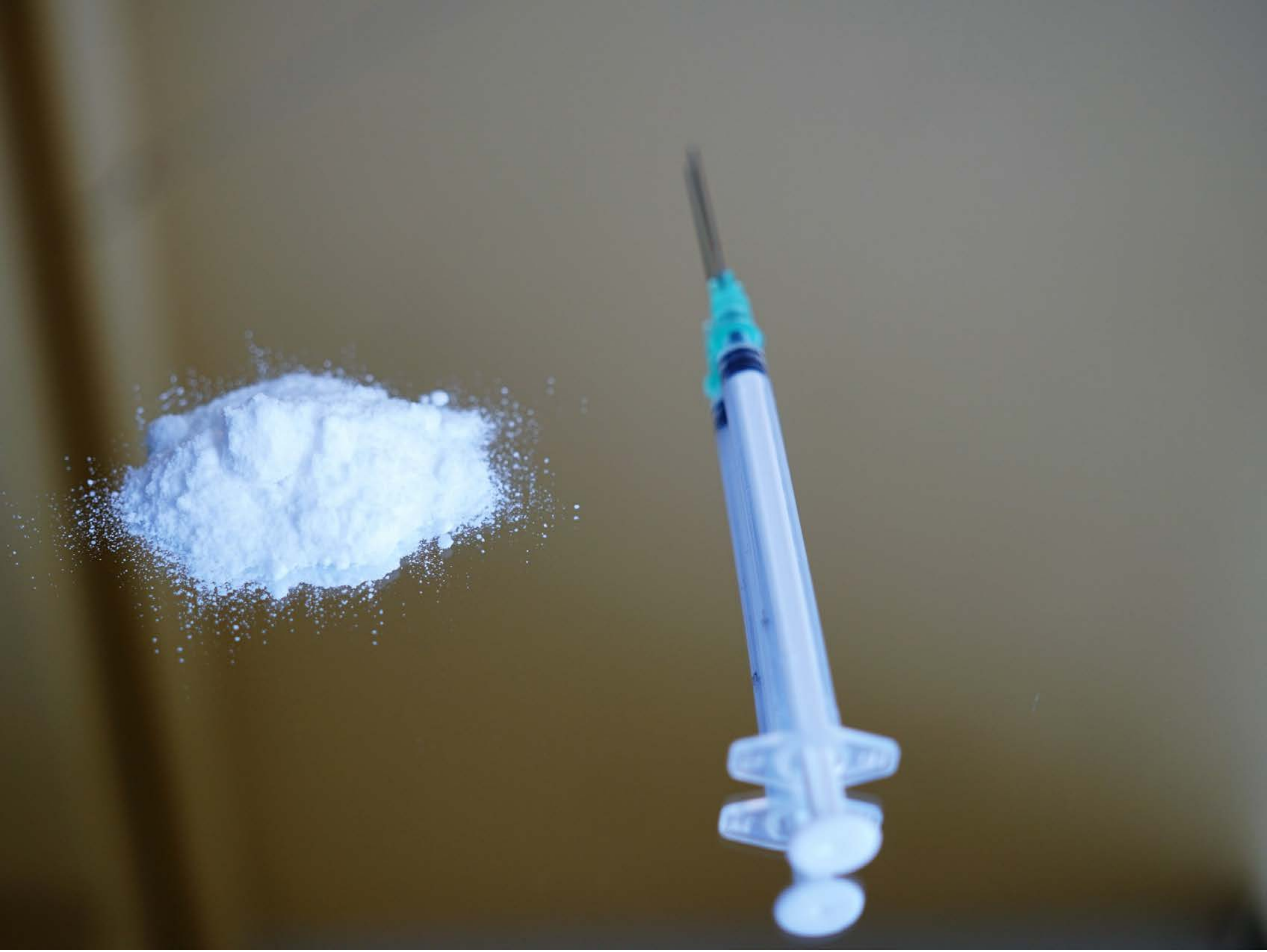

| Category | Craving (0-100) | Valence (1-9) | Arousal (1-9) | Typicality (0-100) | Relatedness | | | | | HSV | | |
|---|---|---|---|---|---|---|---|---|---|---|---|---|
| | | | | | Meth | Opioid | Both | Neither | MethToOpioid | Hue | Saturation | Value |
| Opioid Injection Instrument | 93.54 (10.95) | 4.5 (2.082) | 4.93 (2.92) | 95.11 (9.59) | 0.25 | 0.143 | 0.607 | 0 | -0.107 | 0.21 (0.162) | 0.256 (0.223) | 0.46 (0.181) |

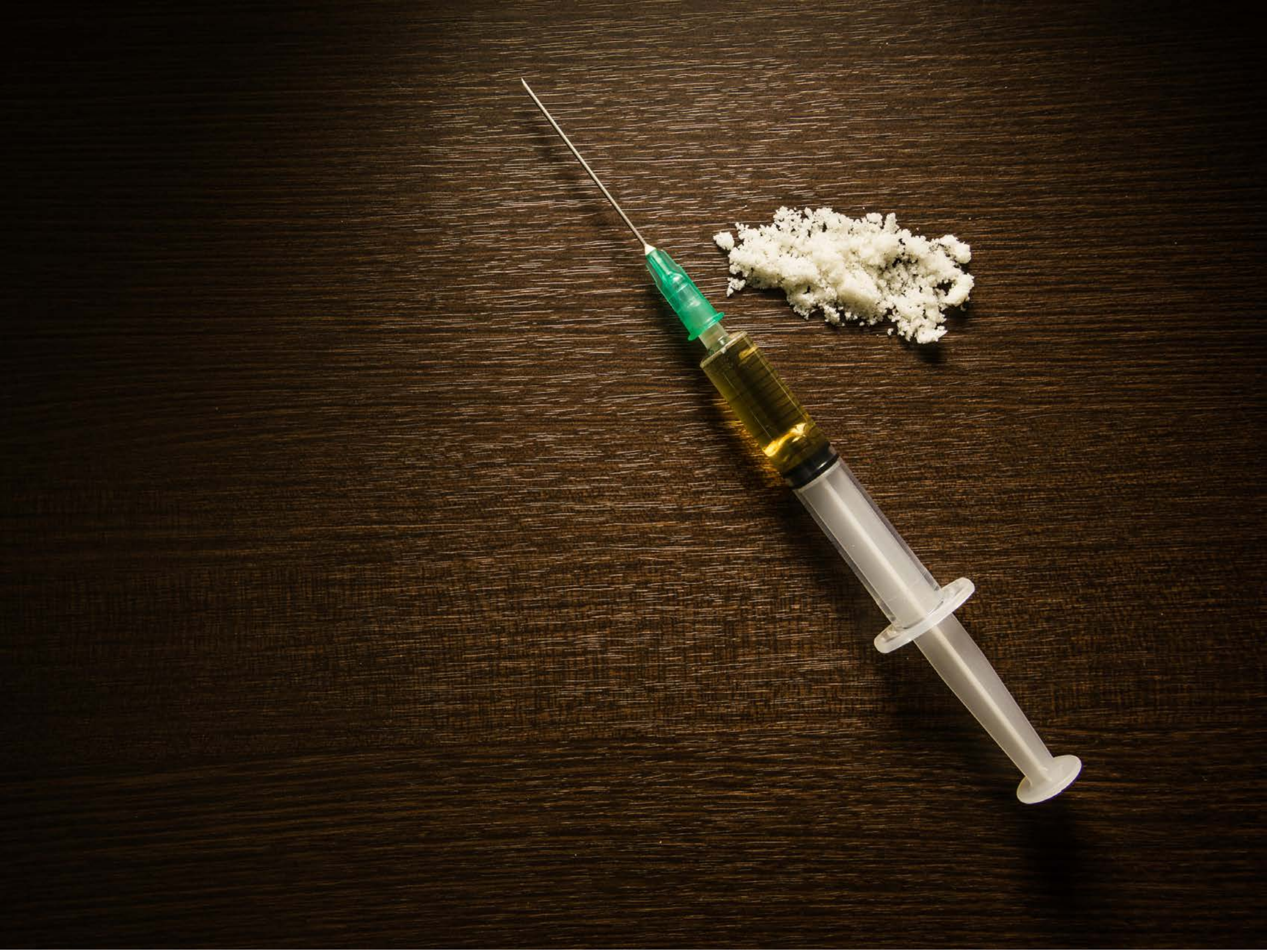

| Category | Craving (0-100) | Valence (1-9) | Arousal (1-9) | Typicality (0-100) | Relatedness | | | | | HSV | | |
|---|---|---|---|---|---|---|---|---|---|---|---|---|
| | | | | | Meth | Opioid | Both | Neither | MethToOpioid | Hue | Saturation | Value |
| Opioid Injection Instrument | 89 (25.14) | 4.29 (1.979) | 4.71 (2.93) | 91.71 (19.82) | 0.321 | 0.179 | 0.5 | 0 | -0.143 | 0.085 (0.039) | 0.701 (0.165) | 0.196 (0.168) |

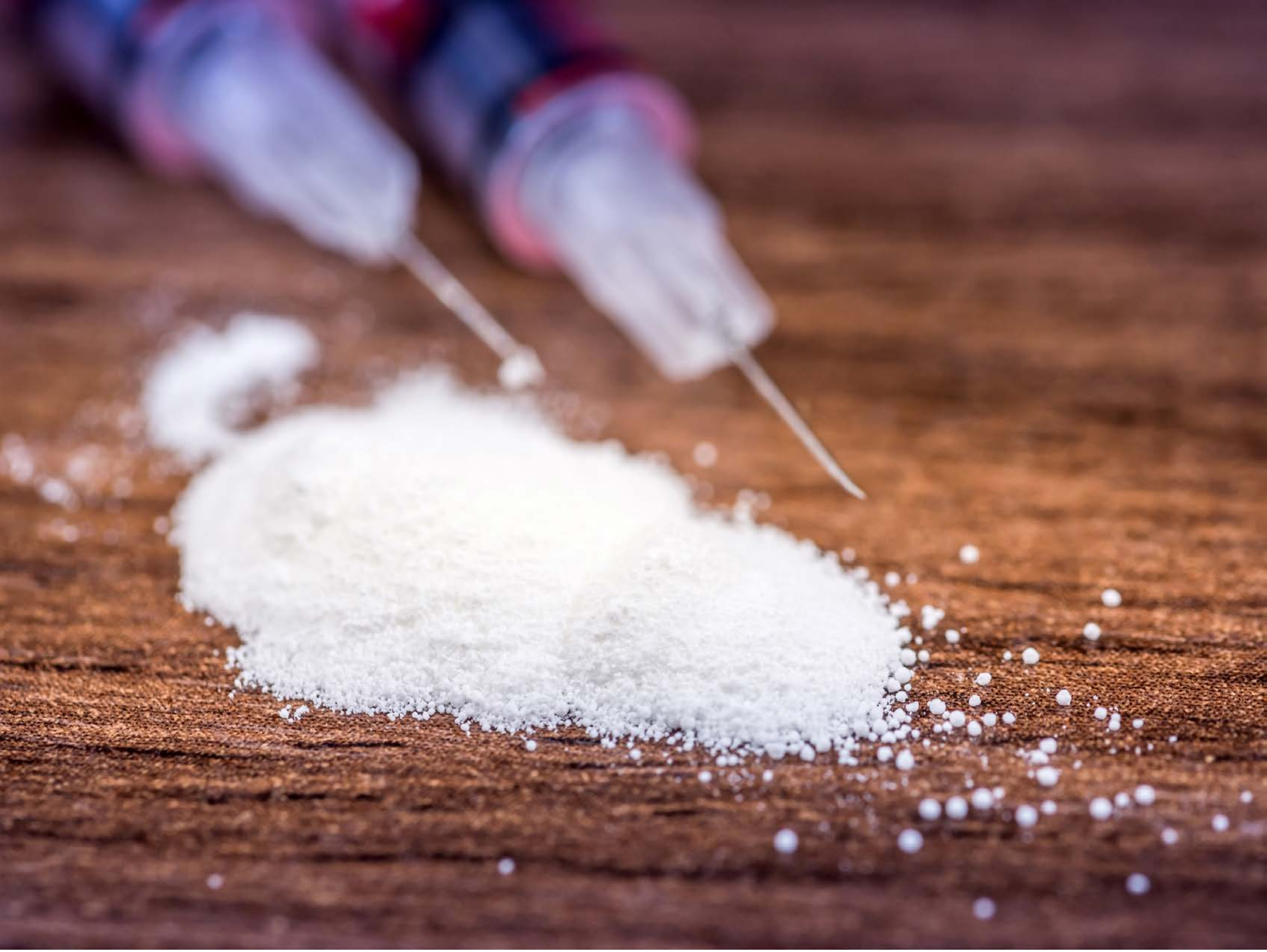

| Category | Craving (0-100) | Valence (1-9) | Arousal (1-9) | Typicality (0-100) | Relatedness | | | | | HSV | | |
|---|---|---|---|---|---|---|---|---|---|---|---|---|
| | | | | | Meth | Opioid | Both | Neither | MethToOpioid | Hue | Saturation | Value |
| Opioid Injection Instrument | 87.63 (17.96) | 4.26 (1.852) | 4.26 (2.65) | 87.44 (18.16) | 0.333 | 0.296 | 0.333 | 0.037 | -0.037 | 0.394 (0.406) | 0.38 (0.205) | 0.579 (0.219) |

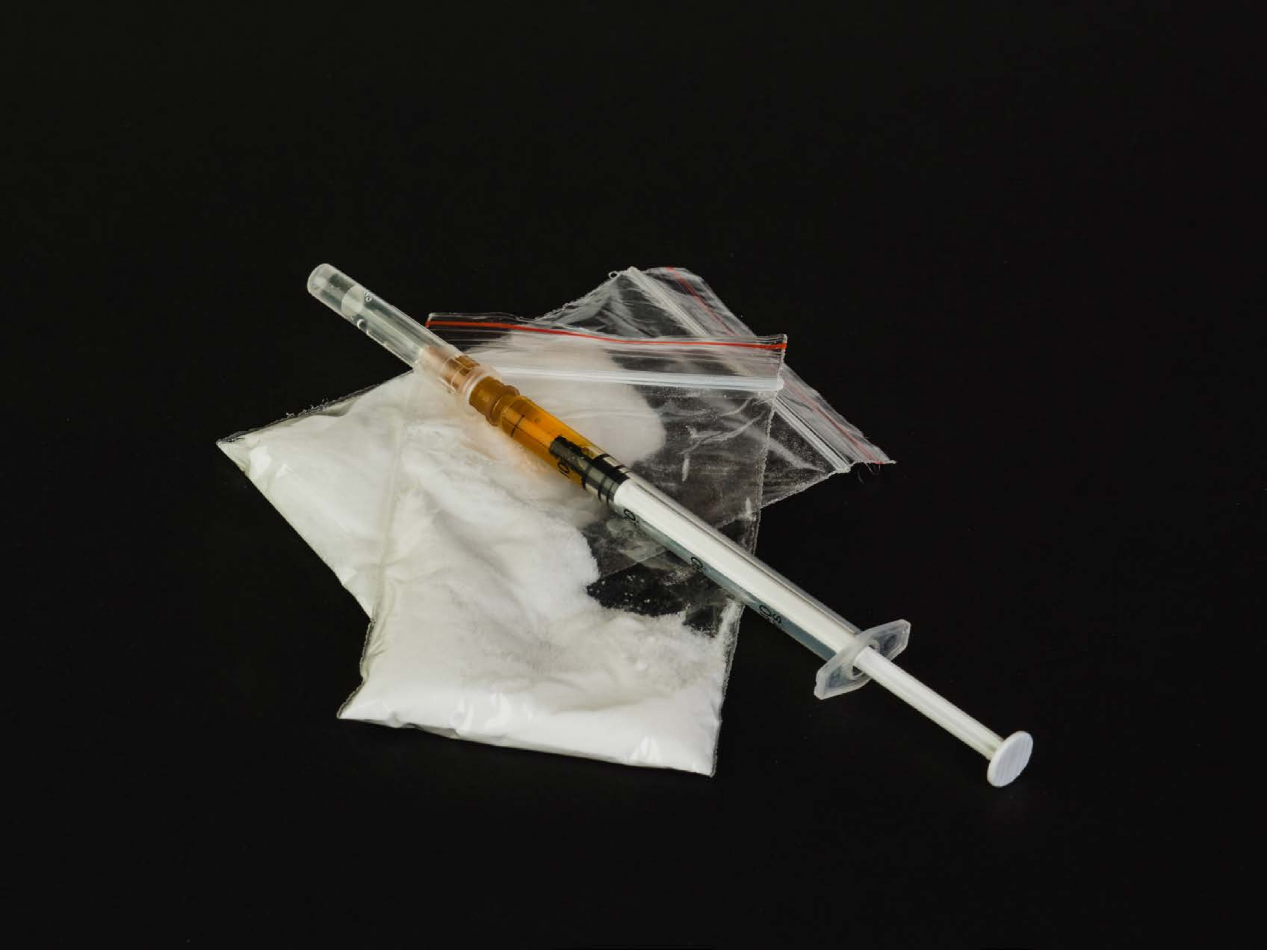

| Category | Craving (0-100) | Valence (1-9) | Arousal (1-9) | Typicality (0-100) | Relatedness | | | | | HSV | | |
|---|---|---|---|---|---|---|---|---|---|---|---|---|
| | | | | | Meth | Opioid | Both | Neither | MethToOpioid | Hue | Saturation | Value |
| Opioid Injection Instrument | 85.67 (23.29) | 4.26 (1.852) | 4.22 (2.72) | 87.22 (22.63) | 0.185 | 0.407 | 0.407 | 0 | 0.222 | 0.367 (0.389) | 0.125 (0.092) | 0.158 (0.234) |

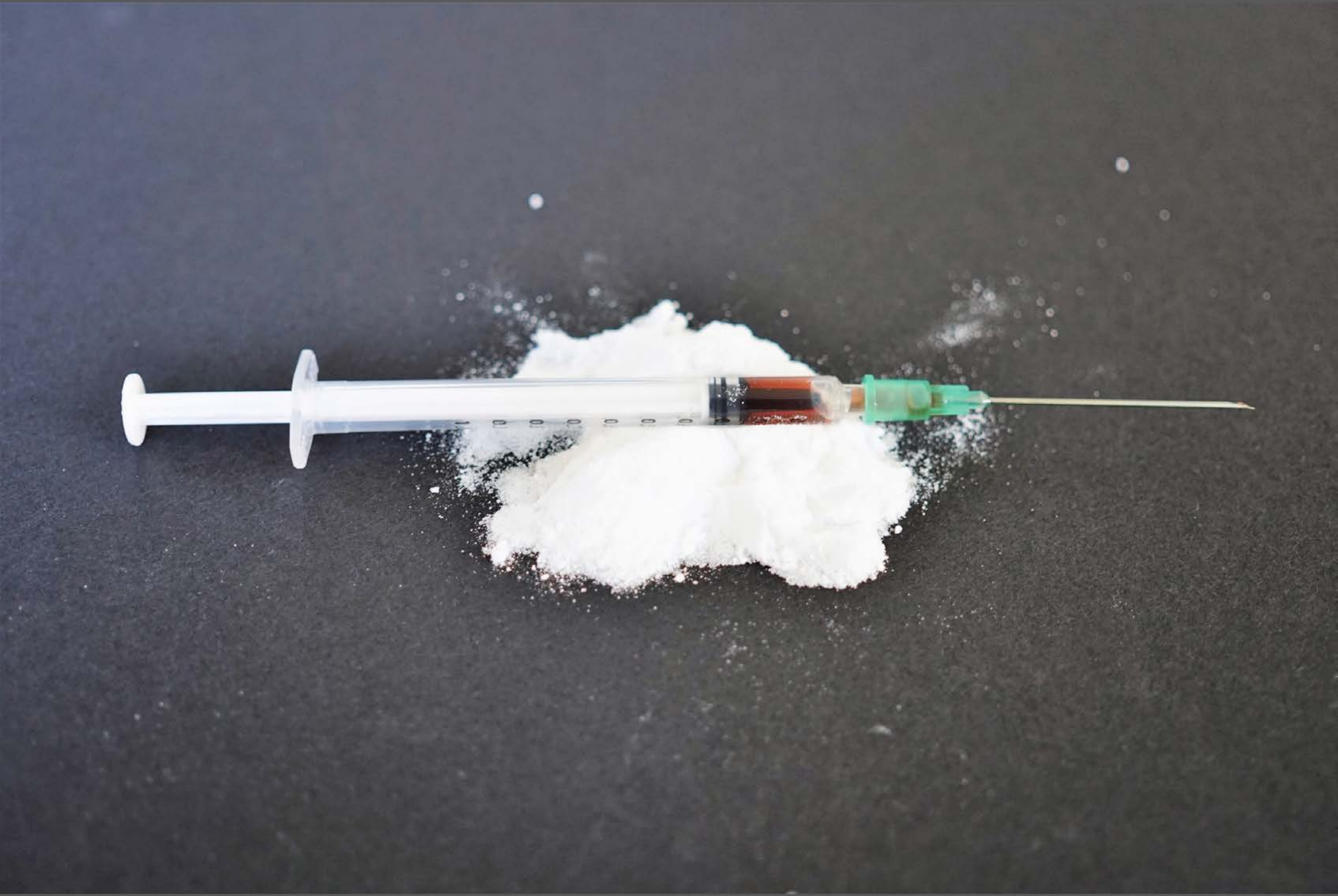

| Category | Craving (0-100) | Valence (1-9) | Arousal (1-9) | Typicality (0-100) | Relatedness | | | | | HSV | | |
|---|---|---|---|---|---|---|---|---|---|---|---|---|
| | | | | | Meth | Opioid | Both | Neither | MethToOpioid | Hue | Saturation | Value |
| Opioid Injection Instrument | 83.85 (19.69) | 4.44 (1.761) | 4.63 (2.69) | 86.07 (18.22) | 0.111 | 0.481 | 0.407 | 0 | 0.370 | 0.383 (0.32) | 0.075 (0.075) | 0.402 (0.174) |

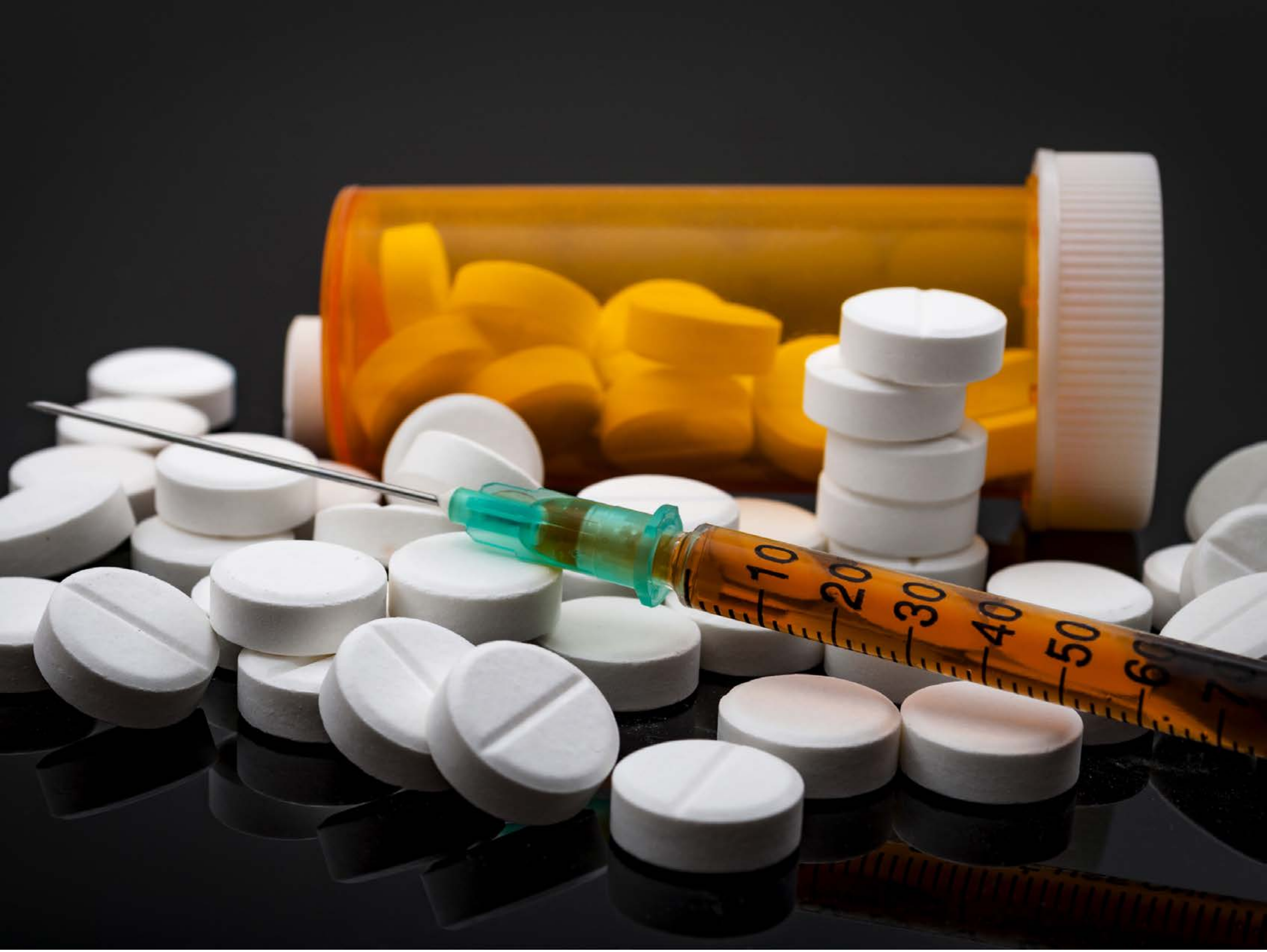


| Category | Craving (0-100) | Valence (1-9) | Arousal (1-9) | Typicality (0-100) | Relatedness | | | | | HSV | | |
|---|---|---|---|---|---|---|---|---|---|---|---|---|
| | | | | | Meth | Opioid | Both | Neither | MethToOpioid | Hue | Saturation | Value |
| Opioid Injection Instrument | 86.85 (18.07) | 4.56 (1.908) | 4.63 (2.68) | 85.93 (18.9) | 0 | 0.889 | 0.111 | 0 | 0.889 | 0.222 (0.313) | 0.266 (0.385) | 0.382 (0.32) |

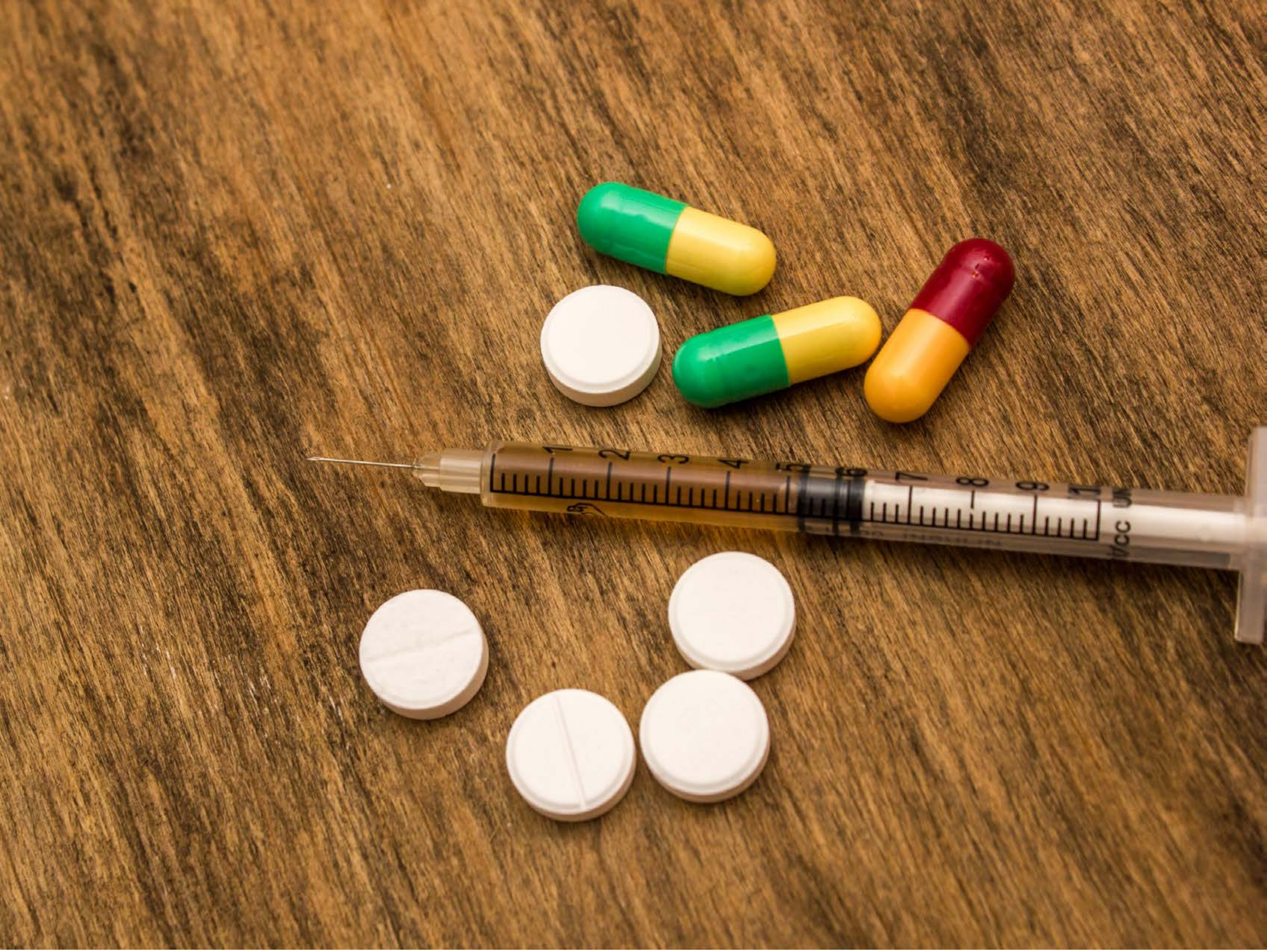

| Category | Craving (0-100) | Valence (1-9) | Arousal (1-9) | Typicality (0-100) | Relatedness | | | | | HSV | | |
|---|---|---|---|---|---|---|---|---|---|---|---|---|
| | | | | | Meth | Opioid | Both | Neither | MethToOpioid | Hue | Saturation | Value |
| Opioid Injection Instrument | 90.75 (15.66) | 4.43 (1.709) | 4.68 (2.78) | 91.75 (13.19) | 0.036 | 0.714 | 0.25 | 0 | 0.679 | 0.094 (0.074) | 0.641 (0.164) | 0.583 (0.173) |

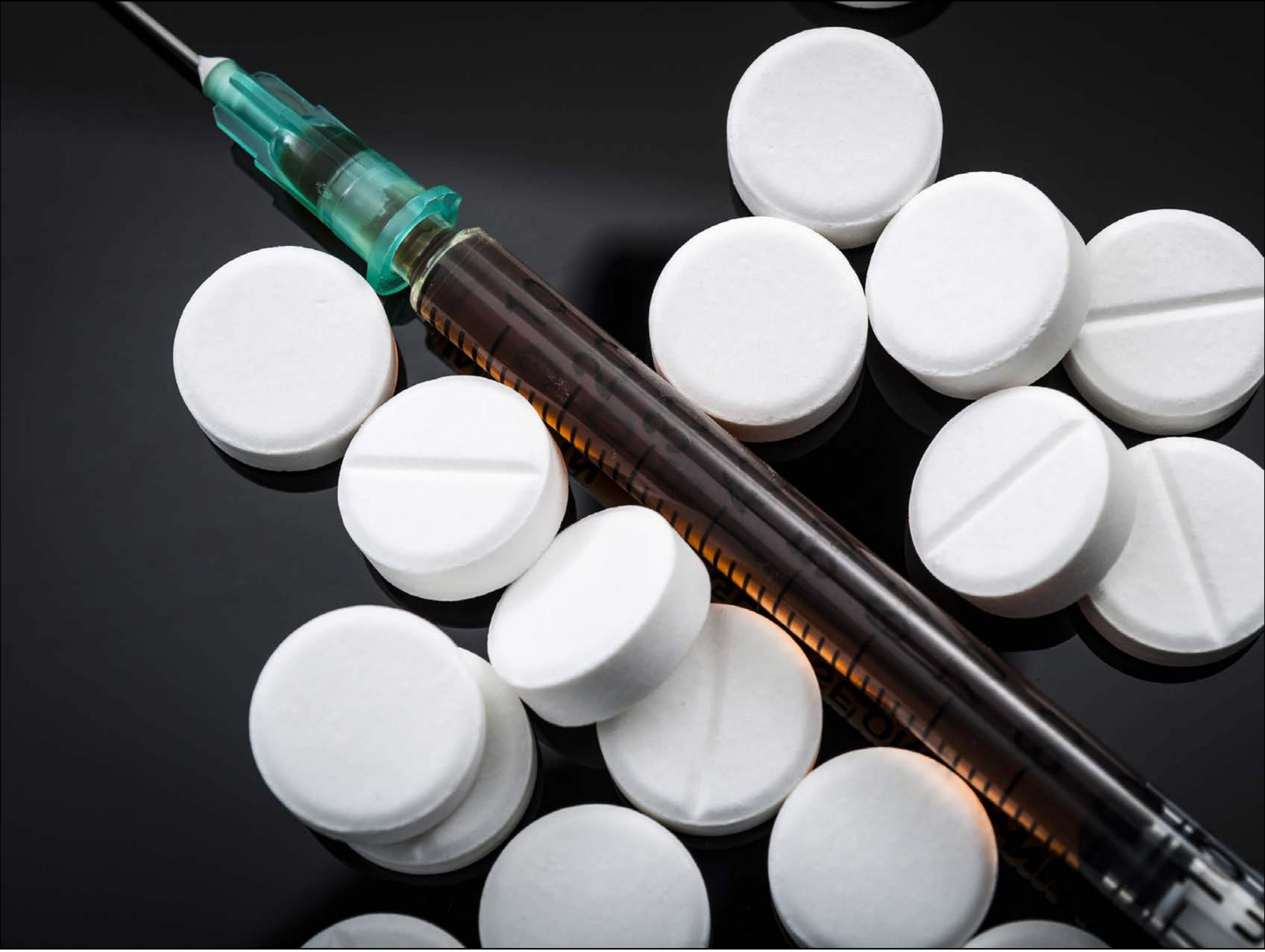

| Category | Craving (0-100) | Valence (1-9) | Arousal (1-9) | Typicality (0-100) | Relatedness | | | | | HSV | | |
|---|---|---|---|---|---|---|---|---|---|---|---|---|
| | | | | | Meth | Opioid | Both | Neither | MethToOpioid | Hue | Saturation | Value |
| Opioid Injection Instrument | 81.81 (23.6) | 4.81 (1.57) | 4.52 (2.41) | 87.41 (17.86) | 0 | 0.889 | 0.111 | 0 | 0.889 | 0.437 (0.298) | 0.085 (0.148) | 0.432 (0.357) |

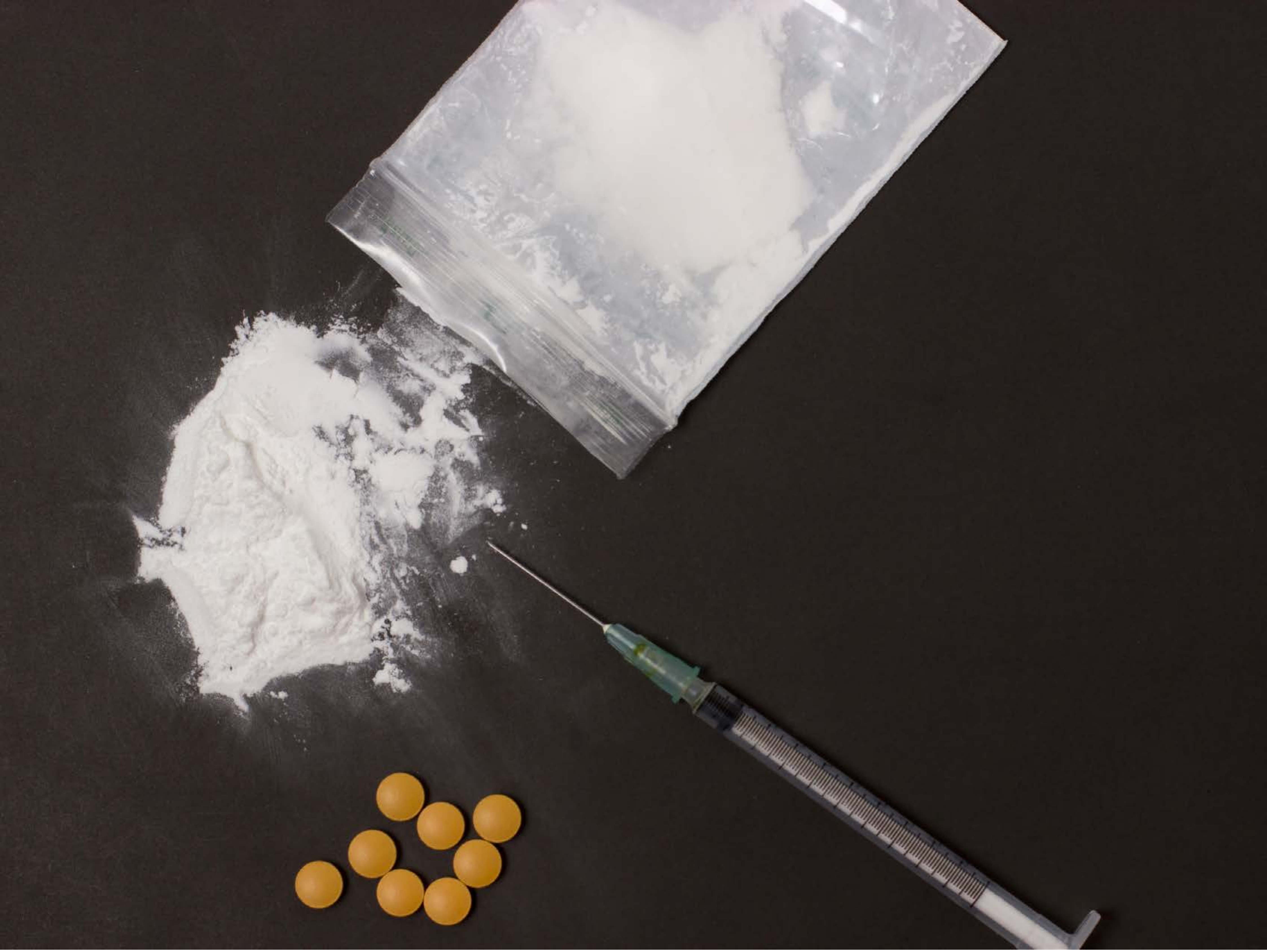

| Category | Craving (0-100) | Valence (1-9) | Arousal (1-9) | Typicality (0-100) | Relatedness | | | | | HSV | | |
|---|---|---|---|---|---|---|---|---|---|---|---|---|
| | | | | | Meth | Opioid | Both | Neither | MethToOpioid | Hue | Saturation | Value |
| Opioid Injection Instrument | 83.15 (20.98) | 4.52 (1.847) | 4.67 (2.67) | 84.89 (20.77) | 0 | 0.593 | 0.407 | 0 | 0.593 | 0.164 (0.281) | 0.18 (0.115) | 0.318 (0.262) |

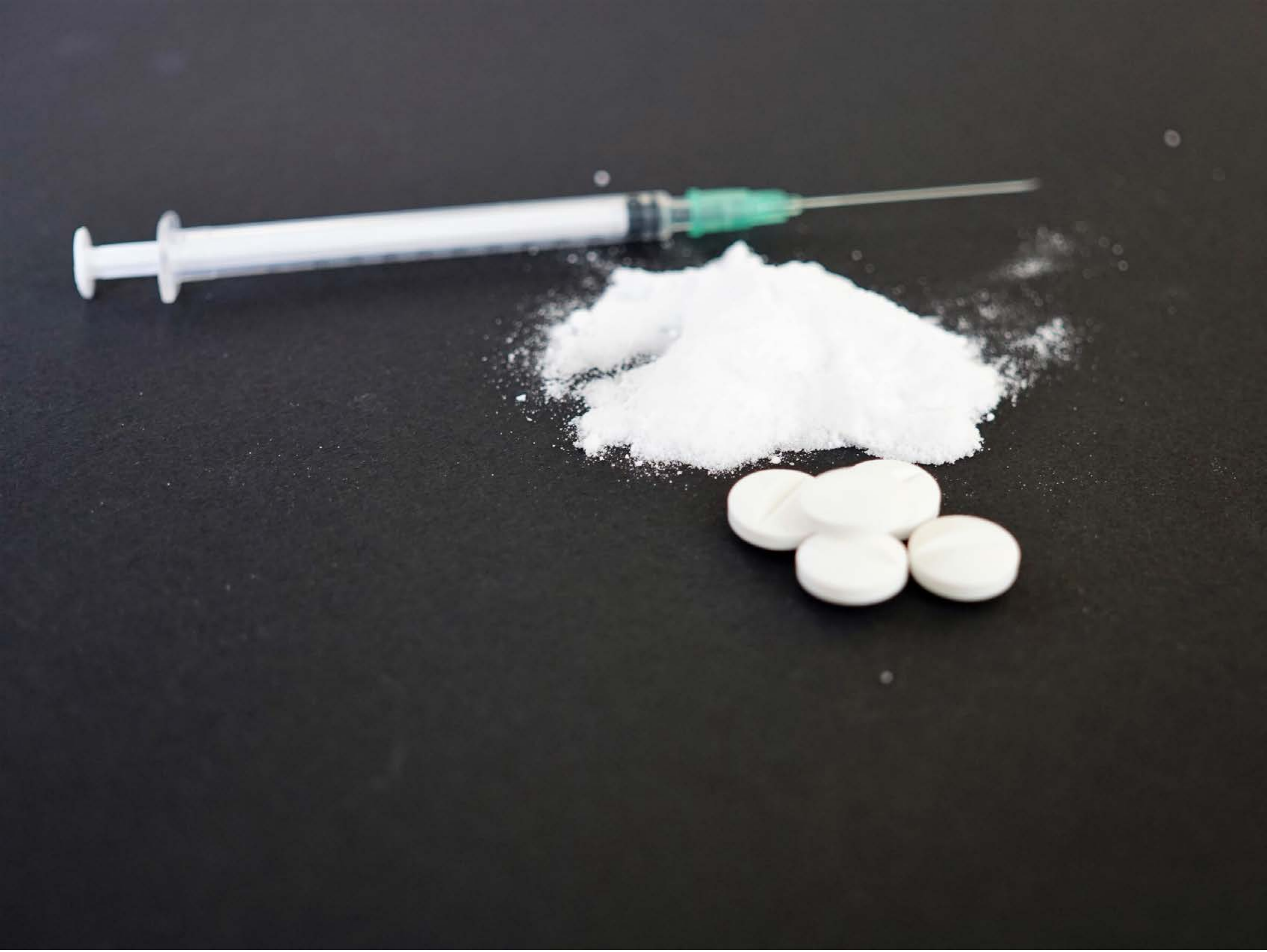

| Category | Craving (0-100) | Valence (1-9) | Arousal (1-9) | Typicality (0-100) | Relatedness | | | | | HSV | | |
|---|---|---|---|---|---|---|---|---|---|---|---|---|
| | | | | | Meth | Opioid | Both | Neither | MethToOpioid | Hue | Saturation | Value |
| Opioid Injection Instrument | 87.18 (21.7) | 4.46 (1.688) | 4.5 (2.44) | 88.75 (22.55) | 0 | 0.5 | 0.5 | 0 | 0.5 | 0.346 (0.322) | 0.09 (0.067) | 0.295 (0.23) |

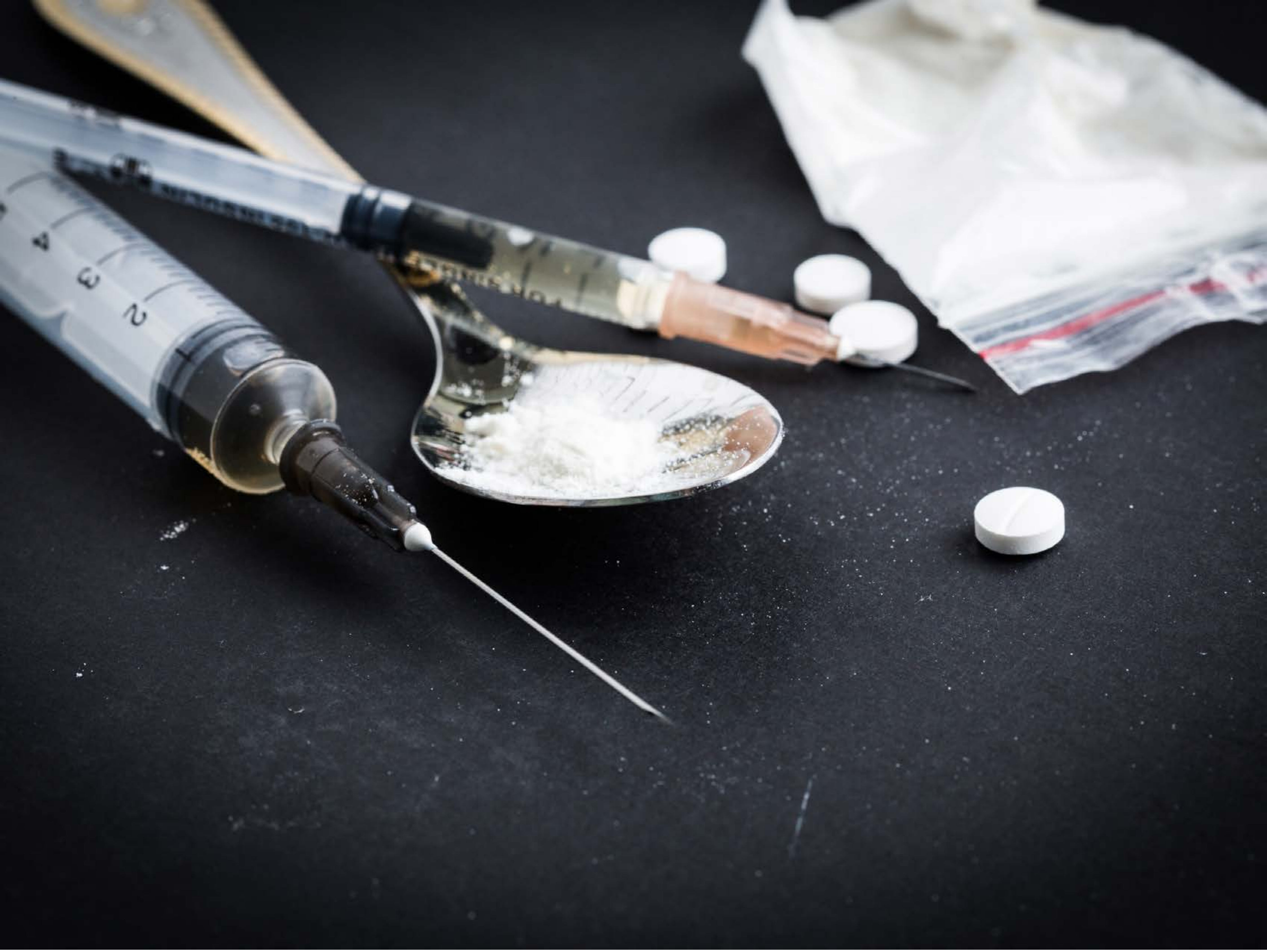


| Category | Craving (0-100) | Valence (1-9) | Arousal (1-9) | Typicality (0-100) | Relatedness | | | | | HSV | | |
|---|---|---|---|---|---|---|---|---|---|---|---|---|
| | | | | | Meth | Opioid | Both | Neither | MethToOpioid | Hue | Saturation | Value |
| Opioid Injection Instrument | 93.68 (11.48) | 4.29 (2.07) | 4.93 (2.75) | 95.36 (10.12) | 0 | 0.393 | 0.607 | 0 | 0.393 | 0.571 (0.165) | 0.158 (0.089) | 0.346 (0.295) |

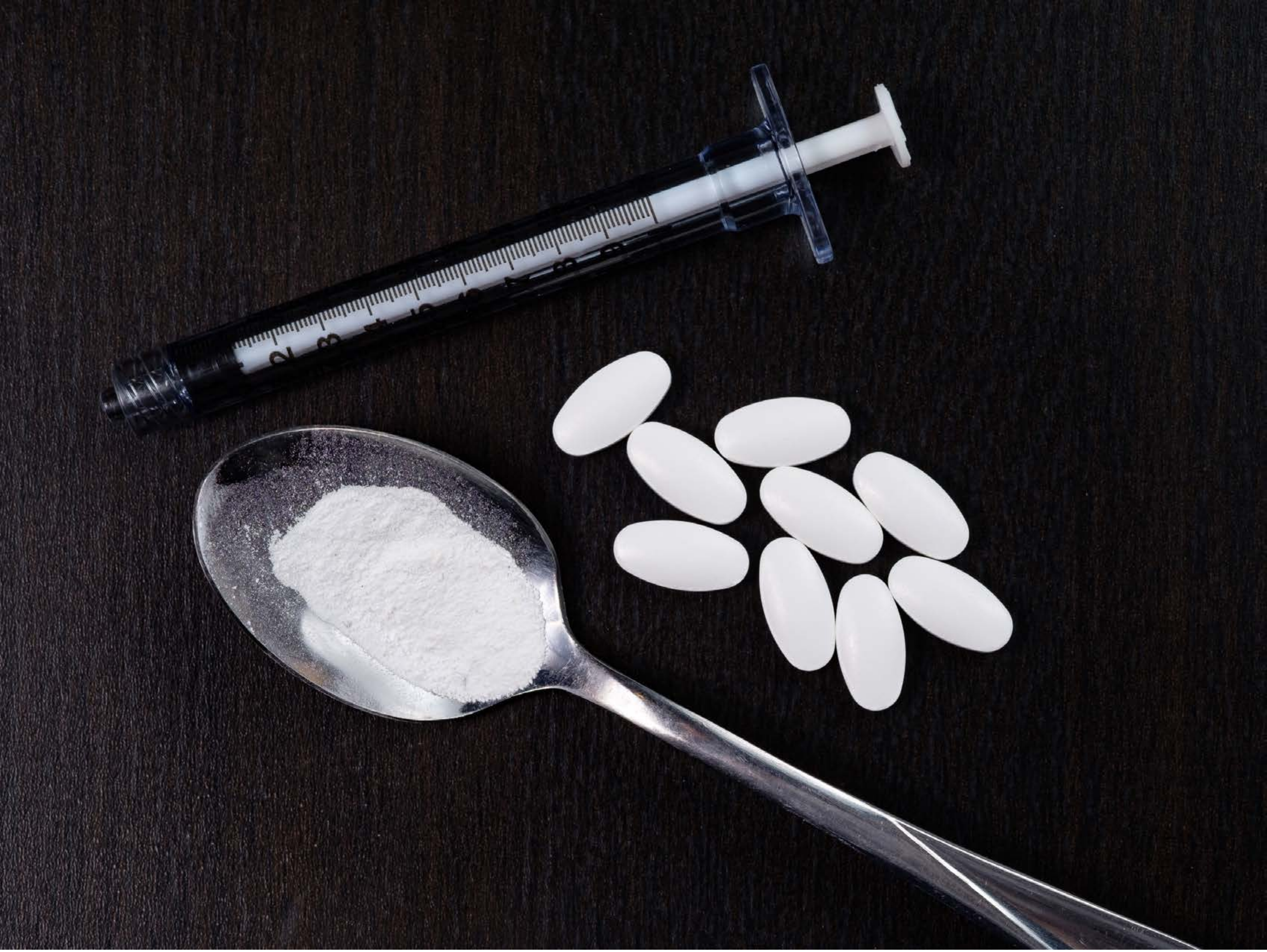

| Category | Craving (0-100) | Valence (1-9) | Arousal (1-9) | Typicality (0-100) | Relatedness | | | | | HSV | | |
|---|---|---|---|---|---|---|---|---|---|---|---|---|
| | | | | | Meth | Opioid | Both | Neither | MethToOpioid | Hue | Saturation | Value |
| Opioid Injection Instrument | 91.89 (11.89) | 4.29 (1.863) | 4.89 (2.85) | 90.11 (13.87) | 0.036 | 0.571 | 0.393 | 0 | 0.536 | 0.679 (0.305) | 0.191 (0.119) | 0.191 (0.257) |

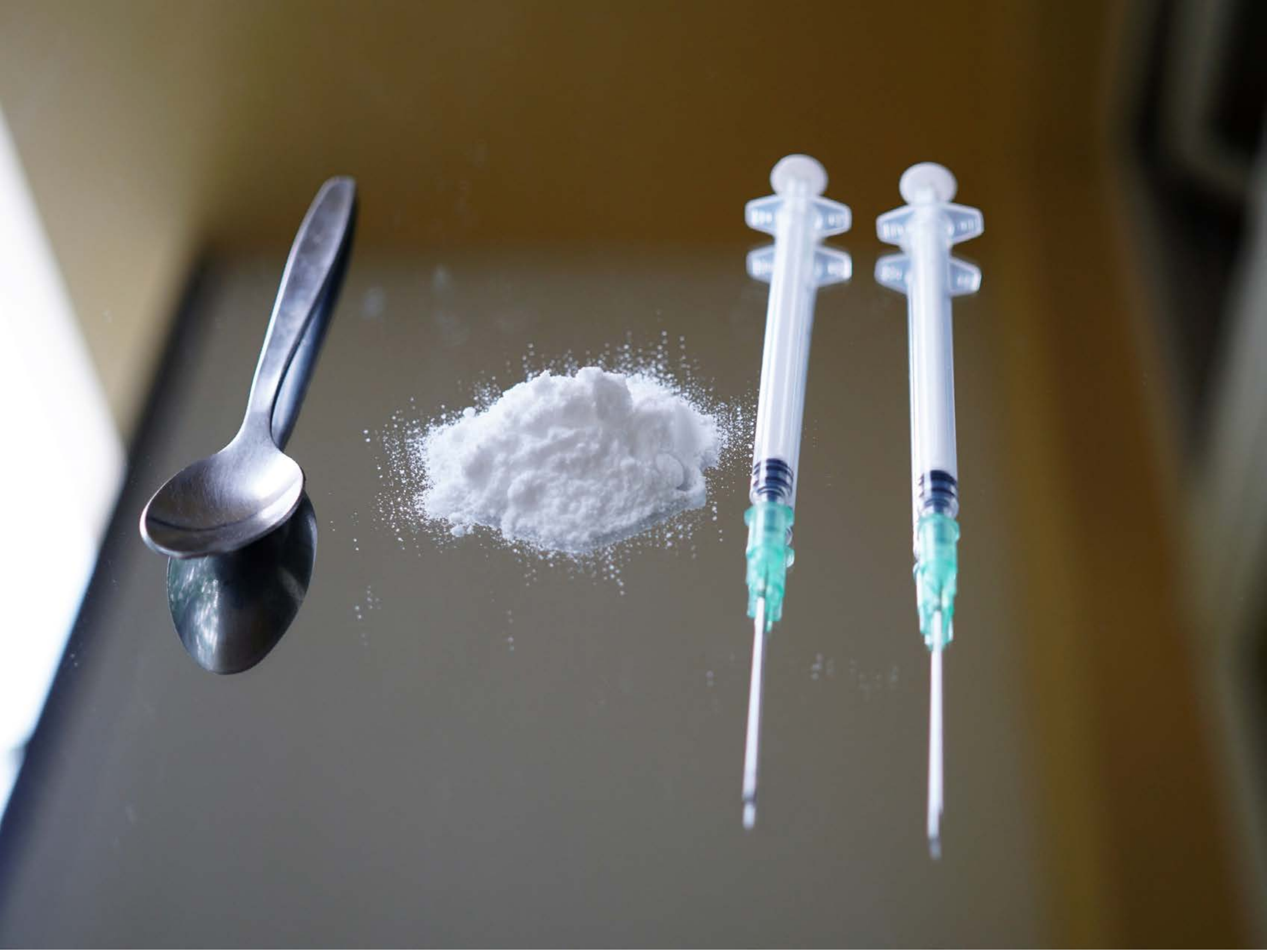

| Category | Craving (0-100) | Valence (1-9) | Arousal (1-9) | Typicality (0-100) | Relatedness | | | | | HSV | | |
|---|---|---|---|---|---|---|---|---|---|---|---|---|
| | | | | | Meth | Opioid | Both | Neither | MethToOpioid | Hue | Saturation | Value |
| Opioid Injection Instrument | 92.79 (13.12) | 4.39 (1.912) | 4.86 (2.77) | 92.64 (12.64) | 0.286 | 0.071 | 0.643 | 0 | -0.214 | 0.236 (0.226) | 0.312 (0.283) | 0.43 (0.202) |

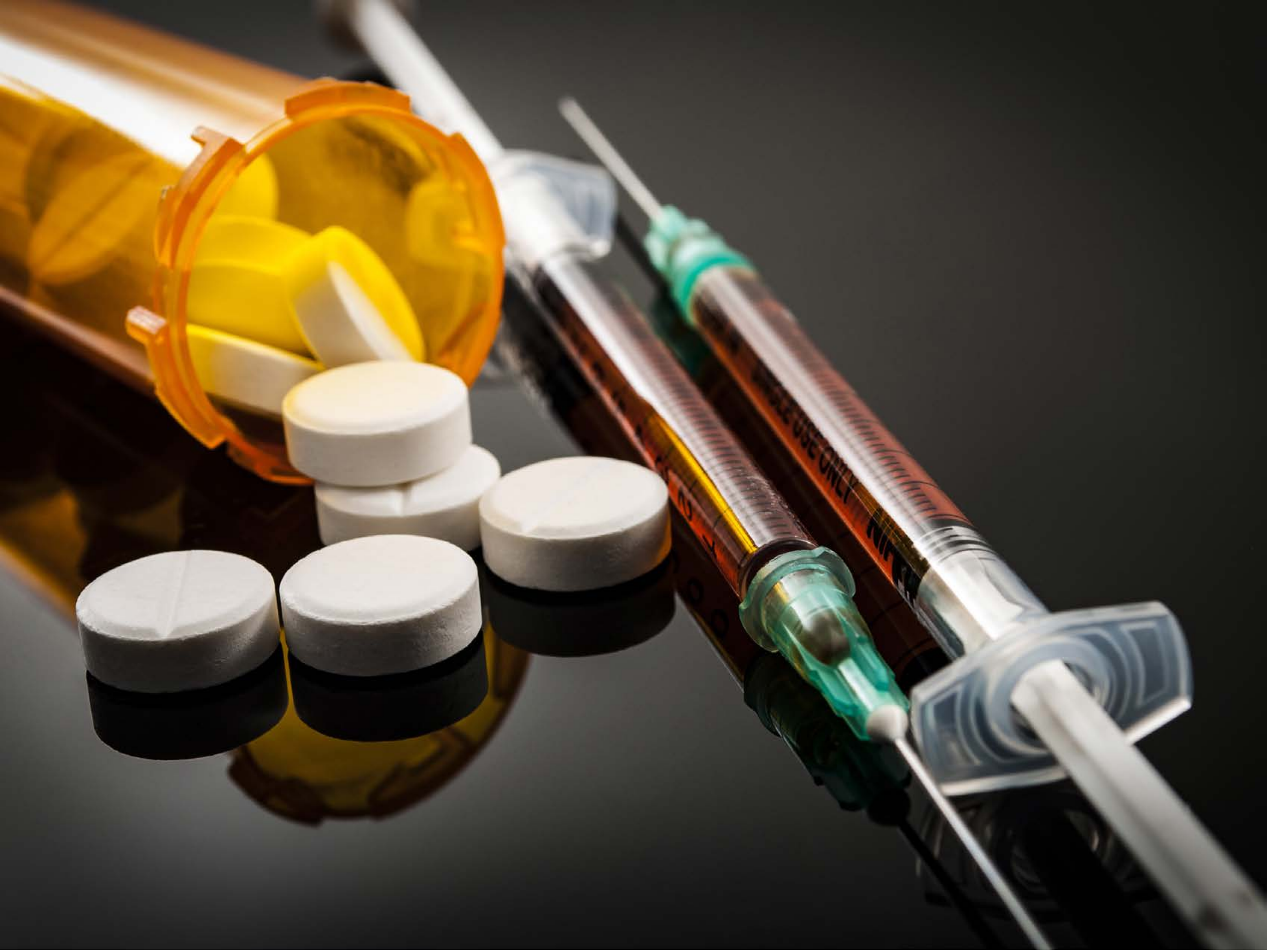

| Category | Craving (0-100) | Valence (1-9) | Arousal (1-9) | Typicality (0-100) | Relatedness | | | | | HSV | | |
|---|---|---|---|---|---|---|---|---|---|---|---|---|
| | | | | | Meth | Opioid | Both | Neither | MethToOpioid | Hue | Saturation | Value |
| Opioid Injection Instrument | 88.56 (15.7) | 4.41 (1.803) | 4.56 (2.49) | 89.67 (15.85) | 0 | 0.778 | 0.222 | 0 | 0.778 | 0.125 (0.15) | 0.291 (0.346) | 0.349 (0.294) |

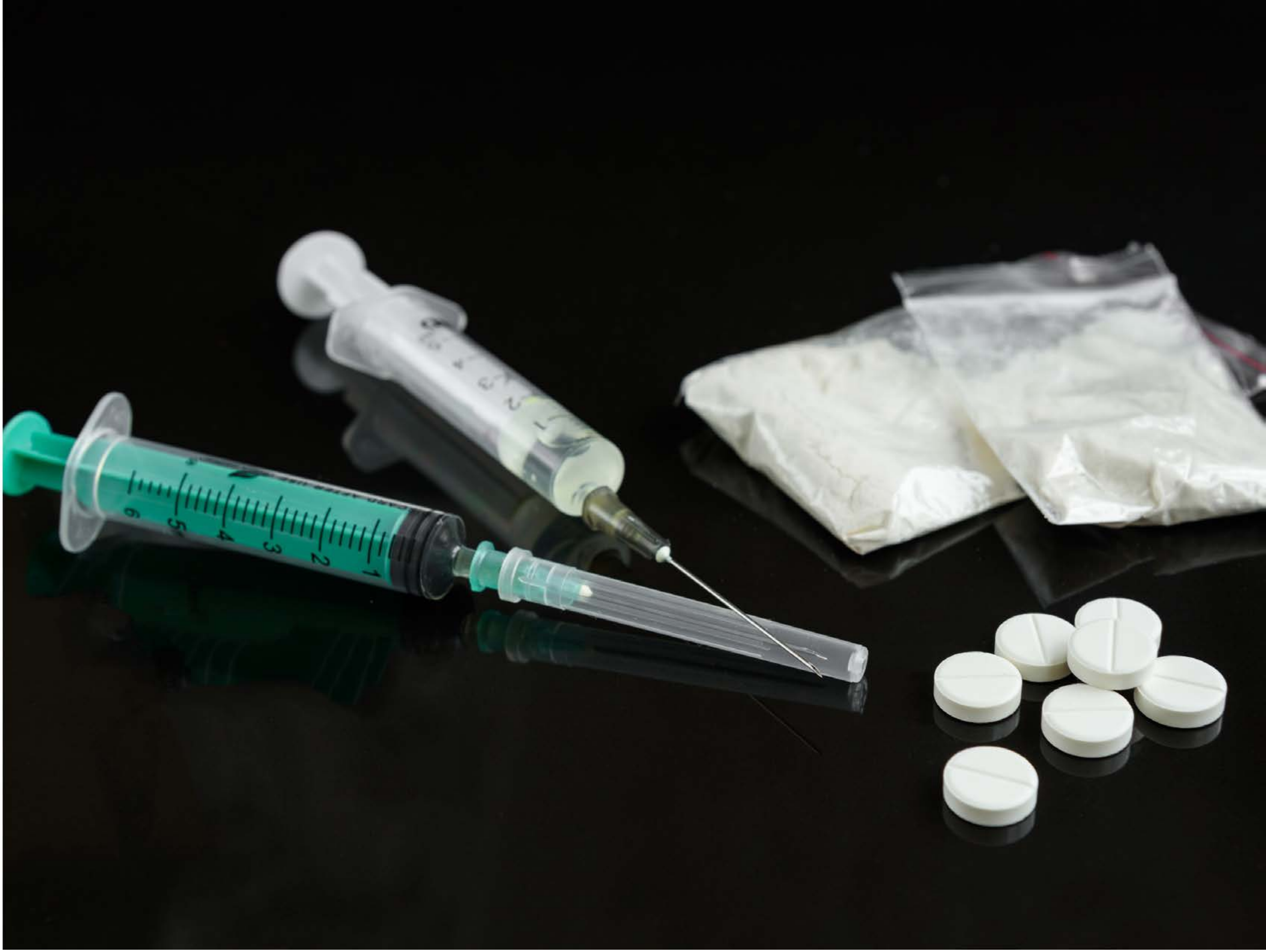

| Category | Craving (0-100) | Valence (1-9) | Arousal (1-9) | Typicality (0-100) | Relatedness | | | | | HSV | | |
|---|---|---|---|---|---|---|---|---|---|---|---|---|
| | | | | | Meth | Opioid | Both | Neither | MethToOpioid | Hue | Saturation | Value |
| Opioid Injection Instrument | 86.04 (23.07) | 4.39 (1.95) | 4.57 (2.85) | 88.86 (21.07) | 0 | 0.286 | 0.714 | 0 | 0.286 | 0.225 (0.216) | 0.244 (0.212) | 0.193 (0.287) |

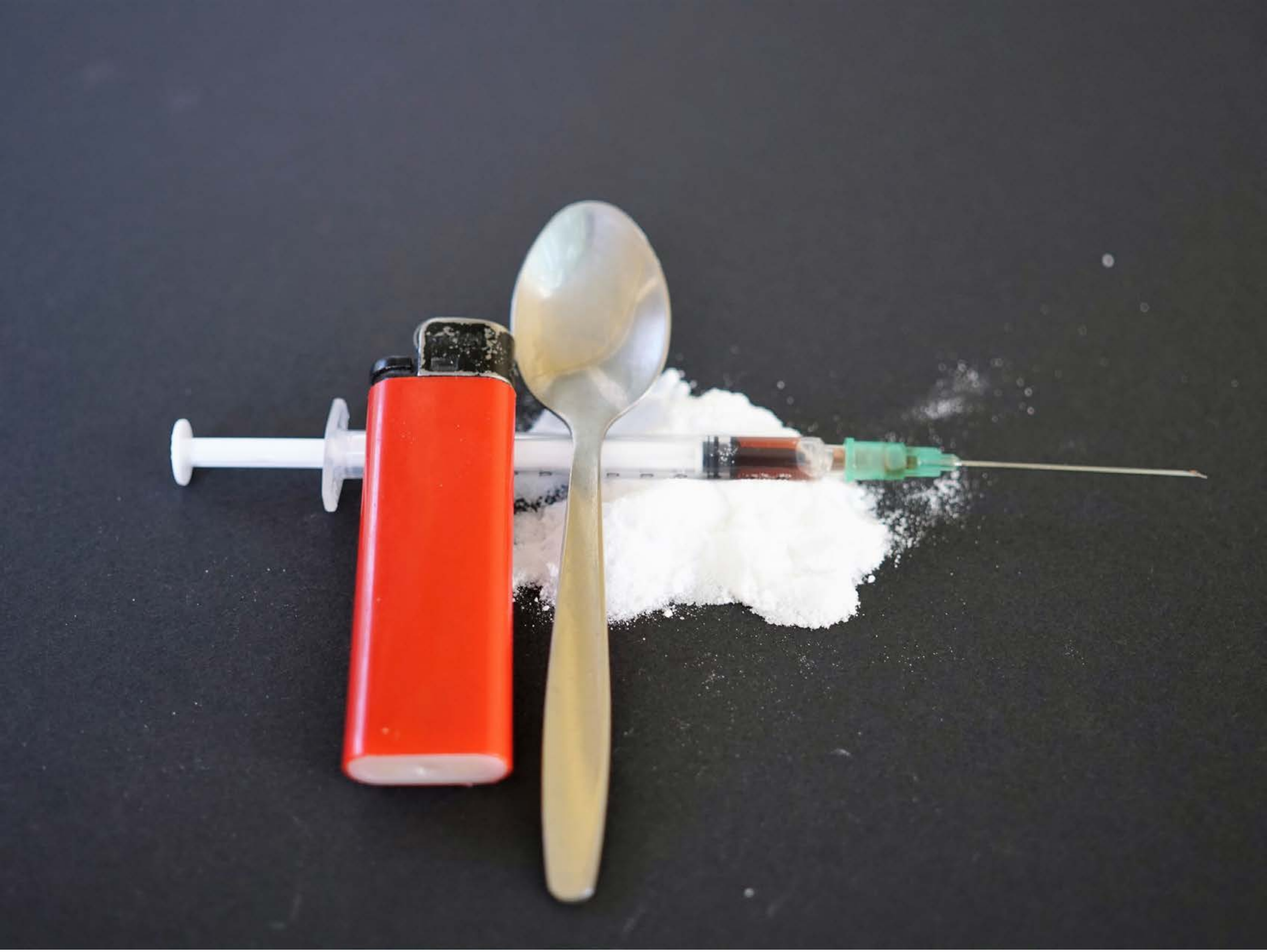

| Category | Craving (0-100) | Valence (1-9) | Arousal (1-9) | Typicality (0-100) | Relatedness | | | | | HSV | | |
|---|---|---|---|---|---|---|---|---|---|---|---|---|
| | | | | | Meth | Opioid | Both | Neither | MethToOpioid | Hue | Saturation | Value |
| Opioid Injection Instrument | 84.19 (20.21) | 4.15 (2.161) | 4.74 (2.43) | 86.07 (19.07) | 0.111 | 0.407 | 0.481 | 0 | 0.296 | 0.482 (0.27) | 0.142 (0.182) | 0.369 (0.23) |

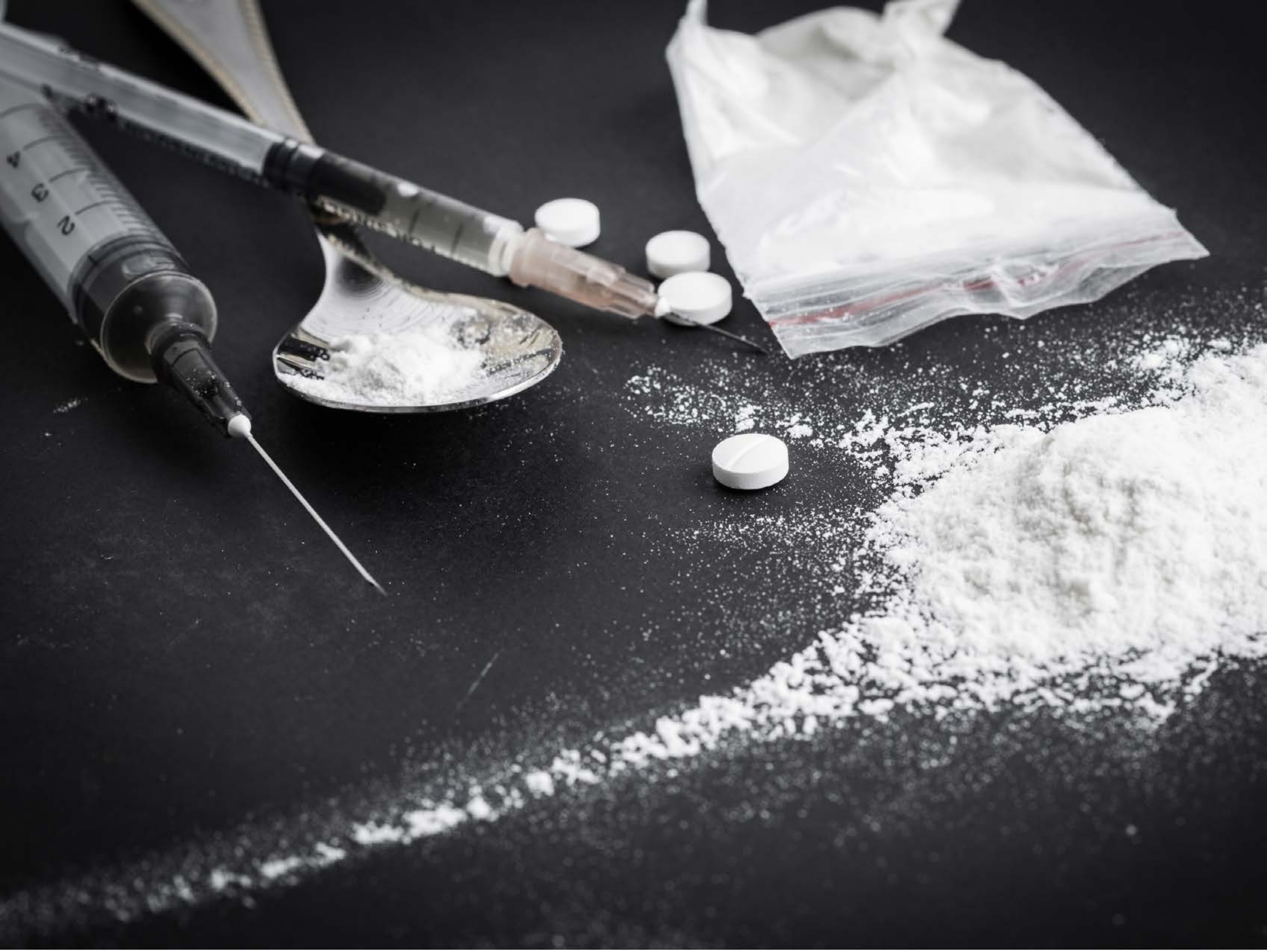

| Category | Craving (0-100) | Valence (1-9) | Arousal (1-9) | Typicality (0-100) | Relatedness | | | | | HSV | | |
|---|---|---|---|---|---|---|---|---|---|---|---|---|
| | | | | | Meth | Opioid | Both | Neither | MethToOpioid | Hue | Saturation | Value |
| Opioid Injection Instrument | 93.71 (11.01) | 4.5 (2.064) | 5.11 (2.79) | 95.14 (10.38) | 0 | 0.429 | 0.571 | 0 | 0.429 | 0.518 (0.257) | 0.044 (0.032) | 0.379 (0.304) |

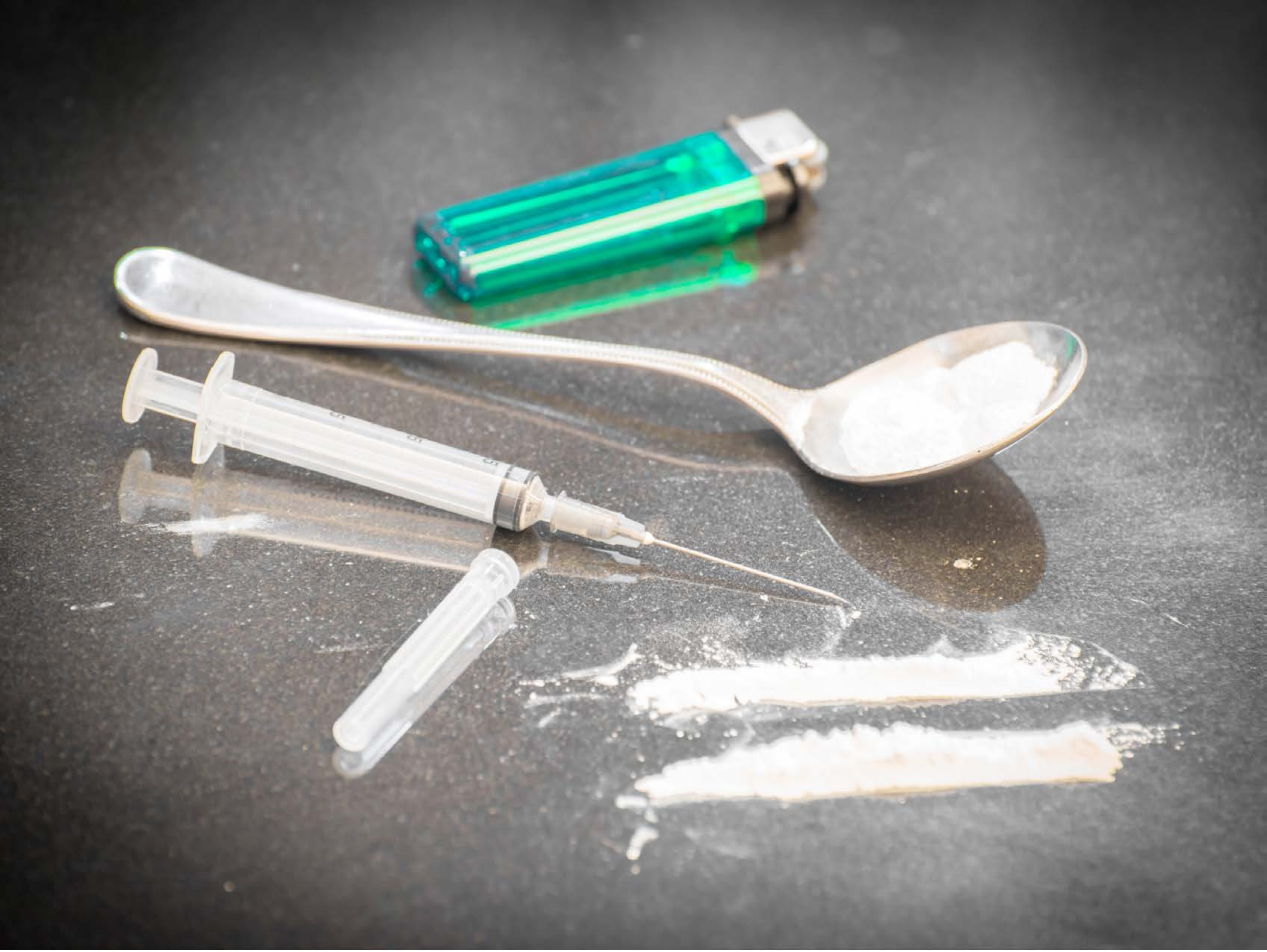

| Category | Craving (0-100) | Valence (1-9) | Arousal (1-9) | Typicality (0-100) | Relatedness | | | | | HSV | | |
|---|---|---|---|---|---|---|---|---|---|---|---|---|
| | | | | | Meth | Opioid | Both | Neither | MethToOpioid | Hue | Saturation | Value |
| Opioid Injection Instrument | 92.93 (11.66) | 4.25 (1.713) | 4.71 (2.83) | 94.07 (10.9) | 0.143 | 0.214 | 0.643 | 0 | 0.071 | 0.117 (0.237) | 0.062 (0.158) | 0.538 (0.219) |

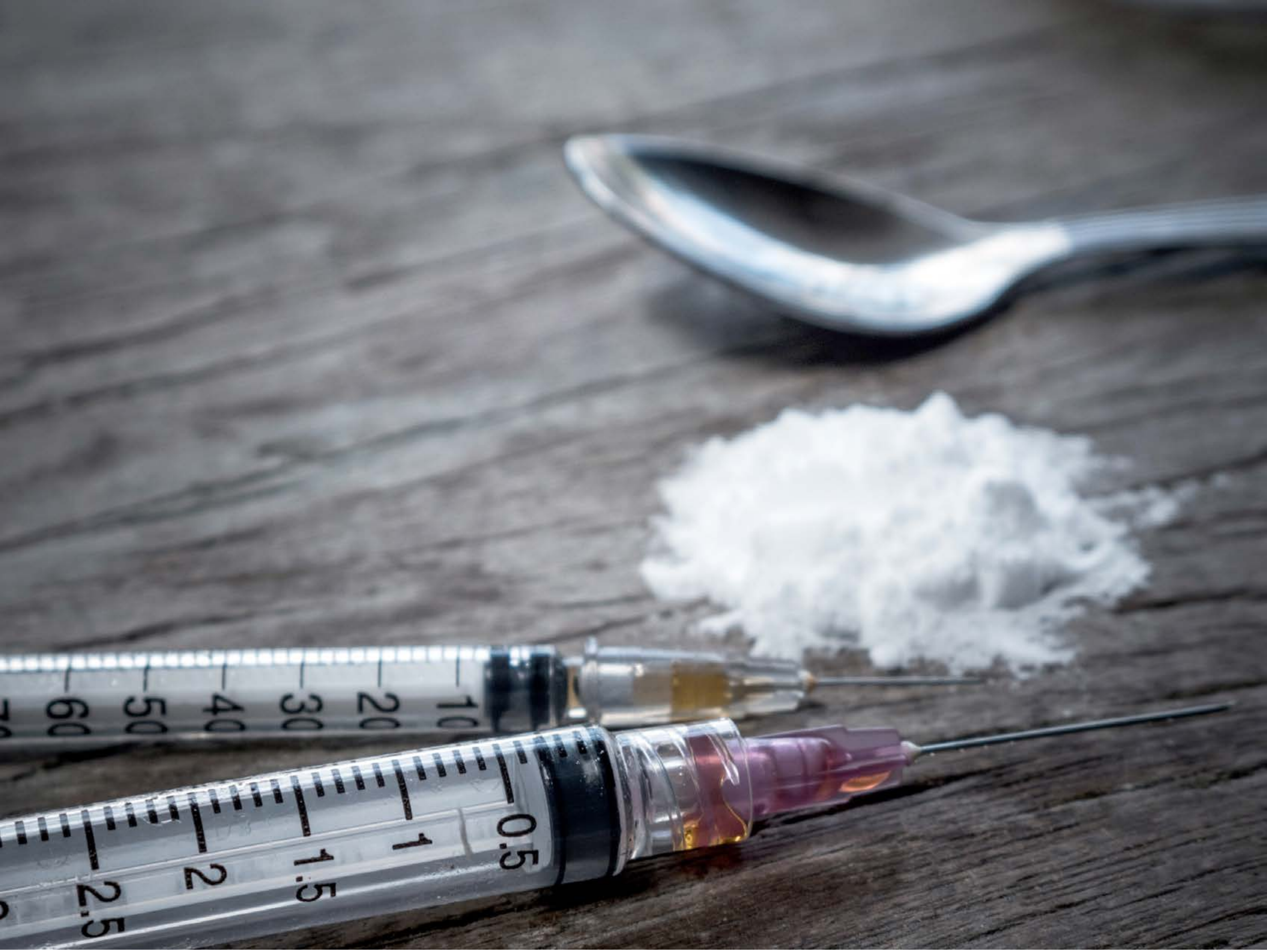


| Category | Craving (0-100) | Valence (1-9) | Arousal (1-9) | Typicality (0-100) | Relatedness | | | | | HSV | | |
|---|---|---|---|---|---|---|---|---|---|---|---|---|
| | | | | | Meth | Opioid | Both | Neither | MethToOpioid | Hue | Saturation | Value |
| Opioid Injection Instrument | 88.93 (15.12) | 4.44 (1.968) | 4.85 (2.68) | 89.44 (15.44) | 0.111 | 0.222 | 0.667 | 0 | 0.111 | 0.593 (0.187) | 0.096 (0.114) | 0.499 (0.21) |

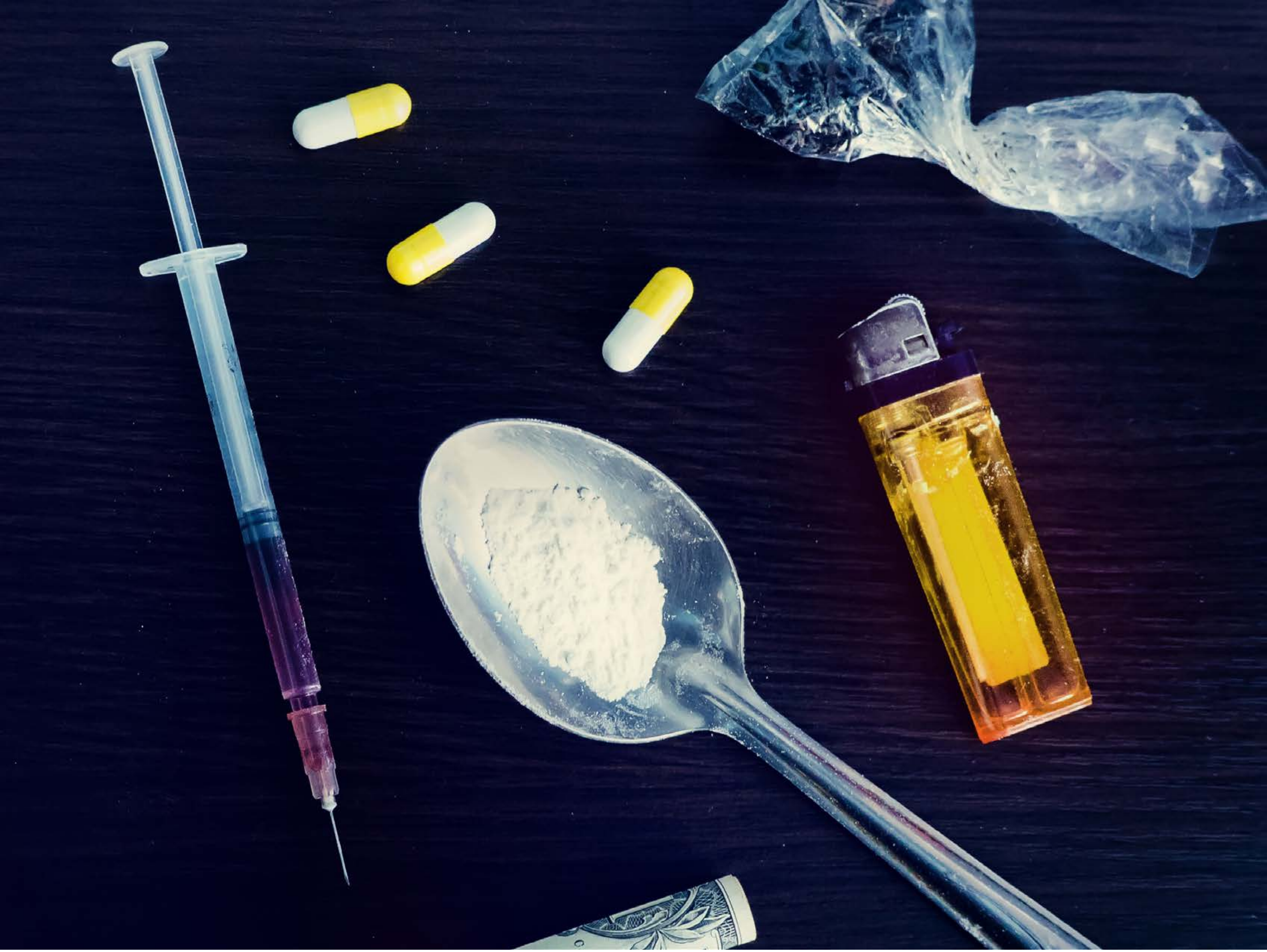

| Category | Craving (0-100) | Valence (1-9) | Arousal (1-9) | Typicality (0-100) | Relatedness | | | | | HSV | | |
|---|---|---|---|---|---|---|---|---|---|---|---|---|
| | | | | | Meth | Opioid | Both | Neither | MethToOpioid | Hue | Saturation | Value |
| Opioid Injection Instrument | 83.96 (24.47) | 4.59 (2.024) | 4.52 (2.87) | 89.41 (16.93) | 0.037 | 0.481 | 0.481 | 0 | 0.444 | 0.608 (0.157) | 0.701 (0.249) | 0.269 (0.24) |

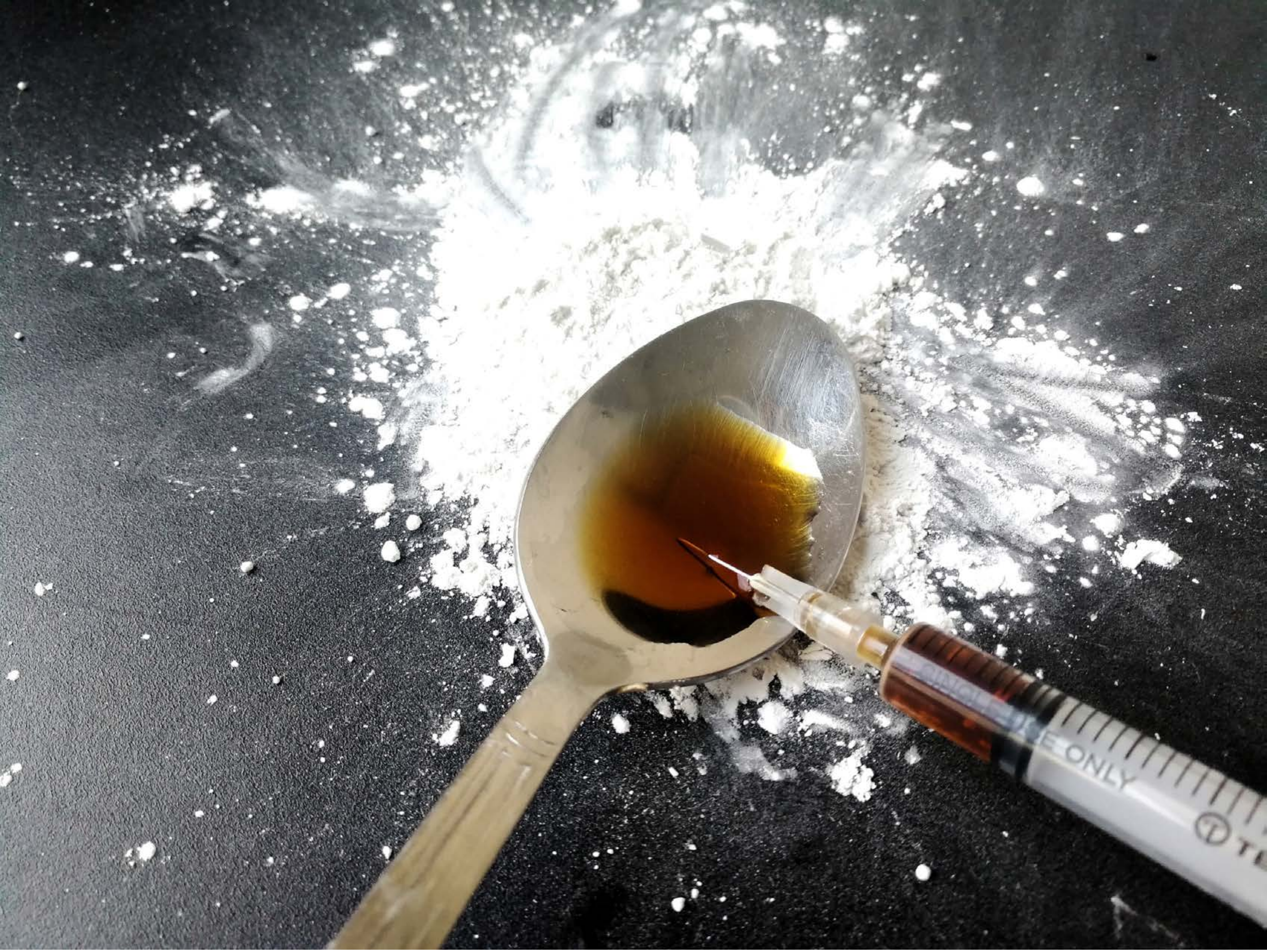


| Category | Craving (0-100) | Valence (1-9) | Arousal (1-9) | Typicality (0-100) | Relatedness | | | | | HSV | | |
|---|---|---|---|---|---|---|---|---|---|---|---|---|
| | | | | | Meth | Opioid | Both | Neither | MethToOpioid | Hue | Saturation | Value |
| Opioid Injection Instrument | 95.43 (10.05) | 4.14 (1.976) | 4.96 (2.9) | 95 (10.4) | 0.143 | 0.571 | 0.286 | 0 | 0.429 | 0.392 (0.265) | 0.097 (0.172) | 0.498 (0.29) |

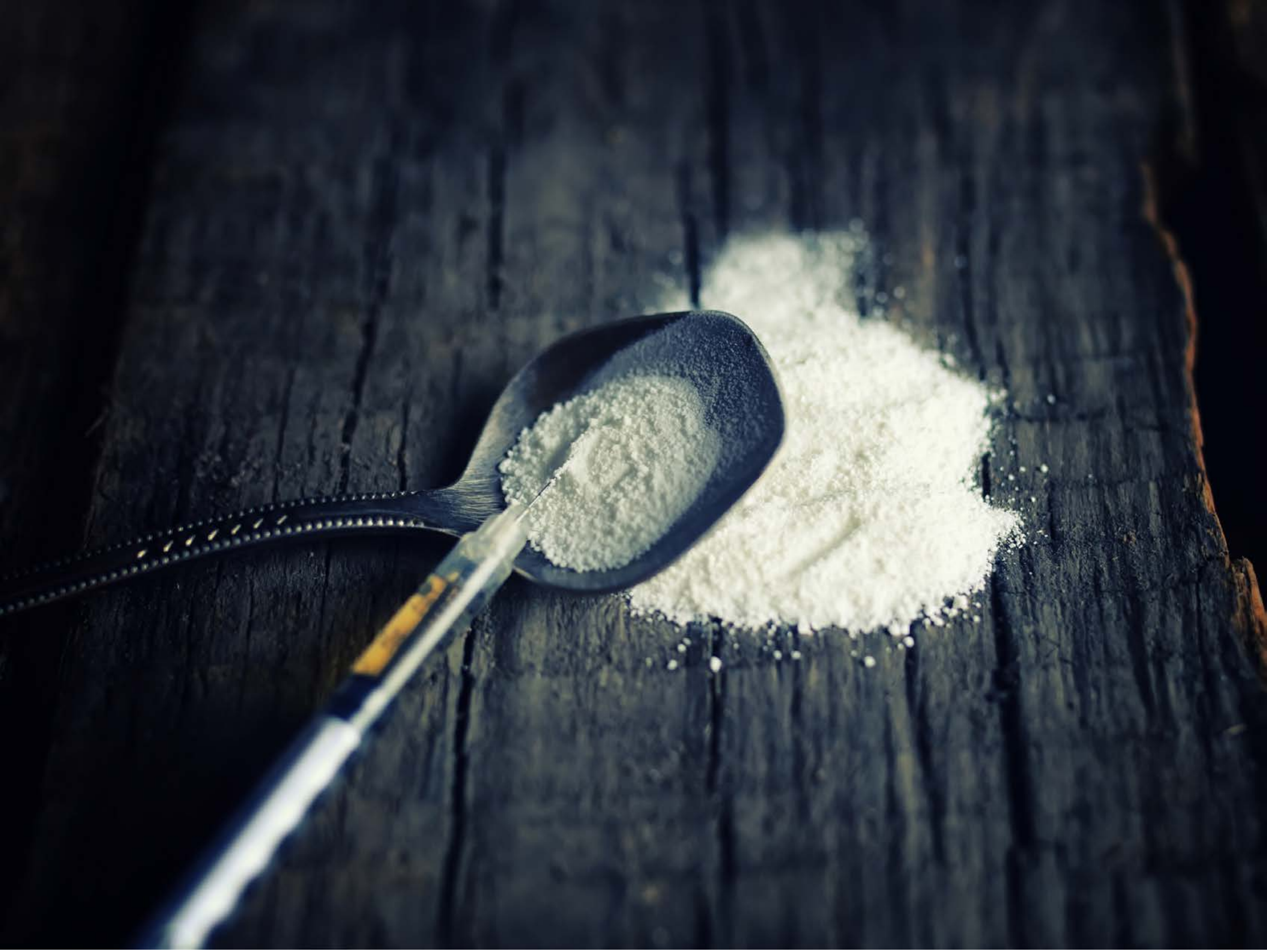

| Category | Craving (0-100) | Valence (1-9) | Arousal (1-9) | Typicality (0-100) | Relatedness | | | | | HSV | | |
|---|---|---|---|---|---|---|---|---|---|---|---|---|
| | | | | | Meth | Opioid | Both | Neither | MethToOpioid | Hue | Saturation | Value |
| Opioid Injection Instrument | 93.96 (10.81) | 4.46 (2.027) | 4.86 (2.93) | 94.79 (10.28) | 0.107 | 0.25 | 0.643 | 0 | 0.143 | 0.565 (0.141) | 0.4 (0.206) | 0.253 (0.227) |

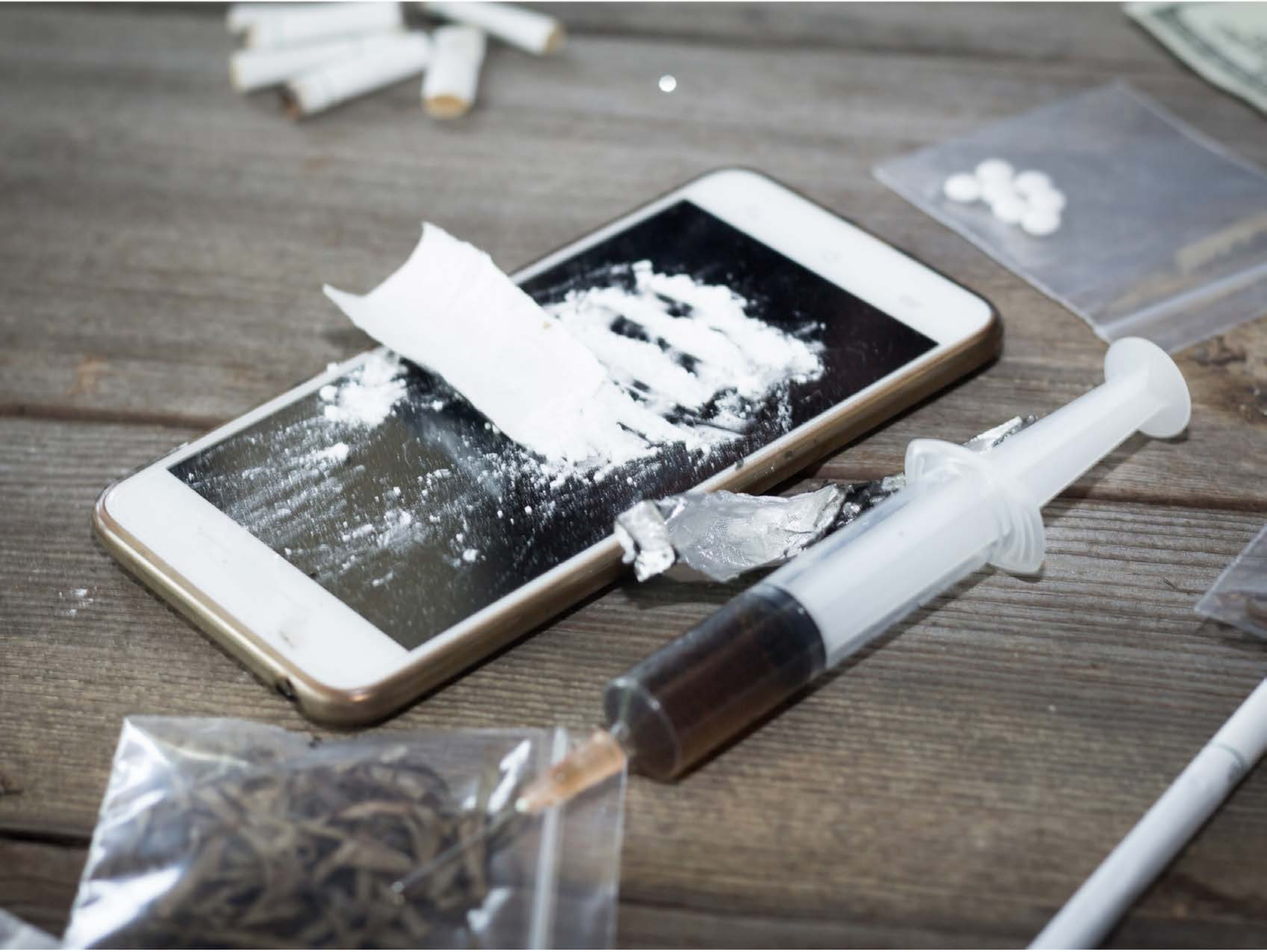

| Category | Craving (0-100) | Valence (1-9) | Arousal (1-9) | Typicality (0-100) | Relatedness | | | | | HSV | | |
|---|---|---|---|---|---|---|---|---|---|---|---|---|
| | | | | | Meth | Opioid | Both | Neither | MethToOpioid | Hue | Saturation | Value |
| Opioid Injection Instrument | 87 (17.75) | 4.33 (1.941) | 4.56 (2.58) | 86.48 (18.93) | 0.148 | 0.370 | 0.481 | 0 | 0.222 | 0.316 (0.265) | 0.1 (0.09) | 0.534 (0.221) |

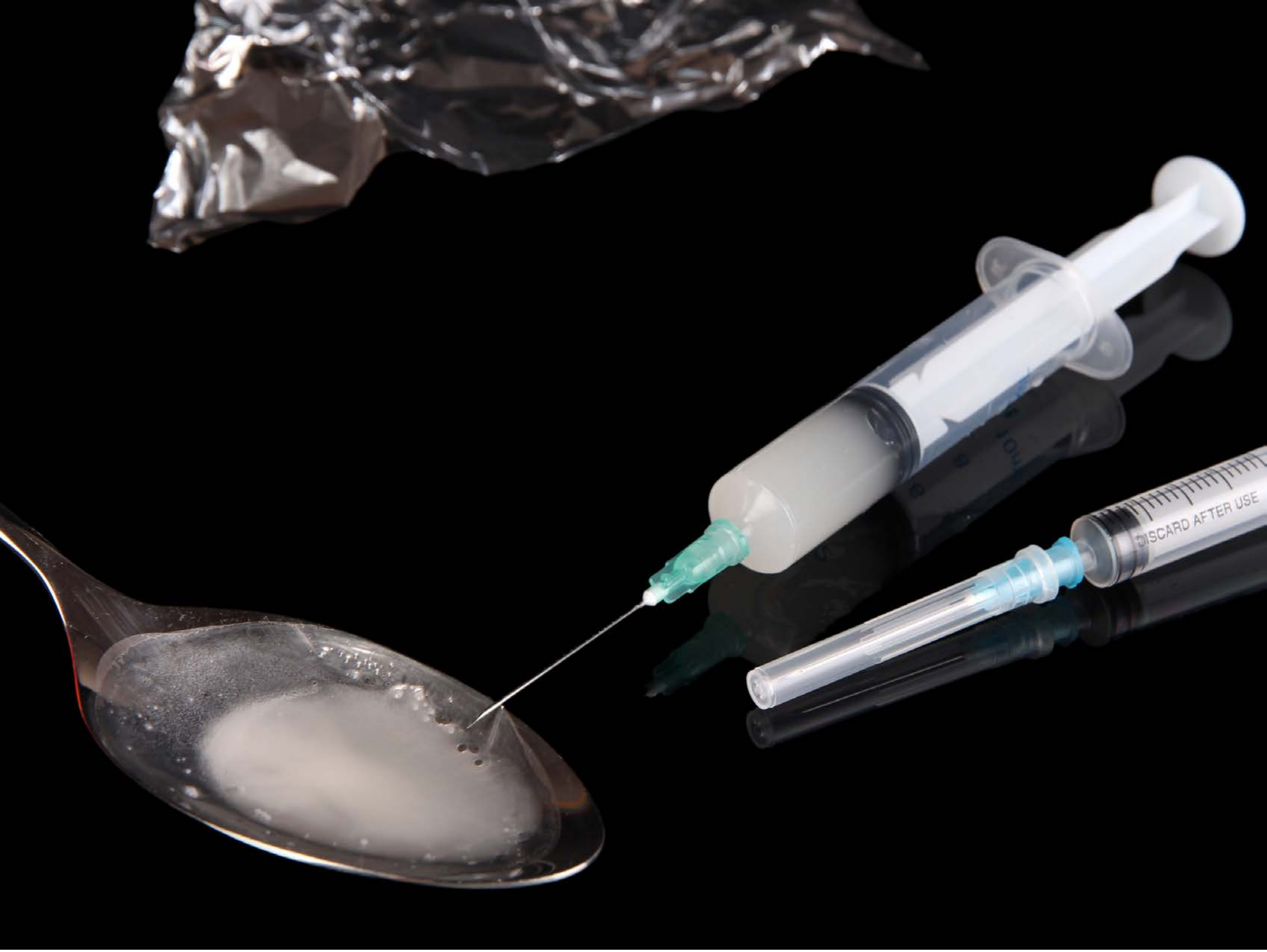


| Category | Craving (0-100) | Valence (1-9) | Arousal (1-9) | Typicality (0-100) | Relatedness | | | | | HSV | | |
|---|---|---|---|---|---|---|---|---|---|---|---|---|
| | | | | | Meth | Opioid | Both | Neither | MethToOpioid | Hue | Saturation | Value |
| Opioid Injection Instrument | 95 (6.64) | 4.25 (2.012) | 5.18 (3.09) | 96.75 (5.3) | 0.429 | 0.071 | 0.5 | 0 | -0.357 | 0.152 (0.297) | 0.046 (0.115) | 0.141 (0.256) |

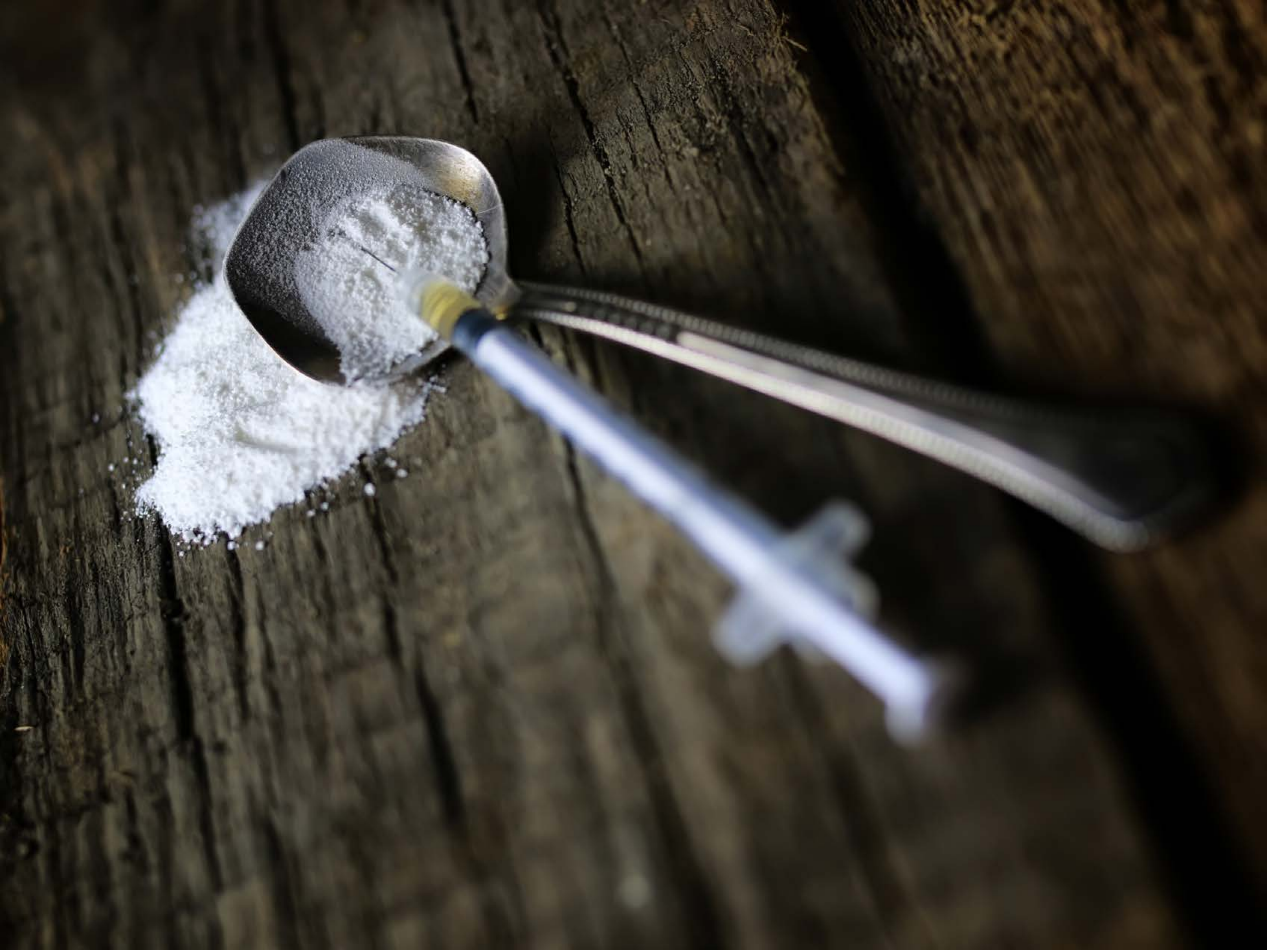

| Category | Craving (0-100) | Valence (1-9) | Arousal (1-9) | Typicality (0-100) | Relatedness | | | | | HSV | | |
|---|---|---|---|---|---|---|---|---|---|---|---|---|
| | | | | | Meth | Opioid | Both | Neither | MethToOpioid | Hue | Saturation | Value |
| Opioid Injection Instrument | 86.93 (17.4) | 4.44 (2.006) | 4.74 (2.75) | 87.19 (16.99) | 0.148 | 0.407 | 0.407 | 0.037 | 0.259 | 0.179 (0.179) | 0.333 (0.195) | 0.249 (0.213) |

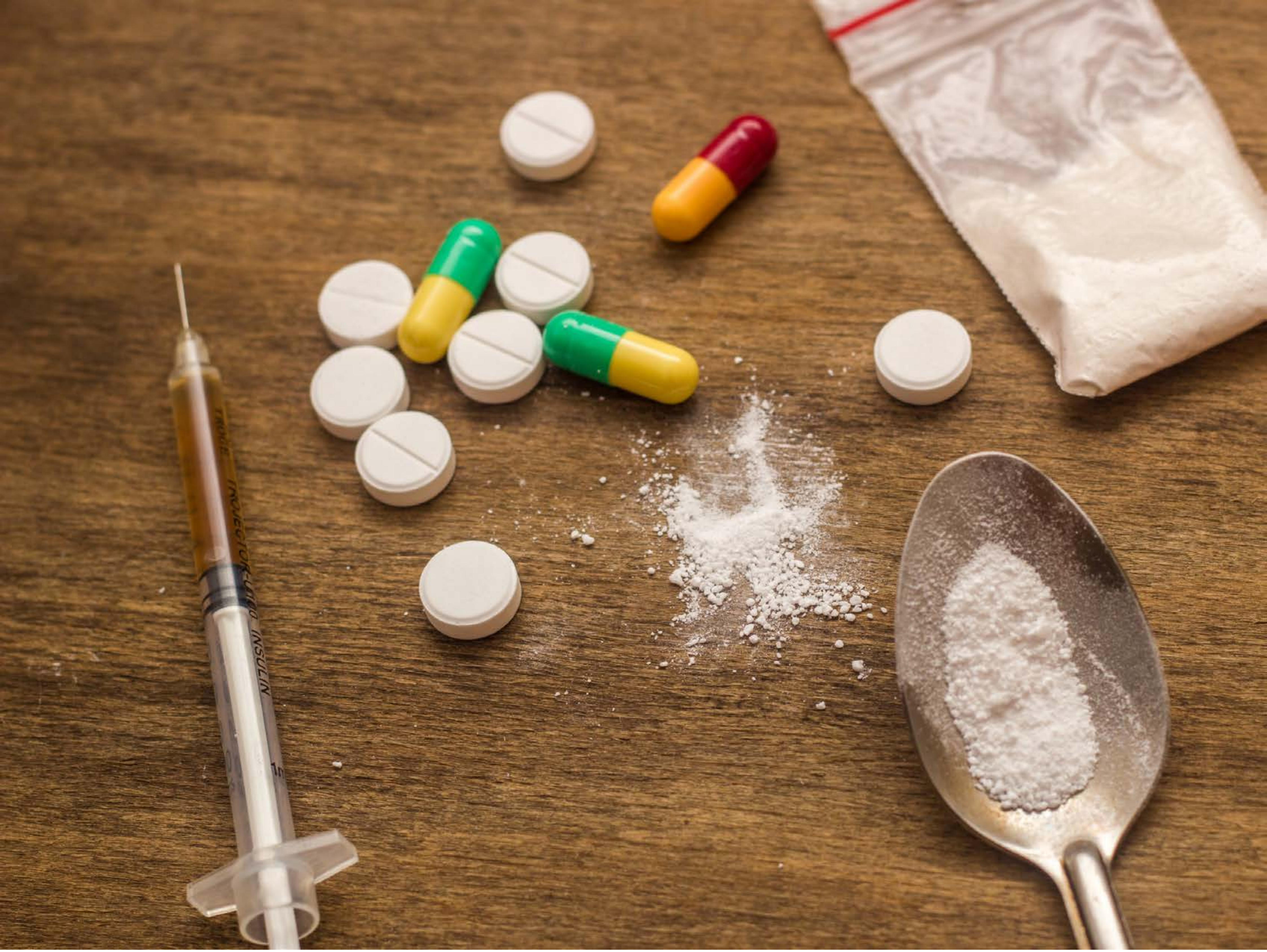


| Category | Craving (0-100) | Valence (1-9) | Arousal (1-9) | Typicality (0-100) | Relatedness | | | | | HSV | | |
|---|---|---|---|---|---|---|---|---|---|---|---|---|
| | | | | | Meth | Opioid | Both | Neither | MethToOpioid | Hue | Saturation | Value |
| Opioid Injection Instrument | 89.93 (14.6) | 4.29 (1.997) | 4.68 (2.91) | 91 (15.82) | 0 | 0.429 | 0.571 | 0 | 0.429 | 0.085 (0.056) | 0.501 (0.207) | 0.58 (0.178) |

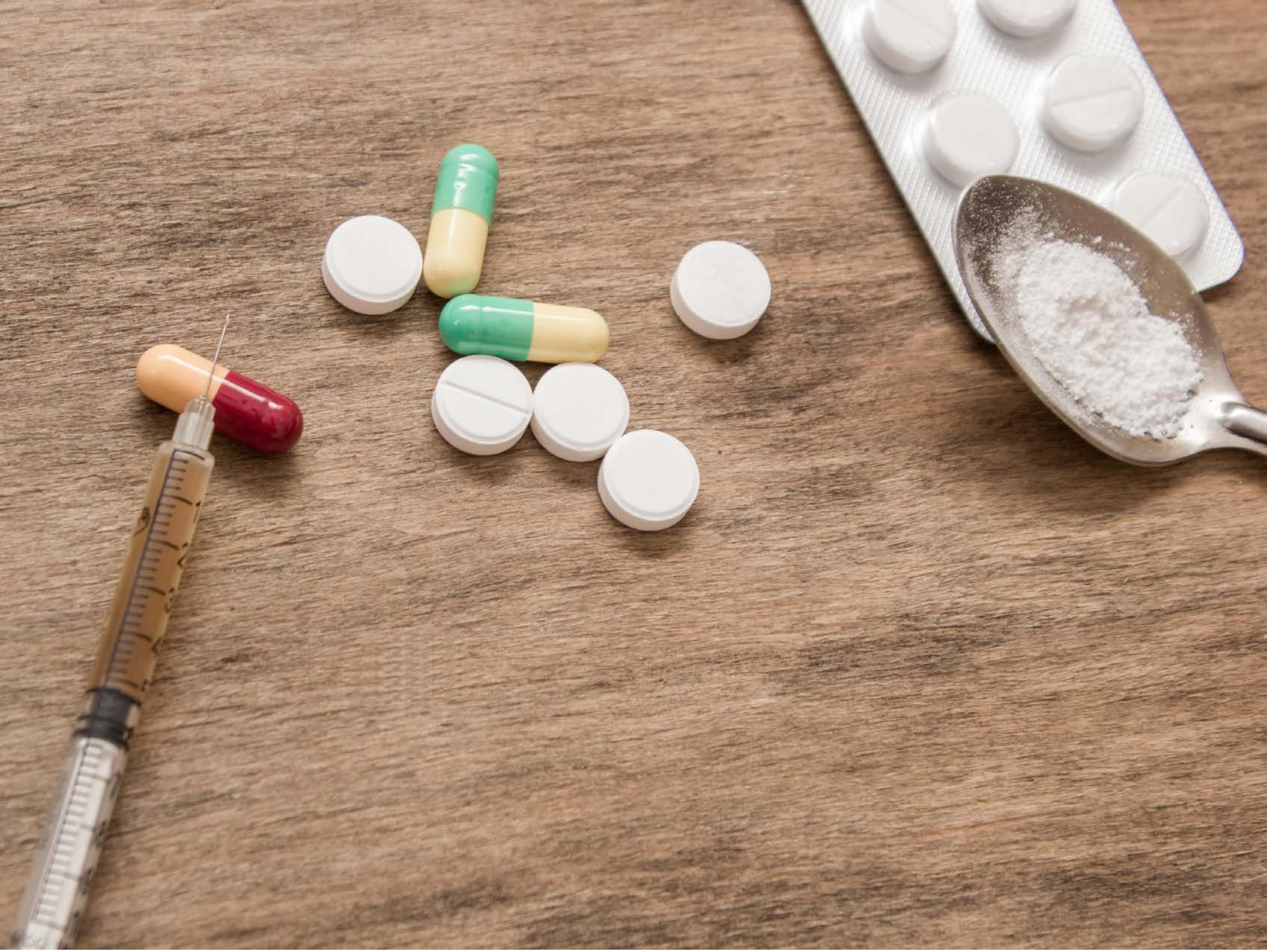

| Category | Craving (0-100) | Valence (1-9) | Arousal (1-9) | Typicality (0-100) | Relatedness | | | | | HSV | | |
|---|---|---|---|---|---|---|---|---|---|---|---|---|
| | | | | | Meth | Opioid | Both | Neither | MethToOpioid | Hue | Saturation | Value |
| Opioid Injection Instrument | 84.67 (23.89) | 4.41 (2.062) | 4.56 (2.79) | 85.81 (23.7) | 0 | 0.667 | 0.333 | 0 | 0.667 | 0.087 (0.092) | 0.32 (0.13) | 0.677 (0.132) |

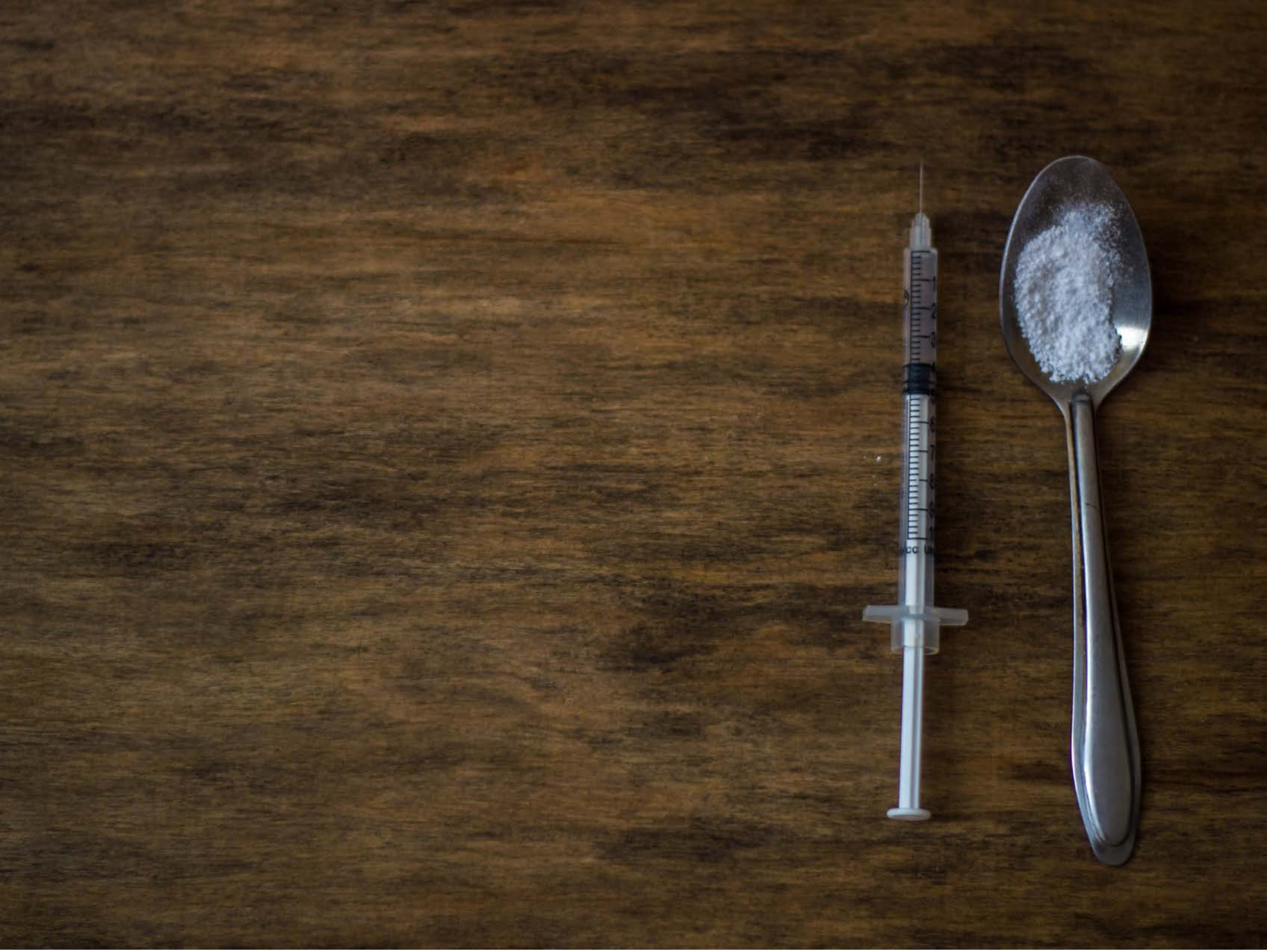

| Category | Craving (0-100) | Valence (1-9) | Arousal (1-9) | Typicality (0-100) | Relatedness | | | | | HSV | | |
|---|---|---|---|---|---|---|---|---|---|---|---|---|
| | | | | | Meth | Opioid | Both | Neither | MethToOpioid | Hue | Saturation | Value |
| Opioid Injection Instrument | 89.67 (15.92) | 4.41 (2.099) | 4.7 (2.93) | 91.63 (14.48) | 0.333 | 0.148 | 0.481 | 0.037 | -0.185 | 0.111 (0.119) | 0.537 (0.126) | 0.29 (0.101) |

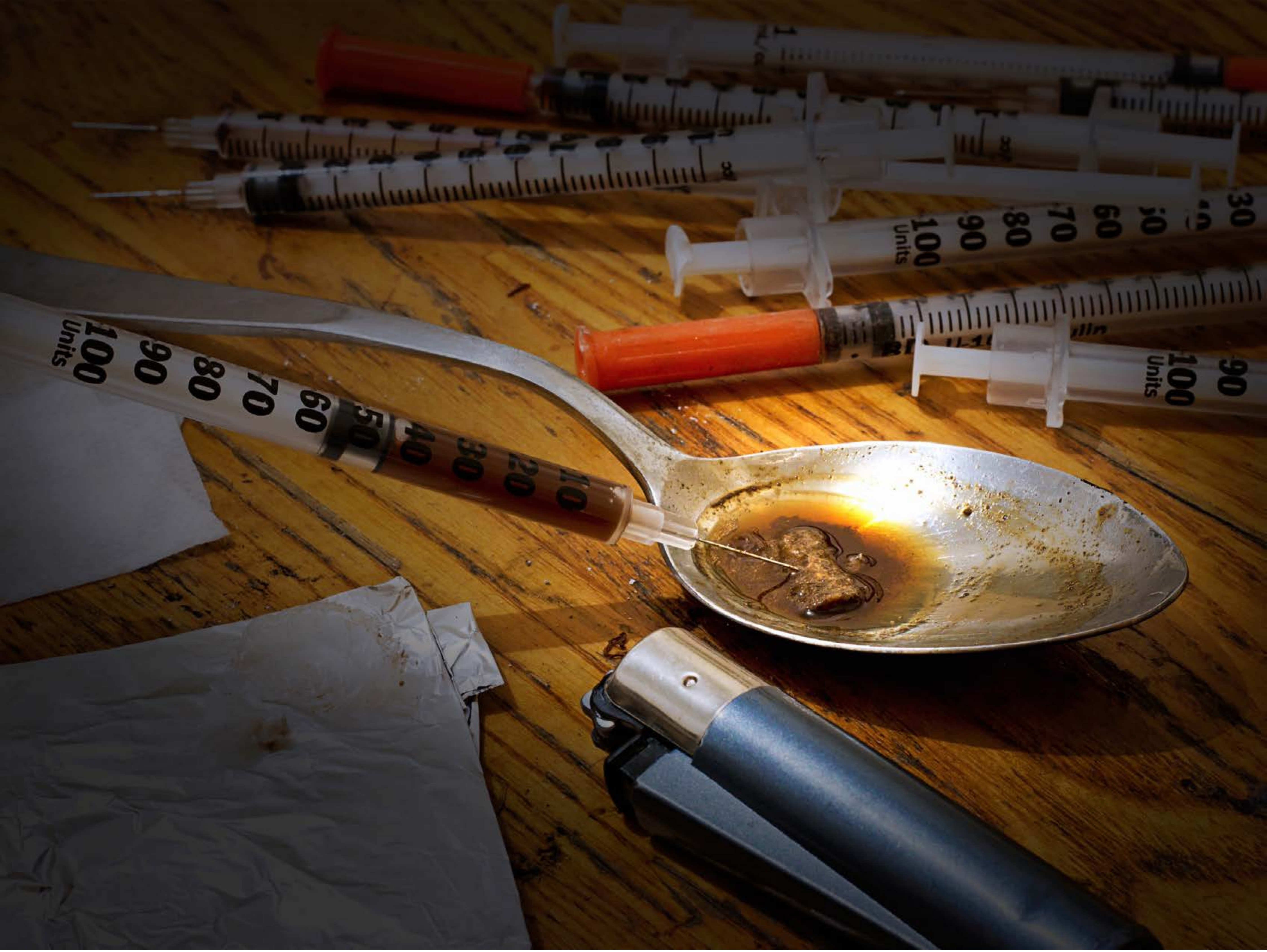


| Category | Craving (0-100) | Valence (1-9) | Arousal (1-9) | Typicality (0-100) | Relatedness | | | | | HSV | | |
|---|---|---|---|---|---|---|---|---|---|---|---|---|
| | | | | | Meth | Opioid | Both | Neither | MethToOpioid | Hue | Saturation | Value |
| Opioid Injection Instrument | 86.67 (18.84) | 4.26 (1.913) | 4.96 (2.65) | 87.3 (18.95) | 0.074 | 0.704 | 0.222 | 0 | 0.630 | 0.258 (0.306) | 0.507 (0.357) | 0.298 (0.201) |

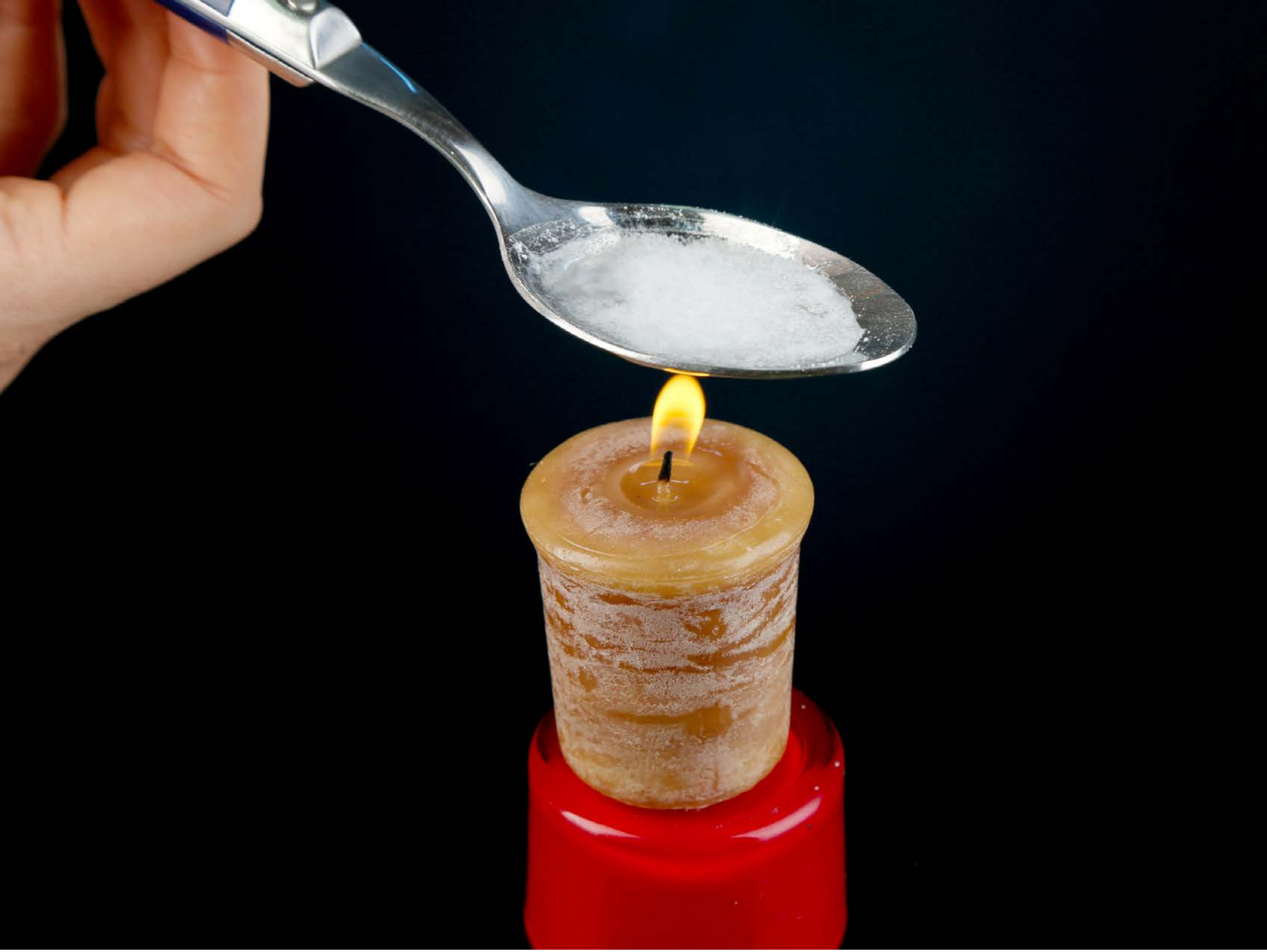

| Category | Craving (0-100) | Valence (1-9) | Arousal (1-9) | Typicality (0-100) | Relatedness | | | | | HSV | | |
|---|---|---|---|---|---|---|---|---|---|---|---|---|
| | | | | | Meth | Opioid | Both | Neither | MethToOpioid | Hue | Saturation | Value |
| Opioid Injection Instrument | 92.46 (13.42) | 4.54 (2.009) | 4.71 (2.84) | 94.18 (11.24) | 0.107 | 0.464 | 0.429 | 0 | 0.357 | 0.256 (0.302) | 0.437 (0.448) | 0.191 (0.297) |

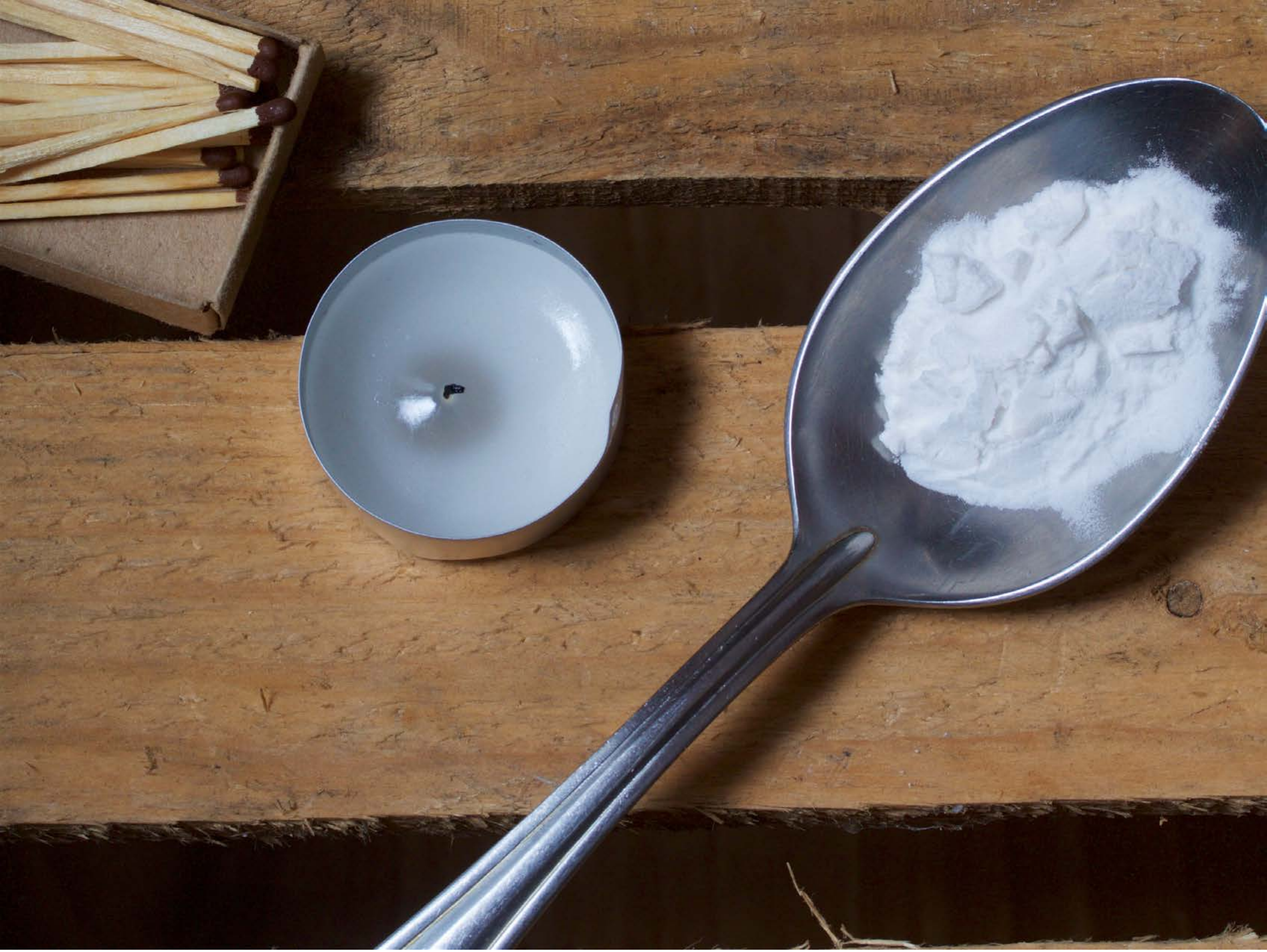

| Category | Craving (0-100) | Valence (1-9) | Arousal (1-9) | Typicality (0-100) | Relatedness | | | | | HSV | | |
|---|---|---|---|---|---|---|---|---|---|---|---|---|
| | | | | | Meth | Opioid | Both | Neither | MethToOpioid | Hue | Saturation | Value |
| Opioid Injection Instrument | 76.96 (31.13) | 4.56 (1.577) | 4.52 (2.5) | 82.19 (24.81) | 0.148 | 0.556 | 0.185 | 0.111 | 0.407 | 0.217 (0.245) | 0.466 (0.24) | 0.48 (0.241) |

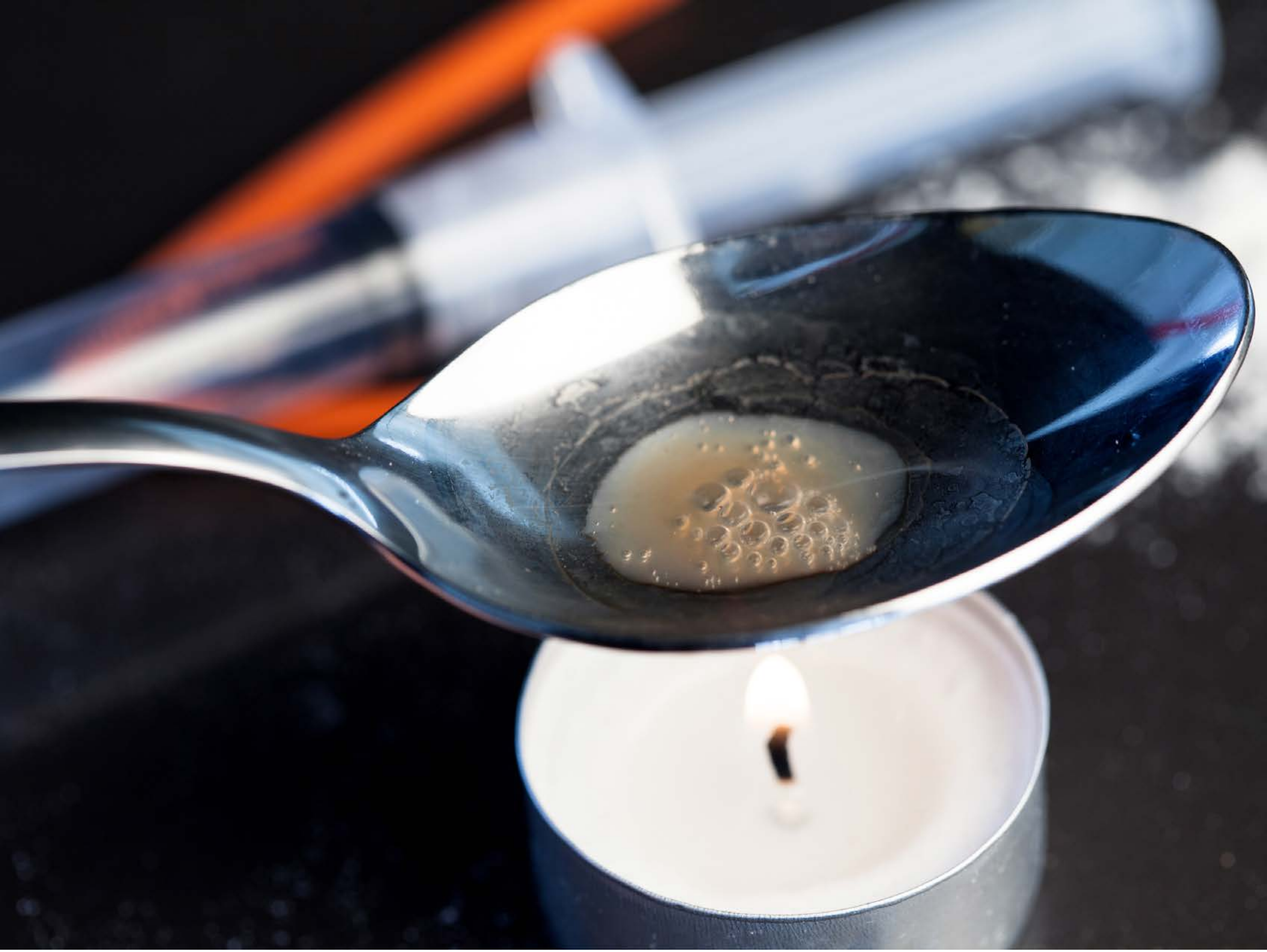

| Category | Craving (0-100) | Valence (1-9) | Arousal (1-9) | Typicality (0-100) | Relatedness | | | | | HSV | | |
|---|---|---|---|---|---|---|---|---|---|---|---|---|
| | | | | | Meth | Opioid | Both | Neither | MethToOpioid | Hue | Saturation | Value |
| Opioid Injection Instrument | 93.54 (10.03) | 4.14 (1.957) | 4.79 (2.71) | 93.79 (9.19) | 0.071 | 0.5 | 0.429 | 0 | 0.429 | 0.532 (0.293) | 0.275 (0.273) | 0.426 (0.346) |

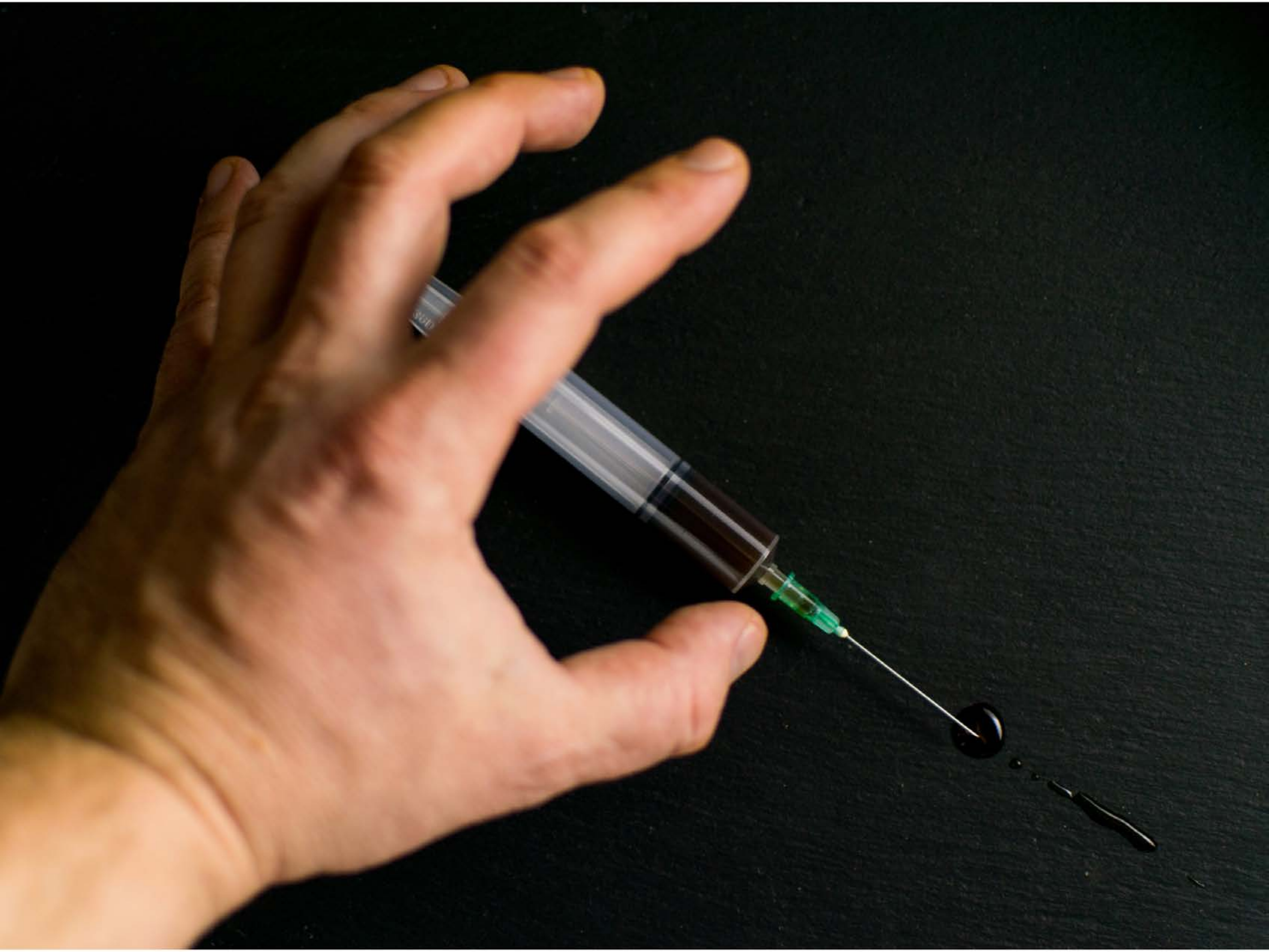

| Category | Craving (0-100) | Valence (1-9) | Arousal (1-9) | Typicality (0-100) | Relatedness | | | | | HSV | | |
|---|---|---|---|---|---|---|---|---|---|---|---|---|
| | | | | | Meth | Opioid | Both | Neither | MethToOpioid | Hue | Saturation | Value |
| Opioid Instrument Hand | 91.29 (13.93) | 4.36 (1.85) | 4.71 (2.85) | 92 (13.84) | 0.071 | 0.179 | 0.75 | 0 | 0.107 | 0.251 (0.191) | 0.375 (0.268) | 0.27 (0.323) |

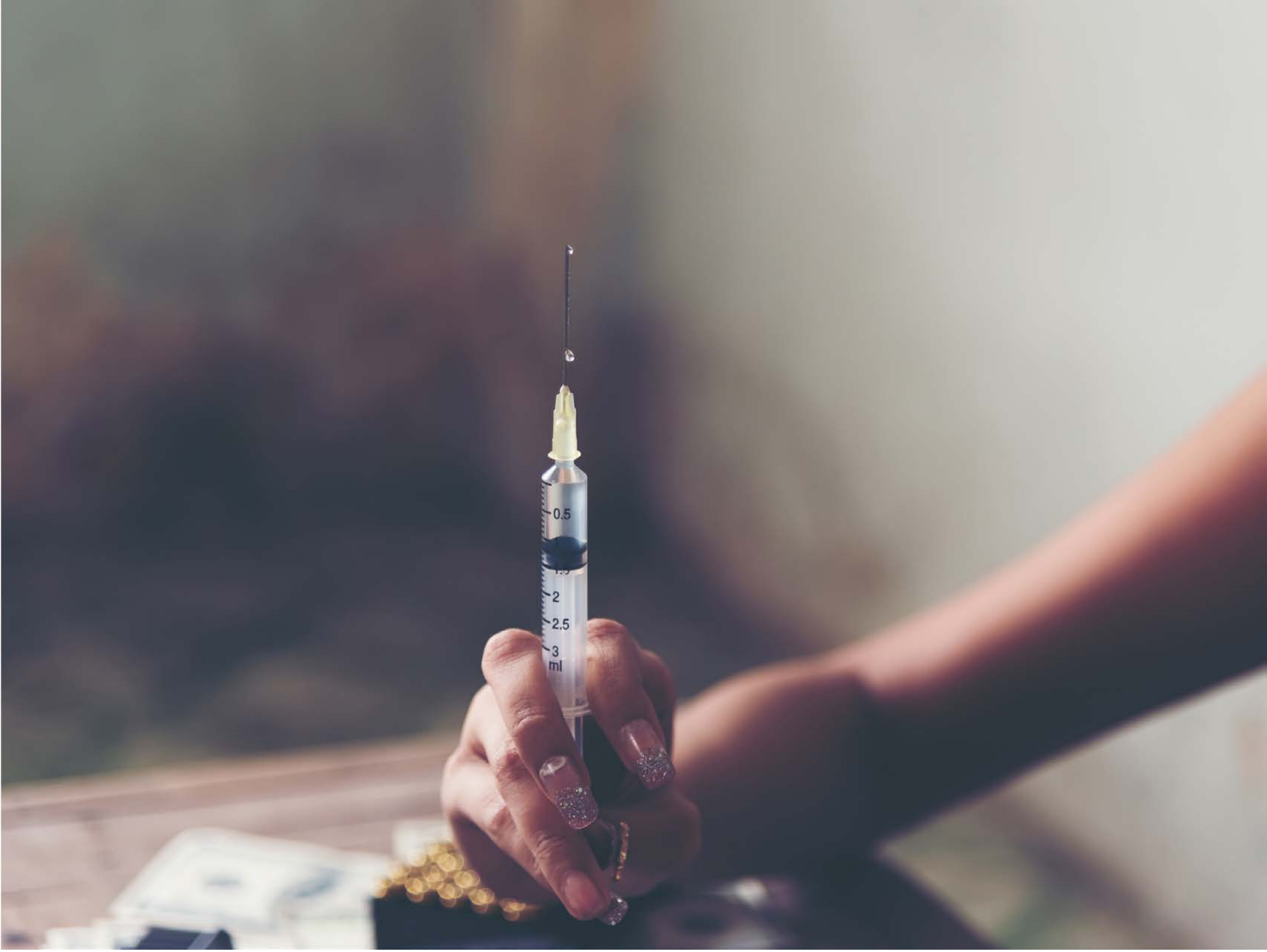


| Category | Craving (0-100) | Valence (1-9) | Arousal (1-9) | Typicality (0-100) | Relatedness | | | | | HSV | | |
|---|---|---|---|---|---|---|---|---|---|---|---|---|
| | | | | | Meth | Opioid | Both | Neither | MethToOpioid | Hue | Saturation | Value |
| Opioid Instrument Hand | 88.85 (14.81) | 4.44 (2.063) | 4.85 (3.02) | 89.7 (16.01) | 0.148 | 0.074 | 0.778 | 0 | -0.074 | 0.396 (0.327) | 0.133 (0.093) | 0.523 (0.214) |

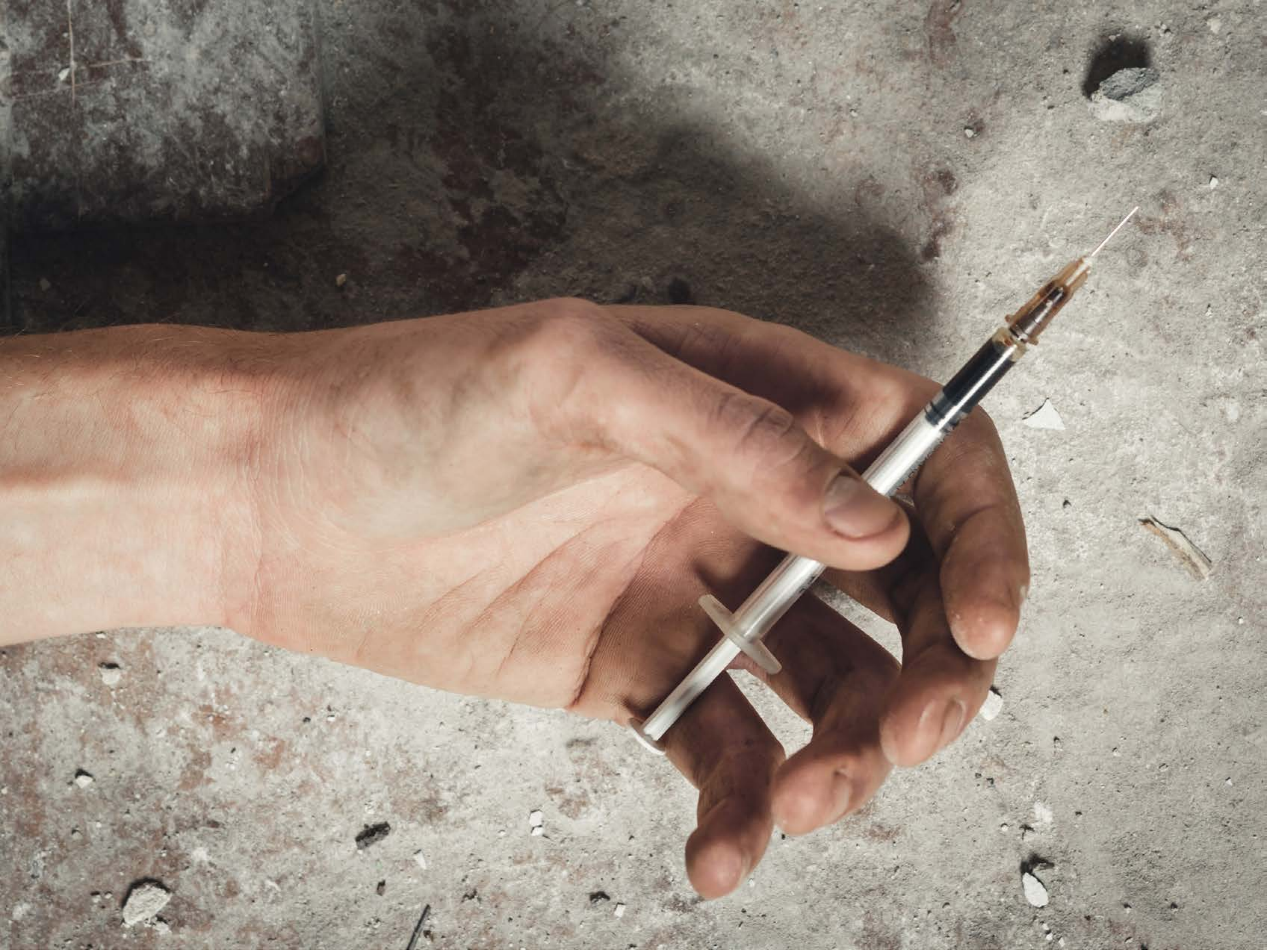

| Category | Craving (0-100) | Valence (1-9) | Arousal (1-9) | Typicality (0-100) | Relatedness | | | | | HSV | | |
|---|---|---|---|---|---|---|---|---|---|---|---|---|
| | | | | | Meth | Opioid | Both | Neither | MethToOpioid | Hue | Saturation | Value |
| Opioid Instrument Hand | 89.44 (15.19) | 4.15 (1.975) | 4.74 (2.63) | 88.96 (16.29) | 0.037 | 0.333 | 0.593 | 0.037 | 0.296 | 0.115 (0.137) | 0.178 (0.123) | 0.592 (0.228) |

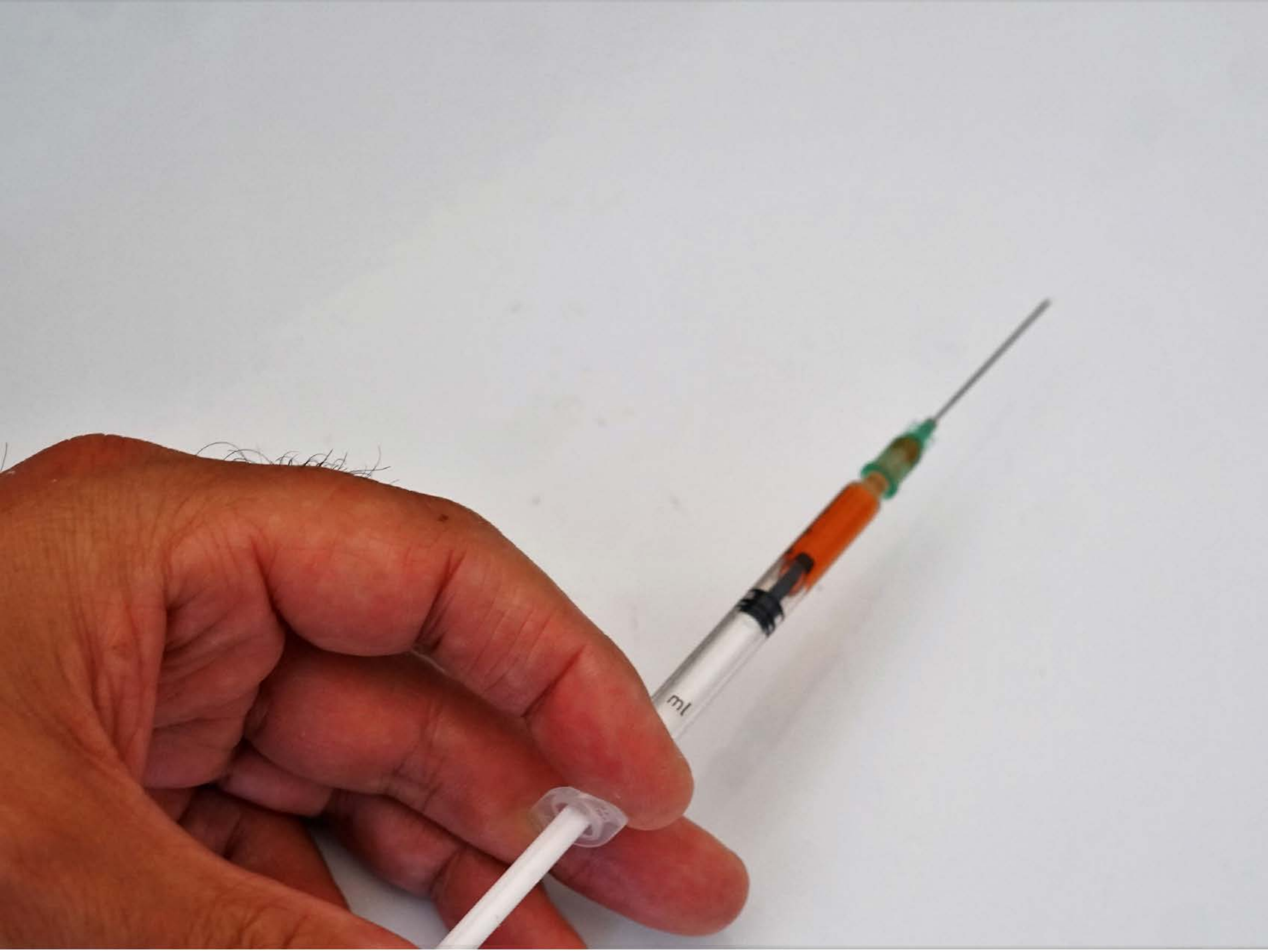


| | | | | | Relatedness | | | | | HSV | | |
|---|---|---|---|---|---|---|---|---|---|---|---|---|
| **Category** | **Craving (0-100)** | **Valence (1-9)** | **Arousal (1-9)** | **Typicality (0-100)** | Meth | Opioid | Both | Neither | MethToOpioid | Hue | Saturation | Value |
| Opioid Instrument Hand | 81.33 (21.15) | 4.3 (1.564) | 4.41 (2.37) | 82.15 (21.16) | 0.074 | 0.148 | 0.741 | 0.037 | 0.074 | 0.369 (0.35) | 0.188 (0.301) | 0.733 (0.143) |

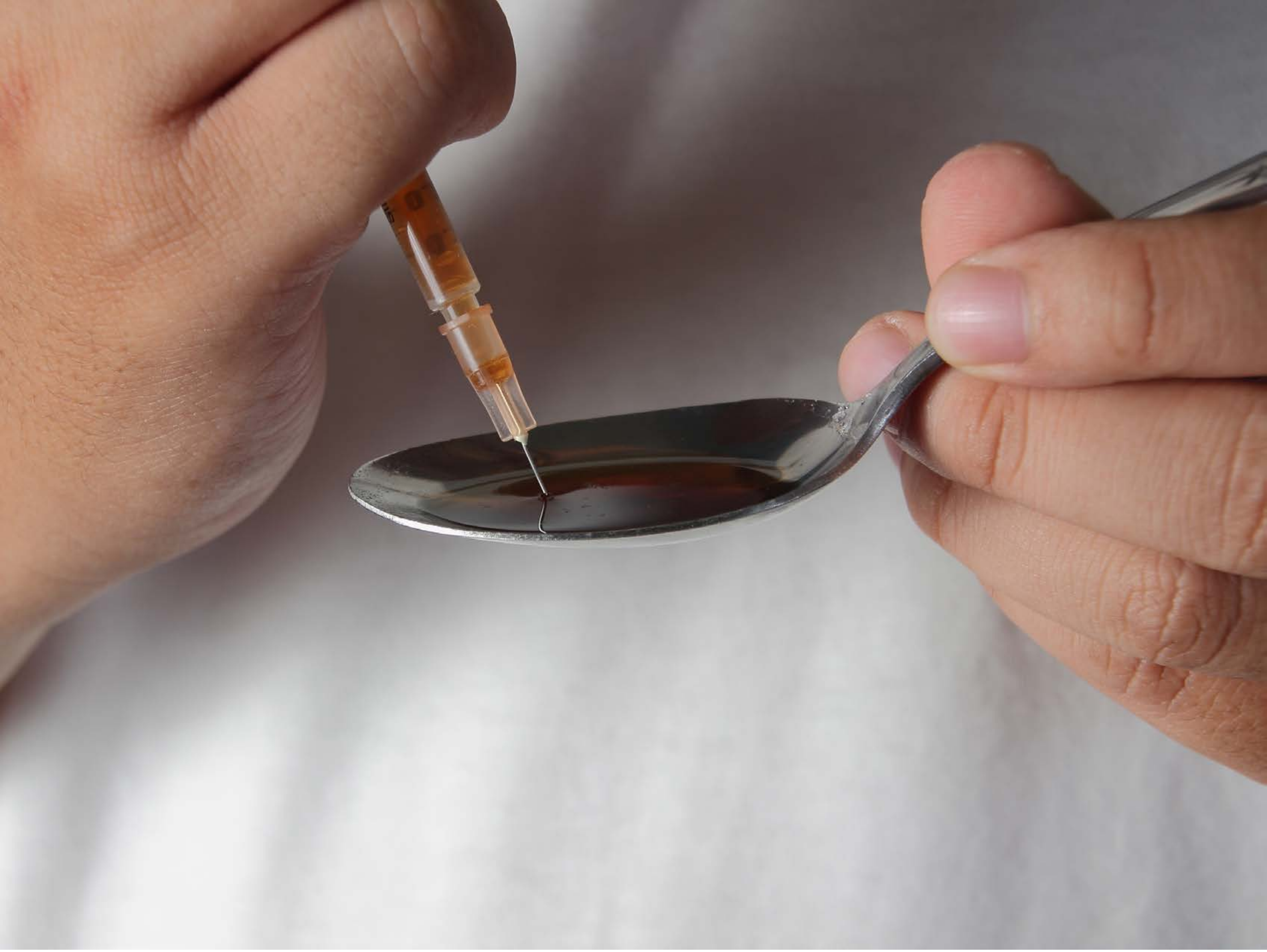

| Category | Craving (0-100) | Valence (1-9) | Arousal (1-9) | Typicality (0-100) | Relatedness | | | | | HSV | | |
|---|---|---|---|---|---|---|---|---|---|---|---|---|
| | | | | | Meth | Opioid | Both | Neither | MethToOpioid | Hue | Saturation | Value |
| Opioid Instrument Hand | 91.32 (20.15) | 4.07 (2.071) | 4.64 (2.79) | 92.57 (19.67) | 0.071 | 0.429 | 0.5 | 0 | 0.357 | 0.21 (0.321) | 0.182 (0.185) | 0.612 (0.2) |

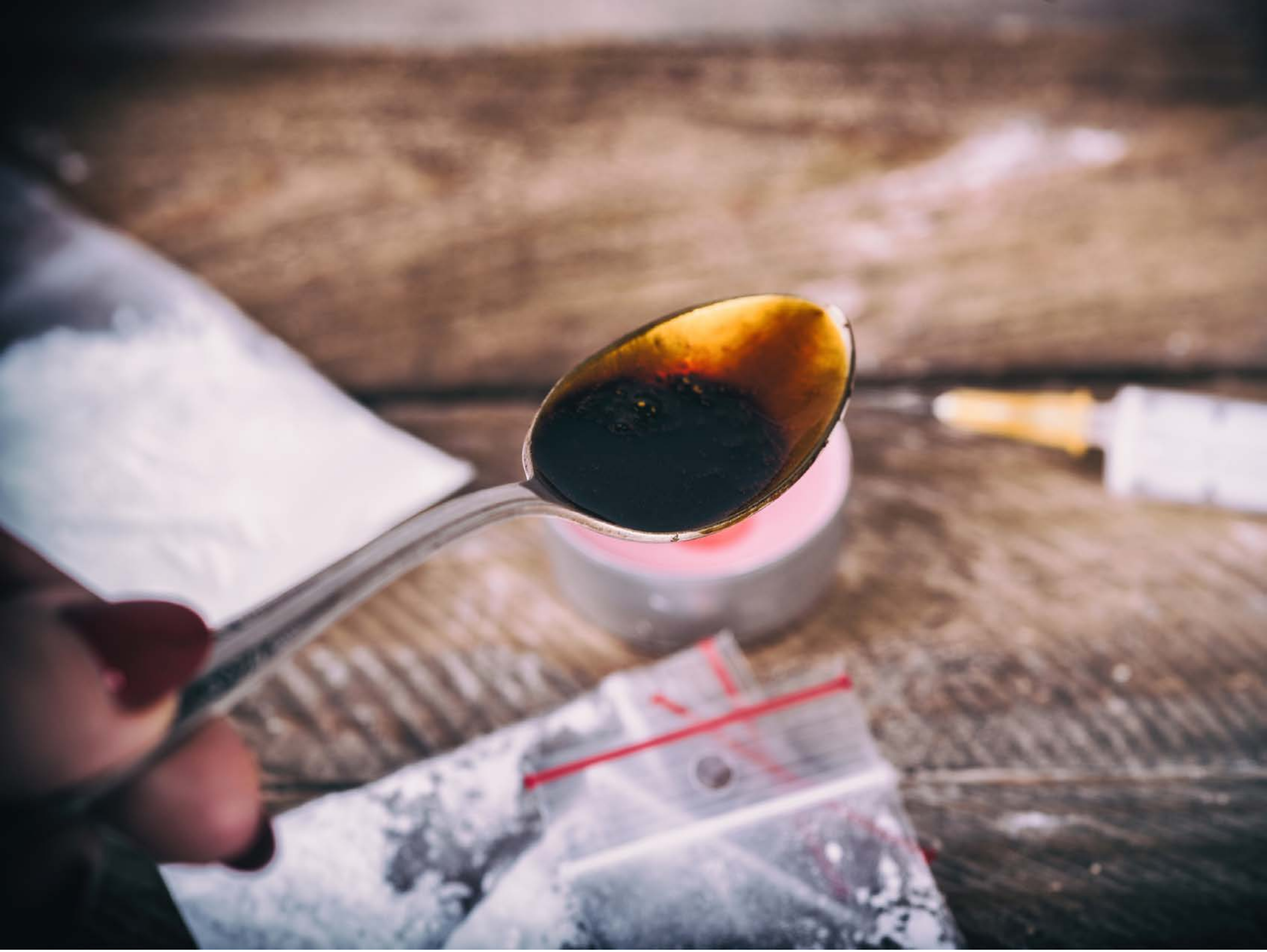

| Category | Craving (0-100) | Valence (1-9) | Arousal (1-9) | Typicality (0-100) | Relatedness | | | | | HSV | | |
|---|---|---|---|---|---|---|---|---|---|---|---|---|
| | | | | | Meth | Opioid | Both | Neither | MethToOpioid | Hue | Saturation | Value |
| Opioid Instrument Hand | 85.44 (22.66) | 4.56 (1.805) | 4.81 (2.6) | 88.37 (16.86) | 0.074 | 0.704 | 0.222 | 0 | 0.630 | 0.393 (0.384) | 0.24 (0.152) | 0.584 (0.237) |

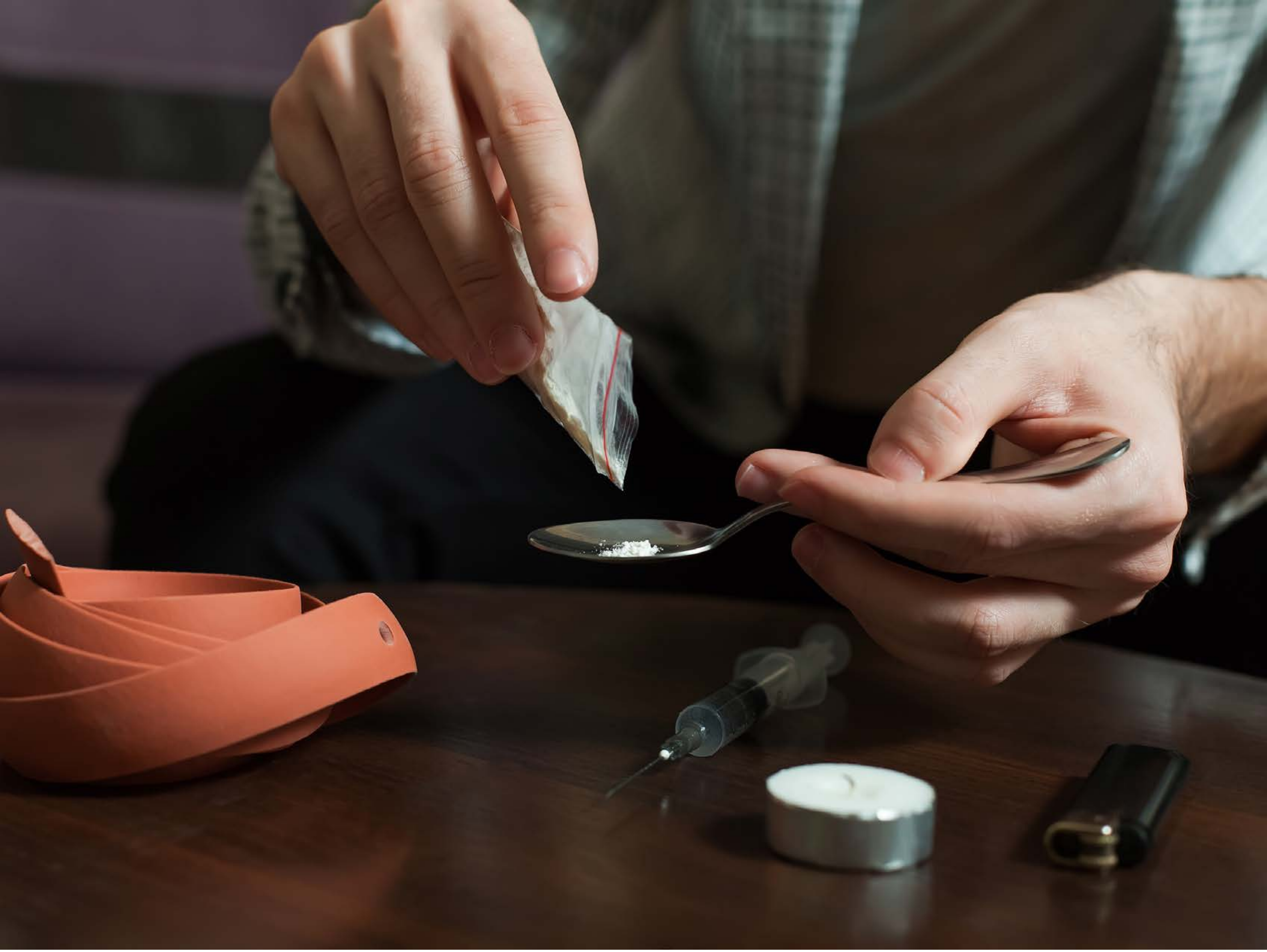

| Category | Craving (0-100) | Valence (1-9) | Arousal (1-9) | Typicality (0-100) | Relatedness | | | | | HSV | | |
|---|---|---|---|---|---|---|---|---|---|---|---|---|
| | | | | | Meth | Opioid | Both | Neither | MethToOpioid | Hue | Saturation | Value |
| Opioid Instrument Hand | 91.18 (15.32) | 4.36 (1.66) | 4.82 (2.76) | 92.89 (13.15) | 0.107 | 0.429 | 0.464 | 0 | 0.321 | 0.2 (0.28) | 0.383 (0.213) | 0.277 (0.233) |

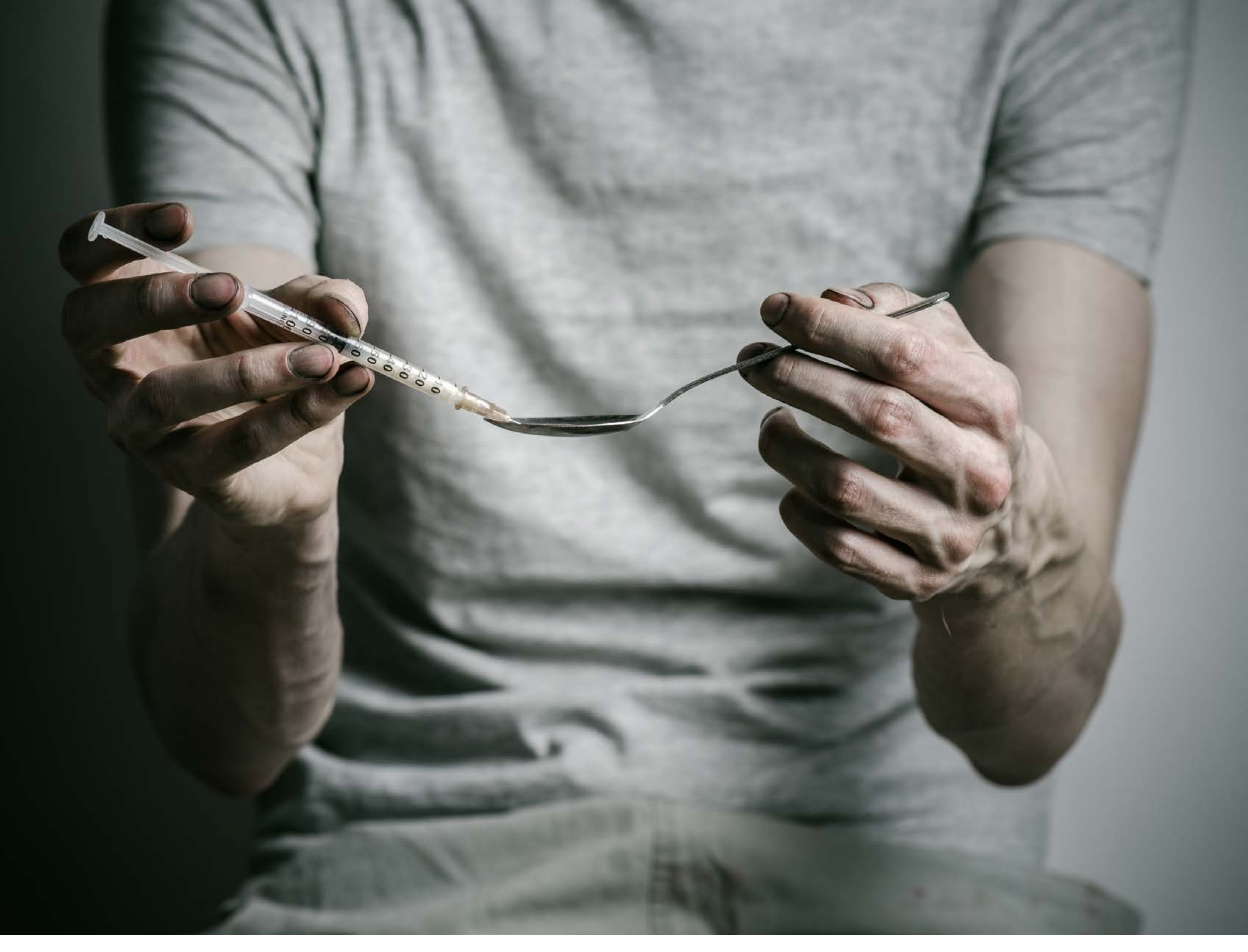

| Category | Craving (0-100) | Valence (1-9) | Arousal (1-9) | Typicality (0-100) | Relatedness | | | | | HSV | | |
|---|---|---|---|---|---|---|---|---|---|---|---|---|
| | | | | | Meth | Opioid | Both | Neither | MethToOpioid | Hue | Saturation | Value |
| Opioid Instrument Hand | 92.79 (11.62) | 4.36 (1.929) | 4.75 (2.81) | 94.14 (10.91) | 0.107 | 0.071 | 0.821 | 0 | -0.036 | 0.367 (0.161) | 0.135 (0.139) | 0.46 (0.238) |

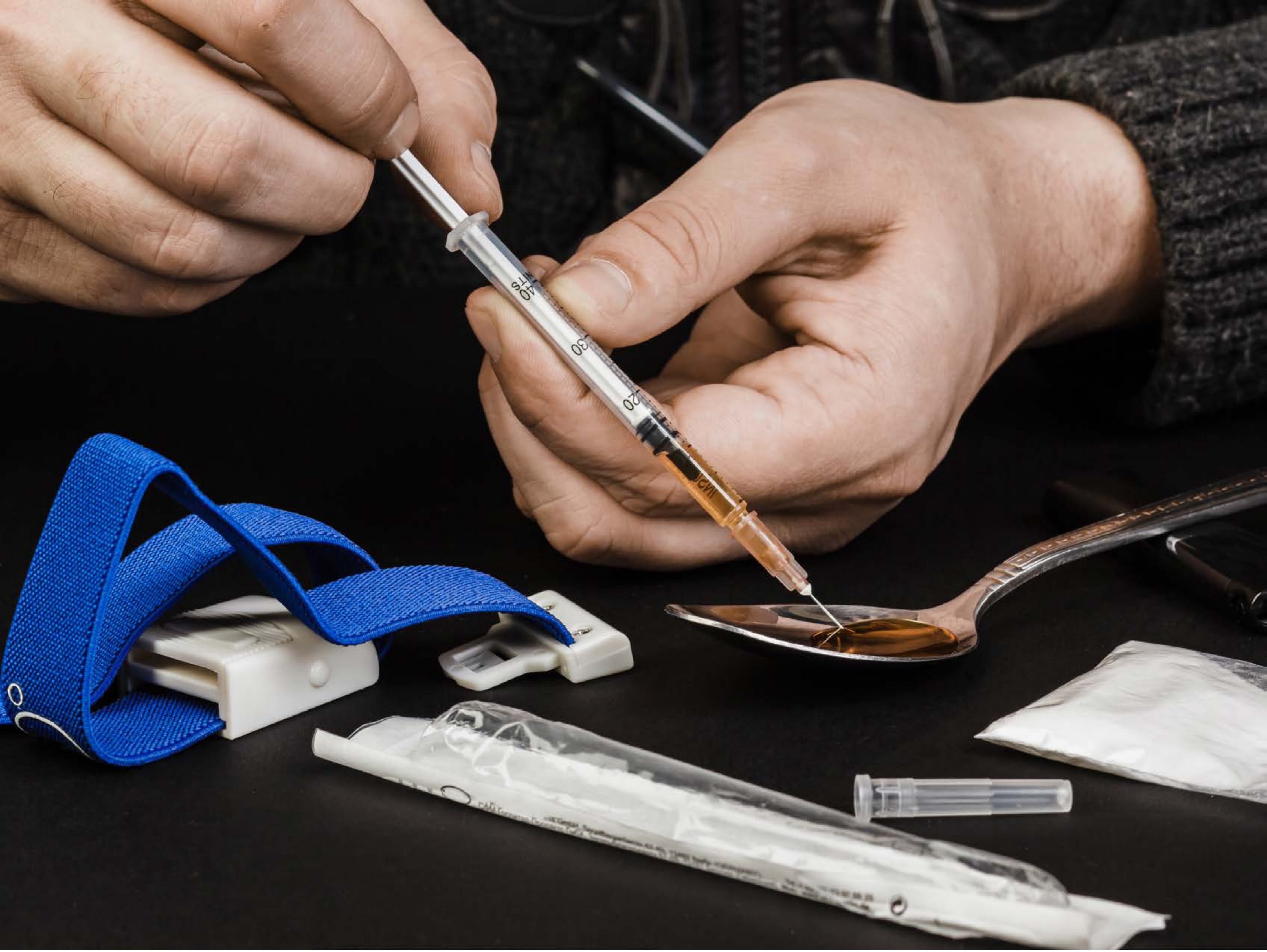

| | | | | | Relatedness | | | | | HSV | | |
|---|---|---|---|---|---|---|---|---|---|---|---|---|
| **Category** | **Craving (0-100)** | **Valence (1-9)** | **Arousal (1-9)** | **Typicality (0-100)** | Meth | Opioid | Both | Neither | MethToOpioid | Hue | Saturation | Value |
| Opioid Instrument Hand | 91.61 (14.5) | 4.5 (1.895) | 4.79 (2.71) | 94.32 (10.05) | 0.071 | 0.357 | 0.571 | 0 | 0.286 | 0.273 (0.355) | 0.235 (0.229) | 0.366 (0.325) |

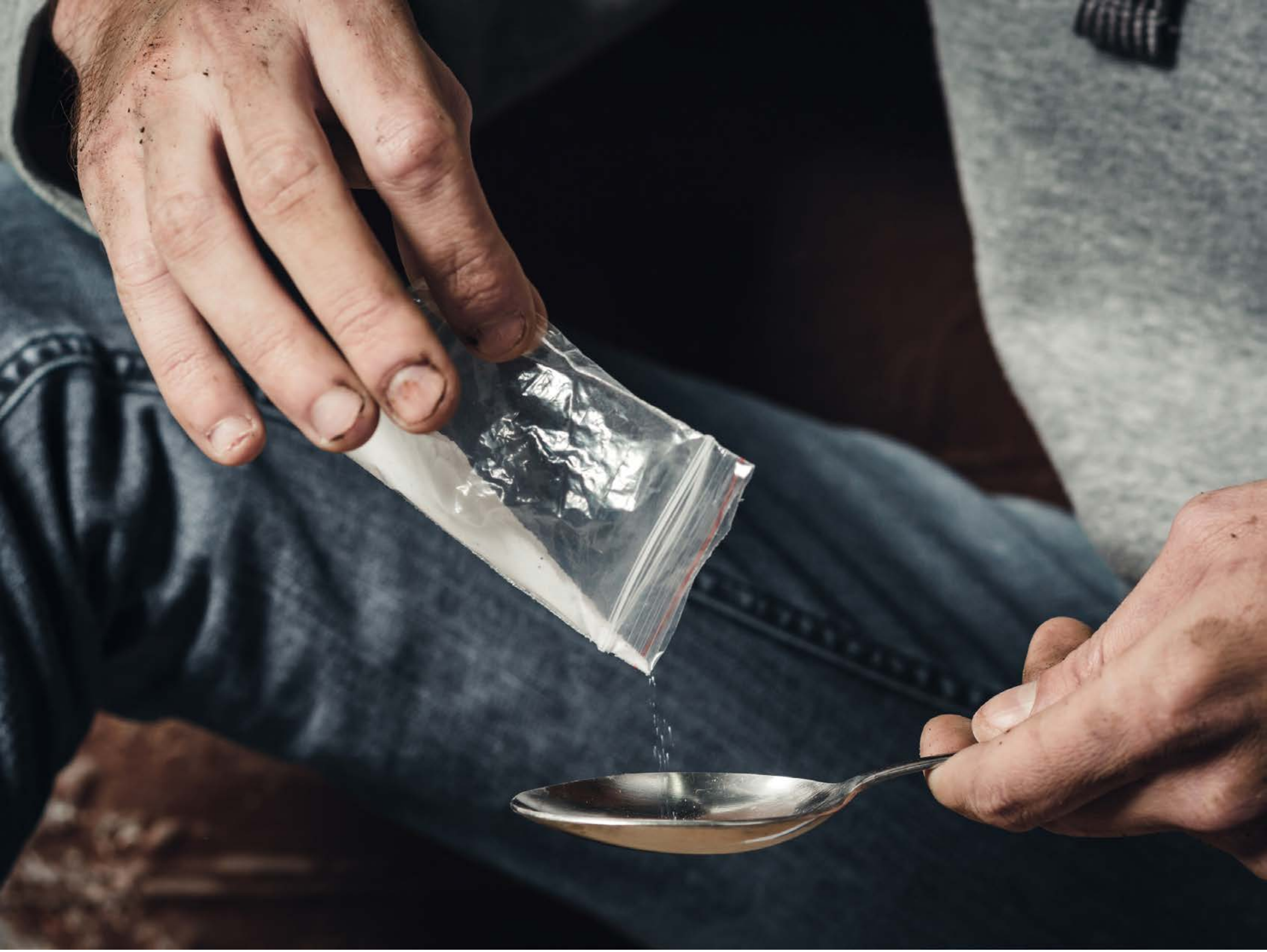

| Category | Craving (0-100) | Valence (1-9) | Arousal (1-9) | Typicality (0-100) | Relatedness | | | | | HSV | | |
|---|---|---|---|---|---|---|---|---|---|---|---|---|
| | | | | | Meth | Opioid | Both | Neither | MethToOpioid | Hue | Saturation | Value |
| Opioid Instrument Hand | 85.67 (18.3) | 4.41 (1.866) | 4.93 (2.57) | 86.93 (18.49) | 0.370 | 0.222 | 0.407 | 0 | -0.148 | 0.442 (0.275) | 0.267 (0.132) | 0.349 (0.269) |

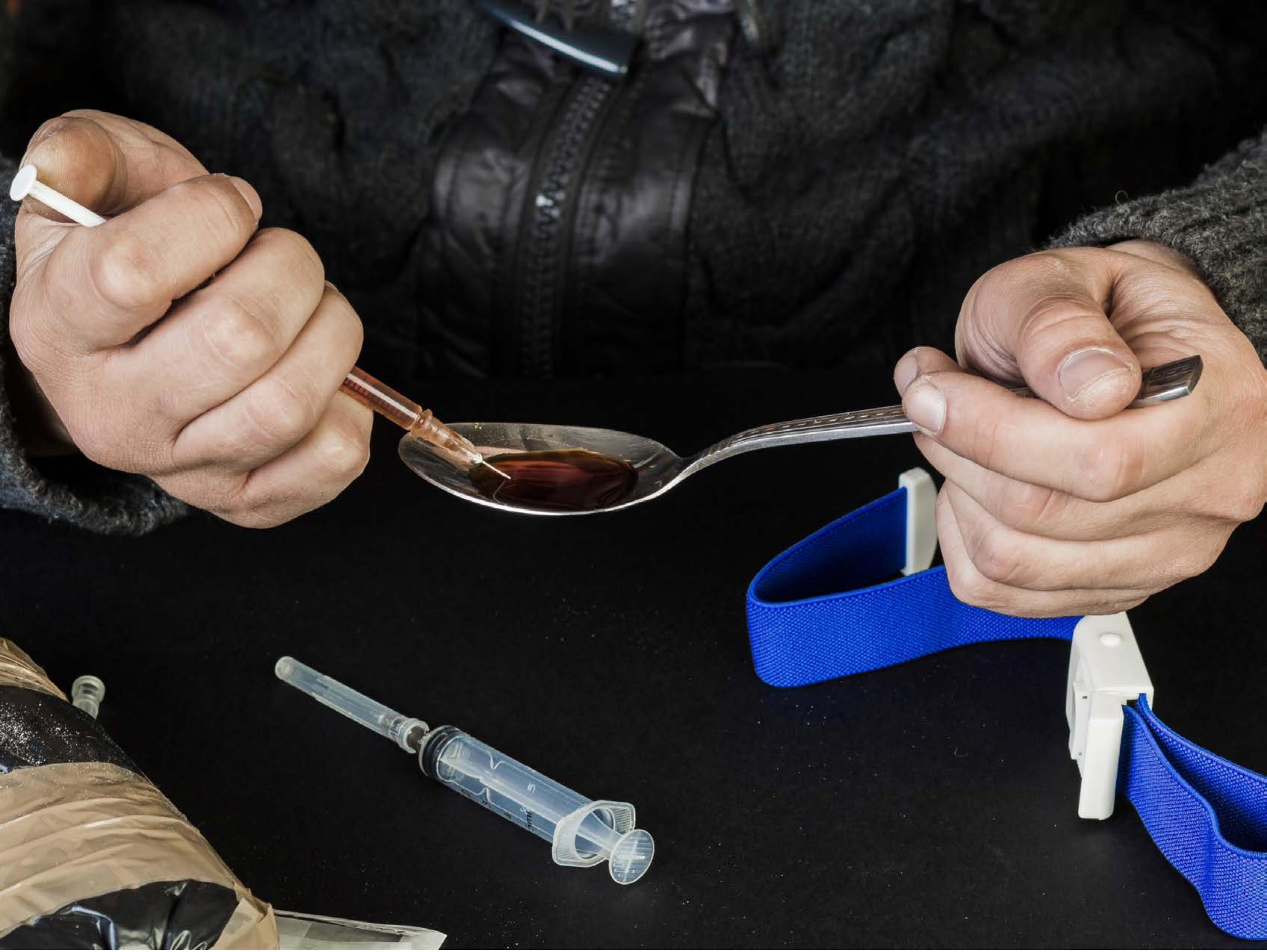

| Category | Craving (0-100) | Valence (1-9) | Arousal (1-9) | Typicality (0-100) | Relatedness | | | | | HSV | | |
|---|---|---|---|---|---|---|---|---|---|---|---|---|
| | | | | | Meth | Opioid | Both | Neither | MethToOpioid | Hue | Saturation | Value |
| Opioid Instrument Hand | 90.37 (13.36) | 4.56 (2.044) | 4.85 (2.76) | 92.7 (11.97) | 0.111 | 0.630 | 0.259 | 0 | 0.519 | 0.235 (0.308) | 0.165 (0.216) | 0.269 (0.297) |

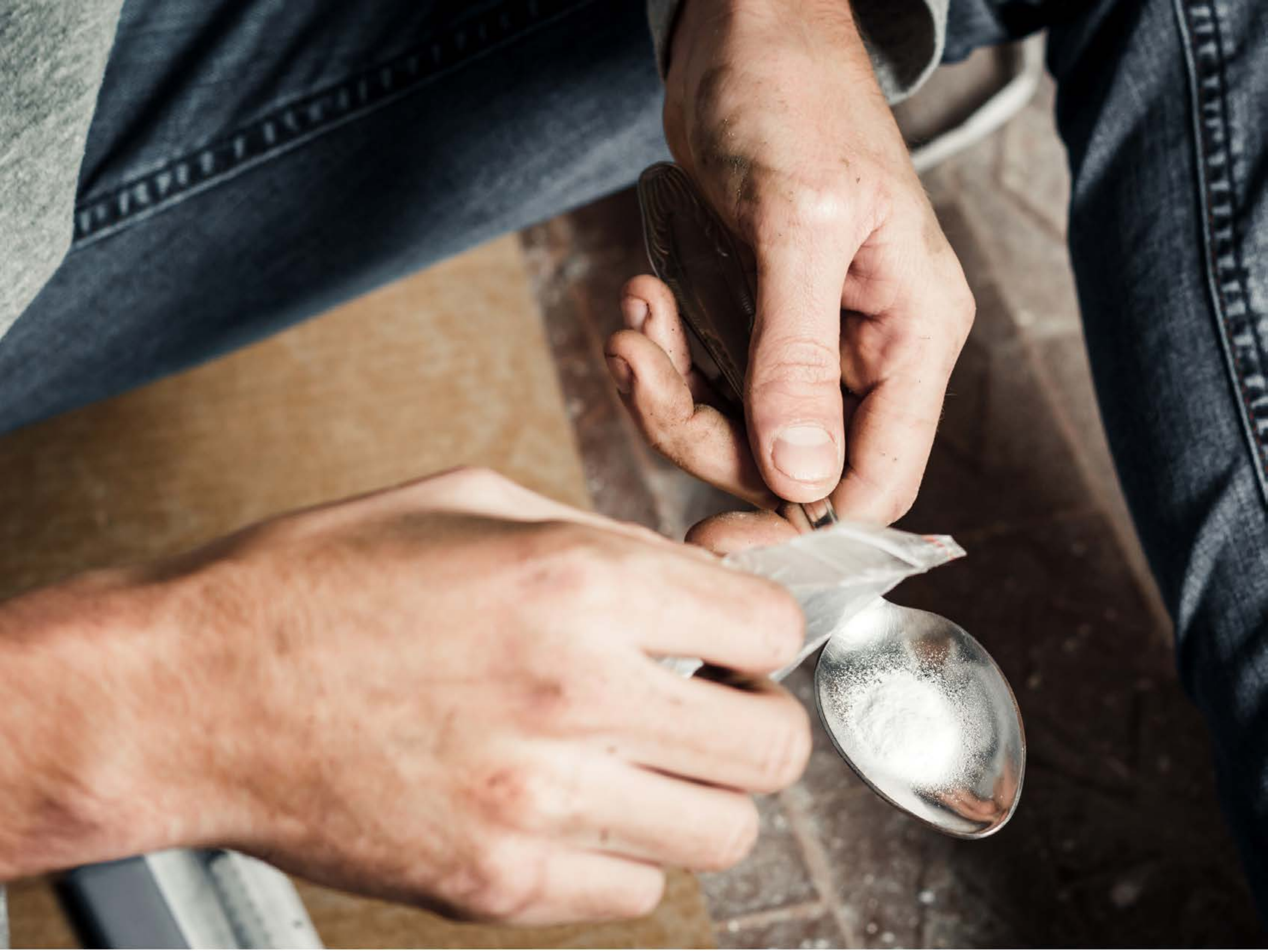

| Category | Craving (0-100) | Valence (1-9) | Arousal (1-9) | Typicality (0-100) | Relatedness | | | | | HSV | | |
|---|---|---|---|---|---|---|---|---|---|---|---|---|
| | | | | | Meth | Opioid | Both | Neither | MethToOpioid | Hue | Saturation | Value |
| Opioid Instrument Hand | 90.29 (14.88) | 4.5 (1.934) | 4.57 (2.92) | 92.18 (13.5) | 0.214 | 0.107 | 0.643 | 0.036 | -0.107 | 0.211 (0.232) | 0.335 (0.169) | 0.5 (0.316) |

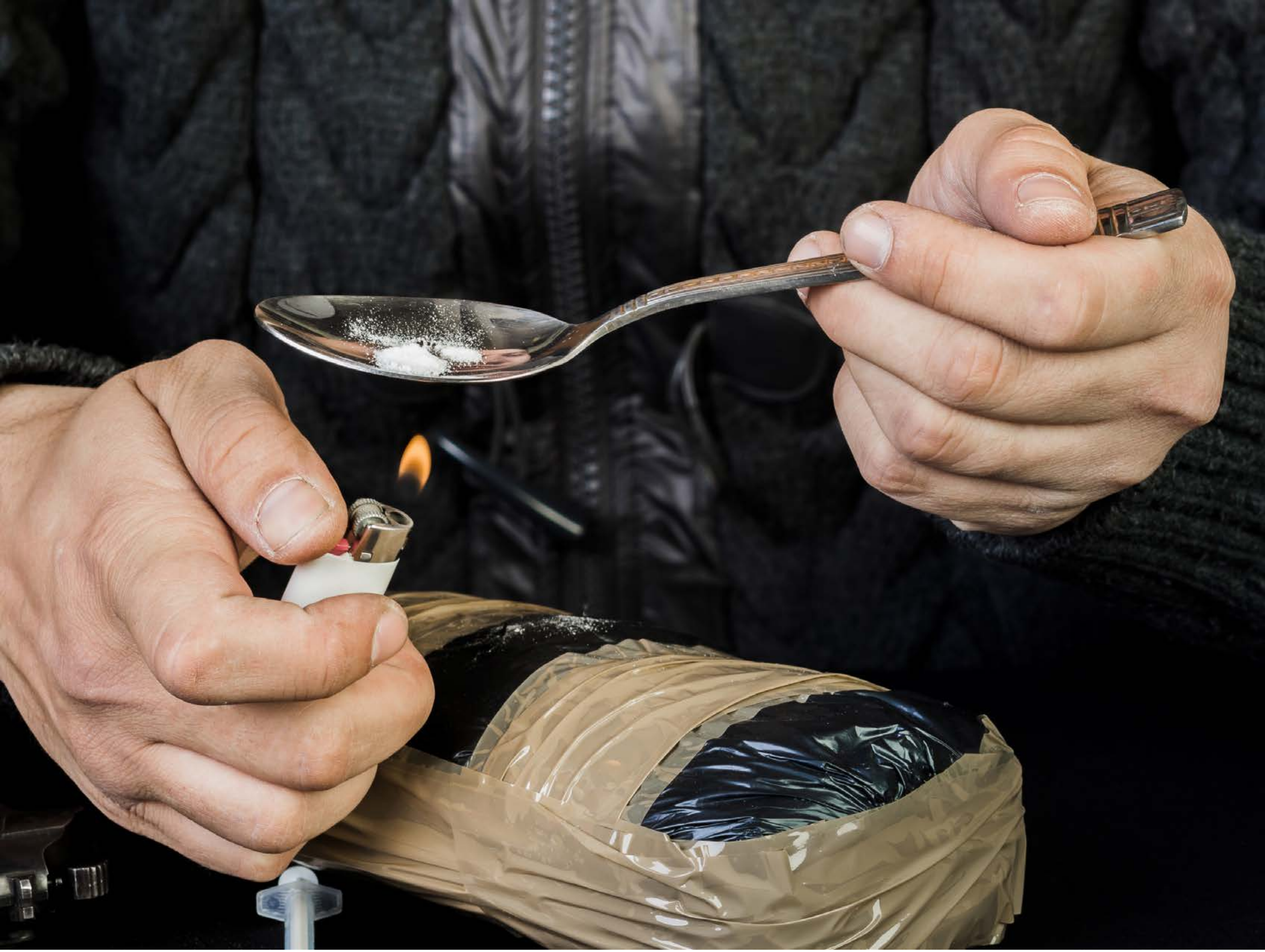

| Category | Craving (0-100) | Valence (1-9) | Arousal (1-9) | Typicality (0-100) | Relatedness | | | | | HSV | | |
|---|---|---|---|---|---|---|---|---|---|---|---|---|
| | | | | | Meth | Opioid | Both | Neither | MethToOpioid | Hue | Saturation | Value |
| Opioid Instrument Hand | 90.57 (14.83) | 4.46 (2.117) | 4.46 (2.96) | 90.07 (17.13) | 0.107 | 0.429 | 0.464 | 0 | 0.321 | 0.274 (0.283) | 0.221 (0.187) | 0.329 (0.313) |

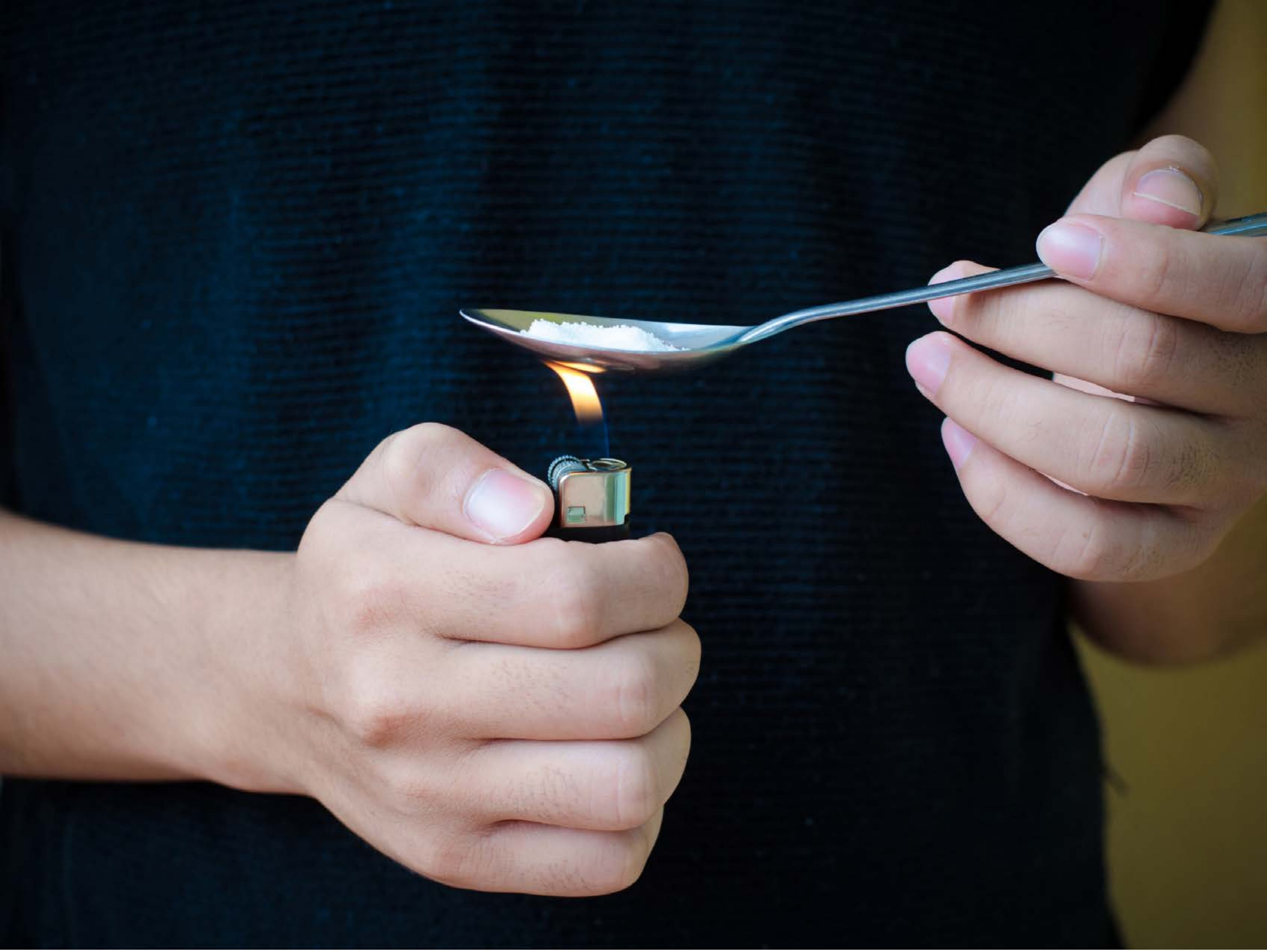

| Category | Craving (0-100) | Valence (1-9) | Arousal (1-9) | Typicality (0-100) | Relatedness | | | | | HSV | | |
|---|---|---|---|---|---|---|---|---|---|---|---|---|
| | | | | | Meth | Opioid | Both | Neither | MethToOpioid | Hue | Saturation | Value |
| Opioid Instrument Hand | 91.64 (13.88) | 4.29 (1.761) | 4.79 (2.79) | 93.36 (13.47) | 0.214 | 0.321 | 0.464 | 0 | 0.107 | 0.443 (0.27) | 0.576 (0.282) | 0.281 (0.284) |

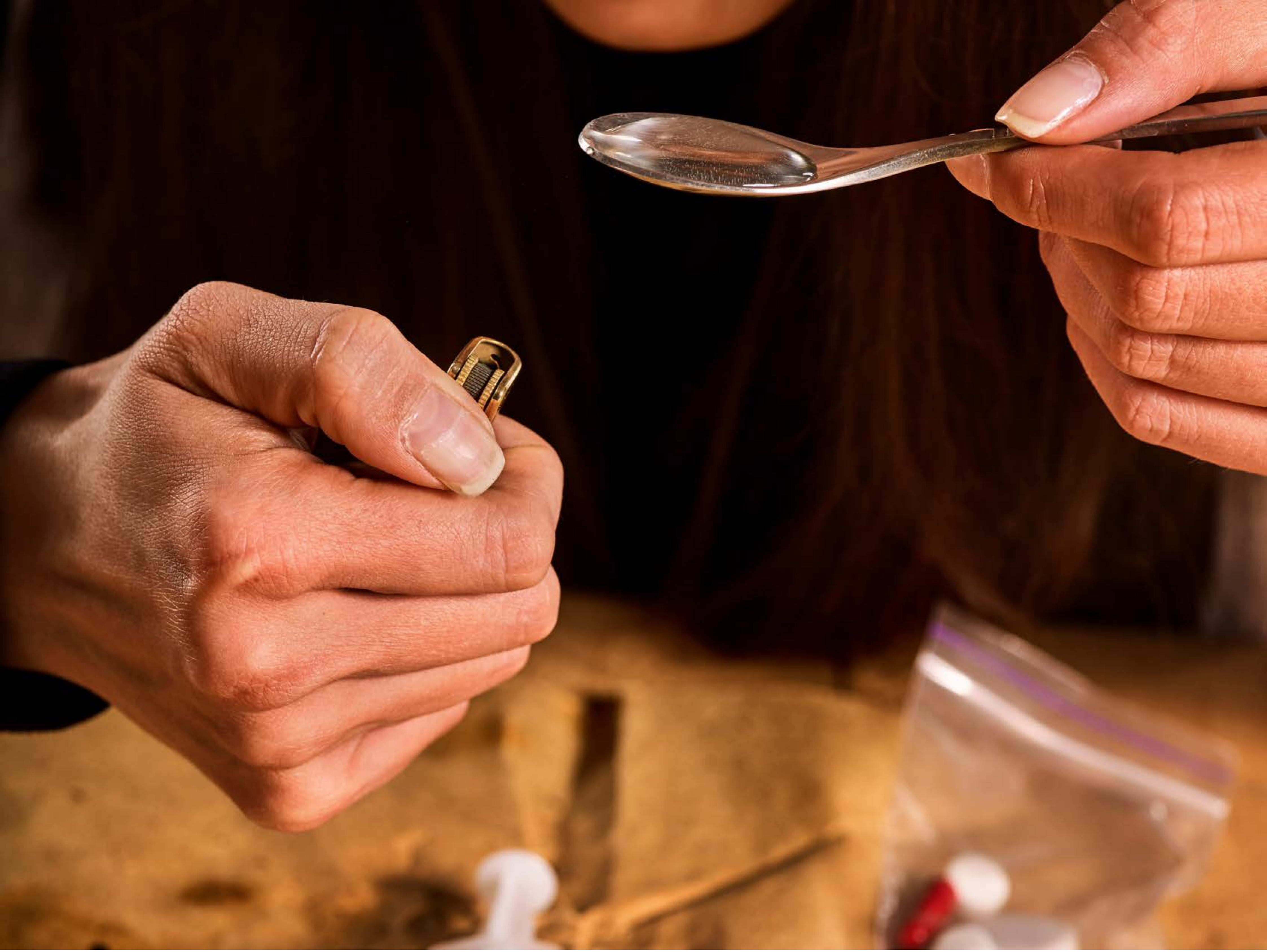

| Category | Craving (0-100) | Valence (1-9) | Arousal (1-9) | Typicality (0-100) | Relatedness | | | | | HSV | | |
|---|---|---|---|---|---|---|---|---|---|---|---|---|
| | | | | | Meth | Opioid | Both | Neither | MethToOpioid | Hue | Saturation | Value |
| Opioid Instrument Hand | 85.78 (17.96) | 4.63 (1.69) | 4.63 (2.48) | 85.67 (18.01) | 0.074 | 0.407 | 0.519 | 0 | 0.333 | 0.094 (0.209) | 0.741 (0.247) | 0.429 (0.339) |

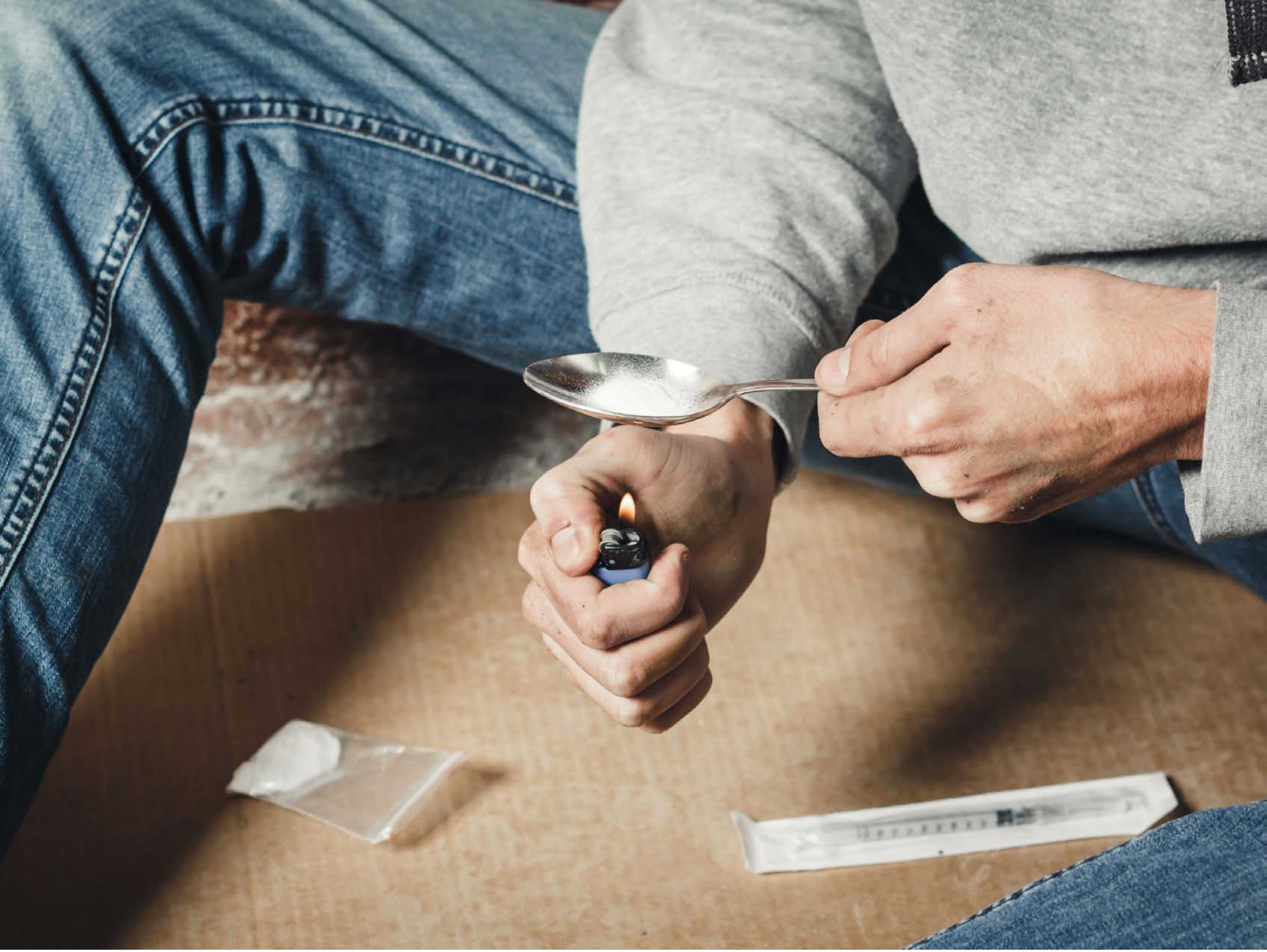

| Category | Craving (0-100) | Valence (1-9) | Arousal (1-9) | Typicality (0-100) | Relatedness | | | | | HSV | | |
|---|---|---|---|---|---|---|---|---|---|---|---|---|
| | | | | | Meth | Opioid | Both | Neither | MethToOpioid | Hue | Saturation | Value |
| Opioid Instrument Hand | 91.07 (13.76) | 4.52 (1.929) | 4.56 (2.89) | 89.93 (16.45) | 0.148 | 0.444 | 0.407 | 0 | 0.296 | 0.244 (0.22) | 0.277 (0.168) | 0.555 (0.242) |

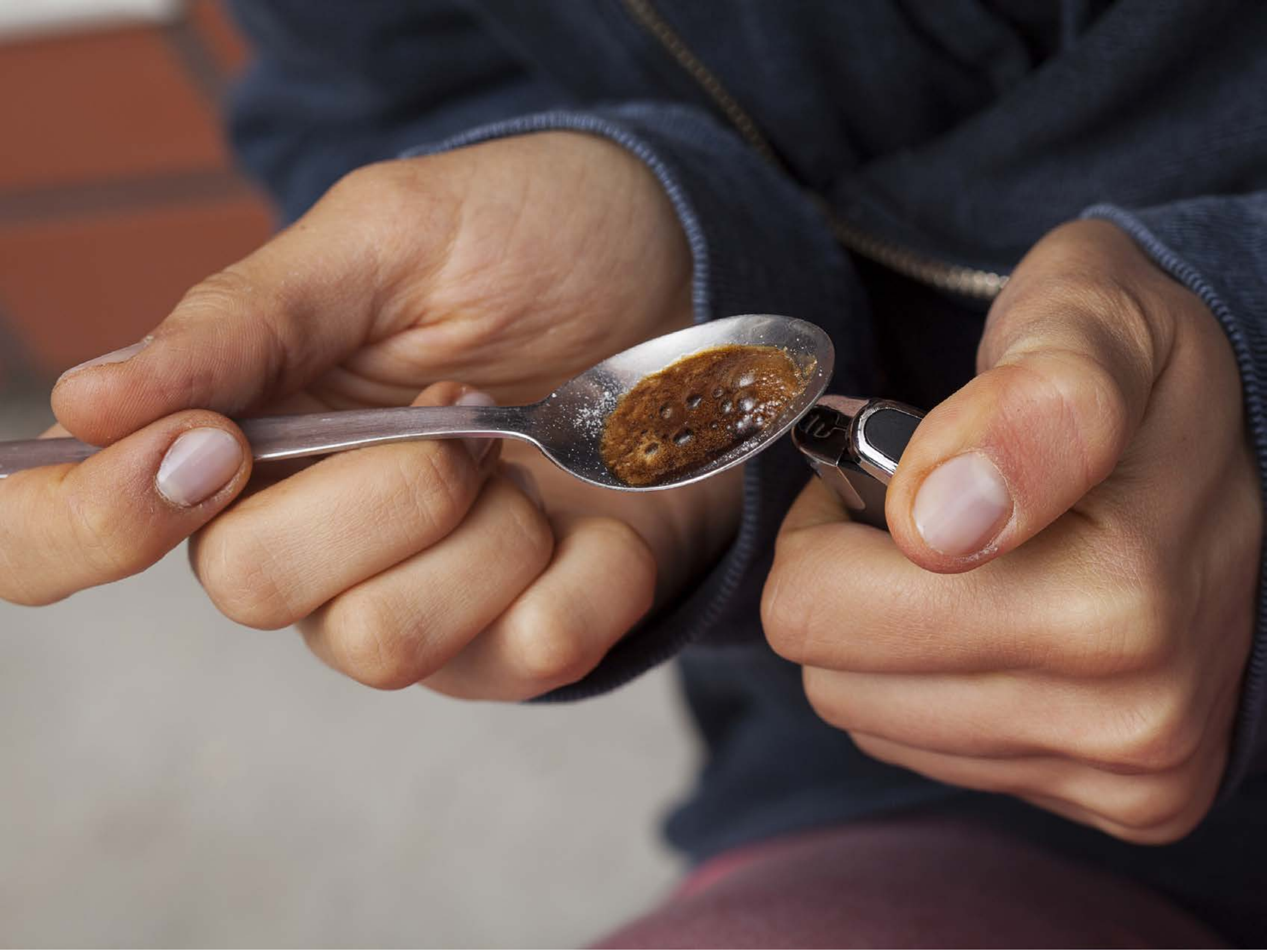

| Category | Craving (0-100) | Valence (1-9) | Arousal (1-9) | Typicality (0-100) | Relatedness | | | | | HSV | | |
|---|---|---|---|---|---|---|---|---|---|---|---|---|
| | | | | | Meth | Opioid | Both | Neither | MethToOpioid | Hue | Saturation | Value |
| Opioid Instrument Hand | 90.61 (20.79) | 4.14 (2.013) | 4.75 (2.9) | 89.36 (21.45) | 0.071 | 0.75 | 0.143 | 0.036 | 0.679 | 0.287 (0.326) | 0.31 (0.168) | 0.458 (0.241) |

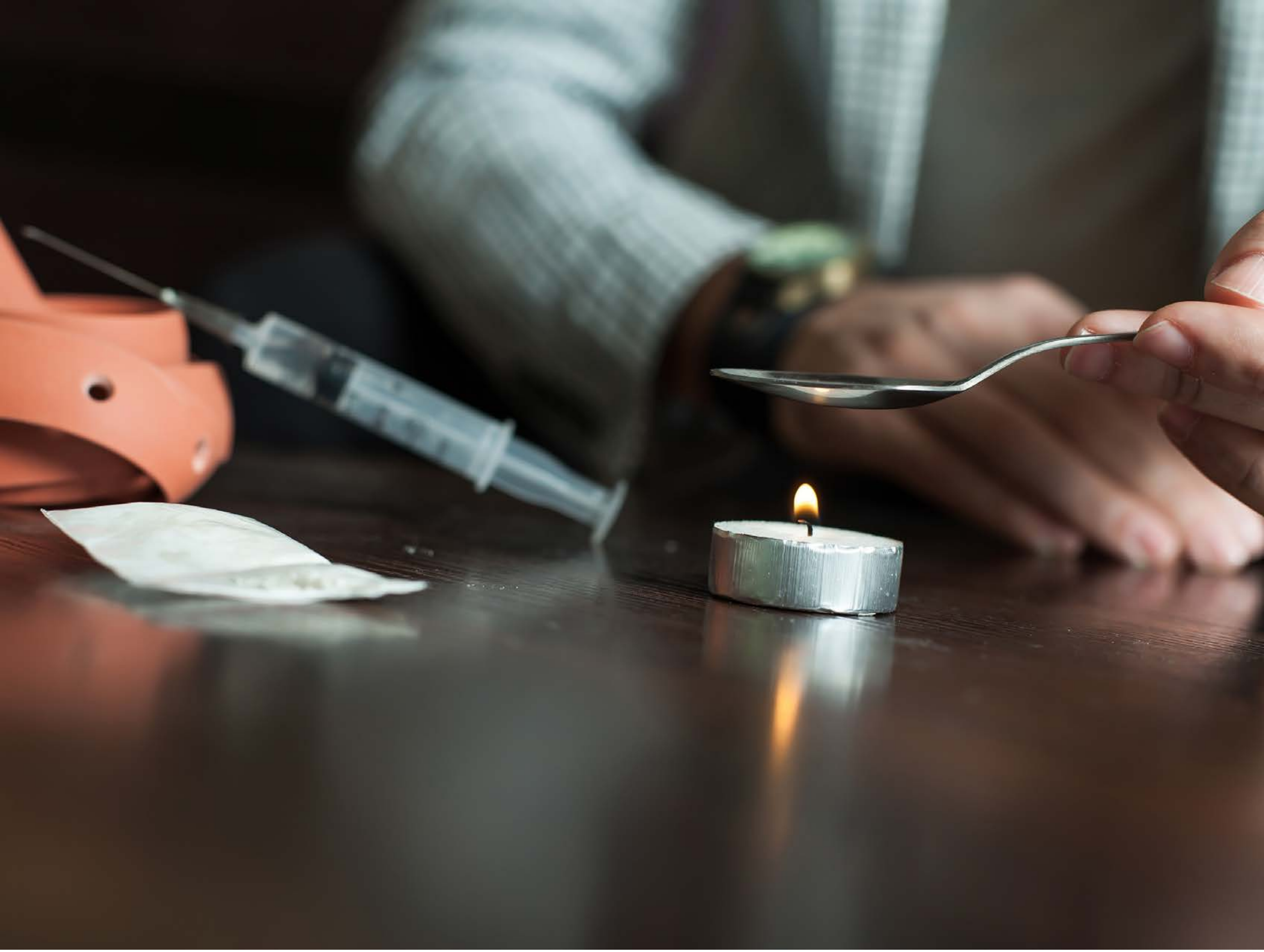

| Category | Craving (0-100) | Valence (1-9) | Arousal (1-9) | Typicality (0-100) | Relatedness | | | | | HSV | | |
|---|---|---|---|---|---|---|---|---|---|---|---|---|
| | | | | | Meth | Opioid | Both | Neither | MethToOpioid | Hue | Saturation | Value |
| Opioid Instrument Hand | 93.93 (11.7) | 4.39 (1.812) | 4.57 (2.82) | 93.93 (11.45) | 0.107 | 0.429 | 0.464 | 0 | 0.321 | 0.18 (0.194) | 0.269 (0.164) | 0.326 (0.23) |

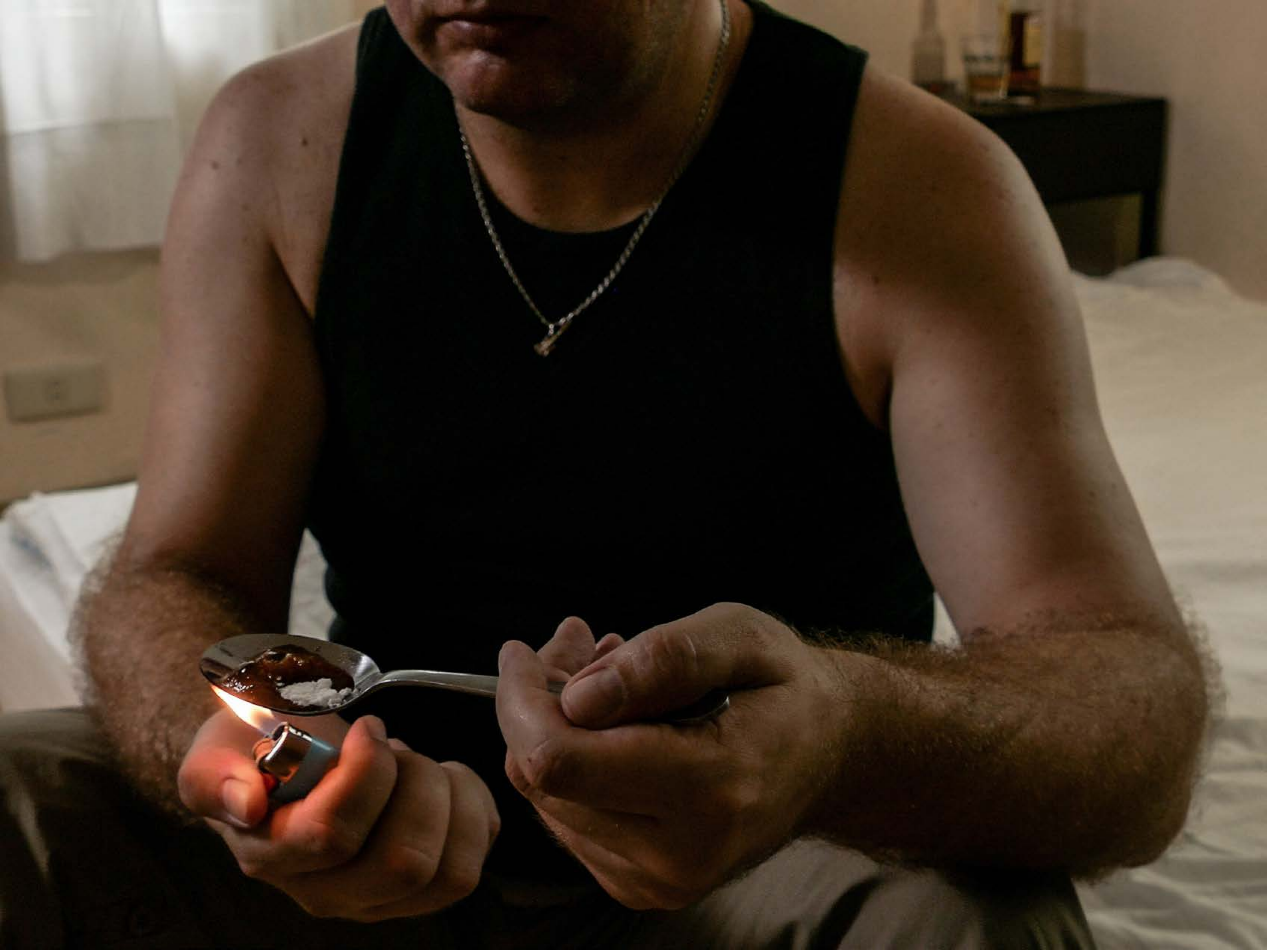

| Category | Craving (0-100) | Valence (1-9) | Arousal (1-9) | Typicality (0-100) | Relatedness | | | | | HSV | | |
|---|---|---|---|---|---|---|---|---|---|---|---|---|
| | | | | | Meth | Opioid | Both | Neither | MethToOpioid | Hue | Saturation | Value |
| Opioid Instrument Hand | 88.52 (15.71) | 4.3 (1.54) | 4.63 (2.6) | 88.78 (17.08) | 0.111 | 0.704 | 0.185 | 0 | 0.593 | 0.102 (0.108) | 0.485 (0.328) | 0.261 (0.235) |

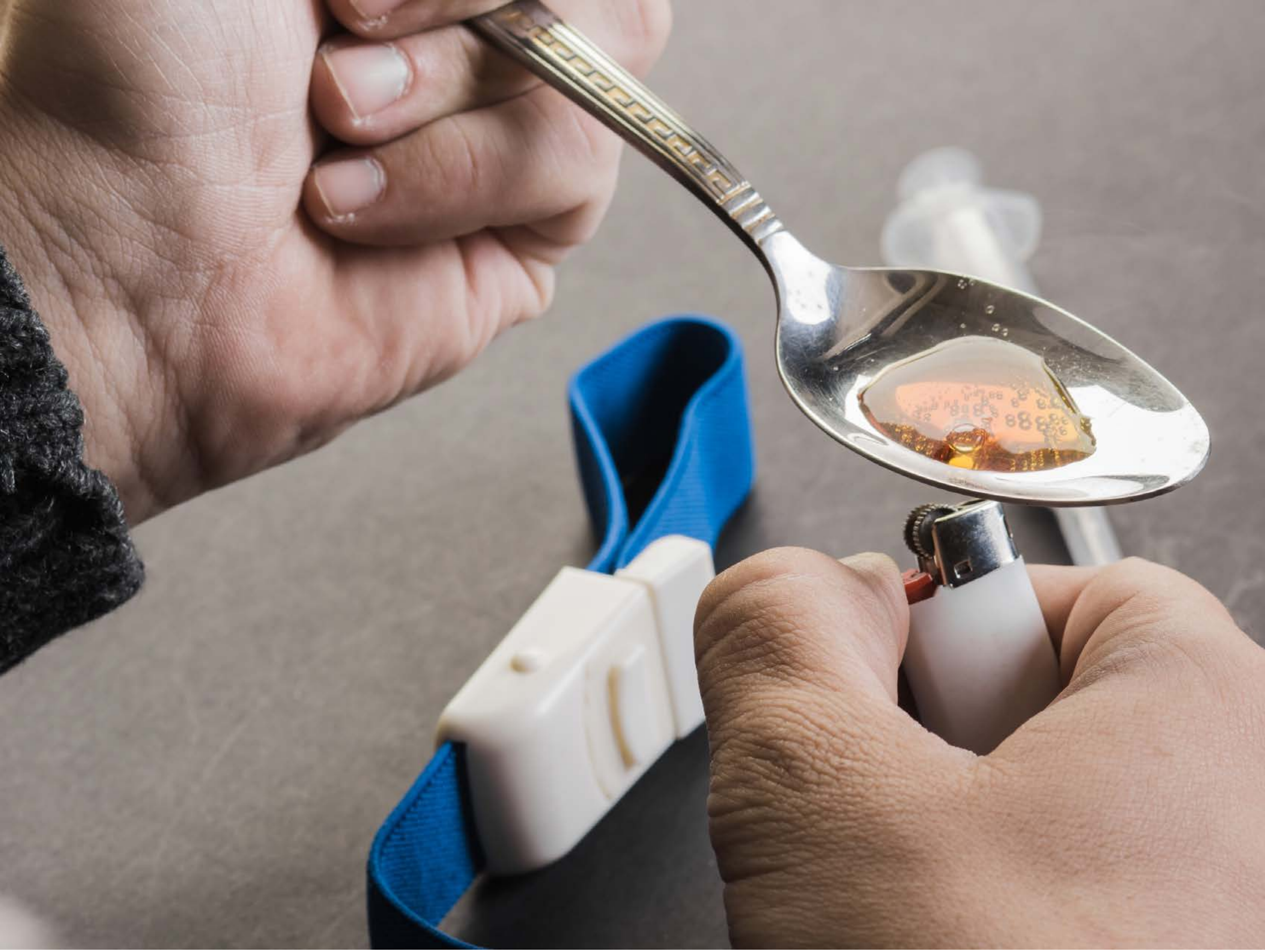

| Category | Craving (0-100) | Valence (1-9) | Arousal (1-9) | Typicality (0-100) | Relatedness | | | | | HSV | | |
|---|---|---|---|---|---|---|---|---|---|---|---|---|
| | | | | | Meth | Opioid | Both | Neither | MethToOpioid | Hue | Saturation | Value |
| Opioid Instrument Hand | 89.67 (14.45) | 4.22 (1.805) | 4.7 (2.57) | 88.33 (16.36) | 0.074 | 0.630 | 0.296 | 0 | 0.556 | 0.116 (0.194) | 0.194 (0.189) | 0.588 (0.201) |

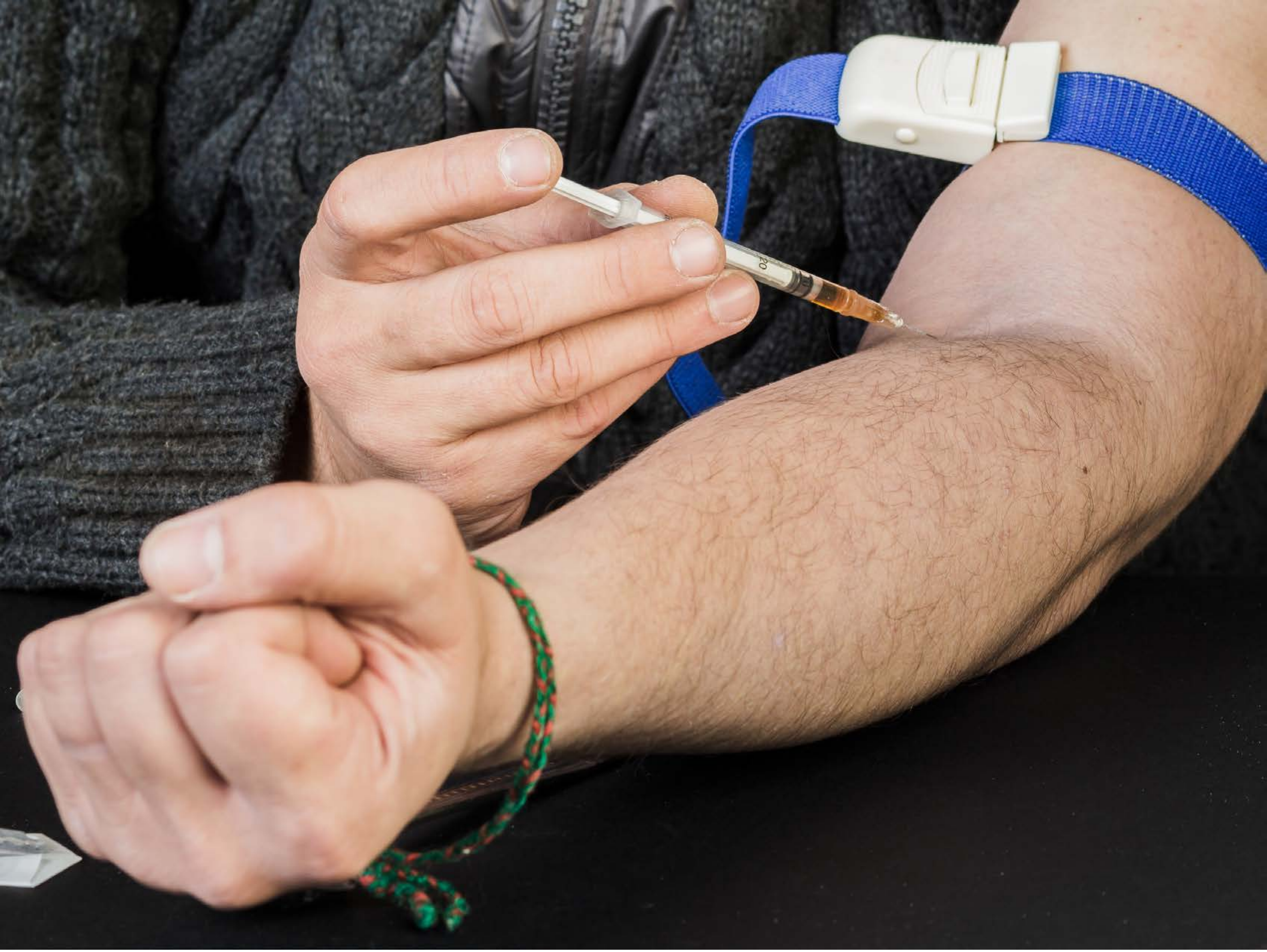

| Category | Craving (0-100) | Valence (1-9) | Arousal (1-9) | Typicality (0-100) | Relatedness | | | | | HSV | | |
|---|---|---|---|---|---|---|---|---|---|---|---|---|
| | | | | | Meth | Opioid | Both | Neither | MethToOpioid | Hue | Saturation | Value |
| Opioid Instrument Hand | 90 (15.4) | 4.22 (2.006) | 4.78 (2.67) | 89.85 (16.26) | 0.111 | 0.185 | 0.704 | 0 | 0.074 | 0.24 (0.31) | 0.215 (0.175) | 0.487 (0.34) |

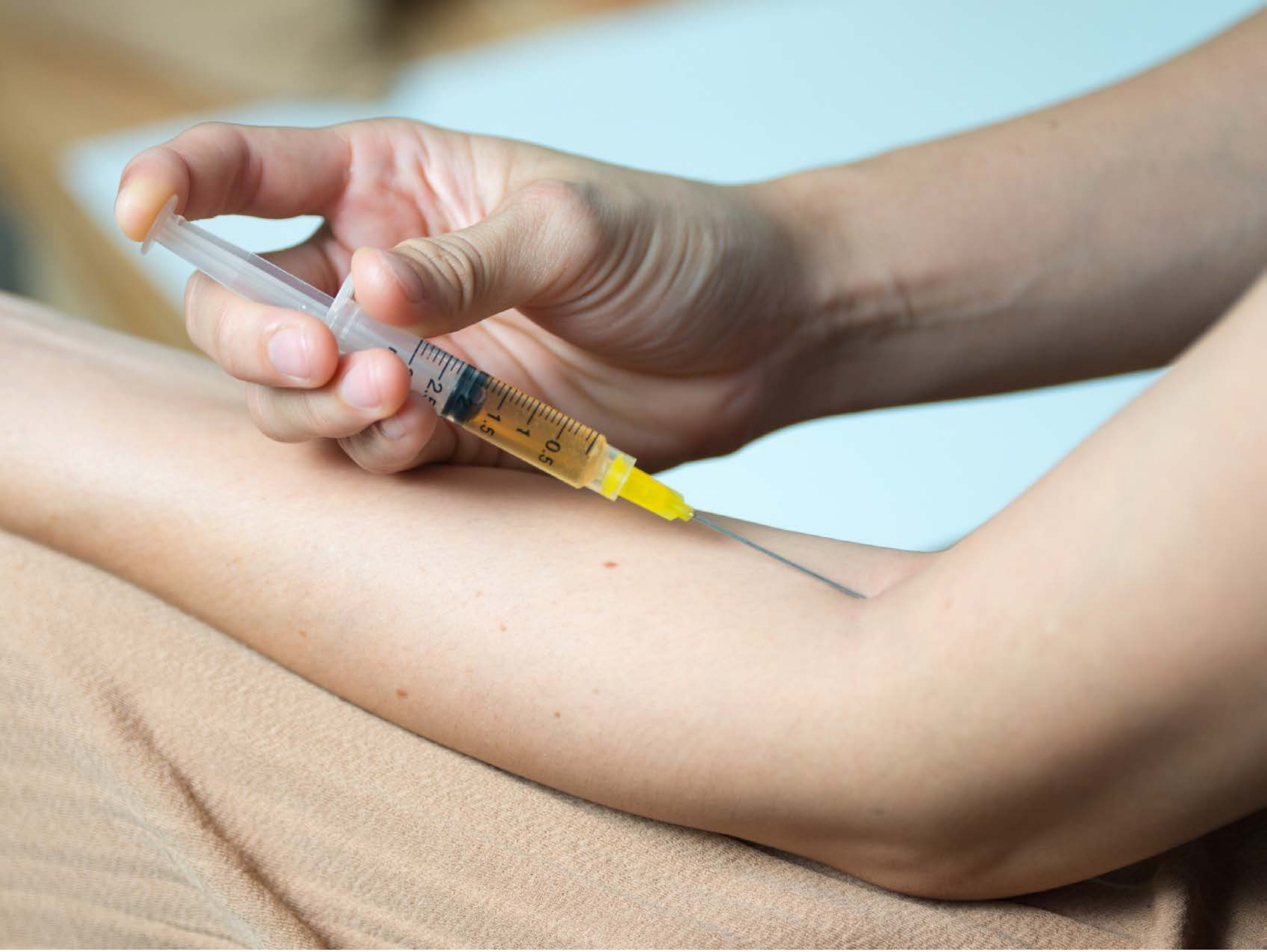


| | | | | | Relatedness | | | | | HSV | | |
|---|---|---|---|---|---|---|---|---|---|---|---|---|
| **Category** | **Craving (0-100)** | **Valence (1-9)** | **Arousal (1-9)** | **Typicality (0-100)** | Meth | Opioid | Both | Neither | MethToOpioid | Hue | Saturation | Value |
| Opioid Instrument Hand | 87.52 (18.3) | 4.3 (2.072) | 4.67 (2.88) | 88.89 (17.57) | 0.037 | 0.148 | 0.815 | 0 | 0.111 | 0.145 (0.172) | 0.244 (0.122) | 0.744 (0.206) |

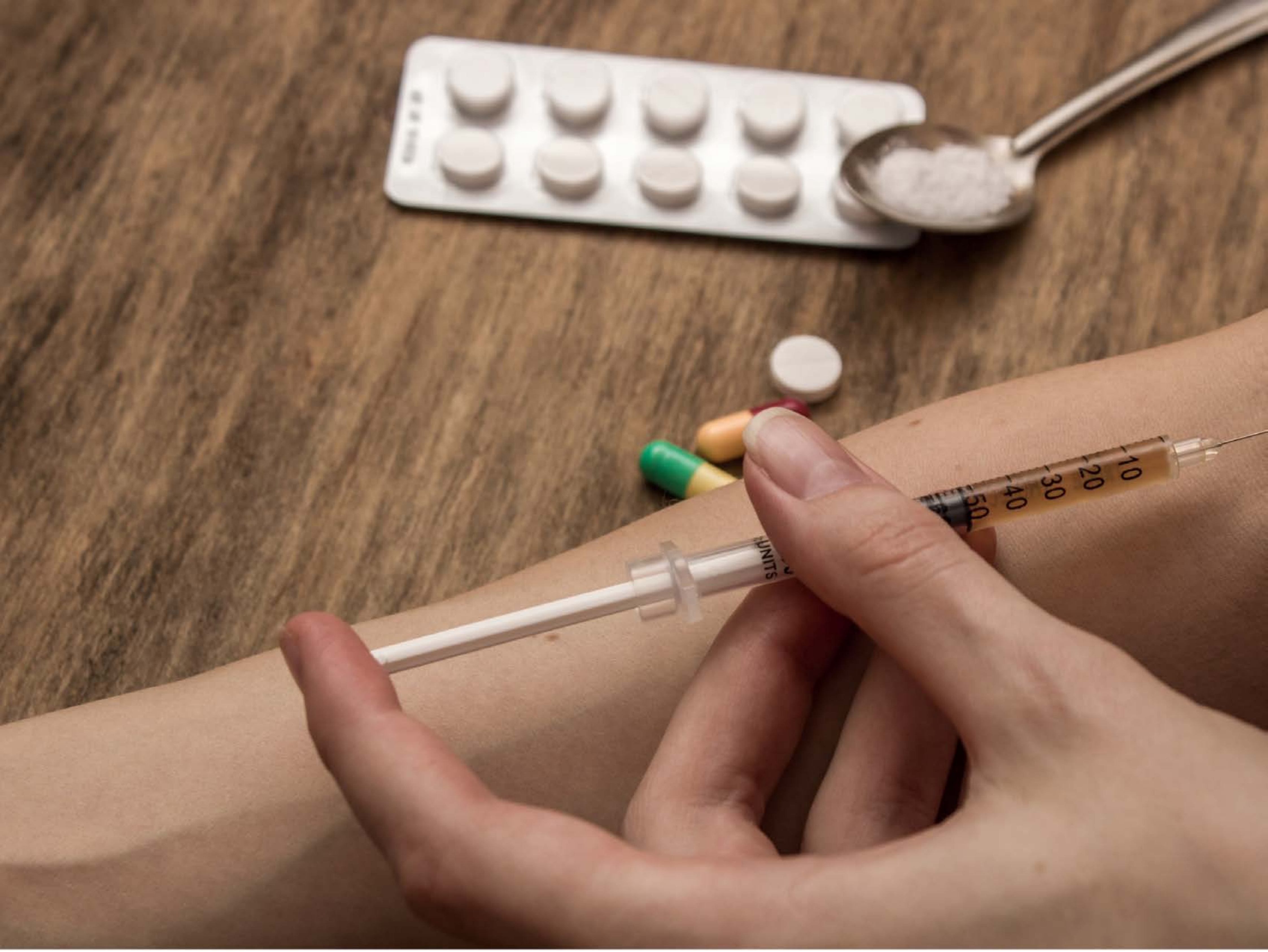


| Category | Craving (0-100) | Valence (1-9) | Arousal (1-9) | Typicality (0-100) | Relatedness | | | | | HSV | | |
|---|---|---|---|---|---|---|---|---|---|---|---|---|
| | | | | | Meth | Opioid | Both | Neither | MethToOpioid | Hue | Saturation | Value |
| Opioid Instrument Hand | 95.39 (9.92) | 4.43 (2.08) | 5.25 (3) | 94.82 (10.58) | 0 | 0.536 | 0.464 | 0 | 0.536 | 0.069 (0.054) | 0.406 (0.14) | 0.574 (0.17) |

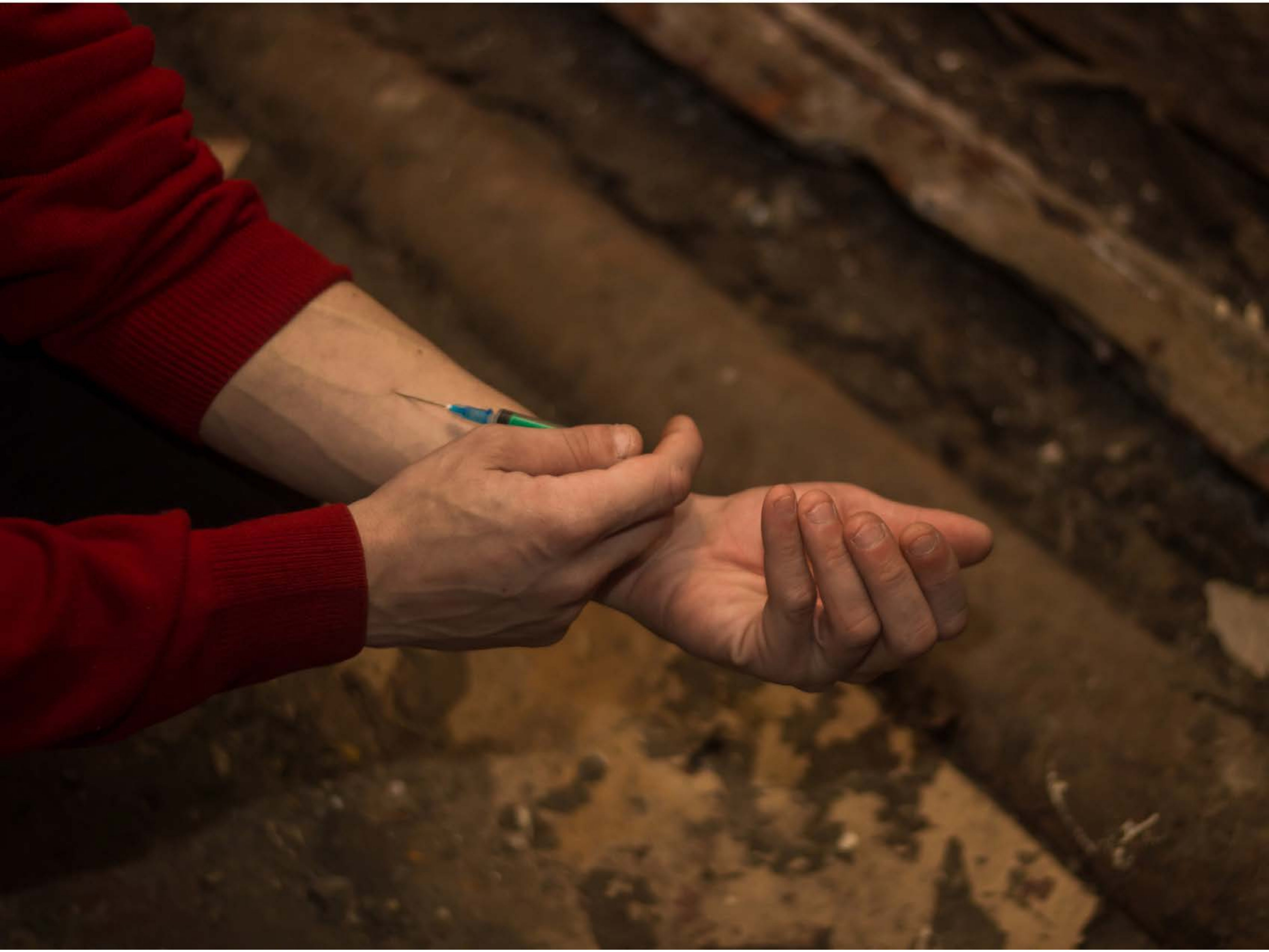

| Category | Craving (0-100) | Valence (1-9) | Arousal (1-9) | Typicality (0-100) | Relatedness | | | | | HSV | | |
|---|---|---|---|---|---|---|---|---|---|---|---|---|
| | | | | | Meth | Opioid | Both | Neither | MethToOpioid | Hue | Saturation | Value |
| Opioid Instrument Hand | 87.19 (17.94) | 4.07 (2.218) | 4.93 (2.84) | 89.48 (15.99) | 0.111 | 0.037 | 0.852 | 0 | -0.074 | 0.179 (0.297) | 0.654 (0.151) | 0.274 (0.162) |

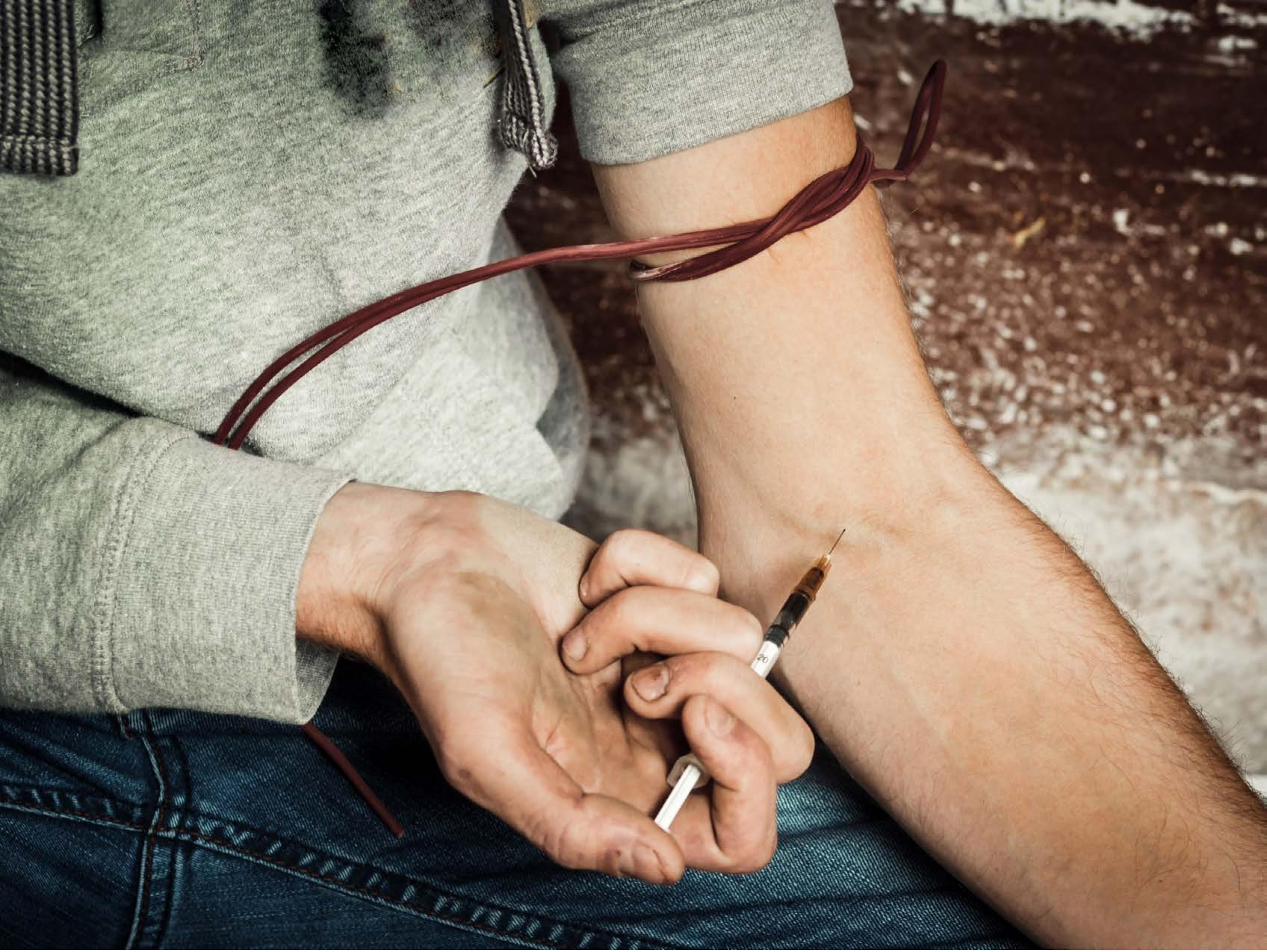

| Category | Craving (0-100) | Valence (1-9) | Arousal (1-9) | Typicality (0-100) | Relatedness | | | | | HSV | | |
|---|---|---|---|---|---|---|---|---|---|---|---|---|
| | | | | | Meth | Opioid | Both | Neither | MethToOpioid | Hue | Saturation | Value |
| Opioid Instrument Hand | 95.5 (9.95) | 4 (2.073) | 5.36 (2.93) | 94.25 (11.13) | 0.107 | 0.214 | 0.679 | 0 | 0.107 | 0.184 (0.189) | 0.314 (0.204) | 0.566 (0.275) |

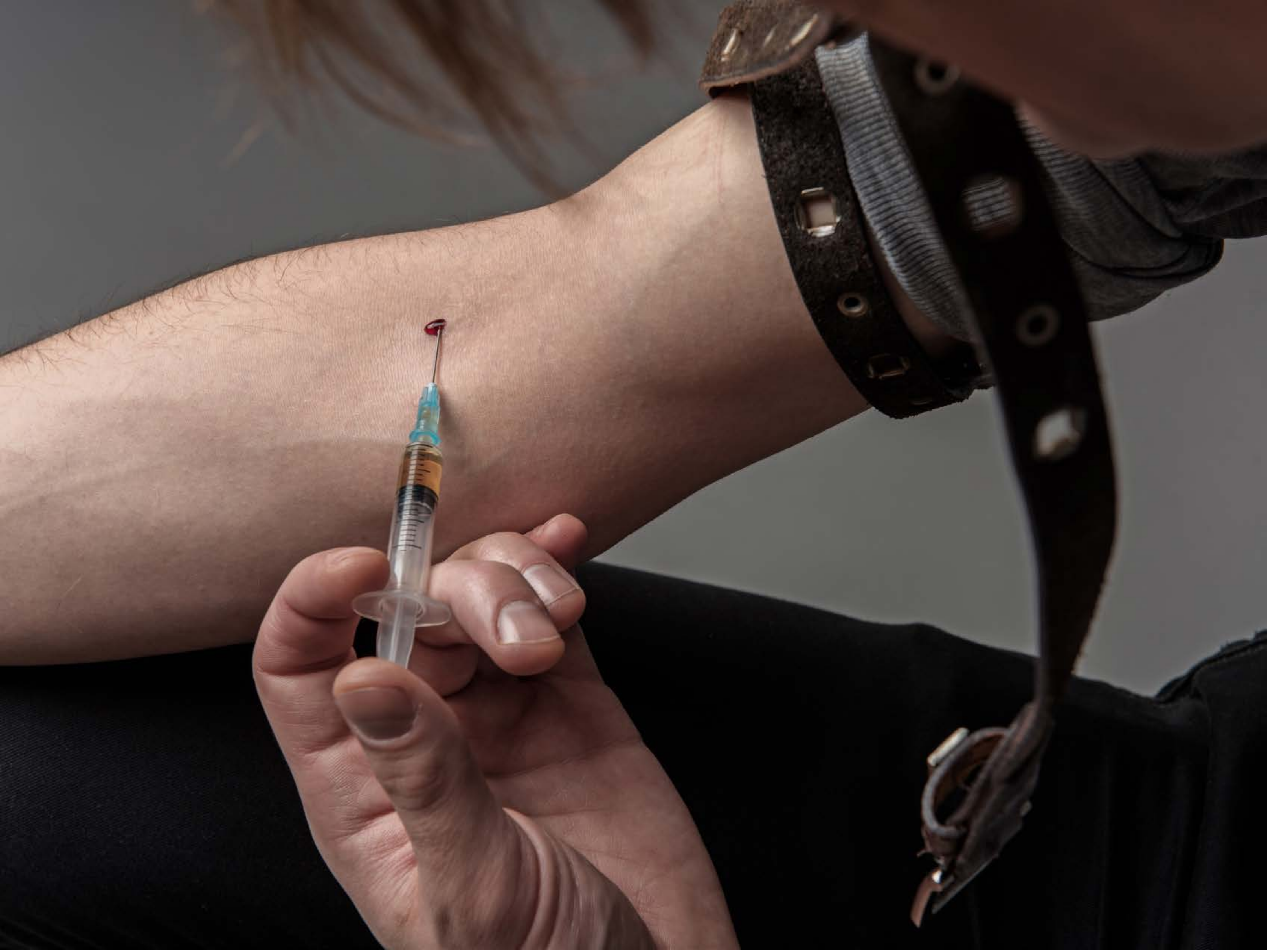

| Category | Craving (0-100) | Valence (1-9) | Arousal (1-9) | Typicality (0-100) | Relatedness | | | | | HSV | | |
|---|---|---|---|---|---|---|---|---|---|---|---|---|
| | | | | | Meth | Opioid | Both | Neither | MethToOpioid | Hue | Saturation | Value |
| Opioid Instrument Hand | 92.82 (12.11) | 3.86 (2.206) | 4.82 (2.91) | 93.82 (10.97) | 0.107 | 0.143 | 0.75 | 0 | 0.036 | 0.181 (0.309) | 0.221 (0.201) | 0.317 (0.249) |

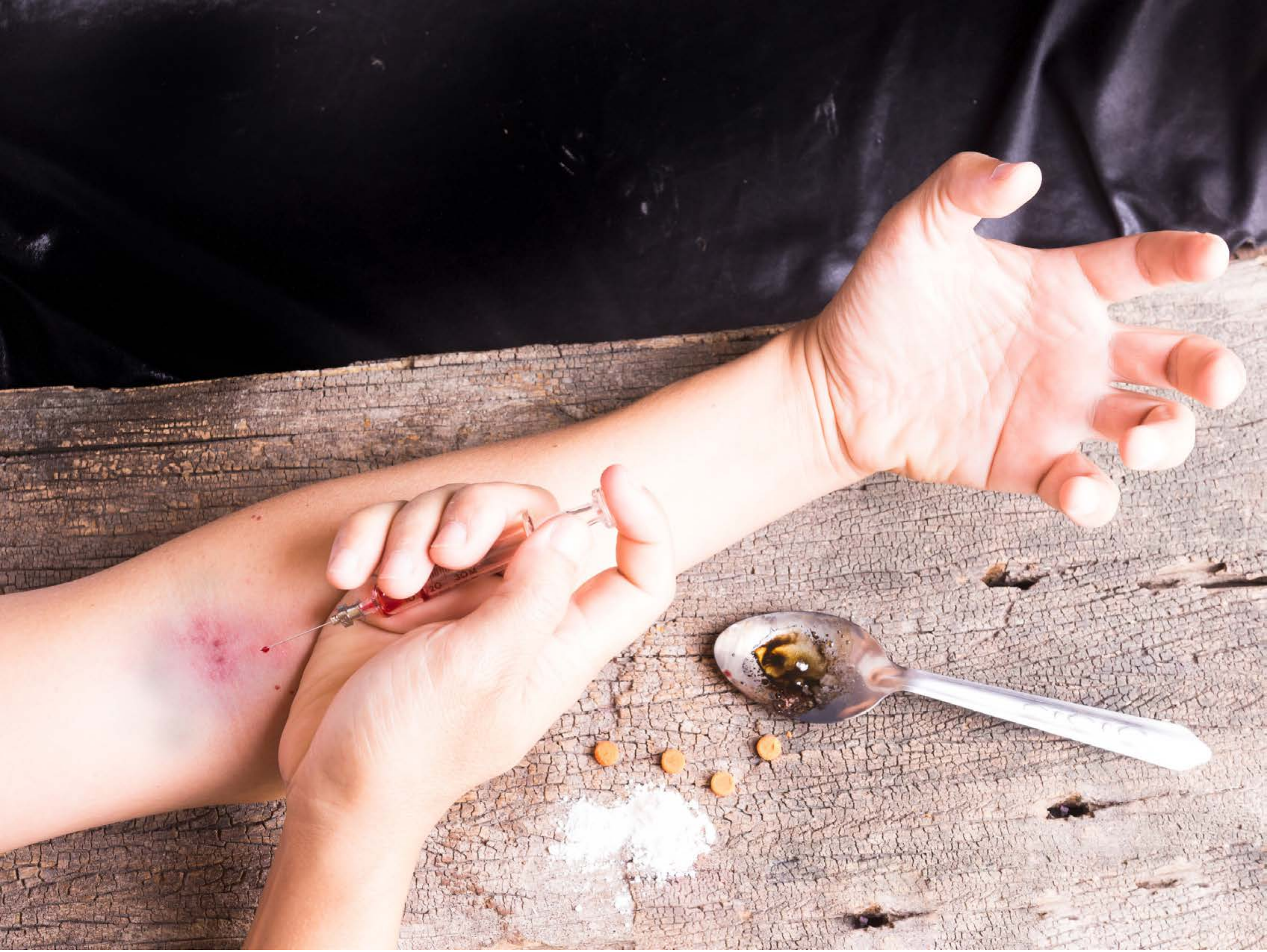

| Category | Craving (0-100) | Valence (1-9) | Arousal (1-9) | Typicality (0-100) | Relatedness | | | | | HSV | | |
|---|---|---|---|---|---|---|---|---|---|---|---|---|
| | | | | | Meth | Opioid | Both | Neither | MethToOpioid | Hue | Saturation | Value |
| Opioid Instrument Hand | 88 (17.9) | 3.85 (2.107) | 4.81 (2.57) | 89.59 (15.94) | 0.037 | 0.741 | 0.222 | 0 | 0.704 | 0.302 (0.37) | 0.242 (0.175) | 0.588 (0.385) |

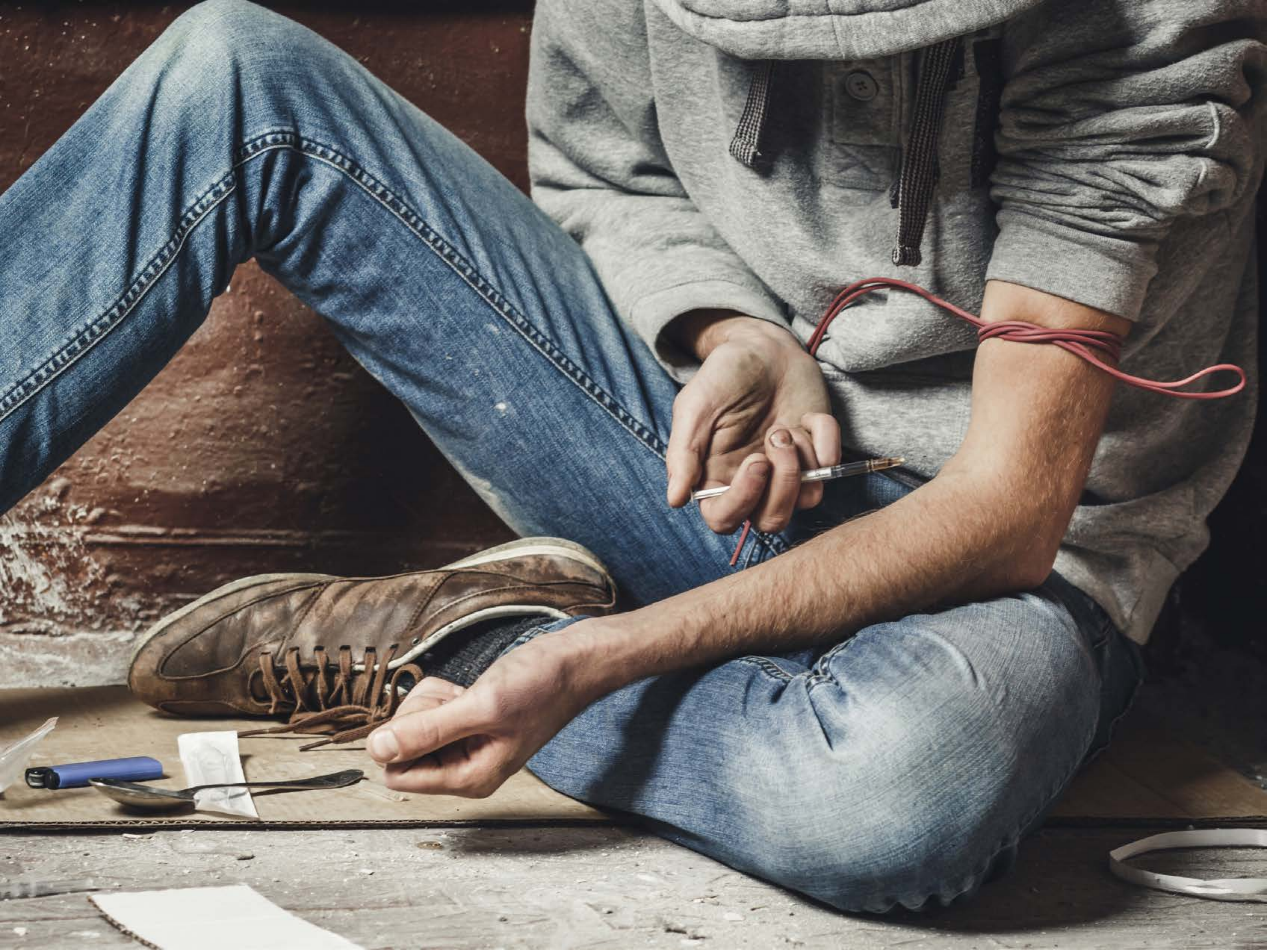

| Category | Craving (0-100) | Valence (1-9) | Arousal (1-9) | Typicality (0-100) | Relatedness | | | | | HSV | | |
|---|---|---|---|---|---|---|---|---|---|---|---|---|
| | | | | | Meth | Opioid | Both | Neither | MethToOpioid | Hue | Saturation | Value |
| Opioid Instrument Hand | 89.46 (21.55) | 4.11 (2.043) | 4.86 (2.8) | 92.75 (13.34) | 0.143 | 0.214 | 0.643 | 0 | 0.071 | 0.32 (0.277) | 0.193 (0.138) | 0.467 (0.242) |

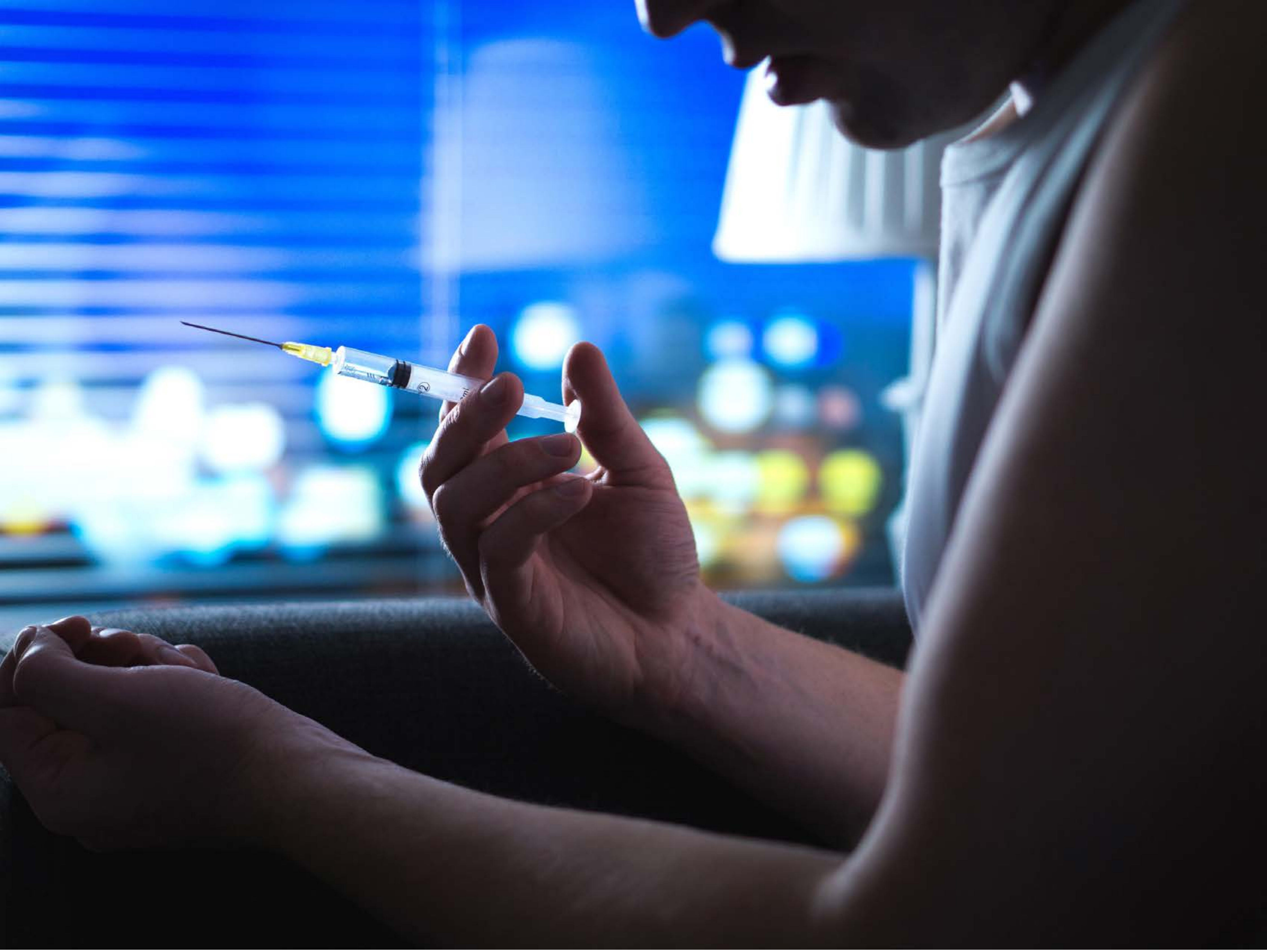

| Category | Craving (0-100) | Valence (1-9) | Arousal (1-9) | Typicality (0-100) | Relatedness | | | | | HSV | | |
|---|---|---|---|---|---|---|---|---|---|---|---|---|
| | | | | | Meth | Opioid | Both | Neither | MethToOpioid | Hue | Saturation | Value |
| Opioid Instrument Hand | 93.86 (10.92) | 4.43 (2.098) | 4.61 (2.83) | 92.07 (20.61) | 0.143 | 0 | 0.821 | 0.036 | -0.143 | 0.625 (0.307) | 0.36 (0.304) | 0.49 (0.39) |

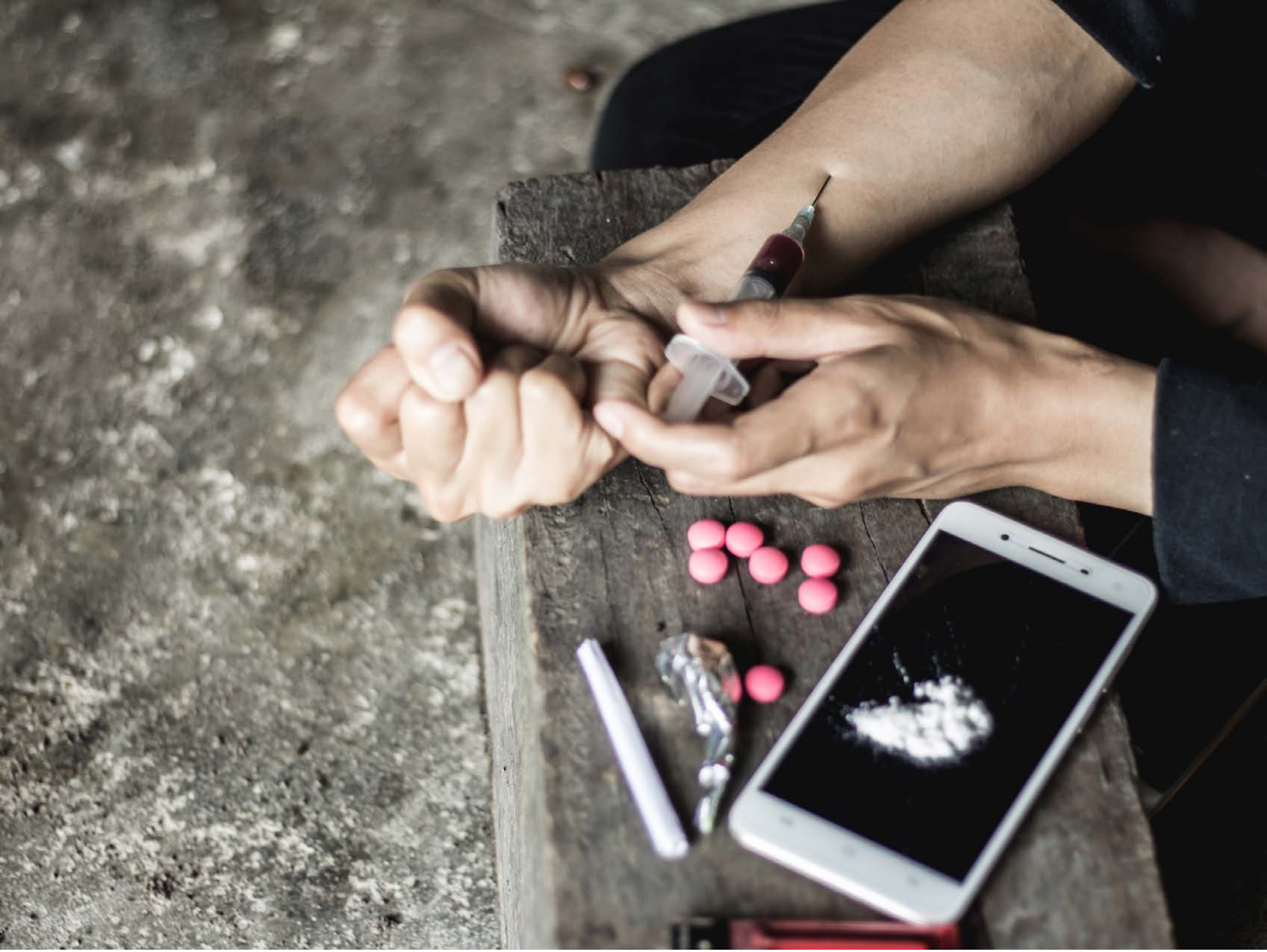

| Category | Craving (0-100) | Valence (1-9) | Arousal (1-9) | Typicality (0-100) | Relatedness | | | | | HSV | | |
|---|---|---|---|---|---|---|---|---|---|---|---|---|
| | | | | | Meth | Opioid | Both | Neither | MethToOpioid | Hue | Saturation | Value |
| Opioid Instrument Hand | 90.79 (15.16) | 4.29 (2.123) | 5.18 (2.51) | 92.07 (14.02) | 0 | 0.643 | 0.357 | 0 | 0.643 | 0.176 (0.211) | 0.14 (0.107) | 0.449 (0.259) |

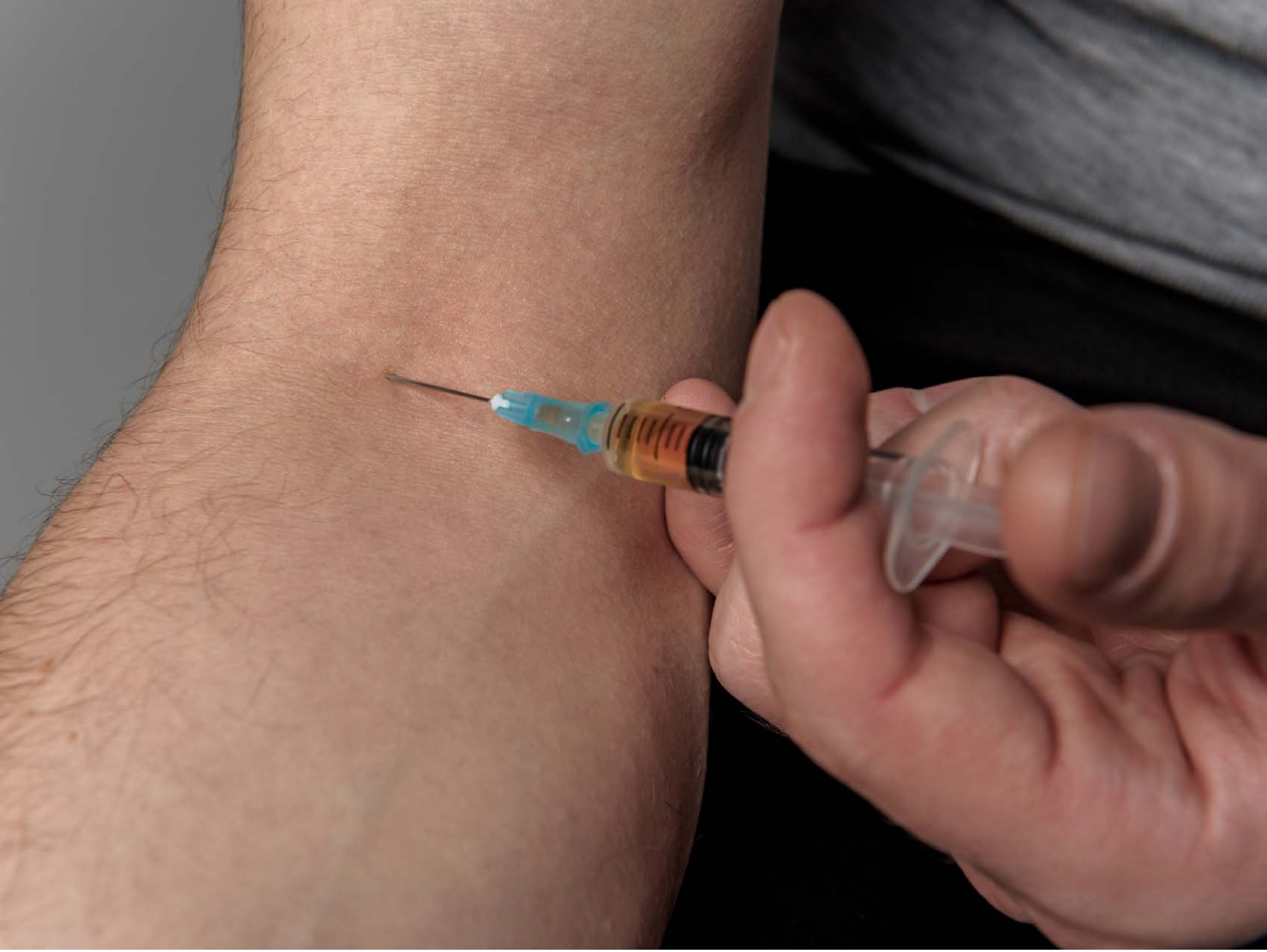

| Category | Craving (0-100) | Valence (1-9) | Arousal (1-9) | Typicality (0-100) | Relatedness | | | | | HSV | | |
|---|---|---|---|---|---|---|---|---|---|---|---|---|
| | | | | | Meth | Opioid | Both | Neither | MethToOpioid | Hue | Saturation | Value |
| Opioid Instrument Hand | 84.85 (20.77) | 4.15 (1.68) | 4.33 (2.5) | 85.7 (20.47) | 0.111 | 0.148 | 0.704 | 0.037 | 0.037 | 0.107 (0.217) | 0.283 (0.176) | 0.508 (0.24) |

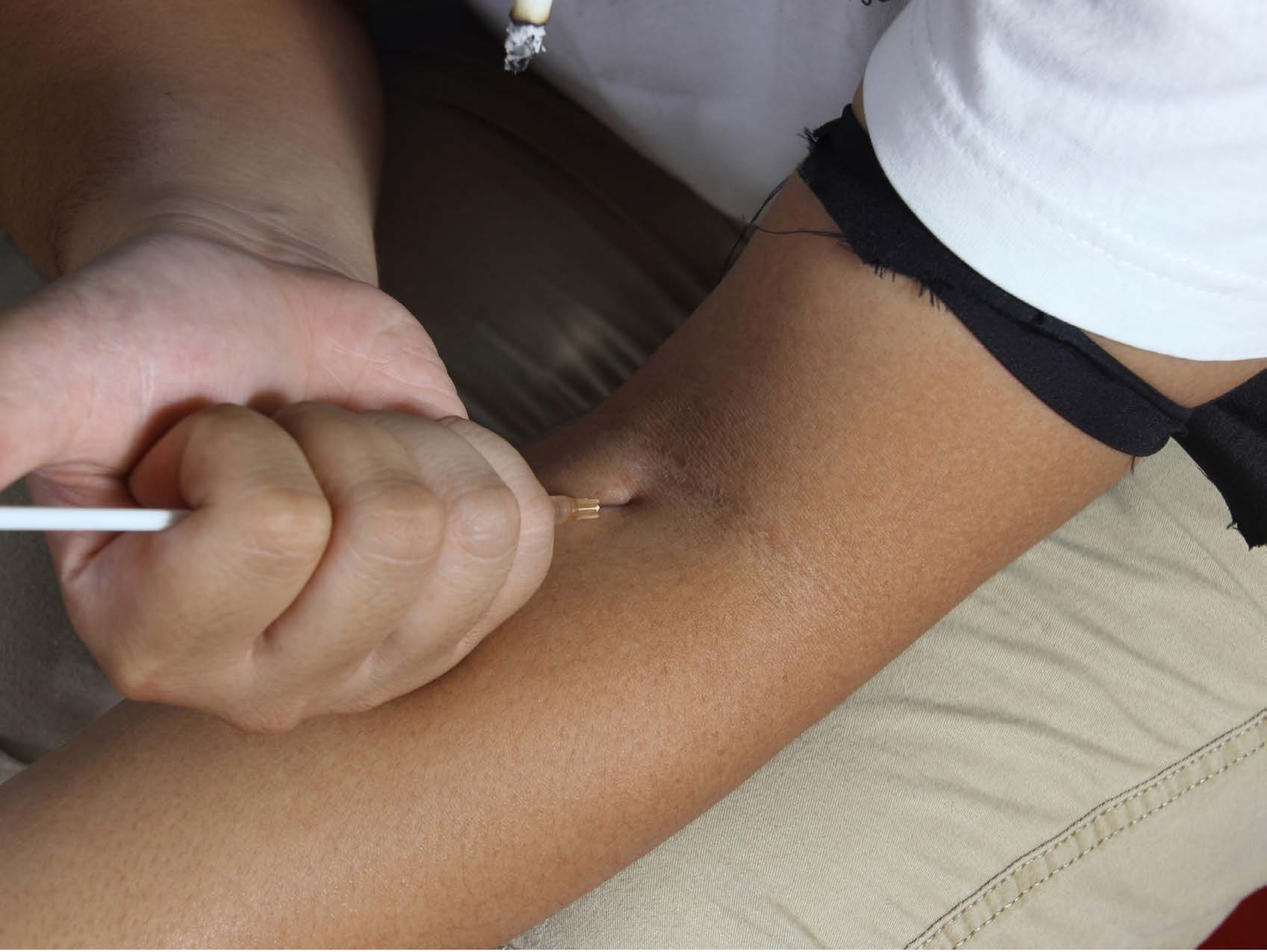

| Category | Craving (0-100) | Valence (1-9) | Arousal (1-9) | Typicality (0-100) | Relatedness | | | | | HSV | | |
|---|---|---|---|---|---|---|---|---|---|---|---|---|
| | | | | | Meth | Opioid | Both | Neither | MethToOpioid | Hue | Saturation | Value |
| Opioid Instrument Hand | 90.59 (15.88) | 4.19 (2.167) | 5.11 (2.89) | 91.04 (15.79) | 0.111 | 0.037 | 0.852 | 0 | -0.074 | 0.178 (0.224) | 0.281 (0.152) | 0.56 (0.236) |

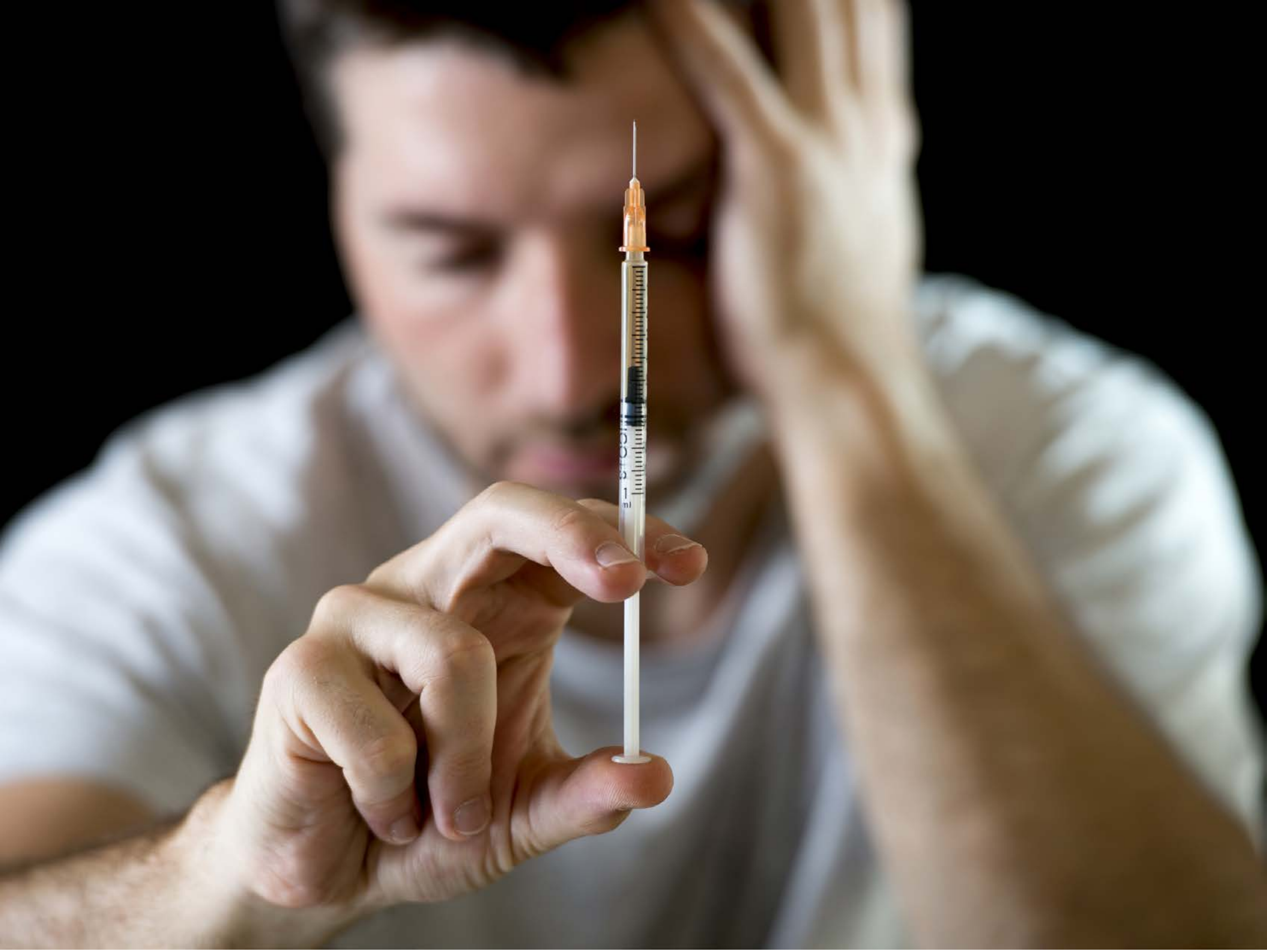

| Category | Craving (0-100) | Valence (1-9) | Arousal (1-9) | Typicality (0-100) | Relatedness | | | | | HSV | | |
|---|---|---|---|---|---|---|---|---|---|---|---|---|
| | | | | | Meth | Opioid | Both | Neither | MethToOpioid | Hue | Saturation | Value |
| Opioid Face Activities | 88.86 (22.33) | 4.04 (2.099) | 4.93 (2.75) | 90.39 (16.41) | 0.107 | 0.179 | 0.714 | 0 | 0.071 | 0.152 (0.213) | 0.22 (0.239) | 0.445 (0.303) |

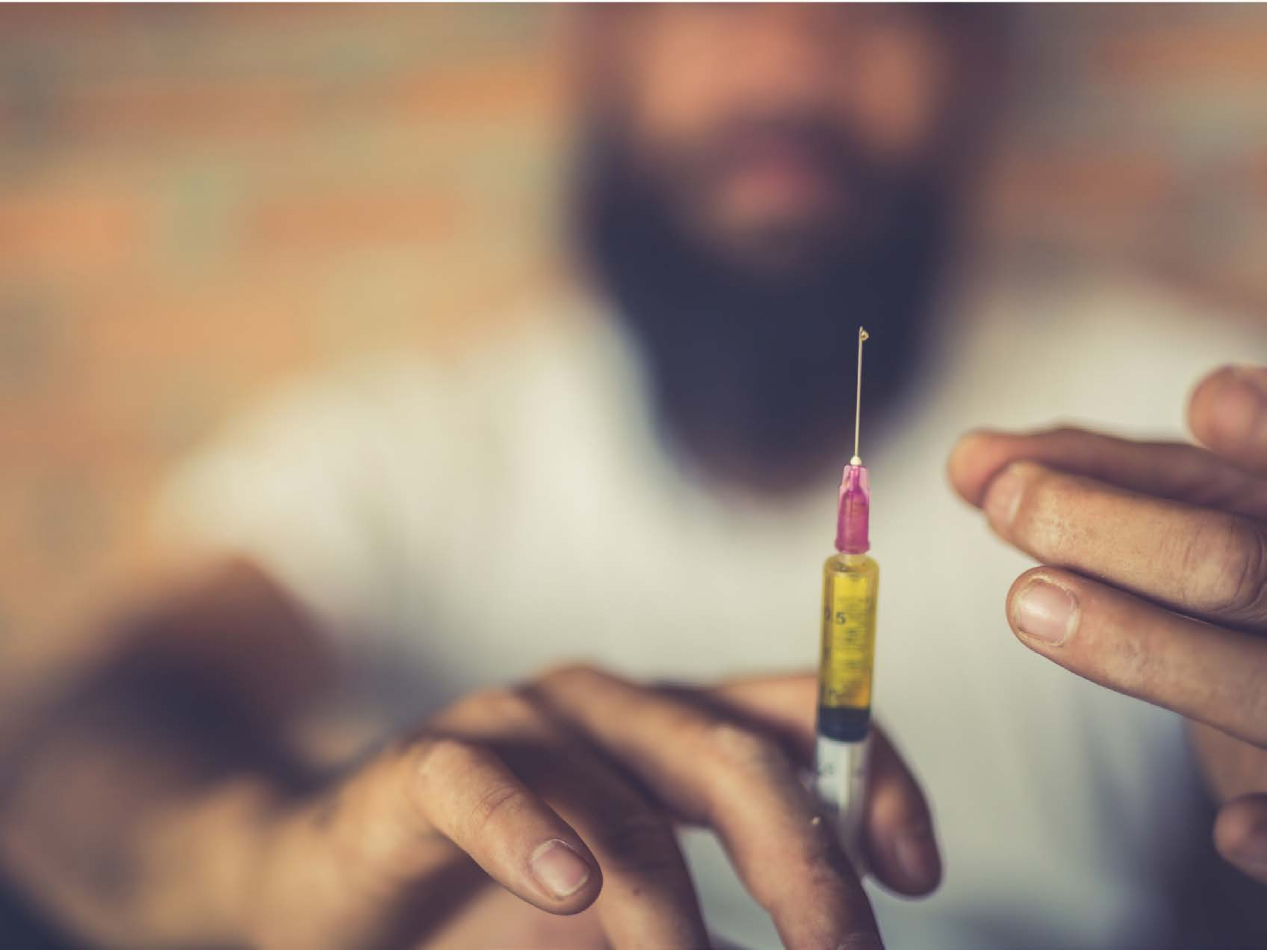

| Category | Craving (0-100) | Valence (1-9) | Arousal (1-9) | Typicality (0-100) | Relatedness | | | | | HSV | | |
|---|---|---|---|---|---|---|---|---|---|---|---|---|
| | | | | | Meth | Opioid | Both | Neither | MethToOpioid | Hue | Saturation | Value |
| Opioid Face Activities | 89.93 (13.6) | 4.63 (1.944) | 4.78 (2.59) | 89.63 (14.37) | 0.111 | 0.111 | 0.778 | 0 | 0 | 0.278 (0.328) | 0.271 (0.119) | 0.615 (0.19) |

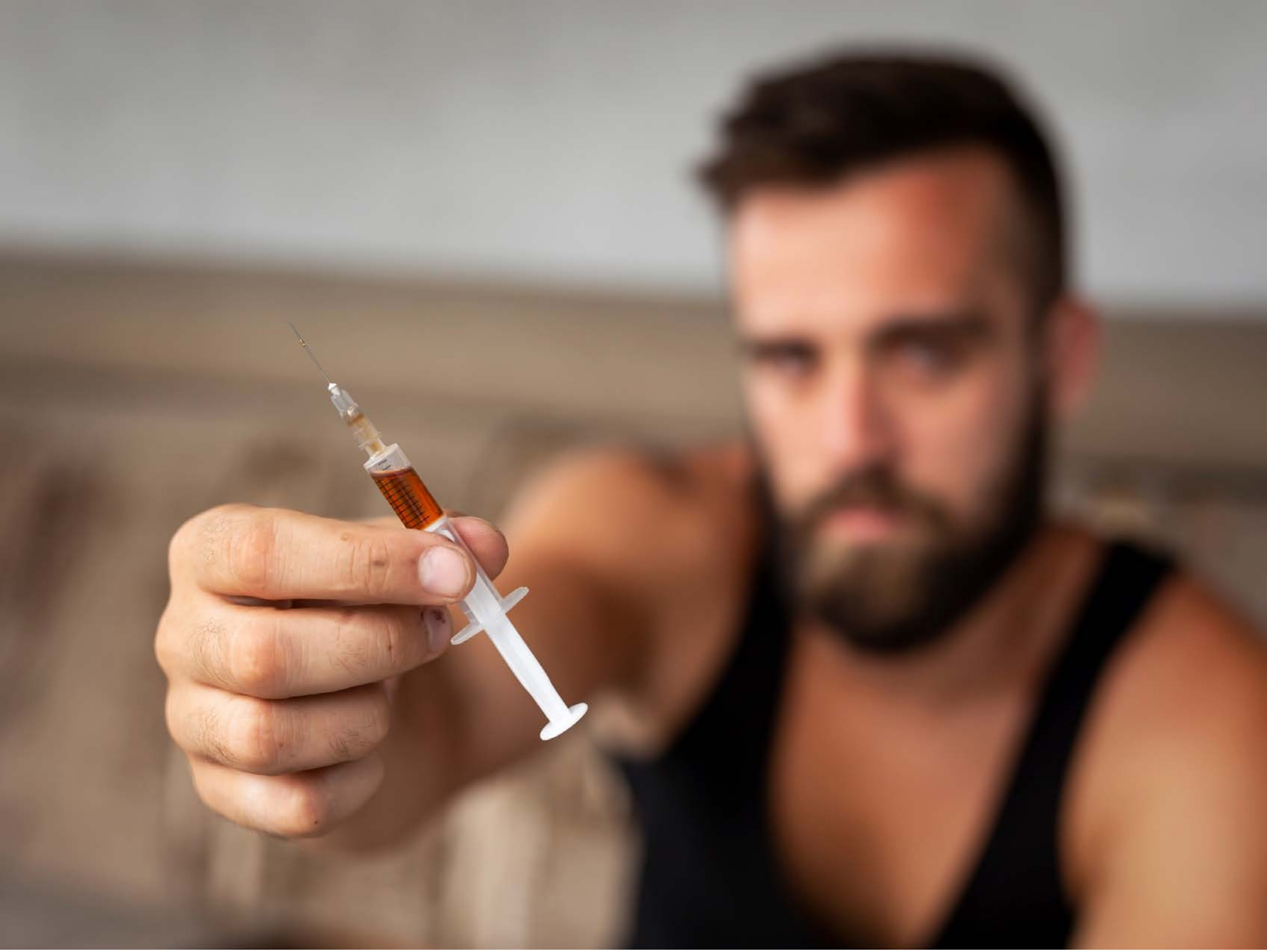

| Category | Craving (0-100) | Valence (1-9) | Arousal (1-9) | Typicality (0-100) | Relatedness | | | | | HSV | | |
|---|---|---|---|---|---|---|---|---|---|---|---|---|
| | | | | | Meth | Opioid | Both | Neither | MethToOpioid | Hue | Saturation | Value |
| Opioid Face Activities | 91 (19.21) | 4.07 (2.035) | 4.89 (2.99) | 92.14 (19.36) | 0.036 | 0.25 | 0.679 | 0.036 | 0.214 | 0.104 (0.158) | 0.345 (0.235) | 0.538 (0.22) |

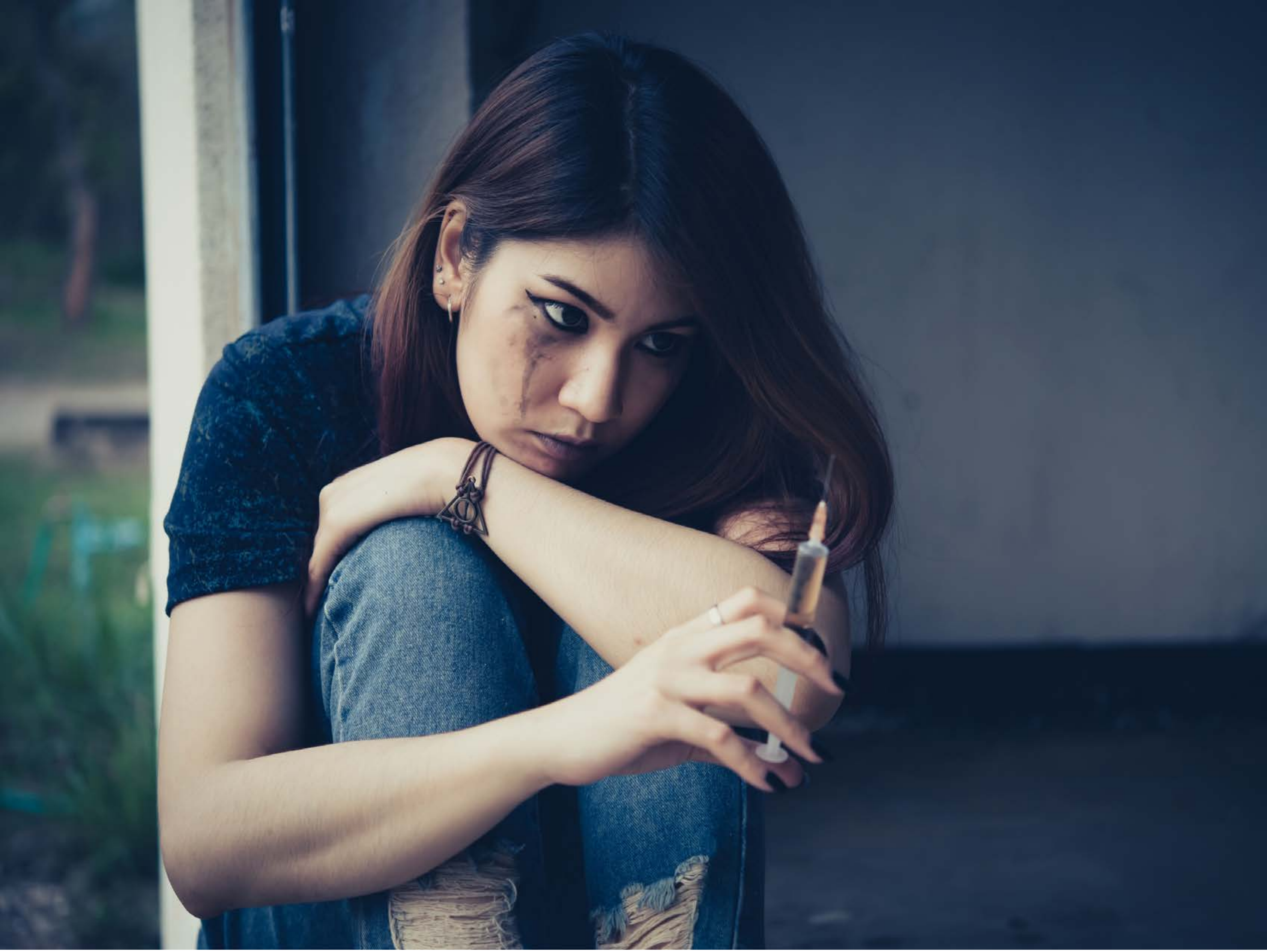

| Category | Craving (0-100) | Valence (1-9) | Arousal (1-9) | Typicality (0-100) | Relatedness | | | | | HSV | | |
|---|---|---|---|---|---|---|---|---|---|---|---|---|
| | | | | | Meth | Opioid | Both | Neither | MethToOpioid | Hue | Saturation | Value |
| Opioid Face Activities | 87.59 (15.96) | 4.19 (2.149) | 4.56 (2.74) | 88.63 (16.57) | 0.074 | 0.111 | 0.778 | 0.037 | 0.037 | 0.519 (0.197) | 0.447 (0.234) | 0.385 (0.2) |

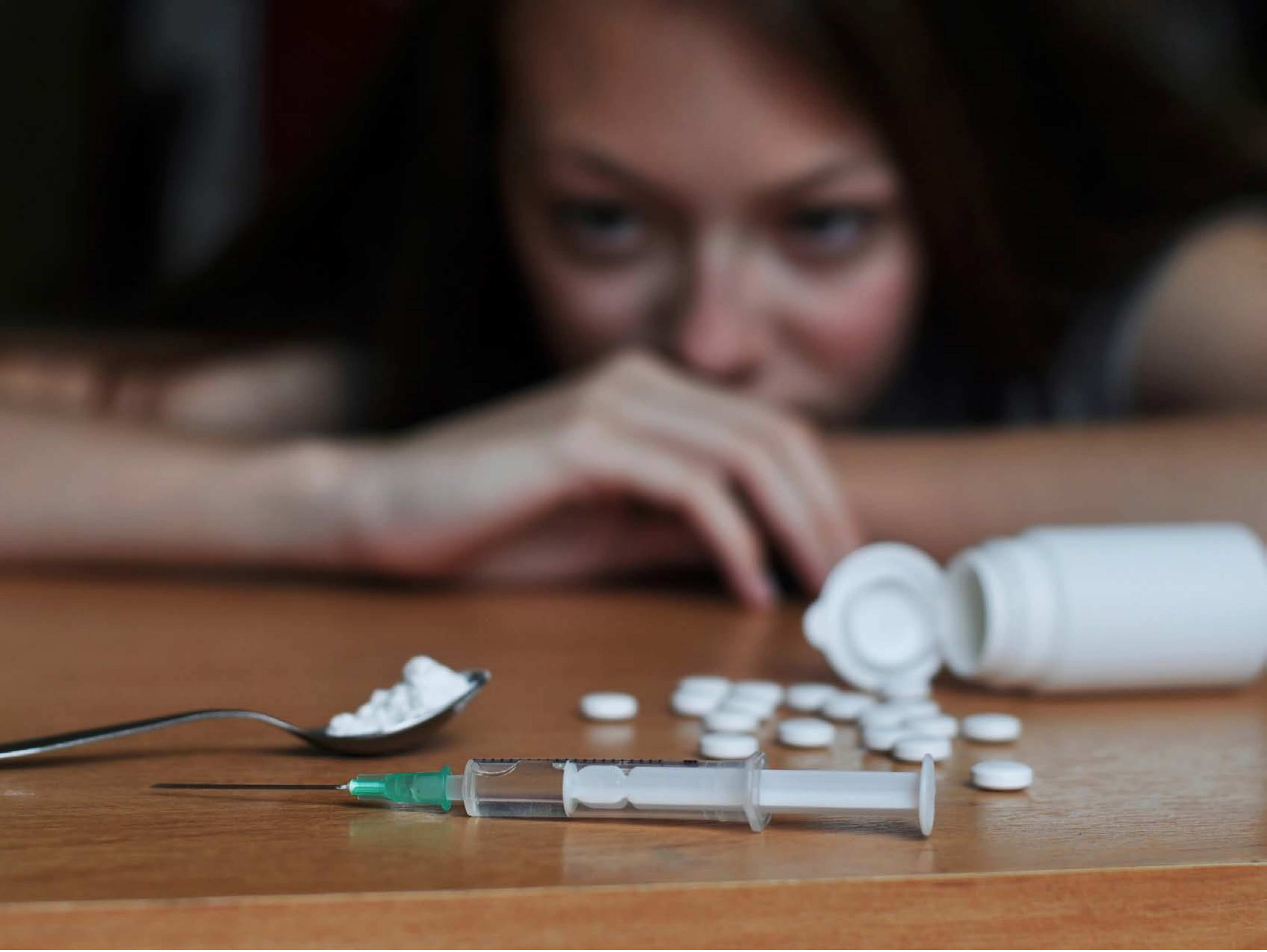

| Category | Craving (0-100) | Valence (1-9) | Arousal (1-9) | Typicality (0-100) | Relatedness | | | | | HSV | | |
|---|---|---|---|---|---|---|---|---|---|---|---|---|
| | | | | | Meth | Opioid | Both | Neither | MethToOpioid | Hue | Saturation | Value |
| Opioid Face Activities | 85.57 (19.36) | 4.61 (1.595) | 4.39 (2.73) | 91.46 (13.64) | 0 | 0.571 | 0.429 | 0 | 0.571 | 0.169 (0.25) | 0.366 (0.206) | 0.381 (0.253) |

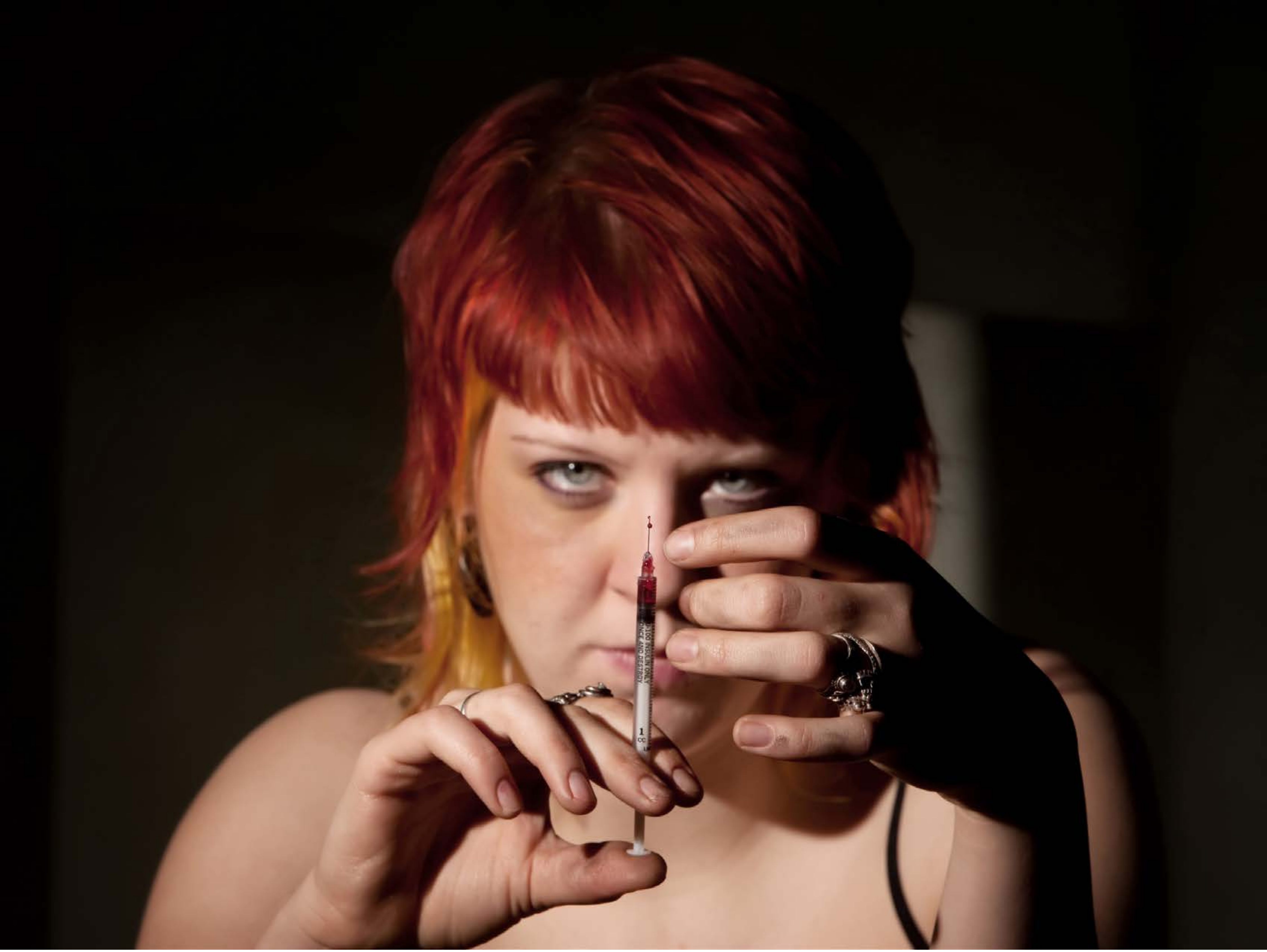

| Category | Craving (0-100) | Valence (1-9) | Arousal (1-9) | Typicality (0-100) | Relatedness | | | | | HSV | | |
|---|---|---|---|---|---|---|---|---|---|---|---|---|
| | | | | | Meth | Opioid | Both | Neither | MethToOpioid | Hue | Saturation | Value |
| Opioid Face Activities | 87.96 (17.41) | 4.33 (2.038) | 4.81 (2.54) | 87.89 (18.22) | 0.111 | 0.148 | 0.741 | 0 | 0.037 | 0.156 (0.268) | 0.662 (0.288) | 0.227 (0.287) |

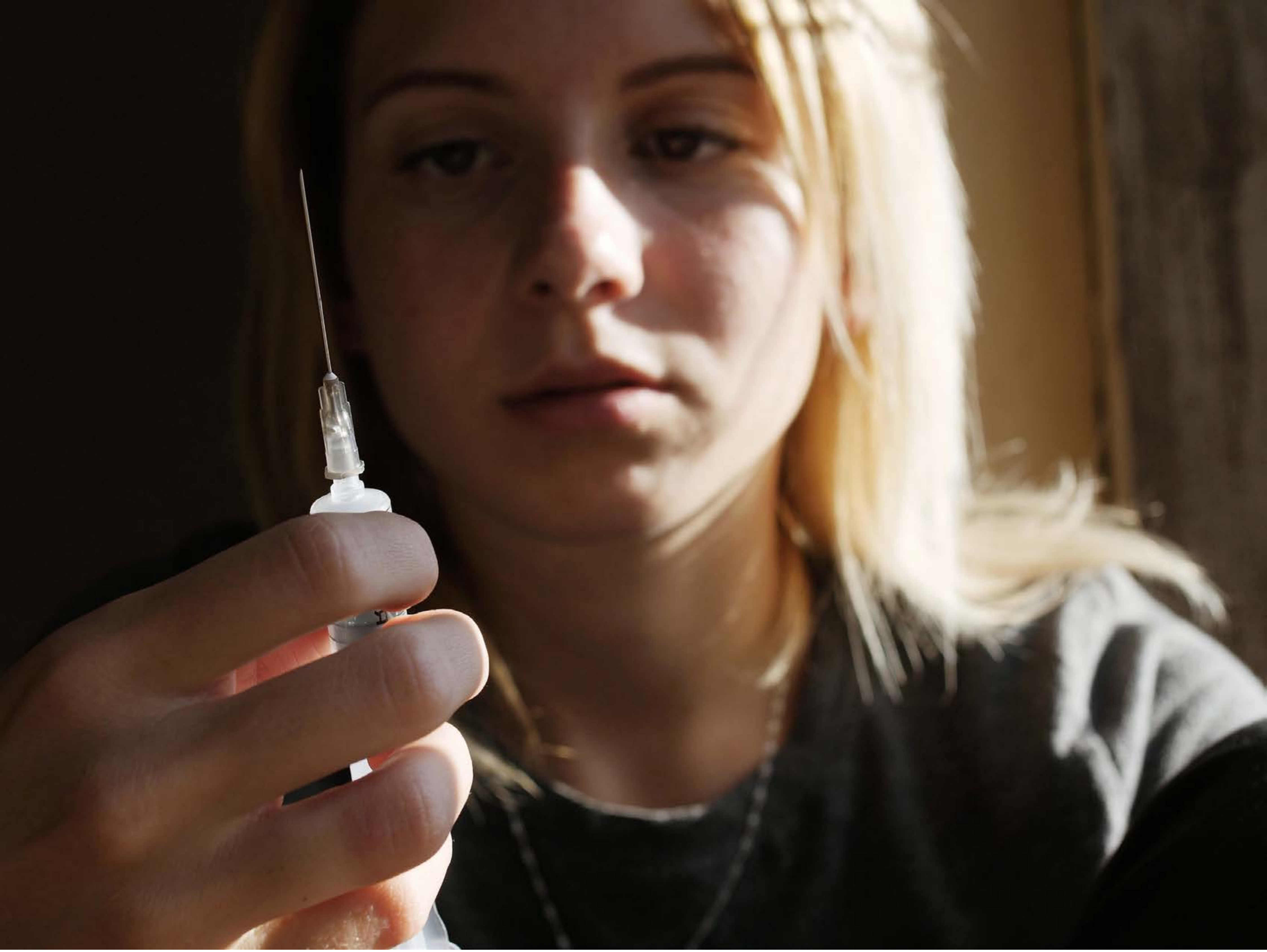

| Category | Craving (0-100) | Valence (1-9) | Arousal (1-9) | Typicality (0-100) | Relatedness | | | | | HSV | | |
|---|---|---|---|---|---|---|---|---|---|---|---|---|
| | | | | | Meth | Opioid | Both | Neither | MethToOpioid | Hue | Saturation | Value |
| Opioid Face Activities | 87.37 (18.41) | 4.41 (2.135) | 4.48 (2.68) | 89.93 (16.64) | 0.148 | 0.037 | 0.815 | 0 | -0.111 | 0.088 (0.069) | 0.431 (0.213) | 0.339 (0.282) |

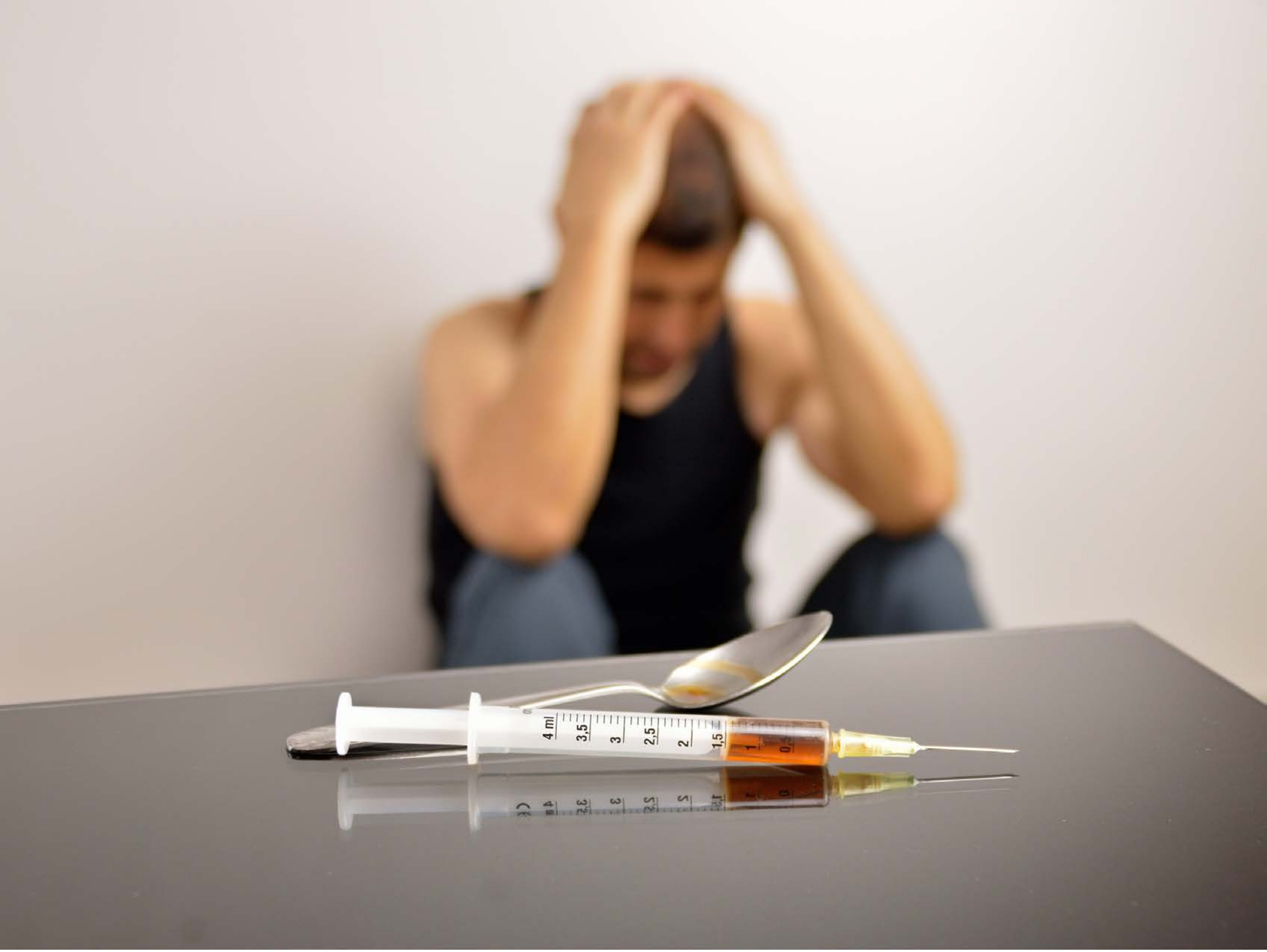


| Category | Craving (0-100) | Valence (1-9) | Arousal (1-9) | Typicality (0-100) | Relatedness | | | | | HSV | | |
|---|---|---|---|---|---|---|---|---|---|---|---|---|
| | | | | | Meth | Opioid | Both | Neither | MethToOpioid | Hue | Saturation | Value |
| Opioid Face Activities | 86.26 (19.21) | 4.07 (1.88) | 4.59 (2.63) | 84.52 (21.41) | 0.111 | 0.407 | 0.481 | 0 | 0.296 | 0.101 (0.15) | 0.155 (0.163) | 0.703 (0.226) |

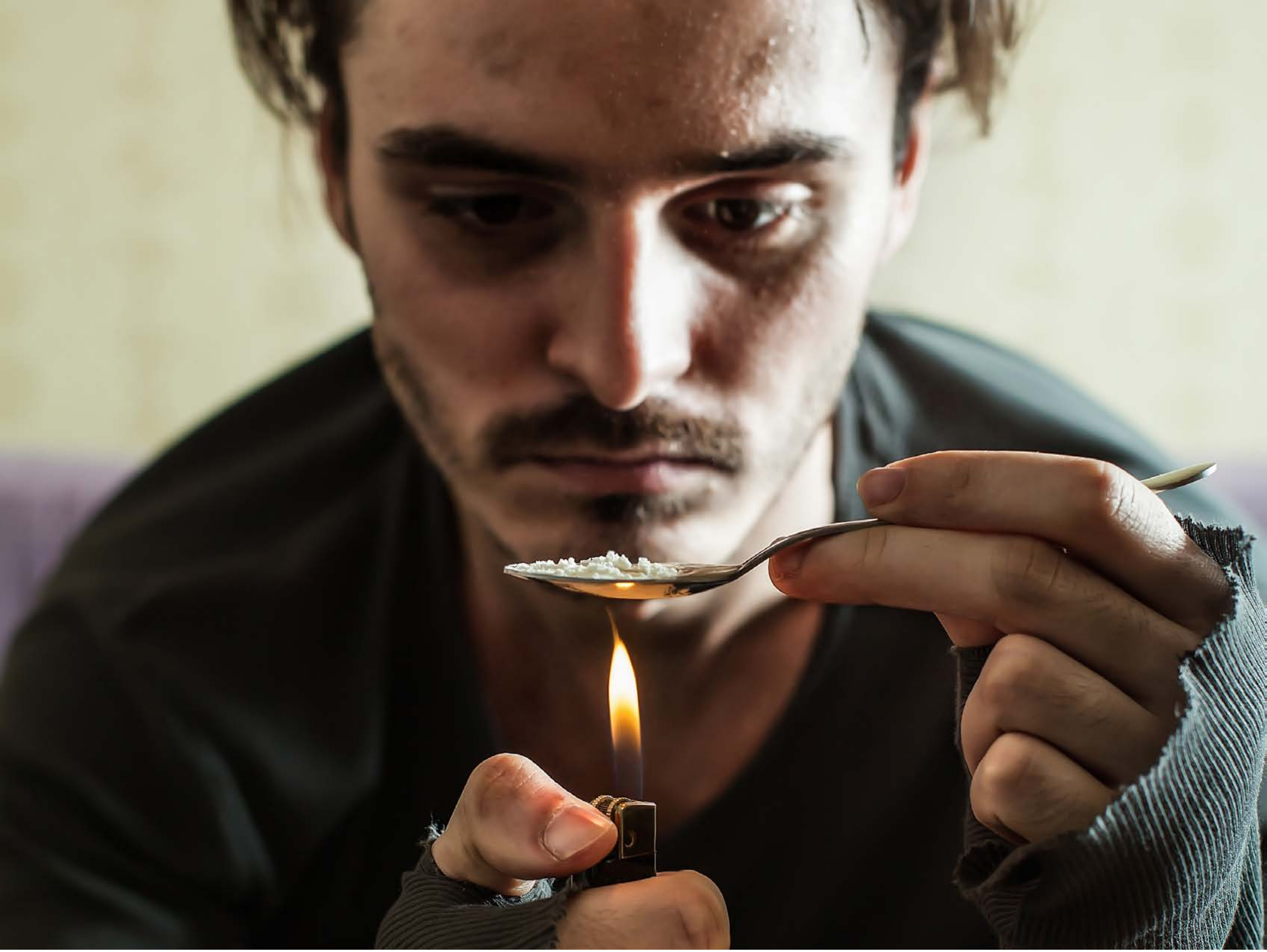

| Category | Craving (0-100) | Valence (1-9) | Arousal (1-9) | Typicality (0-100) | Relatedness | | | | | HSV | | |
|---|---|---|---|---|---|---|---|---|---|---|---|---|
| | | | | | Meth | Opioid | Both | Neither | MethToOpioid | Hue | Saturation | Value |
| Opioid Face Activities | 92.54 (13.96) | 4.25 (1.917) | 4.86 (2.72) | 94.36 (10.13) | 0.143 | 0.464 | 0.393 | 0 | 0.321 | 0.137 (0.142) | 0.325 (0.199) | 0.426 (0.318) |

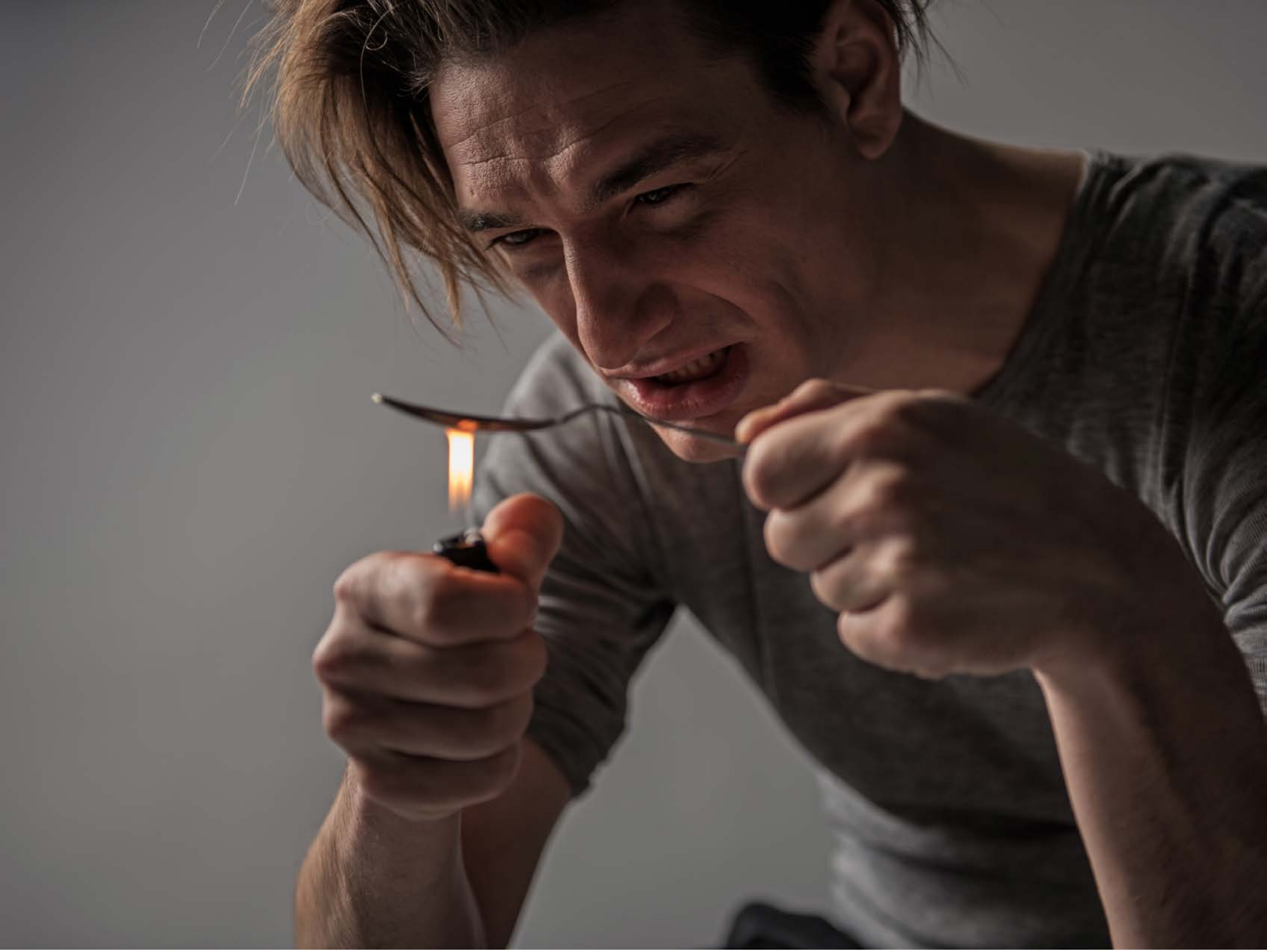

| Category | Craving (0-100) | Valence (1-9) | Arousal (1-9) | Typicality (0-100) | Relatedness | | | | | HSV | | |
|---|---|---|---|---|---|---|---|---|---|---|---|---|
| | | | | | Meth | Opioid | Both | Neither | MethToOpioid | Hue | Saturation | Value |
| Opioid Face Activities | 89.75 (20.56) | 4.43 (1.731) | 4.75 (2.73) | 91.5 (20.28) | 0.143 | 0.357 | 0.5 | 0 | 0.214 | 0.18 (0.309) | 0.229 (0.197) | 0.319 (0.162) |

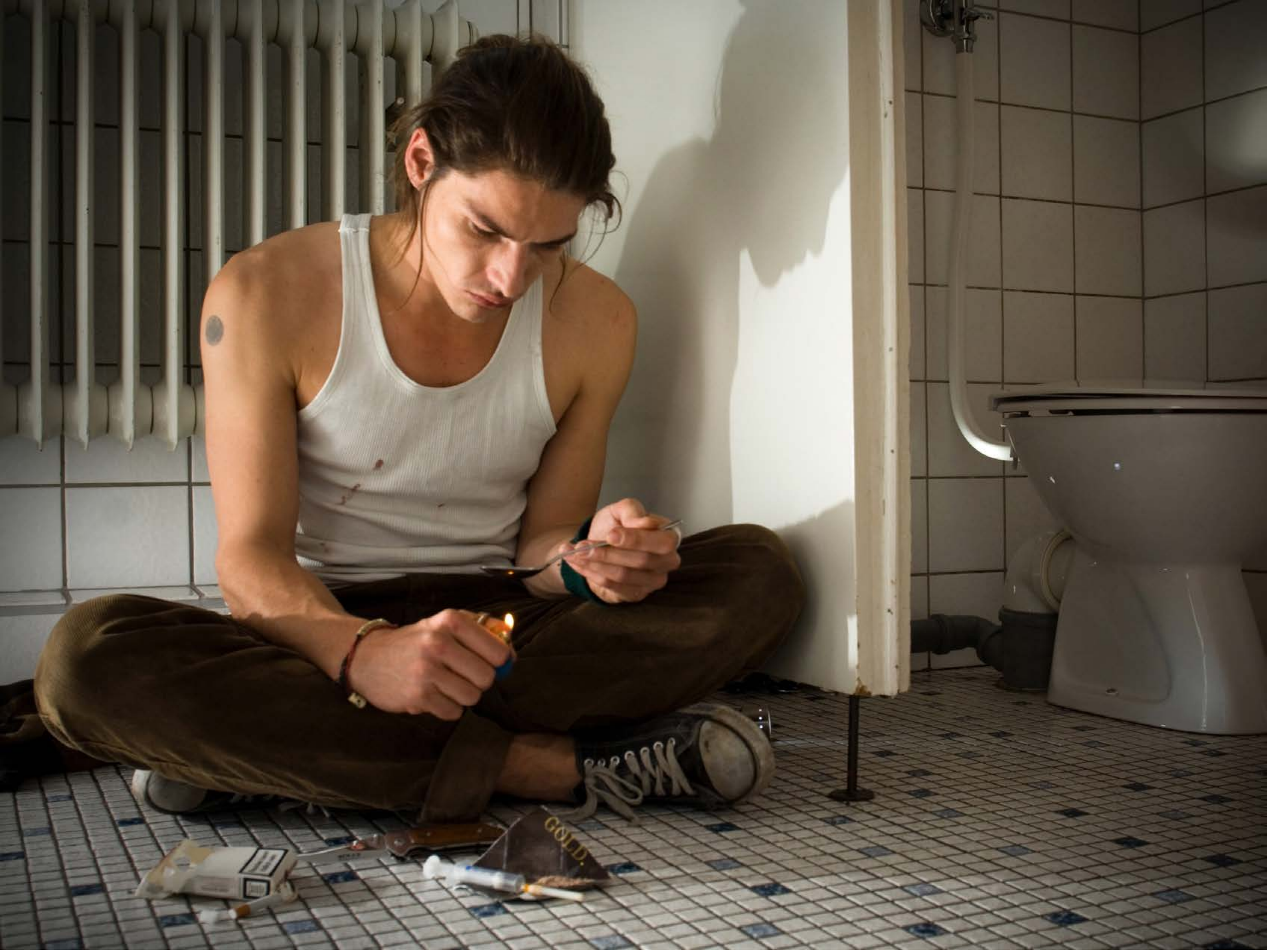


| Category | Craving (0-100) | Valence (1-9) | Arousal (1-9) | Typicality (0-100) | Relatedness | | | | | HSV | | |
|---|---|---|---|---|---|---|---|---|---|---|---|---|
| | | | | | Meth | Opioid | Both | Neither | MethToOpioid | Hue | Saturation | Value |
| Opioid Face Activities | 90.78 (14.5) | 4.07 (2.129) | 4.67 (2.75) | 89.78 (17.9) | 0.111 | 0.519 | 0.370 | 0 | 0.407 | 0.111 (0.066) | 0.318 (0.221) | 0.387 (0.225) |

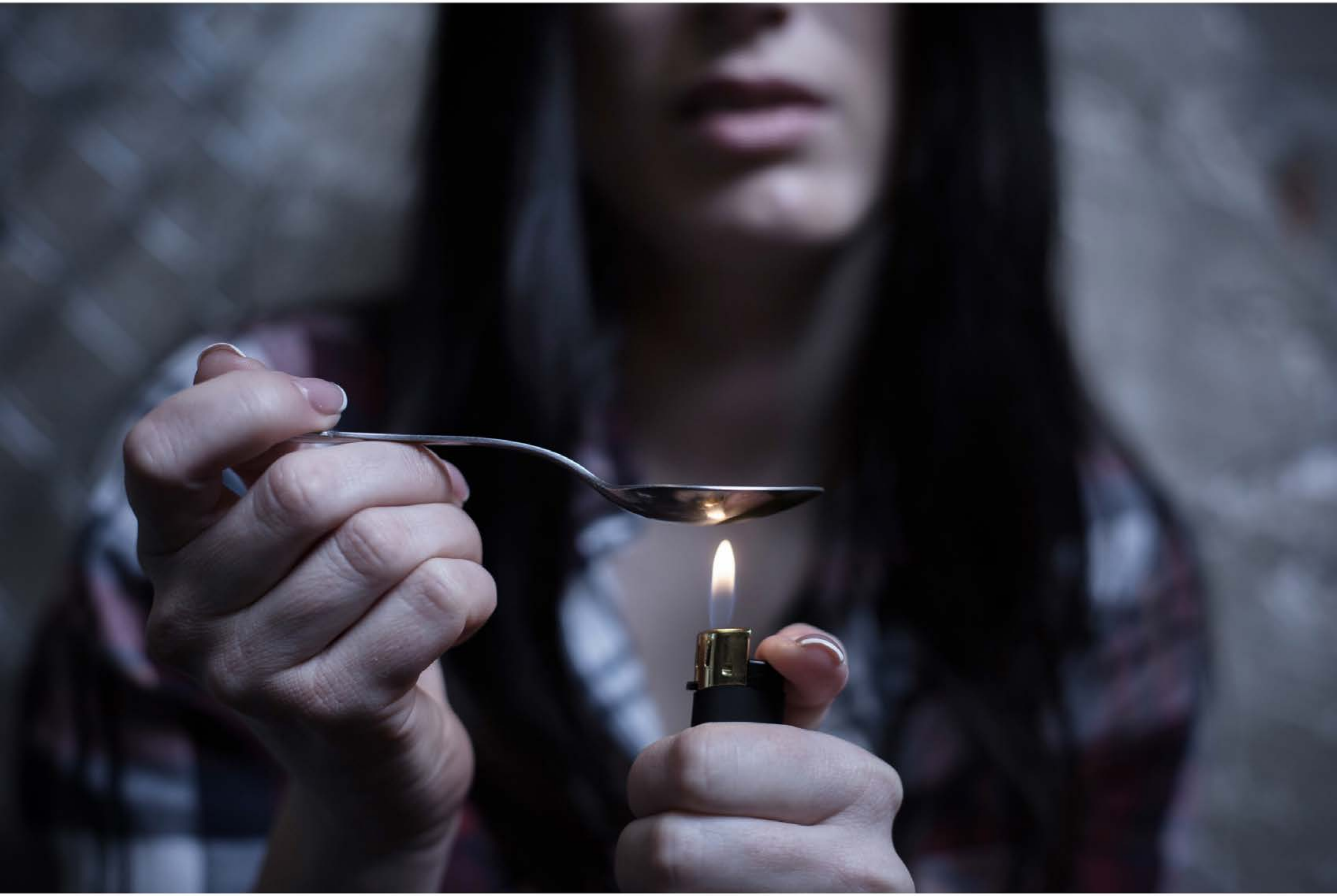

| Category | Craving (0-100) | Valence (1-9) | Arousal (1-9) | Typicality (0-100) | Relatedness | | | | | HSV | | |
|---|---|---|---|---|---|---|---|---|---|---|---|---|
| | | | | | Meth | Opioid | Both | Neither | MethToOpioid | Hue | Saturation | Value |
| Opioid Face Activities | 90.81 (12.46) | 4.37 (1.822) | 4.67 (2.66) | 91.85 (11.8) | 0.074 | 0.444 | 0.481 | 0 | 0.370 | 0.571 (0.297) | 0.214 (0.144) | 0.347 (0.303) |

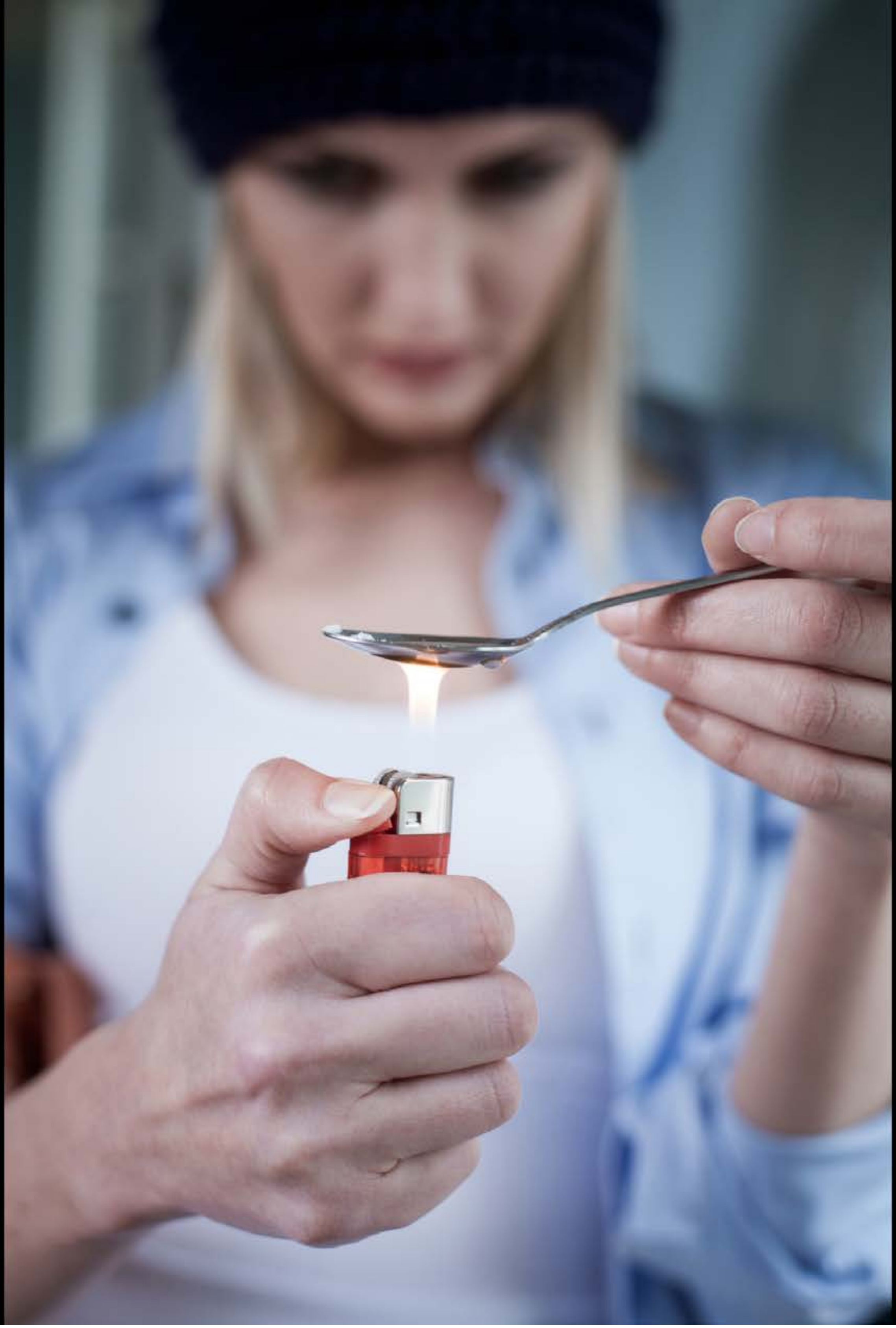

| Category | Craving (0-100) | Valence (1-9) | Arousal (1-9) | Typicality (0-100) | Relatedness | | | | | HSV | | |
|---|---|---|---|---|---|---|---|---|---|---|---|---|
| | | | | | Meth | Opioid | Both | Neither | MethToOpioid | Hue | Saturation | Value |
| Opioid Face Activities | 93.11 (11.47) | 4.39 (1.729) | 4.46 (2.83) | 94.11 (10.35) | 0.036 | 0.429 | 0.536 | 0 | 0.393 | 0.254 (0.338) | 0.13 (0.187) | 0.301 (0.342) |

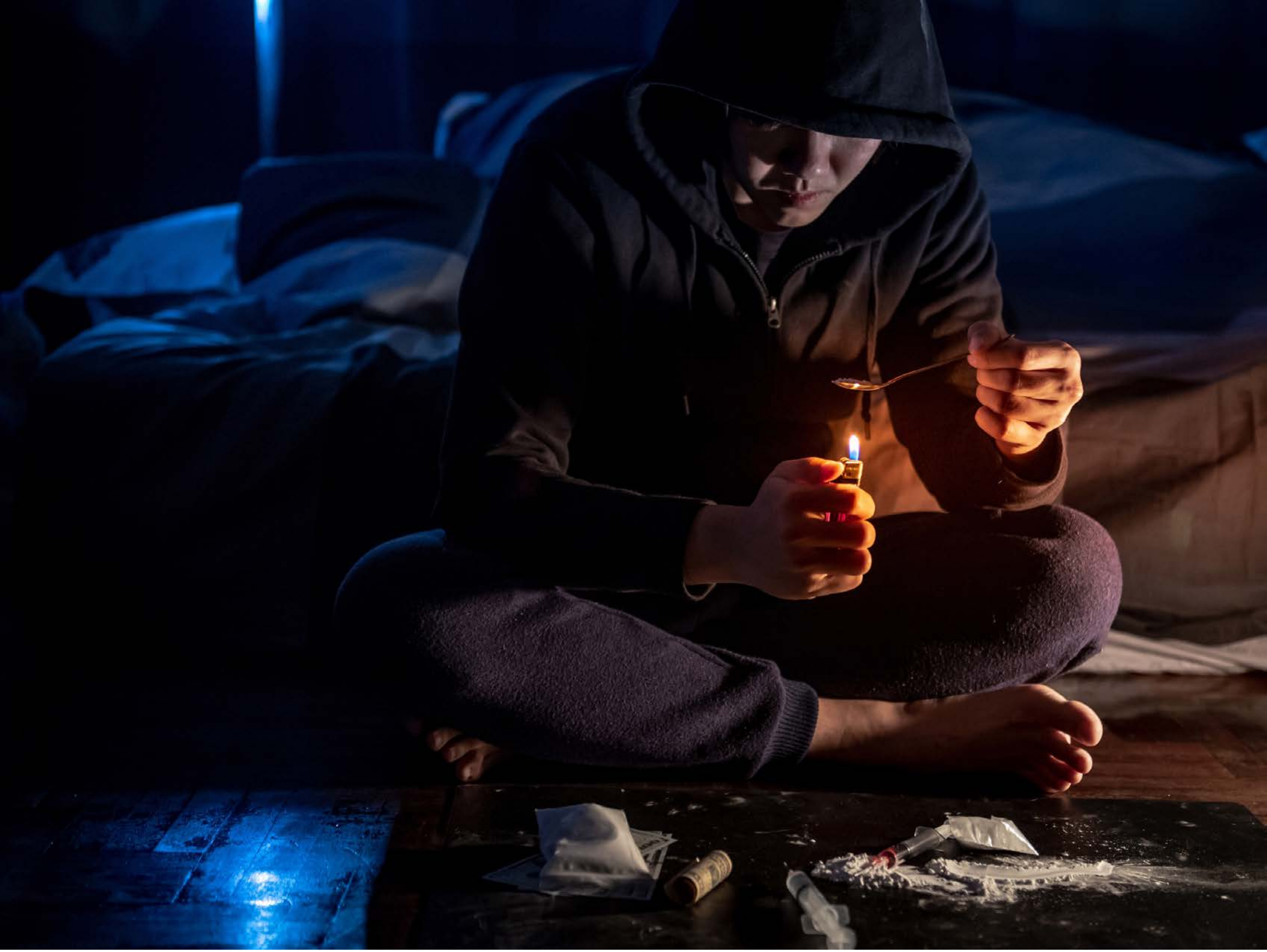

| Category | Craving (0-100) | Valence (1-9) | Arousal (1-9) | Typicality (0-100) | Relatedness | | | | | HSV | | |
|---|---|---|---|---|---|---|---|---|---|---|---|---|
| | | | | | Meth | Opioid | Both | Neither | MethToOpioid | Hue | Saturation | Value |
| Opioid Face Activities | 86.46 (22.44) | 4.25 (1.936) | 4.57 (2.86) | 91.71 (13.62) | 0.107 | 0.393 | 0.5 | 0 | 0.286 | 0.48 (0.329) | 0.695 (0.316) | 0.139 (0.172) |

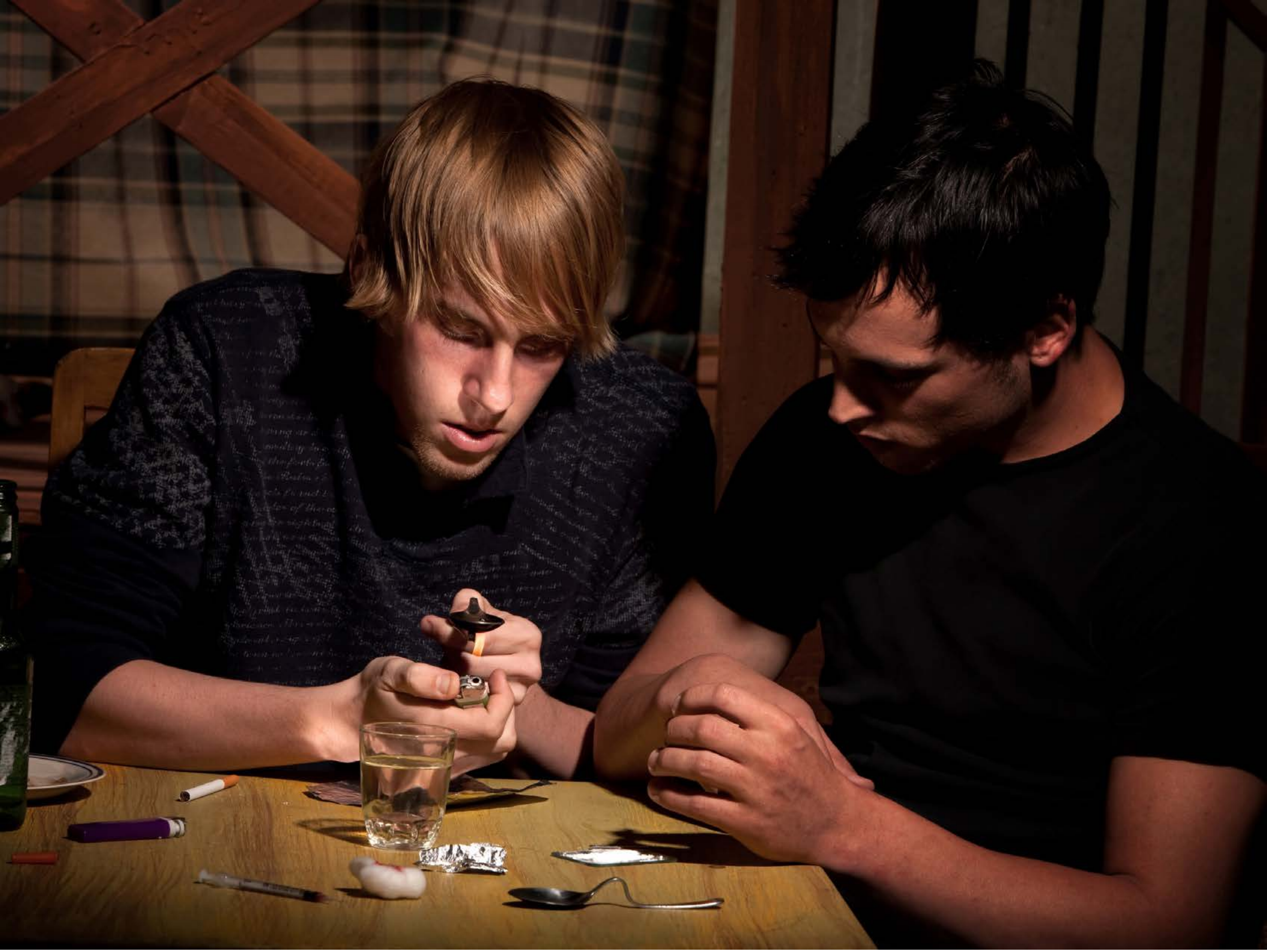

| Category | Craving (0-100) | Valence (1-9) | Arousal (1-9) | Typicality (0-100) | Relatedness | | | | | HSV | | |
|---|---|---|---|---|---|---|---|---|---|---|---|---|
| | | | | | Meth | Opioid | Both | Neither | MethToOpioid | Hue | Saturation | Value |
| Opioid Face Activities | 86.56 (18.87) | 4.3 (2.016) | 4.89 (2.72) | 88.22 (18.27) | 0.037 | 0.407 | 0.556 | 0 | 0.370 | 0.188 (0.31) | 0.466 (0.309) | 0.205 (0.222) |

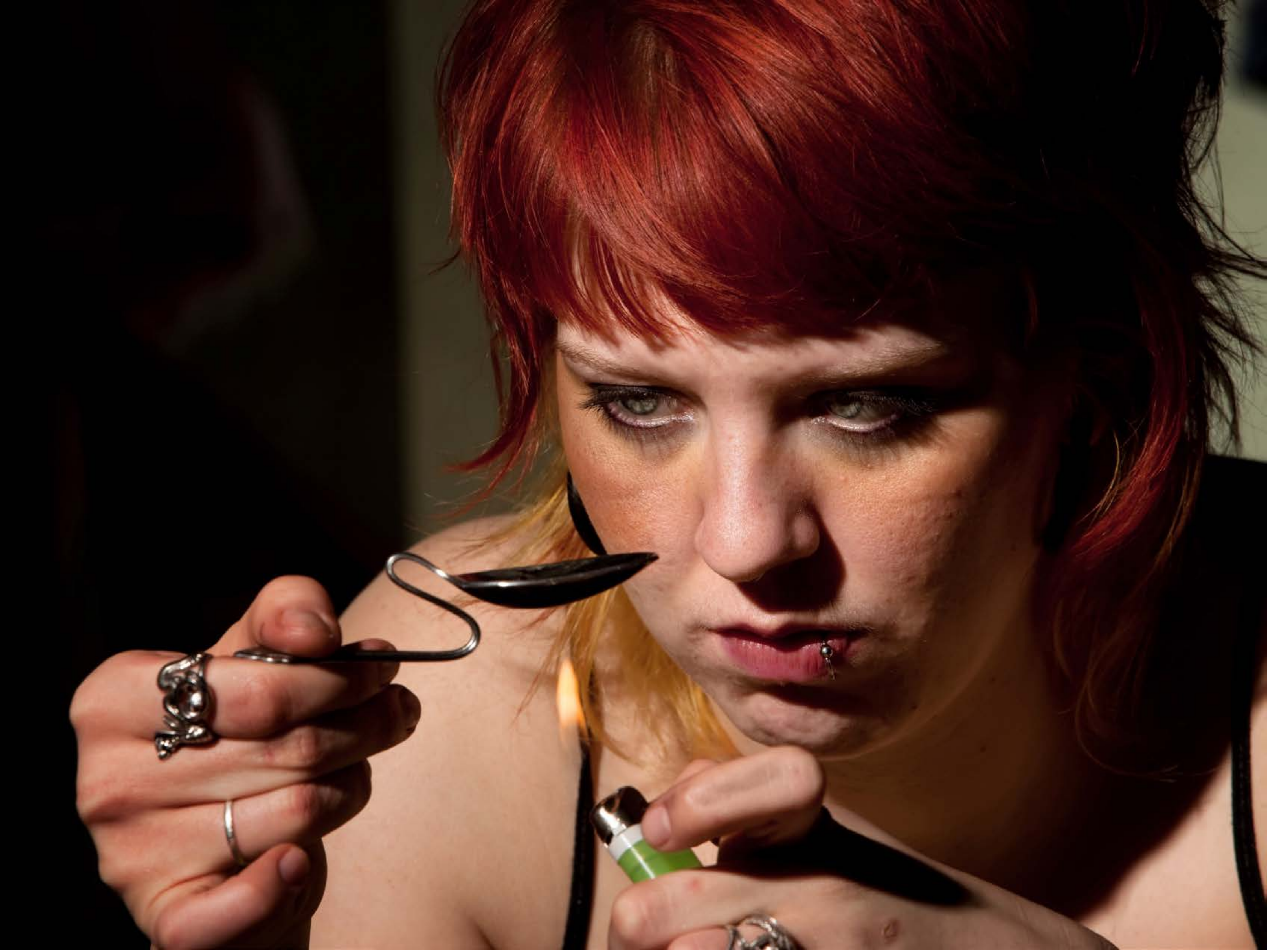

| Category | Craving (0-100) | Valence (1-9) | Arousal (1-9) | Typicality (0-100) | Relatedness | | | | | HSV | | |
|---|---|---|---|---|---|---|---|---|---|---|---|---|
| | | | | | Meth | Opioid | Both | Neither | MethToOpioid | Hue | Saturation | Value |
| Opioid Face Activities | 87.44 (18.43) | 4.37 (1.843) | 4.59 (2.68) | 86.48 (19.4) | 0.074 | 0.481 | 0.407 | 0.037 | 0.407 | 0.191 (0.345) | 0.591 (0.342) | 0.294 (0.301) |

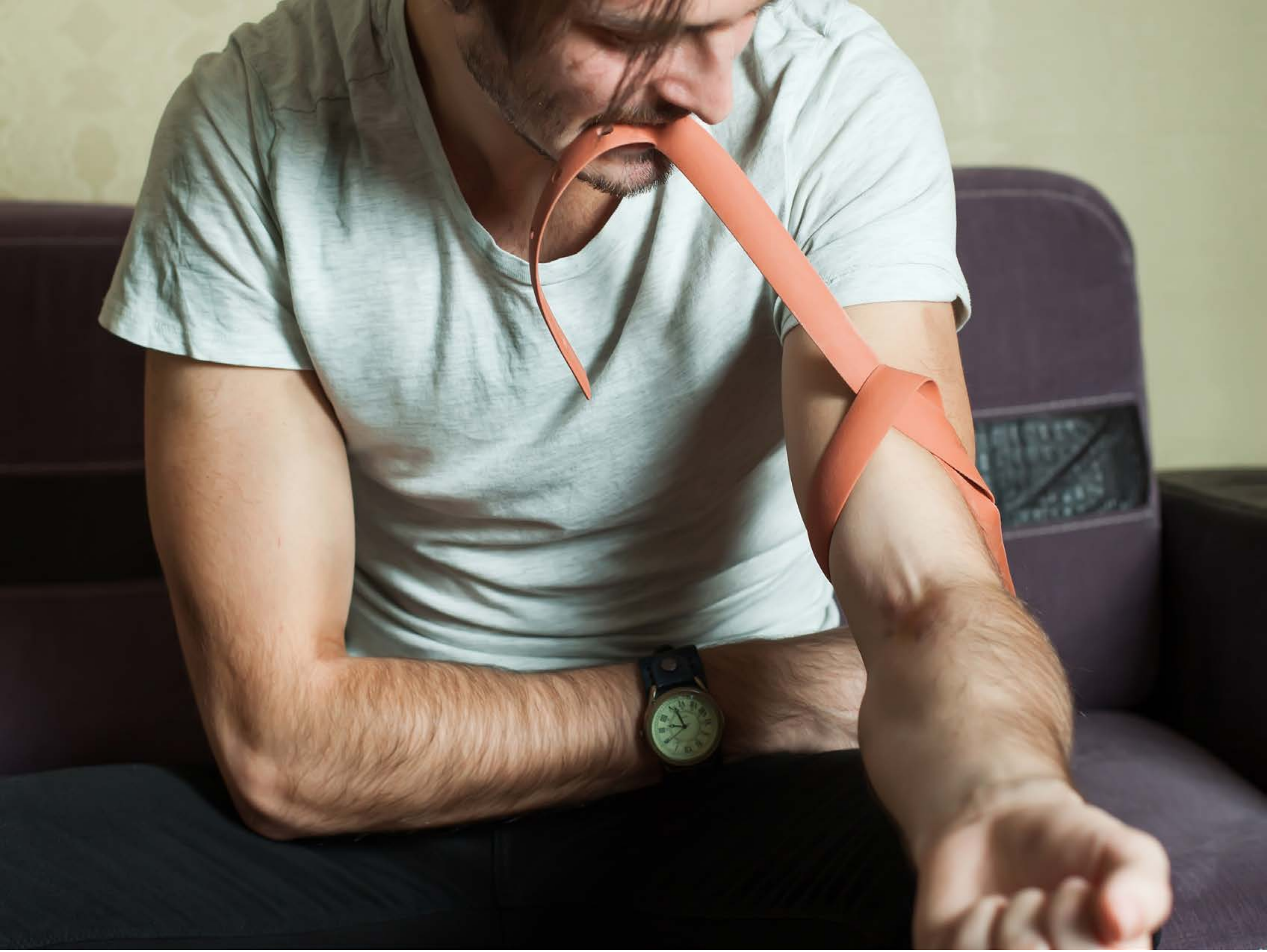

| Category | Craving (0-100) | Valence (1-9) | Arousal (1-9) | Typicality (0-100) | Relatedness | | | | | HSV | | |
|---|---|---|---|---|---|---|---|---|---|---|---|---|
| | | | | | Meth | Opioid | Both | Neither | MethToOpioid | Hue | Saturation | Value |
| Opioid Face Activities | 88.3 (15.23) | 4.33 (2) | 4.78 (2.62) | 86.93 (16.59) | 0.111 | 0 | 0.889 | 0 | -0.111 | 0.284 (0.277) | 0.251 (0.171) | 0.475 (0.305) |

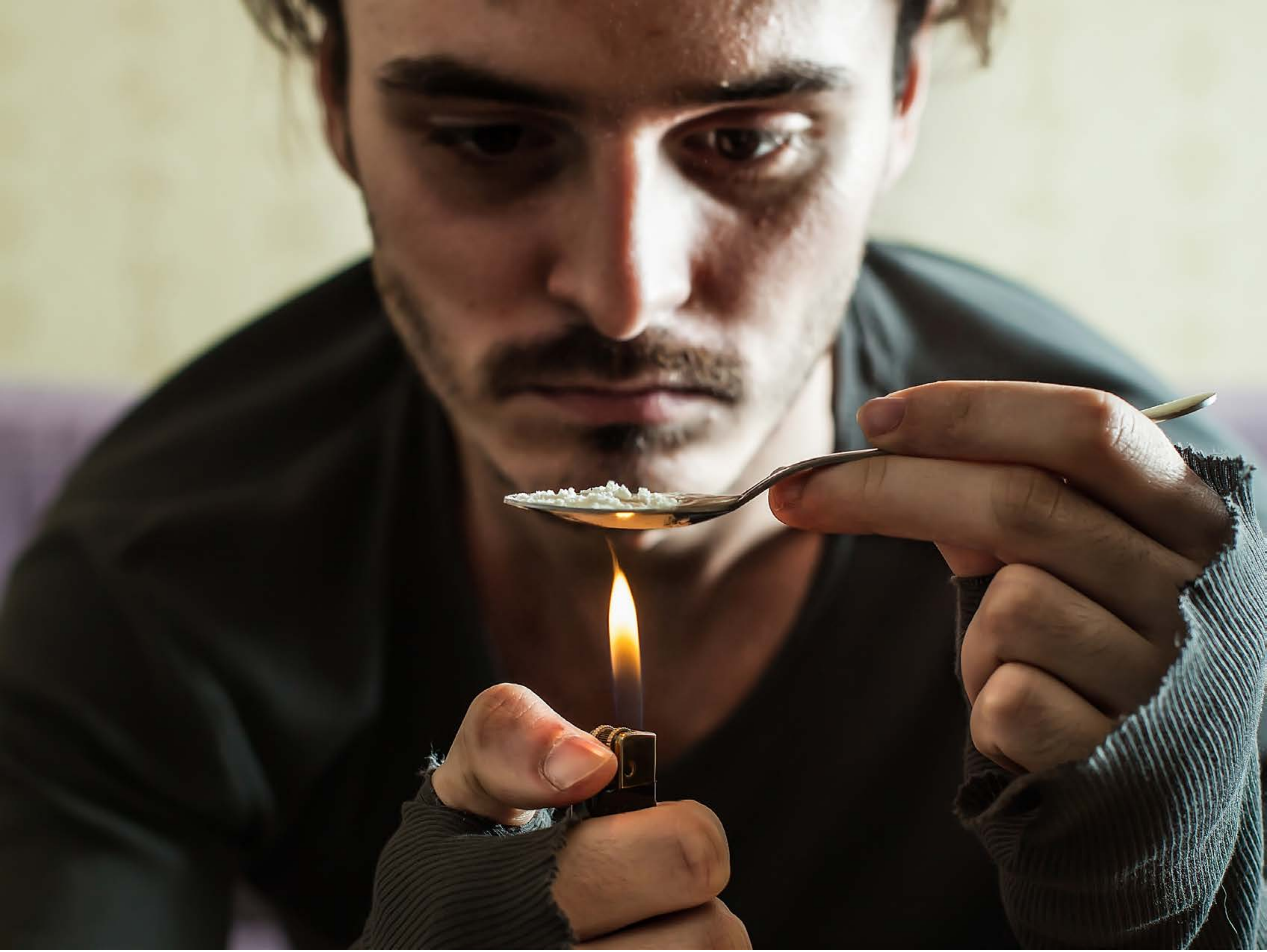

| Category | Craving (0-100) | Valence (1-9) | Arousal (1-9) | Typicality (0-100) | Relatedness | | | | | HSV | | |
|---|---|---|---|---|---|---|---|---|---|---|---|---|
| | | | | | Meth | Opioid | Both | Neither | MethToOpioid | Hue | Saturation | Value |
| Opioid Face Activities | 90.96 (14.13) | 4.18 (2.019) | 5.07 (2.96) | 92.54 (13.69) | 0.179 | 0.464 | 0.357 | 0 | 0.286 | 0.14 (0.15) | 0.329 (0.197) | 0.392 (0.318) |

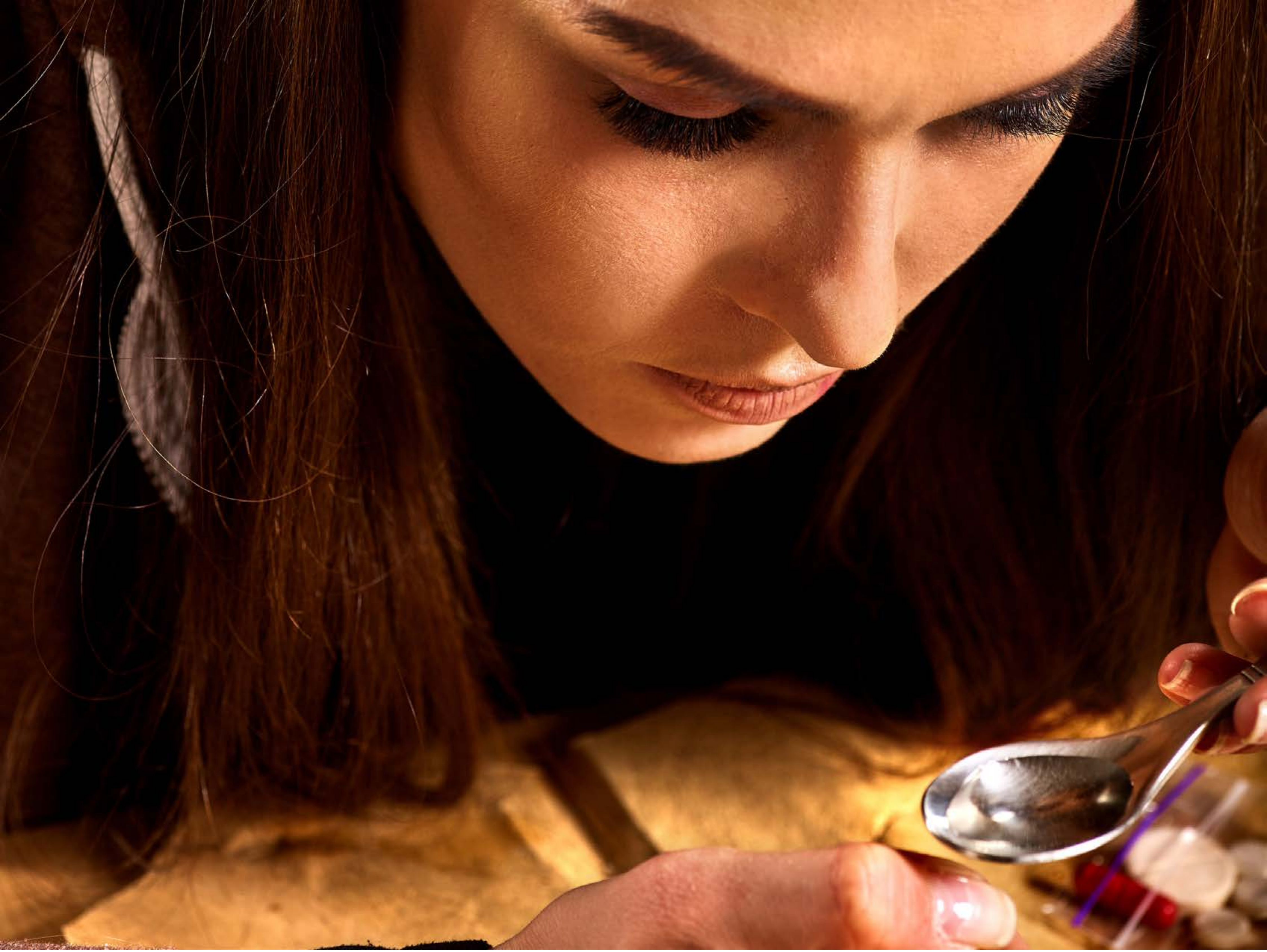

| Category | Craving (0-100) | Valence (1-9) | Arousal (1-9) | Typicality (0-100) | Relatedness | | | | | HSV | | |
|---|---|---|---|---|---|---|---|---|---|---|---|---|
| | | | | | Meth | Opioid | Both | Neither | MethToOpioid | Hue | Saturation | Value |
| Opioid Face Activities | 92.86 (13.39) | 4.46 (1.895) | 4.64 (2.63) | 93.32 (13.03) | 0.25 | 0.179 | 0.571 | 0 | -0.071 | 0.105 (0.239) | 0.748 (0.306) | 0.33 (0.321) |

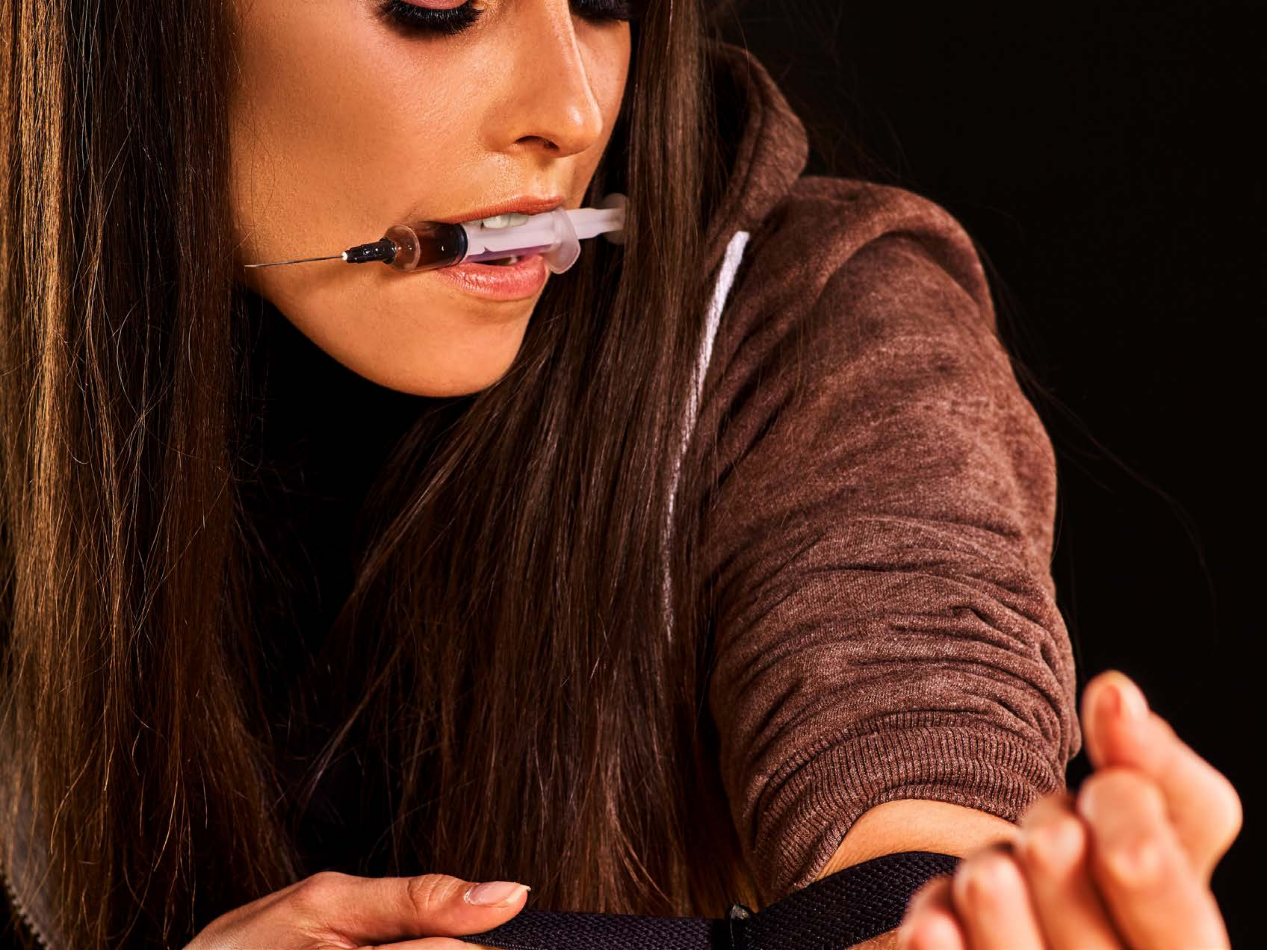

| Category | Craving (0-100) | Valence (1-9) | Arousal (1-9) | Typicality (0-100) | Relatedness | | | | | HSV | | |
|---|---|---|---|---|---|---|---|---|---|---|---|---|
| | | | | | Meth | Opioid | Both | Neither | MethToOpioid | Hue | Saturation | Value |
| Opioid Face Activities | 93.82 (10.78) | 4.36 (2.129) | 5 (2.75) | 94.36 (10.8) | 0.143 | 0.179 | 0.643 | 0.036 | 0.036 | 0.082 (0.195) | 0.688 (0.209) | 0.302 (0.306) |
